# FISH, a new tool for in situ preservation of RNA in tissues of deep-sea mobile fauna

**DOI:** 10.1101/2024.07.23.604794

**Authors:** V. Cueff-Gauchard, J. Aube, J-R. Lagadec, L. Bignon, J-P. Lafontaine, I. Hernandez-Avila, N. Marsaud, B. Shillito, L. Amand, E. G. Roussel, M-A. Cambon

## Abstract

Accessing the metabolic functioning of deep-sea animals in situ remains a technological challenge as the recovery time of samples is incompatible with the short lifespan of such molecules as mRNAs. Tools able to preserve RNA in situ exist but they are incompatible with the study of mobile fauna. Here we describe a new sampling tool, named FISH (Fixer In situ of Homogenized Substrates), implemented on a submersible and equipped with a number of new specific features to collect and preserve in situ tissue of mobile fauna. Connected to the suction pump of a submersible, FISH incorporates a sampling bowl to which two bottles of a preservative reagent are attached, a suction hose, and a support containing a motor connected to the sampling bowl by a magnetic coupling system. We used the deep-sea hydrothermal shrimp *Rimicaris exoculata* from the Mid-Atlantic Ridge as a model to test the suitability of our new tool. FISH was compared to two other sampling methods, which use a metatranscriptomic approach targeting microbial communities associated with cephalothorax symbionts. RNA quality, gene assignment and taxonomic and gene function diversity showed differences between in situ and on-board preservation of tissues. Of the alternative sampling methods tested, the suction sampler was clearly not suitable for RNA-based studies, while pressurized recovery showed results closer to the sample quality obtained with FISH sampling. The FISH sampler has therefore demonstrated to be a cost-effective and reliable tool to efficiently preserve RNA recovered from deep-sea environments.

## INTRODUCTION

The majority of visited hydrothermal fields are located at depths between 500 m, such as Solwara 17 in the back-arc Manus Basin (Massoth et al. 2008), and 4957 m for the Beebe site on the Mid-Cayman Rise (Connelly et al. 2012). These environments are characterized by steep physicochemical gradients controlled by strong spatial and temporal dynamics, creating multiple microhabitats. These ecosystems are sustained by chemolithoautotrophic microbial communities that can teem in abundant and highly diversified meiofauna and macrofauna communities. Accordingly, most endemic fauna (mussels, gastropods and shrimp), harbor symbiotic microbial communities implied in host nutrition called holobionts (Zilber-Rosenberg and Rosenberg 2008; Dubilier et al. 2008). To decipher the in situ metabolic functioning of hydrothermal communities, identifying active metabolic pathways and cellular regulation processes are essential, requiring proteomic or transcriptomic approaches.

One of the main limitations of these remote ecosystems lies in the time between sampling at depth and the recovery of samples on-board the oceanographic ship. Samples can stand for several hours outside the hydrothermal influence after sampling, inducing changes in physical and chemical conditions (e.g. temperature, dioxygen, hydrogen and hydrogen sulfide concentration), and a decrease in pressure during ascent. All these modifications lead to metabolic responses, mRNA rearrangements and degradation, and even cell death of both microbial and animal (host) populations, thereby impairing an accurate understanding of in situ biological processes.

RNA started being used as a proxy for marine microbial metabolic activity in the early 1990s (Kerkhof and Ward 1993; Kemp et al. 1993; Kramer and Singleton 1993; Lee and Kemp 1994; Kerkhof and Kemp 1999) but were limited by the sequencing technology available at the time. Transcriptomics and metatranscriptomics are now widely-used in marine ecology, allowing to better understand the regulation of cellular processes, metabolic pathways and active mechanisms in response to a given environment at a given time (Jiang et al. 2016; Bashiardes et al. 2016; Lavelle and Sokol 2018; Shakya et al. 2019; Mat et al. 2020; Page and Lawley 2022). It is also interesting to use microscopy to study the spatial distribution of expressed metabolic genes within a community, and to link the actors to a taxonomic identification using fluorescent in situ hybridization (Pernthaler and Amann 2004; Pilhofer et al. 2009; Hongo et al. 2016; Takishita et al. 2017; Miyazaki et al. 2020). However, the rapid reorganization and decay of messenger RNA often impair adequate conclusions on in situ expression levels.

The half-lives of mRNAs are indeed relatively short (Rauhut and Klug 1999). Several studies have demonstrated times varying from 1 to 15 min in bacteria, with averages of around 2 to 5 min depending on the bacterial lineage. They can however extend to 20 min during the stationary phase of growth (O’Hara et al. 1995; Bernstein et al. 2002; Hambraeus et al. 2003; Redon et al. 2005; Perwez and Kushner 2006; Steglich et al. 2010; Evguenieva-Hackenberg and Klug 2011; Mohanty and Kushner 2016). In archaea, different studies have described longer mRNA half-lives varying from 2 to 80 min such as in *Haloferax mediterranei* (Hennigan and Reeve 1994; Bini et al. 2002; Jäger et al. 2002; Andersson et al. 2006; Clouet-d’Orval et al. 2018). A study was also carried out at the level of marine microbial communities, where about 80% of the transcripts analyzed were reported to have a half-life between 10 min and 400 min (Steiner et al. 2019). The half-life of mRNA therefore varies according to genes, depending on their location on the chromosome, their function and accessibility to ribonucleases (Mohanty and Kushner 2016). They also depend on the growth phase and on some stress conditions (Takayama and Kjelleberg 2000). The half-lives of mRNA in Bacteria and Archaea remain on average much shorter than in Eukaryotes where they can be preserved for more than 24 hours (Tourrière et al. 2002; Edri and Tuller 2014). Furthermore, marine bacterial cells may contain significantly fewer mRNA molecules, with about 200 molecules of mRNA in a typical marine bacterial cell in coastal seawater *vs* 1800 mRNA molecules in *Escherichia coli* cultures (Moran et al. 2013).

Approaches to identify and quantify in situ deep-sea microbial gene expression remain under scrutiny. Potential biases due to short mRNA lifespan have been suggested in some publications dealing with hydrothermal fluid communities (Wu et al. 2011, 2013; Lesniewski et al. 2012; Baker et al. 2013; Li et al. 2016). In contrast, other studies have analyzed microbial communities metatranscriptomes from hydrothermal chimneys (He et al. 2015), or from hydrothermal animals such as gastropods (Lan et al. 2021), vesicomyids (Hongo et al. 2016), hydrothermal shrimps (Zhu et al. 2020), and sponges in cold seeps (Rubin-Blum et al. 2017, 2019) without addressing this question.

However, as pointed out by Stewart in his review (Stewart 2013), it is critical to ensure that the maximum amount of information from the mRNA is obtained without bias, and should therefore be addressed using new sampling instrumentation (McQuillan and Robidart 2017). For water column or hydrothermal fluid sampling, six different tools have been developed (Feike et al. 2012; Wurzbacher et al. 2012; Akerman et al. 2013; Breier et al. 2014; Taylor et al. 2015; Govindarajan et al. 2015; Edgcomb et al. 2016; Fortunato and Huber 2016; Fortunato et al. 2018; Cron et al. 2020). For in situ fixation, the most commonly used fixative was RNA*later*^®^ (Ambion Inc., Austin, TX), a commercial ammonium sulfate concentrated solution, denser than seawater, which stabilizes cellular RNA by precipitating out RNases, without the need to freeze samples (Mutter et al. 2004; Salehi and Najafi 2014; Menke et al. 2017).

To preserve the RNA of macrofauna in situ, few systems have been developed to date. Some studies on *Bathymodiolus* mussels mention boxes filled with ammonium sulfate-saturated fixative (a type of homemade RNA*later*^®^) (Takishita et al. 2017; Mat et al. 2020). The mussels are simply dropped inside the open box, the fixative remains in the box due to its higher density and then the box is closed before ascent. Two systems have been designed to fix in situ slow-moving animals such as galathea *Shinkai crossnieri,* gastropods *Alviniconcha* or scally foot snail *Chrysomallon squamiferum*. The “In situ Mussel and Snail Homogenizer” ISMACH (Sanders et al. 2013) consists of a box inside which the animals are placed. It allows seawater to be replaced by RNA*later*^®^, before subsequent homogenization. The second system, unnamed, is a suction sampler connected to a plastic bag filled with sulfate salt solution via a hose with a cock valve (Watsuji et al. 2014; Motoki et al. 2020; Sun et al. 2020; Miyazaki et al. 2020). The transfer of the fixative is achieved by diffusion and lasts about 9 minutes.

Here we present the development of a new in situ collection tool named FISH (Fixer In situ of Homogenized Substrates) implemented on any submersible allowing to: (i) capture mobile fauna, (ii) instantly preserve tissues using RNA stabilization reagent such as RNA*later*^®^ or formaldehyde, (iii) homogenize the sample to facilitate tissue impregnation. The biological model used in this methodological study, *Rimicaris exoculata,* a deep-sea hydrothermal shrimp, harbors complex nutritive microbial symbiotic communities, one located in its inflated cephalothoracic cavity (for review (Zbinden and Cambon Bonavita 2020). The aim of this study is to assess the efficiency of FISH to preserve in situ RNA in tissues associated with microbial symbiotic communities. Hence, comparative analyses of the metatranscriptomics of microbial symbiotic communities were performed from samples of *R. exoculata* collected using FISH and two other methods: the submersible suction sampler exposing samples to decompression, and the PERISCOP^©^ pressurized recovery device (PRD), (Shillito et al. 2008, 2023).

## MATERIALS AND PROCEDURES

### Sampling site

Twenty-four in situ deployments of the FISH sampler were carried out at different depths on the Mid-Atlantic Ridge, in the Western Basin of the Mediterranean Sea and in the back-arc basins of the West-Pacific. These deployments took place during trial technical expeditions operated with the Human-Occupied Vehicle (HOV) Nautile (ESSNAUT2017 https://doi.org/10.17600/17009100, ESSNAUT2021 https://doi.org/10.17600/18002379, ESSNAUT2022 https://doi.org/10.17600/18002759), the Remotely-Operated underwater Vehicle (ROV) Victor 6000 (ESSROV2019) and during the oceanographic expeditions HERMINE in 2017 https://doi.org/10.17600/17000200, BICOSE2 in 2018 https://doi.org/10.17600/18000004, and CHUBACARC in 2019 https://doi.org/10.17600/18001111). The *Rimicaris exoculata* shrimp samples used in this study were collected from the Snake Pit hydrothermal field (23°23’N, 44°58’W, -3480 m depth) on the Mid-Atlantic Ridge, on two active sites, “The Beehive” and “The Moose” (Fouquet et al. 1993), during the BICOSE2 expedition (February 2018).

### Different sampling tools

Samples of *R. exoculata* were collected using three different deep-sea sampling tools, including FISH. (i) Samples collected using the submersible’s suction sampler at the end of the dive (Figure S1-a suppl. data) were exposed to a change in environmental factors (e.g. pressure, temperature, chemistry) during the two-hour ascent of the submersible. (ii) Samples were also collected using Periscopette, a sampling cell inserted into the PERISCOP, which maintains in situ pressure during recovery (Figure S1-b suppl. data) (Shillito et al. 2008, 2023). PERISCOP, fixed on an independent shuttle device, was released immediately after in situ closure. PERISCOP’s syntactic foam casing minimizes temperature variation, which may occur when the samplers reach warm surface waters (possibly up to 28°C water temperature at the sea surface).

### FISH instrument design

The sampler FISH consists of several components (Fig. 1a). A 1.7 L (internal volume) removable PVC sampling bowl (in yellow) is connected on one side to a suction pump via the submersible suction system, and on the other side to a transparent flexible sampling tube via a PVC base. This bowl is equipped with blender blades (4) (Moulinex P/N SS98994) connected to a magnetic plate, and a spring-loaded, watertight rotating lid with a rotating handle. The AISI 316L austenitic stainless steel springs, with 45 coils, are 80 mm long at their initial resting position, and stretch to 180 mm with a spring force of 0.348 N/mm. Two 850-mL stainless-steel 316L bottles (in green) equipped with a piston were attached to each side of the sampling bowl, to which they were connected by flexible tubes. These bottles contain the preservative reagent (e.g. RNA*later*^®^). The sampling bowl is then inserted into a PVC base (Fig. 1b), fixed in the basket of the submersible (Fig. 3a – 3h). A hydraulic motor (10) (HPI P/N M3 CBN 1004 CL 20C01 N, capacity = 4.09 cc/rev, maximum pressure = 20 MPa, maximum speed = 5000 RPM) powers the rotation of the blade, and is controlled by the submersible hydraulic power unit (maximum speed = 2000 RPM for HOV Nautile and 2200 RPM for ROV Victor6000). The motor is connected to a magnetic plate (8) allowing the coupling with the blender blades (4) inside the sampling bowl with a coupling force of 1.25 N.m.. The base also allows the junction of the sampling bowl with the sampling tube at the inlet and the suction sampler at the outlet (9). To avoid impeding the use of the submersible’s suction device during the entire dive, a by-pass has been designed to supplant FISH (Fig. 3h).

**Figure 1:**
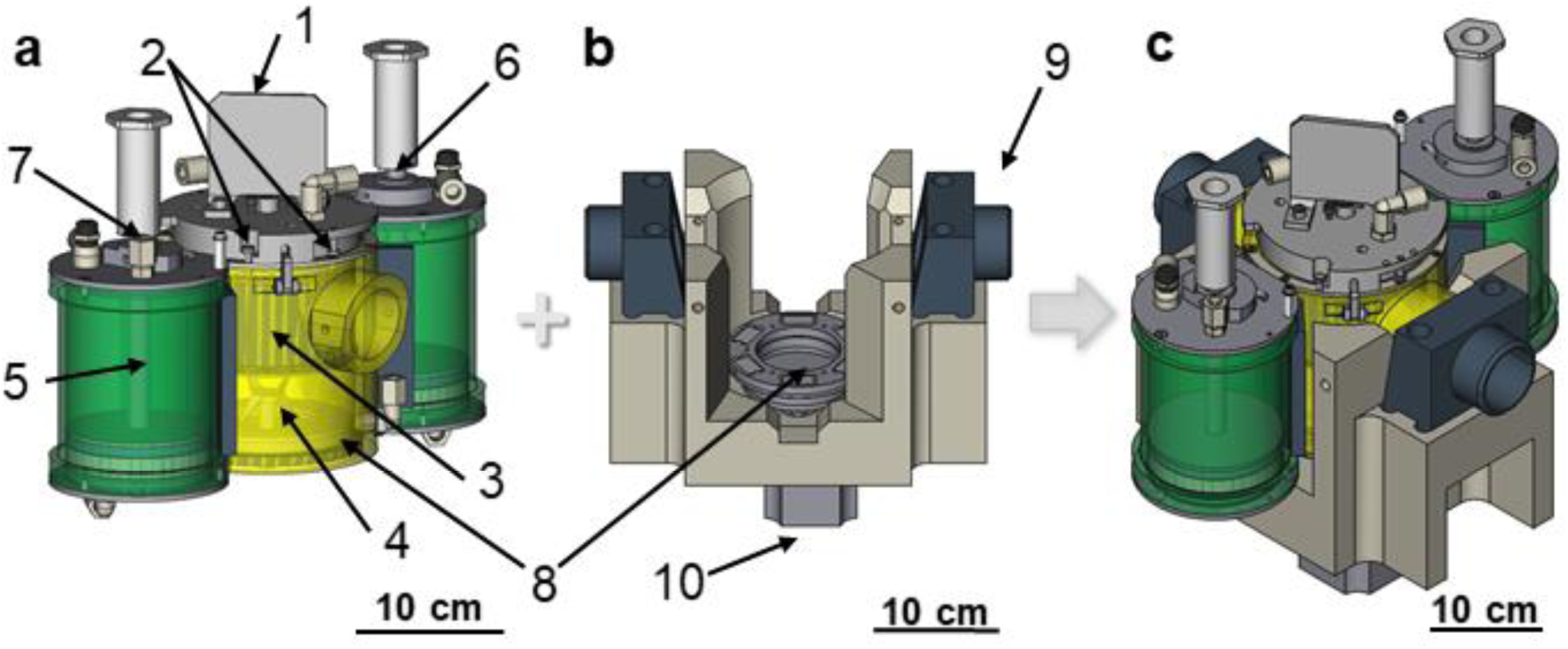
3D visualization of the FISH sampler with (a) complete sampler system including, in yellow, the sampling bowl and in green the fixative bottles, 1: handle, 2: hooks for lid springs, 3: watertight cover incorporating a metal grid, 4: mixer blades, 5: piston, 6: spring, 7: filling plug, 8: magnetic coupling system ; (b) base with magnetic coupling system, 9: end cap for vacuum hose connection, 10: hydraulic motor (c) FISH sampler on its base.

### Sampling system - general principle and operating mode

Before the dive, the support was fixed with brackets (Fig. 2a) onto the submersible’s front basket. Hydraulic cables were connected between the FISH engine and the submersible’s hydraulic system ((Figure S2 suppl. data). A derivation upstream of the sampling bowls of the suction sampler was performed to use the submersible’s suction sampler pump. The suction sampler hose, without its nozzle, was connected onto the back of the support. A transparent flexible hose with a straight metal tip for sampling was placed on the front of the support (Fig. 2b).

**Figure 2:**
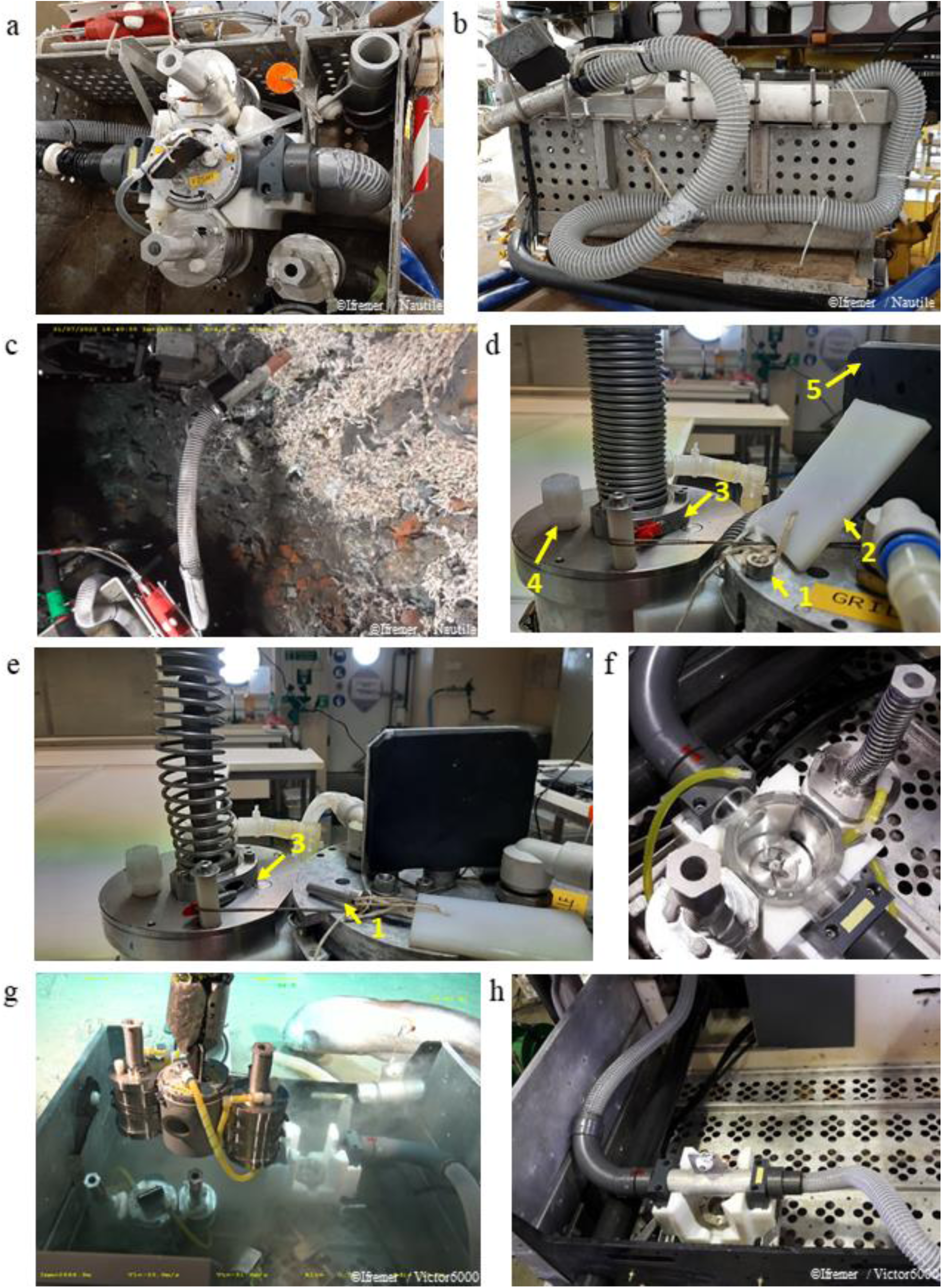
The FISH sampling process. (a) FISH base and system inside submersible’s basket. (b) Suction hose connected to upstream FISH sampler. (c) Shrimp suction through the hose upstream of the sampler FISH. (d) Open position of the sampler with armed fixative bottles: unlocking of the cover by removing the locking pin (1) in a vertical position using the float (2). 1: lid locking pin, 2: float, 3: bottle locking pin, 4: filling plug, 5: grip handle also allows to close the lid (e) Closed position of the sampler with fixative bottles engaged. Spring-loaded closure causes the locking pins (3) of the fixative bottles to be withdrawn simultaneously. Removal of the pistons in the bottles, pushing the fixative into the sampling tank. (f) Mixing part of the sampling system. The submersible’s hydraulic power unit is started to drive the mixing blade thanks to a magnetic coupling downstream of the motor. (g) Removal of the sampling system from the holder to be replaced by another FISH system, (h) View on the by-pass in place for the use of the suction sampler.

FISH boxes were prepared on board the ship in the laboratory. A pin locked the lid in the open position (Fig. 2d, 1) to allow sampling. Then, the fixative bottles were armed, locking the piston in the bottom of the bottle, thanks to the side pins (Fig. 2d, 3). RNA tissue preservation reagent (such as commercial RNA*later*^®^ (Sigma-Aldrich) or homemade buffer 0.019M EDTA, 0.018M sodium citrate, 3.8M ammonium sulphate, pH 5.2, according to Menke (Menke et al. 2017) or formaldehyde 3%) was then transferred into each bottle through the filling plug (Fig. 2d, 4) by means of a small funnel. Then, one of the systems was placed in its support in the front basket of the submersible while the other was placed next to it, or inside the shuttle device, ready to use.

On the seafloor, next to the sampling site, the submersible’s arm first deployed the flexible suction hose to collect mobile fauna (Fig. 2c). The clear flexible pipe made it possible to count the shrimps during the suction phase in order to obtain between 15 and 20 specimens inside the sampler bowl (Fig. 3a). A metal grid fixed inside the lid upstream of the outlet pipe confined the specimens inside the bowl (Fig. 3a).

**Figure 3:**
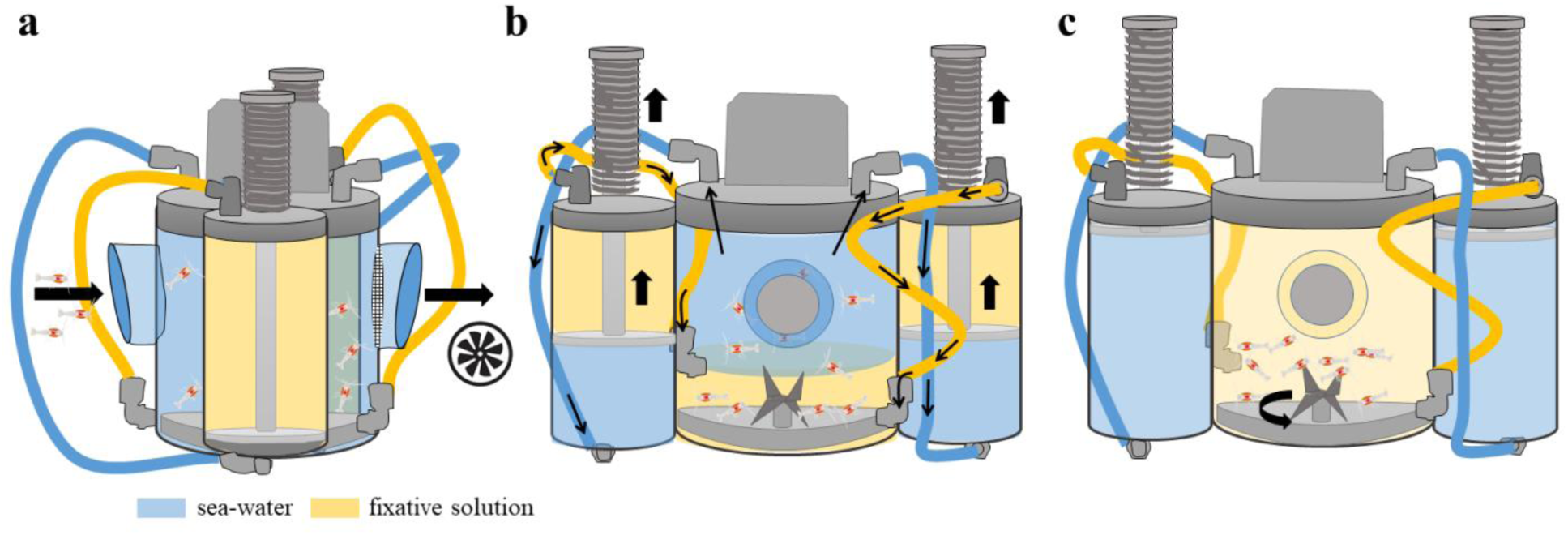
Schematic representation of the fluid flows in FISH during sampling. (a) Side view of FISH: suction of shrimps inside the sampling bowl which are blocked at the outlet by a grid. (b) Front view of FISH the inflow of the fixative solution (in yellow) initially contained in the bottle of fixative inside the sampling bowl which replaces the sea water (in blue) itself sucked back inside the bottles of fixative. (c) Rotation of the blender blades of the magnetic driven plate.

Then the submersible’s arm released the locking pin of the lid (Fig. 2d, 1) by pulling on a float (Fig. 2d, 2) following a vertical movement. Two springs, fixed in the lid, closed it by rotation. The lid rotation led to the simultaneous removal of the locking pins of the two fixative bottles (Fig. 2d and 2e, 3), in order to release the spring-loaded piston of the fixative bottles. This piston then moved upwards, pushing the preservative reagent, which was denser than seawater, from the top of the bottles to the bottom of the sampling bowl (Fig. 3b). The preservative reagent replaced the seawater in the sampling bowl in a few seconds, while the seawater was transferred to the bottom of the fixative bottles (Fig. 3b). Upon contact with the preservative reagent, we observed that the fauna died instantly.

Homogenization followed through rotation of the blade (Fig. 2f, 3c). For this, a driving magnetic plate, linked to the rotating shaft of the hydraulic motor was activated, enabling the rotation of the blender blades of the magnetic-driven plate inside the bowl (Fig 3c). The blender blade rotated at 100 rpm for about 20 seconds facilitating homogenization and complete impregnation of the preservative reagent. It was important not to rotate for too long to avoid tissue damage, which would impair later dissections. The system could be removed from its support to be placed in the independent shuttle vector, between the seafloor and the sea surface, to optimize recovery time on board the research vessel. Another FISH sampler could replace the previous one on the support, to provide additional sampling during the same dive (Fig. 2g). At the end of sampling with FISH, the suction sampler was put back into operation thanks to the use of a by-pass, without loss of suction power (Fig. 2h). A video showing the complete sampling sequence is available as supplementary data.

### Sample processing

On board, shrimps were recovered from the FISH sampling bowl and transferred to a fresh RNA*later*^®^ solution (Sigma-Aldrich). The different organs, such as branchiostegites, scaphognathites and exopodites for the cephalothoracic cavity, were then dissected (Figure S3 suppl. Data) (Cambon-Bonavita et al. 2021), under sterile conditions in Petri dishes filled with RNA*later*^®^. The different tissues were transferred to 1.8 ml cryotubes with RNA*later*^®^ and stored for 24h at 4°C. For long-term preservation, tubes were then transferred at -80°C. For immediate on-board RNA extraction procedures, RNA*later*^®^ was replaced with a TRIzol^TM^ reagent (Invitrogen).

Shrimps recovered from the conventional suction sampler and from PERISCOP were processed on board in the same way as the FISH samples, i.e. transferred to fresh RNA*later*^®^ (Sigma-Aldrich) before further dissection. Tissues for the metagenomics studies were also dissected in fresh RNA*later*^®^ (Sigma-Aldrich) from shrimps collected using the suction sampler on “The Beehive” site of the Snake Pit hydrothermal field. After being flash-frozen with liquid nitrogen, all tubes were stored at -80°C.

### RNA extraction and sequencing

For each shrimp sampled, half of the branchiostegites, scaphognathites and exopodites of cephalothoracic cavity were pooled (Cambon-Bonavita et al. 2021), providing two replicate subsamples per shrimp. For all sampling methods, each cephalothorax sample was grounded with Nucleospin beads (Macherey-Nagel), in 1 mL TRIzol^TM^ reagent (Invitrogen) on a Vortex Genie2 for 10 min at maximum speed. Total RNAs were then extracted with the TRIzol^TM^ method as recommended by the manufacturer with two chloroform purifications. RNA extracts were treated by DNA-free kit DNase Treatment and Removal Reagents (Invitrogen) according to the manufacturer’s recommendations. Concentrations of extracted RNA were measured using a Qubit^TM^ 3.0 Fluorometer (ThermoFisher Scientific) with Qubit^TM^ RNA HS Assay Kit and the Bioanalyser (Agilent) with RNA 6000 Nano Kit (Agilent) to evaluate the quality of RNA through the RNA integrity number (RIN) (Schroeder et al. 2006). However, as the RIN values were defined using standards for prokaryotic RNA, results could have been biased as they were obtained from a mixture of prokaryotic (symbionts) and eukaryotic (host) RNAs. Since the RIN algorithm was unable to differentiate eukaryotic/prokaryotic/chloroplastic ribosomal RNA, this may have created serious quality index underestimation.

Ribodepletion and Illumina libraries were prepared with Illumina^®^ Stranded Total RNA Prep, Ligation with Ribo-Zero Plus at GeT-Biopuces platform (INSA, Toulouse, France) according to Illumina recommendations. Briefly, the use of this kit was carried out in several stages: first the depletion of bacterial and eukaryotic ribosomal RNA, then the fragmentation and denaturation of RNA. Then there was the synthesis of the first strand of cDNA followed by the synthesis of the second strand of cDNA, the adenylation of the 3’ end fragments to allow for ligation of the Illumina adapters to the fragments, followed by the cleaning of the adapter-ligated fragments, the amplification of libraries and finally the cleaning of libraries. The concentration and quality of the final libraries were then checked. The metatranscriptomic sequencing on an Illumina Novaseq 6000 instrument (2 x 150 bases paired-end) was performed at the GeT-PlaGe platform (INRA, Toulouse, France).

### DNA extraction and sequencing

Metatranscriptomic analysis required sequencing the metagenome of specimens from the same sites (i.e. Snake Pit – “The Beehive”), to reduce biases that would have been induced by using metagenomes available in the databases (Cambon-Bonavita et al. 2021). Total DNA of the cephalothorax (branchiostegite, scaphognathite and exopodite) was extracted from four *Rimicaris exoculata* individuals: two males and two females. The Nucleospin soil kit (Macherey-Nagel) was used according to the manufacturer’s recommendations. Nanodrop 2000 (ThermoFisher) was used to evaluate DNA quality while the Qubit^TM^ 3.0 Fluorometer (ThermoFisher Scientific) with Qubit^TM^ DNA HS Assay Kit was used to validate the DNA quantity. Metagenomic sequencing was conducted on an Illumina HiSeq 3000 instrument (2 x 150 bases paired-ended) at the GeT-PlaGe platform (INRA, Toulouse, France) from libraries built with an Illumina TruSeq Nano kit.

### Metatranscriptomic analysis

Snakemake workflow (Köster and Rahmann 2018) was used to evaluate the quality of sequences with FastQ v.0.11.8, to trim adaptors with Cutadapt tool v.1.18 (Martin 2011) and to proceed to ribodepletion with Bowtie2 v.2.3.5 tool (Langmead and Salzberg 2012) and SILVA LSU+SSU Reference sequence databank v.138. Kaiju tool v.1.7.1 (Menzel et al. 2016) was also integrated in the pipeline to classify taxonomy of reads with a graphic interface via Krona v.2.7 (Ondov et al. 2011). This tool taps into the genome data available in the NCBI RefSeq library. The same workflow was applied to metagenomic data without the ribodepletion step. Then the metagenomic Snakemake workflow of anvi’o v6. (Eren et al. 2015) was run. First, illumina-utils v.2.8 to control quality of metatranscriptomic and metagenomics short reads with the “iu-filter-quality-minoche” program with default parameters were used. The four obtained metagenomes were co-assembled with Megahit v.1.2.9 (Li et al. 2015) with the meta-sensitive mode and a minimum contig length of 1000 bp. An anvi’o annotated contig database was next generated to recognize prokaryotic genes using Prodigal v.2.6.3 (Hyatt et al. 2010). Gene functions and metabolic pathways were annotated from the NCBI database of Clusters of Orthologous Genes (COGs) (Galperin et al. 2021) and with eggNOG-mapper v.2.1.8 (Huerta-Cepas et al. 2017; Cantalapiedra et al. 2021) based on precomputed orthology assignments. Simultaneously, short reads of each metatranscriptome were mapped against contigs formed with co-assembly and indexed using bowtie2 v.2.4.2 (Langmead and Salzberg 2012). SAMtools v.1.7 (Li et al. 2009) was then used to generate sorted and indexed BAM files. Individual BAM files were profiled to generate anvi’o profiles using “anvi-profile” which were combined into a single anvi’o profile with the program “anvi-merge”. The function “anvi-summarize” was applied to export functional annotation. Then the function “anvi-profil-blitz” was used to obtain gene-level coverage and detection stats, using the indexed bam-files. Taxonomy assignment on genes was also carried out using the MMSeqs2 v.14.7e284 (Steinegger and Söding 2017) and the UniRef90 database v.2022-01 to compare with the Kaiju tool.

### Statistical analysis

The statistical analysis and visualization of the data obtained according to the different sampling tools were carried out using R software v.4.3.3 (R Core Team 2024) under RStudio v.2023.12.1.402 (Posit team 2024). R Packages Tidyverse (Wickham et al. 2019), ggpubr (Kassambara 2022a), rstatix (Kassambara 2022b), pastecs and FSA (Ogle et al. 2023) were used to analyze data from RNA extractions i.e. concentrations and RIN. For defective RIN values, a RIN value of 0 was assigned. Average and standard errors were calculated for RNA concentration and RIN by separating samples by sampling conditions, i.e. the site associated with the sampling tool. For statistical analysis, data were also separated by the same sampling condition. A number of preliminary tests were carried out: identification of outliers, assumption of normality of the data by Shapiro’s test, assumption of homogeneity of variances by Levene’s test. As some of the samples did not follow a normal distribution, a non-parametric Mann-Whitney test with the “greater” alternative for RIN values was used to compare two by two variances as a function of sampling conditions followed by the non-parametric Kruskal-Wallis test and Dunn post-hoc tests.

A second matrix was generated to include the raw-read data after each cleaning step in order to compare the different sampling conditions, i.e. the sampling tool associated with origin site, integrating the calculation of means and standard deviation. The same statistical tests as above were applied.

Another matrix was created from a file generated with the “anvi-profil-blitz” function to give the total number of transcript read mapping to genes, detected per sample, and the number of different related genes expressed per sample. To obtain this, all values different from zero were replaced by one to deduce the number of related genes expressed per sample whatever the number of copies retrieved per sample. The same R Packages were used to analyze the differences in total transcript abundance and transcript diversity depending on the sampling condition, i.e. the sampling tool associated with site of origin. The same preliminary tests were performed as mentioned above. As some of the samples did not follow a normal distribution, the analysis of variance was then conducted by the non-parametric Mann-Whitney test with the “greater” alternative used to compare two by two variances as a function of sampling conditions. Differential expression analysis was achieved using R packages DESeq2 (Love et al. 2014), phyloseq (McMurdie and Holmes 2013), Tidyverse and vegan (Dixon 2003). A distance matrix was generated from the transcripts detected per sample with the phyloseq tool, integrating data normalization with the variance stabilizing transformation (vst) incorporated in the DESeq2 package. A principal coordinate analysis (PCoA) was performed to represent the different samples according to the Bray-Curtis dissimilarity matrix. Simultaneously, a permutational multivariate analysis of variance (PermANOVA) was run to compare variances with the Adonis2 function based on Bray-Curtis dissimilarities and 9999 permutations to test the significance of the different metadata (site, tool, tool associated with the site of origin, RNA quality). Then, differential expression analysis continued using only the DESeq2 R package outside the phyloseq environment. Initial gene counts were previously filtered by removing those whose total was less than five and vst transformation was performed. Differentially expressed genes with adjusted p-values of 0.05 (padj < 0.05) and absolute log2-fold changes of two were considered significant in this study. Their broad function type was assigned by compiling the results of the COG annotation “COG20_CATEGORY” and the EggNOG-mapper annotation “EGGNOG_COG_CATEGORY”. Figures were obtained thanks to ggplot2 R package (Wickham 2009), ggrepel, RColorBrewer, gridExtra and then refined with Adobe Illustrator.

### Code and data availability

The metatranscriptome raw reads are accessible in the European Nucleotide Archive under Bioproject Accession Number ERP162070 and the metagenomes raw reads are accessible under Bioproject Accession Number ERP162010. The URL https://gitlab.ifremer.fr/vc05320/fish-tool_rimicaris-exoculata-cephalothoracic-epibionts-metatranscriptome provides access to a detailed and reproducible bioinformatic workflow for all bioinformatic and statistical analyses.

## ASSESSMENT

During the BICOSE2 expedition, shrimps from two out of eight deployments of the FISH sampler were used in the present study. The shrimp tissues preserved in situ were collected from two different active sites on the Snake Pit hydrothermal field: “The Beehive” and “The Moose”(Fouquet et al. 1993). While no biometric measurements were made on the collected shrimps, adult individuals collected from large aggregates which appeared to be homogeneous in size but larger at “The Moose” site than at “The Beehive” site. However, biometric measurements performed during another study on the same expedition showed an average carapace length of 14.8 ± 4.8 mm (n=720) at “The Moose” site for all individuals, all life stages and sexes combined, compared with 11.1 ± 3.1 mm (n=1271) at “The Beehive” site (Methou 2019). For each site, the same shrimp aggregate was selected for sampling with FISH and PERISCOP on the one hand, and FISH and suction sampler on the other (Figure S4). Unfortunately, samples could not be collected with all three tools at the same site due to technical and logistic constraints.

The efficiency of the FISH sampler was assessed by comparing the quantity and quality of extracted RNA individually with those obtained with the two other sampling methods. The abundance of genes detected in the metatranscriptomes was also compared as well as the differential expression of genes according to the sampling tool, all genes combined (host and symbionts). On board, shrimp tissues preserved in situ presented a different texture compared to fresh ones, as if “baked” by RNA*later*^®^. Shrimps collected using the suction sampler were not very active, appearing either dead or unhealthy. Shrimps brought up in the PERISCOP were very active, and therefore appeared quite healthy. For a given sampling site, adult individuals of similar size were selected for dissection, whatever the sampling method.

### RNA extraction quality

A different number of extractions had to be performed depending on the sampling method. In some cases, six extractions were not enough to secure at least three replicates satisfying the sequencing platform’s quality requirements. For “The Beehive” site, six extractions were performed from FISH samples, and another six from PERISCOP. For “The Moose” site, nine extractions were required for the FISH samples and 12 for the suction sampler samples. Despite these 12 extractions from the suction sampler samples, only one RNA extract was able to comply with the platform’s requirements, i.e. to achieve RNA integrity qualities via the RNA integrity number (RIN) (Schroeder et al. 2006) greater than eight (Figure 4 and Supplementary Table S1). This means that 92% of the RNA extracted from samples collected using the suction sampler were degraded (RIN<7), even though the RNA concentrations were high. RNA obtained from samples recovered from PERISCOP was compiled with good quality criteria for 50% whereas 33% were of poor quality.

**Figure 4:**
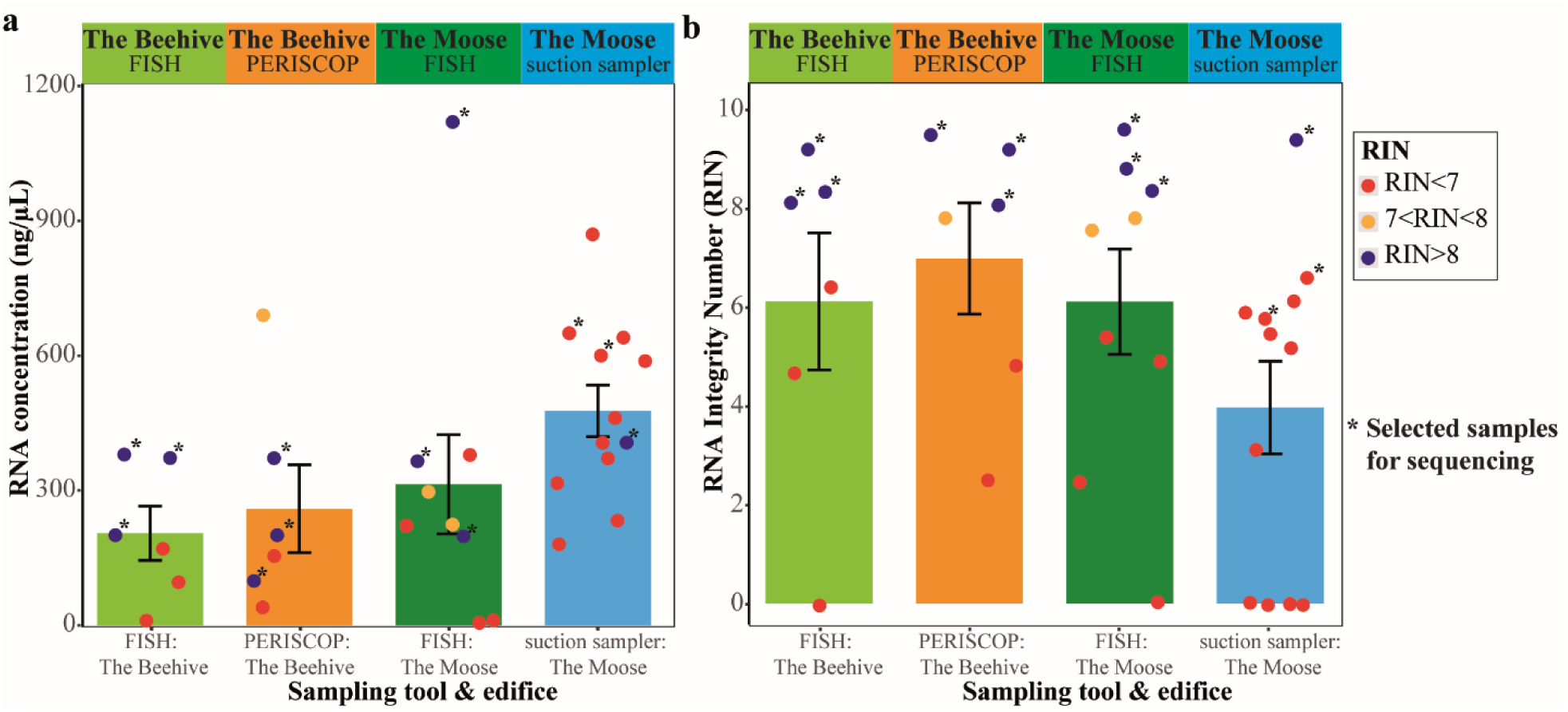
(a) extracted RNA concentrations and (b) RNA integrity Number (RIN) associated with these extracts. The dots represent individual extracts, the barplots the mean of concentrations or of the RIN obtained and the error bar corresponds to the standard error, separated according to the sampling tool and site of origin. The color of the dots vary according to the value of the RIN: in red, RINs <7; in orange, RINs between 7 and 8; and in dark blue, RINs >8. Dots marked with a star have been selected for sequencing.

Extractions from the FISH sampler showed significant disparity in terms of concentration and quality. Indeed, for the “The Moose” site, two extracts out of nine did not reach the minimum concentrations required for sequencing, but one of the extracts obtained from FISH exceeds 1 µg/µL, corresponding to the most concentrated extract of all the experiments. Thus, for FISH sampling on “The Beehive” site, 50% of the extracts reached a sufficient quality for sequencing while 50% of the extracts were of poor quality. On “The Moose” site, RNA extracts were of lower quality with 33% of the extracts displaying RIN >8, 22% with RIN between 7 and 8 and 44% with RIN <7.

Overall, the suction sampler yielded more concentrated RNA but of poorer quality, requiring heavier sampling for fewer exploitable results, demonstrating its unsuitable use for routine transcriptomic approaches. In comparison, samples collected with FISH and PERISCOP appeared to generate RNA of more homogeneous quality and quantity. Statistical tests showed significant differences in RNA concentrations between FISH and the suction sampler on “The Moose” site (Wilcoxon-Mann-Whitney test, W=24, *Pvalue* =0.036) and between FISH from “The Beehive” site and suction sampler from “The Moose” site (Wilcoxon-Mann-Whitney test, W=9, *Pvalue* =0.013). For RIN values, RNA extracts from PERISCOP on “The Beehive” site showed significant differences with quality of RNA extracts from suction sampler on “The Moose” site (Wilcoxon-Mann-Whitney test, W=54, *Pvalue* =0.05). But there were no significant differences on RIN values between FISH and the suction sampler on “The Moose” site (Wilcoxon-Mann-Whitney test; *Padj* > 0.05, Supplementary Table S2).

Three RNA extracts with RIN >8 were selected for sequencing for each site and sampling tool (Figure 4b). For sampling with the suction sampler, two extracts with RIN <7 had to be chosen on the basis of a compromise between highest possible RIN value and RNA concentration to have a triplicate.

### Quality trimming and filtering statistics from sequencing data

Shotgun sequencing of total RNA recovered for this study was successfully completed for all samples. It resulted in 384 million pairs of raw reads with an average of 32.02 ± 1.93 million pairs of raw reads per metatranscriptome (Supplementary Table S3). After rRNA depletion with Bowtie2 and filtration with quality minoche between 15 097 979 and 36 231 982 paired-reads per sample were conserved, representing 87-91% of the initial sequences. As the quantities of sequenced libraries had been normalized beforehand, the number of reads obtained and the qualities were equivalent for all samples, as shown by the statistical analyses, which did not reveal any significant differences between sampling tools (Wilcoxon-Mann-Whitney test; *Padj* = 1, Supplementary Table S4).

With regard to the metagenome sequencing data, a total of 619 million pairs of raw reads per sample were obtained with values ranging from 137 358 894 to 166 733 163 raw reads per sample (Supplementary Table S5). The filtration steps resulted in the retention of between 89% and 91% of the starting reads. Low-quality reads were removed from the data set for all following steps.

Co-assembly of the four samples produced a total of 1 044 858 contigs longer than 1 kbp which recruited between 83% and 90% of metagenomic reads and between 92.96% and 95.61% of metatranscriptomic reads (Supplementary Tables S5 and S3). The results obtained from the “anvi-profile-blitz” function were used to determine the number of reads that mapped onto a gene, corresponding to a total of 223 539 897 genes for the Rimi316Fem sample, 184 946 472 genes for Rimi317-Fem, 237 934 362 genes for Rimi325Mal and 207 291 708 genes for Rimi326Mal.

### Number of total recruited transcripts and number of different genes detected per sample according to the sampling tool and site

As previously indicated, a table listing the number of transcripts detected per sample (only reads mapping to genes) was retrieved using the “anvi-profile-blitz” function. The data were aggregated to obtain the total number of detected transcripts per sample. All values different from zero were replaced by one to deduce the number of related genes expressed per sample whatever the number of copies retrieved per sample (Supplementary Table S6). The two types of data were then compared according to the sampling tool and the sampled site.

The number of total recruited transcripts was higher for samples from the FISH sampler on “The Moose” site compared with the suction sampler on the same site or samples taken on “The Beehive” site, with both the FISH sampler and the PERISCOP (Figure 5a). For FISH sampling on “The Moose”, there was a mean of 33 216 129 ± 10 979 054 total recruited transcripts against 19 097 054 ± 10 979 054 transcripts for the suction sampler, 21 180 038 ± 8 943 098 transcripts for FISH sampling from “The Beehive” and 18 956 230 ± 3 831 351 transcripts from PERISCOP on “The Beehive” site (Supplementary Table S6). On average, compared to the abundance level of detected transcripts using the FISH sampler on “The Moose”, 42.93% fewer transcripts were detected from the samples from the suction sampler on the same site, 42.51% fewer from the PERISCOP samples on “The Beehive” site and 36.24% fewer from the FISH samples on “The Beehive” site. As the standard deviations appeared to be quite large, it was important to validate the observed trends by statistical tests. However, statistical data had to be put into perspective, given the low number of samples per population type (tool associated with the site of origin) (n=3). The non-parametric Wilcoxon or Mann-Whitney test with the alternative “greater” was used to compare variances because of the non-normality distribution of values.

**Figure 5:**
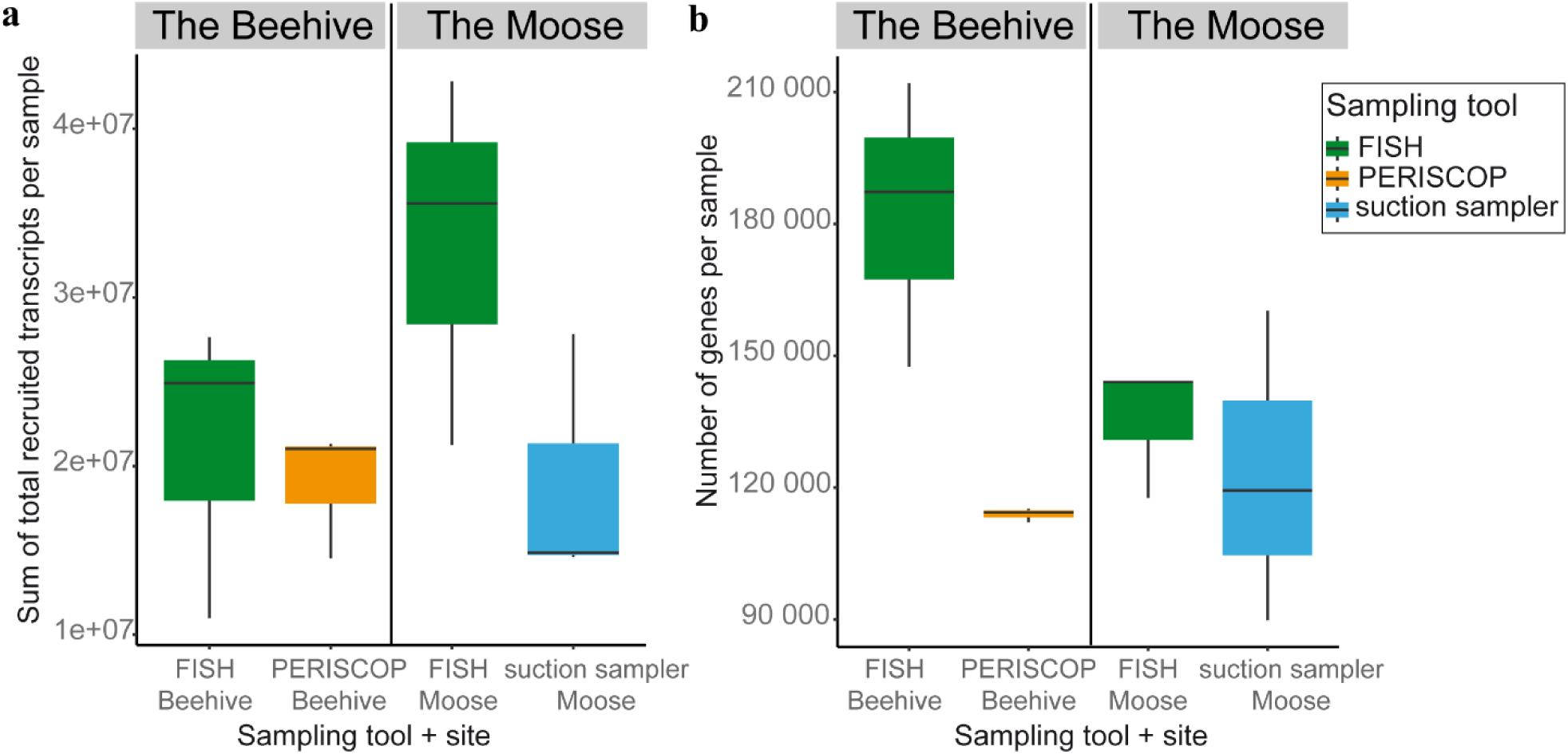
Boxplot of (a) Sum of total recruited transcripts mapped to genes per sample and (b) Number of detected genes per sample, with samples separated by site of origin and sampling tool. In green, sampling with FISH, in orange, sampling with PERISCOP and in blue, sampling with suction sampler.

The number of total recruited transcripts from FISH at “The Beehive” site was not significantly different from that obtained with PERISCOP at the same site (Wilcoxon test, W=6, *Pvalue* =0.35, supplementary table S7), nor from that obtained with the FISH sampler at “The Moose” site (Wilcoxon-Mann-Whitney test, W=2, *Pvalue* =0.9). The number of total recruited transcripts from FISH at “The Moose” site was also not significantly different from that obtained from the suction sampler at this site (Wilcoxon-Mann-Whitney test, W=8, *Pvalue* =0.1).

Looking at the average number of different detected genes (Figure 5b), FISH samples from “The Beehive” site had the highest number of detected genes with an average of 182 257 ± 32 542 genes, compared with 113 901 ± 1 597 genes for PERISCOP at the same site, 135 295 ± 15 239 genes for FISH at “The Moose” site and 123 156 ± 35 348 genes for the suction sampler at “The Moose” site. Compared with the gene diversity detected with the FISH sampler at “The Beehive” site, this represented 37.51% less gene diversity detected with PERISCOP at the same site, 25.77% less for FISH at “The Moose” site and 32.43% less for the suction sampler. Given the large standard deviation, there were no significant differences in the number of detected genes between FISH and the suction sampler on “The Moose” site (Wilcoxon-Mann-Whitney test, W=5, *Pvalue* =0.5, supplementary table S7). In contrast, the number of different genes obtained from FISH on “The Beehive” site was significantly higher than that obtained from PERISCOP (Wilcoxon-Mann-Whitney test, W=9, *Pvalue* =0.05*) or from FISH at “The Moose” (Wilcoxon-Mann-Whitney test, W=9, *Pvalue* =0.05*). As previously indicated, statistical results should be treated with caution, considering the low number of comparative values and some high standard deviations.

### Taxonomic identification of total recruited transcripts

The taxonomy of the total filtered reads was first analyzed with the Kaiju tool which allowed us to identify between 9.06% and 58.87% of reads. The two samples with the fewest sequences identified (taxonomy or function) came from the suction sampler on “The Moose” site (C442_ASPI and C445_ASPI) and the two samples with the highest number of identified reads were from FISH samples on “The Moose” site (C480_FISH and C490_FISH). The number of unidentified reads was very high probably due to the absence of the *Rimicaris exoculata* or any closely related arthropod genomes in the databases, so MMseqs2 software was also used with the “taxonomy” function and the UniRef90 amino acid database on the recruited transcripts.

Unfortunately, this tool did not improve taxonomic identification even if it was more reliable (Table Supplementary data S8). As shown in Figure 6, there were 59.25% ± 3.02% of unidentified genes for the FISH sampler from “The Moose” site, 64.79% ± 3.38% for the FISH sampler from “The Beehive” site, 67.76% ± 2.68% for the PERISCOP and 74.26% ± 12.72% for the suction sampler. The two samples with the less identified genes came from the suction sampler and corresponded to the two RNA with poor quality, probably degraded (C443 and C445). Surprisingly, among the samples from the suction sampler, the C442 sample had the most genes identified with MMSeqs2 (35.6%) while it was the least recognized by Kaiju.

**Figure 6:**
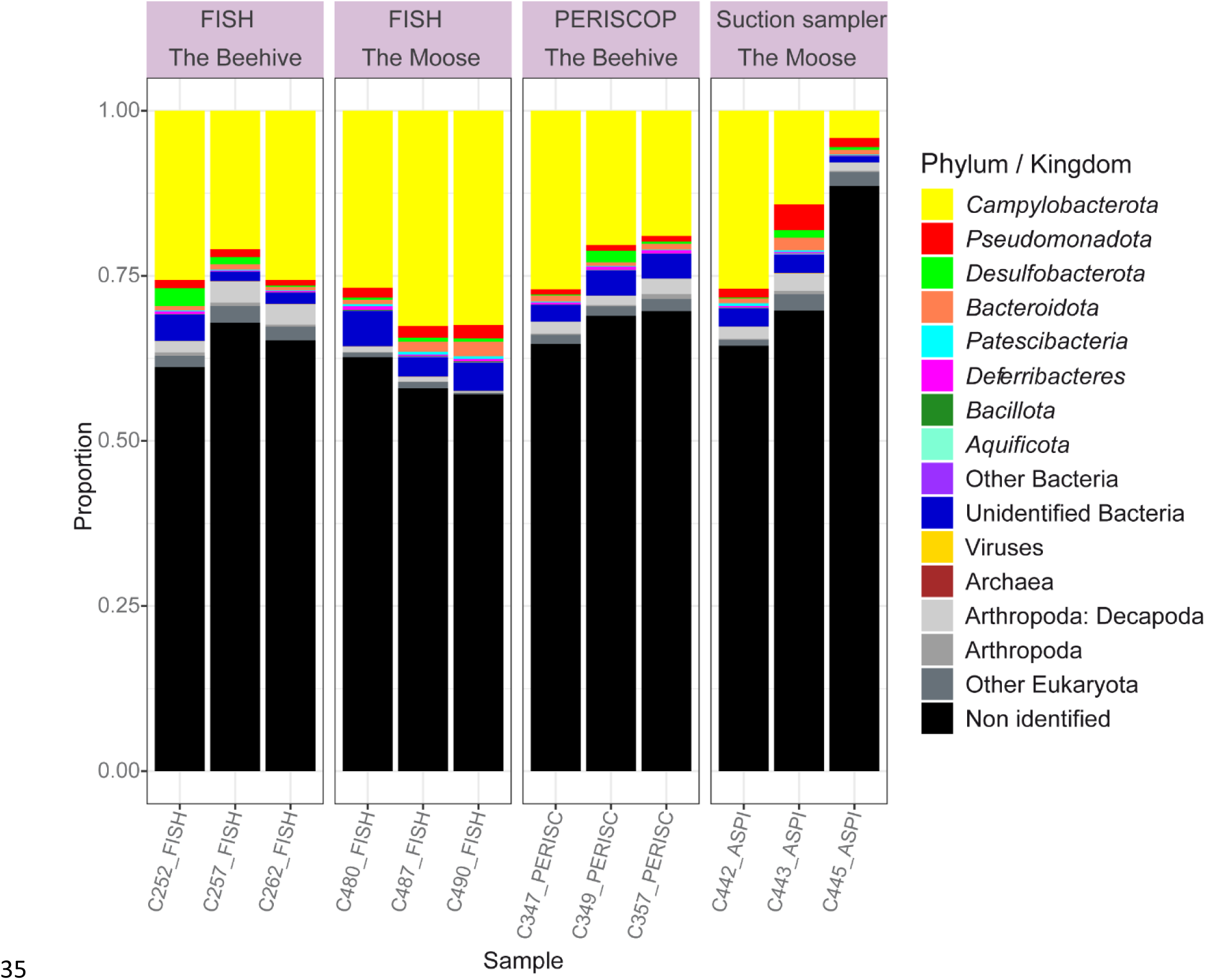
Barplot of taxonomic identification with MMSeqs2 of total recruited transcripts per sample grouped by tool and origin site.

Bacterial groups identified (Figure 6) were similar to those found in previous studies (Zbinden et al. 2008; Guri et al. 2012; Jan et al. 2014; Zbinden and Cambon Bonavita 2020; Cambon-Bonavita et al. 2021) with a majority of *Campylobacteria* representing a mean of 22.97% ± 7.99% of the sequences depending on the samples, followed by *Pseudomonadota (*ex-*Proteobacteria*) composed essentially of *Gammaproteobacteria*, and by *Desulfobacterota* and *Bacteroidota*. Sequences affiliated with Eukaryotes, among which some decapod or arthropod sequences were found but also protists and other Eukaryotes represent on average only 3.55% ± 1.71% of detected sequences.

### Distribution of gene expression data

To reduce biases, the principal coordinate analysis (PCoA) of the differential expression was carried out by separating not only the type of sampling tool but also the site of origin. After normalizing the distance matrix with the variance stabilizing transformation (VST) included in DESeq2 tool, the PCoA (Figure 7) revealed a separation of the data by sampling tool associated with the site of origin. This separation by sampling condition was statistically significant (PermANOVA, R2 = 0.40, Pr(>f) =3 x 10^-4^). Similarly, the separation observed on Axis 1 by site of origin was also statistically significant (PermANOVA, R2 = 0.17, Pr(>f) =0.0021). Finally, Axis 2 appeared to show a separation by RNA quality, which was statistically supported (PermANOVA, R2 = 0.158, Pr(>f) = 0.0312). Indeed, two samples from the suction sampler were isolated from the other points at the top of Figure 7 which corresponded to the two poor quality RNAs (C443 and C445 samples).

**Figure 7:**
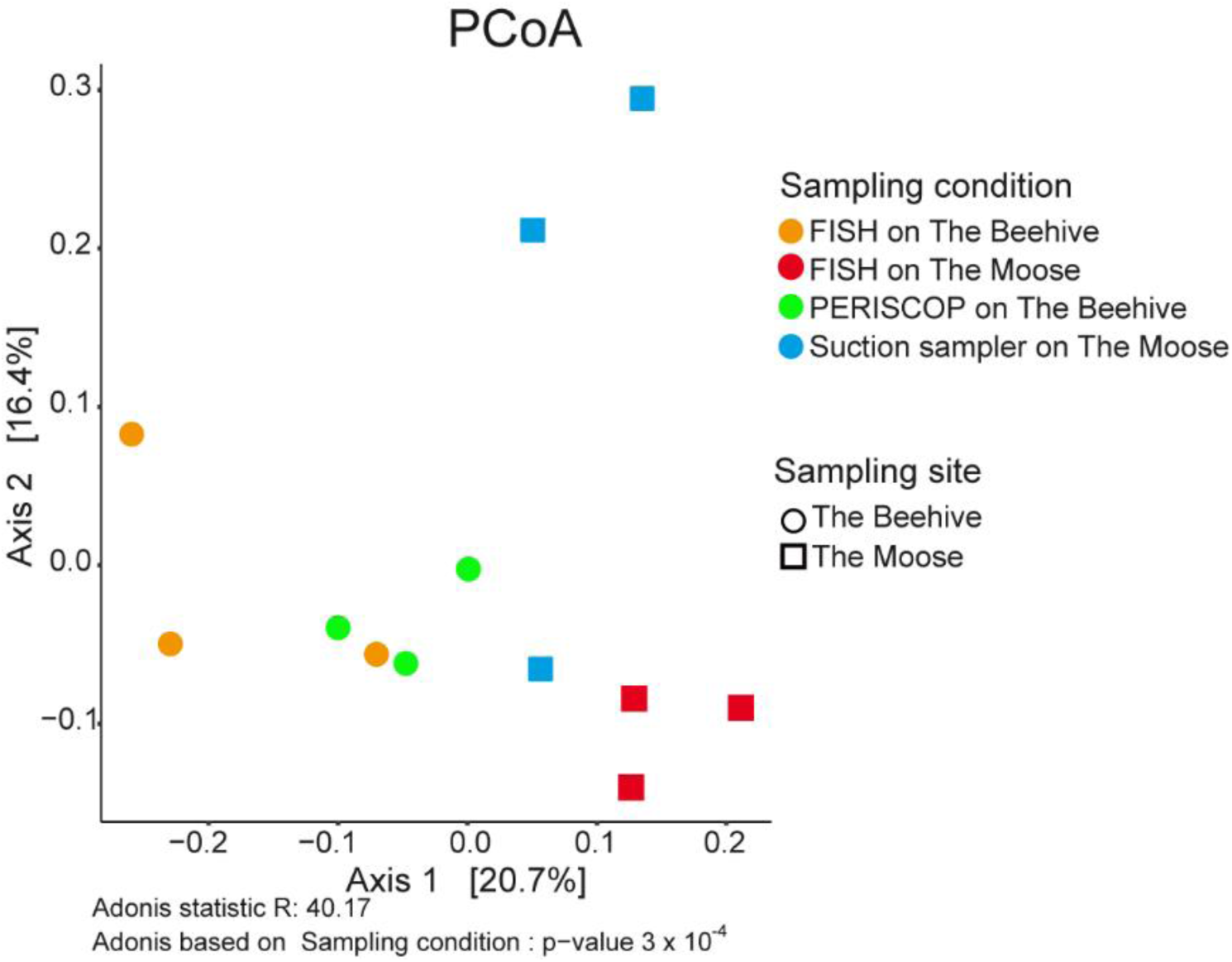
Principal coordinate analysis of gene expression after DeSEQ2 normalization. The samples are represented by colored dots according to the sampling tool associated with site of origin (orange FISH on “The Beehive”, red FISH on “The Moose”, green PERISCOP on “The Beehive” and blue suction sampler on “The Moose”) and by shape according to the site of origin (round for “The Beehive” and square for “The Moose”).

On the sampling site “The Beehive”, one PERISCOP sample was mixed with FISH samples. Gene expression therefore appeared to be more similar between samples taken by PERISCOP and those obtained after in situ stabilization of RNA with FISH tool on “The Beehive” site.

### Differential gene expression according to the type of sampling tool and station of origin

The differential expression analysis gave very variable results depending on the comparisons made. Between the suction sampler and FISH at “The Moose” site, there were 5741 different genes differentially expressed, of which 5025 were over-expressed and 716 were under-expressed in the suction sampler. In contrast, the comparison of expression profiles between the PERISCOP and FISH at “The Beehive” site yielded far fewer numbers of different genes differentially expressed, only 132 of which 81 were over-expressed and 51 were under-expressed with PERISCOP.

To identify genes, COG annotation was coupled with eggNOG-mapper annotation so the “COG20_category”was used with the “EGGNOG_COG_category” and “KEGG class” to compare the data. Of these differentially expressed genes, a very large proportion concerned unidentified genes: 48.6% of the different genes for the suction sampler with FISH comparison and 17.4% for PERISCOP with FISH comparison. Among these numerous unidentified genes, five of them were over-expressed in the suction sampler compared to FISH samples from “The Moose” and contained a very large number of reads (respectively 42 216, 82 883, 114 461, 2 481 748 and 4 259 592 baseMean) (Supplementary Table S9). The analysis of the remaining identified genes, which contained far fewer reads, was therefore difficult due to non-identified read excess. All unidentified genes were removed from Figure 8 to observe the differences in signals.

**Figure 8:**
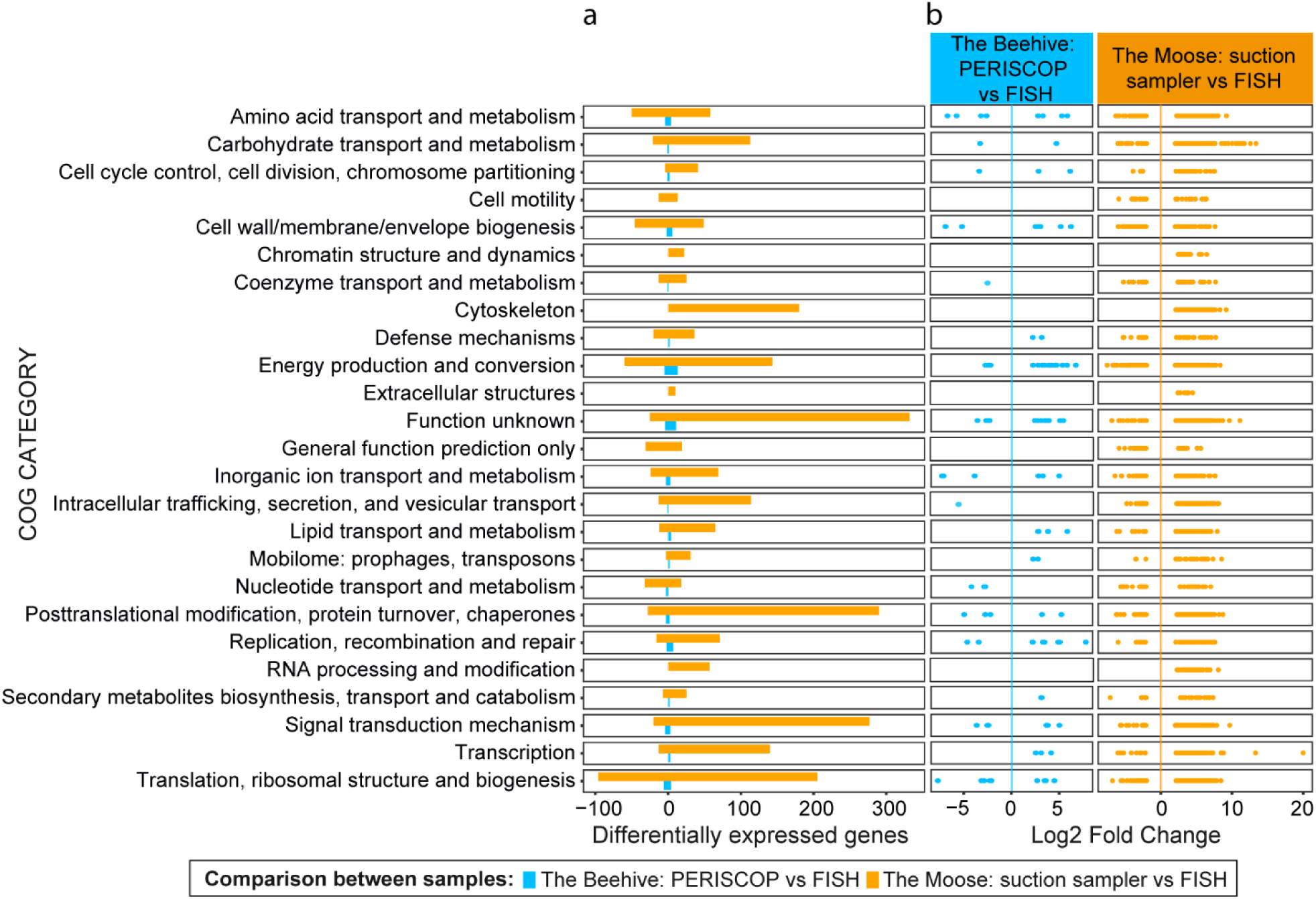
Distribution of differentially expressed genes by COG20 category and by comparative sampling conditions. Only values significantly different between each condition are shown, i.e. with Log2 Fold Change >2 or <-2 and with adjusted p-value <0.05. a: barplots represent sum of different over-expressed genes (abundance >0) and under-expressed genes (abundance <0). b: distribution of values of the Log2 Fold Change, which is a factor expressed on a logarithmic scale (base 2) and represents the difference in expression ratio between the two conditions.

When the suction sampler was compared to FISH (on the “The Moose” site), the categories with a greater variety of over-or under-expressed genes were very diverse (Figure 8a). These included genes involved in the mechanisms of cell synthesis (translation, ribosomal structure and biogenesis: 205 genes over-expressed and 96 under-expressed, transcription: 140 genes over-expressed and 13 under-expressed), in the posttranslational modifications (290 genes over-expressed and 28 under-expressed), in the signal transduction mechanisms (277 genes over-expressed and 20 under-expressed), genes involved in metabolism (carbohydrate transport and metabolism: 113 genes over-expressed and 21 under-expressed, energy production and conversion: 143 genes over-expressed and 60 under-expressed), and other types of genes such as those involved in cytoskeleton (180 genes over-expressed) or intracellular traffic (114 genes over-expressed and 13 under-expressed). In terms of intensity of expression, the greatest differences in expression (Log2 Fold change >10 or <-10, i.e. over-expressed or under-expressed by a factor of at least 2^10^ = 1024) were found in metabolic and transcriptional genes (Figure 8b). This would indicate cellular over-activity with accelerated turnover, probably due to stress, when the animals were collected with the suction sampler, compared with FISH.

As for the differences in genes expressed between sampling with PERISCOP and FISH (on “The Beehive” site), the number of different genes was smaller and mainly found in the energy production and conversion category (13 genes over-expressed and 5 under-expressed). The highest expression differentials were found in DNA replication (Log2 Fold Change = 7.787), translation and biogenesis (Log2 Fold Change = -7.710) and inorganic ion transport and metabolism (Log2 Fold change = -7.249).

## DISCUSSION

A new tool dedicated to in situ RNA preservation of deep-sea mobile fauna is described in the present study. In situ RNA tissue preservation for metatranscriptomic analysis using the new FISH sampler was assessed. The texture of shrimp tissue differed according to the sampling method: translucent and soft for living specimens collected using PERISCOP or suction sampler, and white and hard as they were “baked” for dead specimens recovered from our FISH sampler, suggesting that the RNA*later*^®^ penetrated deeply into the tissues. The exposure of samples to strong physicochemical variations (e.g. pressure, temperature, oxygen or hydrogen sulfide concentrations) before tissue preservation clearly affected the RNA quality as shown by the poor RIN values obtained on samples retrieved using the submersible suction sampler.

The PCoA suggested a separation of the two poor quality RNAs obtained from the suction sampler with the other samples. These samples also resulted in poorer taxonomic identification of transcripts. Given the lack of existing host genomic information in databases, this suggests an enrichment of shrimp sequences in our data at the expense of prokaryotic sequences. This was also supported by the differential expression profiles, which showed that most expressed genes could not be identified from suction sampler specimens. This may be due to chemical modifications, decompression and temperature increase suffered by the shrimp during ascent to the surface, which were recovered unhealthy. Moreover, some of the bacterial mRNA expressed in situ probably degraded during ascent, as they are much more unstable and shorter-lived than eukaryotic RNAs. Finally, unidentified shrimp genes such as stress-related genes may have become over-expressed. Furthermore, genes related to metabolism and posttranslational modifications were over-expressed compared to samples preserved in situ, also stressing the value of in situ preservation of tissue samples.

The data show a significant greater diversity of expressed genes from samples collected with the FISH sampler on “The Beehive” site of 34.71% compared with FISH on “The Moose” site and 60.01% compared with PERISCOP. The number of total recruited transcripts also seemed greater from samples collected with FISH on “The Moose” site (Figure 5), compared to the two other tools (+73.93% compared to suction sampler, +75.23% compared to PERISCOP, +56.83% compared to FISH on “The Beehive”). Unfortunately, due to large standard deviations between samples, statistical tests did not confirm that in situ RNA preservation had higher yields of recruited transcripts mapped on genes. However, statistical test results should be treated with caution, as they are based on only three points for each condition.

Fewer differences were observed between in situ RNA preservation with the FISH sampler and pressurized recovery with PERISCOP from the same site, as shown by the PCoA and differential gene expression analysis. Even if there are differences in “energy production and conversion”, in “inorganic ion transport and metabolism” or in “replication or translation”, ascent into the pressurized enclosure clearly limited the lethal effects of decompression, and possibly caused less disturbance in shrimp metabolism (Ravaux et al. 2019; Shillito et al. 2023). Additionally, PERISCOP’s syntactic foam casing also limited temperature exchange with the water column.

The shrimps were therefore kept in seawater around 15°C and at almost in situ pressure (Shillito et al. 2023). Shrimps were probably less stressed in these conditions (Ravaux et al. 2019) and cellular machinery did not run amok. It is also possible that the half-life of mRNAs was greater at this temperature, close to the natural habitat of shrimps, and at high pressure.

Some of the variance observed between results, in particular RNA concentrations, may have been biased due to the size of the shrimps. As tissues were not weighed before RNA extraction, the higher RNA concentrations obtained with specimens from “The Moose” site could be the consequence of the size of the organs harbored by these larger specimens. Hence, to limit potential sequencing bias, libraries were standardized in order to obtain the same sequenced quantities for each condition. Previous studies (Zbinden et al. 2008; Guri et al. 2012; Jan et al. 2014; Zbinden and Cambon Bonavita 2020; Cambon-Bonavita et al. 2021) show that the shrimp *Rimicaris exoculata* harbors a restricted diversified symbiotic community in the cephalothorax, compared with environmental communities. Symbionts colonize the shrimp’s cephalothorax as early as the juvenile stages, and persist throughout its life, whatever the stage, size or depth of the site of origin (Guéganton et al. 2024). In the present study, all symbiotic partners were retrieved in all samples, as revealed by the taxonomic identification of the expressed genes, indicating an overall homogeneous DNA/RNA extraction not impaired by shrimp size.

Another potential bias of the experimental design was that the number of RNA extracts was not identical for all sampling methods. As the sequencing platform requires a minimum of RIN, RNA, suction sampler specimens were extracted until at least three were obtained with the required RIN. However, only one specimen reached the required standard. If the same number of extractions had been applied in all conditions (i.e. six extractions), none from the suction sampler would have qualified for the platform, suggesting again that exposure of tissues to strong physicochemical variations strongly alter the RNA pool.

The differences observed in the diversity of genes expressed between “The Beehive” and “The Moose” using the FISH sampler could be a consequence of the metagenome assembly used to identify metatranscriptomes only from individuals from “The Beehive” site. On “The Moose” site, the environmental chemical conditions are slightly different (Konn et al. 2022), suggesting that metabolic activities could also be contrasted, potentially introducing some differences between sites. Due to technical constraints, the metagenomes were obtained different shrimps to those used for the metatranscriptomes, introducing a potential additional bias, whatever the sampling method.

Although a number of studies indicate the importance of preserving samples in situ to avoid transcriptional profile changes (Sanders et al. 2013; McQuillan and Robidart 2017; Gao et al. 2019; Sun et al. 2020; Poff et al. 2021), only a few deep-sea studies have compared mRNA datasets between in situ and on-board RNA stabilization methods. In the water column, Feike and colleagues (Feike et al. 2012) and Wurzbacher and collaborators (Wurzbacher et al. 2012) demonstrate a greater number of transcripts with in situ RNA preservation. The results obtained with the FISH sampler also seem to show a greater number of transcripts thanks to in situ RNA preservation, although these are not statistically supported. In contrast to Wurzbacher et al. (Wurzbacher et al. 2012), our results moreover demonstrated differences in the quality of RNA extracts.

Taxonomic and genetic diversity also seemed to be affected by RNA post-preservation on board the ship. For example, a metatranscriptomic study conducted on galathea *Shinkai crosnieri* (Motoki et al. 2020) showed a higher Shannon diversity of OTU with in situ RNA-stabilized samples compared to onboard RNA preservation. Our results also showed a similar trend, with a greater number of different transcripts in the samples preserved in situ than in those recovered with PERISCOP and post-preserved on board, stressing the need for in situ RNA preservation to maintain taxonomic and genetic diversity, Various studies have shown a significant difference in gene expression between in situ RNA preservation and the classical approach (Watsuji et al. 2014; Edgcomb et al. 2016; Olins et al. 2017; Motoki et al. 2020; Miyazaki et al. 2020). Quantitative RT-qPCR approaches used in some studies on symbiotic animal models such as setae of *S. crosnieri* (Watsuji et al. 2014) or gills of the gastropod *Alviniconcha marisindica* (Miyazaki et al. 2020) have demonstrated a higher abundance of some targeted genes like 16S rRNA gene transcripts or functional genes targeting different metabolic pathways for in situ RNA preservation. More holistic metatranscriptomic approaches have revealed variations in gene expression for different gene categories. The study by Motoki et al. on *S. crosnieri* (Motoki et al. 2020), for example, showed significantly different results on PCoA with Weighted Unifrac index between in situ RNA preservation and preservation on board. In the present study, the PCoA on shrimp *R. exoculata* with the Bray-Curtis dissimilarity matrix also showed the influence of the sampling method and the sampling site. Moreover, the use of the Microbial Sampler - In situ Incubation Device (MS-SID) (Edgcomb et al. 2016) highlighted classes of genes differentially expressed for some taxa when fixed in situ compared to samples with Niskin bottles and on-board conditioning. Similarly, the Olins and collaborators study (Olins et al. 2017) revealed statistically significant differences in the expression on genes regarding carbohydrates, RNA metabolism, stress response and fatty acids, lipids and isoprenoids, between the Deep-Sea Environmental Sample Processor (D-ESP) and Niskin bottles. Our results led to similar conclusions, with many genes differentially expressed between the FISH sampler and the suction sampler in different functional categories such as the mechanisms of cell synthesis, metabolism, genes involved in the cytoskeleton and intracellular traffic. Moreover, a greater number of unidentified transcripts were found in specimens sampled with the suction sampler. Various comparative studies have also shown that RNA post-preservation on board the ship leads to major variations in gene expression compared to in situ RNA preservation. This could also bias the relative abundance of some taxa as they could be differentially affected by their proper degradation kinetics of RNA. For example, it seems that rRNA and mRNA of some taxa such as *Methylococcales* and *Sulfurovum* were degraded faster than those of *Thiotrichales* (Motoki et al. 2020). Furthermore, depressurization during ascent causes DNA fragmentation or cell envelope rupture or, for some taxa like methanotrophic or methanogenic bacteria, the release of cell contents into the environment, which also biases DNA analyses (Park and Clark 2002; Chen et al. 2021). All these results showed the added value of in situ preservation to avoid expression shifts related to carbon and energy source depletion, and temperature and hydrostatic pressure changes.

The FISH sampler has been developed at a reasonable cost of *ca.* 6000€. It can be implemented on any submersible using its suction sampler and its hydraulic power system. It is easy to use, assemble/disassemble and clean, and limits the impact on living specimens by restricting sampling to 15-20 individuals. The FISH sampler benefits from the design of existing devices but with improvements of present functions to provide a complete new device for in situ RNA preservation of mobile fauna. A suction function has been added to the ISMACH sampler (Sanders et al. 2013) in order to collect highly mobile animals. Moreover, Miyasaki and collaborators highlighted the incomplete fixation of intracellular RNA of endosymbionts in the absence of gastropod homogenization (Miyazaki et al. 2020). The fixative solution did not reach the interior of the tissues inside the animal, hence the importance of associating a homogenization system. But it was important to develop a homogenization process preserving tissue structure. It was necessary to be able to separate the different organs without crushing the animal, unlike homogenization with ISMACH. In addition, transfer speed of the preservative reagent was improved from nine minutes with the Japanese diffusion system to less than ten seconds with FISH.

To facilitate the implementation of the FISH sampler on the submersible ROV Victor 6000, a future basket directly integrating the position of the FISH sampler and substation connections is under development. This will save time when installing the tool, and take up less space in the basket.

### CONCLUSION AND RECOMMENDATIONS

Obtaining a full deep-sea in situ picture of biological activities is still a challenge. Here, we presented a new sampling tool for in situ RNA preservation of mobile fauna and their associated symbionts in the deep-sea. The FISH sampler combines the benefits of existing systems to create a tool adapted to collect deep-sea mobile animals and efficiently preserve in situ their tissues.

Through metatranscriptomic approaches, differences of gene abundance and gene expression were investigated in the cephalothorax of the hydrothermal shrimp *Rimicaris exoculata* to compare this new sampler FISH to other methods. The results showed differences between in situ and on-board RNA stabilization, whether in terms of RNA quality, abundance of different or taxonomically identified genes and differential expression levels of genes.

The comparison between the samples collected using the submersible’s suction sampler and those collected using FISH revealed a greater number of differentially expressed genes than the comparison of the samples collected using FISH between two geochemically contrasted hydrothermal fields. Therefore we do not recommend the use of the fauna suction samplers developed on most submersibles for gene expression studies. On the other hand, RNA obtained with the PERISCOP pressurized recovery device were relatively comparable to those obtained with FISH, although the genes were less diversified leading to potential bias when interpreting actual in situ biological activities. The FISH sampler is therefore a quite basic and affordable tool, suitable for studies of gene expression using metatranscriptomic.

This work highlights the impact of the sampling tool on results obtained for metatranscriptomic approaches. In situ RNA preservation is key in identifying active members of deep-sea holobiont and characterizing their functions to expand our understanding of the microbiomes or host– symbiont in situ interactions (Lan et al. 2019). The FISH sampler will therefore allow us to compare samples collected from the same hydrothermal field, but which may differ in their gene expression due to different geochemical conditions in environmental microniches, such as the comparison between “The Moose” and “The Beehive” sites. The use of FISH could apply to other animals in other deep-sea environments, such as cold seeps, or animals associated with cold-water corals or abyssal trenches.

## Supporting information

Video of sampling process

## ACKNOWLEDGMENTS

We are grateful to the DSM team at Genavir for their guidance and expertise transfer on HOV Nautile and ROV Victor 6000 submersibles throughout the development of the FISH prototype. We thank the cruise chief scientists of ESSNAUT2017, ESSNAUT2021, ESSNAUT2022, ESSROV2019, HERMINE, BICOSE2 and CHUBACARC (J.-P. Justiniano, M.-A. Cambon, V. Ciausu, Y. Fouquet, S. Hourdez and D. Jollivet), and the captains and crew of R/V *Atalante* and *Pourquoi pas?*, HOV Nautile and ROV Victor 6000 for their technical and logistic assistance sample collection. Further thanks go to the INRAe GeT-PlaGE platform (get.genotoul.fr, Castanet-Tolosan, France) for metagenome and metatranscriptome sequencing. Special thanks to Blandine Trouche for her help with the bioinformatics analyses. We also express our thanks to P. Methou for proofreading and A. Chalm for English language edition.

Funding for the FISH project was provided by the Ifremer REMIMA program, the ANR Carnot EDROME 11 CARN 018-01, within the framework of the Ifremer DEEPECOS 2015 project and Ifremer Merlin project “Pourquoi pas les Abysses?”. Funding information for the PERISCOP device may be found in Shillito et al., 2023 (Shillito et al. 2023).

## Notes

### Competing Interest Statement

The authors have declared no competing interest.

## REFERENCES

1. Akerman, N. H., D. A. Butterfield, and J. A. Huber. 2013. Phylogenetic diversity and functional gene patterns of sulfur-oxidizing subseafloor Epsilonproteobacteria in diffuse hydrothermal vent fluids. Front Microbiol 4. doi:10.3389/fmicb.2013.00185

2. Andersson, A. F., M. Lundgren, S. Eriksson, M. Rosenlund, R. Bernander, and P. Nilsson. 2006. Global analysis of mRNA stability in the archaeon Sulfolobus. Genome Biology 7. doi:10.1186/gb-2006-7-10-r99

3. Baker, B., C. Sheik, C. Taylor, S. Jain, A. Bhasi, J. Cavalcoli, and G. Dick. 2013. Community transcriptomic assembly reveals microbes that contribute to deep-sea carbon and nitrogen cycling. ISME JOURNAL 7: 1962–1973. doi:10.1038/ismej.2013.85

4. Bashiardes, S., G. Zilberman-Schapira, and E. Elinav. 2016. Use of Metatranscriptomics in Microbiome Research. Bioinform Biol Insights 10: 19–25. doi:10.4137/BBI.S34610

5. Bernstein, J. A., A. B. Khodursky, P. H. Lin, S. Lin-Chao, and S. N. Cohen. 2002. Global analysis of mRNA decay and abundance in Escherichia coli at single-gene resolution using two-color fluorescent DNA microarrays. Proceedings of the National Academy of Sciences of the United States of America 99: 9697–9702. doi:10.1073/pnas.112318199

6. Bini, E., V. Dikshit, K. Dirksen, M. Drozda, and P. Blum. 2002. Stability of mRNA in the hyperthermophilic archaeon Sulfolobus solfataricus. Rna 8: 1129–1136. doi:10.1017/s1355838202021052

7. Breier, J. A., C. S. Sheik, D. Gomez-Ibanez, and others. 2014. A large volume particulate and water multi-sampler with in situ preservation for microbial and biogeochemical studies. Deep-Sea Research Part I-Oceanographic Research Papers 94: 195–206. doi:10.1016/j.dsr.2014.08.008

8. Cambon-Bonavita, M.-A., J. Aubé, V. Cueff-Gauchard, and J. Reveillaud. 2021. Niche partitioning in the *Rimicaris exoculata* holobiont: the case of the first symbiotic *Zetaproteobacteria*. Microbiome 9: 87. doi:10.1186/s40168-021-01045-6

9. Cantalapiedra, C. P., A. Hernández-Plaza, I. Letunic, P. Bork, and J. Huerta-Cepas. 2021. eggNOG-mapper v2: Functional Annotation, Orthology Assignments, and Domain Prediction at the Metagenomic Scale. Molecular Biology and Evolution 38: 5825–5829. doi:10.1093/molbev/msab293

10. Chen, H., M. Wang, M. Li, and others. 2021. A glimpse of deep-sea adaptation in chemosynthetic holobionts: Depressurization causes DNA fragmentation and cell death of methanotrophic endosymbionts rather than their deep-sea Bathymodiolinae host. Molecular Ecology 30: 2298– 2312. doi:10.1111/mec.15904

11. Clouet-d’Orval, B., M. Batista, M. Bouvier, Y. Quentin, G. Fichant, A. Marchfelder, and L.-K. Maier. 2018. Insights into RNA-processing pathways and associated RNA-degrading enzymes in Archaea. FEMS Microbiology Reviews 42: 579–613. doi:10.1093/femsre/fuy016

12. Connelly, D. P., J. T. Copley, B. J. Murton, and others. 2012. Hydrothermal vent fields and chemosynthetic biota on the world’s deepest seafloor spreading centre. Nat Commun 3: 620. doi:10.1038/ncomms1636

13. Cron, B., C. Sheik, F. Kafantaris, and others. 2020. Dynamic Biogeochemistry of the Particulate Sulfur Pool in a Buoyant Deep-Sea Hydrothermal Plume. ACS EARTH AND SPACE CHEMISTRY 4: 168–182. doi:10.1021/acsearthspacechem.9b00214

14. Dixon, P. 2003. VEGAN, a package of R functions for community ecology. Journal of Vegetation Science 14: 927–930. doi:10.1111/j.1654-1103.2003.tb02228.x

15. Dubilier, N., C. Bergin, and C. Lott. 2008. Symbiotic diversity in marine animals: the art of harnessing chemosynthesis. Nat Rev Microbiol 6: 725–740. doi:10.1038/nrmicro1992

16. Edgcomb, V. P., C. Taylor, M. G. Pachiadaki, S. Honjo, I. Engstrom, and M. Yakimov. 2016. Comparison of Niskin vs. in situ approaches for analysis of gene expression in deep Mediterranean Sea water samples. Deep-Sea Res Pt Ii 129: 213–222. doi:10.1016/j.dsr2.2014.10.020

17. Edri, S., and T. Tuller. 2014. Quantifying the Effect of Ribosomal Density on mRNA Stability. PLoS One 9. doi:10.1371/journal.pone.0102308

18. Eren, A. M., Ö. C. Esen, C. Quince, J. H. Vineis, H. G. Morrison, M. L. Sogin, and T. O. Delmont. 2015. Anvi’o: an advanced analysis and visualization platform for ’omics data. PeerJ 3: e1319. doi:10.7717/peerj.1319

19. Evguenieva-Hackenberg, E., and G. Klug. 2011. New aspects of RNA processing in prokaryotes. Current Opinion in Microbiology 14: 587–592. doi:10.1016/j.mib.2011.07.025

20. Feike, J., K. Jurgens, J. T. Hollibaugh, S. Kruger, G. Jost, and M. Labrenz. 2012. Measuring unbiased metatranscriptomics in suboxic waters of the central Baltic Sea using a new in situ fixation system. ISME J 6: 461–470. doi:10.1038/ismej.2011.94

21. Fortunato, C., B. Larson, D. Butterfield, and J. Huber. 2018. Spatially distinct, temporally stable microbial populations mediate biogeochemical cycling at and below the seafloor in hydrothermal vent fluids. ENVIRONMENTAL MICROBIOLOGY 20: 769–784. doi:10.1111/1462-2920.14011

22. Fortunato, C. S., and J. A. Huber. 2016. Coupled RNA-SIP and metatranscriptomics of active chemolithoautotrophic communities at a deep-sea hydrothermal vent. ISME J 10: 1925–1938. doi:10.1038/ismej.2015.258

23. Fouquet, Y., W. Amina, P. Cambon, C. Mevel, G. Meyer, and P. Gente. 1993. Tectonic setting and mineralogical and geochemical zonation in the Snake Pit sulfide deposit (Mid-Atlantic Ridge at 23 degrees N). Economic Geology 88: 2018–2036. doi:10.2113/gsecongeo.88.8.2018

24. Galperin, M. Y., Y. I. Wolf, K. S. Makarova, R. Vera Alvarez, D. Landsman, and E. V. Koonin. 2021. COG database update: focus on microbial diversity, model organisms, and widespread pathogens. Nucleic Acids Res 49: D274–D281. doi:10.1093/nar/gkaa1018

25. Gao, Z., J. Huang, G. Cui, and others. 2019. In situ meta-omic insights into the community compositions and ecological roles of hadal microbes in the Mariana Trench. ENVIRONMENTAL MICROBIOLOGY 21: 4092–4108. doi:10.1111/1462-2920.14759

26. Govindarajan, A. F., J. Pineda, M. Purcell, and J. A. Breier. 2015. Species- and stage-specific barnacle larval distributions obtained from AUV sampling and genetic analysis in Buzzards Bay, Massachusetts, USA. Journal of Experimental Marine Biology and Ecology 472: 158–165. doi:10.1016/j.jembe.2015.07.012

27. Guéganton, M., P. Methou, J. Aubé, and others. 2024. Symbiont acquisition strategies in post-settlement stages of two co-occurring deep-sea Rimicaris shrimp. doi:10.22541/au.171309964.46362343/v1

28. Guri, M., L. Durand, V. Cueff-Gauchard, M. Zbinden, P. Crassous, B. Shillito, and M. A. Cambon-Bonavita. 2012. Acquisition of epibiotic bacteria along the life cycle of the hydrothermal shrimp *Rimicaris exoculata*. ISME J 6: 597–609. doi:DOI 10.1038/ismej.2011.133

29. Hambraeus, G., C. von Wachenfeldt, and L. Hederstedt. 2003. Genome-wide survey of mRNA half-lives in Bacillus subtilis identifies extremely stable mRNAs. Mol Gen Genomics 269: 706–714. doi:10.1007/s00438-003-0883-6

30. He, Y., X. Y. Feng, J. Fang, Y. Zhang, and X. Xiao. 2015. Metagenome and Metatranscriptome Revealed a Highly Active and Intensive Sulfur Cycle in an Oil-Immersed Hydrothermal Chimney in Guaymas Basin. Front Microbiol 6. doi:10.3389/fmicb.2015.01236

31. Hennigan, A. N., and J. N. Reeve. 1994. Messenger-RNAs in the methanogenic archaeon Methanococcus vannielii - numbers, half-lives and processing. Molecular Microbiology 11: 655–670. doi:10.1111/j.1365-2958.1994.tb00344.x

32. Hongo, Y., T. Ikuta, Y. Takaki, S. Shimamura, S. Shigenobu, T. Maruyama, and T. Yoshida. 2016. Expression of genes involved in the uptake of inorganic carbon in the gill of a deep-sea vesicomyid clam harboring intracellular thioautotrophic bacteria. Gene 585: 228–240. doi:10.1016/j.gene.2016.03.033

33. Huerta-Cepas, J., K. Forslund, L. P. Coelho, D. Szklarczyk, L. J. Jensen, C. von Mering, and P. Bork. 2017. Fast Genome-Wide Functional Annotation through Orthology Assignment by eggNOG-Mapper. Molecular Biology and Evolution 34: 2115–2122. doi:10.1093/molbev/msx148

34. Hyatt, D., G. L. Chen, P. F. LoCascio, M. L. Land, F. W. Larimer, and L. J. Hauser. 2010. Prodigal: prokaryotic gene recognition and translation initiation site identification. Bmc Bioinformatics 11. doi:Artn 119 10.1186/1471-2105-11-119

35. Jäger, A., R. Samorski, F. Pfeifer, and G. Klug. 2002. Individual gvp transcript segments in Haloferax mediterranei exhibit varying half-lives, which are differentially affected by salt concentration and growth phase. Nucleic Acids Research 30: 5436–5443. doi:10.1093/nar/gkf699

36. Jan, C., J. M. Petersen, J. Werner, and others. 2014. The gill chamber epibiosis of deep-sea shrimp *Rimicaris exoculata*: an in-depth metagenomic investigation and discovery of *Zetaproteobacteria*. Environ Microbiol 16: 2723–2738. doi:10.1111/1462-2920.12406

37. Jiang, Y., X. Xiong, J. Danska, and J. Parkinson. 2016. Metatranscriptomic analysis of diverse microbial communities reveals core metabolic pathways and microbiome-specific functionality. MICROBIOME 4. doi:10.1186/s40168-015-0146-x

38. Kassambara, A. 2022a. ggpubr: “ggplot2” Based Publication Ready Plots.

39. Kassambara, A. 2022b. rstatix: Pipe-Friendly Framework for Basic Statistical Tests.

40. Kemp, P. F., S. Lee, and J. LaRoche. 1993. Estimating the Growth Rate of Slowly Growing Marine Bacteria from RNA Content. Applied and Environmental Microbiology 59: 2594–2601. doi:10.1128/aem.59.8.2594-2601.1993

41. Kerkhof, L., and P. Kemp. 1999. Small ribosomal RNA content in marine Proteobacteria during non-steady-state growth. FEMS Microbiology Ecology 30: 253–260. doi:10.1111/j.1574-6941.1999.tb00653.x

42. Kerkhof, L., and B. B. Ward. 1993. Comparison of Nucleic Acid Hybridization and Fluorometry for Measurement of the Relationship between RNA/DNA Ratio and Growth Rate in a Marine Bacterium. Applied and Environmental Microbiology 59: 1303–1309. doi:10.1128/aem.59.5.1303-1309.1993

43. Konn, C., J. P. Donval, V. Guyader, Y. Germain, A.-S. Alix, E. Roussel, and O. Rouxel. 2022. Extending the dataset of fluid geochemistry of the Menez Gwen, Lucky Strike, Rainbow, TAG and Snake Pit hydrothermal vent fields: Investigation of temporal stability and organic contribution. Deep Sea Research Part I: Oceanographic Research Papers 179: 103630. doi:10.1016/j.dsr.2021.103630

44. Köster, J., and S. Rahmann. 2018. Snakemake-a scalable bioinformatics workflow engine (vol 28, pg 2520, 2012). Bioinformatics 34: 3600–3600. doi:10.1093/bioinformatics/bty350

45. Kramer, J. G., and F. L. Singleton. 1993. Measurement of rRNA Variations in Natural Communities of Microorganisms on the Southeastern U.S. Continental Shelf. Applied and Environmental Microbiology 59: 2430–2436. doi:10.1128/aem.59.8.2430-2436.1993

46. Lan, Y., J. Sun, C. Chen, and others. 2021. Hologenome analysis reveals dual symbiosis in the deep-sea hydrothermal vent snail Gigantopelta aegis. Nat Commun 12: 1165. doi:10.1038/s41467-021-21450-7

47. Lan, Y., J. Sun, W. P. Zhang, and others. 2019. Host-Symbiont Interactions in Deep-Sea Chemosymbiotic Vesicomyid Clams: Insights From Transcriptome Sequencing. Frontiers in Marine Science 6. doi:Artn 680 10.3389/Fmars.2019.00680

48. Langmead, B., and S. L. Salzberg. 2012. Fast gapped-read alignment with Bowtie 2. Nat Methods 9: 357–359. doi:10.1038/nmeth.1923

49. Lavelle, A., and H. Sokol. 2018. Beyond metagenomics, metatranscriptomics illuminates microbiome functionality in IBD. Nat Rev Gastroenterol Hepatol 15: 193–194. doi:10.1038/nrgastro.2018.15

50. Lee, S., and P. F. Kemp. 1994. Single-cell RNA content of natural marine planktonic bacteria measured by hybridization with multiple 16S rRNA-targeted fluorescent probes. Limnology and Oceanography 39: 869–879. doi:10.4319/lo.1994.39.4.0869

51. Lesniewski, R. A., S. Jain, K. Anantharaman, P. D. Schloss, and G. J. Dick. 2012. The metatranscriptome of a deep-sea hydrothermal plume is dominated by water column methanotrophs and lithotrophs. ISME Journal 6: 2257–2268. doi:10.1038/ismej.2012.63

52. Li, D. H., C. M. Liu, R. B. Luo, K. Sadakane, and T. W. Lam. 2015. MEGAHIT: an ultra-fast single-node solution for large and complex metagenomics assembly via succinct de Bruijn graph. Bioinformatics 31: 1674–1676. doi:10.1093/bioinformatics/btv033

53. Li, H., B. Handsaker, A. Wysoker, and others. 2009. The Sequence Alignment/Map format and SAMtools. Bioinformatics 25: 2078–2079. doi:10.1093/bioinformatics/btp352

54. Li, M., S. Jain, and G. Dick. 2016. Genomic and Transcriptomic Resolution of Organic Matter Utilization Among Deep-sea Bacteria in Guaymas Basin Hydrothermal Plumes. FRONTIERS IN MICROBIOLOGY 7. doi:10.3389/fmicb.2016.01125

55. Love, M. I., W. Huber, and S. Anders. 2014. Moderated estimation of fold change and dispersion for RNA-seq data with DESeq2. Genome Biology 15. doi:Artn 550 10.1186/S13059-014-0550-8

56. Martin, M. 2011. Cutadapt removes adapter sequences from high-throughput sequencing reads. EMBnet.journal 17: 3. doi:10.14806/ej.17.1.200

57. Massoth, G., J. Puzic, P. Crowhurst, M. White, K. Nakamura, S. Walker, and E. Baker. 2008. Regional Venting in the Manus Basin, New Britain Back Arc. AGU Fall Meeting Abstracts.

58. Mat, A. M., J. Sarrazin, G. V. Markov, and others. 2020. Biological rhythms in the deep-sea hydrothermal mussel Bathymodiolus azoricus. Nat Commun 11: 3454. doi:10.1038/s41467-020-17284-4

59. McMurdie, P. J., and S. Holmes. 2013. phyloseq: An R Package for Reproducible Interactive Analysis and Graphics of Microbiome Census Data. PLoS One 8. doi:ARTN e61217 10.1371/journal.pone.0061217

60. McQuillan, J. S., and J. C. Robidart. 2017. Molecular-biological sensing in aquatic environments: recent developments and emerging capabilities. Curr Opin Biotech 45: 43–50. doi:10.1016/j.copbio.2016.11.022

61. Menke, S., M. A. F. Gillingham, K. Wilhelm, and S. Sommer. 2017. Home-Made Cost Effective Preservation Buffer Is a Better Alternative to Commercial Preservation Methods for Microbiome Research. Front. Microbiol. 8. doi:10.3389/fmicb.2017.00102

62. Menzel, P., K. L. Ng, and A. Krogh. 2016. Fast and sensitive taxonomic classification for metagenomics with Kaiju. Nat Commun 7: 11257. doi:10.1038/ncomms11257

63. Methou, P. 2019. Lifecycles of two hydrothermal vent shrimps from the Mid-Atlantic Ridge: *Rimicaris exoculata* and *Rimicaris chacei -* Embryonic development, Larva dispersal, Recruitment, Reproduction & Symbioses Acquisition. Université Bretagne Occidentale.

64. Miyazaki, J., T. Ikuta, T. Watsuji, and others. 2020. Dual energy metabolism of the Campylobacterota endosymbiont in the chemosynthetic snail Alviniconcha marisindica. ISME J 14: 1273–1289. doi:10.1038/s41396-020-0605-7

65. Mohanty, B. K., and S. R. Kushner. 2016. Regulation of mRNA Decay in Bacteria. Annu Rev Microbiol 70: 25–44. doi:10.1146/annurev-micro-091014-104515

66. Moran, M. A., B. Satinsky, S. M. Gifford, and others. 2013. Sizing up metatranscriptomics. ISME J 7: 237–243. doi:10.1038/ismej.2012.94

67. Motoki, K., T. Watsuji, Y. Takaki, K. Takai, W. Iwasaki, and J.-B. Raina. 2020. Metatranscriptomics by *In Situ* RNA Stabilization Directly and Comprehensively Revealed Episymbiotic Microbial Communities of Deep-Sea Squat Lobsters. Msystems 5: e00551–20. doi:doi:10.1128/mSystems.00551-20

68. Mutter, G. L., D. Zahrieh, C. M. Liu, D. Neuberg, D. Finkelstein, H. E. Baker, and J. A. Warrington. 2004. Comparison of frozen and RNALater solid tissue storage methods for use in RNA expression microarrays. Bmc Genomics 5. doi:Artn 88 10.1186/1471-2164-5-88

69. Ogle, D. H., J. C. Doll, W. A. Powell, and A. Dinno. 2023. FSA: Simple Fisheries Stock Assessment Methods.

70. O’Hara, E. B., J. A. Chekanova, C. A. Ingle, Z. R. Kushner, E. Peters, and S. R. Kushner. 1995. Polyadenylylation helps regulate mRNA decay in Escherichia coli. Proc Natl Acad Sci U S A 92: 1807–1811.

71. Olins, H. C., D. R. Rogers, C. Preston, and others. 2017. Co-registered Geochemistry and Metatranscriptomics Reveal Unexpected Distributions of Microbial Activity within a Hydrothermal Vent Field. Front Microbiol 8. doi:Artn 1042 10.3389/Fmicb.2017.01042

72. Ondov, B. D., N. H. Bergman, and A. M. Phillippy. 2011. Interactive metagenomic visualization in a Web browser. Bmc Bioinformatics 12: 385. doi:10.1186/1471-2105-12-385

73. Page, T. M., and J. W. Lawley. 2022. The Next Generation Is Here: A Review of Transcriptomic Approaches in Marine Ecology. Front. Mar. Sci. 9. doi:10.3389/fmars.2022.757921

74. Park, C. B., and D. S. Clark. 2002. Rupture of the Cell Envelope by Decompression of the Deep-Sea Methanogen Methanococcus jannaschii. Appl. Environ. Microbiol. 68: 1458–1463.

75. Pernthaler, A., and R. Amann. 2004. Simultaneous fluorescence in situ hybridization of mRNA and rRNA in environmental bacteria. Applied And Environmental Microbiology 70: 5426–5433.

76. Perwez, T., and S. R. Kushner. 2006. RNase Z in Escherichia coli plays a significant role in mRNA decay. Molecular Microbiology 60: 723–737. doi:10.1111/j.1365-2958.2006.05124.x

77. Pilhofer, M., M. Pavlekovic, N. M. Lee, W. Ludwig, and K. H. Schleifer. 2009. Fluorescence in situ hybridization for intracellular localization of nifH mRNA. Systematic and Applied Microbiology 32: 186–92.

78. Poff, K., A. Leu, J. Eppley, D. Karl, and E. DeLong. 2021. Microbial dynamics of elevated carbon flux in the open ocean’s abyss. PROCEEDINGS OF THE NATIONAL ACADEMY OF SCIENCES OF THE UNITED STATES OF AMERICA 118. doi:10.1073/pnas.2018269118

79. Posit team. 2024. RStudio: Integrated Development Environment for R.

80. R Core Team. 2024. R: A Language and Environment for Statistical Computing.

81. Rauhut, R., and G. Klug. 1999. mRNA degradation in bacteria. FEMS Microbiology Reviews 23: 353–370.

82. Ravaux, J., N. Leger, G. Hamel, and B. Shillito. 2019. Assessing a species thermal tolerance through a multiparameter approach: the case study of the deep-sea hydrothermal vent shrimp *Rimicaris exoculata*. Cell Stress Chaperones 24: 647–659. doi:10.1007/s12192-019-01003-0

83. Redon, E., P. Loubière, and M. Cocaign-Bousquet. 2005. Role of mRNA Stability during Genome-wide Adaptation of Lactococcus lactis to Carbon Starvation*. Journal of Biological Chemistry 280: 36380–36385. doi:10.1074/jbc.M506006200

84. Rubin-Blum, M., C. Antony, C. Borowski, and others. 2017. Short-chain alkanes fuel mussel and sponge Cycloclasticus symbionts from deep-sea gas and oil seeps. NATURE MICROBIOLOGY 2. doi:10.1038/nmicrobiol.2017.93

85. Rubin-Blum, M., C. Antony, L. Sayavedra, and others. 2019. Fueled by methane: deep-sea sponges from asphalt seeps gain their nutrition from methane-oxidizing symbionts. ISME JOURNAL 13: 1209– 1225. doi:10.1038/s41396-019-0346-7

86. Salehi, Z., and M. Najafi. 2014. RNA Preservation and Stabilization. Biochem Physiol 3: 126. doi:doi:10.4172/2168-9652.1000126

87. Sanders, J. G., R. A. Beinart, F. J. Stewart, E. F. Delong, and P. R. Girguis. 2013. Metatranscriptomics reveal differences in in situ energy and nitrogen metabolism among hydrothermal vent snail symbionts. ISME J 7: 1556–1567. doi:10.1038/ismej.2013.45

88. Schroeder, A., O. Mueller, S. Stocker, and others. 2006. The RIN: an RNA integrity number for assigning integrity values to RNA measurements. BMC Molecular Biology 7: 3. doi:10.1186/1471-2199-7-3

89. Shakya, M., C.-C. Lo, and P. S. G. Chain. 2019. Advances and Challenges in Metatranscriptomic Analysis. Frontiers in Genetics 10: 904. doi:10.3389/fgene.2019.00904

90. Shillito, B., L. Amand, and G. Hamel. 2023. Update of the PERISCOP system for isobaric sampling of deep-sea fauna. Deep Sea Research Part I: Oceanographic Research Papers 193: 103956. doi:10.1016/j.dsr.2022.103956

91. Shillito, B., G. Hamel, C. Duchi, D. Cottin, J. Sarrazin, P. M. Sarradin, J. Ravaux, and F. Gaill. 2008. Live capture of megafauna from 2300m depth, using a newly designed Pressurized Recovery Device. Deep Sea Res. (I Oceanogr. Res. Pap.) 55: 881–889. doi:10.1016/j.dsr.2008.03.010

92. Steglich, C., D. Lindell, M. Futschik, T. Rector, R. Steen, and S. W. Chisholm. 2010. Short RNA half-lives in the slow-growing marine cyanobacterium Prochlorococcus. Genome Biology 11. doi:10.1186/gb-2010-11-5-r54

93. Steinegger, M., and J. Söding. 2017. MMseqs2 enables sensitive protein sequence searching for the analysis of massive data sets. Nat Biotechnol 35: 1026–1028. doi:10.1038/nbt.3988

94. Steiner, P. A., D. De Corte, J. Geijo, C. Mena, T. Yokokawa, T. Rattei, G. J. Herndl, and E. Sintes. 2019. Highly variable mRNA half-life time within marine bacterial taxa and functional genes. Environmental Microbiology 21: 3873–3884. 10.1111/1462-2920.14737

95. Stewart, F. 2013. Preparation of Microbial Community cDNA for Metatranscriptomic Analysis in Marine Plankton, p. 187–218. *In* E. DeLong [ed.], MICROBIAL METAGENOMICS, METATRANSCRIPTOMICS, AND METAPROTEOMICS.

96. Sun, J., C. Chen, N. Miyamoto, and others. 2020. The Scaly-foot Snail genome and implications for the origins of biomineralised armour. Nat Commun 11: 1657. doi:10.1038/s41467-020-15522-3

97. Takayama, K., and S. Kjelleberg. 2000. The role of RNA stability during bacterial stress responses and starvation. Environmental Microbiology 2: 355–365. 10.1046/j.1462-2920.2000.00119.x

98. Takishita, K., Y. Takaki, Y. Chikaraishi, T. Ikuta, G. Ozawa, T. Yoshida, N. Ohkouchi, and K. Fujikura. 2017. Genomic Evidence that Methanotrophic Endosymbionts Likely Provide Deep-Sea Bathymodiolus Mussels with a Sterol Intermediate in Cholesterol Biosynthesis. Genome Biol Evol 9: 1148–1160. doi:10.1093/gbe/evx082

99. Taylor, C. D., V. P. Edgcomb, K. W. Doherty, I. Engstrom, T. Shanahan, M. G. Pachiadaki, S. J. Molyneaux, and S. Honjo. 2015. Fixation filter, device for the rapid in situ preservation of particulate samples. Deep-Sea Research Part I-Oceanographic Research Papers 96: 69–79. doi:10.1016/j.dsr.2014.09.006

100. Tourrière, H., K. Chebli, and J. Tazi. 2002. mRNA degradation machines in eukaryotic cells. Biochimie 84: 821–837. doi:10.1016/S0300-9084(02)01445-1

101. Watsuji, T. O., A. Yamamoto, Y. Takaki, K. Ueda, S. Kawagucci, and K. Takai. 2014. Diversity and methane oxidation of active epibiotic methanotrophs on live Shinkaia crosnieri. ISME J 8: 1020–1031. doi:10.1038/ismej.2013.226

102. Wickham, H. 2009. ggplot2: Elegant Graphics for Data Analysis, Springer.

103. Wickham, H., M. Averick, J. Bryan, and others. 2019. Welcome to the Tidyverse. Journal of Open Source Software 4: 1686. doi:10.21105/joss.01686

104. Wu, J., W. Gao, R. Johnson, W. Zhang, and D. Meldrum. 2013. Integrated Metagenomic and Metatranscriptomic Analyses of Microbial Communities in the Meso- and Bathypelagic Realm of North Pacific Ocean. MARINE DRUGS 11: 3777–3801. doi:10.3390/md11103777

105. Wu, J. Y., W. M. Gao, W. W. Zhang, and D. R. Meldrum. 2011. Optimization of whole-transcriptome amplification from low cell density deep-sea microbial samples for metatranscriptomic analysis. Journal of Microbiological Methods 84: 88–93. doi:10.1016/j.mimet.2010.10.018

106. Wurzbacher, C., I. Salka, and H. P. Grossart. 2012. Environmental actinorhodopsin expression revealed by a new in situ filtration and fixation sampler. Environmental Microbiology Reports 4: 491–497. doi:10.1111/j.1758-2229.2012.00350.x

107. Zbinden, M., and M.-A. Cambon Bonavita. 2020. *Rimicaris exoculata*: biology and ecology of a shrimp from deep-sea hydrothermal vents associated with ectosymbiotic bacteria. Marine Ecology Progress Series 652: 187–222. 10.3354/meps13467

108. Zbinden, M., B. Shillito, N. Le Bris, C. D. de Montlaur, E. Roussel, F. Guyot, F. Gaill, and M.-A. Cambon-Bonavita. 2008. New insigths on the metabolic diversity among the epibiotic microbial communitiy of the hydrothermal shrimp *Rimicaris exoculata*. Journal of Experimental Marine Biology and Ecology 359: 131–140. doi:10.1016/j.jembe.2008.03.009

109. Zhu, F.-C., J. Sun, G.-Y. Yan, J.-M. Huang, C. Chen, and L.-S. He. 2020. Insights into the strategy of micro-environmental adaptation: Transcriptomic analysis of two alvinocaridid shrimps at a hydrothermal vent. PLOS ONE 15: e0227587. doi:10.1371/journal.pone.0227587

110. Zilber-Rosenberg, I., and E. Rosenberg. 2008. Role of microorganisms in the evolution of animals and plants: the hologenome theory of evolution. FEMS Microbiology Reviews 32: 723–735. doi:10.1111/j.1574-6976.2008.00123.x

