## Supplemental figures for "FISH, a new tool for in situ preservation of RNA in tissues of deep-sea mobile fauna"

a

b

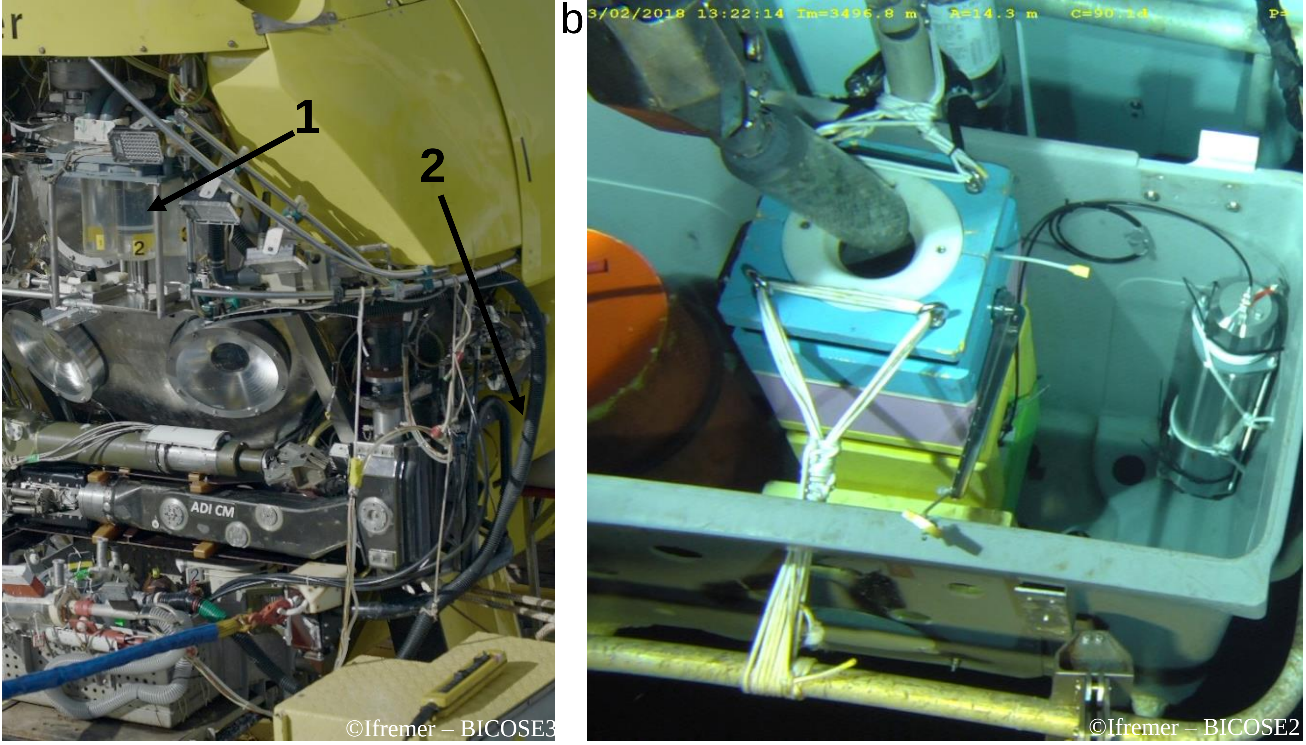


**Figure S1 supplementary data: photographs of two sampling devices. (a) front of the HOV Nautile submersible with 1- suction sampler bowls and 2- suction hose, (b) Periscopette being inserted with submersible arm inside PERISCOP placed in an independent shuttle device.**

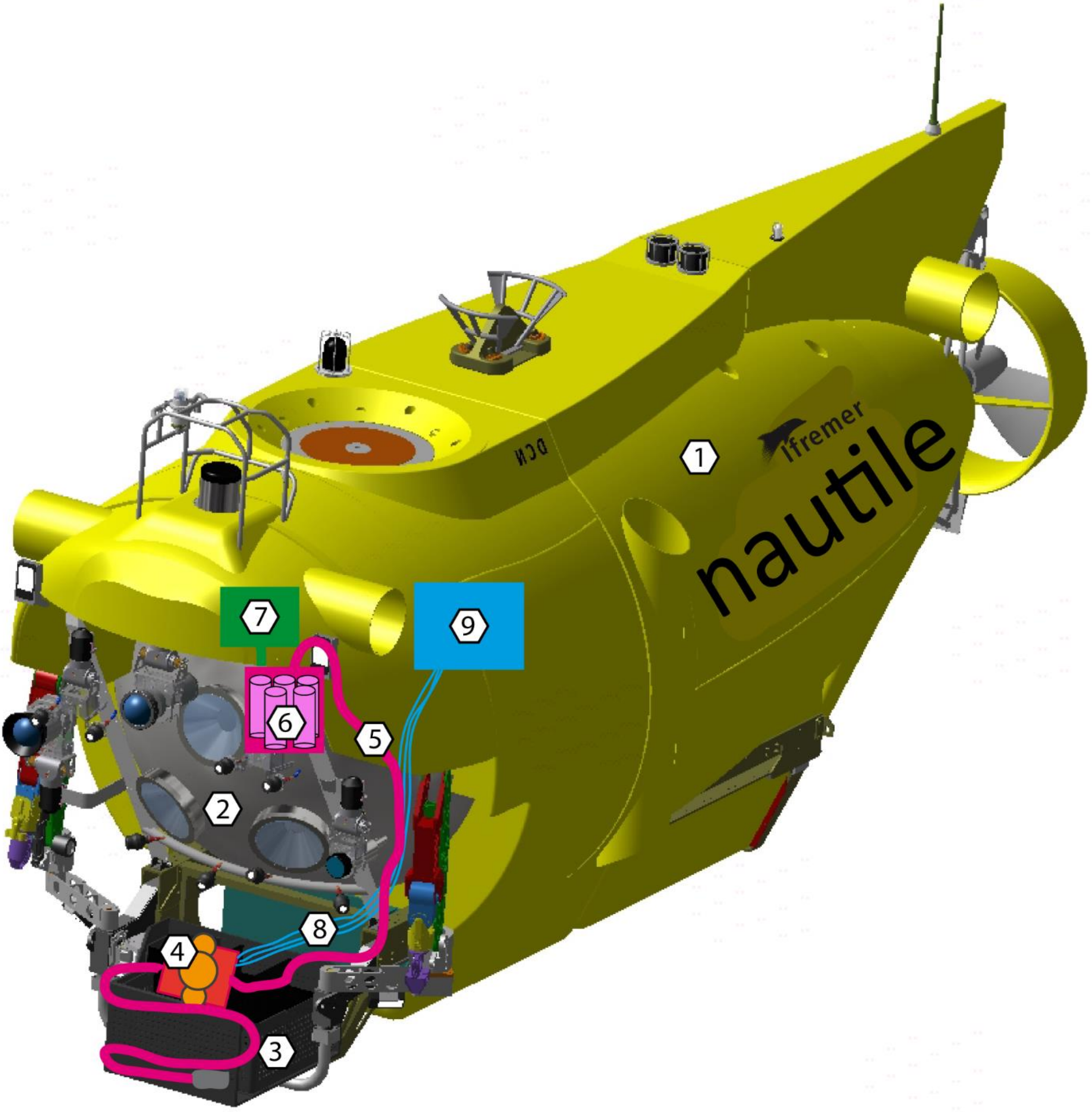


**Figure S2 supplementary data: schematic representation of the FISH system implementation on the submersible and the various connections made. 1: Submersible, 2: Front of submersible, 3: Basket of submersible, 4: FISH system, 5: suction hose, 6: bowls of suction sampler, 7: pump of suction sampler, 8: hydraulic hoses, 9: hydraulic power supply of submersible. Nautile drawing courtesy of V Ciausu (DFO/SM).**

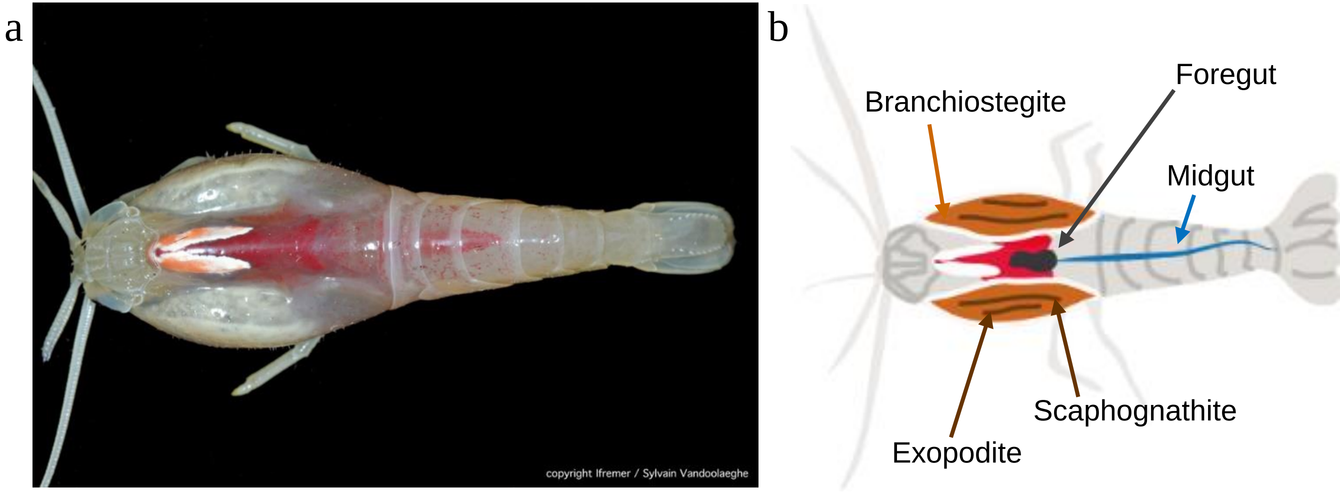


**Figure S3 supplementary data: (a) Picture of shrimp *Rimicaris exoculata* and (b) drawing of the shrimp with the location of the dissected organs: branchiostegite, scaphognathite and exopodite forming the cephalothoracic cavity and foregut and midgut forming the digestive tract. Full length of the shrimp is about 5 cm.**

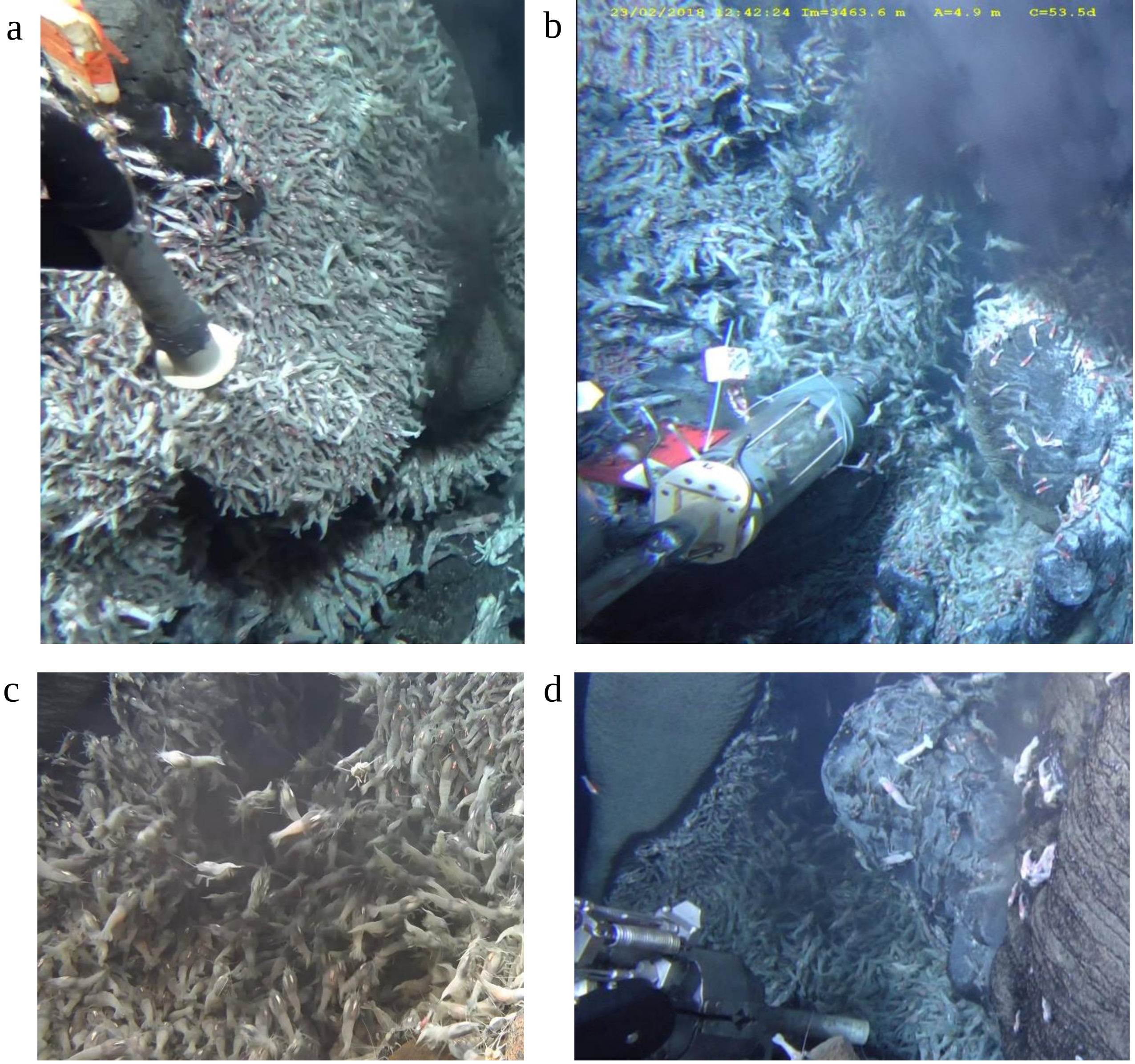


**Figure S4 Supplementary Data: sampled shrimp aggregates (a) FISH1 on Snake Pit – "The Beehive", (b) PERISCOP on Snake Pit – "The Beehive", (c) FISH2 on Snake Pit – "The Moose", (d) suction sampler on Snake Pit – "The Moose"**
