## Supplemental Tables for "FISH, a new tool for in situ preservation of RNA in tissues of deep-sea mobile fauna"

**Table supplementary data S1:** concentrations and RNA Integrity Number (RIN) of extracted RNA[a].

| sample | sampling tool | site | [ ] ng/µL | [ ] average ng/µL | [ ] standard error ng/µL | RNA Integrity Number (RIN) | RIN average | RIN standard error | RIN>8 | 7<RIN<8 | RIN<7 |
|---|---|---|---|---|---|---|---|---|---|---|---|
| ***C257_FISH*** | ***FISH*** | ***The Beehive*** | ***380,00*** | 204,63 | 60,47 | ***8,1*** | 6,117 | 0,7306 | 50% | | 50% |
| ***C262_FISH*** | ***FISH*** | ***The Beehive*** | ***200,00*** | | | ***9,2*** | | | | | |
| C267_FISH | FISH | The Beehive | 10,05 | | | NA | | | | | |
| ***C252_FISH*** | ***FISH*** | ***The Beehive*** | ***372,00*** | | | ***8,3*** | | | | | |
| C253_FISH | FISH | The Beehive | 95,70 | | | 6,4 | | | | | |
| C254_FISH | FISH | The Beehive | 170,00 | | | 4,7 | | | | | |
| ***C347_PERIS-1*** | ***PERISCOP*** | ***The Beehive*** | ***372,00*** | 259,12 | 97,76 | ***9,5*** | 6,983 | 1,1259 | 50% | 17% | 33% |
| C347_PERIS-2 | PERISCOP | The Beehive | 98,90 | | | 8,1 | | | | | |
| C352_PERIS | PERISCOP | The Beehive | 39,80 | | | 2,5 | | | | | |
| ***C357_PERIS*** | ***PERISCOP*** | ***The Beehive*** | ***690,00*** | | | ***7,8*** | | | | | |
| C348_PERIS | PERISCOP | The Beehive | 154,00 | | | 4,8 | | | | | |
| ***C349_PERIS*** | ***PERISCOP*** | ***The Beehive*** | ***200,00*** | | | ***9,2*** | | | | | |
| C480_FISH | FISH | The Moose | 5,99 | 313,39 | 110,17 | NA | 6,111 | 0,7974 | 33% | 22% | 44% |
| C485_FISH | FISH | The Moose | 10,50 | | | 2,5 | | | | | |
| ***C490_FISH*** | ***FISH*** | ***The Moose*** | ***198,00*** | | | ***9,6*** | | | | | |
| ***C480_FISH*** | ***FISH*** | ***The Moose*** | ***365,00*** | | | ***8,4*** | | | | | |
| C481_FISH | FISH | The Moose | 297,00 | | | 7,6 | | | | | |
| C482_FISH | FISH | The Moose | 221,00 | | | 5,4 | | | | | |
| C483_FISH | FISH | The Moose | 224,00 | | | 7,80 | | | | | |
| C486_FISH | FISH | The Moose | 379,00 | | | 4,9 | | | | | |
| ***C487_FISH*** | ***FISH*** | ***The Moose*** | ***1120,00*** | | | ***8,8*** | | | | | |
| C435_ASPI | suction sampler | The Moose | 406,00 | 476,75 | 57,32 | NA | 3,967 | 0,7411 | 8% | | 92% |
| C440_ASPI | suction sampler | The Moose | 316,00 | | | NA | | | | | |
| ***C445_ASPI*** | ***suction sampler*** | ***The Moose*** | ***588,00*** | | | ***6,6*** | | | | | |
| C435_ASPI | suction sampler | The Moose | 371,00 | | | NA | | | | | |
| C436_ASPI | suction sampler | The Moose | 233,00 | | | 3,1 | | | | | |
| C437_ASPI | suction sampler | The Moose | 180,00 | | | 5,90 | | | | | |
| C438_ASPI | suction sampler | The Moose | 870,00 | | | 5,50 | | | | | |
| C439_ASPI | suction sampler | The Moose | 640,00 | | | 6,10 | | | | | |
| C440_ASPI | suction sampler | The Moose | 461,00 | | | 5,20 | | | | | |
| C441_ASPI | suction sampler | The Moose | 600,00 | | | 0,00 | | | | | |
| ***C442_ASPI*** | ***suction sampler*** | ***The Moose*** | ***406,00*** | | | ***9,4*** | | | | | |
| ***C443_ASPI*** | ***suction sampler*** | ***The Moose*** | ***650,00*** | | | ***5,8*** | | | | | |

[a] All extracted RNA are shown in the table with the sample identification and associated metadata such as the sampling tool and the origin edifice, the RNA concentration, the average and standard deviation of RNA concentration grouped by sampling tool and origin, the RNA integrity number (RIN) per sample. The table also shows the percentage of samples classified according to different levels of RNA integrity, grouped by collection tool: good quality (RIN>8), average quality (7<RIN<8) and poor quality with degraded RNA (RIN<7). Selected samples for sequencing are marked in bold and italic.

**Table supplementary data S2:** Statistical results from Wilcoxon-Mann-Whitney test for RNA extracts as a function of concentration and RIN variables

| variable | group1 | group2 | n1 | n2 | statistic | p | p.signif |
|---|---|---|---|---|---|---|---|
| concentration | FISH_The Beehive | FISH_The Moose | 6 | 9 | 23.0 | 0.6889 | ns |
| concentration | FISH_The Beehive | PERISCOP_The Beehive | 6 | 6 | 17.0 | 0.9360 | ns |
| concentration | FISH_The Beehive | suction sampler_The Moose | 6 | 12 | 9.0 | 0.0130 | * |
| concentration | FISH_The Moose | suction sampler_The Moose | 9 | 12 | 24.0 | 0.0360 | * |
| concentration | PERISCOP_The Beehive | suction sampler_The Moose | 6 | 12 | 16.0 | 0.068 | ns |
| RIN | FISH_The Beehive | FISH_The Moose | 6 | 9 | 26.5 | 0.5469 | ns |
| RIN | FISH_The Beehive | PERISCOP_The Beehive | 6 | 6 | 15.0 | 0.7131 | ns |
| RIN | FISH_The Beehive | suction sampler_The Moose | 6 | 12 | 50.0 | 0.1007 | ns |
| RIN | FISH_The Moose | suction sampler_The Moose | 9 | 12 | 73.0 | 0.0929 | ns |
| RIN | PERISCOP_The Beehive | suction sampler_The Moose | 6 | 12 | 54.0 | 0.0497 | * |

**Table supplementary data S3:** Quality trimming and filtering statistics for each metatranscriptome and percent of reads mapped to co-assembly.

| **Sample** | **Edifice** | **Sampling tool** | **RNA [ ] ng/µL** | **Total pairs raw-reads** | **Average of total pair raw-reads** | **Total pairs reads after bowtie2 (rRNA depletion)** | **Average of total pairs reads after bowtie2 (rRNA depletion)** | **Total filtered reads (iu_minoche)** | **Average of total filtered pairs reads (iu_minoche)** | **% of passed pairs (iu_minoche)** | **% of preserved sequences** | **Percent reads mapped assembly** |
|---|---|---|---|---|---|---|---|---|---|---|---|---|
| C252_FISH | The Beehive | FISH | 372 | 36 310 791 | 29 321 030 ± 6 379 392 | 34 110 283 | 27 867 120 ± 5 986 678 | 31 673 007 | 26 091 447 ± 5 496 952 | 92,85% | 87,23% | 95,00% |
| C257_FISH | | | 380 | 16 582 375 | | 15 897 480 | | 15 097 979 | | 94,97% | 91,05% | 93,31% |
| C262_FISH | | | 200 | 35 069 923 | | 33 593 598 | | 31 503 355 | | 93,78% | 89,83% | 93,14% |
| C347_PERISC | | PERISCOP | 372 | 26 273 278 | 31 319 282 ± 4 192 690 | 25 024 095 | 29 542 432 ± 3 886 021 | 23 594 930 | 27 670 372 ± 3 481 603 | 94,29% | 89,81% | 94,33% |
| C349_PERISC | | | 200 | 28 042 341 | | 26 325 110 | | 24 818 538 | | 94,28% | 88,50% | 94,63% |
| C357_PERISC | | | 690 | 39 642 226 | | 37 278 091 | | 34 597 648 | | 92,81% | 87,27% | 95,01% |
| C480_FISH | The Moose | FISH | 365 | 30 755 798 | 31 319 282 ± 4 192 690 | 28 404 174 | 30 999 378 ± 2 007 774 | 26 689 161 | 29 248 834 ± 1 876 054 | 93,96% | 86,78% | 95,15% |
| C487_FISH | | | 1120 | 31 007 559 | | 29 643 272 | | 28 152 792 | | 94,97% | 90,79% | 94,87% |
| C490_FISH | | | 198 | 37 442 430 | | 34 950 687 | | 32 904 548 | | 94,15% | 87,88% | 95,37% |
| C442_ASPI | | suction sampler | 406 | 33 058 864 | 34 361 652 ± 3 480 554 | 31 393 634 | 32 440 492 ± 3 207 964 | 29 496 983 | 30 543 457 ± 3 027 734 | 93,96% | 89,23% | 92,96% |
| C443_ASPI | | | 650 | 29 091 067 | | 27 482 026 | | 25 901 407 | | 94,25% | 89,04% | 95,61% |
| C445_ASPI | | | 588 | 40 935 024 | | 38 445 815 | | 36 231 982 | | 94,24% | 88,51% | 94,05% |
| | | **TOTAL** | | **384 211 676** | **32 017 640 ± 1 934 808** | **362 548 265** | **30 212 355 ± 1 796 842** | **340 662 330** | **28 388 528 ± 1 659 859** | | | |

**Table supplementary data S4:** Statistical results from Wilcoxon-Mann-Whitney test on metatranscriptomes reads according to different sequence cleaning steps.

| variable | group1 | group2 | n1 | n2 | statistic | p | p.adj | p.adj.signif |
|---|---|---|---|---|---|---|---|---|
| Initial total pairs raw reads | FISH_The Beehive | FISH_The Moose | 3 | 3 | 4 | 1 | 1 | ns |
| Initial total pairs raw reads | FISH_The Beehive | PERISCOP_The Beehive | 3 | 3 | 4 | 1 | 1 | ns |
| Initial total pairs raw reads | FISH_The Moose | suction sampler_The Moose | 3 | 3 | 4 | 1 | 1 | ns |
| Total pairs reads after bowtie2 (rRNA depletion) | FISH_The Beehive | FISH_The Moose | 3 | 3 | 4 | 1 | 1 | ns |
| Total pairs reads after bowtie2 (rRNA depletion) | FISH_The Beehive | PERISCOP_The Beehive | 3 | 3 | 4 | 1 | 1 | ns |
| Total pairs reads after bowtie2 (rRNA depletion) | FISH_The Moose | suction sampler_The Moose | 3 | 3 | 4 | 1 | 1 | ns |
| Total pairs reads after quality minoche filtration | FISH_The Beehive | FISH_The Moose | 3 | 3 | 4 | 1 | 1 | ns |
| Total pairs reads after quality minoche filtration | FISH_The Beehive | PERISCOP_The Beehive | 3 | 3 | 4 | 1 | 1 | ns |
| Total pairs reads after quality minoche filtration | FISH_The Moose | suction sampler_The Moose | 3 | 3 | 4 | 1 | 1 | ns |

**Table supplementary data S5**: Quality trimming and filtering statistics for each metagenome and percent of reads mapped to co-assembly

| Sample | line | read | Num Raw reads / line | Total pairs raw-reads / line | Total raw reads / sample | Total pairs raw-reads / sample | Total filtered reads (iu_minoche) | Percent of preserved reads | Num reads mapped assembly | Percent reads mapped assembly | Sum of total detected genes |
|---|---|---|---|---|---|---|---|---|---|---|---|
| Rimi316-Fem | L1 | R1 | 69 043 798 | 69 043 798 | 274 717 788 | 137 358 894 | 250 839 538 | 91,31% | 226 494 675 | 90,29% | 223 539 897 |
| | | R2 | 69 043 798 | | | | | | | | |
| | L2 | R1 | 68 315 096 | 68 315 096 | | | | | | | |
| | | R2 | 68 315 096 | | | | | | | | |
| Rimi317-Fem | L1 | R1 | 82 467 078 | 82 467 078 | 326 638 990 | 163 319 495 | 291 139 334 | 89,13% | 241 264 038 | 82,87% | 184 946 472 |
| | | R2 | 82 467 078 | | | | | | | | |
| | L2 | R1 | 80 852 417 | 80 852 417 | | | | | | | |
| | | R2 | 80 852 417 | | | | | | | | |
| Rimi325-Mal | L1 | R1 | 84 290 796 | 84 290 796 | 333 466 326 | 166 733 163 | 298 749 528 | 89,59% | 261 535 229 | 87,54% | 237 934 362 |
| | | R2 | 84 290 796 | | | | | | | | |
| | L2 | R1 | 82 442 367 | 82 442 367 | | | | | | | |
| | | R2 | 82 442 367 | | | | | | | | |
| Rimi326-Mal | L1 | R1 | 76 818 227 | 76 818 227 | 302 773 514 | 151 386 757 | 269 692 912 | 89,07% | 228 891 605 | 84,87% | 207 291 708 |
| | | R2 | 76 818 227 | | | | | | | | |
| | L2 | R1 | 74 568 530 | 74 568 530 | | | | | | | |
| | | R2 | 74 568 530 | | | | | | | | |
| **TOTAL** | | | **1 237 596 618** | **618 798 309** | **1 237 596 618** | **618 798 309** | **1 110 421 312** | | **958 185 547** | | |

**Table supplementary data S6:** total abundance of recruited transcripts and diversity of expressed genes detected per sample associated with mean ± standard deviation per tool/edifice group

| Sample | Edifice | Sampling tool | Sum of total recruited transcripts | Mean of total recruited transcripts abundance ± SD | Sum of detected genes | Mean of detected genes ± SD |
|---|---|---|---|---|---|---|
| C252_FISH | The Beehive | FISH | 24 918 445 | 21 180 038 ± 8 943 098 | 147 497 | 182 257 ± 32 542 |
| C257_FISH | | | 10 974 334 | | 187 275 | |
| C262_FISH | | | 27 647 335 | | 211 999 | |
| C347_PERISC | | PERISCOP | 21 312 655 | 18 956 230 ± 3 831 351 | 115 207 | 113 901 ± 1 597 |
| C349_PERISC | | | 14 535 380 | | 112 120 | |
| C357_PERISC | | | 21 020 656 | | 114 375 | |
| C480_FISH | The Moose | FISH | 21 249 426 | 33 216 129 ± 10 979 054 | 117 699 | 135 295 ± 15 239 |
| C487_FISH | | | 35 574 823 | | 144 162 | |
| C490_FISH | | | 42 824 137 | | 144 024 | |
| C442_ASPI | | suction sampler | 27 836 047 | 19 097 325 ± 7 569 075 | 119 394 | 123 156 ± 35 348 |
| C443_ASPI | | | 14 858 162 | | 160 235 | |
| C445_ASPI | | | 14 597 767 | | 89 840 | |

**Table supplementary data S7:** Statistical results from Wilcoxon-Mann-Whitney test on reads mapped to genes

| variable | group1 | group2 | n1 | n2 | statistic | p | p. signif |
|---|---|---|---|---|---|---|---|
| total of recruited transcripts mapped to genes | FISH_The Beehive | FISH_The Moose | 3 | 3 | 2 | 0.90 | ns |
| total of recruited transcripts mapped to genes | FISH_The Beehive | PERISCOP_The Beehive | 3 | 3 | 6 | 0.35 | ns |
| total of recruited transcripts mapped to genes | FISH_The Moose | suction sampler_The Moose | 3 | 3 | 8 | 0.10 | ns |
| detected genes | FISH_The Beehive | FISH_The Moose | 3 | 3 | 9 | 0.05 | * |
| detected genes | FISH_The Beehive | PERISCOP_The Beehive | 3 | 3 | 9 | 0.05 | * |
| detected genes | FISH_The Moose | suction sampler_The Moose | 3 | 3 | 5 | 0.50 | ns |

**Table supplementary data S8a:** taxonomic identification with MMSeqs2 per sample expressed in read count

| Edifice + tool | The Beehive: FISH | | | The Beehive: PERISCOP | | | The Moose: FISH | | | The Moose: suction sampler | | |
|---|---|---|---|---|---|---|---|---|---|---|---|---|
| **phylum** | **C252** | **C257** | **C262** | **C347** | **C349** | **C357** | **C480** | **C487** | **C490** | **C442** | **C443** | **C445** |
| *Campylobacterota* | 6 390 690 | 2 302 557 | 7 089 731 | 5 761 882 | 2 954 162 | 3 989 297 | 5 693 229 | 11 596 687 | 13 891 481 | 7 505 190 | 2 111 346 | 604 762 |
| *Desulfobacterota* | 669 797 | 119 695 | 55 374 | 36 177 | 255 264 | 66 097 | 65 697 | 214 957 | 214 796 | 39 998 | 170 109 | 60 276 |
| *Pseudomonadota* | 302 812 | 128 000 | 224 956 | 167 922 | 125 078 | 175 853 | 319 026 | 627 903 | 865 358 | 364 734 | 572 281 | 199 388 |
| *Bacteroidota* | 171 968 | 91 262 | 143 729 | 200 404 | 84 309 | 192 275 | 147 026 | 544 551 | 950 929 | 208 949 | 288 417 | 104 356 |
| *Deferribacteres* | 94 854 | 11 815 | 57 032 | 59 691 | 67 171 | 96 744 | 116 950 | 77 684 | 169 916 | 75 557 | 28 014 | 12 905 |
| *Patescibacteria* | 34 344 | 12 404 | 28 218 | 21 818 | 13 107 | 17 111 | 63 170 | 148 284 | 167 441 | 102 445 | 34 925 | 12 958 |
| *Bacillota* | 16 301 | 4 318 | 8 208 | 7 985 | 7 519 | 7 651 | 26 515 | 37 791 | 54 216 | 16 790 | 8 795 | 2 635 |
| *Aquificota* | 4 766 | 3 430 | 4 590 | 3 594 | 3 100 | 4 464 | 5 693 | 13 026 | 16 664 | 8 928 | 16 006 | 5 858 |
| *Verrucomicrobiota* | 3 725 | 362 | 734 | 793 | 1 924 | 878 | 4 072 | 1 684 | 2 964 | 924 | 3 158 | 1 599 |
| *Cyanobacteriota* | 2 581 | 743 | 1 809 | 918 | 1 310 | 822 | 2 020 | 3 778 | 5 342 | 1 619 | 1 561 | 690 |
| *Planctomycetota* | 2 040 | 514 | 610 | 445 | 877 | 507 | 745 | 1 514 | 2 047 | 613 | 1 588 | 675 |
| *Calditrichota* | 1 346 | 554 | 1 622 | 1 773 | 779 | 1 220 | 1 297 | 3 934 | 3 434 | 2 241 | 760 | 237 |
| *Chloroflexota* | 757 | 282 | 377 | 241 | 335 | 262 | 413 | 683 | 1 191 | 397 | 712 | 254 |
| *Actinomycetota* | 752 | 407 | 891 | 545 | 353 | 344 | 373 | 697 | 981 | 663 | 472 | 186 |
| *Spirochaetota* | 720 | 228 | 610 | 457 | 414 | 385 | 582 | 1 298 | 1 624 | 1 058 | 387 | 133 |
| Other Bacteria | 4 585 | 1 537 | 3 219 | 3 155 | 2 599 | 2 327 | 3 526 | 7 707 | 10 142 | 4 506 | 4 594 | 1 226 |
| Unidentified Bacteria | 990 071 | 151 163 | 471 198 | 541 744 | 554 183 | 788 593 | 1 128 231 | 1 034 270 | 1 817 953 | 764 618 | 410 822 | 135 613 |
| Archaea | 7 122 | 7 280 | 9 691 | 4 795 | 4 358 | 3 922 | 6 340 | 11 834 | 13 693 | 8 895 | 6 375 | 2 028 |
| Viruses | 773 | 349 | 491 | 172 | 290 | 174 | 664 | 1 375 | 2 034 | 1 986 | 505 | 119 |
| Arthropoda_Decapoda | 415 255 | 350 664 | 849 590 | 375 794 | 201 113 | 484 691 | 181 242 | 254 258 | 80 838 | 503 869 | 394 364 | 181 108 |
| Other Arthropoda | 133 868 | 57 094 | 95 548 | 43 391 | 27 580 | 158 177 | 18 305 | 36 610 | 19 988 | 41 785 | 78 031 | 30 859 |
| Other Eukaryota | 416 978 | 274 878 | 562 797 | 291 869 | 208 775 | 387 804 | 142 339 | 331 209 | 88 890 | 250 944 | 360 410 | 303 051 |
| Non identified | 15 252 340 | 7 454 798 | 18 036 310 | 13 787 090 | 10 020 780 | 14 641 058 | 13 321 971 | 20 623 089 | 24 442 215 | 17 929 338 | 10 364 530 | 12 936 851 |
| **TOTAL** | **24 918 445** | **10 974 334** | **27 647 335** | **21 312 655** | **14 535 380** | **21 020 656** | **21 249 426** | **35 574 823** | **42 824 137** | **27 836 047** | **14 858 162** | **14 597 767** |

**Table supplementary data S8b:** taxonomic identification with MMSeqs2 per sample expressed in percentage of the total genes with associated average and standard deviation

| Edifice + tool | The Beehive: FISH | | | | The Beehive: PERISCOP | | | | The Moose: FISH | | | | The Moose: suction sampler | | | | average ± SD |
|---|---|---|---|---|---|---|---|---|---|---|---|---|---|---|---|---|---|
| **phylum** | **C252** | **C257** | **C262** | **average ± SD** | **C347** | **C349** | **C357** | **average ± SD** | **C480** | **C487** | **C490** | **average ± SD** | **C442** | **C443** | **C445** | **average ± SD** | |
| *Campylobacterota* | 25,646% | 20,981% | 25,643% | 24,09% ± 2,693% | 27,035% | 20,324% | 18,978% | 22,112% ± 4,316% | 26,792% | 32,598% | 32,438% | 30,61% ± 3,307% | 26,962% | 14,210% | 4,143% | 15,105% ± 11,436% | 22,979% ± 7,988% |
| *Desulfobacterota* | 2,688% | 1,091% | 0,200% | 1,326% ± 1,26% | 0,170% | 1,756% | 0,314% | 0,747% ± 0,877% | 0,309% | 0,604% | 0,502% | 0,472% ± 0,15% | 0,144% | 1,145% | 0,413% | 0,567% ± 0,518% | 0,778% ± 0,776% |
| *Pseudomonadota* | 1,215% | 1,166% | 0,814% | 1,065% ± 0,219% | 0,788% | 0,861% | 0,837% | 0,828% ± 0,037% | 1,501% | 1,765% | 2,021% | 1,762% ± 0,26% | 1,310% | 3,852% | 1,366% | 2,176% ± 1,451% | 1,458% ± 0,849% |
| *Bacteroidota* | 0,690% | 0,832% | 0,520% | 0,681% ± 0,156% | 0,940% | 0,580% | 0,915% | 0,812% ± 0,201% | 0,692% | 1,531% | 2,221% | 1,481% ± 0,766% | 0,751% | 1,941% | 0,715% | 1,136% ± 0,698% | 1,027% ± 0,558% |
| *Deferribacteres* | 0,381% | 0,108% | 0,206% | 0,232% ± 0,138% | 0,280% | 0,462% | 0,460% | 0,401% ± 0,105% | 0,550% | 0,218% | 0,397% | 0,389% ± 0,166% | 0,271% | 0,189% | 0,088% | 0,183% ± 0,092% | 0,301% ± 0,148% |
| *Patescibacteria* | 0,138% | 0,113% | 0,102% | 0,118% ± 0,018% | 0,102% | 0,090% | 0,081% | 0,091% ± 0,011% | 0,297% | 0,417% | 0,391% | 0,368% ± 0,063% | 0,368% | 0,235% | 0,089% | 0,231% ± 0,14% | 0,202% ± 0,132% |
| *Bacillota* | 0,065% | 0,039% | 0,030% | 0,045% ± 0,018% | 0,037% | 0,052% | 0,036% | 0,042% ± 0,009% | 0,125% | 0,106% | 0,127% | 0,119% ± 0,011% | 0,060% | 0,059% | 0,018% | 0,046% ± 0,024% | 0,063% ± 0,037% |
| *Aquificota* | 0,019% | 0,031% | 0,017% | 0,022% ± 0,008% | 0,017% | 0,021% | 0,021% | 0,02% ± 0,003% | 0,027% | 0,037% | 0,039% | 0,034% ± 0,006% | 0,032% | 0,108% | 0,040% | 0,06% ± 0,042% | 0,034% ± 0,025% |
| *Verrucomicrobiota* | 0,015% | 0,003% | 0,003% | 0,007% ± 0,007% | 0,004% | 0,013% | 0,004% | 0,007% ± 0,005% | 0,019% | 0,005% | 0,007% | 0,01% ± 0,008% | 0,003% | 0,021% | 0,011% | 0,012% ± 0,009% | 0,009% ± 0,007% |
| *Cyanobacteriota* | 0,010% | 0,007% | 0,007% | 0,008% ± 0,002% | 0,004% | 0,009% | 0,004% | 0,006% ± 0,003% | 0,010% | 0,011% | 0,012% | 0,011% ± 0,001% | 0,006% | 0,011% | 0,005% | 0,007% ± 0,003% | 0,008% ± 0,003% |
| *Planctomycetota* | 0,008% | 0,005% | 0,002% | 0,005% ± 0,003% | 0,002% | 0,006% | 0,002% | 0,004% ± 0,002% | 0,004% | 0,004% | 0,005% | 0,004% ± 0,001% | 0,002% | 0,011% | 0,005% | 0,006% ± 0,004% | 0,005% ± 0,003% |
| *Calditrichota* | 0,005% | 0,005% | 0,006% | 0,005% ± 0% | 0,008% | 0,005% | 0,006% | 0,006% ± 0,002% | 0,006% | 0,011% | 0,008% | 0,008% ± 0,002% | 0,008% | 0,005% | 0,002% | 0,005% ± 0,003% | 0,006% ± 0,002% |
| *Chloroflexota* | 0,003% | 0,003% | 0,001% | 0,002% ± 0,001% | 0,001% | 0,002% | 0,001% | 0,002% ± 0,001% | 0,002% | 0,002% | 0,003% | 0,002% ± 0% | 0,001% | 0,005% | 0,002% | 0,003% ± 0,002% | 0,002% ± 0,001% |
| *Actinomycetota* | 0,003% | 0,004% | 0,003% | 0,003% ± 0% | 0,003% | 0,002% | 0,002% | 0,002% ± 0% | 0,002% | 0,002% | 0,002% | 0,002% ± 0% | 0,002% | 0,003% | 0,001% | 0,002% ± 0,001% | 0,002% ± 0,001% |
| *Spirochaetota* | 0,003% | 0,002% | 0,002% | 0,002% ± 0% | 0,002% | 0,003% | 0,002% | 0,002% ± 0,001% | 0,003% | 0,004% | 0,004% | 0,003% ± 0,001% | 0,004% | 0,003% | 0,001% | 0,002% ± 0,001% | 0,003% ± 0,001% |
| Other Bacteria | 0,018% | 0,014% | 0,012% | 0,015% ± 0,003% | 0,015% | 0,018% | 0,011% | 0,015% ± 0,003% | 0,017% | 0,022% | 0,024% | 0,021% ± 0,004% | 0,016% | 0,031% | 0,008% | 0,019% ± 0,011% | 0,017% ± 0,006% |
| Unidentified Bacteria | 3,973% | 1,377% | 1,704% | 2,352% ± 1,414% | 2,542% | 3,813% | 3,752% | 3,369% ± 0,717% | 5,309% | 2,907% | 4,245% | 4,154% ± 1,204% | 2,747% | 2,765% | 0,929% | 2,147% ± 1,055% | 3,005% ± 1,279% |
| Archaea | 0,029% | 0,066% | 0,035% | 0,043% ± 0,02% | 0,022% | 0,030% | 0,019% | 0,024% ± 0,006% | 0,030% | 0,033% | 0,032% | 0,032% ± 0,002% | 0,032% | 0,043% | 0,014% | 0,03% ± 0,015% | 0,032% ± 0,013% |
| Viruses | 0,003% | 0,003% | 0,002% | 0,003% ± 0,001% | 0,001% | 0,002% | 0,001% | 0,001% ± 0,001% | 0,003% | 0,004% | 0,005% | 0,004% ± 0,001% | 0,007% | 0,003% | 0,001% | 0,004% ± 0,003% | 0,003% ± 0,002% |
| Arthropoda_Decapoda | 1,666% | 3,195% | 3,073% | 2,645% ± 0,85% | 1,763% | 1,384% | 2,306% | 1,818% ± 0,463% | 0,853% | 0,715% | 0,189% | 0,585% ± 0,35% | 1,810% | 2,654% | 1,241% | 1,902% ± 0,711% | 1,737% ± 0,938% |
| Other Arthropoda | 0,537% | 0,520% | 0,346% | 0,468% ± 0,106% | 0,204% | 0,190% | 0,752% | 0,382% ± 0,321% | 0,086% | 0,103% | 0,047% | 0,079% ± 0,029% | 0,150% | 0,525% | 0,211% | 0,296% ± 0,201% | 0,306% ± 0,226% |
| Other Eukaryota | 1,673% | 2,505% | 2,036% | 2,071% ± 0,417% | 1,369% | 1,436% | 1,845% | 1,55% ± 0,257% | 0,670% | 0,931% | 0,208% | 0,603% ± 0,366% | 0,902% | 2,426% | 2,076% | 1,801% ± 0,798% | 1,506% ± 0,72% |
| Non identified | 61,209% | 67,929% | 65,237% | 64,792% ± 3,382% | 64,690% | 68,941% | 69,651% | 67,76% ± 2,683% | 62,693% | 57,971% | 57,076% | 59,247% ± 3,018% | 64,411% | 69,756% | 88,622% | 74,263% ± 12,719% | 66,515% ± 8,153% |
| Identified | 38,791% | 32,071% | 34,763% | 35,208% ± 3,382% | 35,310% | 31,059% | 30,349% | 32,24% ± 2,683% | 37,307% | 42,029% | 42,924% | 40,753% ± 3,018% | 35,589% | 30,244% | 11,378% | 25,737% ± 12,719% | 33,485% ± 8,153% |

**Table supplementary data S9:** list of differential exprimed genes with base Mean value, log2Fold Change value and COG20 category

| gene_callers_id | baseMean | log2FoldChange | lfcSE | stat | pvalue | padj | comparative_samples | COG20_CATEGORY |
|---|---|---|---|---|---|---|---|---|
| gene_330014 | 12,82639698 | -6,702649136 | 1,548313972 | -4,328998678 | 1,50E-05 | 0,01126733 | The Beehive: PERISCOP vs FISH | Amino acid transport and metabolism |
| gene_43099 | 11,80369951 | -5,731140219 | 1,514174837 | -3,784992379 | 0,000153713 | 0,041873641 | The Beehive: PERISCOP vs FISH | Amino acid transport and metabolism |
| gene_1875242 | 25,88776078 | -3,18499761 | 0,848131062 | -3,755313007 | 0,000173125 | 0,043237512 | The Beehive: PERISCOP vs FISH | Amino acid transport and metabolism |
| gene_1120289 | 66,14517247 | -2,615110737 | 0,690067065 | -3,789647224 | 0,000150861 | 0,041873641 | The Beehive: PERISCOP vs FISH | Amino acid transport and metabolism |
| gene_1944979 | 69,64852261 | 2,803537559 | 0,742809107 | 3,774236925 | 0,000160498 | 0,042255157 | The Beehive: PERISCOP vs FISH | Amino acid transport and metabolism |
| gene_641988 | 74,98146844 | 3,300135669 | 0,728882036 | 4,527667722 | 5,96E-06 | 0,00765937 | The Beehive: PERISCOP vs FISH | Amino acid transport and metabolism |
| gene_804225 | 16,05853464 | 5,283161095 | 1,335215173 | 3,956786292 | 7,60E-05 | 0,032431851 | The Beehive: PERISCOP vs FISH | Amino acid transport and metabolism |
| gene_242608 | 15,27522735 | 5,862557798 | 1,204006634 | 4,869207223 | 1,12E-06 | 0,002082292 | The Beehive: PERISCOP vs FISH | Amino acid transport and metabolism |
| gene_1363971 | 44,17550053 | -3,276746143 | 0,753580618 | -4,348235696 | 1,37E-05 | 0,010839327 | The Beehive: PERISCOP vs FISH | Carbohydrate transport and metabolism |
| gene_1115130 | 14,97149071 | 4,709427428 | 1,245647918 | 3,780705094 | 0,000156385 | 0,041873641 | The Beehive: PERISCOP vs FISH | Carbohydrate transport and metabolism |
| gene_907749 | 38,10514111 | -3,371201511 | 0,769135753 | -4,383103368 | 1,17E-05 | 0,010838058 | The Beehive: PERISCOP vs FISH | Cell cycle control, cell division, chromosome partitioning |
| gene_1313271 | 177,9735987 | 2,858437712 | 0,706927578 | 4,043466122 | 5,27E-05 | 0,025210625 | The Beehive: PERISCOP vs FISH | Cell cycle control, cell division, chromosome partitioning |
| gene_331339 | 43,31615295 | 6,149585796 | 1,430117576 | 4,300056094 | 1,71E-05 | 0,012260589 | The Beehive: PERISCOP vs FISH | Cell cycle control, cell division, chromosome partitioning |
| gene_328426 | 12,35128142 | -6,910330276 | 1,633355782 | -4,230756307 | 2,33E-05 | 0,013745153 | The Beehive: PERISCOP vs FISH | Cell wall/membrane/envelope biogenesis |
| gene_328421 | 22,33775979 | -5,186233588 | 1,093849005 | -4,741270105 | 2,12E-06 | 0,003195144 | The Beehive: PERISCOP vs FISH | Cell wall/membrane/envelope biogenesis |
| gene_635649 | 66,25656412 | 2,536339255 | 0,675869602 | 3,752705032 | 0,000174937 | 0,043237512 | The Beehive: PERISCOP vs FISH | Cell wall/membrane/envelope biogenesis |
| gene_90814 | 131,6421273 | 2,610339247 | 0,669922584 | 3,896478953 | 9,76E-05 | 0,036276675 | The Beehive: PERISCOP vs FISH | Cell wall/membrane/envelope biogenesis |
| gene_240346 | 3805,331305 | 2,754244806 | 0,62722219 | 4,391178839 | 1,13E-05 | 0,010793104 | The Beehive: PERISCOP vs FISH | Cell wall/membrane/envelope biogenesis |
| gene_1175427 | 159,7994155 | 3,043449194 | 0,788886836 | 3,857903385 | 0,000114364 | 0,038850501 | The Beehive: PERISCOP vs FISH | Cell wall/membrane/envelope biogenesis |
| gene_397613 | 53,75905901 | 5,167460589 | 1,294417235 | 3,992113554 | 6,55E-05 | 0,028983093 | The Beehive: PERISCOP vs FISH | Cell wall/membrane/envelope biogenesis |
| gene_982011 | 31,82609738 | 6,237326777 | 1,533636548 | 4,067017563 | 4,76E-05 | 0,02440368 | The Beehive: PERISCOP vs FISH | Cell wall/membrane/envelope biogenesis |
| gene_283998 | 1424,267723 | -2,508002464 | 0,555483588 | -4,514989315 | 6,33E-06 | 0,007680627 | The Beehive: PERISCOP vs FISH | Coenzyme transport and metabolism |
| gene_240343 | 7367,858598 | 2,234116635 | 0,581804143 | 3,839980624 | 0,000123044 | 0,040270798 | The Beehive: PERISCOP vs FISH | Defense mechanisms |
| gene_655984 | 62,36614606 | 3,162431346 | 0,834667279 | 3,788852666 | 0,000151345 | 0,041873641 | The Beehive: PERISCOP vs FISH | Defense mechanisms |
| gene_904341 | 217,4517414 | -2,707533306 | 0,55972329 | -4,837271125 | 1,32E-06 | 0,002310393 | The Beehive: PERISCOP vs FISH | Energy production and conversion |
| gene_907748 | 260,2339004 | -2,440162779 | 0,639557936 | -3,815389728 | 0,000135968 | 0,041873641 | The Beehive: PERISCOP vs FISH | Energy production and conversion |
| gene_907746 | 307,8233655 | -2,394513621 | 0,623506035 | -3,840401677 | 0,000122833 | 0,040270798 | The Beehive: PERISCOP vs FISH | Energy production and conversion |
| gene_664933 | 485,5673443 | -2,340431894 | 0,606209095 | -3,860766712 | 0,000113032 | 0,038850501 | The Beehive: PERISCOP vs FISH | Energy production and conversion |
| gene_404679 | 1663,797977 | -2,187868424 | 0,578404167 | -3,782594509 | 0,000155202 | 0,041873641 | The Beehive: PERISCOP vs FISH | Energy production and conversion |
| gene_1049435 | 216,8867051 | 2,252369771 | 0,570701616 | 3,946667937 | 7,92E-05 | 0,032514672 | The Beehive: PERISCOP vs FISH | Energy production and conversion |
| gene_677560 | 11982,07974 | 2,790378685 | 0,685435619 | 4,070956644 | 4,68E-05 | 0,02440368 | The Beehive: PERISCOP vs FISH | Energy production and conversion |
| gene_677561 | 12141,09429 | 3,243860503 | 0,678336191 | 4,782083793 | 1,73E-06 | 0,002884729 | The Beehive: PERISCOP vs FISH | Energy production and conversion |
| gene_381046 | 100,6046623 | 3,420142563 | 0,799166232 | 4,279638487 | 1,87E-05 | 0,012527261 | The Beehive: PERISCOP vs FISH | Energy production and conversion |
| gene_381048 | 92,91234109 | 3,464652936 | 0,771508108 | 4,490753757 | 7,10E-06 | 0,007780965 | The Beehive: PERISCOP vs FISH | Energy production and conversion |
| gene_1856084 | 92,76803164 | 3,88917254 | 0,866300806 | 4,489401963 | 7,14E-06 | 0,007780965 | The Beehive: PERISCOP vs FISH | Energy production and conversion |
| gene_381045 | 515,6433177 | 3,99667571 | 0,778281009 | 5,135260482 | 2,82E-07 | 0,000890144 | The Beehive: PERISCOP vs FISH | Energy production and conversion |
| gene_170279 | 47,64059301 | 4,231442552 | 1,077958197 | 3,925423604 | 8,66E-05 | 0,03380357 | The Beehive: PERISCOP vs FISH | Energy production and conversion |
| gene_228362 | 28,82768744 | 4,570828355 | 1,142656509 | 4,000177061 | 6,33E-05 | 0,028566891 | The Beehive: PERISCOP vs FISH | Energy production and conversion |
| gene_579523 | 36,080737 | 4,719841521 | 0,845176929 | 5,584441979 | 2,34E-08 | 0,000246901 | The Beehive: PERISCOP vs FISH | Energy production and conversion |
| gene_1451498 | 22,15121932 | 5,348396031 | 1,205513359 | 4,436612825 | 9,14E-06 | 0,009623801 | The Beehive: PERISCOP vs FISH | Energy production and conversion |
| gene_579522 | 21,45902609 | 5,823222813 | 1,113817241 | 5,22816724 | 1,71E-07 | 0,000672421 | The Beehive: PERISCOP vs FISH | Energy production and conversion |
| gene_1786364 | 26,18676062 | 6,743404935 | 1,163102976 | 5,797771199 | 6,72E-09 | 0,000106156 | The Beehive: PERISCOP vs FISH | Energy production and conversion |
| gene_1156488 | 52,57267808 | -3,548790456 | 0,869900855 | -4,07953439 | 4,51E-05 | 0,024163822 | The Beehive: PERISCOP vs FISH | Function unknown |
| gene_980493 | 114,1237671 | -2,644595498 | 0,70514864 | -3,750408565 | 0,000176547 | 0,043237512 | The Beehive: PERISCOP vs FISH | Function unknown |
| gene_1074196 | 823,9944644 | -2,44156341 | 0,627757644 | -3,889340786 | 0,000100517 | 0,03692593 | The Beehive: PERISCOP vs FISH | Function unknown |
| gene_280111 | 216,2243246 | -2,274829497 | 0,601595276 | -3,781328726 | 0,000155994 | 0,041873641 | The Beehive: PERISCOP vs FISH | Function unknown |
| gene_1161021 | 1775,439274 | 2,444303884 | 0,609533378 | 4,010123108 | 6,07E-05 | 0,027786775 | The Beehive: PERISCOP vs FISH | Function unknown |
| gene_494739 | 189,1495153 | 2,491449596 | 0,631088866 | 3,947858582 | 7,89E-05 | 0,032514672 | The Beehive: PERISCOP vs FISH | Function unknown |
| gene_1857519 | 1179,388965 | 2,707767802 | 0,700939523 | 3,863054821 | 0,000111978 | 0,038850501 | The Beehive: PERISCOP vs FISH | Function unknown |
| gene_155647 | 327,0706708 | 2,746283499 | 0,553960126 | 4,957547246 | 7,14E-07 | 0,001409614 | The Beehive: PERISCOP vs FISH | Function unknown |
| gene_1206116 | 164,909513 | 2,753145787 | 0,649038861 | 4,241881271 | 2,22E-05 | 0,013466756 | The Beehive: PERISCOP vs FISH | Function unknown |
| gene_342830 | 55,78436756 | 3,101535272 | 0,713282638 | 4,348255668 | 1,37E-05 | 0,010839327 | The Beehive: PERISCOP vs FISH | Function unknown |
| gene_1175426 | 121,9259385 | 3,497958833 | 0,692181927 | 5,05352523 | 4,34E-07 | 0,000978771 | The Beehive: PERISCOP vs FISH | Function unknown |
| gene_1087298 | 593,9678921 | 3,698819061 | 0,852897168 | 4,336770247 | 1,45E-05 | 0,011141666 | The Beehive: PERISCOP vs FISH | Function unknown |
| gene_618098 | 29,80684855 | 3,926412558 | 0,92489486 | 4,245252868 | 2,18E-05 | 0,013466756 | The Beehive: PERISCOP vs FISH | Function unknown |
| gene_332013 | 35,22983387 | 5,083092388 | 1,326032662 | 3,833308586 | 0,000126431 | 0,040306637 | The Beehive: PERISCOP vs FISH | Function unknown |
| gene_192454 | 15,80766089 | 5,431404038 | 1,361228134 | 3,990076242 | 6,61E-05 | 0,028983093 | The Beehive: PERISCOP vs FISH | Function unknown |
| gene_1128777 | 13,30676872 | -7,248997598 | 1,599755472 | -4,531316021 | 5,86E-06 | 0,00765937 | The Beehive: PERISCOP vs FISH | Inorganic ion transport and metabolism |
| gene_501651 | 13,10211231 | -7,129899459 | 1,581835985 | -4,507356974 | 6,56E-06 | 0,007680627 | The Beehive: PERISCOP vs FISH | Inorganic ion transport and metabolism |
| gene_1078741 | 28,7763891 | -3,847534182 | 0,899809449 | -4,275943296 | 1,90E-05 | 0,012527261 | The Beehive: PERISCOP vs FISH | Inorganic ion transport and metabolism |
| gene_149497 | 275,6563355 | 2,835855791 | 0,700017724 | 4,051119987 | 5,10E-05 | 0,024775259 | The Beehive: PERISCOP vs FISH | Inorganic ion transport and metabolism |

| | | | | | | | | |
|---|---|---|---|---|---|---|---|---|
| gene_381047 | 154,9317266 | 3,305122953 | 0,874501738 | 3,779435546 | 0,000157184 | 0,041873641 | The Beehive: PERISCOP vs FISH | Inorganic ion transport and metabolism |
| gene_304426 | 14,15935173 | 4,999630166 | 1,145108946 | 4,366073798 | 1,26E-05 | 0,010839327 | The Beehive: PERISCOP vs FISH | Inorganic ion transport and metabolism |
| gene_180283 | 15,58862074 | -5,588962314 | 1,375929524 | -4,061953913 | 4,87E-05 | 0,02440368 | The Beehive: PERISCOP vs FISH | Intracellular trafficking, secretion, and vesicular transport |
| gene_1491065 | 274,0332214 | 2,798756633 | 0,72782292 | 3,845381283 | 0,000120365 | 0,040270798 | The Beehive: PERISCOP vs FISH | Lipid transport and metabolism |
| gene_255021 | 116,2017597 | 2,841737358 | 0,748449985 | 3,796830001 | 0,000146558 | 0,041873641 | The Beehive: PERISCOP vs FISH | Lipid transport and metabolism |
| gene_1013532 | 13,97953564 | 3,843516323 | 1,022176383 | 3,760130235 | 0,000169825 | 0,043237512 | The Beehive: PERISCOP vs FISH | Lipid transport and metabolism |
| gene_158950 | 18,67537431 | 5,859085501 | 1,555111952 | 3,767629394 | 0,000164805 | 0,042854033 | The Beehive: PERISCOP vs FISH | Lipid transport and metabolism |
| gene_463780 | 464,929082 | 2,269539803 | 0,541470091 | 4,191440749 | 2,77E-05 | 0,015922216 | The Beehive: PERISCOP vs FISH | Mobilome: prophages, transposons |
| gene_463781 | 702,6072166 | 2,770584551 | 0,542572959 | 5,106381558 | 3,28E-07 | 0,000943155 | The Beehive: PERISCOP vs FISH | Mobilome: prophages, transposons |
| gene_1004895 | 182,2999916 | 6,470765337 | 1,486215156 | 4,353855034 | 1,34E-05 | 0,010839327 | The Beehive: PERISCOP vs FISH | NA |
| gene_1005093 | 88,17383114 | -3,803714326 | 0,967061117 | -3,93327191 | 8,38E-05 | 0,03380357 | The Beehive: PERISCOP vs FISH | NA |
| gene_1054377 | 114,010381 | 2,50823187 | 0,642156561 | 3,905950703 | 9,39E-05 | 0,035324735 | The Beehive: PERISCOP vs FISH | NA |
| gene_1088960 | 240,6709556 | 2,288698635 | 0,585977381 | 3,905779829 | 9,39E-05 | 0,035324735 | The Beehive: PERISCOP vs FISH | NA |
| gene_1194279 | 27,30448763 | 6,433000937 | 1,627657328 | 3,952306684 | 7,74E-05 | 0,032514672 | The Beehive: PERISCOP vs FISH | NA |
| gene_1228936 | 166,990548 | -2,922531488 | 0,764971955 | -3,820442654 | 0,000133212 | 0,041669095 | The Beehive: PERISCOP vs FISH | NA |
| gene_1371055 | 197,5864405 | -2,361674856 | 0,582804198 | -4,052261232 | 5,07E-05 | 0,024775259 | The Beehive: PERISCOP vs FISH | NA |
| gene_155648 | 280,6002734 | 3,108777443 | 0,568089863 | 5,472333945 | 4,44E-08 | 0,000310744 | The Beehive: PERISCOP vs FISH | NA |
| gene_1696998 | 26,68865168 | -21,38121721 | 4,249865076 | -5,031034356 | 4,88E-07 | 0,00102749 | The Beehive: PERISCOP vs FISH | NA |
| gene_171204 | 15,72336973 | -4,691474419 | 1,215589459 | -3,859423413 | 0,000113655 | 0,038850501 | The Beehive: PERISCOP vs FISH | NA |
| gene_260640 | 33,31314636 | -3,765710447 | 1,013552071 | -3,715359628 | 0,000202915 | 0,048936566 | The Beehive: PERISCOP vs FISH | NA |
| gene_320247 | 19,06929533 | -5,738119926 | 1,384506757 | -4,144522874 | 3,41E-05 | 0,018873878 | The Beehive: PERISCOP vs FISH | NA |
| gene_331338 | 760,0332587 | 3,567049058 | 0,613473607 | 5,814511033 | 6,08E-09 | 0,000106156 | The Beehive: PERISCOP vs FISH | NA |
| gene_391201 | 35,05422528 | -2,844789306 | 0,755267152 | -3,766600068 | 0,000165486 | 0,042854033 | The Beehive: PERISCOP vs FISH | NA |
| gene_393293 | 130,8885394 | 4,007821677 | 0,734806792 | 5,454252359 | 4,92E-08 | 0,000310744 | The Beehive: PERISCOP vs FISH | NA |
| gene_397813 | 24,807505 | 5,983539147 | 1,485967242 | 4,026696536 | 5,66E-05 | 0,026280712 | The Beehive: PERISCOP vs FISH | NA |
| gene_422786 | 1077,336769 | -2,428678178 | 0,618379837 | -3,927486044 | 8,58E-05 | 0,03380357 | The Beehive: PERISCOP vs FISH | NA |
| gene_618097 | 13,1783547 | 6,503416537 | 1,511553151 | 4,302472946 | 1,69E-05 | 0,012260589 | The Beehive: PERISCOP vs FISH | NA |
| gene_65404 | 162,2082341 | 2,752453923 | 0,643228248 | 4,279124763 | 1,88E-05 | 0,012527261 | The Beehive: PERISCOP vs FISH | NA |
| gene_703712 | 11,08098744 | -6,258596635 | 1,616233505 | -3,872334423 | 0,000107798 | 0,038850501 | The Beehive: PERISCOP vs FISH | NA |
| gene_731613 | 251,4722894 | -3,471673973 | 0,924043541 | -3,757045874 | 0,000171931 | 0,043237512 | The Beehive: PERISCOP vs FISH | NA |
| gene_925347 | 170,8521449 | 3,089269165 | 0,650855032 | 4,746478113 | 2,07E-06 | 0,003195144 | The Beehive: PERISCOP vs FISH | NA |
| gene_931390 | 53,66740246 | 3,747699459 | 0,828357955 | 4,524251187 | 6,06E-06 | 0,00765937 | The Beehive: PERISCOP vs FISH | NA |
| gene_89393 | 28,44728333 | -4,180561092 | 1,101527679 | -3,795239257 | 0,000147501 | 0,041873641 | The Beehive: PERISCOP vs FISH | Nucleotide transport and metabolism |
| gene_791198 | 61,35800484 | -2,862270389 | 0,708865421 | -4,037819175 | 5,40E-05 | 0,025439633 | The Beehive: PERISCOP vs FISH | Nucleotide transport and metabolism |
| gene_988777 | 69,57768231 | -2,724523595 | 0,719847266 | -3,784863433 | 0,000153793 | 0,041873641 | The Beehive: PERISCOP vs FISH | Nucleotide transport and metabolism |
| gene_1663221 | 15,57613486 | -4,953402241 | 1,281516438 | -3,865266254 | 0,000110968 | 0,038850501 | The Beehive: PERISCOP vs FISH | Posttranslational modification, protein turnover, chaperones |
| gene_1023468 | 99,55132797 | -2,730167765 | 0,722537571 | -3,77858242 | 0,000157724 | 0,041873641 | The Beehive: PERISCOP vs FISH | Posttranslational modification, protein turnover, chaperones |
| gene_554756 | 341,0409833 | -2,206378209 | 0,557041619 | -3,960885743 | 7,47E-05 | 0,032316722 | The Beehive: PERISCOP vs FISH | Posttranslational modification, protein turnover, chaperones |
| gene_470653 | 42,02309324 | 3,200133577 | 0,835064407 | 3,832199706 | 0,000127003 | 0,040306637 | The Beehive: PERISCOP vs FISH | Posttranslational modification, protein turnover, chaperones |
| gene_1203978 | 47,53127343 | 5,227842039 | 1,111689682 | 4,70260912 | 2,57E-06 | 0,003688596 | The Beehive: PERISCOP vs FISH | Posttranslational modification, protein turnover, chaperones |
| gene_195253 | 17,48999375 | -4,616431192 | 1,117992882 | -4,129213401 | 3,64E-05 | 0,019827681 | The Beehive: PERISCOP vs FISH | Replication, recombination and repair |
| gene_921319 | 66,42635243 | -3,428843917 | 0,806713281 | -4,250387341 | 2,13E-05 | 0,013466756 | The Beehive: PERISCOP vs FISH | Replication, recombination and repair |
| gene_817362 | 364,0178505 | 2,198887673 | 0,579218673 | 3,796299699 | 0,000146872 | 0,041873641 | The Beehive: PERISCOP vs FISH | Replication, recombination and repair |
| gene_1618833 | 1019,915593 | 2,25234915 | 0,514532919 | 4,377463653 | 1,20E-05 | 0,010838058 | The Beehive: PERISCOP vs FISH | Replication, recombination and repair |
| gene_779191 | 51,13485044 | 3,309003698 | 0,861991922 | 3,838787366 | 0,000123643 | 0,040270798 | The Beehive: PERISCOP vs FISH | Replication, recombination and repair |
| gene_775672 | 38,47788915 | 3,461668119 | 0,81359335 | 4,254789103 | 2,09E-05 | 0,013466756 | The Beehive: PERISCOP vs FISH | Replication, recombination and repair |
| gene_293095 | 34,14302994 | 4,886994012 | 0,912457219 | 5,355860974 | 8,51E-08 | 0,000448357 | The Beehive: PERISCOP vs FISH | Replication, recombination and repair |
| gene_1174029 | 28,7692047 | 5,050386742 | 1,14038195 | 4,428680008 | 9,48E-06 | 0,009662522 | The Beehive: PERISCOP vs FISH | Replication, recombination and repair |
| gene_407800 | 18,15717158 | 7,78715726 | 1,495416041 | 5,207351699 | 1,92E-07 | 0,000672421 | The Beehive: PERISCOP vs FISH | Replication, recombination and repair |
| gene_1786365 | 93,43602526 | 3,114430242 | 0,612866523 | 5,081743132 | 3,74E-07 | 0,000978771 | The Beehive: PERISCOP vs FISH | Secondary metabolites biosynthesis, transport and catabolism |
| gene_1786366 | 40,83826516 | 3,172108432 | 0,836604161 | 3,791647926 | 0,000149651 | 0,041873641 | The Beehive: PERISCOP vs FISH | Secondary metabolites biosynthesis, transport and catabolism |
| gene_1181929 | 115,429648 | -3,645051773 | 0,928634785 | -3,925172555 | 8,67E-05 | 0,03380357 | The Beehive: PERISCOP vs FISH | Signal transduction mechanisms |
| gene_1430521 | 46,94458025 | -2,515891726 | 0,665826908 | -3,778597255 | 0,000157714 | 0,041873641 | The Beehive: PERISCOP vs FISH | Signal transduction mechanisms |
| gene_988779 | 80,20496131 | -2,510901586 | 0,617831836 | -4,064053417 | 4,82E-05 | 0,02440368 | The Beehive: PERISCOP vs FISH | Signal transduction mechanisms |
| gene_1430522 | 144,4448771 | -2,418752299 | 0,645980449 | -3,744311926 | 0,000180889 | 0,043960156 | The Beehive: PERISCOP vs FISH | Signal transduction mechanisms |
| gene_215511 | 60,29137953 | 3,682430857 | 0,843771696 | 4,364250273 | 1,28E-05 | 0,010839327 | The Beehive: PERISCOP vs FISH | Signal transduction mechanisms |
| gene_1856083 | 40,5308481 | 3,816436202 | 0,902486087 | 4,228803368 | 2,35E-05 | 0,013745153 | The Beehive: PERISCOP vs FISH | Signal transduction mechanisms |
| gene_407796 | 14,69192421 | 5,040303634 | 1,2892319 | 3,909539962 | 9,25E-05 | 0,035324735 | The Beehive: PERISCOP vs FISH | Signal transduction mechanisms |
| gene_30369 | 68,9361762 | 2,565613691 | 0,677318591 | 3,787897933 | 0,000151927 | 0,041873641 | The Beehive: PERISCOP vs FISH | Transcription |
| gene_607909 | 81,35401111 | 3,117868632 | 0,831301129 | 3,750588716 | 0,00017642 | 0,043237512 | The Beehive: PERISCOP vs FISH | Transcription |
| gene_173139 | 20,20734838 | 4,144520272 | 0,994107961 | 4,169084683 | 3,06E-05 | 0,017253465 | The Beehive: PERISCOP vs FISH | Transcription |
| gene_79125 | 45,78036028 | -7,710096246 | 1,457837855 | -5,288720019 | 1,23E-07 | 0,000555925 | The Beehive: PERISCOP vs FISH | Translation, ribosomal structure and biogenesis |
| gene_778551 | 55,69096637 | -3,163122106 | 0,719162012 | -4,39834426 | 1,09E-05 | 0,01076925 | The Beehive: PERISCOP vs FISH | Translation, ribosomal structure and biogenesis |
| gene_696758 | 40,21595376 | -2,896937421 | 0,769572663 | -3,764345541 | 0,000166986 | 0,042890925 | The Beehive: PERISCOP vs FISH | Translation, ribosomal structure and biogenesis |
| gene_124686 | 90,584743 | -2,838691871 | 0,661374596 | -4,292109021 | 1,77E-05 | 0,012425457 | The Beehive: PERISCOP vs FISH | Translation, ribosomal structure and biogenesis |

| | | | | | | | | |
|---|---|---|---|---|---|---|---|---|
| gene_377145 | 1629,361261 | -2,266163561 | 0,594356736 | -3,812800332 | 0,000137401 | 0,041873641 | The Beehive: PERISCOP vs FISH | Translation, ribosomal structure and biogenesis |
| gene_310317 | 4445,468419 | -2,095217208 | 0,551836799 | -3,796805889 | 0,000146572 | 0,041873641 | The Beehive: PERISCOP vs FISH | Translation, ribosomal structure and biogenesis |
| gene_1277832 | 101,0418339 | 2,68407546 | 0,700605044 | 3,831082125 | 0,000127581 | 0,040306637 | The Beehive: PERISCOP vs FISH | Translation, ribosomal structure and biogenesis |
| gene_275658 | 129,819859 | 3,474691186 | 0,686325618 | 5,062744411 | 4,13E-07 | 0,000978771 | The Beehive: PERISCOP vs FISH | Translation, ribosomal structure and biogenesis |
| gene_728550 | 80,46073784 | 3,585648545 | 0,929040806 | 3,859516742 | 0,000113611 | 0,038850501 | The Beehive: PERISCOP vs FISH | Translation, ribosomal structure and biogenesis |
| gene_154893 | 12,72187482 | 4,5041844 | 1,213562917 | 3,711537603 | 0,000206004 | 0,049305198 | The Beehive: PERISCOP vs FISH | Translation, ribosomal structure and biogenesis |
| gene_1441696 | 12,42408994 | -6,337089792 | 1,565990118 | -4,046698455 | 5,19E-05 | 0,003223638 | The Moose: suction sampler vs FISH | Amino acid transport and metabolism |
| gene_54972 | 4,928503211 | -6,07251227 | 1,918039529 | -3,165999541 | 0,00154551 | 0,028384599 | The Moose: suction sampler vs FISH | Amino acid transport and metabolism |
| gene_219410 | 4,627233894 | -5,834640069 | 1,894849652 | -3,079210038 | 0,002075503 | 0,033957522 | The Moose: suction sampler vs FISH | Amino acid transport and metabolism |
| gene_1120300 | 16,8368425 | -5,257383329 | 1,273252802 | -4,129096219 | 3,64E-05 | 0,002525787 | The Moose: suction sampler vs FISH | Amino acid transport and metabolism |
| gene_89574 | 15,29484153 | -4,930616964 | 1,238172906 | -3,982171584 | 6,83E-05 | 0,003872931 | The Moose: suction sampler vs FISH | Amino acid transport and metabolism |
| gene_599481 | 7,206533409 | -4,380580404 | 1,503128406 | -2,914308843 | 0,00356477 | 0,046569649 | The Moose: suction sampler vs FISH | Amino acid transport and metabolism |
| gene_1653560 | 9,1389108 | -4,277887295 | 1,412925905 | -3,027679852 | 0,00246439 | 0,037427034 | The Moose: suction sampler vs FISH | Amino acid transport and metabolism |
| gene_467240 | 10,10920064 | -4,224790196 | 1,334382478 | -3,166101373 | 0,001544969 | 0,028381761 | The Moose: suction sampler vs FISH | Amino acid transport and metabolism |
| gene_351146 | 91,81090198 | -4,199537544 | 0,857951133 | -4,894844685 | 9,84E-07 | 0,000220298 | The Moose: suction sampler vs FISH | Amino acid transport and metabolism |
| gene_322058 | 14,92654659 | -4,01291986 | 1,090580836 | -3,679617071 | 0,000233584 | 0,008653593 | The Moose: suction sampler vs FISH | Amino acid transport and metabolism |
| gene_351145 | 11,85942825 | -3,991608522 | 1,120650257 | -3,561868206 | 0,000368225 | 0,011547524 | The Moose: suction sampler vs FISH | Amino acid transport and metabolism |
| gene_89570 | 117,0444104 | -3,931243321 | 0,621995331 | -6,320374327 | 2,61E-10 | 7,06E-07 | The Moose: suction sampler vs FISH | Amino acid transport and metabolism |
| gene_1380996 | 36,86730873 | -3,907968458 | 1,256850695 | -3,109333889 | 0,001875097 | 0,031967616 | The Moose: suction sampler vs FISH | Amino acid transport and metabolism |
| gene_1198998 | 14,09508986 | -3,671103903 | 1,252307271 | -2,931472162 | 0,003373596 | 0,045066162 | The Moose: suction sampler vs FISH | Amino acid transport and metabolism |
| gene_1120290 | 12,93849382 | -3,65057572 | 1,183643201 | -3,084185941 | 0,0020411 | 0,033619327 | The Moose: suction sampler vs FISH | Amino acid transport and metabolism |
| gene_1181670 | 20,5766113 | -3,532199496 | 1,003459587 | -3,520021673 | 0,000431512 | 0,012753467 | The Moose: suction sampler vs FISH | Amino acid transport and metabolism |
| gene_403609 | 28,96091081 | -3,461515913 | 0,914601684 | -3,784725058 | 0,000153879 | 0,006570706 | The Moose: suction sampler vs FISH | Amino acid transport and metabolism |
| gene_629505 | 17,5770017 | -3,420927273 | 1,118624927 | -3,058153981 | 0,002227051 | 0,035228466 | The Moose: suction sampler vs FISH | Amino acid transport and metabolism |
| gene_873933 | 586,3424722 | -3,370932545 | 0,75225682 | -4,481092698 | 7,43E-06 | 0,000881125 | The Moose: suction sampler vs FISH | Amino acid transport and metabolism |
| gene_848653 | 13,02433378 | -3,345248291 | 1,153269306 | -2,900665328 | 0,003723713 | 0,047879576 | The Moose: suction sampler vs FISH | Amino acid transport and metabolism |
| gene_951970 | 86,73260247 | -3,294652386 | 0,751321096 | -4,385145583 | 1,16E-05 | 0,001172573 | The Moose: suction sampler vs FISH | Amino acid transport and metabolism |
| gene_589827 | 21,10337748 | -3,221308641 | 0,965287485 | -3,337149492 | 0,000846424 | 0,019703841 | The Moose: suction sampler vs FISH | Amino acid transport and metabolism |
| gene_128624 | 23,47082861 | -3,214403961 | 0,969632386 | -3,315074875 | 0,000916186 | 0,020621902 | The Moose: suction sampler vs FISH | Amino acid transport and metabolism |
| gene_24052 | 100,9624919 | -2,970982224 | 0,690813691 | -4,300699684 | 1,70E-05 | 0,001501167 | The Moose: suction sampler vs FISH | Amino acid transport and metabolism |
| gene_36890 | 68,49410176 | -2,946517991 | 0,733806727 | -4,015386996 | 5,93E-05 | 0,003529424 | The Moose: suction sampler vs FISH | Amino acid transport and metabolism |
| gene_308301 | 349,7251989 | -2,791735301 | 0,650279471 | -4,293131533 | 1,76E-05 | 0,001531225 | The Moose: suction sampler vs FISH | Amino acid transport and metabolism |
| gene_1508228 | 130,0572728 | -2,73974634 | 0,692200776 | -3,958022633 | 7,56E-05 | 0,004163874 | The Moose: suction sampler vs FISH | Amino acid transport and metabolism |
| gene_829492 | 41,30037512 | -2,64709655 | 0,78014008 | -3,39310416 | 0,000691054 | 0,017165926 | The Moose: suction sampler vs FISH | Amino acid transport and metabolism |
| gene_411971 | 59,95123573 | -2,63144892 | 0,766859822 | -3,431460151 | 0,000600341 | 0,015635485 | The Moose: suction sampler vs FISH | Amino acid transport and metabolism |
| gene_629503 | 117,7939079 | -2,559364805 | 0,725711408 | -3,52669777 | 0,000420777 | 0,01257792 | The Moose: suction sampler vs FISH | Amino acid transport and metabolism |
| gene_589836 | 65,34540114 | -2,511990763 | 0,77563398 | -3,23862908 | 0,001201057 | 0,024359814 | The Moose: suction sampler vs FISH | Amino acid transport and metabolism |
| gene_951971 | 24,54171472 | -2,493991267 | 0,784580127 | -3,178759161 | 0,001479069 | 0,027669446 | The Moose: suction sampler vs FISH | Amino acid transport and metabolism |
| gene_1604241 | 78,74721794 | -2,485179928 | 0,703786647 | -3,531155271 | 0,000413749 | 0,01242854 | The Moose: suction sampler vs FISH | Amino acid transport and metabolism |
| gene_1224779 | 34,4770065 | -2,437596159 | 0,769722534 | -3,166850459 | 0,001540995 | 0,028358408 | The Moose: suction sampler vs FISH | Amino acid transport and metabolism |
| gene_791203 | 56,86186283 | -2,39947449 | 0,771794438 | -3,108955404 | 0,001877501 | 0,031993742 | The Moose: suction sampler vs FISH | Amino acid transport and metabolism |
| gene_553460 | 219,1723909 | -2,359647708 | 0,687385444 | -3,432786842 | 0,000597412 | 0,0155979 | The Moose: suction sampler vs FISH | Amino acid transport and metabolism |
| gene_87660 | 4042,105806 | -2,348252038 | 0,592970348 | -3,960150869 | 7,49E-05 | 0,004142476 | The Moose: suction sampler vs FISH | Amino acid transport and metabolism |
| gene_1802818 | 81,66446378 | -2,332420302 | 0,685189387 | -3,404051999 | 0,000663941 | 0,016737807 | The Moose: suction sampler vs FISH | Amino acid transport and metabolism |
| gene_774962 | 2560,871546 | -2,182508757 | 0,575796863 | -3,790414465 | 0,000150396 | 0,006479324 | The Moose: suction sampler vs FISH | Amino acid transport and metabolism |
| gene_1120289 | 66,14517247 | -2,16739291 | 0,691720093 | -3,133338082 | 0,001728302 | 0,030426115 | The Moose: suction sampler vs FISH | Amino acid transport and metabolism |
| gene_791217 | 76,14175145 | -2,162154965 | 0,736984626 | -2,93378571 | 0,003348553 | 0,04493566 | The Moose: suction sampler vs FISH | Amino acid transport and metabolism |
| gene_70409 | 132,6657276 | -2,114788601 | 0,663637935 | -3,186660211 | 0,001439258 | 0,027180835 | The Moose: suction sampler vs FISH | Amino acid transport and metabolism |
| gene_1160776 | 34,03757023 | -2,094479717 | 0,710533019 | -2,947758459 | 0,003200871 | 0,043776571 | The Moose: suction sampler vs FISH | Amino acid transport and metabolism |
| gene_829491 | 132,0694322 | -2,088875993 | 0,709983583 | -2,942146891 | 0,003259453 | 0,044192451 | The Moose: suction sampler vs FISH | Amino acid transport and metabolism |
| gene_431767 | 4047,08436 | -2,07962623 | 0,548741956 | -3,789807226 | 0,000150764 | 0,006482951 | The Moose: suction sampler vs FISH | Amino acid transport and metabolism |
| gene_198820 | 995,4050886 | -2,070038301 | 0,579015754 | -3,575098407 | 0,000350096 | 0,011179485 | The Moose: suction sampler vs FISH | Amino acid transport and metabolism |
| gene_265219 | 7845,657106 | -2,065754895 | 0,571465116 | -3,614839884 | 0,000300533 | 0,010214099 | The Moose: suction sampler vs FISH | Amino acid transport and metabolism |
| gene_108564 | 88,66508212 | -2,063818862 | 0,663986632 | -3,108223513 | 0,001882157 | 0,032028098 | The Moose: suction sampler vs FISH | Amino acid transport and metabolism |
| gene_745443 | 125,194804 | -2,057355517 | 0,699693809 | -2,940365472 | 0,003278253 | 0,044349108 | The Moose: suction sampler vs FISH | Amino acid transport and metabolism |
| gene_87663 | 3992,050355 | -2,048921593 | 0,670631311 | -3,05521314 | 0,002249006 | 0,035423176 | The Moose: suction sampler vs FISH | Amino acid transport and metabolism |
| gene_1891564 | 286,2897242 | 2,237851213 | 0,678231539 | 3,299538703 | 0,000968439 | 0,021308263 | The Moose: suction sampler vs FISH | Amino acid transport and metabolism |
| gene_641988 | 74,98146844 | 2,274198693 | 0,673876685 | 3,374799491 | 0,000738695 | 0,01799451 | The Moose: suction sampler vs FISH | Amino acid transport and metabolism |
| gene_723697 | 77,57808372 | 2,311431247 | 0,736144523 | 3,139915022 | 0,001689968 | 0,030024126 | The Moose: suction sampler vs FISH | Amino acid transport and metabolism |
| gene_110600 | 158,4264345 | 2,340392647 | 0,739203298 | 3,166101471 | 0,001544969 | 0,028381761 | The Moose: suction sampler vs FISH | Amino acid transport and metabolism |
| gene_1901488 | 100,37643 | 2,506056977 | 0,696557206 | 3,597776257 | 0,00032095 | 0,01061808 | The Moose: suction sampler vs FISH | Amino acid transport and metabolism |
| gene_428023 | 29,75445574 | 2,624939447 | 0,805097597 | 3,260399058 | 0,001112556 | 0,023266414 | The Moose: suction sampler vs FISH | Amino acid transport and metabolism |
| gene_271409 | 14,53444184 | 2,720555227 | 0,921060783 | 2,953719533 | 0,003139692 | 0,043306684 | The Moose: suction sampler vs FISH | Amino acid transport and metabolism |
| gene_298402 | 49,87570627 | 2,815007924 | 0,814273682 | 3,457078358 | 0,000546066 | 0,01475021 | The Moose: suction sampler vs FISH | Amino acid transport and metabolism |
| gene_1646605 | 45,58592431 | 2,886493159 | 0,827960775 | 3,486268007 | 0,00048981 | 0,013772625 | The Moose: suction sampler vs FISH | Amino acid transport and metabolism |

| | | | | | | | | |
|---|---|---|---|---|---|---|---|---|
| gene_266395 | 11,62507687 | 2,96952138 | 0,926239057 | 3,205998882 | 0,001345945 | 0,026068813 | The Moose: suction sampler vs FISH | Amino acid transport and metabolism |
| gene_1503956 | 42,92808241 | 3,0166406 | 0,887205617 | 3,40015949 | 0,000673466 | 0,01689814 | The Moose: suction sampler vs FISH | Amino acid transport and metabolism |
| gene_1699828 | 13,48805616 | 3,107201641 | 1,048737461 | 2,962802185 | 0,003048524 | 0,042546401 | The Moose: suction sampler vs FISH | Amino acid transport and metabolism |
| gene_1613020 | 7,687477124 | 3,204437692 | 1,097642318 | 2,919382424 | 0,003507257 | 0,046082171 | The Moose: suction sampler vs FISH | Amino acid transport and metabolism |
| gene_1141617 | 51,67847964 | 3,286103587 | 1,041449744 | 3,155316525 | 0,001603241 | 0,029057674 | The Moose: suction sampler vs FISH | Amino acid transport and metabolism |
| gene_430185 | 231,0909489 | 3,376753856 | 0,940787601 | 3,589283969 | 0,000331587 | 0,010843906 | The Moose: suction sampler vs FISH | Amino acid transport and metabolism |
| gene_146855 | 56,19389619 | 3,500549192 | 1,014975171 | 3,448901305 | 0,000562872 | 0,015060169 | The Moose: suction sampler vs FISH | Amino acid transport and metabolism |
| gene_306390 | 23,85638262 | 3,530986799 | 1,132343042 | 3,118301315 | 0,001818967 | 0,031351249 | The Moose: suction sampler vs FISH | Amino acid transport and metabolism |
| gene_1547095 | 30,75089469 | 3,538777394 | 1,019952897 | 3,469549824 | 0,000521331 | 0,014321278 | The Moose: suction sampler vs FISH | Amino acid transport and metabolism |
| gene_1229784 | 6,537341482 | 3,58677553 | 1,236011999 | 2,901893778 | 0,003709143 | 0,047775868 | The Moose: suction sampler vs FISH | Amino acid transport and metabolism |
| gene_876303 | 96,67108936 | 3,623651012 | 0,865757109 | 4,185528449 | 2,85E-05 | 0,002149728 | The Moose: suction sampler vs FISH | Amino acid transport and metabolism |
| gene_338660 | 15,01737278 | 3,729867994 | 1,171966355 | 3,182572586 | 0,001459729 | 0,027426404 | The Moose: suction sampler vs FISH | Amino acid transport and metabolism |
| gene_1886443 | 69,8753473 | 3,730759876 | 0,894617312 | 4,170229914 | 3,04E-05 | 0,002237113 | The Moose: suction sampler vs FISH | Amino acid transport and metabolism |
| gene_1093419 | 174,8540169 | 3,776775223 | 0,796277963 | 4,743036225 | 2,11E-06 | 0,000359605 | The Moose: suction sampler vs FISH | Amino acid transport and metabolism |
| gene_1203740 | 12,55030619 | 3,784286901 | 1,17615887 | 3,217496375 | 0,001293147 | 0,025471058 | The Moose: suction sampler vs FISH | Amino acid transport and metabolism |
| gene_1290837 | 9,461275065 | 3,79397464 | 1,172230548 | 3,236543057 | 0,00120987 | 0,02444536 | The Moose: suction sampler vs FISH | Amino acid transport and metabolism |
| gene_824777 | 156,8446252 | 3,795848949 | 0,934980108 | 4,059817867 | 4,91E-05 | 0,003123845 | The Moose: suction sampler vs FISH | Amino acid transport and metabolism |
| gene_563727 | 15,76564636 | 3,960047991 | 1,083665937 | 3,654306975 | 0,000257878 | 0,009225657 | The Moose: suction sampler vs FISH | Amino acid transport and metabolism |
| gene_25545 | 54,84484721 | 4,047621845 | 1,001635603 | 4,041012351 | 5,32E-05 | 0,003279206 | The Moose: suction sampler vs FISH | Amino acid transport and metabolism |
| gene_338136 | 6,489607495 | 4,180923961 | 1,36281687 | 3,067854568 | 0,002156015 | 0,034679924 | The Moose: suction sampler vs FISH | Amino acid transport and metabolism |
| gene_1809366 | 10,9119346 | 4,195947052 | 1,396942834 | 3,003664109 | 0,002667496 | 0,039220187 | The Moose: suction sampler vs FISH | Amino acid transport and metabolism |
| gene_1231405 | 13,54118529 | 4,347639691 | 1,274674113 | 3,410785272 | 0,000647761 | 0,016467565 | The Moose: suction sampler vs FISH | Amino acid transport and metabolism |
| gene_1203741 | 8,666361015 | 4,568471214 | 1,478606881 | 3,089713211 | 0,002003498 | 0,033216015 | The Moose: suction sampler vs FISH | Amino acid transport and metabolism |
| gene_313983 | 28,35215419 | 4,730511547 | 1,317715388 | 3,589934208 | 0,000330761 | 0,010831687 | The Moose: suction sampler vs FISH | Amino acid transport and metabolism |
| gene_1729173 | 5,503786209 | 4,930026531 | 1,691343382 | 2,91485844 | 0,003558499 | 0,046527013 | The Moose: suction sampler vs FISH | Amino acid transport and metabolism |
| gene_1325528 | 10,51085781 | 4,964728168 | 1,51720013 | 3,272296165 | 0,001066778 | 0,022683339 | The Moose: suction sampler vs FISH | Amino acid transport and metabolism |
| gene_64838 | 6,30953774 | 5,175657472 | 1,559026687 | 3,319800433 | 0,000900818 | 0,02049231 | The Moose: suction sampler vs FISH | Amino acid transport and metabolism |
| gene_1502027 | 5,693556911 | 5,363989178 | 1,702246435 | 3,151123756 | 0,001626436 | 0,029328152 | The Moose: suction sampler vs FISH | Amino acid transport and metabolism |
| gene_1079659 | 5,227761363 | 5,413094517 | 1,82648008 | 2,963675639 | 0,003039885 | 0,042472454 | The Moose: suction sampler vs FISH | Amino acid transport and metabolism |
| gene_520920 | 4,287095186 | 5,620742225 | 1,790742526 | 3,13877743 | 0,001696543 | 0,030082389 | The Moose: suction sampler vs FISH | Amino acid transport and metabolism |
| gene_773400 | 3,83048958 | 5,623274389 | 1,850304487 | 3,039107579 | 0,002372801 | 0,036523603 | The Moose: suction sampler vs FISH | Amino acid transport and metabolism |
| gene_1114593 | 7,653139566 | 5,630303779 | 1,776010197 | 3,17019789 | 0,001523352 | 0,028160723 | The Moose: suction sampler vs FISH | Amino acid transport and metabolism |
| gene_306391 | 6,259261759 | 5,648322795 | 1,627850332 | 3,469804739 | 0,000520837 | 0,014316191 | The Moose: suction sampler vs FISH | Amino acid transport and metabolism |
| gene_381459 | 4,185570421 | 5,879721047 | 1,791328572 | 3,282324158 | 0,001029552 | 0,022148632 | The Moose: suction sampler vs FISH | Amino acid transport and metabolism |
| gene_563726 | 4,130733649 | 5,999700046 | 1,990846784 | 3,013642282 | 0,00258132 | 0,038557052 | The Moose: suction sampler vs FISH | Amino acid transport and metabolism |
| gene_459706 | 5,610942098 | 6,13118897 | 1,952080791 | 3,14084796 | 0,001684595 | 0,029972233 | The Moose: suction sampler vs FISH | Amino acid transport and metabolism |
| gene_206289 | 4,792581708 | 6,246998209 | 1,780937397 | 3,507702304 | 0,000451995 | 0,013116056 | The Moose: suction sampler vs FISH | Amino acid transport and metabolism |
| gene_500745 | 29,59142688 | 6,328441747 | 1,329876986 | 4,758667016 | 1,95E-06 | 0,000344767 | The Moose: suction sampler vs FISH | Amino acid transport and metabolism |
| gene_597074 | 7,141917568 | 6,40551939 | 1,757302772 | 3,645085805 | 0,000267303 | 0,009447565 | The Moose: suction sampler vs FISH | Amino acid transport and metabolism |
| gene_1085926 | 5,232999333 | 6,431032298 | 1,902732418 | 3,379893167 | 0,00072514 | 0,017746723 | The Moose: suction sampler vs FISH | Amino acid transport and metabolism |
| gene_1948504 | 3,937418201 | 6,485312164 | 2,223281052 | 2,917000601 | 0,003534151 | 0,046284899 | The Moose: suction sampler vs FISH | Amino acid transport and metabolism |
| gene_1162983 | 4,160503495 | 6,638517927 | 1,909472214 | 3,476624525 | 0,000507768 | 0,014088798 | The Moose: suction sampler vs FISH | Amino acid transport and metabolism |
| gene_1366092 | 3,534135515 | 6,715526265 | 2,311652277 | 2,905076309 | 0,003671636 | 0,0474274 | The Moose: suction sampler vs FISH | Amino acid transport and metabolism |
| gene_1285530 | 11,22276791 | 7,007945424 | 1,631904831 | 4,294334628 | 1,75E-05 | 0,001530091 | The Moose: suction sampler vs FISH | Amino acid transport and metabolism |
| gene_1774485 | 8,186644701 | 7,181963022 | 1,943297814 | 3,695760356 | 0,00021923 | 0,008269812 | The Moose: suction sampler vs FISH | Amino acid transport and metabolism |
| gene_1520935 | 5,128725108 | 7,479208463 | 2,207091719 | 3,388716653 | 0,000702205 | 0,017341449 | The Moose: suction sampler vs FISH | Amino acid transport and metabolism |
| gene_878183 | 28,4212425 | 7,735009177 | 2,057212449 | 3,759946708 | 0,00016995 | 0,007032082 | The Moose: suction sampler vs FISH | Amino acid transport and metabolism |
| gene_1513920 | 6,952063871 | 8,04191156 | 2,058176922 | 3,907298481 | 9,33E-05 | 0,004790289 | The Moose: suction sampler vs FISH | Amino acid transport and metabolism |
| gene_760927 | 16,28574829 | 9,217380638 | 1,644398565 | 5,605320289 | 2,08E-08 | 1,53E-05 | The Moose: suction sampler vs FISH | Amino acid transport and metabolism |
| gene_498652 | 4,525016907 | -6,043563392 | 1,911739361 | -3,161290455 | 0,001570718 | 0,02868807 | The Moose: suction sampler vs FISH | Carbohydrate transport and metabolism |
| gene_89573 | 21,72809875 | -5,632975409 | 1,287880967 | -4,373832329 | 1,22E-05 | 0,001206793 | The Moose: suction sampler vs FISH | Carbohydrate transport and metabolism |
| gene_501019 | 5,059424167 | -5,429547285 | 1,704842214 | -3,18477994 | 0,001448641 | 0,027306255 | The Moose: suction sampler vs FISH | Carbohydrate transport and metabolism |
| gene_603346 | 32,6264198 | -4,924434427 | 1,131358986 | -4,352671864 | 1,34E-05 | 0,001295729 | The Moose: suction sampler vs FISH | Carbohydrate transport and metabolism |
| gene_400613 | 9,754730953 | -4,702662862 | 1,417657983 | -3,317205504 | 0,000909227 | 0,020578794 | The Moose: suction sampler vs FISH | Carbohydrate transport and metabolism |
| gene_89572 | 29,64705164 | -4,687040804 | 0,98732468 | -4,747213249 | 2,06E-06 | 0,000355568 | The Moose: suction sampler vs FISH | Carbohydrate transport and metabolism |
| gene_832032 | 98,79144943 | -4,538284336 | 0,798805344 | -5,681339481 | 1,34E-08 | 1,08E-05 | The Moose: suction sampler vs FISH | Carbohydrate transport and metabolism |
| gene_89571 | 236,7258376 | -4,455706855 | 0,606348163 | -7,348429705 | 2,01E-13 | 2,95E-09 | The Moose: suction sampler vs FISH | Carbohydrate transport and metabolism |
| gene_1380997 | 7,597293642 | -4,066198473 | 1,411072903 | -2,881635997 | 0,003956165 | 0,049499236 | The Moose: suction sampler vs FISH | Carbohydrate transport and metabolism |
| gene_138944 | 31,38224443 | -3,304545776 | 1,068091663 | -3,093878448 | 0,001975584 | 0,032945002 | The Moose: suction sampler vs FISH | Carbohydrate transport and metabolism |
| gene_138945 | 99,43691943 | -3,186003354 | 0,735005946 | -4,334663374 | 1,46E-05 | 0,001350346 | The Moose: suction sampler vs FISH | Carbohydrate transport and metabolism |
| gene_1039867 | 25,1298542 | -2,733043489 | 0,875212747 | -3,122719016 | 0,001791887 | 0,031046275 | The Moose: suction sampler vs FISH | Carbohydrate transport and metabolism |
| gene_138943 | 66,04093717 | -2,729455293 | 0,905247461 | -3,015148245 | 0,002568537 | 0,038435586 | The Moose: suction sampler vs FISH | Carbohydrate transport and metabolism |
| gene_829493 | 45,98362856 | -2,549397939 | 0,818862526 | -3,113340588 | 0,001849824 | 0,031680238 | The Moose: suction sampler vs FISH | Carbohydrate transport and metabolism |
| gene_571451 | 573,7815055 | -2,434252021 | 0,600182377 | -4,055853875 | 5,00E-05 | 0,003159937 | The Moose: suction sampler vs FISH | Carbohydrate transport and metabolism |
| gene_791198 | 61,35800484 | -2,362764616 | 0,720360383 | -3,279975792 | 0,00103816 | 0,022281606 | The Moose: suction sampler vs FISH | Carbohydrate transport and metabolism |

| | | | | | | | | |
|---|---|---|---|---|---|---|---|---|
| gene_364928 | 33,74958451 | -2,228393375 | 0,677246948 | -3,290370496 | 0,001000555 | 0,021744748 | The Moose: suction sampler vs FISH | Carbohydrate transport and metabolism |
| gene_473522 | 48,26084954 | -2,207509365 | 0,749121689 | -2,94679676 | 0,003210842 | 0,043881701 | The Moose: suction sampler vs FISH | Carbohydrate transport and metabolism |
| gene_24051 | 103,2074428 | -2,089658049 | 0,689517692 | -3,030608314 | 0,002440616 | 0,03717118 | The Moose: suction sampler vs FISH | Carbohydrate transport and metabolism |
| gene_1323892 | 3285,863095 | -2,063983156 | 0,642709039 | -3,211380316 | 0,00132099 | 0,025815906 | The Moose: suction sampler vs FISH | Carbohydrate transport and metabolism |
| gene_1362175 | 201,6335185 | -2,003076335 | 0,624104769 | -3,209519355 | 0,001329571 | 0,025870505 | The Moose: suction sampler vs FISH | Carbohydrate transport and metabolism |
| gene_123067 | 273,122022 | 2,028944991 | 0,701845778 | 2,890870126 | 0,003841768 | 0,048690024 | The Moose: suction sampler vs FISH | Carbohydrate transport and metabolism |
| gene_1113087 | 147,3017305 | 2,048024714 | 0,685722058 | 2,986668855 | 0,00282035 | 0,040618761 | The Moose: suction sampler vs FISH | Carbohydrate transport and metabolism |
| gene_277302 | 77,31169776 | 2,176740812 | 0,687251628 | 3,167312704 | 0,001538548 | 0,028348882 | The Moose: suction sampler vs FISH | Carbohydrate transport and metabolism |
| gene_418814 | 534,9744714 | 2,374865572 | 0,742549459 | 3,198259111 | 0,0013826 | 0,026497228 | The Moose: suction sampler vs FISH | Carbohydrate transport and metabolism |
| gene_774873 | 513,0182248 | 2,449596379 | 0,826166317 | 2,965016037 | 0,003026672 | 0,042390139 | The Moose: suction sampler vs FISH | Carbohydrate transport and metabolism |
| gene_293161 | 51,95799162 | 2,557841006 | 0,8328517 | 3,07118423 | 0,002132116 | 0,034462996 | The Moose: suction sampler vs FISH | Carbohydrate transport and metabolism |
| gene_107106 | 58,50696725 | 2,717156035 | 0,83855894 | 3,240268401 | 0,001194172 | 0,024268398 | The Moose: suction sampler vs FISH | Carbohydrate transport and metabolism |
| gene_516923 | 99,47774941 | 2,757573105 | 0,93820906 | 2,939188314 | 0,003290731 | 0,044468761 | The Moose: suction sampler vs FISH | Carbohydrate transport and metabolism |
| gene_602329 | 49,04507269 | 2,784132708 | 0,81925967 | 3,398351963 | 0,000677931 | 0,016970943 | The Moose: suction sampler vs FISH | Carbohydrate transport and metabolism |
| gene_472456 | 38,60909836 | 2,83204204 | 0,871617873 | 3,249178485 | 0,001157388 | 0,023837461 | The Moose: suction sampler vs FISH | Carbohydrate transport and metabolism |
| gene_757655 | 23,44551524 | 2,838534586 | 0,908281279 | 3,125171302 | 0,001777015 | 0,030868708 | The Moose: suction sampler vs FISH | Carbohydrate transport and metabolism |
| gene_1764239 | 236,4491348 | 2,871590776 | 0,804927933 | 3,567512891 | 0,000360386 | 0,011403931 | The Moose: suction sampler vs FISH | Carbohydrate transport and metabolism |
| gene_716384 | 10,14631321 | 3,033111373 | 0,988511813 | 3,06836128 | 0,002152362 | 0,034648127 | The Moose: suction sampler vs FISH | Carbohydrate transport and metabolism |
| gene_15288 | 48,54036175 | 3,052263718 | 0,886742681 | 3,442107596 | 0,000577201 | 0,015287594 | The Moose: suction sampler vs FISH | Carbohydrate transport and metabolism |
| gene_472476 | 38,58198102 | 3,079019449 | 0,815997214 | 3,773321031 | 0,000161089 | 0,006804675 | The Moose: suction sampler vs FISH | Carbohydrate transport and metabolism |
| gene_329858 | 13,48819396 | 3,108317707 | 1,040480882 | 2,987385701 | 0,002813745 | 0,040552491 | The Moose: suction sampler vs FISH | Carbohydrate transport and metabolism |
| gene_1220994 | 33,44959987 | 3,143577265 | 1,074236264 | 2,926336944 | 0,003429792 | 0,045535325 | The Moose: suction sampler vs FISH | Carbohydrate transport and metabolism |
| gene_976379 | 194,9741304 | 3,172760488 | 0,755242342 | 4,200983327 | 2,66E-05 | 0,002048123 | The Moose: suction sampler vs FISH | Carbohydrate transport and metabolism |
| gene_219132 | 50,79459706 | 3,194599987 | 0,949771893 | 3,363544458 | 0,000769484 | 0,01845682 | The Moose: suction sampler vs FISH | Carbohydrate transport and metabolism |
| gene_1184782 | 20,12110657 | 3,210550112 | 0,999375225 | 3,212557238 | 0,001315589 | 0,025759386 | The Moose: suction sampler vs FISH | Carbohydrate transport and metabolism |
| gene_103917 | 12,76320248 | 3,267686556 | 1,080719002 | 3,023622747 | 0,002497677 | 0,037784114 | The Moose: suction sampler vs FISH | Carbohydrate transport and metabolism |
| gene_1391512 | 47,93642732 | 3,295997873 | 0,908963046 | 3,626107669 | 0,000287725 | 0,009911817 | The Moose: suction sampler vs FISH | Carbohydrate transport and metabolism |
| gene_655955 | 16,03517673 | 3,3924842 | 1,122679338 | 3,02177486 | 0,002512974 | 0,037931428 | The Moose: suction sampler vs FISH | Carbohydrate transport and metabolism |
| gene_1177662 | 28,49450566 | 3,432097614 | 1,010440535 | 3,396634927 | 0,000682199 | 0,017024844 | The Moose: suction sampler vs FISH | Carbohydrate transport and metabolism |
| gene_1635363 | 484,9921814 | 3,441152396 | 0,817880379 | 4,207403046 | 2,58E-05 | 0,002007468 | The Moose: suction sampler vs FISH | Carbohydrate transport and metabolism |
| gene_1417424 | 13,70810076 | 3,471190981 | 1,178563029 | 2,945273945 | 0,003226688 | 0,044000023 | The Moose: suction sampler vs FISH | Carbohydrate transport and metabolism |
| gene_1766143 | 20,2552763 | 3,518432696 | 0,900601466 | 3,906758792 | 9,35E-05 | 0,004794295 | The Moose: suction sampler vs FISH | Carbohydrate transport and metabolism |
| gene_999239 | 75,2676936 | 3,603764405 | 0,872848367 | 4,128740501 | 3,65E-05 | 0,00252731 | The Moose: suction sampler vs FISH | Carbohydrate transport and metabolism |
| gene_1483432 | 16,68514551 | 3,662259485 | 1,151426796 | 3,180627287 | 0,001469566 | 0,027575945 | The Moose: suction sampler vs FISH | Carbohydrate transport and metabolism |
| gene_1442355 | 26,24932856 | 3,688489587 | 1,062346342 | 3,472021734 | 0,000516555 | 0,014230438 | The Moose: suction sampler vs FISH | Carbohydrate transport and metabolism |
| gene_813165 | 47,07645642 | 3,774449837 | 0,934646369 | 4,038372117 | 5,38E-05 | 0,003306916 | The Moose: suction sampler vs FISH | Carbohydrate transport and metabolism |
| gene_1503055 | 11,46383433 | 3,77501893 | 1,152461788 | 3,2756131 | 0,001054329 | 0,022529876 | The Moose: suction sampler vs FISH | Carbohydrate transport and metabolism |
| gene_1792430 | 11,94519042 | 3,86435505 | 1,20971409 | 3,19443667 | 0,00140104 | 0,026706306 | The Moose: suction sampler vs FISH | Carbohydrate transport and metabolism |
| gene_1253355 | 138,1379165 | 4,11961318 | 0,886601259 | 4,646523043 | 3,38E-06 | 0,000495866 | The Moose: suction sampler vs FISH | Carbohydrate transport and metabolism |
| gene_1177663 | 9,656844199 | 4,126043716 | 1,299984787 | 3,173916923 | 0,001503967 | 0,02797844 | The Moose: suction sampler vs FISH | Carbohydrate transport and metabolism |
| gene_1865660 | 12,53570283 | 4,172145623 | 1,204425724 | 3,464012382 | 0,000532182 | 0,014507026 | The Moose: suction sampler vs FISH | Carbohydrate transport and metabolism |
| gene_982805 | 23,18240474 | 4,183028792 | 1,035879051 | 4,038144018 | 5,39E-05 | 0,003306916 | The Moose: suction sampler vs FISH | Carbohydrate transport and metabolism |
| gene_1291393 | 8,64895226 | 4,209425831 | 1,355690803 | 3,105004343 | 0,001902762 | 0,032229792 | The Moose: suction sampler vs FISH | Carbohydrate transport and metabolism |
| gene_1200531 | 31,64404324 | 4,212292907 | 1,065258945 | 3,954243169 | 7,68E-05 | 0,004205003 | The Moose: suction sampler vs FISH | Carbohydrate transport and metabolism |
| gene_207289 | 200,0811885 | 4,218102736 | 0,973177615 | 4,334360625 | 1,46E-05 | 0,001350505 | The Moose: suction sampler vs FISH | Carbohydrate transport and metabolism |
| gene_1941209 | 16,78791663 | 4,224377626 | 1,127026118 | 3,748251758 | 0,000178071 | 0,007226628 | The Moose: suction sampler vs FISH | Carbohydrate transport and metabolism |
| gene_1791108 | 25,94323808 | 4,228808814 | 1,070580565 | 3,950014554 | 7,81E-05 | 0,004249399 | The Moose: suction sampler vs FISH | Carbohydrate transport and metabolism |
| gene_1822431 | 22,47279473 | 4,257649286 | 1,096037144 | 3,884584853 | 0,000102505 | 0,00506993 | The Moose: suction sampler vs FISH | Carbohydrate transport and metabolism |
| gene_1200530 | 221,6933227 | 4,26065803 | 0,881611556 | 4,83280647 | 1,35E-06 | 0,000271629 | The Moose: suction sampler vs FISH | Carbohydrate transport and metabolism |
| gene_1200528 | 125,3070123 | 4,272072252 | 0,987497897 | 4,32615833 | 1,52E-05 | 0,001387794 | The Moose: suction sampler vs FISH | Carbohydrate transport and metabolism |
| gene_1766144 | 33,12034352 | 4,282864491 | 0,972211272 | 4,405281665 | 1,06E-05 | 0,001103728 | The Moose: suction sampler vs FISH | Carbohydrate transport and metabolism |
| gene_463812 | 15,70100145 | 4,294187833 | 1,463660846 | 2,933868077 | 0,003347665 | 0,044932242 | The Moose: suction sampler vs FISH | Carbohydrate transport and metabolism |
| gene_1914761 | 5,841131389 | 4,354233048 | 1,450758105 | 3,001350144 | 0,002687853 | 0,039361021 | The Moose: suction sampler vs FISH | Carbohydrate transport and metabolism |
| gene_78157 | 16,0988708 | 4,393165011 | 1,408759914 | 3,118462534 | 0,001817973 | 0,031347689 | The Moose: suction sampler vs FISH | Carbohydrate transport and metabolism |
| gene_918096 | 6,699840331 | 4,402526924 | 1,391565705 | 3,163721921 | 0,001557655 | 0,028536295 | The Moose: suction sampler vs FISH | Carbohydrate transport and metabolism |
| gene_1047973 | 12,32558321 | 4,403455425 | 1,133099993 | 3,886201971 | 0,000101825 | 0,005059893 | The Moose: suction sampler vs FISH | Carbohydrate transport and metabolism |
| gene_1389352 | 1191,284647 | 4,491203803 | 0,888005053 | 5,057633158 | 4,24E-07 | 0,000123229 | The Moose: suction sampler vs FISH | Carbohydrate transport and metabolism |
| gene_1599611 | 36,48439802 | 4,510509061 | 0,955008454 | 4,723004325 | 2,32E-06 | 0,000386144 | The Moose: suction sampler vs FISH | Carbohydrate transport and metabolism |
| gene_1626919 | 19,55090889 | 4,620267059 | 1,075445988 | 4,296140495 | 1,74E-05 | 0,001523216 | The Moose: suction sampler vs FISH | Carbohydrate transport and metabolism |
| gene_1000791 | 14,20248207 | 4,785538029 | 1,224007848 | 3,909728223 | 9,24E-05 | 0,00475899 | The Moose: suction sampler vs FISH | Carbohydrate transport and metabolism |
| gene_435787 | 7,072496107 | 4,791142271 | 1,602261233 | 2,990237904 | 0,002787602 | 0,040294324 | The Moose: suction sampler vs FISH | Carbohydrate transport and metabolism |
| gene_844281 | 28,98095944 | 4,834500676 | 0,993321379 | 4,867005561 | 1,13E-06 | 0,000241903 | The Moose: suction sampler vs FISH | Carbohydrate transport and metabolism |
| gene_1200532 | 125,1982025 | 4,843245595 | 1,020810572 | 4,744509636 | 2,09E-06 | 0,000358666 | The Moose: suction sampler vs FISH | Carbohydrate transport and metabolism |
| gene_1200529 | 58,24829406 | 4,906775228 | 0,976439766 | 5,025169394 | 5,03E-07 | 0,000142073 | The Moose: suction sampler vs FISH | Carbohydrate transport and metabolism |
| gene_633199 | 28,96266966 | 4,956334453 | 1,114938429 | 4,445388486 | 8,77E-06 | 0,000973348 | The Moose: suction sampler vs FISH | Carbohydrate transport and metabolism |

| | | | | | | | | |
|---|---|---|---|---|---|---|---|---|
| gene_22887 | 10,25458092 | 5,023565786 | 1,704376128 | 2,94745139 | 0,003204051 | 0,043805201 | The Moose: suction sampler vs FISH | Carbohydrate transport and metabolism |
| gene_1941211 | 66,20734813 | 5,038119523 | 0,8818189 | 5,713326762 | 1,11E-08 | 9,60E-06 | The Moose: suction sampler vs FISH | Carbohydrate transport and metabolism |
| gene_1253354 | 47,51155927 | 5,04473848 | 1,006586754 | 5,011727462 | 5,39E-07 | 0,000145124 | The Moose: suction sampler vs FISH | Carbohydrate transport and metabolism |
| gene_1540701 | 10,66848326 | 5,050732647 | 1,615959156 | 3,125532368 | 0,001774835 | 0,030852902 | The Moose: suction sampler vs FISH | Carbohydrate transport and metabolism |
| gene_1364527 | 9,359173014 | 5,095758084 | 1,536628135 | 3,316194706 | 0,000912522 | 0,020601212 | The Moose: suction sampler vs FISH | Carbohydrate transport and metabolism |
| gene_1120412 | 17,8810557 | 5,107345364 | 1,343285327 | 3,802129942 | 0,000143457 | 0,006311059 | The Moose: suction sampler vs FISH | Carbohydrate transport and metabolism |
| gene_1314866 | 6,82578831 | 5,115510885 | 1,653458188 | 3,093825367 | 0,001975938 | 0,032945002 | The Moose: suction sampler vs FISH | Carbohydrate transport and metabolism |
| gene_633198 | 28,74124262 | 5,174985417 | 1,20838693 | 4,282556597 | 1,85E-05 | 0,001574195 | The Moose: suction sampler vs FISH | Carbohydrate transport and metabolism |
| gene_533857 | 4,459921737 | 5,217989274 | 1,775966375 | 2,938112652 | 0,00330217 | 0,044566386 | The Moose: suction sampler vs FISH | Carbohydrate transport and metabolism |
| gene_1323083 | 6,906487382 | 5,223264942 | 1,606511898 | 3,251307972 | 0,001148753 | 0,023726153 | The Moose: suction sampler vs FISH | Carbohydrate transport and metabolism |
| gene_247151 | 7,888447336 | 5,26494254 | 1,708368634 | 3,081853901 | 0,002057158 | 0,033770217 | The Moose: suction sampler vs FISH | Carbohydrate transport and metabolism |
| gene_1385387 | 10,10862789 | 5,269814721 | 1,551177326 | 3,397299993 | 0,000680543 | 0,016998304 | The Moose: suction sampler vs FISH | Carbohydrate transport and metabolism |
| gene_1590507 | 12,23244783 | 5,366194485 | 1,379220334 | 3,890744904 | 9,99E-05 | 0,004993997 | The Moose: suction sampler vs FISH | Carbohydrate transport and metabolism |
| gene_1068279 | 31,57819297 | 5,373830764 | 1,122004844 | 4,789489807 | 1,67E-06 | 0,000310112 | The Moose: suction sampler vs FISH | Carbohydrate transport and metabolism |
| gene_1016166 | 9,788164599 | 5,393545187 | 1,446704039 | 3,72816073 | 0,000192882 | 0,007640908 | The Moose: suction sampler vs FISH | Carbohydrate transport and metabolism |
| gene_1691258 | 7,245161897 | 5,546082268 | 1,666109502 | 3,328762162 | 0,000872329 | 0,020084068 | The Moose: suction sampler vs FISH | Carbohydrate transport and metabolism |
| gene_397595 | 10,45400155 | 5,573194267 | 1,573608431 | 3,541665232 | 0,00039761 | 0,012172759 | The Moose: suction sampler vs FISH | Carbohydrate transport and metabolism |
| gene_1689774 | 33,17852959 | 5,591546583 | 1,084291785 | 5,156865211 | 2,51E-07 | 8,66E-05 | The Moose: suction sampler vs FISH | Carbohydrate transport and metabolism |
| gene_1197 | 6,768105235 | 5,639076857 | 1,763664696 | 3,197363348 | 0,001386901 | 0,026542102 | The Moose: suction sampler vs FISH | Carbohydrate transport and metabolism |
| gene_973409 | 5,726117101 | 5,69940623 | 1,804111367 | 3,159121069 | 0,001582457 | 0,028811002 | The Moose: suction sampler vs FISH | Carbohydrate transport and metabolism |
| gene_633202 | 63,21836858 | 5,808847519 | 0,984987651 | 5,897381062 | 3,69E-09 | 4,24E-06 | The Moose: suction sampler vs FISH | Carbohydrate transport and metabolism |
| gene_1250000 | 5,136429395 | 6,087432256 | 1,78679765 | 3,406895154 | 0,000657064 | 0,016646443 | The Moose: suction sampler vs FISH | Carbohydrate transport and metabolism |
| gene_516132 | 5,891455572 | 6,13938014 | 1,95155741 | 3,14588754 | 0,001655837 | 0,029656462 | The Moose: suction sampler vs FISH | Carbohydrate transport and metabolism |
| gene_1366624 | 6,213085218 | 6,156598962 | 2,04562103 | 3,009647864 | 0,002615507 | 0,038835543 | The Moose: suction sampler vs FISH | Carbohydrate transport and metabolism |
| gene_345782 | 4,620751597 | 6,309396999 | 2,076548514 | 3,038405776 | 0,002378335 | 0,036573867 | The Moose: suction sampler vs FISH | Carbohydrate transport and metabolism |
| gene_914839 | 1955,908765 | 6,395829617 | 1,650626799 | 3,874788427 | 0,000106717 | 0,005204421 | The Moose: suction sampler vs FISH | Carbohydrate transport and metabolism |
| gene_533104 | 3,02636259 | 6,511223193 | 2,165505907 | 3,006790778 | 0,002640214 | 0,03898959 | The Moose: suction sampler vs FISH | Carbohydrate transport and metabolism |
| gene_964300 | 5,620836076 | 6,556388614 | 1,804354871 | 3,633646972 | 0,000279443 | 0,009722265 | The Moose: suction sampler vs FISH | Carbohydrate transport and metabolism |
| gene_1860374 | 4,461898726 | 6,633706767 | 2,086085744 | 3,179977998 | 0,001472862 | 0,027593959 | The Moose: suction sampler vs FISH | Carbohydrate transport and metabolism |
| gene_1667280 | 8,649289607 | 6,73074368 | 1,725296982 | 3,901208748 | 9,57E-05 | 0,004864833 | The Moose: suction sampler vs FISH | Carbohydrate transport and metabolism |
| gene_1285096 | 5,999886132 | 7,074377732 | 1,926572481 | 3,672001859 | 0,000240658 | 0,008818332 | The Moose: suction sampler vs FISH | Carbohydrate transport and metabolism |
| gene_980887 | 10,30554106 | 7,157187775 | 1,696879051 | 4,217853812 | 2,47E-05 | 0,001941519 | The Moose: suction sampler vs FISH | Carbohydrate transport and metabolism |
| gene_1098076 | 4,200557406 | 7,164565218 | 2,333609142 | 3,070165045 | 0,002139405 | 0,034518587 | The Moose: suction sampler vs FISH | Carbohydrate transport and metabolism |
| gene_985614 | 6,234911933 | 7,177074907 | 1,960899257 | 3,660093644 | 0,000252123 | 0,009085058 | The Moose: suction sampler vs FISH | Carbohydrate transport and metabolism |
| gene_985056 | 14,89900365 | 7,474231638 | 1,590206496 | 4,700164198 | 2,60E-06 | 0,000417772 | The Moose: suction sampler vs FISH | Carbohydrate transport and metabolism |
| gene_1381475 | 5,930286518 | 7,519617044 | 1,861425614 | 4,039708591 | 5,35E-05 | 0,003294723 | The Moose: suction sampler vs FISH | Carbohydrate transport and metabolism |
| gene_914840 | 184,9973135 | 7,61145348 | 1,724198021 | 4,414489165 | 1,01E-05 | 0,001071481 | The Moose: suction sampler vs FISH | Carbohydrate transport and metabolism |
| gene_1833135 | 9,239747857 | 8,499865657 | 2,903281943 | 2,927674895 | 0,003415069 | 0,045397242 | The Moose: suction sampler vs FISH | Carbohydrate transport and metabolism |
| gene_1064859 | 371,1362622 | 8,528948966 | 1,07337433 | 7,945922242 | 1,93E-15 | 7,08E-11 | The Moose: suction sampler vs FISH | Carbohydrate transport and metabolism |
| gene_1080284 | 69,13217855 | 8,921821373 | 2,395150114 | 3,724952903 | 0,000195352 | 0,007701345 | The Moose: suction sampler vs FISH | Carbohydrate transport and metabolism |
| gene_1185620 | 175,1595009 | 9,026248773 | 1,23396787 | 7,314816695 | 2,58E-13 | 3,15E-09 | The Moose: suction sampler vs FISH | Carbohydrate transport and metabolism |
| gene_1080287 | 214,9084965 | 9,442552066 | 2,320946047 | 4,068406536 | 4,73E-05 | 0,003058801 | The Moose: suction sampler vs FISH | Carbohydrate transport and metabolism |
| gene_1102150 | 32,51033463 | 10,02811257 | 1,734124267 | 5,782810818 | 7,35E-09 | 7,29E-06 | The Moose: suction sampler vs FISH | Carbohydrate transport and metabolism |
| gene_868878 | 37,69703284 | 10,55576939 | 2,733358594 | 3,86183116 | 0,00011254 | 0,005402304 | The Moose: suction sampler vs FISH | Carbohydrate transport and metabolism |
| gene_1399620 | 80,15888923 | 10,61922638 | 2,486030918 | 4,271558453 | 1,94E-05 | 0,00162006 | The Moose: suction sampler vs FISH | Carbohydrate transport and metabolism |
| gene_921158 | 91,66545718 | 10,88584083 | 2,168306917 | 5,020433565 | 5,16E-07 | 0,00014331 | The Moose: suction sampler vs FISH | Carbohydrate transport and metabolism |
| gene_1080286 | 61,49130455 | 11,24146257 | 2,809231292 | 4,001615177 | 6,29E-05 | 0,003658384 | The Moose: suction sampler vs FISH | Carbohydrate transport and metabolism |
| gene_535259 | 66,98614106 | 11,36943752 | 3,346661149 | 3,397247889 | 0,000680673 | 0,016998304 | The Moose: suction sampler vs FISH | Carbohydrate transport and metabolism |
| gene_921159 | 224,4599097 | 11,54663721 | 2,624727388 | 4,3991758 | 1,09E-05 | 0,001124706 | The Moose: suction sampler vs FISH | Carbohydrate transport and metabolism |
| gene_79924 | 74,54607299 | 11,55773645 | 3,543472084 | 3,26169818 | 0,00110747 | 0,023246106 | The Moose: suction sampler vs FISH | Carbohydrate transport and metabolism |
| gene_1635359 | 85,45080563 | 11,73492993 | 3,578811599 | 3,279001871 | 0,001041749 | 0,022345587 | The Moose: suction sampler vs FISH | Carbohydrate transport and metabolism |
| gene_307405 | 155,2747599 | 12,57135537 | 3,310092762 | 3,797886125 | 0,000145935 | 0,006362888 | The Moose: suction sampler vs FISH | Carbohydrate transport and metabolism |
| gene_1351360 | 257,8844853 | 13,37596591 | 4,247162 | 3,149389148 | 0,001636122 | 0,029448747 | The Moose: suction sampler vs FISH | Carbohydrate transport and metabolism |
| gene_474670 | 11,84348453 | -3,895044428 | 1,203711055 | -3,235863301 | 0,001212754 | 0,024471718 | The Moose: suction sampler vs FISH | Cell cycle control, cell division, chromosome partitioning |
| gene_829482 | 23,16477097 | -2,864993527 | 0,969572357 | -2,9549043 | 0,003127661 | 0,043219384 | The Moose: suction sampler vs FISH | Cell cycle control, cell division, chromosome partitioning |
| gene_614020 | 51,56987665 | -2,8176749 | 0,718507742 | -3,921565118 | 8,80E-05 | 0,004582648 | The Moose: suction sampler vs FISH | Cell cycle control, cell division, chromosome partitioning |
| gene_280108 | 30,26621584 | -2,530969701 | 0,820093676 | -3,086195852 | 0,002027352 | 0,033448147 | The Moose: suction sampler vs FISH | Cell cycle control, cell division, chromosome partitioning |
| gene_1313271 | 177,9735987 | 2,117469858 | 0,692853397 | 3,056158585 | 0,002241926 | 0,035387554 | The Moose: suction sampler vs FISH | Cell cycle control, cell division, chromosome partitioning |
| gene_1104177 | 158,6807669 | 2,186270796 | 0,734144026 | 2,977986225 | 0,00290149 | 0,041278765 | The Moose: suction sampler vs FISH | Cell cycle control, cell division, chromosome partitioning |
| gene_1135102 | 85,29433315 | 2,543524339 | 0,706535757 | 3,599993791 | 0,000318225 | 0,010580484 | The Moose: suction sampler vs FISH | Cell cycle control, cell division, chromosome partitioning |
| gene_163808 | 29,80741136 | 2,565395351 | 0,863442839 | 2,971123549 | 0,002967124 | 0,041835364 | The Moose: suction sampler vs FISH | Cell cycle control, cell division, chromosome partitioning |
| gene_236728 | 60,76530026 | 2,659731798 | 0,72671563 | 3,659934765 | 0,000252279 | 0,009085058 | The Moose: suction sampler vs FISH | Cell cycle control, cell division, chromosome partitioning |
| gene_395043 | 14,50193932 | 2,776686836 | 0,855825544 | 3,244454264 | 0,001176759 | 0,024038351 | The Moose: suction sampler vs FISH | Cell cycle control, cell division, chromosome partitioning |
| gene_1630162 | 20,57376571 | 2,866860345 | 0,904884985 | 3,168204127 | 0,001533838 | 0,028297594 | The Moose: suction sampler vs FISH | Cell cycle control, cell division, chromosome partitioning |
| gene_146848 | 48,69484379 | 2,924039144 | 0,744615196 | 3,926913069 | 8,60E-05 | 0,004520338 | The Moose: suction sampler vs FISH | Cell cycle control, cell division, chromosome partitioning |

| | | | | | | | | |
|---|---|---|---|---|---|---|---|---|
| gene_1918024 | 13,11309227 | 3,057699742 | 1,013700798 | 3,016373024 | 0,002558183 | 0,038346672 | The Moose: suction sampler vs FISH | Cell cycle control, cell division, chromosome partitioning |
| gene_1909191 | 19,75680291 | 3,184768851 | 0,861181736 | 3,698137939 | 0,000217187 | 0,008226554 | The Moose: suction sampler vs FISH | Cell cycle control, cell division, chromosome partitioning |
| gene_233213 | 259,7988558 | 3,211472964 | 0,88356321 | 3,634683888 | 0,000278322 | 0,009701631 | The Moose: suction sampler vs FISH | Cell cycle control, cell division, chromosome partitioning |
| gene_1098509 | 13,65066678 | 3,314163226 | 1,016931188 | 3,258984743 | 0,001118117 | 0,023337123 | The Moose: suction sampler vs FISH | Cell cycle control, cell division, chromosome partitioning |
| gene_146847 | 16,13286565 | 3,339423161 | 1,026302067 | 3,253840432 | 0,001138562 | 0,023598077 | The Moose: suction sampler vs FISH | Cell cycle control, cell division, chromosome partitioning |
| gene_188312 | 10,90650368 | 3,449049463 | 1,144673754 | 3,013128809 | 0,002585691 | 0,038583118 | The Moose: suction sampler vs FISH | Cell cycle control, cell division, chromosome partitioning |
| gene_1375560 | 35,50436791 | 3,463069782 | 0,962272834 | 3,598843967 | 0,000319635 | 0,010599955 | The Moose: suction sampler vs FISH | Cell cycle control, cell division, chromosome partitioning |
| gene_581677 | 370,3177316 | 3,527632812 | 0,718351112 | 4,910736199 | 9,07E-07 | 0,000208904 | The Moose: suction sampler vs FISH | Cell cycle control, cell division, chromosome partitioning |
| gene_1366275 | 246,1588514 | 3,588133665 | 0,798040253 | 4,496181303 | 6,92E-06 | 0,00083664 | The Moose: suction sampler vs FISH | Cell cycle control, cell division, chromosome partitioning |
| gene_1316132 | 14,22678596 | 3,844952975 | 1,21705521 | 3,159226421 | 0,001581885 | 0,02880773 | The Moose: suction sampler vs FISH | Cell cycle control, cell division, chromosome partitioning |
| gene_1496850 | 11,83108353 | 3,938138521 | 1,164589421 | 3,381568173 | 0,000720733 | 0,017685616 | The Moose: suction sampler vs FISH | Cell cycle control, cell division, chromosome partitioning |
| gene_459558 | 10,15075299 | 3,945409303 | 1,188559838 | 3,319487313 | 0,000901829 | 0,02049231 | The Moose: suction sampler vs FISH | Cell cycle control, cell division, chromosome partitioning |
| gene_1239425 | 15,23041214 | 4,032985607 | 1,170000027 | 3,446996166 | 0,000566857 | 0,015137809 | The Moose: suction sampler vs FISH | Cell cycle control, cell division, chromosome partitioning |
| gene_1202953 | 13,27829391 | 4,089422265 | 1,179925056 | 3,465832211 | 0,000528593 | 0,014464425 | The Moose: suction sampler vs FISH | Cell cycle control, cell division, chromosome partitioning |
| gene_547389 | 34,00538275 | 4,10864515 | 0,991328431 | 4,144585206 | 3,40E-05 | 0,002418722 | The Moose: suction sampler vs FISH | Cell cycle control, cell division, chromosome partitioning |
| gene_783457 | 10,36817375 | 4,156775423 | 1,44230797 | 2,882030406 | 0,003951216 | 0,049454169 | The Moose: suction sampler vs FISH | Cell cycle control, cell division, chromosome partitioning |
| gene_1918026 | 9,356932881 | 4,327892681 | 1,381324943 | 3,133145972 | 0,001729434 | 0,030426115 | The Moose: suction sampler vs FISH | Cell cycle control, cell division, chromosome partitioning |
| gene_1492259 | 7,348247202 | 4,411581694 | 1,481758127 | 2,977261682 | 0,002908357 | 0,041340185 | The Moose: suction sampler vs FISH | Cell cycle control, cell division, chromosome partitioning |
| gene_1484046 | 13,23511891 | 4,435725325 | 1,291482195 | 3,434600447 | 0,000593428 | 0,015532547 | The Moose: suction sampler vs FISH | Cell cycle control, cell division, chromosome partitioning |
| gene_1918027 | 10,56730337 | 4,525526109 | 1,208409529 | 3,74502683 | 0,000180374 | 0,007282905 | The Moose: suction sampler vs FISH | Cell cycle control, cell division, chromosome partitioning |
| gene_524681 | 8,452825252 | 4,671339694 | 1,481809345 | 3,15245663 | 0,001619029 | 0,029230473 | The Moose: suction sampler vs FISH | Cell cycle control, cell division, chromosome partitioning |
| gene_334631 | 10,01040371 | 4,854258458 | 1,46378926 | 3,316227677 | 0,000912415 | 0,020601212 | The Moose: suction sampler vs FISH | Cell cycle control, cell division, chromosome partitioning |
| gene_1918025 | 9,184836297 | 4,906104313 | 1,484812092 | 3,304192053 | 0,000952506 | 0,021103066 | The Moose: suction sampler vs FISH | Cell cycle control, cell division, chromosome partitioning |
| gene_1249294 | 4,843615753 | 4,964072716 | 1,611925801 | 3,079591327 | 0,002072848 | 0,033938676 | The Moose: suction sampler vs FISH | Cell cycle control, cell division, chromosome partitioning |
| gene_1490380 | 7,955749116 | 5,432721621 | 1,495070906 | 3,633755162 | 0,000279326 | 0,009722265 | The Moose: suction sampler vs FISH | Cell cycle control, cell division, chromosome partitioning |
| gene_79876 | 8,761879905 | 5,435550259 | 1,573272615 | 3,454932227 | 0,000550431 | 0,014817027 | The Moose: suction sampler vs FISH | Cell cycle control, cell division, chromosome partitioning |
| gene_24802 | 8,377060018 | 5,558926587 | 1,772394679 | 3,136393183 | 0,001710397 | 0,030233485 | The Moose: suction sampler vs FISH | Cell cycle control, cell division, chromosome partitioning |
| gene_1375302 | 6,297548785 | 6,262718797 | 1,681627822 | 3,724200274 | 0,000195935 | 0,00771607 | The Moose: suction sampler vs FISH | Cell cycle control, cell division, chromosome partitioning |
| gene_560604 | 3,066931508 | 6,378307845 | 2,145292245 | 2,973165013 | 0,002947459 | 0,041690887 | The Moose: suction sampler vs FISH | Cell cycle control, cell division, chromosome partitioning |
| gene_1821164 | 4,293885174 | 6,414305185 | 1,769351718 | 3,625229014 | 0,000288705 | 0,009931604 | The Moose: suction sampler vs FISH | Cell cycle control, cell division, chromosome partitioning |
| gene_1626603 | 8,442089517 | 6,94736181 | 1,693418352 | 4,102566741 | 4,09E-05 | 0,002745566 | The Moose: suction sampler vs FISH | Cell cycle control, cell division, chromosome partitioning |
| gene_640594 | 8,598381982 | 7,421919307 | 1,71782954 | 4,32052141 | 1,56E-05 | 0,001407946 | The Moose: suction sampler vs FISH | Cell cycle control, cell division, chromosome partitioning |
| gene_1473765 | 6,218347136 | 7,514609914 | 2,073300735 | 3,624466912 | 0,000289558 | 0,009951611 | The Moose: suction sampler vs FISH | Cell cycle control, cell division, chromosome partitioning |
| gene_1700837 | 11,42152664 | -5,908322453 | 1,514043062 | -3,902347695 | 9,53E-05 | 0,004858804 | The Moose: suction sampler vs FISH | Cell motility |
| gene_90803 | 11,00492244 | -4,066747891 | 1,297470801 | -3,134365636 | 0,001722261 | 0,0303701 | The Moose: suction sampler vs FISH | Cell motility |
| gene_808560 | 16,75251861 | -3,989774103 | 1,18214808 | -3,375020582 | 0,000738102 | 0,017991995 | The Moose: suction sampler vs FISH | Cell motility |
| gene_56377 | 38,21986714 | -3,861940965 | 0,862575665 | -4,477219936 | 7,56E-06 | 0,000886907 | The Moose: suction sampler vs FISH | Cell motility |
| gene_90801 | 10,27144133 | -3,766202893 | 1,277164843 | -2,948877677 | 0,003189302 | 0,04368487 | The Moose: suction sampler vs FISH | Cell motility |
| gene_1545061 | 13,18551776 | -3,737560683 | 1,223751464 | -3,054182809 | 0,002256745 | 0,035496378 | The Moose: suction sampler vs FISH | Cell motility |
| gene_375993 | 14,08396755 | -3,442995604 | 1,179378225 | -2,919331161 | 0,003507834 | 0,046082171 | The Moose: suction sampler vs FISH | Cell motility |
| gene_446823 | 17,52047611 | -3,001837572 | 1,012124177 | -2,965878734 | 0,003018195 | 0,04234409 | The Moose: suction sampler vs FISH | Cell motility |
| gene_1406714 | 149,8156031 | -2,826213118 | 0,641119383 | -4,40824781 | 1,04E-05 | 0,001091829 | The Moose: suction sampler vs FISH | Cell motility |
| gene_559148 | 49,67528502 | -2,500101341 | 0,780868917 | -3,201691458 | 0,001366232 | 0,026302214 | The Moose: suction sampler vs FISH | Cell motility |
| gene_44361 | 1777,378489 | -2,379405495 | 0,59170012 | -4,021303047 | 5,79E-05 | 0,003478546 | The Moose: suction sampler vs FISH | Cell motility |
| gene_1052323 | 773,2638838 | -2,203842031 | 0,599899498 | -3,673685403 | 0,000239077 | 0,008795979 | The Moose: suction sampler vs FISH | Cell motility |
| gene_1005536 | 81,70896769 | -2,035689383 | 0,684689302 | -2,973157866 | 0,002947528 | 0,041690887 | The Moose: suction sampler vs FISH | Cell motility |
| gene_1059294 | 57,85502757 | 2,102201547 | 0,720700255 | 2,916887474 | 0,003535433 | 0,046293434 | The Moose: suction sampler vs FISH | Cell motility |
| gene_1059293 | 32,85495727 | 2,358706516 | 0,755751764 | 3,121006962 | 0,001802338 | 0,03113919 | The Moose: suction sampler vs FISH | Cell motility |
| gene_611025 | 8,714452029 | 2,985268256 | 1,00619809 | 2,96687927 | 0,003008391 | 0,042263055 | The Moose: suction sampler vs FISH | Cell motility |
| gene_1909607 | 16,37187515 | 3,538324474 | 1,180717415 | 2,996758096 | 0,002728671 | 0,039731815 | The Moose: suction sampler vs FISH | Cell motility |
| gene_754484 | 123,4369807 | 3,991184653 | 0,870294684 | 4,586015203 | 4,52E-06 | 0,000616755 | The Moose: suction sampler vs FISH | Cell motility |
| gene_241622 | 32,60450676 | 4,104352178 | 1,329517844 | 3,087098227 | 0,002021208 | 0,033374015 | The Moose: suction sampler vs FISH | Cell motility |
| gene_1017750 | 6,65964489 | 4,371330795 | 1,486892313 | 2,939910818 | 0,003283067 | 0,044389705 | The Moose: suction sampler vs FISH | Cell motility |
| gene_99838 | 29,21100187 | 4,847873169 | 1,387256754 | 3,494575285 | 0,000474817 | 0,013490488 | The Moose: suction sampler vs FISH | Cell motility |
| gene_258814 | 1109,766503 | 5,775690768 | 1,847380834 | 3,12642129 | 0,001769479 | 0,030814148 | The Moose: suction sampler vs FISH | Cell motility |
| gene_846253 | 53,90453686 | 5,803534804 | 1,334379077 | 4,349239959 | 1,37E-05 | 0,00130134 | The Moose: suction sampler vs FISH | Cell motility |
| gene_259044 | 54,12515514 | 6,191505531 | 1,907135505 | 3,246494816 | 0,001168356 | 0,023920816 | The Moose: suction sampler vs FISH | Cell motility |
| gene_1059449 | 4,190027483 | 6,362650383 | 1,922953248 | 3,30879099 | 0,000936998 | 0,020910905 | The Moose: suction sampler vs FISH | Cell motility |
| gene_598600 | 5,130585233 | 6,377330807 | 2,049257553 | 3,112020155 | 0,001858118 | 0,031751858 | The Moose: suction sampler vs FISH | Cell motility |
| gene_302142 | 8,571770844 | -5,981019717 | 1,719074012 | -3,47921013 | 0,000502894 | 0,014018422 | The Moose: suction sampler vs FISH | Cell wall/membrane/envelope biogenesis |
| gene_302141 | 8,914757502 | -5,931179403 | 1,730363513 | -3,427707159 | 0,000608702 | 0,015808379 | The Moose: suction sampler vs FISH | Cell wall/membrane/envelope biogenesis |
| gene_1419614 | 5,390037956 | -5,4216338 | 1,671789392 | -3,243012443 | 0,00118273 | 0,024122643 | The Moose: suction sampler vs FISH | Cell wall/membrane/envelope biogenesis |
| gene_907755 | 9,597580968 | -5,250061751 | 1,566715495 | -3,350998805 | 0,000805206 | 0,018985037 | The Moose: suction sampler vs FISH | Cell wall/membrane/envelope biogenesis |
| gene_1492175 | 22,27784243 | -5,101838459 | 1,259098421 | -4,05197749 | 5,08E-05 | 0,003196245 | The Moose: suction sampler vs FISH | Cell wall/membrane/envelope biogenesis |
| gene_716040 | 21,96501058 | -5,020818287 | 1,462573116 | -3,432866523 | 0,000597236 | 0,0155979 | The Moose: suction sampler vs FISH | Cell wall/membrane/envelope biogenesis |

| | | | | | | | | |
|---|---|---|---|---|---|---|---|---|
| gene_328420 | 26,56930894 | -4,716722942 | 1,085379648 | -4,345689501 | 1,39E-05 | 0,001314293 | The Moose: suction sampler vs FISH | Cell wall/membrane/envelope biogenesis |
| gene_1433902 | 19,85050208 | -4,513143851 | 1,25885514 | -3,585117704 | 0,000336926 | 0,010927321 | The Moose: suction sampler vs FISH | Cell wall/membrane/envelope biogenesis |
| gene_334867 | 19,23883761 | -4,349695154 | 1,316367319 | -3,304317185 | 0,00095208 | 0,021103066 | The Moose: suction sampler vs FISH | Cell wall/membrane/envelope biogenesis |
| gene_685765 | 38,51584144 | -4,244795117 | 1,060597806 | -4,002266544 | 6,27E-05 | 0,003654111 | The Moose: suction sampler vs FISH | Cell wall/membrane/envelope biogenesis |
| gene_778552 | 14,19886897 | -4,1875442 | 1,34871049 | -3,104850322 | 0,001903753 | 0,032232343 | The Moose: suction sampler vs FISH | Cell wall/membrane/envelope biogenesis |
| gene_328426 | 12,35128142 | -4,048717824 | 1,379407814 | -2,935113012 | 0,003334262 | 0,044841602 | The Moose: suction sampler vs FISH | Cell wall/membrane/envelope biogenesis |
| gene_328421 | 22,33775979 | -3,946656556 | 1,053097588 | -3,747664607 | 0,000178489 | 0,007226628 | The Moose: suction sampler vs FISH | Cell wall/membrane/envelope biogenesis |
| gene_421140 | 13,63737409 | -3,719443555 | 1,292480664 | -2,877755666 | 0,004005152 | 0,049832018 | The Moose: suction sampler vs FISH | Cell wall/membrane/envelope biogenesis |
| gene_335930 | 13,23249948 | -3,662127481 | 1,15652162 | -3,166501531 | 0,001542845 | 0,028370623 | The Moose: suction sampler vs FISH | Cell wall/membrane/envelope biogenesis |
| gene_326084 | 17,00874305 | -3,448285632 | 1,082892524 | -3,184328598 | 0,001450902 | 0,02733047 | The Moose: suction sampler vs FISH | Cell wall/membrane/envelope biogenesis |
| gene_988784 | 15,55104946 | -3,418896765 | 1,023509369 | -3,340366847 | 0,000836678 | 0,01953889 | The Moose: suction sampler vs FISH | Cell wall/membrane/envelope biogenesis |
| gene_1252386 | 82,23859399 | -3,380331459 | 0,797349682 | -4,239459217 | 2,24E-05 | 0,001808352 | The Moose: suction sampler vs FISH | Cell wall/membrane/envelope biogenesis |
| gene_767265 | 16,1812259 | -3,305577303 | 1,089181166 | -3,034919632 | 0,002405998 | 0,03686804 | The Moose: suction sampler vs FISH | Cell wall/membrane/envelope biogenesis |
| gene_1734821 | 1066,742716 | -3,164991766 | 0,676957467 | -4,675318496 | 2,93E-06 | 0,000446293 | The Moose: suction sampler vs FISH | Cell wall/membrane/envelope biogenesis |
| gene_176367 | 488,970295 | -3,148709305 | 0,647607819 | -4,862061901 | 1,16E-06 | 0,000245986 | The Moose: suction sampler vs FISH | Cell wall/membrane/envelope biogenesis |
| gene_321485 | 526,154058 | -3,13856503 | 0,774295966 | -4,053443603 | 5,05E-05 | 0,003181724 | The Moose: suction sampler vs FISH | Cell wall/membrane/envelope biogenesis |
| gene_780629 | 621,3773275 | -3,110427197 | 0,773087514 | -4,023383045 | 5,74E-05 | 0,003456439 | The Moose: suction sampler vs FISH | Cell wall/membrane/envelope biogenesis |
| gene_271556 | 12,56594937 | -3,098954982 | 1,068436093 | -2,900458907 | 0,003726167 | 0,047894352 | The Moose: suction sampler vs FISH | Cell wall/membrane/envelope biogenesis |
| gene_328419 | 36,34933818 | -2,944867089 | 0,79351442 | -3,711170223 | 0,000206303 | 0,007983112 | The Moose: suction sampler vs FISH | Cell wall/membrane/envelope biogenesis |
| gene_209425 | 46,90622327 | -2,924727983 | 0,882031434 | -3,315899946 | 0,000913485 | 0,020610638 | The Moose: suction sampler vs FISH | Cell wall/membrane/envelope biogenesis |
| gene_175131 | 46,70612983 | -2,793407893 | 0,782607018 | -3,569362184 | 0,000357851 | 0,011357993 | The Moose: suction sampler vs FISH | Cell wall/membrane/envelope biogenesis |
| gene_175130 | 53,51162544 | -2,742473184 | 0,774960837 | -3,538853903 | 0,000401868 | 0,012225229 | The Moose: suction sampler vs FISH | Cell wall/membrane/envelope biogenesis |
| gene_25663 | 91,07957517 | -2,724817847 | 0,764683833 | -3,563326081 | 0,000366185 | 0,011508122 | The Moose: suction sampler vs FISH | Cell wall/membrane/envelope biogenesis |
| gene_846827 | 6381,370474 | -2,705238411 | 0,676886956 | -3,996588187 | 6,43E-05 | 0,00371408 | The Moose: suction sampler vs FISH | Cell wall/membrane/envelope biogenesis |
| gene_340517 | 413,7784281 | -2,692466004 | 0,538380686 | -5,001044937 | 5,70E-07 | 0,000150103 | The Moose: suction sampler vs FISH | Cell wall/membrane/envelope biogenesis |
| gene_539930 | 84,34961234 | -2,637030725 | 0,83072003 | -3,174391648 | 0,001501509 | 0,027946872 | The Moose: suction sampler vs FISH | Cell wall/membrane/envelope biogenesis |
| gene_258231 | 16,32231863 | -2,602112704 | 0,891175167 | -2,91986671 | 0,003501811 | 0,046077151 | The Moose: suction sampler vs FISH | Cell wall/membrane/envelope biogenesis |
| gene_14487 | 189,3544757 | -2,572709054 | 0,716454758 | -3,590888366 | 0,000329553 | 0,010805359 | The Moose: suction sampler vs FISH | Cell wall/membrane/envelope biogenesis |
| gene_242820 | 1287,137484 | -2,555544924 | 0,715028681 | -3,57404534 | 0,000351508 | 0,01120022 | The Moose: suction sampler vs FISH | Cell wall/membrane/envelope biogenesis |
| gene_916476 | 32,937623 | -2,553208705 | 0,799796104 | -3,19232451 | 0,001411327 | 0,026839693 | The Moose: suction sampler vs FISH | Cell wall/membrane/envelope biogenesis |
| gene_578119 | 573,5672543 | -2,526886535 | 0,744788215 | -3,392758484 | 0,000691926 | 0,01717422 | The Moose: suction sampler vs FISH | Cell wall/membrane/envelope biogenesis |
| gene_1323883 | 547,2597651 | -2,472490554 | 0,594122408 | -4,161584416 | 3,16E-05 | 0,002284654 | The Moose: suction sampler vs FISH | Cell wall/membrane/envelope biogenesis |
| gene_1162037 | 517,2153987 | -2,432001904 | 0,575201425 | -4,228087411 | 2,36E-05 | 0,001869328 | The Moose: suction sampler vs FISH | Cell wall/membrane/envelope biogenesis |
| gene_68936 | 34,08011704 | -2,376352867 | 0,725205673 | -3,276798508 | 0,001049913 | 0,02247474 | The Moose: suction sampler vs FISH | Cell wall/membrane/envelope biogenesis |
| gene_277063 | 257,2347379 | -2,288773163 | 0,785551515 | -2,913587611 | 0,003573016 | 0,046635886 | The Moose: suction sampler vs FISH | Cell wall/membrane/envelope biogenesis |
| gene_400785 | 4081,044967 | -2,184342479 | 0,627494253 | -3,481055749 | 0,000499442 | 0,013957982 | The Moose: suction sampler vs FISH | Cell wall/membrane/envelope biogenesis |
| gene_1764721 | 160,1002215 | -2,175527015 | 0,684997756 | -3,175962249 | 0,001493404 | 0,027838336 | The Moose: suction sampler vs FISH | Cell wall/membrane/envelope biogenesis |
| gene_88559 | 54,73246959 | -2,150008476 | 0,662269174 | -3,24642692 | 0,001168634 | 0,023920816 | The Moose: suction sampler vs FISH | Cell wall/membrane/envelope biogenesis |
| gene_129396 | 492,3123271 | -2,115395098 | 0,709878502 | -2,979939655 | 0,002883052 | 0,041123662 | The Moose: suction sampler vs FISH | Cell wall/membrane/envelope biogenesis |
| gene_57796 | 146,4416207 | -2,075173119 | 0,586409865 | -3,538775935 | 0,000401987 | 0,012225229 | The Moose: suction sampler vs FISH | Cell wall/membrane/envelope biogenesis |
| gene_1115744 | 451,3722791 | 2,025447878 | 0,675982393 | 2,996302711 | 0,00273275 | 0,039760916 | The Moose: suction sampler vs FISH | Cell wall/membrane/envelope biogenesis |
| gene_1768169 | 72,18348598 | 2,057736071 | 0,638807116 | 3,221216573 | 0,001276476 | 0,025283391 | The Moose: suction sampler vs FISH | Cell wall/membrane/envelope biogenesis |
| gene_1717265 | 43,45260066 | 2,064875732 | 0,689120852 | 2,996391309 | 0,002731956 | 0,039760916 | The Moose: suction sampler vs FISH | Cell wall/membrane/envelope biogenesis |
| gene_1164375 | 25,69642482 | 2,221849734 | 0,772325158 | 2,876832006 | 0,004016894 | 0,049918912 | The Moose: suction sampler vs FISH | Cell wall/membrane/envelope biogenesis |
| gene_1032237 | 50,36265369 | 2,31146229 | 0,746514907 | 3,096337754 | 0,001959271 | 0,032755968 | The Moose: suction sampler vs FISH | Cell wall/membrane/envelope biogenesis |
| gene_1191561 | 100,6667196 | 2,426211074 | 0,831849597 | 2,916646331 | 0,003538167 | 0,046312722 | The Moose: suction sampler vs FISH | Cell wall/membrane/envelope biogenesis |
| gene_378697 | 66,06992867 | 2,475310256 | 0,71635163 | 3,455440249 | 0,000549395 | 0,014796594 | The Moose: suction sampler vs FISH | Cell wall/membrane/envelope biogenesis |
| gene_1084 | 44,55004692 | 2,608665519 | 0,881622375 | 2,958937513 | 0,003087017 | 0,042924263 | The Moose: suction sampler vs FISH | Cell wall/membrane/envelope biogenesis |
| gene_167654 | 42,68747645 | 2,709331279 | 0,720237026 | 3,761721738 | 0,000168748 | 0,007017937 | The Moose: suction sampler vs FISH | Cell wall/membrane/envelope biogenesis |
| gene_813170 | 60,63431696 | 2,722454663 | 0,843404352 | 3,227935279 | 0,001246872 | 0,024884919 | The Moose: suction sampler vs FISH | Cell wall/membrane/envelope biogenesis |
| gene_494836 | 121,400283 | 2,766103942 | 0,755983617 | 3,65894694 | 0,000253254 | 0,009097339 | The Moose: suction sampler vs FISH | Cell wall/membrane/envelope biogenesis |
| gene_1121042 | 22,00166803 | 2,783297167 | 0,814965644 | 3,415232517 | 0,000637276 | 0,016313951 | The Moose: suction sampler vs FISH | Cell wall/membrane/envelope biogenesis |
| gene_462373 | 22,8759292 | 2,783748229 | 0,888852797 | 3,131843921 | 0,001737122 | 0,030507633 | The Moose: suction sampler vs FISH | Cell wall/membrane/envelope biogenesis |
| gene_1715978 | 39,44777919 | 2,823598616 | 0,90356496 | 3,124953647 | 0,001778331 | 0,030878692 | The Moose: suction sampler vs FISH | Cell wall/membrane/envelope biogenesis |
| gene_496919 | 170,5756515 | 2,827634842 | 0,939496379 | 3,009734689 | 0,00261476 | 0,038835543 | The Moose: suction sampler vs FISH | Cell wall/membrane/envelope biogenesis |
| gene_431562 | 16,88910572 | 3,010989007 | 1,044786146 | 2,881918964 | 0,003952614 | 0,049463233 | The Moose: suction sampler vs FISH | Cell wall/membrane/envelope biogenesis |
| gene_1749916 | 38,1839411 | 3,014603941 | 0,787996253 | 3,825657712 | 0,000130423 | 0,005936145 | The Moose: suction sampler vs FISH | Cell wall/membrane/envelope biogenesis |
| gene_316376 | 54,63598773 | 3,102109112 | 0,954147047 | 3,25118557 | 0,001149248 | 0,023728034 | The Moose: suction sampler vs FISH | Cell wall/membrane/envelope biogenesis |
| gene_462374 | 23,67640768 | 3,227511249 | 0,848082513 | 3,805657114 | 0,000141428 | 0,006253579 | The Moose: suction sampler vs FISH | Cell wall/membrane/envelope biogenesis |
| gene_819647 | 15,77416152 | 3,251478512 | 1,105683134 | 2,94069649 | 0,003274752 | 0,044326241 | The Moose: suction sampler vs FISH | Cell wall/membrane/envelope biogenesis |
| gene_21563 | 10,36680655 | 3,310502778 | 1,123683872 | 2,946115773 | 0,003217919 | 0,04392122 | The Moose: suction sampler vs FISH | Cell wall/membrane/envelope biogenesis |
| gene_980779 | 3213,633087 | 3,337043842 | 0,681382355 | 4,89746149 | 9,71E-07 | 0,000219392 | The Moose: suction sampler vs FISH | Cell wall/membrane/envelope biogenesis |
| gene_1640157 | 78,55209556 | 3,383503756 | 1,088415345 | 3,10865128 | 0,001879434 | 0,032001019 | The Moose: suction sampler vs FISH | Cell wall/membrane/envelope biogenesis |
| gene_819646 | 11,83815371 | 3,461231878 | 1,065291511 | 3,249093644 | 0,001157734 | 0,023837887 | The Moose: suction sampler vs FISH | Cell wall/membrane/envelope biogenesis |
| gene_621410 | 13,28448507 | 3,6266308 | 1,240881729 | 2,922624062 | 0,003470953 | 0,045841423 | The Moose: suction sampler vs FISH | Cell wall/membrane/envelope biogenesis |

| | | | | | | | | |
|---|---|---|---|---|---|---|---|---|
| gene_1009515 | 71,59706133 | 3,698612194 | 1,092727028 | 3,384754015 | 0,000712421 | 0,017536627 | The Moose: suction sampler vs FISH | Cell wall/membrane/envelope biogenesis |
| gene_1229541 | 14,04606653 | 3,723109508 | 1,238173006 | 3,006938036 | 0,002638935 | 0,038986234 | The Moose: suction sampler vs FISH | Cell wall/membrane/envelope biogenesis |
| gene_120946 | 31,02579588 | 3,811109397 | 1,239952855 | 3,07359218 | 0,002114983 | 0,034313001 | The Moose: suction sampler vs FISH | Cell wall/membrane/envelope biogenesis |
| gene_466430 | 47,11635688 | 3,819121148 | 0,828444345 | 4,609991212 | 4,03E-06 | 0,000563339 | The Moose: suction sampler vs FISH | Cell wall/membrane/envelope biogenesis |
| gene_119838 | 24,77485934 | 3,820421051 | 1,256321538 | 3,040958016 | 0,002358267 | 0,03638717 | The Moose: suction sampler vs FISH | Cell wall/membrane/envelope biogenesis |
| gene_21564 | 45,35423528 | 3,897383783 | 0,883098607 | 4,413305321 | 1,02E-05 | 0,001072742 | The Moose: suction sampler vs FISH | Cell wall/membrane/envelope biogenesis |
| gene_539140 | 31,78005246 | 4,067263403 | 1,280355438 | 3,176667419 | 0,001489778 | 0,027784847 | The Moose: suction sampler vs FISH | Cell wall/membrane/envelope biogenesis |
| gene_190537 | 6,410790122 | 4,133788318 | 1,400709787 | 2,951209705 | 0,003165319 | 0,043486135 | The Moose: suction sampler vs FISH | Cell wall/membrane/envelope biogenesis |
| gene_401272 | 12,1458314 | 4,450903903 | 1,155686447 | 3,851307519 | 0,000117489 | 0,005552747 | The Moose: suction sampler vs FISH | Cell wall/membrane/envelope biogenesis |
| gene_908384 | 11,0059151 | 4,533492153 | 1,549783695 | 2,925241869 | 0,003441886 | 0,045647426 | The Moose: suction sampler vs FISH | Cell wall/membrane/envelope biogenesis |
| gene_1349398 | 7,199060293 | 4,791218888 | 1,48099852 | 3,235127398 | 0,001215885 | 0,024511793 | The Moose: suction sampler vs FISH | Cell wall/membrane/envelope biogenesis |
| gene_951450 | 10,31373111 | 4,855534677 | 1,269219579 | 3,825606503 | 0,000130451 | 0,005936145 | The Moose: suction sampler vs FISH | Cell wall/membrane/envelope biogenesis |
| gene_925253 | 9,852144099 | 5,440120189 | 1,666017694 | 3,265343585 | 0,001093314 | 0,023047768 | The Moose: suction sampler vs FISH | Cell wall/membrane/envelope biogenesis |
| gene_1653169 | 37,55809926 | 5,655945237 | 1,331917173 | 4,24646919 | 2,17E-05 | 0,001762401 | The Moose: suction sampler vs FISH | Cell wall/membrane/envelope biogenesis |
| gene_158878 | 7,160641822 | 5,724097142 | 1,492740918 | 3,834621984 | 0,000125758 | 0,005802755 | The Moose: suction sampler vs FISH | Cell wall/membrane/envelope biogenesis |
| gene_997484 | 5,480302796 | 6,172320296 | 1,763667725 | 3,499707008 | 0,00046577 | 0,013354236 | The Moose: suction sampler vs FISH | Cell wall/membrane/envelope biogenesis |
| gene_46509 | 9,266703184 | 6,22469958 | 1,843970412 | 3,375704696 | 0,000736269 | 0,017959241 | The Moose: suction sampler vs FISH | Cell wall/membrane/envelope biogenesis |
| gene_1927994 | 3,670865561 | 6,288572461 | 2,093665953 | 3,003617865 | 0,002667902 | 0,039220187 | The Moose: suction sampler vs FISH | Cell wall/membrane/envelope biogenesis |
| gene_176179 | 6,087898625 | 6,338269745 | 1,912398327 | 3,314304167 | 0,000918715 | 0,020672497 | The Moose: suction sampler vs FISH | Cell wall/membrane/envelope biogenesis |
| gene_1653170 | 29,29042826 | 6,458638231 | 1,399030255 | 4,616510763 | 3,90E-06 | 0,000553313 | The Moose: suction sampler vs FISH | Cell wall/membrane/envelope biogenesis |
| gene_1170550 | 11,53961319 | 6,600714499 | 1,507720857 | 4,377942023 | 1,20E-05 | 0,001193907 | The Moose: suction sampler vs FISH | Cell wall/membrane/envelope biogenesis |
| gene_473084 | 4,774793092 | 6,619939552 | 1,996614472 | 3,315582275 | 0,000914524 | 0,020610638 | The Moose: suction sampler vs FISH | Cell wall/membrane/envelope biogenesis |
| gene_462709 | 3,779304954 | 6,807622079 | 2,01198849 | 3,383529336 | 0,000715606 | 0,017589577 | The Moose: suction sampler vs FISH | Cell wall/membrane/envelope biogenesis |
| gene_1267235 | 4,830846791 | 7,597472867 | 2,000551015 | 3,797690141 | 0,000146051 | 0,006362888 | The Moose: suction sampler vs FISH | Cell wall/membrane/envelope biogenesis |
| gene_333992 | 44,32870264 | 2,410183045 | 0,816341353 | 2,95242062 | 0,003152931 | 0,043383106 | The Moose: suction sampler vs FISH | Chromatin structure and dynamics |
| gene_772698 | 43,71452944 | 2,506439976 | 0,800755761 | 3,130092967 | 0,00174751 | 0,030638783 | The Moose: suction sampler vs FISH | Chromatin structure and dynamics |
| gene_710068 | 31,53068835 | 2,64756323 | 0,808692935 | 3,273879511 | 0,001060818 | 0,022622371 | The Moose: suction sampler vs FISH | Chromatin structure and dynamics |
| gene_219490 | 65,614433 | 2,778211024 | 0,704944899 | 3,941032877 | 8,11E-05 | 0,004338943 | The Moose: suction sampler vs FISH | Chromatin structure and dynamics |
| gene_387930 | 16,60877335 | 2,949663661 | 1,001882149 | 2,944122384 | 0,003238719 | 0,044090776 | The Moose: suction sampler vs FISH | Chromatin structure and dynamics |
| gene_1095838 | 16,7476328 | 3,001536512 | 0,975725718 | 3,076209283 | 0,002096506 | 0,034164169 | The Moose: suction sampler vs FISH | Chromatin structure and dynamics |
| gene_109309 | 655,7325137 | 3,046215473 | 0,82106036 | 3,710099307 | 0,000207178 | 0,008008502 | The Moose: suction sampler vs FISH | Chromatin structure and dynamics |
| gene_948554 | 111,1317031 | 3,144389312 | 0,707045742 | 4,447221908 | 8,70E-06 | 0,000967043 | The Moose: suction sampler vs FISH | Chromatin structure and dynamics |
| gene_1078412 | 18,09604127 | 3,34533604 | 0,99144343 | 3,374207685 | 0,000740285 | 0,018021288 | The Moose: suction sampler vs FISH | Chromatin structure and dynamics |
| gene_109307 | 833,1653302 | 3,368677224 | 0,847318738 | 3,975690696 | 7,02E-05 | 0,003947106 | The Moose: suction sampler vs FISH | Chromatin structure and dynamics |
| gene_1247884 | 252,0483663 | 3,421369455 | 0,866255106 | 3,949609568 | 7,83E-05 | 0,004249399 | The Moose: suction sampler vs FISH | Chromatin structure and dynamics |
| gene_219893 | 13,7901856 | 3,470245122 | 1,161114323 | 2,98871959 | 0,002801491 | 0,040431423 | The Moose: suction sampler vs FISH | Chromatin structure and dynamics |
| gene_1209342 | 33,38363595 | 3,475333137 | 0,915066767 | 3,797901161 | 0,000145926 | 0,006362888 | The Moose: suction sampler vs FISH | Chromatin structure and dynamics |
| gene_950360 | 98,99321669 | 3,543765456 | 0,730676725 | 4,849977203 | 1,23E-06 | 0,000254879 | The Moose: suction sampler vs FISH | Chromatin structure and dynamics |
| gene_607358 | 123,0599172 | 3,546535288 | 0,766765057 | 4,625322 | 3,74E-06 | 0,000535471 | The Moose: suction sampler vs FISH | Chromatin structure and dynamics |
| gene_970785 | 8,973942072 | 3,788994401 | 1,236614676 | 3,064005688 | 0,002183948 | 0,034922715 | The Moose: suction sampler vs FISH | Chromatin structure and dynamics |
| gene_1114986 | 109,9000876 | 4,005995616 | 0,842825671 | 4,753053631 | 2,00E-06 | 0,00035038 | The Moose: suction sampler vs FISH | Chromatin structure and dynamics |
| gene_109308 | 425,2368654 | 4,185604986 | 0,84141869 | 4,974461626 | 6,54E-07 | 0,000164687 | The Moose: suction sampler vs FISH | Chromatin structure and dynamics |
| gene_971914 | 3,817116343 | 5,384282654 | 1,851339682 | 2,908316992 | 0,003633798 | 0,047112049 | The Moose: suction sampler vs FISH | Chromatin structure and dynamics |
| gene_278909 | 7,138711437 | 5,718204652 | 1,746066146 | 3,274907234 | 0,001056967 | 0,022573109 | The Moose: suction sampler vs FISH | Chromatin structure and dynamics |
| gene_715268 | 6,049986439 | 6,42282997 | 1,658756574 | 3,872075065 | 0,000107913 | 0,005238367 | The Moose: suction sampler vs FISH | Chromatin structure and dynamics |
| gene_1826140 | 3,218779583 | 6,444670193 | 2,19583734 | 2,934948812 | 0,003336027 | 0,044841602 | The Moose: suction sampler vs FISH | Chromatin structure and dynamics |
| gene_585256 | 5,771523797 | -5,230882086 | 1,751423155 | -2,986646642 | 0,002820555 | 0,040618761 | The Moose: suction sampler vs FISH | Coenzyme transport and metabolism |
| gene_88186 | 18,0887782 | -4,504535589 | 1,270714492 | -3,544884093 | 0,000392786 | 0,012065308 | The Moose: suction sampler vs FISH | Coenzyme transport and metabolism |
| gene_1259245 | 10,09300847 | -4,472302478 | 1,38545921 | -3,228028978 | 0,001246463 | 0,024884919 | The Moose: suction sampler vs FISH | Coenzyme transport and metabolism |
| gene_968567 | 8,819199357 | -3,927672353 | 1,360469716 | -2,886997268 | 0,003889376 | 0,049005872 | The Moose: suction sampler vs FISH | Coenzyme transport and metabolism |
| gene_585257 | 11,94838717 | -3,84583286 | 1,297039478 | -2,965085431 | 0,003025989 | 0,042388661 | The Moose: suction sampler vs FISH | Coenzyme transport and metabolism |
| gene_1064480 | 15,16459305 | -3,076360012 | 1,050766204 | -2,92773026 | 0,003414461 | 0,045397242 | The Moose: suction sampler vs FISH | Coenzyme transport and metabolism |
| gene_832598 | 31,8498044 | -2,644428417 | 0,819861196 | -3,225458685 | 0,00125771 | 0,025026416 | The Moose: suction sampler vs FISH | Coenzyme transport and metabolism |
| gene_980492 | 16,48449567 | -2,568013302 | 0,887241596 | -2,89437884 | 0,003799095 | 0,048366186 | The Moose: suction sampler vs FISH | Coenzyme transport and metabolism |
| gene_849788 | 184,2230733 | -2,379119791 | 0,559156107 | -4,254840034 | 2,09E-05 | 0,001720199 | The Moose: suction sampler vs FISH | Coenzyme transport and metabolism |
| gene_1526395 | 97,85171413 | -2,335622899 | 0,664276904 | -3,516038095 | 0,000438038 | 0,012889307 | The Moose: suction sampler vs FISH | Coenzyme transport and metabolism |
| gene_283998 | 1424,267723 | -2,322420885 | 0,555619632 | -4,179875494 | 2,92E-05 | 0,002176993 | The Moose: suction sampler vs FISH | Coenzyme transport and metabolism |
| gene_1663150 | 26,54273223 | -2,25429787 | 0,778555677 | -2,895487035 | 0,003785707 | 0,048279428 | The Moose: suction sampler vs FISH | Coenzyme transport and metabolism |
| gene_632976 | 606,0637171 | -2,042513666 | 0,558170297 | -3,659301965 | 0,000252903 | 0,00909469 | The Moose: suction sampler vs FISH | Coenzyme transport and metabolism |
| gene_358502 | 79,01794977 | 2,393982997 | 0,778413885 | 3,075462864 | 0,002101761 | 0,034196685 | The Moose: suction sampler vs FISH | Coenzyme transport and metabolism |
| gene_223782 | 139,95133 | 2,521164049 | 0,875384434 | 2,880064976 | 0,003975932 | 0,049611339 | The Moose: suction sampler vs FISH | Coenzyme transport and metabolism |
| gene_1099241 | 37,8723157 | 2,609594342 | 0,896588104 | 2,910583277 | 0,003607548 | 0,046911539 | The Moose: suction sampler vs FISH | Coenzyme transport and metabolism |
| gene_193032 | 23,1487991 | 2,682920495 | 0,904948149 | 2,964722892 | 0,003029557 | 0,042414374 | The Moose: suction sampler vs FISH | Coenzyme transport and metabolism |
| gene_619636 | 17,79682986 | 2,9247347 | 0,891274469 | 3,28151967 | 0,001032493 | 0,022172943 | The Moose: suction sampler vs FISH | Coenzyme transport and metabolism |
| gene_332503 | 155,9454896 | 2,975458934 | 0,773488532 | 3,846804199 | 0,000119669 | 0,0056304 | The Moose: suction sampler vs FISH | Coenzyme transport and metabolism |

| | | | | | | | | |
|---|---|---|---|---|---|---|---|---|
| gene_137488 | 12,50667476 | 3,066654999 | 1,027318674 | 2,985105864 | 0,002834802 | 0,040773117 | The Moose: suction sampler vs FISH | Coenzyme transport and metabolism |
| gene_1906025 | 33,00781098 | 3,133903909 | 0,951532743 | 3,29353239 | 0,000989369 | 0,021574892 | The Moose: suction sampler vs FISH | Coenzyme transport and metabolism |
| gene_1090676 | 24,74887057 | 3,181621574 | 0,949927858 | 3,349329686 | 0,000810073 | 0,019075294 | The Moose: suction sampler vs FISH | Coenzyme transport and metabolism |
| gene_763837 | 24,16856562 | 3,208077055 | 0,907112327 | 3,536581921 | 0,000405341 | 0,012276393 | The Moose: suction sampler vs FISH | Coenzyme transport and metabolism |
| gene_1128841 | 20,25721823 | 3,228951142 | 0,920112015 | 3,509302227 | 0,000449284 | 0,01306321 | The Moose: suction sampler vs FISH | Coenzyme transport and metabolism |
| gene_1732518 | 6,554081348 | 3,452219853 | 1,168799026 | 2,953647099 | 0,003140429 | 0,043306684 | The Moose: suction sampler vs FISH | Coenzyme transport and metabolism |
| gene_1455469 | 13,23525218 | 3,64963192 | 1,099894304 | 3,318166032 | 0,000906106 | 0,020546144 | The Moose: suction sampler vs FISH | Coenzyme transport and metabolism |
| gene_469485 | 17,91472441 | 3,998842827 | 1,154923979 | 3,462429475 | 0,000535322 | 0,014561755 | The Moose: suction sampler vs FISH | Coenzyme transport and metabolism |
| gene_555094 | 14,85401594 | 4,016198575 | 0,939431635 | 4,275136608 | 1,91E-05 | 0,001601538 | The Moose: suction sampler vs FISH | Coenzyme transport and metabolism |
| gene_1509949 | 10,1753344 | 4,11458288 | 1,169054557 | 3,519581577 | 0,000432228 | 0,012764372 | The Moose: suction sampler vs FISH | Coenzyme transport and metabolism |
| gene_821133 | 15,74532211 | 4,319122042 | 1,25683 | 3,436520487 | 0,000589238 | 0,015472495 | The Moose: suction sampler vs FISH | Coenzyme transport and metabolism |
| gene_432477 | 13,78106487 | 4,524480927 | 1,128910181 | 4,00783074 | 6,13E-05 | 0,003606207 | The Moose: suction sampler vs FISH | Coenzyme transport and metabolism |
| gene_696127 | 3,928502251 | 5,562640675 | 1,847657381 | 3,010645119 | 0,002606933 | 0,038790953 | The Moose: suction sampler vs FISH | Coenzyme transport and metabolism |
| gene_1242233 | 4,192865397 | 5,79513939 | 1,848485573 | 3,135074179 | 0,001718107 | 0,030313307 | The Moose: suction sampler vs FISH | Coenzyme transport and metabolism |
| gene_370182 | 4,27977357 | 5,812895369 | 1,817591712 | 3,198130434 | 0,001383217 | 0,026497228 | The Moose: suction sampler vs FISH | Coenzyme transport and metabolism |
| gene_560761 | 3,731754519 | 6,090730447 | 1,993236003 | 3,055699594 | 0,002245361 | 0,035391185 | The Moose: suction sampler vs FISH | Coenzyme transport and metabolism |
| gene_148263 | 3,733180928 | 6,783717542 | 2,262168687 | 2,998767325 | 0,002710742 | 0,039541305 | The Moose: suction sampler vs FISH | Coenzyme transport and metabolism |
| gene_1437266 | 3,577345816 | 6,800048794 | 2,154524488 | 3,156171505 | 0,001598548 | 0,029003306 | The Moose: suction sampler vs FISH | Coenzyme transport and metabolism |
| gene_1608717 | 10,27038899 | 7,683260915 | 1,790333671 | 4,291524557 | 1,77E-05 | 0,001540172 | The Moose: suction sampler vs FISH | Coenzyme transport and metabolism |
| gene_192121 | 382,5890607 | 2,090280448 | 0,68906309 | 3,033510975 | 0,002417259 | 0,036971179 | The Moose: suction sampler vs FISH | Cytoskeleton |
| gene_1454674 | 753,3588197 | 2,172171382 | 0,682569318 | 3,182345478 | 0,001460874 | 0,027440901 | The Moose: suction sampler vs FISH | Cytoskeleton |
| gene_1173409 | 188,8870603 | 2,343039045 | 0,678302762 | 3,45426729 | 0,00055179 | 0,014833906 | The Moose: suction sampler vs FISH | Cytoskeleton |
| gene_87543 | 45,96066644 | 2,34373422 | 0,805207279 | 2,910721599 | 0,003605952 | 0,046907389 | The Moose: suction sampler vs FISH | Cytoskeleton |
| gene_1021493 | 70,41288337 | 2,364192609 | 0,716732117 | 3,298572161 | 0,000971779 | 0,021343394 | The Moose: suction sampler vs FISH | Cytoskeleton |
| gene_1088054 | 54,46753376 | 2,487348853 | 0,772932133 | 3,218068893 | 0,001290568 | 0,025452679 | The Moose: suction sampler vs FISH | Cytoskeleton |
| gene_1361207 | 60,45745472 | 2,552146589 | 0,80746191 | 3,160702143 | 0,001573894 | 0,028704895 | The Moose: suction sampler vs FISH | Cytoskeleton |
| gene_939093 | 30,95324006 | 2,577226139 | 0,887017691 | 2,905495758 | 0,003666718 | 0,047420693 | The Moose: suction sampler vs FISH | Cytoskeleton |
| gene_410365 | 243,1594056 | 2,593077396 | 0,682044265 | 3,80191951 | 0,000143579 | 0,006311059 | The Moose: suction sampler vs FISH | Cytoskeleton |
| gene_1685007 | 40,78423905 | 2,617021417 | 0,785701418 | 3,33080908 | 0,00086594 | 0,01999338 | The Moose: suction sampler vs FISH | Cytoskeleton |
| gene_1594166 | 21,98500564 | 2,650746595 | 0,883499362 | 3,000281278 | 0,002697304 | 0,03943505 | The Moose: suction sampler vs FISH | Cytoskeleton |
| gene_1021492 | 90,55752428 | 2,653308774 | 0,725331904 | 3,658061585 | 0,00025413 | 0,009118015 | The Moose: suction sampler vs FISH | Cytoskeleton |
| gene_354989 | 41,36639618 | 2,683686108 | 0,894725397 | 2,999452252 | 0,002704655 | 0,039498414 | The Moose: suction sampler vs FISH | Cytoskeleton |
| gene_1886453 | 32,87432645 | 2,785829232 | 0,912809428 | 3,051928634 | 0,002273762 | 0,035649607 | The Moose: suction sampler vs FISH | Cytoskeleton |
| gene_1385254 | 35,19883181 | 2,788164536 | 0,906890956 | 3,07442093 | 0,002109116 | 0,034252904 | The Moose: suction sampler vs FISH | Cytoskeleton |
| gene_187440 | 54,35480988 | 2,80002709 | 0,814142331 | 3,439235358 | 0,00058336 | 0,015389678 | The Moose: suction sampler vs FISH | Cytoskeleton |
| gene_1701251 | 368,0187784 | 2,816314231 | 0,959091393 | 2,936439896 | 0,003320031 | 0,044733024 | The Moose: suction sampler vs FISH | Cytoskeleton |
| gene_599256 | 21,34297314 | 2,823016103 | 0,847307763 | 3,331748188 | 0,000863023 | 0,019957409 | The Moose: suction sampler vs FISH | Cytoskeleton |
| gene_187438 | 696,1486807 | 2,828132622 | 0,724291003 | 3,904691086 | 9,43E-05 | 0,004827305 | The Moose: suction sampler vs FISH | Cytoskeleton |
| gene_186441 | 18,14101724 | 2,851957302 | 0,990915083 | 2,878104645 | 0,004000724 | 0,049803793 | The Moose: suction sampler vs FISH | Cytoskeleton |
| gene_187439 | 18,74851425 | 2,862924201 | 0,941121315 | 3,042035235 | 0,002349844 | 0,036325886 | The Moose: suction sampler vs FISH | Cytoskeleton |
| gene_793391 | 25,46337938 | 2,930860207 | 1,004385366 | 2,918063431 | 0,003522127 | 0,046218083 | The Moose: suction sampler vs FISH | Cytoskeleton |
| gene_1885841 | 60,71227086 | 2,93723243 | 0,73316635 | 4,006229187 | 6,17E-05 | 0,003613435 | The Moose: suction sampler vs FISH | Cytoskeleton |
| gene_154457 | 95,05903889 | 2,944149928 | 0,807573175 | 3,645675734 | 0,00026669 | 0,009438241 | The Moose: suction sampler vs FISH | Cytoskeleton |
| gene_181579 | 47,27512825 | 2,959291569 | 0,749456168 | 3,948585249 | 7,86E-05 | 0,004254853 | The Moose: suction sampler vs FISH | Cytoskeleton |
| gene_1255140 | 30,15446733 | 2,988475805 | 0,941801179 | 3,173149355 | 0,001507949 | 0,028016083 | The Moose: suction sampler vs FISH | Cytoskeleton |
| gene_1845065 | 52,486107 | 2,991622896 | 0,83135642 | 3,598484143 | 0,000320077 | 0,010602098 | The Moose: suction sampler vs FISH | Cytoskeleton |
| gene_1412836 | 12,16219503 | 2,999562034 | 0,962575725 | 3,116182922 | 0,001832086 | 0,031519688 | The Moose: suction sampler vs FISH | Cytoskeleton |
| gene_1250731 | 94,50144128 | 3,014451539 | 0,942633664 | 3,197903548 | 0,001384306 | 0,026511169 | The Moose: suction sampler vs FISH | Cytoskeleton |
| gene_1221460 | 35,43111003 | 3,025805659 | 0,830535336 | 3,643199186 | 0,00026927 | 0,009494267 | The Moose: suction sampler vs FISH | Cytoskeleton |
| gene_991121 | 52,62344252 | 3,057055567 | 0,871683395 | 3,507071014 | 0,000453068 | 0,013131648 | The Moose: suction sampler vs FISH | Cytoskeleton |
| gene_1475003 | 23,41675511 | 3,140903682 | 1,067466136 | 2,942391871 | 0,003256875 | 0,044186664 | The Moose: suction sampler vs FISH | Cytoskeleton |
| gene_931655 | 15,69646629 | 3,151889046 | 0,980103324 | 3,215874254 | 0,001300478 | 0,025552062 | The Moose: suction sampler vs FISH | Cytoskeleton |
| gene_117377 | 30,48475577 | 3,193466654 | 0,815456338 | 3,916171233 | 9,00E-05 | 0,004669667 | The Moose: suction sampler vs FISH | Cytoskeleton |
| gene_1670895 | 87,15173703 | 3,210553251 | 0,793480023 | 4,046167715 | 5,21E-05 | 0,003226804 | The Moose: suction sampler vs FISH | Cytoskeleton |
| gene_675563 | 23,47024871 | 3,220769087 | 1,061470776 | 3,034251305 | 0,002411335 | 0,036919008 | The Moose: suction sampler vs FISH | Cytoskeleton |
| gene_1109808 | 68,65388543 | 3,241039011 | 0,748826662 | 4,328156535 | 1,50E-05 | 0,001380423 | The Moose: suction sampler vs FISH | Cytoskeleton |
| gene_212483 | 112,7102328 | 3,266069376 | 0,891668736 | 3,662873042 | 0,000249402 | 0,009018878 | The Moose: suction sampler vs FISH | Cytoskeleton |
| gene_1670894 | 24,83236079 | 3,281399916 | 1,023049665 | 3,207468835 | 0,001339086 | 0,0259702 | The Moose: suction sampler vs FISH | Cytoskeleton |
| gene_921311 | 18,93075576 | 3,309760016 | 1,008235574 | 3,282724891 | 0,001028089 | 0,022130137 | The Moose: suction sampler vs FISH | Cytoskeleton |
| gene_446326 | 40,79852509 | 3,315047957 | 1,056721145 | 3,137107622 | 0,001706235 | 0,0301817 | The Moose: suction sampler vs FISH | Cytoskeleton |
| gene_1147158 | 10,73709161 | 3,382342494 | 1,074018022 | 3,149241842 | 0,001636947 | 0,029448747 | The Moose: suction sampler vs FISH | Cytoskeleton |
| gene_1293671 | 28,74609836 | 3,385321207 | 1,058481054 | 3,198282286 | 0,001382489 | 0,026497228 | The Moose: suction sampler vs FISH | Cytoskeleton |
| gene_185256 | 14,065579 | 3,429922514 | 1,086117544 | 3,157966219 | 0,00158874 | 0,028868131 | The Moose: suction sampler vs FISH | Cytoskeleton |
| gene_550880 | 21,69399646 | 3,489306881 | 1,181882626 | 2,952329448 | 0,003153862 | 0,043383106 | The Moose: suction sampler vs FISH | Cytoskeleton |
| gene_433521 | 19,87124278 | 3,521785617 | 0,969032572 | 3,634331518 | 0,000278702 | 0,009705686 | The Moose: suction sampler vs FISH | Cytoskeleton |

| | | | | | | | | |
|---|---|---|---|---|---|---|---|---|
| gene_21385 | 33,23392101 | 3,529145614 | 0,946137445 | 3,730055958 | 0,000191437 | 0,007614874 | The Moose: suction sampler vs FISH | Cytoskeleton |
| gene_1809633 | 23,49270007 | 3,539205592 | 0,938256986 | 3,772106839 | 0,000161875 | 0,006828778 | The Moose: suction sampler vs FISH | Cytoskeleton |
| gene_614700 | 20,19650713 | 3,583154546 | 0,960849661 | 3,729152116 | 0,000192125 | 0,007625297 | The Moose: suction sampler vs FISH | Cytoskeleton |
| gene_1709949 | 12,98593198 | 3,601312513 | 1,171420284 | 3,074312919 | 0,00210988 | 0,034252904 | The Moose: suction sampler vs FISH | Cytoskeleton |
| gene_1903116 | 19,61589881 | 3,628744704 | 1,119032545 | 3,242751716 | 0,001183813 | 0,024131325 | The Moose: suction sampler vs FISH | Cytoskeleton |
| gene_117209 | 26,4164195 | 3,661540276 | 1,194377099 | 3,065648427 | 0,002171985 | 0,034799623 | The Moose: suction sampler vs FISH | Cytoskeleton |
| gene_567702 | 98,89674386 | 3,66278254 | 0,742407261 | 4,933656675 | 8,07E-07 | 0,000194338 | The Moose: suction sampler vs FISH | Cytoskeleton |
| gene_1393949 | 15,86147279 | 3,714456378 | 0,982355171 | 3,781174556 | 0,00015609 | 0,006645822 | The Moose: suction sampler vs FISH | Cytoskeleton |
| gene_1425732 | 41,34749399 | 3,752712214 | 1,067090729 | 3,516769579 | 0,000436833 | 0,012867033 | The Moose: suction sampler vs FISH | Cytoskeleton |
| gene_1809635 | 13,95859834 | 3,758943979 | 1,027508175 | 3,658310533 | 0,000253883 | 0,009113617 | The Moose: suction sampler vs FISH | Cytoskeleton |
| gene_741827 | 14,76512937 | 3,773427076 | 1,210983092 | 3,116003105 | 0,001833203 | 0,031531528 | The Moose: suction sampler vs FISH | Cytoskeleton |
| gene_1548254 | 9,61074313 | 3,779841803 | 1,17690156 | 3,211689008 | 0,001319571 | 0,025804086 | The Moose: suction sampler vs FISH | Cytoskeleton |
| gene_130395 | 8,721831336 | 3,82818924 | 1,225707063 | 3,123249718 | 0,001788659 | 0,0310258 | The Moose: suction sampler vs FISH | Cytoskeleton |
| gene_258042 | 1060,064375 | 3,830789431 | 0,816367399 | 4,692482128 | 2,70E-06 | 0,000427235 | The Moose: suction sampler vs FISH | Cytoskeleton |
| gene_699105 | 18,26308014 | 3,837094329 | 1,115260353 | 3,440536837 | 0,000580561 | 0,015337446 | The Moose: suction sampler vs FISH | Cytoskeleton |
| gene_1898003 | 13,85492426 | 3,851144428 | 1,207775005 | 3,188627361 | 0,0014295 | 0,027061083 | The Moose: suction sampler vs FISH | Cytoskeleton |
| gene_376532 | 13,79244814 | 3,890335734 | 1,087939657 | 3,575874552 | 0,000349059 | 0,011160923 | The Moose: suction sampler vs FISH | Cytoskeleton |
| gene_989357 | 227,6579509 | 3,911673162 | 0,816500771 | 4,790777056 | 1,66E-06 | 0,000309039 | The Moose: suction sampler vs FISH | Cytoskeleton |
| gene_939958 | 160,5423626 | 3,929645862 | 0,854615198 | 4,598146477 | 4,26E-06 | 0,00058848 | The Moose: suction sampler vs FISH | Cytoskeleton |
| gene_1944028 | 13,08205099 | 3,942484308 | 1,284069616 | 3,07030418 | 0,002138409 | 0,034518033 | The Moose: suction sampler vs FISH | Cytoskeleton |
| gene_569598 | 406,5920168 | 3,954313085 | 0,883042784 | 4,47805379 | 7,53E-06 | 0,000886832 | The Moose: suction sampler vs FISH | Cytoskeleton |
| gene_770532 | 18,29798137 | 3,985225094 | 1,091429353 | 3,651381632 | 0,000260833 | 0,009281445 | The Moose: suction sampler vs FISH | Cytoskeleton |
| gene_1586255 | 37,55562294 | 3,985447107 | 0,805924902 | 4,945184219 | 7,61E-07 | 0,000185618 | The Moose: suction sampler vs FISH | Cytoskeleton |
| gene_21383 | 25,07076383 | 4,013457046 | 1,01961353 | 3,936253226 | 8,28E-05 | 0,004398391 | The Moose: suction sampler vs FISH | Cytoskeleton |
| gene_1415377 | 59,4287112 | 4,01919299 | 0,925219326 | 4,34404349 | 1,40E-05 | 0,001322235 | The Moose: suction sampler vs FISH | Cytoskeleton |
| gene_939956 | 22,62710373 | 4,10394351 | 1,400308908 | 2,930741557 | 0,00338154 | 0,045114839 | The Moose: suction sampler vs FISH | Cytoskeleton |
| gene_109860 | 7,750583463 | 4,143955392 | 1,334194237 | 3,105961093 | 0,001896617 | 0,032162783 | The Moose: suction sampler vs FISH | Cytoskeleton |
| gene_627246 | 13,64851143 | 4,158770623 | 1,095400466 | 3,796575547 | 0,000146709 | 0,006383305 | The Moose: suction sampler vs FISH | Cytoskeleton |
| gene_1594165 | 11,66063322 | 4,190986521 | 1,098420131 | 3,815467689 | 0,000135925 | 0,006113186 | The Moose: suction sampler vs FISH | Cytoskeleton |
| gene_1666160 | 84,29350271 | 4,199956644 | 0,853622002 | 4,920159785 | 8,65E-07 | 0,000202263 | The Moose: suction sampler vs FISH | Cytoskeleton |
| gene_492851 | 17,96903669 | 4,23062334 | 1,045477241 | 4,046595348 | 5,20E-05 | 0,003223638 | The Moose: suction sampler vs FISH | Cytoskeleton |
| gene_1228236 | 26,49063774 | 4,239695039 | 1,334397675 | 3,177235032 | 0,001486865 | 0,027756766 | The Moose: suction sampler vs FISH | Cytoskeleton |
| gene_699102 | 16,61195587 | 4,26635313 | 1,277284386 | 3,340174808 | 0,000837257 | 0,019546192 | The Moose: suction sampler vs FISH | Cytoskeleton |
| gene_590040 | 25,4305427 | 4,266713981 | 0,996469997 | 4,28182885 | 1,85E-05 | 0,001574414 | The Moose: suction sampler vs FISH | Cytoskeleton |
| gene_1421624 | 9,54494024 | 4,267209458 | 1,267700141 | 3,366103167 | 0,000762382 | 0,018358406 | The Moose: suction sampler vs FISH | Cytoskeleton |
| gene_1424874 | 362,423147 | 4,275975677 | 0,947411396 | 4,513325148 | 6,38E-06 | 0,000791755 | The Moose: suction sampler vs FISH | Cytoskeleton |
| gene_563124 | 44,27872289 | 4,294345202 | 0,980377411 | 4,380297987 | 1,19E-05 | 0,001189643 | The Moose: suction sampler vs FISH | Cytoskeleton |
| gene_1112907 | 11,3568882 | 4,317591546 | 1,380007686 | 3,1286721 | 0,001755981 | 0,030685572 | The Moose: suction sampler vs FISH | Cytoskeleton |
| gene_1372322 | 195,835689 | 4,321841448 | 0,863376464 | 5,005743878 | 5,56E-07 | 0,000148617 | The Moose: suction sampler vs FISH | Cytoskeleton |
| gene_516320 | 8,883697195 | 4,355030236 | 1,345813779 | 3,235982797 | 0,001212247 | 0,024471718 | The Moose: suction sampler vs FISH | Cytoskeleton |
| gene_1868181 | 111,9159452 | 4,355186644 | 0,930472003 | 4,680620833 | 2,86E-06 | 0,000439201 | The Moose: suction sampler vs FISH | Cytoskeleton |
| gene_819168 | 17,74125732 | 4,412821787 | 1,13424498 | 3,890536759 | 0,000100023 | 0,004993997 | The Moose: suction sampler vs FISH | Cytoskeleton |
| gene_939952 | 8,538382882 | 4,41451484 | 1,490518163 | 2,961731664 | 0,003059143 | 0,042657819 | The Moose: suction sampler vs FISH | Cytoskeleton |
| gene_776516 | 461,0597575 | 4,428724438 | 0,757067504 | 5,849840887 | 4,92E-09 | 5,34E-06 | The Moose: suction sampler vs FISH | Cytoskeleton |
| gene_252217 | 1647,416592 | 4,450891139 | 0,866171253 | 5,138580995 | 2,77E-07 | 9,20E-05 | The Moose: suction sampler vs FISH | Cytoskeleton |
| gene_1050618 | 685,7522391 | 4,451566119 | 0,835215122 | 5,329843774 | 9,83E-08 | 4,64E-05 | The Moose: suction sampler vs FISH | Cytoskeleton |
| gene_872377 | 16,65988467 | 4,465228632 | 1,160084729 | 3,849053883 | 0,000118575 | 0,005596874 | The Moose: suction sampler vs FISH | Cytoskeleton |
| gene_939199 | 442,8569912 | 4,509907894 | 0,830172814 | 5,432492871 | 5,56E-08 | 3,27E-05 | The Moose: suction sampler vs FISH | Cytoskeleton |
| gene_699103 | 305,34537 | 4,516841011 | 0,86445451 | 5,225076578 | 1,74E-07 | 6,52E-05 | The Moose: suction sampler vs FISH | Cytoskeleton |
| gene_445931 | 14,08208784 | 4,551674309 | 1,358114672 | 3,351465383 | 0,000803851 | 0,018965252 | The Moose: suction sampler vs FISH | Cytoskeleton |
| gene_1118117 | 7,975528948 | 4,558503503 | 1,445847537 | 3,152824476 | 0,00161699 | 0,029225655 | The Moose: suction sampler vs FISH | Cytoskeleton |
| gene_1684085 | 7,763441798 | 4,564309406 | 1,417259698 | 3,22051732 | 0,001279595 | 0,025297396 | The Moose: suction sampler vs FISH | Cytoskeleton |
| gene_1218851 | 16,83410132 | 4,570931178 | 1,15449119 | 3,959260337 | 7,52E-05 | 0,00414858 | The Moose: suction sampler vs FISH | Cytoskeleton |
| gene_209439 | 113,4673698 | 4,575431599 | 0,891908935 | 5,12993134 | 2,90E-07 | 9,38E-05 | The Moose: suction sampler vs FISH | Cytoskeleton |
| gene_150817 | 9,517704233 | 4,646200194 | 1,372905677 | 3,384209324 | 0,000713836 | 0,017563703 | The Moose: suction sampler vs FISH | Cytoskeleton |
| gene_1121942 | 41,41727148 | 4,653806783 | 0,873305222 | 5,328957926 | 9,88E-08 | 4,64E-05 | The Moose: suction sampler vs FISH | Cytoskeleton |
| gene_954779 | 7,344226019 | 4,74152735 | 1,503922046 | 3,152774681 | 0,001617266 | 0,029225655 | The Moose: suction sampler vs FISH | Cytoskeleton |
| gene_385435 | 7,01601554 | 4,751586793 | 1,39588125 | 3,404005027 | 0,000664055 | 0,016737807 | The Moose: suction sampler vs FISH | Cytoskeleton |
| gene_1856946 | 39,96643779 | 4,761517697 | 1,028649297 | 4,628902884 | 3,68E-06 | 0,000530432 | The Moose: suction sampler vs FISH | Cytoskeleton |
| gene_812408 | 16,77270918 | 4,772518281 | 1,143583822 | 4,173299927 | 3,00E-05 | 0,002211595 | The Moose: suction sampler vs FISH | Cytoskeleton |
| gene_1508354 | 6,49070202 | 4,798371961 | 1,630460093 | 2,942955784 | 0,003250948 | 0,044160858 | The Moose: suction sampler vs FISH | Cytoskeleton |
| gene_1418030 | 18,4617466 | 4,817764131 | 1,240064208 | 3,885092482 | 0,000102291 | 0,005069338 | The Moose: suction sampler vs FISH | Cytoskeleton |
| gene_1283376 | 10,72041581 | 4,820595808 | 1,230814792 | 3,916589108 | 8,98E-05 | 0,004668182 | The Moose: suction sampler vs FISH | Cytoskeleton |
| gene_1598869 | 9,221478602 | 4,891945438 | 1,328737338 | 3,681649712 | 0,00023173 | 0,008608692 | The Moose: suction sampler vs FISH | Cytoskeleton |
| gene_1551019 | 8,591313286 | 4,910574193 | 1,567451282 | 3,132840075 | 0,001731237 | 0,030426115 | The Moose: suction sampler vs FISH | Cytoskeleton |

| | | | | | | | | |
|---|---|---|---|---|---|---|---|---|
| gene_201139 | 42,15499512 | 4,982339357 | 1,014065343 | 4,91323305 | 8,96E-07 | 0,000207561 | The Moose: suction sampler vs FISH | Cytoskeleton |
| gene_515095 | 39,96979907 | 4,98695801 | 1,15714847 | 4,309695894 | 1,63E-05 | 0,001453599 | The Moose: suction sampler vs FISH | Cytoskeleton |
| gene_776518 | 107,484543 | 5,013278997 | 0,847859958 | 5,91286208 | 3,36E-09 | 4,05E-06 | The Moose: suction sampler vs FISH | Cytoskeleton |
| gene_785757 | 100,1210651 | 5,089245787 | 0,947956931 | 5,368646634 | 7,93E-08 | 4,10E-05 | The Moose: suction sampler vs FISH | Cytoskeleton |
| gene_1912633 | 36,84367519 | 5,115034195 | 1,297465441 | 3,94232789 | 8,07E-05 | 0,004319688 | The Moose: suction sampler vs FISH | Cytoskeleton |
| gene_699486 | 7,045496479 | 5,175398093 | 1,735364503 | 2,982311834 | 0,002860804 | 0,040967641 | The Moose: suction sampler vs FISH | Cytoskeleton |
| gene_1002137 | 406,1692898 | 5,216218916 | 0,922731986 | 5,653016252 | 1,58E-08 | 1,22E-05 | The Moose: suction sampler vs FISH | Cytoskeleton |
| gene_1634348 | 4,849420988 | 5,24333295 | 1,768907883 | 2,964163934 | 0,003035066 | 0,042459122 | The Moose: suction sampler vs FISH | Cytoskeleton |
| gene_776519 | 105,2250311 | 5,255973289 | 0,921765434 | 5,702072456 | 1,18E-08 | 9,88E-06 | The Moose: suction sampler vs FISH | Cytoskeleton |
| gene_822741 | 2024,08958 | 5,30103087 | 0,87784801 | 6,038665928 | 1,55E-09 | 2,39E-06 | The Moose: suction sampler vs FISH | Cytoskeleton |
| gene_1901881 | 6,963334805 | 5,338096095 | 1,497692403 | 3,56421391 | 0,000364948 | 0,011490192 | The Moose: suction sampler vs FISH | Cytoskeleton |
| gene_182223 | 13,04378242 | 5,344389612 | 1,283187368 | 4,164933155 | 3,11E-05 | 0,002267936 | The Moose: suction sampler vs FISH | Cytoskeleton |
| gene_939957 | 8,157756835 | 5,403113535 | 1,663110358 | 3,248800363 | 0,001158928 | 0,023840915 | The Moose: suction sampler vs FISH | Cytoskeleton |
| gene_1332552 | 6,940611476 | 5,462724324 | 1,661651095 | 3,287527895 | 0,001010712 | 0,021852142 | The Moose: suction sampler vs FISH | Cytoskeleton |
| gene_1669450 | 4,716374235 | 5,468100368 | 1,785635231 | 3,062271775 | 0,002196639 | 0,035034128 | The Moose: suction sampler vs FISH | Cytoskeleton |
| gene_939955 | 6,967115601 | 5,483333299 | 1,875862022 | 2,923100546 | 0,003465646 | 0,045802481 | The Moose: suction sampler vs FISH | Cytoskeleton |
| gene_464037 | 5,343093747 | 5,486983021 | 1,839387486 | 2,98304901 | 0,002853923 | 0,040907188 | The Moose: suction sampler vs FISH | Cytoskeleton |
| gene_1333986 | 11,10505021 | 5,492843798 | 1,329813878 | 4,130535775 | 3,62E-05 | 0,00251477 | The Moose: suction sampler vs FISH | Cytoskeleton |
| gene_1247569 | 4,035068918 | 5,504359858 | 1,892631374 | 2,908310585 | 0,003633872 | 0,047112049 | The Moose: suction sampler vs FISH | Cytoskeleton |
| gene_1002138 | 79,55591763 | 5,541720533 | 0,971792394 | 5,702576565 | 1,18E-08 | 9,88E-06 | The Moose: suction sampler vs FISH | Cytoskeleton |
| gene_1424873 | 75,5037892 | 5,663951466 | 1,092541833 | 5,184196426 | 2,17E-07 | 7,75E-05 | The Moose: suction sampler vs FISH | Cytoskeleton |
| gene_694180 | 5,005484105 | 5,723815573 | 1,903559032 | 3,006902059 | 0,002639248 | 0,038986234 | The Moose: suction sampler vs FISH | Cytoskeleton |
| gene_715564 | 4,063457539 | 5,732396665 | 1,903812633 | 3,011008838 | 0,002603813 | 0,038774742 | The Moose: suction sampler vs FISH | Cytoskeleton |
| gene_1115360 | 15,93999828 | 5,795062537 | 1,34386136 | 4,312247311 | 1,62E-05 | 0,001447674 | The Moose: suction sampler vs FISH | Cytoskeleton |
| gene_1700944 | 2484,766009 | 5,847123601 | 0,886987157 | 6,592117545 | 4,34E-11 | 1,87E-07 | The Moose: suction sampler vs FISH | Cytoskeleton |
| gene_1073583 | 24,21545153 | 5,927845644 | 1,166067797 | 5,08362006 | 3,70E-07 | 0,000113796 | The Moose: suction sampler vs FISH | Cytoskeleton |
| gene_1476609 | 517,5690651 | 5,942344229 | 1,558385161 | 3,813142206 | 0,000137211 | 0,006144799 | The Moose: suction sampler vs FISH | Cytoskeleton |
| gene_1776552 | 136,0643742 | 5,968747789 | 1,893291957 | 3,152576531 | 0,001618364 | 0,029225655 | The Moose: suction sampler vs FISH | Cytoskeleton |
| gene_711004 | 15,08375214 | 5,975304573 | 1,556919825 | 3,837901269 | 0,00012409 | 0,005771891 | The Moose: suction sampler vs FISH | Cytoskeleton |
| gene_1189857 | 148,411172 | 6,004625982 | 1,201492557 | 4,997638935 | 5,80E-07 | 0,000151608 | The Moose: suction sampler vs FISH | Cytoskeleton |
| gene_727088 | 6,172499431 | 6,127380788 | 1,682888631 | 3,640990066 | 0,000271592 | 0,009538157 | The Moose: suction sampler vs FISH | Cytoskeleton |
| gene_566948 | 3,347321027 | 6,154539566 | 2,017504921 | 3,050569791 | 0,002284076 | 0,035723316 | The Moose: suction sampler vs FISH | Cytoskeleton |
| gene_1709954 | 21,15246373 | 6,16898125 | 1,239611203 | 4,976545256 | 6,47E-07 | 0,000164687 | The Moose: suction sampler vs FISH | Cytoskeleton |
| gene_1551020 | 6,151763856 | 6,297510464 | 1,833848785 | 3,434040209 | 0,000594656 | 0,0155536 | The Moose: suction sampler vs FISH | Cytoskeleton |
| gene_1443208 | 31,38686628 | 6,318029259 | 1,965582286 | 3,214329568 | 0,001307495 | 0,025650268 | The Moose: suction sampler vs FISH | Cytoskeleton |
| gene_1721634 | 116,6530177 | 6,381465082 | 2,048239141 | 3,115585947 | 0,001835798 | 0,03156138 | The Moose: suction sampler vs FISH | Cytoskeleton |
| gene_810461 | 4,091478787 | 6,448527269 | 2,149246583 | 3,000366417 | 0,00269655 | 0,03943505 | The Moose: suction sampler vs FISH | Cytoskeleton |
| gene_842033 | 4,107829809 | 6,506287419 | 2,105101866 | 3,090723315 | 0,001996696 | 0,033147553 | The Moose: suction sampler vs FISH | Cytoskeleton |
| gene_1439694 | 158,7850653 | 6,56481549 | 1,745246454 | 3,761540654 | 0,00016887 | 0,007019046 | The Moose: suction sampler vs FISH | Cytoskeleton |
| gene_909773 | 28,9759903 | 6,574926288 | 1,428838687 | 4,601587532 | 4,19E-06 | 0,000584211 | The Moose: suction sampler vs FISH | Cytoskeleton |
| gene_1639139 | 77,99141641 | 6,629448121 | 1,741374229 | 3,80702092 | 0,000140651 | 0,00623422 | The Moose: suction sampler vs FISH | Cytoskeleton |
| gene_653010 | 75,43454994 | 6,633049352 | 2,279468757 | 2,909910185 | 0,003615326 | 0,046971103 | The Moose: suction sampler vs FISH | Cytoskeleton |
| gene_393943 | 4,756999356 | 6,646479532 | 1,842774346 | 3,606778848 | 0,000310022 | 0,010401801 | The Moose: suction sampler vs FISH | Cytoskeleton |
| gene_735561 | 5,573560913 | 6,666620773 | 1,876643272 | 3,552417698 | 0,000381708 | 0,011838925 | The Moose: suction sampler vs FISH | Cytoskeleton |
| gene_980371 | 11,60468253 | 6,73107683 | 1,827626056 | 3,682961735 | 0,00023054 | 0,008577504 | The Moose: suction sampler vs FISH | Cytoskeleton |
| gene_1700942 | 462,2163252 | 6,789562683 | 0,917544966 | 7,399705664 | 1,36E-13 | 2,51E-09 | The Moose: suction sampler vs FISH | Cytoskeleton |
| gene_1719700 | 3,922931885 | 6,789976449 | 1,974141095 | 3,439458539 | 0,000582879 | 0,01538252 | The Moose: suction sampler vs FISH | Cytoskeleton |
| gene_1776550 | 190,6532757 | 6,80864048 | 1,568940599 | 4,339641975 | 1,43E-05 | 0,001337845 | The Moose: suction sampler vs FISH | Cytoskeleton |
| gene_714371 | 99,03359941 | 6,822264687 | 1,716186826 | 3,975245926 | 7,03E-05 | 0,003950777 | The Moose: suction sampler vs FISH | Cytoskeleton |
| gene_1721635 | 10,28925579 | 6,883256128 | 1,89089903 | 3,640202897 | 0,000272423 | 0,009545863 | The Moose: suction sampler vs FISH | Cytoskeleton |
| gene_1269615 | 95,84376693 | 6,896516405 | 1,948443961 | 3,539499489 | 0,000400887 | 0,01221018 | The Moose: suction sampler vs FISH | Cytoskeleton |
| gene_1915893 | 155,3684206 | 6,936010152 | 1,975392671 | 3,511205775 | 0,000446079 | 0,013006062 | The Moose: suction sampler vs FISH | Cytoskeleton |
| gene_895575 | 10,04093911 | 6,959524764 | 1,802566311 | 3,860898055 | 0,000112971 | 0,005408838 | The Moose: suction sampler vs FISH | Cytoskeleton |
| gene_1295759 | 5,06370519 | 6,964935388 | 1,945909107 | 3,579270667 | 0,000344554 | 0,011099036 | The Moose: suction sampler vs FISH | Cytoskeleton |
| gene_1189856 | 91,96321369 | 7,009279694 | 2,14931294 | 3,261172239 | 0,001109526 | 0,023254434 | The Moose: suction sampler vs FISH | Cytoskeleton |
| gene_1772005 | 5,88600618 | 7,029316543 | 1,729586644 | 4,06415982 | 4,82E-05 | 0,003086725 | The Moose: suction sampler vs FISH | Cytoskeleton |
| gene_1863727 | 6,750833767 | 7,10740525 | 1,619576783 | 4,388433648 | 1,14E-05 | 0,001161387 | The Moose: suction sampler vs FISH | Cytoskeleton |
| gene_1395395 | 5,992559114 | 7,108693577 | 1,960522038 | 3,62591873 | 0,000287936 | 0,009914418 | The Moose: suction sampler vs FISH | Cytoskeleton |
| gene_54245 | 27,56970835 | 7,138408241 | 2,433654373 | 2,933205438 | 0,003354818 | 0,044952336 | The Moose: suction sampler vs FISH | Cytoskeleton |
| gene_895576 | 21,89014671 | 7,193113155 | 1,655244976 | 4,345648686 | 1,39E-05 | 0,001314293 | The Moose: suction sampler vs FISH | Cytoskeleton |
| gene_80280 | 46,41827551 | 7,312121002 | 2,104822446 | 3,473984713 | 0,00051279 | 0,014178245 | The Moose: suction sampler vs FISH | Cytoskeleton |
| gene_523058 | 33,30339091 | 7,372855806 | 2,129032109 | 3,463008272 | 0,000534172 | 0,01454663 | The Moose: suction sampler vs FISH | Cytoskeleton |
| gene_1086684 | 5,85695931 | 7,393851083 | 1,922309212 | 3,846338058 | 0,000119896 | 0,005637506 | The Moose: suction sampler vs FISH | Cytoskeleton |
| gene_314137 | 119,1458362 | 7,509155466 | 2,235074847 | 3,359688592 | 0,000780304 | 0,018637206 | The Moose: suction sampler vs FISH | Cytoskeleton |
| gene_1084954 | 234,1718311 | 7,537153421 | 2,008653623 | 3,752341037 | 0,000175191 | 0,007164197 | The Moose: suction sampler vs FISH | Cytoskeleton |

| | | | | | | | | |
|---|---|---|---|---|---|---|---|---|
| gene_55019 | 13,18606252 | 7,621850186 | 1,761757925 | 4,326275523 | 1,52E-05 | 0,001387794 | The Moose: suction sampler vs FISH | Cytoskeleton |
| gene_1721633 | 99,65974955 | 8,303388904 | 1,945450335 | 4,268106338 | 1,97E-05 | 0,001636034 | The Moose: suction sampler vs FISH | Cytoskeleton |
| gene_1088754 | 20,97573059 | 8,305087546 | 1,761530377 | 4,714700157 | 2,42E-06 | 0,00039596 | The Moose: suction sampler vs FISH | Cytoskeleton |
| gene_321523 | 161,0742749 | 9,182662788 | 1,796641092 | 5,111016792 | 3,20E-07 | 0,000102768 | The Moose: suction sampler vs FISH | Cytoskeleton |
| gene_347625 | 6,401687535 | -5,286660089 | 1,813286927 | -2,915512161 | 0,003551053 | 0,046448275 | The Moose: suction sampler vs FISH | Defense mechanisms |
| gene_209505 | 4,952877744 | -5,178053704 | 1,779227665 | -2,910281694 | 0,003611031 | 0,046940354 | The Moose: suction sampler vs FISH | Defense mechanisms |
| gene_874803 | 14,24864924 | -4,187252414 | 1,226127387 | -3,415022337 | 0,000637768 | 0,016320264 | The Moose: suction sampler vs FISH | Defense mechanisms |
| gene_642125 | 18,86560672 | -3,090316056 | 1,023199673 | -3,020247306 | 0,002525684 | 0,037994426 | The Moose: suction sampler vs FISH | Defense mechanisms |
| gene_915744 | 22,79311727 | -3,039069559 | 0,928114381 | -3,274455843 | 0,001058657 | 0,022589498 | The Moose: suction sampler vs FISH | Defense mechanisms |
| gene_220291 | 31,34788954 | -2,972828399 | 0,945090919 | -3,145547523 | 0,001657763 | 0,029678718 | The Moose: suction sampler vs FISH | Defense mechanisms |
| gene_1605370 | 24,9574167 | -2,713728439 | 0,916181075 | -2,962000104 | 0,003056477 | 0,042636836 | The Moose: suction sampler vs FISH | Defense mechanisms |
| gene_980494 | 158,6216888 | -2,674577265 | 0,685152642 | -3,903622496 | 9,48E-05 | 0,004839992 | The Moose: suction sampler vs FISH | Defense mechanisms |
| gene_273828 | 39,90636205 | -2,649713576 | 0,880399004 | -3,009673529 | 0,002615286 | 0,038835543 | The Moose: suction sampler vs FISH | Defense mechanisms |
| gene_280925 | 23,22527859 | -2,514970153 | 0,847816539 | -2,966408458 | 0,003013 | 0,042287372 | The Moose: suction sampler vs FISH | Defense mechanisms |
| gene_785241 | 21,7150533 | -2,498466701 | 0,753591455 | -3,315412726 | 0,000915079 | 0,02061142 | The Moose: suction sampler vs FISH | Defense mechanisms |
| gene_176366 | 317,000368 | -2,48569709 | 0,706004767 | -3,520793635 | 0,000430257 | 0,012732417 | The Moose: suction sampler vs FISH | Defense mechanisms |
| gene_1751729 | 146,9968944 | -2,366550204 | 0,722031217 | -3,277628649 | 0,00104683 | 0,022415289 | The Moose: suction sampler vs FISH | Defense mechanisms |
| gene_729806 | 46,02515867 | -2,307761707 | 0,731805179 | -3,153519234 | 0,001613146 | 0,029181652 | The Moose: suction sampler vs FISH | Defense mechanisms |
| gene_474671 | 179,791419 | -2,30074878 | 0,710454148 | -3,2384198 | 0,001201938 | 0,024365538 | The Moose: suction sampler vs FISH | Defense mechanisms |
| gene_751641 | 189,9057383 | -2,276991293 | 0,597743467 | -3,809311888 | 0,000139354 | 0,006191688 | The Moose: suction sampler vs FISH | Defense mechanisms |
| gene_284578 | 41,64167608 | -2,276107406 | 0,711888389 | -3,19728126 | 0,001387296 | 0,026542102 | The Moose: suction sampler vs FISH | Defense mechanisms |
| gene_284514 | 215,66386 | -2,130148809 | 0,593981705 | -3,586219561 | 0,000335507 | 0,010917666 | The Moose: suction sampler vs FISH | Defense mechanisms |
| gene_869875 | 135,1185208 | -2,078788878 | 0,63912419 | -3,252558596 | 0,00114371 | 0,023661913 | The Moose: suction sampler vs FISH | Defense mechanisms |
| gene_637232 | 94,80101186 | -2,073317029 | 0,662491916 | -3,129573327 | 0,001750604 | 0,030666956 | The Moose: suction sampler vs FISH | Defense mechanisms |
| gene_763779 | 29,19973571 | 2,193281335 | 0,742382064 | 2,954383521 | 0,003132944 | 0,043275647 | The Moose: suction sampler vs FISH | Defense mechanisms |
| gene_213244 | 46,12327314 | 2,483713536 | 0,831514003 | 2,986977402 | 0,002817505 | 0,040598723 | The Moose: suction sampler vs FISH | Defense mechanisms |
| gene_110940 | 30,58150157 | 2,543651709 | 0,87912659 | 2,893385023 | 0,003811138 | 0,048435547 | The Moose: suction sampler vs FISH | Defense mechanisms |
| gene_45012 | 46,5211547 | 2,580473608 | 0,790234976 | 3,26545102 | 0,001092899 | 0,023045642 | The Moose: suction sampler vs FISH | Defense mechanisms |
| gene_348541 | 32,65861609 | 2,590298839 | 0,828000881 | 3,128376913 | 0,001757746 | 0,030695208 | The Moose: suction sampler vs FISH | Defense mechanisms |
| gene_1200355 | 54,26076433 | 2,724551127 | 0,763914516 | 3,566565459 | 0,00036169 | 0,011430447 | The Moose: suction sampler vs FISH | Defense mechanisms |
| gene_328222 | 39,70823502 | 2,731036193 | 0,813224561 | 3,358280511 | 0,00078429 | 0,018701511 | The Moose: suction sampler vs FISH | Defense mechanisms |
| gene_586659 | 14,08510584 | 3,161035718 | 0,919615084 | 3,43734653 | 0,000587444 | 0,015452235 | The Moose: suction sampler vs FISH | Defense mechanisms |
| gene_285857 | 14,49363517 | 3,477190183 | 1,127732176 | 3,083347497 | 0,00204686 | 0,033673581 | The Moose: suction sampler vs FISH | Defense mechanisms |
| gene_344068 | 11,53388963 | 3,498422332 | 1,081653686 | 3,234327565 | 0,001219295 | 0,024539704 | The Moose: suction sampler vs FISH | Defense mechanisms |
| gene_1378774 | 20,82263393 | 3,501607366 | 1,112642174 | 3,147110049 | 0,001648929 | 0,029566792 | The Moose: suction sampler vs FISH | Defense mechanisms |
| gene_402570 | 44,23480393 | 3,532066101 | 0,862607686 | 4,094637872 | 4,23E-05 | 0,002811398 | The Moose: suction sampler vs FISH | Defense mechanisms |
| gene_809320 | 10,59725092 | 3,539315963 | 1,178971332 | 3,002037342 | 0,002681793 | 0,039329926 | The Moose: suction sampler vs FISH | Defense mechanisms |
| gene_591771 | 9,163688219 | 3,574589448 | 1,218356028 | 2,933944894 | 0,003346836 | 0,044929334 | The Moose: suction sampler vs FISH | Defense mechanisms |
| gene_1039827 | 16,34305131 | 3,598012222 | 1,092609821 | 3,293044006 | 0,00099109 | 0,021599578 | The Moose: suction sampler vs FISH | Defense mechanisms |
| gene_516507 | 30,32975466 | 3,646835954 | 1,096593869 | 3,325603084 | 0,000882275 | 0,02022603 | The Moose: suction sampler vs FISH | Defense mechanisms |
| gene_285856 | 13,74281565 | 3,684793139 | 1,147686577 | 3,210626674 | 0,001324459 | 0,02582757 | The Moose: suction sampler vs FISH | Defense mechanisms |
| gene_1746960 | 26,3152717 | 3,744774657 | 0,948570556 | 3,94780824 | 7,89E-05 | 0,004260333 | The Moose: suction sampler vs FISH | Defense mechanisms |
| gene_862798 | 10,62588829 | 3,750949396 | 1,292850627 | 2,901301448 | 0,003716162 | 0,047824342 | The Moose: suction sampler vs FISH | Defense mechanisms |
| gene_1853851 | 16,89556372 | 3,792404891 | 1,120447026 | 3,38472485 | 0,000712496 | 0,017536627 | The Moose: suction sampler vs FISH | Defense mechanisms |
| gene_1746959 | 64,22994276 | 3,853600427 | 0,780494632 | 4,93738236 | 7,92E-07 | 0,000191922 | The Moose: suction sampler vs FISH | Defense mechanisms |
| gene_591770 | 12,66920852 | 3,929493267 | 1,29691807 | 3,029870088 | 0,00244659 | 0,037233686 | The Moose: suction sampler vs FISH | Defense mechanisms |
| gene_1529428 | 7,49729208 | 3,997602842 | 1,34859009 | 2,964283122 | 0,00303389 | 0,042450764 | The Moose: suction sampler vs FISH | Defense mechanisms |
| gene_541854 | 7,693305434 | 4,531325423 | 1,50861819 | 3,0036264 | 0,002667827 | 0,039220187 | The Moose: suction sampler vs FISH | Defense mechanisms |
| gene_1839662 | 42,37293758 | 4,546549005 | 1,026057189 | 4,431087324 | 9,38E-06 | 0,001020169 | The Moose: suction sampler vs FISH | Defense mechanisms |
| gene_1637007 | 9,142647204 | 4,555613169 | 1,473804968 | 3,091055647 | 0,001994463 | 0,033118539 | The Moose: suction sampler vs FISH | Defense mechanisms |
| gene_1874213 | 55,00012399 | 4,708758265 | 0,970010459 | 4,854337621 | 1,21E-06 | 0,000251778 | The Moose: suction sampler vs FISH | Defense mechanisms |
| gene_850880 | 31,38983928 | 5,519859719 | 1,081370081 | 5,10450568 | 3,32E-07 | 0,000104323 | The Moose: suction sampler vs FISH | Defense mechanisms |
| gene_90337 | 8,418758948 | 5,523891997 | 1,557356791 | 3,546966264 | 0,000389694 | 0,011992803 | The Moose: suction sampler vs FISH | Defense mechanisms |
| gene_1674683 | 4,56955162 | 5,903432238 | 1,912014917 | 3,087545074 | 0,002018172 | 0,03334637 | The Moose: suction sampler vs FISH | Defense mechanisms |
| gene_465189 | 3,503848187 | 6,08723737 | 1,983074865 | 3,069595343 | 0,00214349 | 0,034569303 | The Moose: suction sampler vs FISH | Defense mechanisms |
| gene_1801898 | 8,206448211 | 6,113745999 | 1,648808168 | 3,70797896 | 0,00020892 | 0,008046858 | The Moose: suction sampler vs FISH | Defense mechanisms |
| gene_582253 | 4,431614073 | 6,232050226 | 1,970282064 | 3,163024391 | 0,001561392 | 0,028588655 | The Moose: suction sampler vs FISH | Defense mechanisms |
| gene_1018157 | 4,295712245 | 6,607574218 | 1,837468693 | 3,596020027 | 0,000323123 | 0,010665961 | The Moose: suction sampler vs FISH | Defense mechanisms |
| gene_629491 | 12,34396076 | 7,081223125 | 1,742278911 | 4,064345312 | 4,82E-05 | 0,003086725 | The Moose: suction sampler vs FISH | Defense mechanisms |
| gene_154283 | 6,77734138 | 7,710147418 | 1,671271063 | 4,613343455 | 3,96E-06 | 0,000556444 | The Moose: suction sampler vs FISH | Defense mechanisms |
| gene_1380995 | 24,6386988 | -7,516697236 | 1,581110086 | -4,754063176 | 1,99E-06 | 0,000349466 | The Moose: suction sampler vs FISH | Energy production and conversion |
| gene_222558 | 17,00425545 | -6,722228639 | 1,507756227 | -4,458432018 | 8,26E-06 | 0,000940188 | The Moose: suction sampler vs FISH | Energy production and conversion |
| gene_589829 | 14,45227919 | -6,54938199 | 1,509929566 | -4,33754139 | 1,44E-05 | 0,001337845 | The Moose: suction sampler vs FISH | Energy production and conversion |
| gene_197218 | 42,78789379 | -6,482133824 | 1,168565969 | -5,547084199 | 2,90E-08 | 1,96E-05 | The Moose: suction sampler vs FISH | Energy production and conversion |
| gene_600136 | 7,885082604 | -6,247606606 | 1,848559586 | -3,379716106 | 0,000725607 | 0,017752243 | The Moose: suction sampler vs FISH | Energy production and conversion |

| | | | | | | | | |
|---|---|---|---|---|---|---|---|---|
| gene_11903 | 5,942393988 | -6,137199924 | 1,807598865 | -3,395222271 | 0,000685729 | 0,017089708 | The Moose: suction sampler vs FISH | Energy production and conversion |
| gene_1374011 | 6,095550538 | -5,60293834 | 1,725716861 | -3,246730948 | 0,001167387 | 0,023920816 | The Moose: suction sampler vs FISH | Energy production and conversion |
| gene_701459 | 3,660384843 | -5,549661689 | 1,819125776 | -3,050730062 | 0,002282857 | 0,035723316 | The Moose: suction sampler vs FISH | Energy production and conversion |
| gene_222557 | 16,4622309 | -5,295469352 | 1,308601943 | -4,04666169 | 5,20E-05 | 0,003223638 | The Moose: suction sampler vs FISH | Energy production and conversion |
| gene_844111 | 9,442101174 | -4,987074515 | 1,454478843 | -3,428770752 | 0,000606321 | 0,015758826 | The Moose: suction sampler vs FISH | Energy production and conversion |
| gene_147759 | 7,968311804 | -4,642919957 | 1,585293927 | -2,928743924 | 0,003403347 | 0,045330064 | The Moose: suction sampler vs FISH | Energy production and conversion |
| gene_147760 | 45,01650471 | -4,338376009 | 1,038324654 | -4,178246169 | 2,94E-05 | 0,002186829 | The Moose: suction sampler vs FISH | Energy production and conversion |
| gene_959023 | 38,47473209 | -4,267765555 | 1,205828205 | -3,539281582 | 0,000401218 | 0,012211948 | The Moose: suction sampler vs FISH | Energy production and conversion |
| gene_525558 | 8,386535587 | -4,076368328 | 1,376264694 | -2,961907217 | 0,003057399 | 0,042641601 | The Moose: suction sampler vs FISH | Energy production and conversion |
| gene_1114198 | 55,52298458 | -4,051846314 | 1,10846846 | -3,65535553 | 0,000256826 | 0,009196768 | The Moose: suction sampler vs FISH | Energy production and conversion |
| gene_403417 | 19,05455365 | -4,049906719 | 1,207903312 | -3,352840148 | 0,000799869 | 0,018928075 | The Moose: suction sampler vs FISH | Energy production and conversion |
| gene_861084 | 136,504664 | -4,016738879 | 0,744500703 | -5,395211668 | 6,84E-08 | 3,72E-05 | The Moose: suction sampler vs FISH | Energy production and conversion |
| gene_861082 | 75,82898766 | -4,016064519 | 0,758225626 | -5,296661546 | 1,18E-07 | 5,15E-05 | The Moose: suction sampler vs FISH | Energy production and conversion |
| gene_1352477 | 143,3294019 | -3,984407301 | 1,097461616 | -3,630566429 | 0,0002828 | 0,009811166 | The Moose: suction sampler vs FISH | Energy production and conversion |
| gene_861083 | 215,7794024 | -3,826200999 | 0,718273001 | -5,32694532 | 9,99E-08 | 4,64E-05 | The Moose: suction sampler vs FISH | Energy production and conversion |
| gene_1031137 | 106,9695827 | -3,822321494 | 0,956805769 | -3,994877138 | 6,47E-05 | 0,003734434 | The Moose: suction sampler vs FISH | Energy production and conversion |
| gene_484732 | 69,97753855 | -3,746817038 | 1,25572501 | -2,983787859 | 0,002847041 | 0,040863967 | The Moose: suction sampler vs FISH | Energy production and conversion |
| gene_1405416 | 226,17178 | -3,580724118 | 1,103359486 | -3,245292368 | 0,001173301 | 0,02399028 | The Moose: suction sampler vs FISH | Energy production and conversion |
| gene_28282 | 15,82428392 | -3,576540696 | 1,063284582 | -3,363672112 | 0,000769129 | 0,01845682 | The Moose: suction sampler vs FISH | Energy production and conversion |
| gene_28281 | 54,52400602 | -3,475530347 | 0,904576161 | -3,842164427 | 0,000121954 | 0,005708679 | The Moose: suction sampler vs FISH | Energy production and conversion |
| gene_914189 | 14,40053034 | -3,24876839 | 1,121025569 | -2,898032372 | 0,003755118 | 0,048064598 | The Moose: suction sampler vs FISH | Energy production and conversion |
| gene_28283 | 80,54628025 | -3,171302167 | 0,80708943 | -3,929307026 | 8,52E-05 | 0,004497449 | The Moose: suction sampler vs FISH | Energy production and conversion |
| gene_28280 | 81,6658753 | -3,149305679 | 0,814111639 | -3,868395348 | 0,000109554 | 0,005293545 | The Moose: suction sampler vs FISH | Energy production and conversion |
| gene_1430223 | 18,71154432 | -3,116416902 | 1,012514324 | -3,077899077 | 0,002084655 | 0,034069312 | The Moose: suction sampler vs FISH | Energy production and conversion |
| gene_115225 | 15,60091092 | -3,095390044 | 0,997049318 | -3,104550586 | 0,001905683 | 0,032242091 | The Moose: suction sampler vs FISH | Energy production and conversion |
| gene_1430221 | 23,76477179 | -3,068396446 | 0,949653579 | -3,231069215 | 0,001233281 | 0,024687463 | The Moose: suction sampler vs FISH | Energy production and conversion |
| gene_326081 | 38,53925058 | -2,997602593 | 0,918354468 | -3,26410193 | 0,001098117 | 0,023109217 | The Moose: suction sampler vs FISH | Energy production and conversion |
| gene_98698 | 118,6433732 | -2,989398004 | 0,633890385 | -4,71595417 | 2,41E-06 | 0,000394407 | The Moose: suction sampler vs FISH | Energy production and conversion |
| gene_568421 | 16,5722104 | -2,983621869 | 0,98498095 | -3,029116317 | 0,002452702 | 0,037289539 | The Moose: suction sampler vs FISH | Energy production and conversion |
| gene_832597 | 85,48771935 | -2,886328922 | 0,716274636 | -4,029639998 | 5,59E-05 | 0,003387952 | The Moose: suction sampler vs FISH | Energy production and conversion |
| gene_438933 | 38,53066747 | -2,855439569 | 0,740308024 | -3,857096612 | 0,000114742 | 0,005461423 | The Moose: suction sampler vs FISH | Energy production and conversion |
| gene_404679 | 1663,797977 | -2,814321417 | 0,579168465 | -4,859244915 | 1,18E-06 | 0,000247371 | The Moose: suction sampler vs FISH | Energy production and conversion |
| gene_1232076 | 22,39842016 | -2,773930755 | 0,917364702 | -3,023803673 | 0,002496184 | 0,037769306 | The Moose: suction sampler vs FISH | Energy production and conversion |
| gene_1445059 | 50,75743322 | -2,752603568 | 0,635808788 | -4,329294627 | 1,50E-05 | 0,00137675 | The Moose: suction sampler vs FISH | Energy production and conversion |
| gene_1153099 | 160,9689797 | -2,731660969 | 0,846614752 | -3,226569063 | 0,00125284 | 0,024963328 | The Moose: suction sampler vs FISH | Energy production and conversion |
| gene_861081 | 27,6657039 | -2,633533832 | 0,860967708 | -3,058806744 | 0,002222205 | 0,035212473 | The Moose: suction sampler vs FISH | Energy production and conversion |
| gene_904341 | 217,4517414 | -2,570561178 | 0,565528709 | -4,545412353 | 5,48E-06 | 0,000701538 | The Moose: suction sampler vs FISH | Energy production and conversion |
| gene_55090 | 307,1332887 | -2,463990364 | 0,573856056 | -4,293742897 | 1,76E-05 | 0,001530565 | The Moose: suction sampler vs FISH | Energy production and conversion |
| gene_907746 | 307,8233655 | -2,446774985 | 0,62843937 | -3,893414544 | 9,88E-05 | 0,00497608 | The Moose: suction sampler vs FISH | Energy production and conversion |
| gene_1430220 | 82,53275464 | -2,431985262 | 0,760680874 | -3,197116352 | 0,001388089 | 0,026542102 | The Moose: suction sampler vs FISH | Energy production and conversion |
| gene_907748 | 260,2339004 | -2,431465888 | 0,644261395 | -3,774036295 | 0,000160627 | 0,006799584 | The Moose: suction sampler vs FISH | Energy production and conversion |
| gene_438930 | 323,3970448 | -2,383124348 | 0,575785197 | -4,138912153 | 3,49E-05 | 0,002454294 | The Moose: suction sampler vs FISH | Energy production and conversion |
| gene_664933 | 485,5673443 | -2,366763773 | 0,606724331 | -3,900888182 | 9,58E-05 | 0,0048661 | The Moose: suction sampler vs FISH | Energy production and conversion |
| gene_438932 | 80,13310499 | -2,340595127 | 0,679549467 | -3,444333693 | 0,000572469 | 0,015238755 | The Moose: suction sampler vs FISH | Energy production and conversion |
| gene_237309 | 1722,136541 | -2,294681496 | 0,610269178 | -3,7601137 | 0,000169836 | 0,007031351 | The Moose: suction sampler vs FISH | Energy production and conversion |
| gene_907747 | 268,5824356 | -2,288120174 | 0,736061736 | -3,108598183 | 0,001879772 | 0,032001019 | The Moose: suction sampler vs FISH | Energy production and conversion |
| gene_1181686 | 4549,619229 | -2,255226585 | 0,579664275 | -3,890573704 | 0,000100007 | 0,004993997 | The Moose: suction sampler vs FISH | Energy production and conversion |
| gene_985774 | 415,7025449 | -2,247943029 | 0,755946355 | -2,973680626 | 0,002942512 | 0,041640225 | The Moose: suction sampler vs FISH | Energy production and conversion |
| gene_119919 | 214,0842505 | -2,178685577 | 0,735222698 | -2,963300212 | 0,003043596 | 0,042505587 | The Moose: suction sampler vs FISH | Energy production and conversion |
| gene_361274 | 94,95689945 | -2,1573157 | 0,712698621 | -3,026967691 | 0,002470204 | 0,037492065 | The Moose: suction sampler vs FISH | Energy production and conversion |
| gene_438927 | 114,5783007 | -2,116759997 | 0,576472186 | -3,671920429 | 0,000240735 | 0,008818332 | The Moose: suction sampler vs FISH | Energy production and conversion |
| gene_438925 | 269,2708045 | -2,095418846 | 0,557943835 | -3,755608922 | 0,000172921 | 0,007107441 | The Moose: suction sampler vs FISH | Energy production and conversion |
| gene_632988 | 2249,326887 | -2,088623411 | 0,585333213 | -3,568263963 | 0,000359354 | 0,011381105 | The Moose: suction sampler vs FISH | Energy production and conversion |
| gene_1400284 | 36,98250984 | -2,085104297 | 0,698132 | -2,986690622 | 0,00282015 | 0,040618761 | The Moose: suction sampler vs FISH | Energy production and conversion |
| gene_566652 | 2819,606172 | -2,005557826 | 0,607430886 | -3,301705382 | 0,00096099 | 0,021202975 | The Moose: suction sampler vs FISH | Energy production and conversion |
| gene_141 | 2494,824278 | 2,013889781 | 0,691370884 | 2,91289354 | 0,003580967 | 0,04668988 | The Moose: suction sampler vs FISH | Energy production and conversion |
| gene_99904 | 158,3141168 | 2,085147336 | 0,690609786 | 3,019284374 | 0,002533726 | 0,038094062 | The Moose: suction sampler vs FISH | Energy production and conversion |
| gene_138 | 9810,210983 | 2,257147935 | 0,739533736 | 3,052123013 | 0,00227229 | 0,035644662 | The Moose: suction sampler vs FISH | Energy production and conversion |
| gene_370465 | 261,9805663 | 2,322467978 | 0,759836162 | 3,056537835 | 0,002239092 | 0,035360455 | The Moose: suction sampler vs FISH | Energy production and conversion |
| gene_145 | 7029,405422 | 2,350212301 | 0,750167685 | 3,132915942 | 0,00173079 | 0,030426115 | The Moose: suction sampler vs FISH | Energy production and conversion |
| gene_146 | 2087,640689 | 2,41039106 | 0,727747 | 3,312127784 | 0,000925892 | 0,02076659 | The Moose: suction sampler vs FISH | Energy production and conversion |
| gene_410432 | 33,3008841 | 2,436923744 | 0,779600377 | 3,125862707 | 0,001772843 | 0,030843646 | The Moose: suction sampler vs FISH | Energy production and conversion |
| gene_140 | 10576,87137 | 2,446706522 | 0,733150448 | 3,337250256 | 0,000846117 | 0,019702943 | The Moose: suction sampler vs FISH | Energy production and conversion |
| gene_1116540 | 109,8838692 | 2,512389883 | 0,749603306 | 3,351625936 | 0,000803385 | 0,018965252 | The Moose: suction sampler vs FISH | Energy production and conversion |
| gene_56456 | 31,91133395 | 2,605740486 | 0,88470749 | 2,945313015 | 0,003226281 | 0,044000023 | The Moose: suction sampler vs FISH | Energy production and conversion |

| | | | | | | | | |
|---|---|---|---|---|---|---|---|---|
| gene_1022976 | 23,98311635 | 2,732712705 | 0,829440323 | 3,294646557 | 0,000985455 | 0,021527467 | The Moose: suction sampler vs FISH | Energy production and conversion |
| gene_808996 | 40,43529859 | 2,733037353 | 0,854406724 | 3,198754501 | 0,001380226 | 0,026474467 | The Moose: suction sampler vs FISH | Energy production and conversion |
| gene_391445 | 62,46760843 | 2,758696814 | 0,767920364 | 3,592425653 | 0,000327614 | 0,010765826 | The Moose: suction sampler vs FISH | Energy production and conversion |
| gene_440590 | 233,364292 | 2,774652507 | 0,592079107 | 4,686286809 | 2,78E-06 | 0,0004329 | The Moose: suction sampler vs FISH | Energy production and conversion |
| gene_381046 | 100,6046623 | 2,807010381 | 0,789728238 | 3,55440042 | 0,000378842 | 0,01178985 | The Moose: suction sampler vs FISH | Energy production and conversion |
| gene_1221972 | 22,46881548 | 2,928845657 | 0,989917262 | 2,958677224 | 0,003089625 | 0,042944271 | The Moose: suction sampler vs FISH | Energy production and conversion |
| gene_25544 | 47,2392807 | 2,955995026 | 0,861429367 | 3,431500178 | 0,000600253 | 0,015635485 | The Moose: suction sampler vs FISH | Energy production and conversion |
| gene_888492 | 28,31064388 | 2,963281584 | 0,951860502 | 3,113146914 | 0,001851038 | 0,031687986 | The Moose: suction sampler vs FISH | Energy production and conversion |
| gene_1549622 | 10,74156271 | 3,03393226 | 1,052038823 | 2,883859601 | 0,003928339 | 0,049252677 | The Moose: suction sampler vs FISH | Energy production and conversion |
| gene_834923 | 13,8321304 | 3,050738428 | 1,058787455 | 2,881351129 | 0,003959742 | 0,049527126 | The Moose: suction sampler vs FISH | Energy production and conversion |
| gene_577859 | 22,25229136 | 3,118252891 | 0,955889965 | 3,262146278 | 0,001105721 | 0,023216026 | The Moose: suction sampler vs FISH | Energy production and conversion |
| gene_1349171 | 57,59574762 | 3,120672031 | 0,843235627 | 3,700830385 | 0,000214895 | 0,008176582 | The Moose: suction sampler vs FISH | Energy production and conversion |
| gene_1428149 | 291,9537894 | 3,125537328 | 0,852134683 | 3,667891226 | 0,000244559 | 0,008927261 | The Moose: suction sampler vs FISH | Energy production and conversion |
| gene_1609437 | 163,8219122 | 3,152066854 | 0,736426644 | 4,280218376 | 1,87E-05 | 0,001579829 | The Moose: suction sampler vs FISH | Energy production and conversion |
| gene_616447 | 24,4378942 | 3,154716297 | 1,091756183 | 2,88957951 | 0,003857574 | 0,048780915 | The Moose: suction sampler vs FISH | Energy production and conversion |
| gene_57853 | 17,07625316 | 3,157774659 | 1,002719256 | 3,149211149 | 0,001637119 | 0,029448747 | The Moose: suction sampler vs FISH | Energy production and conversion |
| gene_859612 | 198,2520811 | 3,172215914 | 0,854017963 | 3,714460409 | 0,000203638 | 0,007909376 | The Moose: suction sampler vs FISH | Energy production and conversion |
| gene_966796 | 24,03824819 | 3,240101657 | 1,043298099 | 3,105633625 | 0,001898718 | 0,032177526 | The Moose: suction sampler vs FISH | Energy production and conversion |
| gene_211619 | 21,89132444 | 3,249933132 | 0,988075818 | 3,289153598 | 0,001004892 | 0,021789908 | The Moose: suction sampler vs FISH | Energy production and conversion |
| gene_56452 | 40,35755934 | 3,260220416 | 0,881801145 | 3,697228602 | 0,000217966 | 0,008239072 | The Moose: suction sampler vs FISH | Energy production and conversion |
| gene_190667 | 89,51909116 | 3,282261815 | 0,927999475 | 3,53692206 | 0,000404819 | 0,012270495 | The Moose: suction sampler vs FISH | Energy production and conversion |
| gene_146856 | 77,54989021 | 3,288567664 | 0,990602289 | 3,319765865 | 0,00090093 | 0,02049231 | The Moose: suction sampler vs FISH | Energy production and conversion |
| gene_442820 | 968,3250361 | 3,328408721 | 0,795509994 | 4,183993596 | 2,86E-05 | 0,002158827 | The Moose: suction sampler vs FISH | Energy production and conversion |
| gene_258993 | 37,73176278 | 3,362829439 | 1,016082749 | 3,309601943 | 0,000934287 | 0,020887457 | The Moose: suction sampler vs FISH | Energy production and conversion |
| gene_512938 | 186,0868452 | 3,363045425 | 0,898447909 | 3,74317241 | 0,000181711 | 0,007327963 | The Moose: suction sampler vs FISH | Energy production and conversion |
| gene_680122 | 25,55490606 | 3,369599111 | 0,957060366 | 3,520780121 | 0,000430279 | 0,012732417 | The Moose: suction sampler vs FISH | Energy production and conversion |
| gene_1678187 | 10,53372215 | 3,406458693 | 1,177211742 | 2,893666936 | 0,003807718 | 0,048417221 | The Moose: suction sampler vs FISH | Energy production and conversion |
| gene_1806082 | 52,76445443 | 3,44450706 | 0,977106446 | 3,525211686 | 0,000423145 | 0,012611664 | The Moose: suction sampler vs FISH | Energy production and conversion |
| gene_1037026 | 55,2062646 | 3,459503753 | 0,933718507 | 3,705082127 | 0,000211322 | 0,008092056 | The Moose: suction sampler vs FISH | Energy production and conversion |
| gene_952377 | 26,72974518 | 3,472701329 | 1,014403064 | 3,423393968 | 0,000618444 | 0,015965413 | The Moose: suction sampler vs FISH | Energy production and conversion |
| gene_272668 | 13,41823256 | 3,527871664 | 1,169158167 | 3,017446026 | 0,002549144 | 0,038255395 | The Moose: suction sampler vs FISH | Energy production and conversion |
| gene_1542416 | 155,5694647 | 3,563198869 | 0,975760341 | 3,651715202 | 0,000260495 | 0,009278386 | The Moose: suction sampler vs FISH | Energy production and conversion |
| gene_1798902 | 16,53167438 | 3,575263346 | 1,192552538 | 2,997992316 | 0,002717645 | 0,039626257 | The Moose: suction sampler vs FISH | Energy production and conversion |
| gene_59029 | 20,18692276 | 3,585627395 | 1,165236992 | 3,077165778 | 0,00208979 | 0,034092895 | The Moose: suction sampler vs FISH | Energy production and conversion |
| gene_377691 | 9,177145998 | 3,587154856 | 1,231997756 | 2,911656973 | 0,003595172 | 0,0468273 | The Moose: suction sampler vs FISH | Energy production and conversion |
| gene_1839443 | 26,21359769 | 3,601614622 | 1,200315962 | 3,000555467 | 0,002694877 | 0,03943505 | The Moose: suction sampler vs FISH | Energy production and conversion |
| gene_928547 | 103,3902837 | 3,603731583 | 0,852101159 | 4,229229763 | 2,34E-05 | 0,001865013 | The Moose: suction sampler vs FISH | Energy production and conversion |
| gene_1542414 | 516,1789299 | 3,638334796 | 0,865320145 | 4,204611228 | 2,62E-05 | 0,002024043 | The Moose: suction sampler vs FISH | Energy production and conversion |
| gene_381045 | 515,6433177 | 3,718170237 | 0,775631767 | 4,793731243 | 1,64E-06 | 0,000308493 | The Moose: suction sampler vs FISH | Energy production and conversion |
| gene_663096 | 15,59224536 | 3,732246433 | 1,134567344 | 3,289576817 | 0,001003382 | 0,021783436 | The Moose: suction sampler vs FISH | Energy production and conversion |
| gene_953492 | 12,78318189 | 3,737715088 | 1,259922097 | 2,966623965 | 0,00301089 | 0,042273902 | The Moose: suction sampler vs FISH | Energy production and conversion |
| gene_1806081 | 57,04407448 | 3,741712788 | 0,899654104 | 4,159057101 | 3,20E-05 | 0,002301018 | The Moose: suction sampler vs FISH | Energy production and conversion |
| gene_1667224 | 19,79361754 | 3,804726187 | 1,096654685 | 3,469393091 | 0,000521636 | 0,014321278 | The Moose: suction sampler vs FISH | Energy production and conversion |
| gene_1087997 | 9,618580014 | 3,833229159 | 1,282822577 | 2,98812106 | 0,002806983 | 0,040478873 | The Moose: suction sampler vs FISH | Energy production and conversion |
| gene_115816 | 64,52357278 | 3,839546643 | 0,87086885 | 4,408868963 | 1,04E-05 | 0,001090258 | The Moose: suction sampler vs FISH | Energy production and conversion |
| gene_680121 | 38,01118938 | 3,885782745 | 0,982739553 | 3,95403109 | 7,68E-05 | 0,004205597 | The Moose: suction sampler vs FISH | Energy production and conversion |
| gene_1454211 | 12,02825198 | 3,936497444 | 1,302948664 | 3,021222213 | 0,002517565 | 0,037936859 | The Moose: suction sampler vs FISH | Energy production and conversion |
| gene_1313119 | 28,57755294 | 3,977500253 | 1,076214278 | 3,695825574 | 0,000219173 | 0,008269812 | The Moose: suction sampler vs FISH | Energy production and conversion |
| gene_179448 | 16,49639971 | 3,993809853 | 1,282306749 | 3,114551066 | 0,001842251 | 0,031613107 | The Moose: suction sampler vs FISH | Energy production and conversion |
| gene_370420 | 120,9369605 | 4,039887609 | 0,918264269 | 4,399482531 | 1,09E-05 | 0,001124706 | The Moose: suction sampler vs FISH | Energy production and conversion |
| gene_502326 | 113,0367789 | 4,063865887 | 1,125904766 | 3,609422404 | 0,00030688 | 0,01032331 | The Moose: suction sampler vs FISH | Energy production and conversion |
| gene_649468 | 14,21389261 | 4,068185496 | 1,105915659 | 3,678567586 | 0,000234548 | 0,008678259 | The Moose: suction sampler vs FISH | Energy production and conversion |
| gene_764412 | 45,61170962 | 4,095959842 | 0,927909224 | 4,414181619 | 1,01E-05 | 0,001071481 | The Moose: suction sampler vs FISH | Energy production and conversion |
| gene_1330737 | 8,523684677 | 4,105747694 | 1,248181858 | 3,289382607 | 0,001004074 | 0,021787013 | The Moose: suction sampler vs FISH | Energy production and conversion |
| gene_905279 | 388,6594805 | 4,123053548 | 0,889179044 | 4,636921638 | 3,54E-06 | 0,000515334 | The Moose: suction sampler vs FISH | Energy production and conversion |
| gene_1858343 | 13,01053766 | 4,127759578 | 1,237624665 | 3,335227307 | 0,000852297 | 0,01978737 | The Moose: suction sampler vs FISH | Energy production and conversion |
| gene_535556 | 105,7738678 | 4,136239684 | 0,883933286 | 4,679357309 | 2,88E-06 | 0,000439641 | The Moose: suction sampler vs FISH | Energy production and conversion |
| gene_1771470 | 8,57034763 | 4,159671034 | 1,412274039 | 2,945371026 | 0,003225676 | 0,044000023 | The Moose: suction sampler vs FISH | Energy production and conversion |
| gene_353448 | 19,32377378 | 4,174308816 | 1,130259631 | 3,693230033 | 0,000221424 | 0,008343999 | The Moose: suction sampler vs FISH | Energy production and conversion |
| gene_978855 | 34,55828474 | 4,180757355 | 1,015270924 | 4,117873621 | 3,82E-05 | 0,002619796 | The Moose: suction sampler vs FISH | Energy production and conversion |
| gene_1542415 | 75,28384616 | 4,252647237 | 1,066108328 | 3,988944769 | 6,64E-05 | 0,003784463 | The Moose: suction sampler vs FISH | Energy production and conversion |
| gene_432691 | 14,59355431 | 4,267865668 | 1,411845611 | 3,02289828 | 0,002503664 | 0,037843505 | The Moose: suction sampler vs FISH | Energy production and conversion |
| gene_1394492 | 16,6578573 | 4,31493958 | 1,088633003 | 3,963631056 | 7,38E-05 | 0,004111117 | The Moose: suction sampler vs FISH | Energy production and conversion |
| gene_1151695 | 23,16244802 | 4,348073002 | 1,132640938 | 3,838880317 | 0,000123597 | 0,005759871 | The Moose: suction sampler vs FISH | Energy production and conversion |
| gene_371362 | 8,22330813 | 4,352155985 | 1,37424222 | 3,166949698 | 0,00154047 | 0,028358408 | The Moose: suction sampler vs FISH | Energy production and conversion |

| | | | | | | | | |
|---|---|---|---|---|---|---|---|---|
| gene_1508152 | 13,86776311 | 4,402647835 | 1,267066227 | 3,474678546 | 0,000511466 | 0,014165535 | The Moose: suction sampler vs FISH | Energy production and conversion |
| gene_177380 | 17,31365321 | 4,4043126 | 1,158620743 | 3,801341058 | 0,000143915 | 0,006317896 | The Moose: suction sampler vs FISH | Energy production and conversion |
| gene_50636 | 11,15541707 | 4,427268958 | 1,36498305 | 3,243460757 | 0,001180871 | 0,024091404 | The Moose: suction sampler vs FISH | Energy production and conversion |
| gene_1104525 | 32,89531545 | 4,461441495 | 1,136113778 | 3,926931951 | 8,60E-05 | 0,004520338 | The Moose: suction sampler vs FISH | Energy production and conversion |
| gene_160760 | 14,24664234 | 4,470687445 | 1,442948518 | 3,098300037 | 0,001946343 | 0,03263058 | The Moose: suction sampler vs FISH | Energy production and conversion |
| gene_1846333 | 7,285591422 | 4,500535656 | 1,350737109 | 3,33191087 | 0,000862519 | 0,019952028 | The Moose: suction sampler vs FISH | Energy production and conversion |
| gene_1271082 | 8,695453296 | 4,518714325 | 1,50483383 | 3,002799535 | 0,002675086 | 0,039302195 | The Moose: suction sampler vs FISH | Energy production and conversion |
| gene_1187975 | 7,120119787 | 4,519100733 | 1,376041892 | 3,2841302 | 0,001022977 | 0,022039457 | The Moose: suction sampler vs FISH | Energy production and conversion |
| gene_350061 | 40,60966499 | 4,560383578 | 1,046363373 | 4,358317288 | 1,31E-05 | 0,001269942 | The Moose: suction sampler vs FISH | Energy production and conversion |
| gene_370418 | 142,3834803 | 4,605540509 | 0,923875008 | 4,985025537 | 6,20E-07 | 0,000158543 | The Moose: suction sampler vs FISH | Energy production and conversion |
| gene_576572 | 12,58681777 | 4,608204539 | 1,305265092 | 3,530474054 | 0,000414816 | 0,012450404 | The Moose: suction sampler vs FISH | Energy production and conversion |
| gene_1499847 | 19,59438184 | 4,686127607 | 1,298246837 | 3,609581377 | 0,000306692 | 0,01032308 | The Moose: suction sampler vs FISH | Energy production and conversion |
| gene_26356 | 93,36181504 | 4,699879891 | 0,974589845 | 4,822418288 | 1,42E-06 | 0,000279769 | The Moose: suction sampler vs FISH | Energy production and conversion |
| gene_1087996 | 13,12509341 | 4,740011757 | 1,063338836 | 4,457668241 | 8,29E-06 | 0,00094055 | The Moose: suction sampler vs FISH | Energy production and conversion |
| gene_1437915 | 7,107545608 | 4,747254884 | 1,557933921 | 3,047147776 | 0,002310241 | 0,035993986 | The Moose: suction sampler vs FISH | Energy production and conversion |
| gene_286894 | 131,8768047 | 4,753719246 | 0,885966076 | 5,365577053 | 8,07E-08 | 4,14E-05 | The Moose: suction sampler vs FISH | Energy production and conversion |
| gene_299961 | 65,91377264 | 4,817464695 | 0,920348942 | 5,234389346 | 1,66E-07 | 6,33E-05 | The Moose: suction sampler vs FISH | Energy production and conversion |
| gene_681656 | 82,0797257 | 4,854228403 | 0,92318141 | 5,258152243 | 1,46E-07 | 5,81E-05 | The Moose: suction sampler vs FISH | Energy production and conversion |
| gene_1251823 | 28,01340046 | 4,899957343 | 1,00275325 | 4,886503576 | 1,03E-06 | 0,000226853 | The Moose: suction sampler vs FISH | Energy production and conversion |
| gene_849114 | 106,1089259 | 5,025790937 | 1,101337435 | 4,5633525 | 5,03E-06 | 0,000660549 | The Moose: suction sampler vs FISH | Energy production and conversion |
| gene_1251326 | 8,078378821 | 5,047947314 | 1,519256576 | 3,322643057 | 0,000891689 | 0,020370179 | The Moose: suction sampler vs FISH | Energy production and conversion |
| gene_1192657 | 23,33724103 | 5,096334071 | 1,313835226 | 3,878975057 | 0,000104898 | 0,005142989 | The Moose: suction sampler vs FISH | Energy production and conversion |
| gene_603015 | 22,32425607 | 5,09690313 | 1,020144864 | 4,996254269 | 5,85E-07 | 0,000151703 | The Moose: suction sampler vs FISH | Energy production and conversion |
| gene_199949 | 6,482616845 | 5,135471587 | 1,598185338 | 3,213314167 | 0,001312127 | 0,025713863 | The Moose: suction sampler vs FISH | Energy production and conversion |
| gene_16344 | 67,98554758 | 5,137818574 | 1,090944026 | 4,709516211 | 2,48E-06 | 0,000403469 | The Moose: suction sampler vs FISH | Energy production and conversion |
| gene_1121444 | 82,66649749 | 5,174658345 | 0,946315373 | 5,468217566 | 4,55E-08 | 2,78E-05 | The Moose: suction sampler vs FISH | Energy production and conversion |
| gene_466280 | 13,09526137 | 5,198040572 | 1,328979929 | 3,911301035 | 9,18E-05 | 0,004738068 | The Moose: suction sampler vs FISH | Energy production and conversion |
| gene_877223 | 6,357243809 | 5,202307935 | 1,63008956 | 3,191424607 | 0,001415731 | 0,026885368 | The Moose: suction sampler vs FISH | Energy production and conversion |
| gene_1645557 | 11,11955022 | 5,304219506 | 1,650710372 | 3,213295073 | 0,001312214 | 0,025713863 | The Moose: suction sampler vs FISH | Energy production and conversion |
| gene_1573554 | 6,635290115 | 5,320043056 | 1,725391332 | 3,083383438 | 0,002046613 | 0,033673581 | The Moose: suction sampler vs FISH | Energy production and conversion |
| gene_1781137 | 11,65558337 | 5,351552277 | 1,427248392 | 3,749559158 | 0,000177146 | 0,00720569 | The Moose: suction sampler vs FISH | Energy production and conversion |
| gene_934509 | 1208,774149 | 5,353933273 | 0,974447137 | 5,49432911 | 3,92E-08 | 2,46E-05 | The Moose: suction sampler vs FISH | Energy production and conversion |
| gene_730774 | 8,422776622 | 5,362569248 | 1,608032539 | 3,334863642 | 0,000853412 | 0,019803751 | The Moose: suction sampler vs FISH | Energy production and conversion |
| gene_436275 | 11,39179783 | 5,374628242 | 1,509740303 | 3,559968711 | 0,000370899 | 0,011601651 | The Moose: suction sampler vs FISH | Energy production and conversion |
| gene_16343 | 10,34529625 | 5,385400445 | 1,505456046 | 3,577255184 | 0,000347221 | 0,01114583 | The Moose: suction sampler vs FISH | Energy production and conversion |
| gene_1786311 | 11,93031741 | 5,512336341 | 1,334689438 | 4,130051669 | 3,63E-05 | 0,00251769 | The Moose: suction sampler vs FISH | Energy production and conversion |
| gene_1434043 | 9,008323098 | 5,587456362 | 1,50424754 | 3,714452716 | 0,000203644 | 0,007909376 | The Moose: suction sampler vs FISH | Energy production and conversion |
| gene_1786364 | 26,18676062 | 5,637359901 | 1,273574972 | 4,426406003 | 9,58E-06 | 0,001029043 | The Moose: suction sampler vs FISH | Energy production and conversion |
| gene_849113 | 45,29910444 | 5,675946639 | 1,114709499 | 5,091861729 | 3,55E-07 | 0,000110082 | The Moose: suction sampler vs FISH | Energy production and conversion |
| gene_1250406 | 40,79063116 | 5,676463727 | 0,996183302 | 5,69821208 | 1,21E-08 | 9,99E-06 | The Moose: suction sampler vs FISH | Energy production and conversion |
| gene_199948 | 4,974608729 | 5,696845556 | 1,822154587 | 3,126433727 | 0,001769404 | 0,030814148 | The Moose: suction sampler vs FISH | Energy production and conversion |
| gene_350060 | 44,41759571 | 5,748861662 | 1,14879653 | 5,004247064 | 5,61E-07 | 0,000148858 | The Moose: suction sampler vs FISH | Energy production and conversion |
| gene_1857433 | 18,61681234 | 5,793337597 | 1,173716125 | 4,935893333 | 7,98E-07 | 0,000192756 | The Moose: suction sampler vs FISH | Energy production and conversion |
| gene_1898238 | 7,951447307 | 5,84560446 | 1,536115056 | 3,805447018 | 0,000141548 | 0,006255124 | The Moose: suction sampler vs FISH | Energy production and conversion |
| gene_1361086 | 4,475426029 | 5,895055565 | 1,989032431 | 2,963780517 | 0,003038849 | 0,042472454 | The Moose: suction sampler vs FISH | Energy production and conversion |
| gene_337239 | 4,659085553 | 5,94403198 | 1,834775003 | 3,239651712 | 0,001196758 | 0,024295584 | The Moose: suction sampler vs FISH | Energy production and conversion |
| gene_1725240 | 4,718557605 | 5,955591011 | 1,957387269 | 3,042622738 | 0,002345262 | 0,036262681 | The Moose: suction sampler vs FISH | Energy production and conversion |
| gene_1013652 | 74,54066184 | 5,957759589 | 0,964141396 | 6,179342171 | 6,44E-10 | 1,24E-06 | The Moose: suction sampler vs FISH | Energy production and conversion |
| gene_97564 | 3,207465383 | 5,961117812 | 2,041915915 | 2,919374774 | 0,003507343 | 0,046082171 | The Moose: suction sampler vs FISH | Energy production and conversion |
| gene_382737 | 5,438209002 | 6,037999235 | 1,776064803 | 3,399650297 | 0,000674721 | 0,016904601 | The Moose: suction sampler vs FISH | Energy production and conversion |
| gene_610957 | 4,75554504 | 6,059959692 | 2,014309434 | 3,0084552 | 0,002625795 | 0,038881355 | The Moose: suction sampler vs FISH | Energy production and conversion |
| gene_927716 | 3,605615359 | 6,083060913 | 1,934002662 | 3,145321893 | 0,001659042 | 0,029688834 | The Moose: suction sampler vs FISH | Energy production and conversion |
| gene_50282 | 6,07425989 | 6,118938164 | 1,798153528 | 3,402900847 | 0,000666745 | 0,016775973 | The Moose: suction sampler vs FISH | Energy production and conversion |
| gene_1312523 | 3,425097656 | 6,13579766 | 1,962055769 | 3,12722898 | 0,001764624 | 0,030769903 | The Moose: suction sampler vs FISH | Energy production and conversion |
| gene_1583326 | 11,86283263 | 6,175988665 | 1,542565041 | 4,003713621 | 6,24E-05 | 0,003640481 | The Moose: suction sampler vs FISH | Energy production and conversion |
| gene_702895 | 4,795719376 | 6,248581013 | 2,051730648 | 3,045517217 | 0,002322805 | 0,036082789 | The Moose: suction sampler vs FISH | Energy production and conversion |
| gene_138352 | 4,847133629 | 6,315037636 | 1,891347854 | 3,338908611 | 0,000841082 | 0,019610568 | The Moose: suction sampler vs FISH | Energy production and conversion |
| gene_13321 | 3,565447056 | 6,521961312 | 2,168377138 | 3,007761518 | 0,002631796 | 0,038938807 | The Moose: suction sampler vs FISH | Energy production and conversion |
| gene_50637 | 6,093725039 | 6,554799671 | 1,872858819 | 3,499889903 | 0,00046545 | 0,013354236 | The Moose: suction sampler vs FISH | Energy production and conversion |
| gene_890191 | 7,836804242 | 6,689983026 | 1,747087497 | 3,829220367 | 0,00012855 | 0,005886125 | The Moose: suction sampler vs FISH | Energy production and conversion |
| gene_140888 | 10,18823786 | 6,789680696 | 1,724993715 | 3,936061122 | 8,28E-05 | 0,004398728 | The Moose: suction sampler vs FISH | Energy production and conversion |
| gene_769394 | 4,479455319 | 6,852735551 | 2,321009064 | 2,952481168 | 0,003152313 | 0,043383106 | The Moose: suction sampler vs FISH | Energy production and conversion |
| gene_132139 | 3,462709933 | 6,887837055 | 2,129020186 | 3,235214536 | 0,001215514 | 0,024511793 | The Moose: suction sampler vs FISH | Energy production and conversion |
| gene_1047996 | 4,030663732 | 7,257992763 | 2,0205858 | 3,592024036 | 0,00032812 | 0,010777613 | The Moose: suction sampler vs FISH | Energy production and conversion |
| gene_1497561 | 6,489263404 | 7,586893241 | 1,686837162 | 4,497703402 | 6,87E-06 | 0,000833891 | The Moose: suction sampler vs FISH | Energy production and conversion |

| | | | | | | | | |
|---|---|---|---|---|---|---|---|---|
| gene_512939 | 6,300289245 | 7,594125751 | 2,068699639 | 3,67096586 | 0,000241636 | 0,008843513 | The Moose: suction sampler vs FISH | Energy production and conversion |
| gene_939541 | 9,373821975 | 7,695874866 | 1,669027265 | 4,610994097 | 4,01E-06 | 0,000561697 | The Moose: suction sampler vs FISH | Energy production and conversion |
| gene_1657013 | 28,85235357 | 8,338277604 | 1,62448489 | 5,13287483 | 2,85E-07 | 9,36E-05 | The Moose: suction sampler vs FISH | Energy production and conversion |
| gene_1399214 | 125,5668904 | 2,392089893 | 0,823236383 | 2,905714497 | 0,003664156 | 0,047400856 | The Moose: suction sampler vs FISH | Extracellular structures |
| gene_856921 | 31,33966946 | 2,719094693 | 0,913001917 | 2,978191659 | 0,002899546 | 0,041270768 | The Moose: suction sampler vs FISH | Extracellular structures |
| gene_603502 | 15,59002535 | 3,294682495 | 0,976912978 | 3,372544504 | 0,00074477 | 0,0180868 | The Moose: suction sampler vs FISH | Extracellular structures |
| gene_1860 | 16,29078384 | 3,325638939 | 1,01050729 | 3,29105883 | 0,00099811 | 0,021713927 | The Moose: suction sampler vs FISH | Extracellular structures |
| gene_831491 | 19,46151093 | 3,533843222 | 1,163103199 | 3,038288627 | 0,00237926 | 0,03658043 | The Moose: suction sampler vs FISH | Extracellular structures |
| gene_1668434 | 17,61684546 | 3,670498196 | 1,057026434 | 3,472475311 | 0,000515682 | 0,014214983 | The Moose: suction sampler vs FISH | Extracellular structures |
| gene_1196742 | 63,86989338 | 3,701024937 | 0,832693657 | 4,444641681 | 8,80E-06 | 0,000973567 | The Moose: suction sampler vs FISH | Extracellular structures |
| gene_1196741 | 8,990968735 | 3,79265211 | 1,282185851 | 2,957958167 | 0,003096841 | 0,04300388 | The Moose: suction sampler vs FISH | Extracellular structures |
| gene_1001708 | 14,27498831 | 3,885423739 | 1,164865226 | 3,335513544 | 0,00085142 | 0,019776264 | The Moose: suction sampler vs FISH | Extracellular structures |
| gene_109987 | 8,690861215 | 4,45732061 | 1,453650377 | 3,066294812 | 0,002167295 | 0,03478518 | The Moose: suction sampler vs FISH | Extracellular structures |
| gene_616399 | 11,44880703 | -6,890809998 | 1,598577017 | -4,310589934 | 1,63E-05 | 0,001451729 | The Moose: suction sampler vs FISH | Function unknown |
| gene_404376 | 9,895060046 | -6,863143826 | 1,894549718 | -3,622572562 | 0,000291688 | 0,009987411 | The Moose: suction sampler vs FISH | Function unknown |
| gene_178079 | 4,422453779 | -5,912270045 | 2,048084198 | -2,886731927 | 0,003892657 | 0,049030393 | The Moose: suction sampler vs FISH | Function unknown |
| gene_42905 | 5,210087868 | -5,726244642 | 1,842491782 | -3,107880696 | 0,001884341 | 0,032041914 | The Moose: suction sampler vs FISH | Function unknown |
| gene_850075 | 5,073206706 | -5,700998255 | 1,828926897 | -3,117127461 | 0,001826226 | 0,031440967 | The Moose: suction sampler vs FISH | Function unknown |
| gene_1240684 | 6,353250864 | -5,310109206 | 1,608515857 | -3,301247659 | 0,000962559 | 0,021223395 | The Moose: suction sampler vs FISH | Function unknown |
| gene_1425974 | 21,81576811 | -4,753142178 | 1,145254808 | -4,150292271 | 3,32E-05 | 0,002372324 | The Moose: suction sampler vs FISH | Function unknown |
| gene_1229459 | 15,31869999 | -4,586213701 | 1,160884709 | -3,950619442 | 7,79E-05 | 0,004243868 | The Moose: suction sampler vs FISH | Function unknown |
| gene_1044493 | 13,29536744 | -4,102202503 | 1,328368655 | -3,088150633 | 0,002014064 | 0,033315968 | The Moose: suction sampler vs FISH | Function unknown |
| gene_937929 | 55,54815471 | -4,017090318 | 1,101167792 | -3,648026776 | 0,000264262 | 0,009376198 | The Moose: suction sampler vs FISH | Function unknown |
| gene_342524 | 24,27204904 | -3,825022619 | 0,924344752 | -4,138090914 | 3,50E-05 | 0,002454294 | The Moose: suction sampler vs FISH | Function unknown |
| gene_1878812 | 16,93060708 | -3,721667244 | 1,190988109 | -3,124856761 | 0,001778917 | 0,030879823 | The Moose: suction sampler vs FISH | Function unknown |
| gene_256665 | 217,6372149 | -3,671128075 | 0,821486307 | -4,468885292 | 7,86E-06 | 0,000907996 | The Moose: suction sampler vs FISH | Function unknown |
| gene_937933 | 438,7136135 | -3,006534697 | 0,932721489 | -3,223400267 | 0,001266784 | 0,02517969 | The Moose: suction sampler vs FISH | Function unknown |
| gene_24056 | 76,94637196 | -2,679864391 | 0,702854738 | -3,812828235 | 0,000137386 | 0,006145279 | The Moose: suction sampler vs FISH | Function unknown |
| gene_923668 | 48,46947358 | -2,586480291 | 0,738707857 | -3,501357492 | 0,000462894 | 0,013327042 | The Moose: suction sampler vs FISH | Function unknown |
| gene_1074197 | 645,1687847 | -2,308456754 | 0,584867154 | -3,946976227 | 7,91E-05 | 0,004267813 | The Moose: suction sampler vs FISH | Function unknown |
| gene_990698 | 44,20428241 | -2,3025075 | 0,754153853 | -3,053100492 | 0,002264901 | 0,035551536 | The Moose: suction sampler vs FISH | Function unknown |
| gene_133350 | 289,5705836 | -2,281105276 | 0,767448179 | -2,972324828 | 0,002955538 | 0,041752163 | The Moose: suction sampler vs FISH | Function unknown |
| gene_109465 | 56,49205285 | -2,212985356 | 0,736077205 | -3,006458211 | 0,002643104 | 0,03898959 | The Moose: suction sampler vs FISH | Function unknown |
| gene_266110 | 352,1746577 | -2,18249866 | 0,600352657 | -3,635361043 | 0,000277592 | 0,009687135 | The Moose: suction sampler vs FISH | Function unknown |
| gene_47221 | 32,78052571 | -2,133327038 | 0,69568901 | -3,066495241 | 0,002165843 | 0,034777068 | The Moose: suction sampler vs FISH | Function unknown |
| gene_280112 | 81,52409866 | -2,124685545 | 0,706666026 | -3,006633216 | 0,002641583 | 0,03898959 | The Moose: suction sampler vs FISH | Function unknown |
| gene_87656 | 52,42590486 | -2,057337786 | 0,663636647 | -3,100096711 | 0,001934575 | 0,032541813 | The Moose: suction sampler vs FISH | Function unknown |
| gene_467212 | 238,5640968 | -2,030552623 | 0,564863217 | -3,594768716 | 0,00032468 | 0,010695092 | The Moose: suction sampler vs FISH | Function unknown |
| gene_1331157 | 228,0261609 | 2,097054532 | 0,687115903 | 3,051966228 | 0,002273477 | 0,035649607 | The Moose: suction sampler vs FISH | Function unknown |
| gene_38378 | 43,29693397 | 2,190422668 | 0,750118602 | 2,920101784 | 0,003499171 | 0,046073252 | The Moose: suction sampler vs FISH | Function unknown |
| gene_516565 | 92,29317482 | 2,277176705 | 0,781032071 | 2,915599485 | 0,003550059 | 0,046443549 | The Moose: suction sampler vs FISH | Function unknown |
| gene_461836 | 44,19232984 | 2,288061001 | 0,771789804 | 2,964616776 | 0,003030602 | 0,042420919 | The Moose: suction sampler vs FISH | Function unknown |
| gene_1358134 | 188,8770938 | 2,326518846 | 0,74144841 | 3,137802732 | 0,001702194 | 0,030139256 | The Moose: suction sampler vs FISH | Function unknown |
| gene_1404350 | 43,94500419 | 2,346690476 | 0,754000509 | 3,112319486 | 0,001856235 | 0,03174722 | The Moose: suction sampler vs FISH | Function unknown |
| gene_1592592 | 26,07828032 | 2,380751394 | 0,817849085 | 2,910991084 | 0,003602843 | 0,046875253 | The Moose: suction sampler vs FISH | Function unknown |
| gene_325081 | 110,5095265 | 2,419458249 | 0,689321106 | 3,509914651 | 0,000448251 | 0,013040214 | The Moose: suction sampler vs FISH | Function unknown |
| gene_1690690 | 30,32757559 | 2,420207617 | 0,820067539 | 2,951229627 | 0,003165115 | 0,043486135 | The Moose: suction sampler vs FISH | Function unknown |
| gene_663221 | 40,47734145 | 2,431955489 | 0,749136236 | 3,246346086 | 0,001168806 | 0,023920816 | The Moose: suction sampler vs FISH | Function unknown |
| gene_756908 | 62,96351931 | 2,456791261 | 0,810443251 | 3,031416769 | 0,00243409 | 0,037135802 | The Moose: suction sampler vs FISH | Function unknown |
| gene_208722 | 65,96786766 | 2,47463875 | 0,775693231 | 3,190228623 | 0,001421603 | 0,02697923 | The Moose: suction sampler vs FISH | Function unknown |
| gene_530568 | 30,3636311 | 2,528079909 | 0,839245387 | 3,012325059 | 0,002592548 | 0,03865402 | The Moose: suction sampler vs FISH | Function unknown |
| gene_1237016 | 105,8091485 | 2,528474392 | 0,840721089 | 3,007506799 | 0,002634002 | 0,038947916 | The Moose: suction sampler vs FISH | Function unknown |
| gene_603310 | 23,92812138 | 2,532844254 | 0,813541155 | 3,113357249 | 0,00184972 | 0,031680238 | The Moose: suction sampler vs FISH | Function unknown |
| gene_462702 | 42,15207704 | 2,552792211 | 0,729022163 | 3,501666121 | 0,000462359 | 0,013322059 | The Moose: suction sampler vs FISH | Function unknown |
| gene_1583643 | 57,22813569 | 2,564932713 | 0,765307297 | 3,351506933 | 0,00080373 | 0,018965252 | The Moose: suction sampler vs FISH | Function unknown |
| gene_475243 | 193,6222217 | 2,574267759 | 0,818651143 | 3,144523501 | 0,001663575 | 0,029742284 | The Moose: suction sampler vs FISH | Function unknown |
| gene_1468622 | 23,16187725 | 2,589083545 | 0,769463914 | 3,364788781 | 0,000766023 | 0,01840749 | The Moose: suction sampler vs FISH | Function unknown |
| gene_941049 | 60,15034266 | 2,60982929 | 0,837119757 | 3,117629548 | 0,001823118 | 0,031394819 | The Moose: suction sampler vs FISH | Function unknown |
| gene_546296 | 57,54692848 | 2,611823848 | 0,822479339 | 3,175549493 | 0,00149553 | 0,027863827 | The Moose: suction sampler vs FISH | Function unknown |
| gene_265530 | 20,07167705 | 2,642816442 | 0,862018915 | 3,065845072 | 0,002170558 | 0,034799519 | The Moose: suction sampler vs FISH | Function unknown |
| gene_986010 | 28,62351123 | 2,664703006 | 0,848123723 | 3,141880051 | 0,001678668 | 0,029910181 | The Moose: suction sampler vs FISH | Function unknown |
| gene_1604761 | 36,62621154 | 2,672880446 | 0,785101772 | 3,404501865 | 0,000662848 | 0,016737807 | The Moose: suction sampler vs FISH | Function unknown |
| gene_458452 | 26,63192708 | 2,693434673 | 0,87957554 | 3,062198244 | 0,002197179 | 0,035034722 | The Moose: suction sampler vs FISH | Function unknown |
| gene_1325735 | 26,69081271 | 2,704685334 | 0,853013253 | 3,17074245 | 0,001520499 | 0,028136316 | The Moose: suction sampler vs FISH | Function unknown |
| gene_1284619 | 19,99296608 | 2,717248841 | 0,840184353 | 3,234110264 | 0,001220223 | 0,024539704 | The Moose: suction sampler vs FISH | Function unknown |

| | | | | | | | | |
|---|---|---|---|---|---|---|---|---|
| gene_1247462 | 59,95304135 | 2,718475553 | 0,771759587 | 3,522438334 | 0,000427597 | 0,012694431 | The Moose: suction sampler vs FISH | Function unknown |
| gene_766451 | 20,77946634 | 2,719848606 | 0,913184149 | 2,978422928 | 0,002897359 | 0,04124763 | The Moose: suction sampler vs FISH | Function unknown |
| gene_1228064 | 30,62529944 | 2,720408554 | 0,89330294 | 3,045337065 | 0,002324197 | 0,036082789 | The Moose: suction sampler vs FISH | Function unknown |
| gene_413882 | 22,78772339 | 2,749376331 | 0,935650621 | 2,93846471 | 0,003298422 | 0,044556301 | The Moose: suction sampler vs FISH | Function unknown |
| gene_253180 | 93,07864009 | 2,7531957 | 0,851392264 | 3,233757008 | 0,001221734 | 0,024539704 | The Moose: suction sampler vs FISH | Function unknown |
| gene_488235 | 46,79068467 | 2,779496686 | 0,822720176 | 3,378422902 | 0,000729029 | 0,017818141 | The Moose: suction sampler vs FISH | Function unknown |
| gene_1734391 | 50,6157766 | 2,785379622 | 0,742942648 | 3,749117956 | 0,000177458 | 0,007212714 | The Moose: suction sampler vs FISH | Function unknown |
| gene_349782 | 25,89852767 | 2,789504171 | 0,801754619 | 3,479249268 | 0,000502821 | 0,014018422 | The Moose: suction sampler vs FISH | Function unknown |
| gene_1399593 | 14,76242231 | 2,793535696 | 0,921614902 | 3,031131213 | 0,002436393 | 0,037163223 | The Moose: suction sampler vs FISH | Function unknown |
| gene_98670 | 38,22444656 | 2,812971292 | 0,808802228 | 3,477947012 | 0,00050527 | 0,014054379 | The Moose: suction sampler vs FISH | Function unknown |
| gene_1705591 | 19,44359997 | 2,814194642 | 0,904661992 | 3,110769178 | 0,001866007 | 0,031864273 | The Moose: suction sampler vs FISH | Function unknown |
| gene_1809001 | 24,08374258 | 2,815323897 | 0,941649092 | 2,989780292 | 0,002791782 | 0,040338857 | The Moose: suction sampler vs FISH | Function unknown |
| gene_676497 | 33,31341226 | 2,816966432 | 0,853153812 | 3,301827164 | 0,000960572 | 0,021202975 | The Moose: suction sampler vs FISH | Function unknown |
| gene_1273850 | 16,94598562 | 2,818250501 | 0,929849135 | 3,030868553 | 0,002438514 | 0,03717118 | The Moose: suction sampler vs FISH | Function unknown |
| gene_657636 | 62,73432132 | 2,830773572 | 0,70163953 | 4,034512667 | 5,47E-05 | 0,003323902 | The Moose: suction sampler vs FISH | Function unknown |
| gene_745710 | 78,59948211 | 2,832794197 | 0,703718192 | 4,025466767 | 5,69E-05 | 0,003434312 | The Moose: suction sampler vs FISH | Function unknown |
| gene_273221 | 298,7582186 | 2,835197744 | 0,852420071 | 3,326057001 | 0,000880839 | 0,020204007 | The Moose: suction sampler vs FISH | Function unknown |
| gene_628826 | 50,11675253 | 2,842218144 | 0,709190342 | 4,007694375 | 6,13E-05 | 0,003606207 | The Moose: suction sampler vs FISH | Function unknown |
| gene_1468621 | 12,79538491 | 2,84496423 | 0,925727516 | 3,073219908 | 0,002117624 | 0,034325508 | The Moose: suction sampler vs FISH | Function unknown |
| gene_528759 | 19,79767225 | 2,847741695 | 0,931912443 | 3,055803917 | 0,00224458 | 0,035391185 | The Moose: suction sampler vs FISH | Function unknown |
| gene_813153 | 39,30618461 | 2,856222527 | 0,920090462 | 3,104284464 | 0,001907398 | 0,032256247 | The Moose: suction sampler vs FISH | Function unknown |
| gene_1200090 | 89,81539837 | 2,874514166 | 0,713293759 | 4,029916328 | 5,58E-05 | 0,003386768 | The Moose: suction sampler vs FISH | Function unknown |
| gene_866104 | 34,50814035 | 2,883133188 | 0,873662046 | 3,300055439 | 0,000966657 | 0,021281818 | The Moose: suction sampler vs FISH | Function unknown |
| gene_341329 | 17,7335192 | 2,930761822 | 0,98294158 | 2,98162361 | 0,002867243 | 0,041011106 | The Moose: suction sampler vs FISH | Function unknown |
| gene_523145 | 45,347862 | 2,946234202 | 0,871472124 | 3,380755529 | 0,000722868 | 0,017704153 | The Moose: suction sampler vs FISH | Function unknown |
| gene_1592747 | 14,09822644 | 2,94936203 | 0,993376614 | 2,969027043 | 0,002987443 | 0,042033095 | The Moose: suction sampler vs FISH | Function unknown |
| gene_661371 | 242,2187596 | 2,964619398 | 0,761908972 | 3,89104146 | 9,98E-05 | 0,004993997 | The Moose: suction sampler vs FISH | Function unknown |
| gene_316712 | 10,39364222 | 2,966233103 | 1,02990107 | 2,880114595 | 0,003975307 | 0,049611339 | The Moose: suction sampler vs FISH | Function unknown |
| gene_33724 | 57,68077855 | 2,970517613 | 0,771120259 | 3,852210573 | 0,000117056 | 0,005535866 | The Moose: suction sampler vs FISH | Function unknown |
| gene_1136447 | 15,47583244 | 2,982467704 | 1,030889979 | 2,89309991 | 0,003814599 | 0,04847115 | The Moose: suction sampler vs FISH | Function unknown |
| gene_253179 | 63,75203489 | 2,984735928 | 0,980364333 | 3,044517051 | 0,002330542 | 0,03615688 | The Moose: suction sampler vs FISH | Function unknown |
| gene_399887 | 10,85285507 | 3,003416698 | 1,007604008 | 2,980751042 | 0,002875424 | 0,041078691 | The Moose: suction sampler vs FISH | Function unknown |
| gene_109264 | 16,64279318 | 3,0050388 | 0,894802278 | 3,358327167 | 0,000784157 | 0,018701511 | The Moose: suction sampler vs FISH | Function unknown |
| gene_973271 | 26,62929093 | 3,018714902 | 0,786441863 | 3,838446353 | 0,000123815 | 0,005765353 | The Moose: suction sampler vs FISH | Function unknown |
| gene_1687595 | 16,11782983 | 3,02278435 | 0,886655364 | 3,409198739 | 0,00065154 | 0,016535022 | The Moose: suction sampler vs FISH | Function unknown |
| gene_441124 | 21,96374362 | 3,039796192 | 0,948220331 | 3,20579099 | 0,001346918 | 0,026080775 | The Moose: suction sampler vs FISH | Function unknown |
| gene_1422351 | 26,69869489 | 3,053040765 | 0,989248267 | 3,08622301 | 0,002027167 | 0,033448147 | The Moose: suction sampler vs FISH | Function unknown |
| gene_253360 | 84,97761006 | 3,05450977 | 0,732621686 | 4,169286587 | 3,06E-05 | 0,002241904 | The Moose: suction sampler vs FISH | Function unknown |
| gene_564933 | 47,86182179 | 3,061613175 | 0,731824816 | 4,183532871 | 2,87E-05 | 0,002158827 | The Moose: suction sampler vs FISH | Function unknown |
| gene_45647 | 19,30547577 | 3,065664269 | 0,88328887 | 3,470738026 | 0,00051903 | 0,014277216 | The Moose: suction sampler vs FISH | Function unknown |
| gene_1389590 | 24,58329444 | 3,065701017 | 0,951878602 | 3,220684876 | 0,001278847 | 0,025289765 | The Moose: suction sampler vs FISH | Function unknown |
| gene_220495 | 18,37430603 | 3,067409219 | 0,963723029 | 3,182874255 | 0,001458209 | 0,027418889 | The Moose: suction sampler vs FISH | Function unknown |
| gene_1535110 | 28,05247105 | 3,068835822 | 0,813428371 | 3,772717956 | 0,000161479 | 0,006815983 | The Moose: suction sampler vs FISH | Function unknown |
| gene_370942 | 12,57408393 | 3,074720465 | 0,997454379 | 3,082567513 | 0,002052232 | 0,033717846 | The Moose: suction sampler vs FISH | Function unknown |
| gene_1018965 | 10,88379054 | 3,07758595 | 1,067528057 | 2,882908726 | 0,003940216 | 0,049366967 | The Moose: suction sampler vs FISH | Function unknown |
| gene_484762 | 181,5039569 | 3,083342953 | 0,742616105 | 4,152001191 | 3,30E-05 | 0,002358711 | The Moose: suction sampler vs FISH | Function unknown |
| gene_435067 | 15,25362519 | 3,09162406 | 1,013976349 | 3,049010032 | 0,002295968 | 0,03582481 | The Moose: suction sampler vs FISH | Function unknown |
| gene_211362 | 16,53617408 | 3,104323262 | 0,909985929 | 3,411396993 | 0,000646309 | 0,01644774 | The Moose: suction sampler vs FISH | Function unknown |
| gene_131905 | 16,27539476 | 3,12017055 | 0,992037405 | 3,145214621 | 0,00165965 | 0,029688834 | The Moose: suction sampler vs FISH | Function unknown |
| gene_1730475 | 19,725613 | 3,123908607 | 0,85260674 | 3,663950167 | 0,000248355 | 0,008995475 | The Moose: suction sampler vs FISH | Function unknown |
| gene_48047 | 25,86721278 | 3,127089003 | 0,962022363 | 3,250536706 | 0,001151874 | 0,023755291 | The Moose: suction sampler vs FISH | Function unknown |
| gene_106783 | 31,49810013 | 3,129290064 | 0,986558714 | 3,171924811 | 0,001514322 | 0,028058455 | The Moose: suction sampler vs FISH | Function unknown |
| gene_546804 | 21,44430971 | 3,131981258 | 0,834159801 | 3,754653791 | 0,000173581 | 0,007124112 | The Moose: suction sampler vs FISH | Function unknown |
| gene_1628716 | 17,57006923 | 3,13569761 | 1,089513547 | 2,878071246 | 0,004001148 | 0,049803793 | The Moose: suction sampler vs FISH | Function unknown |
| gene_357653 | 25,20339575 | 3,140818567 | 1,023289639 | 3,069334868 | 0,00214536 | 0,034584272 | The Moose: suction sampler vs FISH | Function unknown |
| gene_9997 | 14,9008713 | 3,174263799 | 1,078643934 | 2,942828213 | 0,003252288 | 0,044160858 | The Moose: suction sampler vs FISH | Function unknown |
| gene_634860 | 173,9904314 | 3,180319806 | 0,834521457 | 3,810950313 | 0,000138434 | 0,00616571 | The Moose: suction sampler vs FISH | Function unknown |
| gene_1401065 | 17,477825 | 3,197210447 | 1,009262723 | 3,167867369 | 0,001535616 | 0,028301954 | The Moose: suction sampler vs FISH | Function unknown |
| gene_186944 | 23,4882719 | 3,20096735 | 0,987360507 | 3,241943879 | 0,001187174 | 0,024172995 | The Moose: suction sampler vs FISH | Function unknown |
| gene_1347148 | 49,11936333 | 3,204599184 | 0,797282811 | 4,019400818 | 5,83E-05 | 0,003492459 | The Moose: suction sampler vs FISH | Function unknown |
| gene_673627 | 78,16839301 | 3,237643955 | 1,081611513 | 2,993351973 | 0,002759313 | 0,039995611 | The Moose: suction sampler vs FISH | Function unknown |
| gene_1736459 | 448,0281785 | 3,239950447 | 0,719086862 | 4,50564545 | 6,62E-06 | 0,000812703 | The Moose: suction sampler vs FISH | Function unknown |
| gene_611393 | 22,84379198 | 3,242709068 | 0,962173299 | 3,370192325 | 0,000751157 | 0,018201502 | The Moose: suction sampler vs FISH | Function unknown |
| gene_1627046 | 27,65043757 | 3,249086164 | 0,866891933 | 3,747971391 | 0,000178271 | 0,007226628 | The Moose: suction sampler vs FISH | Function unknown |
| gene_334771 | 118,1352852 | 3,259353376 | 0,818839897 | 3,980452573 | 6,88E-05 | 0,003886042 | The Moose: suction sampler vs FISH | Function unknown |

| | | | | | | | | |
|---|---|---|---|---|---|---|---|---|
| gene_1889194 | 39,61516513 | 3,269145997 | 0,97244593 | 3,361776624 | 0,000774427 | 0,018551149 | The Moose: suction sampler vs FISH | Function unknown |
| gene_615425 | 16,51881055 | 3,287663643 | 1,120917439 | 2,933011416 | 0,003356915 | 0,044966015 | The Moose: suction sampler vs FISH | Function unknown |
| gene_294811 | 92,88425801 | 3,294125705 | 0,794814137 | 4,144523292 | 3,41E-05 | 0,002418722 | The Moose: suction sampler vs FISH | Function unknown |
| gene_889516 | 209,7549543 | 3,341425676 | 0,750414304 | 4,452774501 | 8,48E-06 | 0,000959286 | The Moose: suction sampler vs FISH | Function unknown |
| gene_641834 | 11,32326788 | 3,342884943 | 1,043033138 | 3,204965231 | 0,001350788 | 0,026125951 | The Moose: suction sampler vs FISH | Function unknown |
| gene_923839 | 67,91390904 | 3,348147284 | 0,757598456 | 4,419421999 | 9,90E-06 | 0,001052955 | The Moose: suction sampler vs FISH | Function unknown |
| gene_28130 | 52,11448042 | 3,359562619 | 0,848044967 | 3,961538301 | 7,45E-05 | 0,004121578 | The Moose: suction sampler vs FISH | Function unknown |
| gene_1283315 | 11,13254893 | 3,361270088 | 1,15114586 | 2,919934133 | 0,003501054 | 0,046077151 | The Moose: suction sampler vs FISH | Function unknown |
| gene_1431254 | 13,65118969 | 3,362769261 | 1,153186306 | 2,916067632 | 0,003544737 | 0,046390444 | The Moose: suction sampler vs FISH | Function unknown |
| gene_1023633 | 21,83674178 | 3,381440012 | 0,903487945 | 3,742650944 | 0,000182089 | 0,007327963 | The Moose: suction sampler vs FISH | Function unknown |
| gene_818983 | 54,68214184 | 3,401268709 | 0,647874424 | 5,249888842 | 1,52E-07 | 6,04E-05 | The Moose: suction sampler vs FISH | Function unknown |
| gene_1109602 | 13,349512 | 3,403683999 | 1,159807313 | 2,934697826 | 0,003338727 | 0,044861465 | The Moose: suction sampler vs FISH | Function unknown |
| gene_1343562 | 32,96877195 | 3,406546319 | 1,022625985 | 3,331175198 | 0,000864802 | 0,019979664 | The Moose: suction sampler vs FISH | Function unknown |
| gene_379177 | 7,174807279 | 3,408930345 | 1,123508065 | 3,034184134 | 0,002411872 | 0,036919533 | The Moose: suction sampler vs FISH | Function unknown |
| gene_1818927 | 11,91850682 | 3,416368134 | 1,176013901 | 2,905040604 | 0,003672055 | 0,0474274 | The Moose: suction sampler vs FISH | Function unknown |
| gene_801346 | 52,53184978 | 3,436989273 | 0,96993701 | 3,543518019 | 0,000394826 | 0,012102938 | The Moose: suction sampler vs FISH | Function unknown |
| gene_479692 | 287,5502733 | 3,447423932 | 1,065024533 | 3,236943212 | 0,001208175 | 0,024431271 | The Moose: suction sampler vs FISH | Function unknown |
| gene_981883 | 31,18964905 | 3,451737243 | 0,969121415 | 3,561718058 | 0,000368436 | 0,0115492 | The Moose: suction sampler vs FISH | Function unknown |
| gene_1727798 | 15,33108108 | 3,465902418 | 1,155964291 | 2,998278099 | 0,002715098 | 0,039596976 | The Moose: suction sampler vs FISH | Function unknown |
| gene_424597 | 67,68808452 | 3,466857158 | 1,057480825 | 3,27841137 | 0,001043931 | 0,022379318 | The Moose: suction sampler vs FISH | Function unknown |
| gene_475783 | 46,37711767 | 3,505977107 | 1,160443535 | 3,021238865 | 0,002517427 | 0,037936859 | The Moose: suction sampler vs FISH | Function unknown |
| gene_921336 | 8,771949298 | 3,517399475 | 1,222987227 | 2,876072126 | 0,004026577 | 0,049980048 | The Moose: suction sampler vs FISH | Function unknown |
| gene_852898 | 11,93437133 | 3,523016497 | 1,202787347 | 2,929043531 | 0,003400068 | 0,045304426 | The Moose: suction sampler vs FISH | Function unknown |
| gene_91106 | 50,56005286 | 3,52502658 | 0,925895196 | 3,80715506 | 0,000140575 | 0,00623422 | The Moose: suction sampler vs FISH | Function unknown |
| gene_499245 | 21,61551823 | 3,535439907 | 0,870026815 | 4,063598781 | 4,83E-05 | 0,003090397 | The Moose: suction sampler vs FISH | Function unknown |
| gene_1187529 | 14,32094774 | 3,552052731 | 1,175217923 | 3,022463036 | 0,002507267 | 0,037874584 | The Moose: suction sampler vs FISH | Function unknown |
| gene_1256386 | 84,77772253 | 3,553096098 | 0,853163669 | 4,164612522 | 3,12E-05 | 0,002267936 | The Moose: suction sampler vs FISH | Function unknown |
| gene_645137 | 18,19318807 | 3,562325128 | 1,090750743 | 3,26593876 | 0,001091018 | 0,023025818 | The Moose: suction sampler vs FISH | Function unknown |
| gene_924236 | 8,757245776 | 3,585302767 | 1,123262431 | 3,191865647 | 0,001413571 | 0,026863397 | The Moose: suction sampler vs FISH | Function unknown |
| gene_471391 | 11,9127762 | 3,586595728 | 1,11517417 | 3,216175398 | 0,001299114 | 0,025546445 | The Moose: suction sampler vs FISH | Function unknown |
| gene_579621 | 23,31987647 | 3,591804391 | 0,923266067 | 3,890324271 | 0,00010011 | 0,004994976 | The Moose: suction sampler vs FISH | Function unknown |
| gene_1546968 | 14,97209988 | 3,599969177 | 1,206956789 | 2,982682735 | 0,00285734 | 0,040939788 | The Moose: suction sampler vs FISH | Function unknown |
| gene_18411 | 17,91982755 | 3,601609637 | 1,088719657 | 3,308114824 | 0,000939263 | 0,020932062 | The Moose: suction sampler vs FISH | Function unknown |
| gene_96699 | 14,81604423 | 3,607578247 | 1,168363939 | 3,087717899 | 0,002016998 | 0,033340635 | The Moose: suction sampler vs FISH | Function unknown |
| gene_788988 | 27,32958724 | 3,620926493 | 1,03244423 | 3,507140035 | 0,000452951 | 0,013131648 | The Moose: suction sampler vs FISH | Function unknown |
| gene_1266955 | 17,59493425 | 3,627799395 | 1,053128406 | 3,444783538 | 0,000571517 | 0,015224903 | The Moose: suction sampler vs FISH | Function unknown |
| gene_1305939 | 12,77400143 | 3,629578512 | 1,198612778 | 3,028149356 | 0,002460564 | 0,037384393 | The Moose: suction sampler vs FISH | Function unknown |
| gene_1599013 | 21,14288625 | 3,638337964 | 0,989820194 | 3,675756453 | 0,000237146 | 0,008752344 | The Moose: suction sampler vs FISH | Function unknown |
| gene_645136 | 29,33728641 | 3,666181967 | 0,862479547 | 4,250746559 | 2,13E-05 | 0,001738681 | The Moose: suction sampler vs FISH | Function unknown |
| gene_522505 | 18,65047782 | 3,674630825 | 1,156217597 | 3,178148158 | 0,00148219 | 0,027699602 | The Moose: suction sampler vs FISH | Function unknown |
| gene_824904 | 62,47475513 | 3,675437301 | 0,773681095 | 4,750584351 | 2,03E-06 | 0,000350538 | The Moose: suction sampler vs FISH | Function unknown |
| gene_300461 | 40,64364894 | 3,676976171 | 1,142575052 | 3,218148485 | 0,00129021 | 0,025452452 | The Moose: suction sampler vs FISH | Function unknown |
| gene_575887 | 32,09392584 | 3,69208669 | 0,880395005 | 4,193670648 | 2,74E-05 | 0,002095522 | The Moose: suction sampler vs FISH | Function unknown |
| gene_1238287 | 22,62774054 | 3,697445084 | 0,996352606 | 3,710980491 | 0,000206458 | 0,007984892 | The Moose: suction sampler vs FISH | Function unknown |
| gene_132781 | 10,60787028 | 3,70354834 | 1,177994206 | 3,143944445 | 0,001666871 | 0,029770449 | The Moose: suction sampler vs FISH | Function unknown |
| gene_1433123 | 7,500046245 | 3,714023674 | 1,232249198 | 3,014019955 | 0,002578108 | 0,03851692 | The Moose: suction sampler vs FISH | Function unknown |
| gene_884016 | 10,23934519 | 3,721595071 | 1,259731531 | 2,95427635 | 0,003134032 | 0,043275647 | The Moose: suction sampler vs FISH | Function unknown |
| gene_1456703 | 26,29588798 | 3,742330576 | 0,978761564 | 3,823536512 | 0,000131551 | 0,005963296 | The Moose: suction sampler vs FISH | Function unknown |
| gene_32209 | 11,40090314 | 3,761022061 | 1,162267888 | 3,235933901 | 0,001212455 | 0,024471718 | The Moose: suction sampler vs FISH | Function unknown |
| gene_702909 | 21,00019759 | 3,815711698 | 0,983152535 | 3,881098368 | 0,000103986 | 0,005111938 | The Moose: suction sampler vs FISH | Function unknown |
| gene_1640829 | 33,43454345 | 3,825734001 | 1,026243297 | 3,727901572 | 0,000193081 | 0,007644645 | The Moose: suction sampler vs FISH | Function unknown |
| gene_677844 | 47,91292283 | 3,826116947 | 0,940574074 | 4,067852873 | 4,74E-05 | 0,003059563 | The Moose: suction sampler vs FISH | Function unknown |
| gene_1099913 | 35,54061238 | 3,860394408 | 0,975860331 | 3,95588824 | 7,63E-05 | 0,004186299 | The Moose: suction sampler vs FISH | Function unknown |
| gene_791579 | 62,68127375 | 3,861853092 | 0,836261003 | 4,617999739 | 3,87E-06 | 0,000552304 | The Moose: suction sampler vs FISH | Function unknown |
| gene_1374272 | 17,9963521 | 3,866911839 | 1,037482678 | 3,727206172 | 0,000193614 | 0,007660085 | The Moose: suction sampler vs FISH | Function unknown |
| gene_875969 | 18,26055083 | 3,874087585 | 1,10141926 | 3,517359579 | 0,000435863 | 0,012851044 | The Moose: suction sampler vs FISH | Function unknown |
| gene_968467 | 21,01362613 | 3,876014456 | 1,061209768 | 3,652448907 | 0,000259751 | 0,00926989 | The Moose: suction sampler vs FISH | Function unknown |
| gene_494063 | 11,26226981 | 3,88628491 | 1,157764284 | 3,356715148 | 0,000788743 | 0,01874733 | The Moose: suction sampler vs FISH | Function unknown |
| gene_1664903 | 9,005259041 | 3,89937268 | 1,274607696 | 3,05927282 | 0,00221875 | 0,03520092 | The Moose: suction sampler vs FISH | Function unknown |
| gene_764175 | 25,83105178 | 3,900383294 | 0,896061603 | 4,352807085 | 1,34E-05 | 0,001295729 | The Moose: suction sampler vs FISH | Function unknown |
| gene_1918317 | 63,29749403 | 3,901730668 | 0,908629082 | 4,294085172 | 1,75E-05 | 0,001530091 | The Moose: suction sampler vs FISH | Function unknown |
| gene_413081 | 537,3366262 | 3,910556426 | 0,819508109 | 4,77183372 | 1,83E-06 | 0,00032805 | The Moose: suction sampler vs FISH | Function unknown |
| gene_834900 | 8,622856768 | 3,918128241 | 1,307690127 | 2,99622071 | 0,002733485 | 0,039760916 | The Moose: suction sampler vs FISH | Function unknown |
| gene_1427765 | 7,270247989 | 3,927741974 | 1,303597403 | 3,0130023 | 0,00258677 | 0,038591365 | The Moose: suction sampler vs FISH | Function unknown |
| gene_1715049 | 11,40573308 | 3,929941064 | 1,256828522 | 3,126871324 | 0,001766772 | 0,030792738 | The Moose: suction sampler vs FISH | Function unknown |

| | | | | | | | | |
|---|---|---|---|---|---|---|---|---|
| gene_959923 | 11,57836272 | 3,947670821 | 1,31147389 | 3,010102489 | 0,002611595 | 0,0388198 | The Moose: suction sampler vs FISH | Function unknown |
| gene_733944 | 291,4946095 | 3,985264719 | 0,762239473 | 5,228363084 | 1,71E-07 | 6,47E-05 | The Moose: suction sampler vs FISH | Function unknown |
| gene_1237019 | 12,12192998 | 3,989754059 | 1,206042547 | 3,308137072 | 0,000939188 | 0,020932062 | The Moose: suction sampler vs FISH | Function unknown |
| gene_1700565 | 12,12737554 | 3,995125504 | 1,249053624 | 3,198522006 | 0,00138134 | 0,026488906 | The Moose: suction sampler vs FISH | Function unknown |
| gene_1680038 | 11,16231297 | 3,996941251 | 1,227718215 | 3,2555852 | 0,001131589 | 0,023505612 | The Moose: suction sampler vs FISH | Function unknown |
| gene_1351396 | 8,105289521 | 4,03434362 | 1,338117136 | 3,014940554 | 0,002570296 | 0,038439302 | The Moose: suction sampler vs FISH | Function unknown |
| gene_524249 | 38,61420371 | 4,043244538 | 0,846762421 | 4,77494565 | 1,80E-06 | 0,000325979 | The Moose: suction sampler vs FISH | Function unknown |
| gene_601066 | 18,28285884 | 4,048151261 | 1,105478655 | 3,661899071 | 0,000250352 | 0,009044338 | The Moose: suction sampler vs FISH | Function unknown |
| gene_149406 | 13,89432907 | 4,080316193 | 1,146261195 | 3,559674017 | 0,000371315 | 0,011606974 | The Moose: suction sampler vs FISH | Function unknown |
| gene_908589 | 12,12884262 | 4,08103608 | 1,082513692 | 3,769962552 | 0,000163272 | 0,006860135 | The Moose: suction sampler vs FISH | Function unknown |
| gene_1816925 | 19,26351603 | 4,096499797 | 1,031578375 | 3,971098945 | 7,15E-05 | 0,004004872 | The Moose: suction sampler vs FISH | Function unknown |
| gene_1101109 | 43,77505691 | 4,128296306 | 0,971993632 | 4,247246249 | 2,16E-05 | 0,001758245 | The Moose: suction sampler vs FISH | Function unknown |
| gene_481156 | 9,357287926 | 4,170934189 | 1,290847517 | 3,231159476 | 0,001232891 | 0,024687463 | The Moose: suction sampler vs FISH | Function unknown |
| gene_741610 | 15,6286628 | 4,191422846 | 1,194896516 | 3,507770581 | 0,000451879 | 0,013116056 | The Moose: suction sampler vs FISH | Function unknown |
| gene_934815 | 10,35232461 | 4,191684241 | 1,18467171 | 3,538266513 | 0,000402763 | 0,012232174 | The Moose: suction sampler vs FISH | Function unknown |
| gene_966546 | 14,52602489 | 4,214117545 | 1,190190076 | 3,540709698 | 0,000399052 | 0,012196229 | The Moose: suction sampler vs FISH | Function unknown |
| gene_1737052 | 24,13184573 | 4,231001506 | 1,072873498 | 3,943616384 | 8,03E-05 | 0,004302805 | The Moose: suction sampler vs FISH | Function unknown |
| gene_889358 | 6,996678136 | 4,258635577 | 1,472531827 | 2,892049938 | 0,003827371 | 0,048578262 | The Moose: suction sampler vs FISH | Function unknown |
| gene_1544565 | 9,284653486 | 4,29555508 | 1,389359666 | 3,09175168 | 0,001989792 | 0,033077161 | The Moose: suction sampler vs FISH | Function unknown |
| gene_1296031 | 37,74945006 | 4,299695904 | 1,0334625 | 4,160475975 | 3,18E-05 | 0,002293513 | The Moose: suction sampler vs FISH | Function unknown |
| gene_1708893 | 9,294645699 | 4,300133955 | 1,229501125 | 3,497462401 | 0,000469707 | 0,013397136 | The Moose: suction sampler vs FISH | Function unknown |
| gene_1782205 | 13,82809193 | 4,302739824 | 1,263352882 | 3,405809956 | 0,000659681 | 0,01668972 | The Moose: suction sampler vs FISH | Function unknown |
| gene_398510 | 6,647364446 | 4,327719186 | 1,424553922 | 3,03794691 | 0,002381959 | 0,03660661 | The Moose: suction sampler vs FISH | Function unknown |
| gene_999240 | 20,80607088 | 4,331795547 | 1,046285743 | 4,140164936 | 3,47E-05 | 0,002448563 | The Moose: suction sampler vs FISH | Function unknown |
| gene_1057510 | 15,86085307 | 4,332578715 | 1,109288287 | 3,905728353 | 9,39E-05 | 0,004811416 | The Moose: suction sampler vs FISH | Function unknown |
| gene_1238328 | 6,84149923 | 4,337448586 | 1,297830084 | 3,342077393 | 0,000831539 | 0,019443604 | The Moose: suction sampler vs FISH | Function unknown |
| gene_135919 | 7,413857289 | 4,34471543 | 1,458071651 | 2,979768124 | 0,002884667 | 0,041138705 | The Moose: suction sampler vs FISH | Function unknown |
| gene_1428700 | 7,879934101 | 4,353855304 | 1,313136373 | 3,315615493 | 0,000914415 | 0,020610638 | The Moose: suction sampler vs FISH | Function unknown |
| gene_1018914 | 6,889503461 | 4,372904915 | 1,341215124 | 3,260405314 | 0,001112531 | 0,023266414 | The Moose: suction sampler vs FISH | Function unknown |
| gene_824629 | 13,87622553 | 4,376359423 | 1,212885597 | 3,608221118 | 0,000308304 | 0,010363096 | The Moose: suction sampler vs FISH | Function unknown |
| gene_1427769 | 7,703244063 | 4,394527695 | 1,445887551 | 3,039328815 | 0,002371059 | 0,036509811 | The Moose: suction sampler vs FISH | Function unknown |
| gene_1779229 | 7,678887194 | 4,397251206 | 1,478700231 | 2,973727273 | 0,002942064 | 0,041640225 | The Moose: suction sampler vs FISH | Function unknown |
| gene_243719 | 6,392500692 | 4,418553404 | 1,401914743 | 3,151798942 | 0,00162268 | 0,029281992 | The Moose: suction sampler vs FISH | Function unknown |
| gene_977080 | 9,20675617 | 4,473763915 | 1,343745538 | 3,329323736 | 0,000870571 | 0,020049897 | The Moose: suction sampler vs FISH | Function unknown |
| gene_1639859 | 8,707605122 | 4,474759463 | 1,444260198 | 3,098305603 | 0,001946306 | 0,03263058 | The Moose: suction sampler vs FISH | Function unknown |
| gene_135896 | 14,00311686 | 4,480969834 | 1,356094252 | 3,304320351 | 0,00095207 | 0,021103066 | The Moose: suction sampler vs FISH | Function unknown |
| gene_133683 | 5,900923887 | 4,49130446 | 1,522730022 | 2,94950805 | 0,003182803 | 0,043652839 | The Moose: suction sampler vs FISH | Function unknown |
| gene_787762 | 57,51090839 | 4,499502126 | 1,040385173 | 4,32484261 | 1,53E-05 | 0,001390906 | The Moose: suction sampler vs FISH | Function unknown |
| gene_402673 | 27,88457854 | 4,50316283 | 0,859814666 | 5,237364525 | 1,63E-07 | 6,30E-05 | The Moose: suction sampler vs FISH | Function unknown |
| gene_1432161 | 31,64687016 | 4,504410417 | 0,89670657 | 5,023282495 | 5,08E-07 | 0,000142073 | The Moose: suction sampler vs FISH | Function unknown |
| gene_1067009 | 15,16807041 | 4,525362973 | 1,275520123 | 3,547856982 | 0,000388379 | 0,011964974 | The Moose: suction sampler vs FISH | Function unknown |
| gene_790171 | 74,42021583 | 4,539231415 | 0,97985325 | 4,632562492 | 3,61E-06 | 0,000522164 | The Moose: suction sampler vs FISH | Function unknown |
| gene_739548 | 37,12960303 | 4,5600502 | 0,961278221 | 4,743736101 | 2,10E-06 | 0,000359199 | The Moose: suction sampler vs FISH | Function unknown |
| gene_1843429 | 7,384812479 | 4,560615904 | 1,561963787 | 2,919796183 | 0,003502604 | 0,046077151 | The Moose: suction sampler vs FISH | Function unknown |
| gene_1853676 | 15,88241321 | 4,565114397 | 1,127883138 | 4,047506559 | 5,18E-05 | 0,003223638 | The Moose: suction sampler vs FISH | Function unknown |
| gene_1778105 | 5,954499508 | 4,572492135 | 1,521099157 | 3,00604475 | 0,0026467 | 0,039033517 | The Moose: suction sampler vs FISH | Function unknown |
| gene_1606366 | 18,69740964 | 4,576877892 | 1,022081481 | 4,477997083 | 7,53E-06 | 0,000886832 | The Moose: suction sampler vs FISH | Function unknown |
| gene_1321691 | 11,96088095 | 4,581011228 | 1,297932746 | 3,529467335 | 0,000416397 | 0,012487662 | The Moose: suction sampler vs FISH | Function unknown |
| gene_1716556 | 12,6485365 | 4,60533975 | 1,366189576 | 3,370937555 | 0,000749128 | 0,018164322 | The Moose: suction sampler vs FISH | Function unknown |
| gene_1930397 | 130,6403047 | 4,616971164 | 1,026382954 | 4,498292912 | 6,85E-06 | 0,00083296 | The Moose: suction sampler vs FISH | Function unknown |
| gene_968468 | 148,6933536 | 4,620117652 | 0,804413738 | 5,743459412 | 9,28E-09 | 8,85E-06 | The Moose: suction sampler vs FISH | Function unknown |
| gene_968466 | 439,6307537 | 4,642631053 | 0,871122487 | 5,329481353 | 9,85E-08 | 4,64E-05 | The Moose: suction sampler vs FISH | Function unknown |
| gene_1192042 | 29,0832049 | 4,665892217 | 1,30283077 | 3,581349415 | 0,000341824 | 0,011035285 | The Moose: suction sampler vs FISH | Function unknown |
| gene_783573 | 65,88513879 | 4,670912309 | 0,856646298 | 5,452556465 | 4,97E-08 | 2,96E-05 | The Moose: suction sampler vs FISH | Function unknown |
| gene_1446328 | 4,661237171 | 4,699535326 | 1,602406823 | 2,932797875 | 0,003359225 | 0,044974217 | The Moose: suction sampler vs FISH | Function unknown |
| gene_916551 | 14,45962611 | 4,762888412 | 1,298702132 | 3,667421725 | 0,000245009 | 0,008930348 | The Moose: suction sampler vs FISH | Function unknown |
| gene_1140428 | 28,05571574 | 4,790423206 | 0,995988904 | 4,809715436 | 1,51E-06 | 0,000292899 | The Moose: suction sampler vs FISH | Function unknown |
| gene_1608547 | 35,16240832 | 4,800352672 | 1,202015504 | 3,993586319 | 6,51E-05 | 0,003740709 | The Moose: suction sampler vs FISH | Function unknown |
| gene_1309995 | 4,65383602 | 4,801624302 | 1,636193468 | 2,93463114 | 0,003339444 | 0,04486178 | The Moose: suction sampler vs FISH | Function unknown |
| gene_883736 | 17,91316832 | 4,822441741 | 1,579411254 | 3,053316056 | 0,002263274 | 0,035551536 | The Moose: suction sampler vs FISH | Function unknown |
| gene_430724 | 7,310097507 | 4,833360248 | 1,484977943 | 3,25483639 | 0,001134577 | 0,023545921 | The Moose: suction sampler vs FISH | Function unknown |
| gene_1879627 | 51,73454566 | 4,837960496 | 0,845833826 | 5,719752922 | 1,07E-08 | 9,48E-06 | The Moose: suction sampler vs FISH | Function unknown |
| gene_1805394 | 24,50086345 | 4,864840767 | 1,300029726 | 3,742099637 | 0,000182489 | 0,007340041 | The Moose: suction sampler vs FISH | Function unknown |
| gene_1192803 | 6,014714634 | 4,889388684 | 1,507093619 | 3,244250139 | 0,001177603 | 0,024038351 | The Moose: suction sampler vs FISH | Function unknown |
| gene_1666549 | 15,76582952 | 4,889506107 | 1,116935993 | 4,377606362 | 1,20E-05 | 0,001194126 | The Moose: suction sampler vs FISH | Function unknown |

| | | | | | | | | |
|---|---|---|---|---|---|---|---|---|
| gene_417198 | 20,87958016 | 4,915831399 | 1,082923465 | 4,539407962 | 5,64E-06 | 0,000716304 | The Moose: suction sampler vs FISH | Function unknown |
| gene_103622 | 8,708382009 | 4,95355649 | 1,713671155 | 2,890610883 | 0,003844939 | 0,04871339 | The Moose: suction sampler vs FISH | Function unknown |
| gene_1879763 | 13,53791489 | 4,990169623 | 1,385090385 | 3,602775442 | 0,000314837 | 0,010520127 | The Moose: suction sampler vs FISH | Function unknown |
| gene_1798391 | 5,31479511 | 5,018513537 | 1,631665411 | 3,075700143 | 0,002100089 | 0,034184632 | The Moose: suction sampler vs FISH | Function unknown |
| gene_1025882 | 25,563238 | 5,023855097 | 1,00970196 | 4,975582197 | 6,51E-07 | 0,000164687 | The Moose: suction sampler vs FISH | Function unknown |
| gene_1200787 | 14,5409007 | 5,040283006 | 1,349618955 | 3,73459708 | 0,000188016 | 0,0075048 | The Moose: suction sampler vs FISH | Function unknown |
| gene_1104137 | 35,94457283 | 5,055318613 | 1,039766105 | 4,861976738 | 1,16E-06 | 0,000245986 | The Moose: suction sampler vs FISH | Function unknown |
| gene_1633413 | 20,73889523 | 5,097287787 | 1,036586519 | 4,91737804 | 8,77E-07 | 0,000203859 | The Moose: suction sampler vs FISH | Function unknown |
| gene_1930396 | 89,68465519 | 5,118152005 | 0,992251022 | 5,15812218 | 2,49E-07 | 8,66E-05 | The Moose: suction sampler vs FISH | Function unknown |
| gene_686938 | 6,449800763 | 5,127108419 | 1,558271083 | 3,290254484 | 0,001000968 | 0,021744748 | The Moose: suction sampler vs FISH | Function unknown |
| gene_1263768 | 9,37949394 | 5,14343367 | 1,577494593 | 3,260507956 | 0,001112128 | 0,023266414 | The Moose: suction sampler vs FISH | Function unknown |
| gene_1453116 | 8,060548514 | 5,187505728 | 1,765430735 | 2,93837964 | 0,003299327 | 0,044559979 | The Moose: suction sampler vs FISH | Function unknown |
| gene_1911673 | 5,943891216 | 5,242454512 | 1,771256314 | 2,95973794 | 0,003079008 | 0,042855864 | The Moose: suction sampler vs FISH | Function unknown |
| gene_951565 | 4,749795433 | 5,298743145 | 1,788401052 | 2,962838307 | 0,003048166 | 0,042546401 | The Moose: suction sampler vs FISH | Function unknown |
| gene_1266257 | 7,305105907 | 5,304829762 | 1,5060068 | 3,522447416 | 0,000427582 | 0,012694431 | The Moose: suction sampler vs FISH | Function unknown |
| gene_1017301 | 12,7477754 | 5,314067433 | 1,342928927 | 3,957072728 | 7,59E-05 | 0,004177327 | The Moose: suction sampler vs FISH | Function unknown |
| gene_1516256 | 6,268820451 | 5,325023484 | 1,742144365 | 3,056591401 | 0,002238692 | 0,035360455 | The Moose: suction sampler vs FISH | Function unknown |
| gene_515675 | 10,21819024 | 5,353625647 | 1,406360804 | 3,806722735 | 0,00014082 | 0,006234468 | The Moose: suction sampler vs FISH | Function unknown |
| gene_809328 | 6,615739797 | 5,368187873 | 1,511601227 | 3,551325427 | 0,000383296 | 0,011875719 | The Moose: suction sampler vs FISH | Function unknown |
| gene_1059200 | 5,501434003 | 5,379951022 | 1,771209191 | 3,037445293 | 0,002385927 | 0,036636926 | The Moose: suction sampler vs FISH | Function unknown |
| gene_1632917 | 6,64142812 | 5,389310222 | 1,623471822 | 3,319620426 | 0,000901399 | 0,02049231 | The Moose: suction sampler vs FISH | Function unknown |
| gene_213491 | 16,46677281 | 5,417013177 | 1,605554165 | 3,373921165 | 0,000741056 | 0,01803053 | The Moose: suction sampler vs FISH | Function unknown |
| gene_875864 | 4,509522184 | 5,47132711 | 1,902203478 | 2,876310117 | 0,004023542 | 0,049950818 | The Moose: suction sampler vs FISH | Function unknown |
| gene_271410 | 16,50527613 | 5,475447362 | 1,305484307 | 4,194188572 | 2,74E-05 | 0,002092916 | The Moose: suction sampler vs FISH | Function unknown |
| gene_127456 | 4,388953292 | 5,479195765 | 1,793466779 | 3,055086288 | 0,002249958 | 0,035430564 | The Moose: suction sampler vs FISH | Function unknown |
| gene_248290 | 5,39467809 | 5,516500811 | 1,678595355 | 3,286379172 | 0,001014843 | 0,021909214 | The Moose: suction sampler vs FISH | Function unknown |
| gene_473252 | 7,254781571 | 5,518871653 | 1,651994922 | 3,340731608 | 0,00083558 | 0,019519447 | The Moose: suction sampler vs FISH | Function unknown |
| gene_673368 | 9,466838382 | 5,55891276 | 1,645978476 | 3,377269412 | 0,000732093 | 0,017875191 | The Moose: suction sampler vs FISH | Function unknown |
| gene_448266 | 14,39031536 | 5,582293215 | 1,222267815 | 4,567160442 | 4,94E-06 | 0,000655485 | The Moose: suction sampler vs FISH | Function unknown |
| gene_278240 | 8,818488291 | 5,594739186 | 1,735245993 | 3,224176404 | 0,001263355 | 0,025118339 | The Moose: suction sampler vs FISH | Function unknown |
| gene_1174571 | 6,170221042 | 5,598118678 | 1,655869084 | 3,380773716 | 0,00072282 | 0,017704153 | The Moose: suction sampler vs FISH | Function unknown |
| gene_1321106 | 44,969172 | 5,647597706 | 0,930855456 | 6,067104907 | 1,30E-09 | 2,17E-06 | The Moose: suction sampler vs FISH | Function unknown |
| gene_377420 | 7,296978323 | 5,661266243 | 1,818032149 | 3,113952769 | 0,00184599 | 0,031647704 | The Moose: suction sampler vs FISH | Function unknown |
| gene_1387171 | 22,75139786 | 5,676696543 | 1,271889653 | 4,463198933 | 8,07E-06 | 0,000926612 | The Moose: suction sampler vs FISH | Function unknown |
| gene_1041255 | 5,56859089 | 5,682118624 | 1,689102065 | 3,363987731 | 0,00076825 | 0,018445273 | The Moose: suction sampler vs FISH | Function unknown |
| gene_1716557 | 11,77909956 | 5,703138394 | 1,459562119 | 3,907431085 | 9,33E-05 | 0,004790289 | The Moose: suction sampler vs FISH | Function unknown |
| gene_703667 | 87,89236722 | 5,719821958 | 1,054992181 | 5,421672371 | 5,90E-08 | 3,35E-05 | The Moose: suction sampler vs FISH | Function unknown |
| gene_194525 | 5,093572118 | 5,732862129 | 1,985041307 | 2,888031654 | 0,003876609 | 0,048920536 | The Moose: suction sampler vs FISH | Function unknown |
| gene_1247932 | 17,66237284 | 5,79815519 | 1,309699988 | 4,427086541 | 9,55E-06 | 0,001029043 | The Moose: suction sampler vs FISH | Function unknown |
| gene_1245891 | 6,78746639 | 5,799036821 | 1,651832225 | 3,510669385 | 0,00044698 | 0,013021992 | The Moose: suction sampler vs FISH | Function unknown |
| gene_1942686 | 5,280786524 | 5,816709895 | 1,92525963 | 3,021259993 | 0,002517251 | 0,037936859 | The Moose: suction sampler vs FISH | Function unknown |
| gene_358703 | 7,537422841 | 5,83802766 | 1,559847982 | 3,742690139 | 0,000182061 | 0,007327963 | The Moose: suction sampler vs FISH | Function unknown |
| gene_1574722 | 4,438745142 | 5,841046523 | 1,739571693 | 3,357749811 | 0,000785797 | 0,018701511 | The Moose: suction sampler vs FISH | Function unknown |
| gene_1631415 | 3,671216654 | 5,852991141 | 1,868392787 | 3,132634198 | 0,001732452 | 0,030440178 | The Moose: suction sampler vs FISH | Function unknown |
| gene_1754388 | 12,78360973 | 5,853802816 | 1,43192403 | 4,088068007 | 4,35E-05 | 0,002871034 | The Moose: suction sampler vs FISH | Function unknown |
| gene_1768561 | 3,573362639 | 5,871726075 | 1,972267195 | 2,977145333 | 0,002909461 | 0,041340185 | The Moose: suction sampler vs FISH | Function unknown |
| gene_750069 | 17,8373675 | 5,889754008 | 1,312588293 | 4,487129773 | 7,22E-06 | 0,000862102 | The Moose: suction sampler vs FISH | Function unknown |
| gene_1905699 | 5,283533951 | 5,916571497 | 2,03875049 | 2,90205767 | 0,003707203 | 0,047759255 | The Moose: suction sampler vs FISH | Function unknown |
| gene_1651841 | 3,349976554 | 5,931995926 | 2,027144861 | 2,926281214 | 0,003430407 | 0,045535325 | The Moose: suction sampler vs FISH | Function unknown |
| gene_136101 | 19,18033852 | 5,933192745 | 1,478088226 | 4,014099185 | 5,97E-05 | 0,003539821 | The Moose: suction sampler vs FISH | Function unknown |
| gene_1049799 | 3,4172879 | 5,94929162 | 1,932720209 | 3,078196002 | 0,002082579 | 0,034042957 | The Moose: suction sampler vs FISH | Function unknown |
| gene_506546 | 4,234637132 | 5,970143926 | 1,760169871 | 3,391799863 | 0,000694351 | 0,017217113 | The Moose: suction sampler vs FISH | Function unknown |
| gene_1661256 | 2,719511081 | 5,994525682 | 2,061603886 | 2,90770003 | 0,003640974 | 0,047187461 | The Moose: suction sampler vs FISH | Function unknown |
| gene_1596726 | 8,774947023 | 6,011954824 | 1,429961373 | 4,204277778 | 2,62E-05 | 0,002024895 | The Moose: suction sampler vs FISH | Function unknown |
| gene_200784 | 4,289142352 | 6,024867876 | 1,919536473 | 3,138709768 | 0,001696934 | 0,030082389 | The Moose: suction sampler vs FISH | Function unknown |
| gene_1040288 | 4,184236501 | 6,045650424 | 1,874038962 | 3,226000392 | 0,001255332 | 0,024995522 | The Moose: suction sampler vs FISH | Function unknown |
| gene_362467 | 5,717281397 | 6,066178435 | 1,925292452 | 3,150782849 | 0,001628335 | 0,029347992 | The Moose: suction sampler vs FISH | Function unknown |
| gene_1567694 | 3,477654374 | 6,070842362 | 2,062638003 | 2,943241787 | 0,003247946 | 0,044160858 | The Moose: suction sampler vs FISH | Function unknown |
| gene_1717764 | 5,621446787 | 6,089266272 | 1,819325531 | 3,346991051 | 0,000816938 | 0,019200015 | The Moose: suction sampler vs FISH | Function unknown |
| gene_124276 | 8,874891727 | 6,131800132 | 1,7390903 | 3,525866444 | 0,0004221 | 0,012602081 | The Moose: suction sampler vs FISH | Function unknown |
| gene_545687 | 3,256512873 | 6,132728845 | 1,933150601 | 3,172400972 | 0,001511841 | 0,028039689 | The Moose: suction sampler vs FISH | Function unknown |
| gene_119624 | 114,202562 | 6,137710425 | 2,022904511 | 3,034107834 | 0,002412482 | 0,036921177 | The Moose: suction sampler vs FISH | Function unknown |
| gene_729801 | 4,498042782 | 6,140286161 | 1,991518919 | 3,083217589 | 0,002047754 | 0,033673581 | The Moose: suction sampler vs FISH | Function unknown |
| gene_661027 | 11,2065276 | 6,141612441 | 1,433564036 | 4,284156332 | 1,83E-05 | 0,001566547 | The Moose: suction sampler vs FISH | Function unknown |
| gene_193455 | 6,935193415 | 6,230062848 | 2,076484477 | 3,000293485 | 0,002697196 | 0,03943505 | The Moose: suction sampler vs FISH | Function unknown |

| | | | | | | | | |
|---|---|---|---|---|---|---|---|---|
| gene_1930770 | 6,411088923 | 6,239674443 | 1,871933812 | 3,333277279 | 0,000858294 | 0,019873068 | The Moose: suction sampler vs FISH | Function unknown |
| gene_1213188 | 5,746259245 | 6,240169488 | 1,76366196 | 3,538189081 | 0,000402881 | 0,012232174 | The Moose: suction sampler vs FISH | Function unknown |
| gene_588947 | 3,152262263 | 6,242339117 | 2,134581543 | 2,924385408 | 0,003451371 | 0,045700744 | The Moose: suction sampler vs FISH | Function unknown |
| gene_1944607 | 4,41991438 | 6,254541354 | 1,927024376 | 3,245699137 | 0,001171626 | 0,023962702 | The Moose: suction sampler vs FISH | Function unknown |
| gene_1355836 | 4,174730301 | 6,268820639 | 2,125224571 | 2,949721514 | 0,003180605 | 0,043638991 | The Moose: suction sampler vs FISH | Function unknown |
| gene_1585203 | 9,777976835 | 6,288145059 | 1,744304608 | 3,604958118 | 0,000312203 | 0,010457499 | The Moose: suction sampler vs FISH | Function unknown |
| gene_865682 | 7,050567971 | 6,309227709 | 1,700828778 | 3,709501972 | 0,000207667 | 0,008014781 | The Moose: suction sampler vs FISH | Function unknown |
| gene_247156 | 11,81057818 | 6,323051457 | 1,722570577 | 3,670706758 | 0,000241881 | 0,008847076 | The Moose: suction sampler vs FISH | Function unknown |
| gene_1072505 | 10,05405032 | 6,345435578 | 1,657721635 | 3,827805251 | 0,000129291 | 0,005910274 | The Moose: suction sampler vs FISH | Function unknown |
| gene_253445 | 3,762300591 | 6,351145449 | 1,797473226 | 3,533374159 | 0,000410291 | 0,012380188 | The Moose: suction sampler vs FISH | Function unknown |
| gene_3013 | 3,465211567 | 6,385073648 | 2,1930592 | 2,911491696 | 0,003597075 | 0,0468273 | The Moose: suction sampler vs FISH | Function unknown |
| gene_1511467 | 3,424237588 | 6,385688714 | 2,143682443 | 2,978840796 | 0,002893411 | 0,04121539 | The Moose: suction sampler vs FISH | Function unknown |
| gene_1349471 | 9,923644592 | 6,426769858 | 1,724497452 | 3,726749409 | 0,000193965 | 0,007660085 | The Moose: suction sampler vs FISH | Function unknown |
| gene_1416799 | 4,97034514 | 6,468557343 | 1,812362943 | 3,569129113 | 0,00035817 | 0,011358285 | The Moose: suction sampler vs FISH | Function unknown |
| gene_710064 | 5,348422402 | 6,510637397 | 1,673029677 | 3,891525348 | 9,96E-05 | 0,004990652 | The Moose: suction sampler vs FISH | Function unknown |
| gene_422183 | 5,825098271 | 6,521179749 | 1,816857652 | 3,589262891 | 0,000331614 | 0,010843906 | The Moose: suction sampler vs FISH | Function unknown |
| gene_1023862 | 6,136023332 | 6,522359103 | 2,179352026 | 2,992797412 | 0,002764332 | 0,040036748 | The Moose: suction sampler vs FISH | Function unknown |
| gene_781719 | 5,602627809 | 6,534948971 | 1,670066382 | 3,912987557 | 9,12E-05 | 0,004715026 | The Moose: suction sampler vs FISH | Function unknown |
| gene_1388949 | 13,85951495 | 6,566973636 | 1,469376998 | 4,469223109 | 7,85E-06 | 0,000907991 | The Moose: suction sampler vs FISH | Function unknown |
| gene_144169 | 3,121450631 | 6,621782439 | 2,201675048 | 3,007611157 | 0,002633098 | 0,038947916 | The Moose: suction sampler vs FISH | Function unknown |
| gene_1480564 | 3,763815592 | 6,653316128 | 2,225627321 | 2,989411599 | 0,002795153 | 0,040371686 | The Moose: suction sampler vs FISH | Function unknown |
| gene_743092 | 15,32393729 | 6,679297038 | 1,269180797 | 5,262683657 | 1,42E-07 | 5,70E-05 | The Moose: suction sampler vs FISH | Function unknown |
| gene_440366 | 3,524154478 | 6,693618027 | 2,319394492 | 2,885933397 | 0,003902547 | 0,049070808 | The Moose: suction sampler vs FISH | Function unknown |
| gene_738727 | 7,183940161 | 6,700186682 | 1,601572097 | 4,183506128 | 2,87E-05 | 0,002158827 | The Moose: suction sampler vs FISH | Function unknown |
| gene_1122784 | 5,043354496 | 6,727180572 | 1,814496871 | 3,707463309 | 0,000209346 | 0,008049569 | The Moose: suction sampler vs FISH | Function unknown |
| gene_49590 | 69,53257258 | 6,734774023 | 1,792352983 | 3,75750429 | 0,000171616 | 0,007073158 | The Moose: suction sampler vs FISH | Function unknown |
| gene_548666 | 7,152250427 | 6,762104153 | 1,954808865 | 3,459215003 | 0,000541752 | 0,014654547 | The Moose: suction sampler vs FISH | Function unknown |
| gene_612902 | 20,49987539 | 6,768926625 | 1,325444548 | 5,106910461 | 3,27E-07 | 0,000104117 | The Moose: suction sampler vs FISH | Function unknown |
| gene_1397340 | 5,124088544 | 6,818487133 | 1,985334227 | 3,434427835 | 0,000593806 | 0,015536905 | The Moose: suction sampler vs FISH | Function unknown |
| gene_1279365 | 7,671941779 | 6,935760024 | 1,704907282 | 4,068115666 | 4,74E-05 | 0,003058801 | The Moose: suction sampler vs FISH | Function unknown |
| gene_1341358 | 3,563081331 | 6,992265455 | 2,302921398 | 3,036258841 | 0,002395336 | 0,036739539 | The Moose: suction sampler vs FISH | Function unknown |
| gene_1096385 | 145,3674483 | 7,046800136 | 1,063619672 | 6,625300679 | 3,47E-11 | 1,59E-07 | The Moose: suction sampler vs FISH | Function unknown |
| gene_1069596 | 5,430103742 | 7,14335615 | 1,975686118 | 3,615633113 | 0,000299614 | 0,010192304 | The Moose: suction sampler vs FISH | Function unknown |
| gene_486997 | 8,952907355 | 7,169225913 | 1,784716717 | 4,017010569 | 5,89E-05 | 0,003516594 | The Moose: suction sampler vs FISH | Function unknown |
| gene_1276815 | 11,60482942 | 7,24577964 | 1,788762683 | 4,050721601 | 5,11E-05 | 0,003205211 | The Moose: suction sampler vs FISH | Function unknown |
| gene_1545496 | 5,716278433 | 7,274157721 | 1,781748383 | 4,082595383 | 4,45E-05 | 0,002917852 | The Moose: suction sampler vs FISH | Function unknown |
| gene_1925689 | 6,055083157 | 7,445884104 | 2,016424001 | 3,692618268 | 0,000221957 | 0,00835194 | The Moose: suction sampler vs FISH | Function unknown |
| gene_1351164 | 32,94330533 | 7,465911354 | 1,410216356 | 5,294160234 | 1,20E-07 | 5,17E-05 | The Moose: suction sampler vs FISH | Function unknown |
| gene_813152 | 5,521564697 | 7,525249348 | 2,034511041 | 3,698799955 | 0,000216621 | 0,008217845 | The Moose: suction sampler vs FISH | Function unknown |
| gene_839989 | 22,10129518 | 7,91922581 | 1,324077732 | 5,980937236 | 2,22E-09 | 3,19E-06 | The Moose: suction sampler vs FISH | Function unknown |
| gene_1480843 | 22,78278796 | 8,212790244 | 1,546893833 | 5,309213902 | 1,10E-07 | 4,93E-05 | The Moose: suction sampler vs FISH | Function unknown |
| gene_1514491 | 16,08826286 | 8,262279096 | 1,580525812 | 5,227550878 | 1,72E-07 | 6,47E-05 | The Moose: suction sampler vs FISH | Function unknown |
| gene_189522 | 10,59897923 | 8,701995593 | 2,682364147 | 3,244151471 | 0,001178011 | 0,024039735 | The Moose: suction sampler vs FISH | Function unknown |
| gene_1080470 | 53,30659783 | 9,613445565 | 1,602366388 | 5,999530216 | 1,98E-09 | 2,91E-06 | The Moose: suction sampler vs FISH | Function unknown |
| gene_493024 | 55,7311026 | 11,11531943 | 2,081133691 | 5,340992497 | 9,24E-08 | 4,56E-05 | The Moose: suction sampler vs FISH | Function unknown |
| gene_1630036 | 7,602756954 | -5,86393818 | 1,767558279 | -3,317535976 | 0,000908152 | 0,02056145 | The Moose: suction sampler vs FISH | General function prediction only |
| gene_1003782 | 9,088933689 | -5,07272279 | 1,574132701 | -3,222550925 | 0,001270546 | 0,025213519 | The Moose: suction sampler vs FISH | General function prediction only |
| gene_873932 | 850,8037851 | -4,287488949 | 0,757257061 | -5,661867241 | 1,50E-08 | 1,17E-05 | The Moose: suction sampler vs FISH | General function prediction only |
| gene_239727 | 36,6803221 | -3,811749089 | 0,965173748 | -3,949287989 | 7,84E-05 | 0,004251782 | The Moose: suction sampler vs FISH | General function prediction only |
| gene_69345 | 84,81415923 | -3,488493286 | 0,71173139 | -4,901418338 | 9,51E-07 | 0,0002167 | The Moose: suction sampler vs FISH | General function prediction only |
| gene_1371540 | 17,06306524 | -3,376558288 | 1,081284168 | -3,122729795 | 0,001791822 | 0,031046275 | The Moose: suction sampler vs FISH | General function prediction only |
| gene_1120301 | 16,09838972 | -3,307137213 | 1,101743524 | -3,001730568 | 0,002684497 | 0,039338159 | The Moose: suction sampler vs FISH | General function prediction only |
| gene_69019 | 23,544797 | -3,231773783 | 1,019392353 | -3,170294316 | 0,001522846 | 0,028160723 | The Moose: suction sampler vs FISH | General function prediction only |
| gene_1235153 | 52,81131633 | -3,108829352 | 0,863758725 | -3,599187204 | 0,000319213 | 0,010599955 | The Moose: suction sampler vs FISH | General function prediction only |
| gene_980493 | 114,1237671 | -3,015228118 | 0,721428675 | -4,179523527 | 2,92E-05 | 0,002178151 | The Moose: suction sampler vs FISH | General function prediction only |
| gene_805846 | 99,39901297 | -3,014206727 | 0,732476068 | -4,115092434 | 3,87E-05 | 0,002639276 | The Moose: suction sampler vs FISH | General function prediction only |
| gene_1665172 | 44,35506975 | -2,901001336 | 0,923706764 | -3,140608525 | 0,001685972 | 0,029973367 | The Moose: suction sampler vs FISH | General function prediction only |
| gene_176362 | 1031,296746 | -2,880026632 | 0,629799094 | -4,572929141 | 4,81E-06 | 0,000644591 | The Moose: suction sampler vs FISH | General function prediction only |
| gene_1593737 | 80,76978318 | -2,763176357 | 0,792887516 | -3,484953792 | 0,000492222 | 0,013823223 | The Moose: suction sampler vs FISH | General function prediction only |
| gene_474108 | 29,52923114 | -2,580423383 | 0,786155931 | -3,282330238 | 0,00102953 | 0,022148632 | The Moose: suction sampler vs FISH | General function prediction only |
| gene_387624 | 56,00345141 | -2,477687453 | 0,764343873 | -3,241587382 | 0,00118866 | 0,024188888 | The Moose: suction sampler vs FISH | General function prediction only |
| gene_821664 | 6386,993874 | -2,424677477 | 0,6466406 | -3,749652401 | 0,00017708 | 0,00720569 | The Moose: suction sampler vs FISH | General function prediction only |
| gene_928072 | 40,49670806 | -2,422231196 | 0,839664221 | -2,884761713 | 0,0039171 | 0,049178022 | The Moose: suction sampler vs FISH | General function prediction only |
| gene_74292 | 445,9016022 | -2,335082627 | 0,679343595 | -3,437263033 | 0,000587625 | 0,015452235 | The Moose: suction sampler vs FISH | General function prediction only |
| gene_1128908 | 224,5683969 | -2,324460367 | 0,665849285 | -3,490970735 | 0,000481269 | 0,013594923 | The Moose: suction sampler vs FISH | General function prediction only |

| | | | | | | | | |
|---|---|---|---|---|---|---|---|---|
| gene_1120285 | 43,06847192 | -2,279686408 | 0,746038363 | -3,055722761 | 0,002245188 | 0,035391185 | The Moose: suction sampler vs FISH | General function prediction only |
| gene_1274176 | 1359,408332 | -2,27704225 | 0,564046124 | -4,036978805 | 5,41E-05 | 0,003307895 | The Moose: suction sampler vs FISH | General function prediction only |
| gene_205577 | 71,56553101 | -2,271919187 | 0,616033531 | -3,687979751 | 0,000226042 | 0,008467069 | The Moose: suction sampler vs FISH | General function prediction only |
| gene_87649 | 2702,909589 | -2,262172155 | 0,591015917 | -3,827599377 | 0,000129399 | 0,005910274 | The Moose: suction sampler vs FISH | General function prediction only |
| gene_440831 | 455,1632249 | -2,241730265 | 0,548087747 | -4,090093742 | 4,31E-05 | 0,002856724 | The Moose: suction sampler vs FISH | General function prediction only |
| gene_1885233 | 845,6595212 | -2,226915322 | 0,584095741 | -3,81258613 | 0,00013752 | 0,006145279 | The Moose: suction sampler vs FISH | General function prediction only |
| gene_1074196 | 823,9944644 | -2,176975455 | 0,627633456 | -3,468545907 | 0,000523283 | 0,014356566 | The Moose: suction sampler vs FISH | General function prediction only |
| gene_440830 | 197,4675151 | -2,173109207 | 0,667175334 | -3,25717858 | 0,001125256 | 0,023425294 | The Moose: suction sampler vs FISH | General function prediction only |
| gene_488576 | 101,126117 | -2,159885605 | 0,684549405 | -3,155193168 | 0,001603919 | 0,029057674 | The Moose: suction sampler vs FISH | General function prediction only |
| gene_553121 | 224,6864783 | -2,067687437 | 0,636662154 | -3,247699624 | 0,00116342 | 0,023888006 | The Moose: suction sampler vs FISH | General function prediction only |
| gene_176368 | 6309,45244 | -2,061163258 | 0,583624362 | -3,531660759 | 0,000412959 | 0,012414964 | The Moose: suction sampler vs FISH | General function prediction only |
| gene_44421 | 35,92987689 | 2,475802051 | 0,854364113 | 2,897830109 | 0,003757541 | 0,048082426 | The Moose: suction sampler vs FISH | General function prediction only |
| gene_388862 | 21,17721004 | 2,49707392 | 0,857029723 | 2,913637477 | 0,003572445 | 0,046635886 | The Moose: suction sampler vs FISH | General function prediction only |
| gene_47331 | 24,07652123 | 2,652466268 | 0,883758323 | 3,001347993 | 0,002687872 | 0,039361021 | The Moose: suction sampler vs FISH | General function prediction only |
| gene_1083809 | 16,32316129 | 2,6837583 | 0,871986363 | 3,077752604 | 0,00208568 | 0,034074598 | The Moose: suction sampler vs FISH | General function prediction only |
| gene_350172 | 14,98798912 | 2,707855297 | 0,905804269 | 2,989448592 | 0,002794815 | 0,040371686 | The Moose: suction sampler vs FISH | General function prediction only |
| gene_717819 | 24,62054373 | 2,717829163 | 0,929354296 | 2,924427395 | 0,003450906 | 0,045700744 | The Moose: suction sampler vs FISH | General function prediction only |
| gene_980213 | 16,7017355 | 2,782928158 | 0,908948676 | 3,061699996 | 0,002200839 | 0,035070651 | The Moose: suction sampler vs FISH | General function prediction only |
| gene_330291 | 36,58325592 | 2,810378523 | 0,954990729 | 2,942833304 | 0,003252235 | 0,044160858 | The Moose: suction sampler vs FISH | General function prediction only |
| gene_848486 | 31,39331919 | 2,856096172 | 0,978052052 | 2,920188314 | 0,003498199 | 0,046068719 | The Moose: suction sampler vs FISH | General function prediction only |
| gene_1079509 | 61,27662906 | 2,905368947 | 0,836571689 | 3,472946771 | 0,000514777 | 0,014209953 | The Moose: suction sampler vs FISH | General function prediction only |
| gene_1705590 | 79,20949474 | 3,204261788 | 0,767474635 | 4,175071909 | 2,98E-05 | 0,002198862 | The Moose: suction sampler vs FISH | General function prediction only |
| gene_186943 | 25,65746173 | 3,255395275 | 0,880357534 | 3,697810433 | 0,000217467 | 0,008227995 | The Moose: suction sampler vs FISH | General function prediction only |
| gene_332013 | 35,22983387 | 3,305463751 | 0,858039375 | 3,852345065 | 0,000116992 | 0,005535866 | The Moose: suction sampler vs FISH | General function prediction only |
| gene_1861082 | 43,94024546 | 3,437957761 | 0,840807826 | 4,088874597 | 4,33E-05 | 0,002865552 | The Moose: suction sampler vs FISH | General function prediction only |
| gene_189644 | 10,47692384 | 3,464857354 | 1,187086008 | 2,918792177 | 0,003513904 | 0,046134909 | The Moose: suction sampler vs FISH | General function prediction only |
| gene_554335 | 15,80831354 | 3,673794007 | 1,168137274 | 3,145001953 | 0,001660857 | 0,029700915 | The Moose: suction sampler vs FISH | General function prediction only |
| gene_332014 | 39,61062957 | 3,721591488 | 0,921874371 | 4,03698335 | 5,41E-05 | 0,003307895 | The Moose: suction sampler vs FISH | General function prediction only |
| gene_1577833 | 6,644685182 | 5,071304143 | 1,655171697 | 3,063914246 | 0,002184615 | 0,034925787 | The Moose: suction sampler vs FISH | General function prediction only |
| gene_186011 | 13,88999002 | 5,617310026 | 1,45306391 | 3,865838239 | 0,000110708 | 0,005338786 | The Moose: suction sampler vs FISH | General function prediction only |
| gene_1289648 | 21,35181648 | -6,442103191 | 1,579137921 | -4,07950636 | 4,51E-05 | 0,002951628 | The Moose: suction sampler vs FISH | Inorganic ion transport and metabolism |
| gene_791214 | 7,583894615 | -5,540725783 | 1,560308043 | -3,551046095 | 0,000383703 | 0,011875719 | The Moose: suction sampler vs FISH | Inorganic ion transport and metabolism |
| gene_1222439 | 11,3743601 | -5,473787485 | 1,649150593 | -3,319155637 | 0,000902901 | 0,02050512 | The Moose: suction sampler vs FISH | Inorganic ion transport and metabolism |
| gene_482061 | 8,431361035 | -4,459830665 | 1,457800173 | -3,059288061 | 0,002218637 | 0,03520092 | The Moose: suction sampler vs FISH | Inorganic ion transport and metabolism |
| gene_632179 | 11,6794492 | -3,894041218 | 1,182174892 | -3,293963732 | 0,000987852 | 0,021549906 | The Moose: suction sampler vs FISH | Inorganic ion transport and metabolism |
| gene_501651 | 13,10211231 | -3,816818446 | 1,297897462 | -2,940770407 | 0,003273971 | 0,044323834 | The Moose: suction sampler vs FISH | Inorganic ion transport and metabolism |
| gene_397547 | 212,6063004 | -3,468403941 | 0,651351576 | -5,324933677 | 1,01E-07 | 4,64E-05 | The Moose: suction sampler vs FISH | Inorganic ion transport and metabolism |
| gene_963149 | 65,2109441 | -3,368531898 | 1,046535462 | -3,218746062 | 0,001287525 | 0,025413125 | The Moose: suction sampler vs FISH | Inorganic ion transport and metabolism |
| gene_1114201 | 22,71781818 | -3,316570494 | 1,066364602 | -3,110165593 | 0,001869825 | 0,031899951 | The Moose: suction sampler vs FISH | Inorganic ion transport and metabolism |
| gene_1003778 | 30,20346348 | -3,100422491 | 1,016208958 | -3,050969491 | 0,002281038 | 0,035717888 | The Moose: suction sampler vs FISH | Inorganic ion transport and metabolism |
| gene_288276 | 13,19983131 | -3,016958802 | 1,020316653 | -2,956884799 | 0,003107642 | 0,043085392 | The Moose: suction sampler vs FISH | Inorganic ion transport and metabolism |
| gene_334865 | 36,56779696 | -2,805962575 | 0,816818552 | -3,435233647 | 0,000592043 | 0,015512882 | The Moose: suction sampler vs FISH | Inorganic ion transport and metabolism |
| gene_248654 | 36,4399116 | -2,800630913 | 0,835790218 | -3,350877833 | 0,000805558 | 0,018987237 | The Moose: suction sampler vs FISH | Inorganic ion transport and metabolism |
| gene_1524226 | 19,48357241 | -2,669678985 | 0,921099294 | -2,898361775 | 0,003751176 | 0,0480412 | The Moose: suction sampler vs FISH | Inorganic ion transport and metabolism |
| gene_53652 | 238,6142767 | -2,607497698 | 0,704434451 | -3,701547665 | 0,000214288 | 0,008171553 | The Moose: suction sampler vs FISH | Inorganic ion transport and metabolism |
| gene_196822 | 3082,825913 | -2,60099864 | 0,767088176 | -3,390742709 | 0,000697035 | 0,017248564 | The Moose: suction sampler vs FISH | Inorganic ion transport and metabolism |
| gene_829538 | 519,886512 | -2,460739094 | 0,617974862 | -3,981940442 | 6,84E-05 | 0,003873709 | The Moose: suction sampler vs FISH | Inorganic ion transport and metabolism |
| gene_232570 | 56,50901377 | -2,339032343 | 0,738933701 | -3,165415706 | 0,001548615 | 0,028420302 | The Moose: suction sampler vs FISH | Inorganic ion transport and metabolism |
| gene_38719 | 91,667595 | -2,289149435 | 0,596871724 | -3,835245234 | 0,000125439 | 0,005797411 | The Moose: suction sampler vs FISH | Inorganic ion transport and metabolism |
| gene_775655 | 1882,727495 | -2,263716418 | 0,596791203 | -3,793146423 | 0,00014875 | 0,006426454 | The Moose: suction sampler vs FISH | Inorganic ion transport and metabolism |
| gene_571435 | 302,4635297 | -2,217597751 | 0,584461481 | -3,79425817 | 0,000148086 | 0,006412819 | The Moose: suction sampler vs FISH | Inorganic ion transport and metabolism |
| gene_475356 | 383,2000684 | -2,146581216 | 0,60811036 | -3,529920485 | 0,000415685 | 0,012471386 | The Moose: suction sampler vs FISH | Inorganic ion transport and metabolism |
| gene_653158 | 69,19966294 | -2,08759979 | 0,722767752 | -2,88834108 | 0,003872797 | 0,048896414 | The Moose: suction sampler vs FISH | Inorganic ion transport and metabolism |
| gene_131510 | 47,36890594 | -2,08211807 | 0,721343208 | -2,886445796 | 0,003896199 | 0,049048664 | The Moose: suction sampler vs FISH | Inorganic ion transport and metabolism |
| gene_356101 | 47,45012414 | 2,124748345 | 0,709876258 | 2,993124958 | 0,002761367 | 0,040015779 | The Moose: suction sampler vs FISH | Inorganic ion transport and metabolism |
| gene_15170 | 33,57261624 | 2,357108858 | 0,75837432 | 3,10810743 | 0,001882896 | 0,032028098 | The Moose: suction sampler vs FISH | Inorganic ion transport and metabolism |
| gene_699930 | 34,61968713 | 2,432189918 | 0,760146191 | 3,199634421 | 0,00137602 | 0,026440739 | The Moose: suction sampler vs FISH | Inorganic ion transport and metabolism |
| gene_1517572 | 60,68037321 | 2,560357654 | 0,708726958 | 3,612615019 | 0,000303125 | 0,010271678 | The Moose: suction sampler vs FISH | Inorganic ion transport and metabolism |
| gene_1378314 | 268,0065869 | 2,580785482 | 0,863819199 | 2,987645429 | 0,002811355 | 0,040526 | The Moose: suction sampler vs FISH | Inorganic ion transport and metabolism |
| gene_333194 | 36,81391235 | 2,615115638 | 0,844171763 | 3,0978478 | 0,001949315 | 0,032644267 | The Moose: suction sampler vs FISH | Inorganic ion transport and metabolism |
| gene_1151056 | 20,48633297 | 2,674468032 | 0,845119159 | 3,164604664 | 0,001552938 | 0,028468833 | The Moose: suction sampler vs FISH | Inorganic ion transport and metabolism |
| gene_522157 | 25,4348307 | 2,742561795 | 0,876873263 | 3,127660417 | 0,001762036 | 0,030732073 | The Moose: suction sampler vs FISH | Inorganic ion transport and metabolism |
| gene_356100 | 65,86533734 | 2,767158979 | 0,726272192 | 3,810085269 | 0,000138919 | 0,006176087 | The Moose: suction sampler vs FISH | Inorganic ion transport and metabolism |
| gene_432201 | 23,47750611 | 2,769182205 | 0,934357371 | 2,963729179 | 0,003039356 | 0,042472454 | The Moose: suction sampler vs FISH | Inorganic ion transport and metabolism |
| gene_1410346 | 16,35674472 | 2,871183759 | 0,938164648 | 3,060426296 | 0,002210222 | 0,035159134 | The Moose: suction sampler vs FISH | Inorganic ion transport and metabolism |

| | | | | | | | | |
|---|---|---|---|---|---|---|---|---|
| gene_450963 | 31,91148382 | 2,876159008 | 0,874263148 | 3,289809268 | 0,001002553 | 0,021771885 | The Moose: suction sampler vs FISH | Inorganic ion transport and metabolism |
| gene_1138738 | 47,33269646 | 2,915491166 | 0,816175165 | 3,572139033 | 0,000354077 | 0,011267422 | The Moose: suction sampler vs FISH | Inorganic ion transport and metabolism |
| gene_1571489 | 642,8537806 | 2,938055833 | 0,912225794 | 3,22075505 | 0,001278534 | 0,025289765 | The Moose: suction sampler vs FISH | Inorganic ion transport and metabolism |
| gene_925572 | 16,22370344 | 2,986707772 | 0,992616214 | 3,008925029 | 0,002621738 | 0,038868298 | The Moose: suction sampler vs FISH | Inorganic ion transport and metabolism |
| gene_450962 | 32,23213371 | 3,01030314 | 0,845592219 | 3,559993897 | 0,000370863 | 0,011601651 | The Moose: suction sampler vs FISH | Inorganic ion transport and metabolism |
| gene_450964 | 20,72909426 | 3,129046972 | 0,913641627 | 3,424807801 | 0,000615234 | 0,015913531 | The Moose: suction sampler vs FISH | Inorganic ion transport and metabolism |
| gene_1612085 | 23,76515731 | 3,129794136 | 0,894199055 | 3,500108973 | 0,000465068 | 0,013354236 | The Moose: suction sampler vs FISH | Inorganic ion transport and metabolism |
| gene_304426 | 14,15935173 | 3,129909981 | 1,060443492 | 2,951510386 | 0,003162239 | 0,043468211 | The Moose: suction sampler vs FISH | Inorganic ion transport and metabolism |
| gene_614041 | 26,63683989 | 3,135192577 | 0,885679314 | 3,539873325 | 0,000400319 | 0,0122099 | The Moose: suction sampler vs FISH | Inorganic ion transport and metabolism |
| gene_743132 | 66,90889816 | 3,136276804 | 0,88249791 | 3,553863153 | 0,000379617 | 0,011793971 | The Moose: suction sampler vs FISH | Inorganic ion transport and metabolism |
| gene_1267076 | 209,8506728 | 3,236828013 | 0,707255579 | 4,576603009 | 4,73E-06 | 0,000635699 | The Moose: suction sampler vs FISH | Inorganic ion transport and metabolism |
| gene_1928898 | 59,29804408 | 3,276639994 | 0,833264466 | 3,932292962 | 8,41E-05 | 0,004452176 | The Moose: suction sampler vs FISH | Inorganic ion transport and metabolism |
| gene_1928897 | 54,22425385 | 3,379023624 | 0,753838008 | 4,482426713 | 7,38E-06 | 0,000878471 | The Moose: suction sampler vs FISH | Inorganic ion transport and metabolism |
| gene_1679660 | 331,7045399 | 3,443296083 | 0,861427786 | 3,997196446 | 6,41E-05 | 0,00371303 | The Moose: suction sampler vs FISH | Inorganic ion transport and metabolism |
| gene_1135002 | 19,64727554 | 3,481605487 | 1,094806298 | 3,180110941 | 0,001472187 | 0,027593959 | The Moose: suction sampler vs FISH | Inorganic ion transport and metabolism |
| gene_1431916 | 16,72179016 | 3,494777774 | 1,122213634 | 3,114182244 | 0,001844555 | 0,031630482 | The Moose: suction sampler vs FISH | Inorganic ion transport and metabolism |
| gene_1893346 | 289,2621957 | 3,566412266 | 0,759845235 | 4,693603516 | 2,68E-06 | 0,000427235 | The Moose: suction sampler vs FISH | Inorganic ion transport and metabolism |
| gene_450961 | 113,9883456 | 3,609772765 | 0,817863547 | 4,413661395 | 1,02E-05 | 0,001072517 | The Moose: suction sampler vs FISH | Inorganic ion transport and metabolism |
| gene_1735345 | 62,87197559 | 3,641262536 | 0,78548004 | 4,635716185 | 3,56E-06 | 0,00051732 | The Moose: suction sampler vs FISH | Inorganic ion transport and metabolism |
| gene_1092750 | 11,00161219 | 3,736794422 | 1,190534949 | 3,138752395 | 0,001696687 | 0,030082389 | The Moose: suction sampler vs FISH | Inorganic ion transport and metabolism |
| gene_222621 | 36,04863841 | 3,74121657 | 1,03079475 | 3,629448606 | 0,000284027 | 0,00984091 | The Moose: suction sampler vs FISH | Inorganic ion transport and metabolism |
| gene_1284018 | 20,30594037 | 3,800286855 | 1,067899328 | 3,558656473 | 0,000372757 | 0,011630041 | The Moose: suction sampler vs FISH | Inorganic ion transport and metabolism |
| gene_406111 | 18,71398048 | 3,843635584 | 1,028815386 | 3,735981826 | 0,000186984 | 0,007479876 | The Moose: suction sampler vs FISH | Inorganic ion transport and metabolism |
| gene_159604 | 22,6000514 | 3,848460569 | 1,16741863 | 3,296555724 | 0,000978782 | 0,021439499 | The Moose: suction sampler vs FISH | Inorganic ion transport and metabolism |
| gene_166550 | 14,74747928 | 3,871845688 | 0,955479419 | 4,052254411 | 5,07E-05 | 0,003195201 | The Moose: suction sampler vs FISH | Inorganic ion transport and metabolism |
| gene_374027 | 179,4373091 | 3,936834961 | 0,858222329 | 4,587197079 | 4,49E-06 | 0,000614417 | The Moose: suction sampler vs FISH | Inorganic ion transport and metabolism |
| gene_781582 | 681,7243258 | 3,963340342 | 0,841940748 | 4,707386296 | 2,51E-06 | 0,000406806 | The Moose: suction sampler vs FISH | Inorganic ion transport and metabolism |
| gene_1209244 | 16,84307541 | 4,088978907 | 1,258819517 | 3,248264626 | 0,001161112 | 0,02386063 | The Moose: suction sampler vs FISH | Inorganic ion transport and metabolism |
| gene_822819 | 742,1097838 | 4,1306972 | 0,772902816 | 5,344394033 | 9,07E-08 | 4,54E-05 | The Moose: suction sampler vs FISH | Inorganic ion transport and metabolism |
| gene_1813354 | 1287,669377 | 4,147865999 | 0,807554487 | 5,136329581 | 2,80E-07 | 9,23E-05 | The Moose: suction sampler vs FISH | Inorganic ion transport and metabolism |
| gene_781581 | 126,9928222 | 4,24156763 | 0,968156279 | 4,381077438 | 1,18E-05 | 0,001188374 | The Moose: suction sampler vs FISH | Inorganic ion transport and metabolism |
| gene_697899 | 34,86612463 | 4,283637031 | 0,990255594 | 4,325789277 | 1,52E-05 | 0,001388331 | The Moose: suction sampler vs FISH | Inorganic ion transport and metabolism |
| gene_519646 | 8,405413053 | 4,291852815 | 1,480009128 | 2,899882665 | 0,003733024 | 0,047956041 | The Moose: suction sampler vs FISH | Inorganic ion transport and metabolism |
| gene_1285639 | 79,04073584 | 4,327846727 | 0,926421483 | 4,671574232 | 2,99E-06 | 0,000450774 | The Moose: suction sampler vs FISH | Inorganic ion transport and metabolism |
| gene_1798992 | 8,97927158 | 4,433028644 | 1,540330758 | 2,877971905 | 0,004002408 | 0,04980631 | The Moose: suction sampler vs FISH | Inorganic ion transport and metabolism |
| gene_737958 | 65,98130418 | 4,554387402 | 0,961209851 | 4,738182197 | 2,16E-06 | 0,000366549 | The Moose: suction sampler vs FISH | Inorganic ion transport and metabolism |
| gene_1527107 | 10,23232902 | 4,556315837 | 1,419347245 | 3,210148786 | 0,001326663 | 0,025852149 | The Moose: suction sampler vs FISH | Inorganic ion transport and metabolism |
| gene_925573 | 8,255614466 | 4,639569617 | 1,265029652 | 3,667557996 | 0,000244878 | 0,008930023 | The Moose: suction sampler vs FISH | Inorganic ion transport and metabolism |
| gene_781583 | 55,45179162 | 4,697921997 | 1,037446527 | 4,528350982 | 5,94E-06 | 0,000747602 | The Moose: suction sampler vs FISH | Inorganic ion transport and metabolism |
| gene_263198 | 55,13982253 | 4,742128643 | 0,9711892 | 4,882806194 | 1,05E-06 | 0,000229981 | The Moose: suction sampler vs FISH | Inorganic ion transport and metabolism |
| gene_593493 | 6,768999276 | 4,79504254 | 1,496746259 | 3,203644245 | 0,001357 | 0,026205737 | The Moose: suction sampler vs FISH | Inorganic ion transport and metabolism |
| gene_104240 | 28,777618 | 4,878221992 | 1,071515399 | 4,552638253 | 5,30E-06 | 0,000686229 | The Moose: suction sampler vs FISH | Inorganic ion transport and metabolism |
| gene_1599283 | 32,89946448 | 5,024127447 | 1,019986037 | 4,925682573 | 8,41E-07 | 0,000199814 | The Moose: suction sampler vs FISH | Inorganic ion transport and metabolism |
| gene_887822 | 12,11503685 | 5,088175928 | 1,209041451 | 4,208437953 | 2,57E-05 | 0,002006993 | The Moose: suction sampler vs FISH | Inorganic ion transport and metabolism |
| gene_1519695 | 42,9928063 | 5,092659488 | 1,018820967 | 4,998581354 | 5,78E-07 | 0,00015149 | The Moose: suction sampler vs FISH | Inorganic ion transport and metabolism |
| gene_169941 | 10,20163568 | 5,515152105 | 1,65303572 | 3,336378057 | 0,000848777 | 0,019733585 | The Moose: suction sampler vs FISH | Inorganic ion transport and metabolism |
| gene_407148 | 6,040616815 | 5,676324141 | 1,649391977 | 3,441464626 | 0,000578574 | 0,015307412 | The Moose: suction sampler vs FISH | Inorganic ion transport and metabolism |
| gene_751790 | 6,021342292 | 5,774554044 | 1,882518732 | 3,067461664 | 0,002158851 | 0,034702746 | The Moose: suction sampler vs FISH | Inorganic ion transport and metabolism |
| gene_182729 | 3,156417337 | 5,853322255 | 1,990663685 | 2,940387318 | 0,003278022 | 0,044349108 | The Moose: suction sampler vs FISH | Inorganic ion transport and metabolism |
| gene_1399733 | 3,241326887 | 6,043542615 | 2,002401071 | 3,018147914 | 0,002543247 | 0,03821018 | The Moose: suction sampler vs FISH | Inorganic ion transport and metabolism |
| gene_1952520 | 3,028509516 | 6,046780961 | 1,996543891 | 3,028624108 | 0,002456701 | 0,03734115 | The Moose: suction sampler vs FISH | Inorganic ion transport and metabolism |
| gene_199561 | 3,441517039 | 6,758610392 | 2,245620965 | 3,009684402 | 0,002615193 | 0,038835543 | The Moose: suction sampler vs FISH | Inorganic ion transport and metabolism |
| gene_751789 | 3,984474926 | 6,795412616 | 2,110666638 | 3,219557506 | 0,001283886 | 0,025375407 | The Moose: suction sampler vs FISH | Inorganic ion transport and metabolism |
| gene_1385135 | 3,704928642 | 6,801959242 | 2,194748967 | 3,099196922 | 0,00194046 | 0,032597686 | The Moose: suction sampler vs FISH | Inorganic ion transport and metabolism |
| gene_1124038 | 5,430043014 | 6,852543403 | 2,042283799 | 3,355333576 | 0,000792693 | 0,01881686 | The Moose: suction sampler vs FISH | Inorganic ion transport and metabolism |
| gene_1594873 | 5,631768 | 6,913663286 | 1,696899513 | 4,074291515 | 4,62E-05 | 0,003005166 | The Moose: suction sampler vs FISH | Inorganic ion transport and metabolism |
| gene_1508153 | 13,50245826 | 7,564696783 | 1,612240363 | 4,692040316 | 2,70E-06 | 0,000427235 | The Moose: suction sampler vs FISH | Inorganic ion transport and metabolism |
| gene_770643 | 6,197305656 | 7,625921403 | 1,97630947 | 3,858667641 | 0,000114007 | 0,005442151 | The Moose: suction sampler vs FISH | Inorganic ion transport and metabolism |
| gene_1171029 | 10,5663115 | -4,799874642 | 1,473546364 | -3,257362482 | 0,001124527 | 0,023421866 | The Moose: suction sampler vs FISH | Intracellular trafficking, secretion, and vesicular transport |
| gene_103832 | 8,68206854 | -4,22607303 | 1,399185596 | -3,0203806 | 0,002524572 | 0,037994269 | The Moose: suction sampler vs FISH | Intracellular trafficking, secretion, and vesicular transport |
| gene_335929 | 18,62933368 | -3,481736612 | 1,070658049 | -3,251959497 | 0,001146124 | 0,023698492 | The Moose: suction sampler vs FISH | Intracellular trafficking, secretion, and vesicular transport |
| gene_600131 | 17,30403453 | -3,10635583 | 1,020604711 | -3,043642457 | 0,002337327 | 0,036208609 | The Moose: suction sampler vs FISH | Intracellular trafficking, secretion, and vesicular transport |
| gene_164129 | 23,27713239 | -3,065924781 | 0,876688794 | -3,497164333 | 0,000470232 | 0,013406908 | The Moose: suction sampler vs FISH | Intracellular trafficking, secretion, and vesicular transport |
| gene_779575 | 52,5869723 | -3,018197342 | 0,875660497 | -3,446766586 | 0,000567339 | 0,015141053 | The Moose: suction sampler vs FISH | Intracellular trafficking, secretion, and vesicular transport |
| gene_326076 | 22,13002459 | -2,814900967 | 0,915009158 | -3,076363709 | 0,002095421 | 0,034154056 | The Moose: suction sampler vs FISH | Intracellular trafficking, secretion, and vesicular transport |

| | | | | | | | | |
|---|---|---|---|---|---|---|---|---|
| gene_1791217 | 20,3180668 | -2,669410214 | 0,801953813 | -3,32863336 | 0,000872732 | 0,02008706 | The Moose: suction sampler vs FISH | Intracellular trafficking, secretion, and vesicular transport |
| gene_335928 | 121,0585568 | -2,511768875 | 0,701441298 | -3,58086825 | 0,000342454 | 0,011045919 | The Moose: suction sampler vs FISH | Intracellular trafficking, secretion, and vesicular transport |
| gene_124687 | 49,87529412 | -2,423955241 | 0,728050029 | -3,329380047 | 0,000870395 | 0,020049897 | The Moose: suction sampler vs FISH | Intracellular trafficking, secretion, and vesicular transport |
| gene_415477 | 173,7482826 | -2,262908845 | 0,627370346 | -3,606974505 | 0,000309788 | 0,010398714 | The Moose: suction sampler vs FISH | Intracellular trafficking, secretion, and vesicular transport |
| gene_879295 | 50,08924502 | -2,022454258 | 0,690851105 | -2,927482119 | 0,003417187 | 0,045400743 | The Moose: suction sampler vs FISH | Intracellular trafficking, secretion, and vesicular transport |
| gene_632987 | 1629,521553 | -2,012615248 | 0,596160791 | -3,375960442 | 0,000735585 | 0,017948517 | The Moose: suction sampler vs FISH | Intracellular trafficking, secretion, and vesicular transport |
| gene_1499043 | 41,40694643 | 2,217854462 | 0,758615243 | 2,923556415 | 0,003460575 | 0,045761962 | The Moose: suction sampler vs FISH | Intracellular trafficking, secretion, and vesicular transport |
| gene_5539 | 45,86799738 | 2,229197423 | 0,710939367 | 3,135566161 | 0,001715227 | 0,030282421 | The Moose: suction sampler vs FISH | Intracellular trafficking, secretion, and vesicular transport |
| gene_330193 | 54,62494301 | 2,395863412 | 0,82054725 | 2,919836014 | 0,003502156 | 0,046077151 | The Moose: suction sampler vs FISH | Intracellular trafficking, secretion, and vesicular transport |
| gene_1815134 | 17,88839822 | 2,593907121 | 0,85243909 | 3,042923712 | 0,002342917 | 0,036249327 | The Moose: suction sampler vs FISH | Intracellular trafficking, secretion, and vesicular transport |
| gene_1258326 | 44,65987812 | 2,597250077 | 0,822509674 | 3,157713714 | 0,001590116 | 0,028878856 | The Moose: suction sampler vs FISH | Intracellular trafficking, secretion, and vesicular transport |
| gene_10795 | 29,90627841 | 2,689653852 | 0,828917686 | 3,244777976 | 0,001175422 | 0,024020281 | The Moose: suction sampler vs FISH | Intracellular trafficking, secretion, and vesicular transport |
| gene_497119 | 25,22473348 | 2,724759979 | 0,824939611 | 3,302981143 | 0,000956628 | 0,021154692 | The Moose: suction sampler vs FISH | Intracellular trafficking, secretion, and vesicular transport |
| gene_657637 | 44,00997973 | 2,743059987 | 0,904653528 | 3,032166351 | 0,002428054 | 0,037089932 | The Moose: suction sampler vs FISH | Intracellular trafficking, secretion, and vesicular transport |
| gene_942528 | 25,47727245 | 2,852751138 | 0,82260376 | 3,467952952 | 0,000524439 | 0,014377268 | The Moose: suction sampler vs FISH | Intracellular trafficking, secretion, and vesicular transport |
| gene_1354528 | 112,5424673 | 2,870351357 | 0,748849942 | 3,833012724 | 0,000126583 | 0,005828286 | The Moose: suction sampler vs FISH | Intracellular trafficking, secretion, and vesicular transport |
| gene_1011379 | 35,97214313 | 2,87658041 | 0,864496653 | 3,327462749 | 0,000876407 | 0,020140086 | The Moose: suction sampler vs FISH | Intracellular trafficking, secretion, and vesicular transport |
| gene_181541 | 23,24341391 | 2,928137946 | 0,926396564 | 3,16078239 | 0,00157346 | 0,028704116 | The Moose: suction sampler vs FISH | Intracellular trafficking, secretion, and vesicular transport |
| gene_1180768 | 33,66855416 | 2,935450267 | 0,766901485 | 3,827675816 | 0,000129359 | 0,005910274 | The Moose: suction sampler vs FISH | Intracellular trafficking, secretion, and vesicular transport |
| gene_1079858 | 12,57073088 | 3,032806855 | 0,998457912 | 3,037490933 | 0,002385566 | 0,036636926 | The Moose: suction sampler vs FISH | Intracellular trafficking, secretion, and vesicular transport |
| gene_1180767 | 22,13629553 | 3,071437777 | 0,840882321 | 3,652636879 | 0,000259561 | 0,009267608 | The Moose: suction sampler vs FISH | Intracellular trafficking, secretion, and vesicular transport |
| gene_64180 | 16,60043404 | 3,104400756 | 1,000396308 | 3,103170945 | 0,00191459 | 0,032303486 | The Moose: suction sampler vs FISH | Intracellular trafficking, secretion, and vesicular transport |
| gene_1781208 | 11,81763069 | 3,122837135 | 1,0812044 | 2,888294881 | 0,003873366 | 0,048896414 | The Moose: suction sampler vs FISH | Intracellular trafficking, secretion, and vesicular transport |
| gene_1079856 | 25,25745772 | 3,185801836 | 0,895082737 | 3,559226096 | 0,000371949 | 0,011615767 | The Moose: suction sampler vs FISH | Intracellular trafficking, secretion, and vesicular transport |
| gene_1144745 | 30,67364838 | 3,246701758 | 0,808030001 | 4,018046056 | 5,87E-05 | 0,003506878 | The Moose: suction sampler vs FISH | Intracellular trafficking, secretion, and vesicular transport |
| gene_294774 | 32,9596439 | 3,255754604 | 1,113136688 | 2,924847091 | 0,003446255 | 0,045679518 | The Moose: suction sampler vs FISH | Intracellular trafficking, secretion, and vesicular transport |
| gene_1130223 | 29,77608649 | 3,279613093 | 0,897556438 | 3,653935234 | 0,000258251 | 0,009234312 | The Moose: suction sampler vs FISH | Intracellular trafficking, secretion, and vesicular transport |
| gene_984570 | 68,85646785 | 3,306129493 | 0,852174268 | 3,879640137 | 0,000104611 | 0,005132374 | The Moose: suction sampler vs FISH | Intracellular trafficking, secretion, and vesicular transport |
| gene_1495662 | 29,77262018 | 3,369457689 | 0,938951382 | 3,588532646 | 0,000332544 | 0,010869479 | The Moose: suction sampler vs FISH | Intracellular trafficking, secretion, and vesicular transport |
| gene_1696408 | 29,61601115 | 3,441695828 | 0,912001379 | 3,773783578 | 0,00016079 | 0,006802101 | The Moose: suction sampler vs FISH | Intracellular trafficking, secretion, and vesicular transport |
| gene_654256 | 31,51423663 | 3,454604665 | 1,165736472 | 2,963452502 | 0,00304209 | 0,042492641 | The Moose: suction sampler vs FISH | Intracellular trafficking, secretion, and vesicular transport |
| gene_64448 | 10,12371449 | 3,463444271 | 1,142425859 | 3,031657805 | 0,002432148 | 0,037129303 | The Moose: suction sampler vs FISH | Intracellular trafficking, secretion, and vesicular transport |
| gene_1949463 | 17,94069414 | 3,490994499 | 1,059568765 | 3,294731419 | 0,000985158 | 0,021527467 | The Moose: suction sampler vs FISH | Intracellular trafficking, secretion, and vesicular transport |
| gene_870627 | 13,43388638 | 3,521885864 | 1,158522221 | 3,03998128 | 0,002365929 | 0,03648239 | The Moose: suction sampler vs FISH | Intracellular trafficking, secretion, and vesicular transport |
| gene_386090 | 39,98792785 | 3,589258507 | 0,899434942 | 3,990570459 | 6,59E-05 | 0,003767389 | The Moose: suction sampler vs FISH | Intracellular trafficking, secretion, and vesicular transport |
| gene_1131614 | 60,75606639 | 3,601662817 | 0,709849287 | 5,073841566 | 3,90E-07 | 0,00011832 | The Moose: suction sampler vs FISH | Intracellular trafficking, secretion, and vesicular transport |
| gene_1075641 | 16,71799271 | 3,624977553 | 1,113974044 | 3,254095166 | 0,001137542 | 0,02358745 | The Moose: suction sampler vs FISH | Intracellular trafficking, secretion, and vesicular transport |
| gene_193665 | 16,99114948 | 3,668651825 | 0,977152284 | 3,754432022 | 0,000173735 | 0,007124492 | The Moose: suction sampler vs FISH | Intracellular trafficking, secretion, and vesicular transport |
| gene_1227809 | 35,36970623 | 3,763131478 | 1,062975663 | 3,540185922 | 0,000399845 | 0,012205583 | The Moose: suction sampler vs FISH | Intracellular trafficking, secretion, and vesicular transport |
| gene_484474 | 33,5060453 | 3,86928381 | 0,997995632 | 3,877054856 | 0,000105729 | 0,005173372 | The Moose: suction sampler vs FISH | Intracellular trafficking, secretion, and vesicular transport |
| gene_1130221 | 10,33231694 | 3,9033979 | 1,224148135 | 3,188664663 | 0,001429316 | 0,027061083 | The Moose: suction sampler vs FISH | Intracellular trafficking, secretion, and vesicular transport |
| gene_1847890 | 66,66970403 | 3,912196529 | 0,864558666 | 4,525079309 | 6,04E-06 | 0,00075667 | The Moose: suction sampler vs FISH | Intracellular trafficking, secretion, and vesicular transport |
| gene_1131612 | 11,9526435 | 3,933606126 | 1,172726259 | 3,354240681 | 0,000795831 | 0,018860863 | The Moose: suction sampler vs FISH | Intracellular trafficking, secretion, and vesicular transport |
| gene_778455 | 9,740655156 | 3,935587492 | 1,283007867 | 3,067469494 | 0,002158795 | 0,034702746 | The Moose: suction sampler vs FISH | Intracellular trafficking, secretion, and vesicular transport |
| gene_296384 | 34,60180286 | 3,937099117 | 0,951571646 | 4,137469977 | 3,51E-05 | 0,002458597 | The Moose: suction sampler vs FISH | Intracellular trafficking, secretion, and vesicular transport |
| gene_79089 | 18,50754017 | 4,009326321 | 1,046957303 | 3,829503179 | 0,000128402 | 0,005883033 | The Moose: suction sampler vs FISH | Intracellular trafficking, secretion, and vesicular transport |
| gene_470933 | 15,68951343 | 4,027299571 | 1,154376678 | 3,488722224 | 0,000485335 | 0,013672975 | The Moose: suction sampler vs FISH | Intracellular trafficking, secretion, and vesicular transport |
| gene_1092724 | 11,95305005 | 4,049939555 | 1,148078085 | 3,527581973 | 0,000419374 | 0,01255131 | The Moose: suction sampler vs FISH | Intracellular trafficking, secretion, and vesicular transport |
| gene_338757 | 24,11738505 | 4,072672788 | 0,980247747 | 4,154738229 | 3,26E-05 | 0,002340324 | The Moose: suction sampler vs FISH | Intracellular trafficking, secretion, and vesicular transport |
| gene_1315716 | 40,50669512 | 4,076668887 | 0,911444945 | 4,47275385 | 7,72E-06 | 0,000898783 | The Moose: suction sampler vs FISH | Intracellular trafficking, secretion, and vesicular transport |
| gene_1491520 | 14,57249632 | 4,10981283 | 1,084794169 | 3,78856464 | 0,00015152 | 0,006500233 | The Moose: suction sampler vs FISH | Intracellular trafficking, secretion, and vesicular transport |
| gene_64272 | 99,96360086 | 4,118532537 | 0,905052748 | 4,55059945 | 5,35E-06 | 0,000691693 | The Moose: suction sampler vs FISH | Intracellular trafficking, secretion, and vesicular transport |
| gene_1784231 | 19,79729846 | 4,181185308 | 0,961784418 | 4,3473207 | 1,38E-05 | 0,001307688 | The Moose: suction sampler vs FISH | Intracellular trafficking, secretion, and vesicular transport |
| gene_283081 | 7,017777958 | 4,183877846 | 1,38098584 | 3,029631243 | 0,002448525 | 0,037247704 | The Moose: suction sampler vs FISH | Intracellular trafficking, secretion, and vesicular transport |
| gene_1009326 | 16,22110217 | 4,22140124 | 1,26947114 | 3,325322732 | 0,000883162 | 0,020232022 | The Moose: suction sampler vs FISH | Intracellular trafficking, secretion, and vesicular transport |
| gene_912939 | 20,05999162 | 4,236296906 | 1,117230396 | 3,791784506 | 0,000149569 | 0,006454213 | The Moose: suction sampler vs FISH | Intracellular trafficking, secretion, and vesicular transport |
| gene_50932 | 10,61924833 | 4,245284713 | 1,469151033 | 2,889617622 | 0,003857107 | 0,048780915 | The Moose: suction sampler vs FISH | Intracellular trafficking, secretion, and vesicular transport |
| gene_1790678 | 17,68680863 | 4,36703403 | 1,113596382 | 3,921559103 | 8,80E-05 | 0,004582648 | The Moose: suction sampler vs FISH | Intracellular trafficking, secretion, and vesicular transport |
| gene_1940135 | 10,20072456 | 4,39181836 | 1,275957005 | 3,441979896 | 0,000577473 | 0,015289299 | The Moose: suction sampler vs FISH | Intracellular trafficking, secretion, and vesicular transport |
| gene_1701227 | 14,46693182 | 4,40458298 | 1,173472929 | 3,753459385 | 0,000174411 | 0,00714705 | The Moose: suction sampler vs FISH | Intracellular trafficking, secretion, and vesicular transport |
| gene_1772987 | 11,66072321 | 4,439304509 | 1,258757076 | 3,526736487 | 0,000420715 | 0,01257792 | The Moose: suction sampler vs FISH | Intracellular trafficking, secretion, and vesicular transport |
| gene_853002 | 11,77145002 | 4,50598358 | 1,157436621 | 3,893071549 | 9,90E-05 | 0,00497608 | The Moose: suction sampler vs FISH | Intracellular trafficking, secretion, and vesicular transport |
| gene_644283 | 79,38541352 | 4,517126251 | 1,241719179 | 3,637800178 | 0,000274977 | 0,009612403 | The Moose: suction sampler vs FISH | Intracellular trafficking, secretion, and vesicular transport |
| gene_2073 | 15,41130271 | 4,544472678 | 1,067220666 | 4,258231519 | 2,06E-05 | 0,001698471 | The Moose: suction sampler vs FISH | Intracellular trafficking, secretion, and vesicular transport |
| gene_283080 | 70,96071943 | 4,548923893 | 0,846091099 | 5,376399657 | 7,60E-08 | 3,96E-05 | The Moose: suction sampler vs FISH | Intracellular trafficking, secretion, and vesicular transport |

| | | | | | | | | |
|---|---|---|---|---|---|---|---|---|
| gene_232807 | 15,08759026 | 4,634107728 | 1,162786348 | 3,985347556 | 6,74E-05 | 0,003836301 | The Moose: suction sampler vs FISH | Intracellular trafficking, secretion, and vesicular transport |
| gene_1701226 | 12,63051613 | 4,688823286 | 1,113258172 | 4,211802262 | 2,53E-05 | 0,001981539 | The Moose: suction sampler vs FISH | Intracellular trafficking, secretion, and vesicular transport |
| gene_24754 | 22,82582509 | 4,745143414 | 1,171348936 | 4,051007575 | 5,10E-05 | 0,003204034 | The Moose: suction sampler vs FISH | Intracellular trafficking, secretion, and vesicular transport |
| gene_1608722 | 28,13860958 | 4,824574168 | 0,884487435 | 5,454655405 | 4,91E-08 | 2,95E-05 | The Moose: suction sampler vs FISH | Intracellular trafficking, secretion, and vesicular transport |
| gene_1065562 | 10,40784802 | 4,826420901 | 1,359688829 | 3,549651067 | 0,000385742 | 0,011918732 | The Moose: suction sampler vs FISH | Intracellular trafficking, secretion, and vesicular transport |
| gene_1509561 | 45,54252424 | 4,874957985 | 1,042832659 | 4,674726997 | 2,94E-06 | 0,000446656 | The Moose: suction sampler vs FISH | Intracellular trafficking, secretion, and vesicular transport |
| gene_1291111 | 10,13687773 | 4,897780633 | 1,302901831 | 3,759132512 | 0,000170504 | 0,007047063 | The Moose: suction sampler vs FISH | Intracellular trafficking, secretion, and vesicular transport |
| gene_1603100 | 9,693658813 | 4,909380612 | 1,329851304 | 3,691676352 | 0,000222781 | 0,008377951 | The Moose: suction sampler vs FISH | Intracellular trafficking, secretion, and vesicular transport |
| gene_1582308 | 9,274681581 | 4,936159135 | 1,388010347 | 3,556284106 | 0,000376137 | 0,011720579 | The Moose: suction sampler vs FISH | Intracellular trafficking, secretion, and vesicular transport |
| gene_435459 | 6,054240542 | 4,988765878 | 1,611098013 | 3,096500547 | 0,001958195 | 0,032745818 | The Moose: suction sampler vs FISH | Intracellular trafficking, secretion, and vesicular transport |
| gene_1126957 | 26,9128764 | 5,030908233 | 1,148819643 | 4,37919761 | 1,19E-05 | 0,001192318 | The Moose: suction sampler vs FISH | Intracellular trafficking, secretion, and vesicular transport |
| gene_1865336 | 31,0584066 | 5,045138844 | 1,115527893 | 4,52264697 | 6,11E-06 | 0,000762817 | The Moose: suction sampler vs FISH | Intracellular trafficking, secretion, and vesicular transport |
| gene_177405 | 8,031485153 | 5,097551915 | 1,467586532 | 3,473425112 | 0,000513861 | 0,014198835 | The Moose: suction sampler vs FISH | Intracellular trafficking, secretion, and vesicular transport |
| gene_1374906 | 5,27406628 | 5,206009592 | 1,737259988 | 2,996678463 | 0,002729384 | 0,039734317 | The Moose: suction sampler vs FISH | Intracellular trafficking, secretion, and vesicular transport |
| gene_700525 | 7,517315824 | 5,219609093 | 1,573944493 | 3,316259954 | 0,000912309 | 0,020601212 | The Moose: suction sampler vs FISH | Intracellular trafficking, secretion, and vesicular transport |
| gene_1029224 | 5,504457789 | 5,225633681 | 1,792782828 | 2,914816898 | 0,003558973 | 0,046527013 | The Moose: suction sampler vs FISH | Intracellular trafficking, secretion, and vesicular transport |
| gene_1601351 | 173,6049339 | 5,257235952 | 1,553629589 | 3,383841289 | 0,000714793 | 0,017575487 | The Moose: suction sampler vs FISH | Intracellular trafficking, secretion, and vesicular transport |
| gene_811069 | 6,057799378 | 5,296435156 | 1,73436734 | 3,053813937 | 0,002259522 | 0,035527361 | The Moose: suction sampler vs FISH | Intracellular trafficking, secretion, and vesicular transport |
| gene_824157 | 8,324149002 | 5,359347844 | 1,574389553 | 3,404079907 | 0,000663873 | 0,016737807 | The Moose: suction sampler vs FISH | Intracellular trafficking, secretion, and vesicular transport |
| gene_677437 | 5,806559927 | 5,385261192 | 1,844372442 | 2,919833906 | 0,00350218 | 0,046077151 | The Moose: suction sampler vs FISH | Intracellular trafficking, secretion, and vesicular transport |
| gene_1131611 | 8,486731254 | 5,501324005 | 1,580298076 | 3,481193889 | 0,000499184 | 0,013957982 | The Moose: suction sampler vs FISH | Intracellular trafficking, secretion, and vesicular transport |
| gene_1940133 | 6,246391011 | 5,580973583 | 1,727259004 | 3,231115641 | 0,00123308 | 0,024687463 | The Moose: suction sampler vs FISH | Intracellular trafficking, secretion, and vesicular transport |
| gene_1365021 | 3,356504482 | 5,613344414 | 1,941887271 | 2,890664405 | 0,003844284 | 0,04871339 | The Moose: suction sampler vs FISH | Intracellular trafficking, secretion, and vesicular transport |
| gene_413425 | 16,23962478 | 5,657672273 | 1,567281402 | 3,609863721 | 0,000306358 | 0,010316578 | The Moose: suction sampler vs FISH | Intracellular trafficking, secretion, and vesicular transport |
| gene_1713827 | 7,628374491 | 5,673327023 | 1,647773723 | 3,443025546 | 0,000575245 | 0,015257613 | The Moose: suction sampler vs FISH | Intracellular trafficking, secretion, and vesicular transport |
| gene_1701079 | 20,04751335 | 5,734098442 | 1,340892624 | 4,276329318 | 1,90E-05 | 0,001596631 | The Moose: suction sampler vs FISH | Intracellular trafficking, secretion, and vesicular transport |
| gene_1800291 | 2,990755087 | 5,755048409 | 1,911131805 | 3,011329932 | 0,002601061 | 0,038749472 | The Moose: suction sampler vs FISH | Intracellular trafficking, secretion, and vesicular transport |
| gene_925798 | 10,33333931 | 5,801496162 | 1,435544407 | 4,041321282 | 5,32E-05 | 0,003277638 | The Moose: suction sampler vs FISH | Intracellular trafficking, secretion, and vesicular transport |
| gene_1012366 | 4,524796602 | 5,847238936 | 2,010179811 | 2,908813879 | 0,003628028 | 0,047077827 | The Moose: suction sampler vs FISH | Intracellular trafficking, secretion, and vesicular transport |
| gene_1348525 | 5,362100523 | 5,898093134 | 1,721406761 | 3,426321581 | 0,000611816 | 0,01587145 | The Moose: suction sampler vs FISH | Intracellular trafficking, secretion, and vesicular transport |
| gene_581243 | 6,150182248 | 5,924847307 | 1,752413468 | 3,380964262 | 0,000722319 | 0,017704153 | The Moose: suction sampler vs FISH | Intracellular trafficking, secretion, and vesicular transport |
| gene_1844714 | 5,64816653 | 5,997510424 | 1,826174958 | 3,284192677 | 0,00102275 | 0,022039457 | The Moose: suction sampler vs FISH | Intracellular trafficking, secretion, and vesicular transport |
| gene_814453 | 8,959057198 | 6,050220893 | 1,807860245 | 3,346619801 | 0,000818033 | 0,019213448 | The Moose: suction sampler vs FISH | Intracellular trafficking, secretion, and vesicular transport |
| gene_853000 | 4,248018933 | 6,16274258 | 2,048458096 | 3,008478715 | 0,002625592 | 0,038881355 | The Moose: suction sampler vs FISH | Intracellular trafficking, secretion, and vesicular transport |
| gene_1602684 | 3,762941818 | 6,212284761 | 1,813934726 | 3,42475651 | 0,000615351 | 0,015913531 | The Moose: suction sampler vs FISH | Intracellular trafficking, secretion, and vesicular transport |
| gene_469882 | 3,556210105 | 6,215229592 | 1,971212752 | 3,152997862 | 0,00161603 | 0,029225655 | The Moose: suction sampler vs FISH | Intracellular trafficking, secretion, and vesicular transport |
| gene_1522941 | 2,684171904 | 6,248323121 | 2,150154405 | 2,905988104 | 0,003660954 | 0,047379514 | The Moose: suction sampler vs FISH | Intracellular trafficking, secretion, and vesicular transport |
| gene_1698993 | 3,978892773 | 6,287849794 | 2,0866146 | 3,013421738 | 0,002583197 | 0,03856156 | The Moose: suction sampler vs FISH | Intracellular trafficking, secretion, and vesicular transport |
| gene_1890798 | 12,18302858 | 6,377796405 | 1,545907515 | 4,125600234 | 3,70E-05 | 0,002550018 | The Moose: suction sampler vs FISH | Intracellular trafficking, secretion, and vesicular transport |
| gene_271687 | 9,463533655 | 6,423669446 | 1,679981955 | 3,823653835 | 0,000131488 | 0,005963296 | The Moose: suction sampler vs FISH | Intracellular trafficking, secretion, and vesicular transport |
| gene_1473807 | 5,773344945 | 6,449036679 | 1,712891783 | 3,764999485 | 0,000166549 | 0,006958033 | The Moose: suction sampler vs FISH | Intracellular trafficking, secretion, and vesicular transport |
| gene_1802917 | 44,11889064 | 6,51992718 | 1,159537777 | 5,622867415 | 1,88E-08 | 1,41E-05 | The Moose: suction sampler vs FISH | Intracellular trafficking, secretion, and vesicular transport |
| gene_784599 | 8,415327463 | 6,779266322 | 1,715594818 | 3,951554441 | 7,76E-05 | 0,004232585 | The Moose: suction sampler vs FISH | Intracellular trafficking, secretion, and vesicular transport |
| gene_1653691 | 4,363874744 | 6,78390371 | 1,950301748 | 3,478386724 | 0,000504442 | 0,014044726 | The Moose: suction sampler vs FISH | Intracellular trafficking, secretion, and vesicular transport |
| gene_701753 | 4,178508423 | 6,788028881 | 1,921181566 | 3,533257346 | 0,000410473 | 0,012380188 | The Moose: suction sampler vs FISH | Intracellular trafficking, secretion, and vesicular transport |
| gene_1091231 | 16,32197966 | 6,820809304 | 1,352876143 | 5,041710093 | 4,61E-07 | 0,00013237 | The Moose: suction sampler vs FISH | Intracellular trafficking, secretion, and vesicular transport |
| gene_1802916 | 3,890542777 | 6,87958717 | 2,195897774 | 3,132926883 | 0,001730726 | 0,030426115 | The Moose: suction sampler vs FISH | Intracellular trafficking, secretion, and vesicular transport |
| gene_917238 | 17,14347825 | 7,044002348 | 1,456822938 | 4,835180834 | 1,33E-06 | 0,000271496 | The Moose: suction sampler vs FISH | Intracellular trafficking, secretion, and vesicular transport |
| gene_1487341 | 4,018126211 | 7,107004956 | 2,161410742 | 3,288132523 | 0,001008544 | 0,021818264 | The Moose: suction sampler vs FISH | Intracellular trafficking, secretion, and vesicular transport |
| gene_1698992 | 6,036815878 | 7,251102114 | 1,872375066 | 3,872676071 | 0,000107647 | 0,005228918 | The Moose: suction sampler vs FISH | Intracellular trafficking, secretion, and vesicular transport |
| gene_903341 | 6,568696756 | 7,397159239 | 1,852615552 | 3,992819357 | 6,53E-05 | 0,003741521 | The Moose: suction sampler vs FISH | Intracellular trafficking, secretion, and vesicular transport |
| gene_912940 | 10,78473748 | 7,550752154 | 1,670455041 | 4,520176819 | 6,18E-06 | 0,000769156 | The Moose: suction sampler vs FISH | Intracellular trafficking, secretion, and vesicular transport |
| gene_761756 | 7,895014569 | 7,938578973 | 1,765934123 | 4,495399274 | 6,94E-06 | 0,00083664 | The Moose: suction sampler vs FISH | Intracellular trafficking, secretion, and vesicular transport |
| gene_1092830 | 10,31731637 | 8,077409304 | 1,569701521 | 5,145824986 | 2,66E-07 | 8,97E-05 | The Moose: suction sampler vs FISH | Intracellular trafficking, secretion, and vesicular transport |
| gene_1091232 | 11,02112298 | 8,122998259 | 1,882521487 | 4,314956464 | 1,60E-05 | 0,001435048 | The Moose: suction sampler vs FISH | Intracellular trafficking, secretion, and vesicular transport |
| gene_1455601 | 5,627666411 | -6,193925597 | 1,708081944 | -3,626246164 | 0,000287571 | 0,009911154 | The Moose: suction sampler vs FISH | Lipid transport and metabolism |
| gene_1708262 | 13,35605016 | -5,799269606 | 1,399799541 | -4,142928639 | 3,43E-05 | 0,002430907 | The Moose: suction sampler vs FISH | Lipid transport and metabolism |
| gene_336959 | 14,37352621 | -3,988745611 | 1,309738472 | -3,045451972 | 0,002323309 | 0,036082789 | The Moose: suction sampler vs FISH | Lipid transport and metabolism |
| gene_1065095 | 26,55906809 | -3,559404453 | 0,95239782 | -3,737308484 | 0,000186001 | 0,007453932 | The Moose: suction sampler vs FISH | Lipid transport and metabolism |
| gene_720381 | 28,62121192 | -3,049963228 | 0,943427646 | -3,232853353 | 0,001225605 | 0,024594134 | The Moose: suction sampler vs FISH | Lipid transport and metabolism |
| gene_1235155 | 52,05203708 | -2,475654067 | 0,709577144 | -3,488914611 | 0,000484986 | 0,013668381 | The Moose: suction sampler vs FISH | Lipid transport and metabolism |
| gene_98697 | 157,11925 | -2,320786718 | 0,692259784 | -3,352479477 | 0,000800912 | 0,0189324 | The Moose: suction sampler vs FISH | Lipid transport and metabolism |
| gene_468420 | 39,39575472 | -2,282541558 | 0,737328499 | -3,095691489 | 0,001963545 | 0,032800264 | The Moose: suction sampler vs FISH | Lipid transport and metabolism |
| gene_208390 | 88,36216283 | -2,231741582 | 0,720391679 | -3,097955801 | 0,001948605 | 0,032644267 | The Moose: suction sampler vs FISH | Lipid transport and metabolism |
| gene_340516 | 3011,979429 | -2,200163773 | 0,546899778 | -4,022974337 | 5,75E-05 | 0,003459606 | The Moose: suction sampler vs FISH | Lipid transport and metabolism |

| | | | | | | | | |
|---|---|---|---|---|---|---|---|---|
| gene_95464 | 201,0933158 | -2,171761757 | 0,712554044 | -3,047855491 | 0,002304807 | 0,035916948 | The Moose: suction sampler vs FISH | Lipid transport and metabolism |
| gene_873930 | 8084,12458 | -2,12148993 | 0,552435222 | -3,840251028 | 0,000122909 | 0,005746034 | The Moose: suction sampler vs FISH | Lipid transport and metabolism |
| gene_574008 | 46,77478291 | 2,093116876 | 0,709037494 | 2,952053866 | 0,003156679 | 0,043408024 | The Moose: suction sampler vs FISH | Lipid transport and metabolism |
| gene_962740 | 45,6253991 | 2,2160012 | 0,752982449 | 2,942965275 | 0,003250849 | 0,044160858 | The Moose: suction sampler vs FISH | Lipid transport and metabolism |
| gene_572542 | 87,06681389 | 2,531156263 | 0,690066031 | 3,667991392 | 0,000244463 | 0,008927261 | The Moose: suction sampler vs FISH | Lipid transport and metabolism |
| gene_1043803 | 32,97961841 | 2,539156722 | 0,76528261 | 3,317933387 | 0,000906861 | 0,020556918 | The Moose: suction sampler vs FISH | Lipid transport and metabolism |
| gene_1852105 | 43,35124072 | 2,677435534 | 0,859632579 | 3,114627806 | 0,001841771 | 0,031612271 | The Moose: suction sampler vs FISH | Lipid transport and metabolism |
| gene_1177903 | 15,46504952 | 2,857575234 | 0,976715004 | 2,92570015 | 0,00343682 | 0,045612081 | The Moose: suction sampler vs FISH | Lipid transport and metabolism |
| gene_1597300 | 66,56392485 | 2,862461272 | 0,771807449 | 3,708776427 | 0,000208263 | 0,008033557 | The Moose: suction sampler vs FISH | Lipid transport and metabolism |
| gene_575570 | 17,99290906 | 3,072974689 | 1,044432924 | 2,942242263 | 0,003258449 | 0,044186999 | The Moose: suction sampler vs FISH | Lipid transport and metabolism |
| gene_604450 | 35,15399139 | 3,129492079 | 0,791267908 | 3,955034756 | 7,65E-05 | 0,004194372 | The Moose: suction sampler vs FISH | Lipid transport and metabolism |
| gene_1796572 | 17,48741621 | 3,2275022 | 1,048692656 | 3,077643562 | 0,002086443 | 0,034075699 | The Moose: suction sampler vs FISH | Lipid transport and metabolism |
| gene_247786 | 10,13367778 | 3,388890785 | 0,996728152 | 3,400015116 | 0,000673821 | 0,01689814 | The Moose: suction sampler vs FISH | Lipid transport and metabolism |
| gene_570642 | 28,237957 | 3,460417497 | 1,193314849 | 2,899836115 | 0,003733578 | 0,047956041 | The Moose: suction sampler vs FISH | Lipid transport and metabolism |
| gene_1612924 | 27,69075088 | 3,500949534 | 0,944596601 | 3,706290631 | 0,000210317 | 0,008061967 | The Moose: suction sampler vs FISH | Lipid transport and metabolism |
| gene_469163 | 239,9062155 | 3,563654777 | 0,754761017 | 4,721567086 | 2,34E-06 | 0,000388006 | The Moose: suction sampler vs FISH | Lipid transport and metabolism |
| gene_469164 | 55,95968433 | 3,574213218 | 0,878567517 | 4,068228278 | 4,74E-05 | 0,003058801 | The Moose: suction sampler vs FISH | Lipid transport and metabolism |
| gene_1652050 | 13,58238913 | 3,602279683 | 1,111940421 | 3,23963372 | 0,001196833 | 0,024295584 | The Moose: suction sampler vs FISH | Lipid transport and metabolism |
| gene_915849 | 17,63810235 | 3,627492138 | 1,110801142 | 3,265653951 | 0,001092116 | 0,023035748 | The Moose: suction sampler vs FISH | Lipid transport and metabolism |
| gene_1225754 | 8,392903063 | 3,629060326 | 1,224245455 | 2,964324117 | 0,003033486 | 0,042450764 | The Moose: suction sampler vs FISH | Lipid transport and metabolism |
| gene_979954 | 46,51583381 | 3,648847003 | 0,854790226 | 4,26870464 | 1,97E-05 | 0,001633498 | The Moose: suction sampler vs FISH | Lipid transport and metabolism |
| gene_347444 | 20,64135363 | 3,66408512 | 0,854441082 | 4,288282948 | 1,80E-05 | 0,001552169 | The Moose: suction sampler vs FISH | Lipid transport and metabolism |
| gene_1196138 | 17,79697157 | 3,716932458 | 1,095434421 | 3,393112712 | 0,000691032 | 0,017165926 | The Moose: suction sampler vs FISH | Lipid transport and metabolism |
| gene_968113 | 8,026707591 | 3,788418558 | 1,143483466 | 3,313050578 | 0,000922843 | 0,020720941 | The Moose: suction sampler vs FISH | Lipid transport and metabolism |
| gene_1330747 | 21,1688993 | 3,804864304 | 0,950489456 | 4,003057875 | 6,25E-05 | 0,003644795 | The Moose: suction sampler vs FISH | Lipid transport and metabolism |
| gene_717820 | 25,07379286 | 3,904890606 | 1,064060303 | 3,669801978 | 0,000242738 | 0,008869614 | The Moose: suction sampler vs FISH | Lipid transport and metabolism |
| gene_1089953 | 33,73559198 | 3,913838635 | 0,81119303 | 4,824793222 | 1,40E-06 | 0,000278767 | The Moose: suction sampler vs FISH | Lipid transport and metabolism |
| gene_1325059 | 15,19588292 | 3,974517868 | 1,194541875 | 3,327231931 | 0,000877133 | 0,020150472 | The Moose: suction sampler vs FISH | Lipid transport and metabolism |
| gene_495589 | 18,87632065 | 4,007070241 | 1,264639146 | 3,168548318 | 0,001532023 | 0,028285424 | The Moose: suction sampler vs FISH | Lipid transport and metabolism |
| gene_1786022 | 65,03865472 | 4,044671127 | 0,971477132 | 4,163423918 | 3,14E-05 | 0,002273029 | The Moose: suction sampler vs FISH | Lipid transport and metabolism |
| gene_1007746 | 148,2524186 | 4,046061007 | 0,903393245 | 4,478737281 | 7,51E-06 | 0,000886605 | The Moose: suction sampler vs FISH | Lipid transport and metabolism |
| gene_413304 | 66,60534254 | 4,047565237 | 0,846949623 | 4,778991721 | 1,76E-06 | 0,000321873 | The Moose: suction sampler vs FISH | Lipid transport and metabolism |
| gene_531692 | 14,42070981 | 4,082843861 | 1,150262718 | 3,549488128 | 0,000385981 | 0,011921095 | The Moose: suction sampler vs FISH | Lipid transport and metabolism |
| gene_1553586 | 12,35652216 | 4,096637326 | 1,32025759 | 3,102907614 | 0,001916295 | 0,032317394 | The Moose: suction sampler vs FISH | Lipid transport and metabolism |
| gene_1775845 | 35,48529619 | 4,102620993 | 0,855945094 | 4,793088973 | 1,64E-06 | 0,000308493 | The Moose: suction sampler vs FISH | Lipid transport and metabolism |
| gene_1762843 | 12,61276338 | 4,135201486 | 1,131598543 | 3,654300822 | 0,000257884 | 0,009225657 | The Moose: suction sampler vs FISH | Lipid transport and metabolism |
| gene_1786021 | 29,92229691 | 4,162997361 | 0,821507474 | 5,067510027 | 4,03E-07 | 0,000119364 | The Moose: suction sampler vs FISH | Lipid transport and metabolism |
| gene_1778697 | 9,578156892 | 4,246005211 | 1,428646371 | 2,972047734 | 0,002958207 | 0,041777252 | The Moose: suction sampler vs FISH | Lipid transport and metabolism |
| gene_515634 | 26,39996235 | 4,280411665 | 1,019189076 | 4,199820982 | 2,67E-05 | 0,002056507 | The Moose: suction sampler vs FISH | Lipid transport and metabolism |
| gene_1482001 | 10,59227823 | 4,321084884 | 1,242306528 | 3,478275923 | 0,00050465 | 0,014044726 | The Moose: suction sampler vs FISH | Lipid transport and metabolism |
| gene_1862024 | 10,1945195 | 4,327808088 | 1,39148488 | 3,110208491 | 0,001869553 | 0,031899951 | The Moose: suction sampler vs FISH | Lipid transport and metabolism |
| gene_140552 | 7,149813051 | 4,568976237 | 1,544490593 | 2,958241545 | 0,003093996 | 0,042988748 | The Moose: suction sampler vs FISH | Lipid transport and metabolism |
| gene_1030266 | 17,78974559 | 4,606667303 | 1,224892809 | 3,760873824 | 0,000169321 | 0,007025858 | The Moose: suction sampler vs FISH | Lipid transport and metabolism |
| gene_596381 | 15,97147592 | 4,667617653 | 1,262840247 | 3,696126777 | 0,000218914 | 0,008266383 | The Moose: suction sampler vs FISH | Lipid transport and metabolism |
| gene_620403 | 6,817178937 | 4,756702988 | 1,552258579 | 3,06437539 | 0,00218125 | 0,034899494 | The Moose: suction sampler vs FISH | Lipid transport and metabolism |
| gene_1196137 | 22,50995617 | 4,801293747 | 1,171698586 | 4,097720867 | 4,17E-05 | 0,002788357 | The Moose: suction sampler vs FISH | Lipid transport and metabolism |
| gene_1024047 | 5,624655662 | 4,994296169 | 1,665310599 | 2,999017824 | 0,002708515 | 0,039532364 | The Moose: suction sampler vs FISH | Lipid transport and metabolism |
| gene_781199 | 7,759173956 | 5,121197446 | 1,656249278 | 3,092045089 | 0,001987827 | 0,03306067 | The Moose: suction sampler vs FISH | Lipid transport and metabolism |
| gene_661726 | 5,68882038 | 5,140376396 | 1,580825081 | 3,251704733 | 0,001147151 | 0,023706063 | The Moose: suction sampler vs FISH | Lipid transport and metabolism |
| gene_788373 | 4,49757401 | 5,512651326 | 1,888773678 | 2,918640486 | 0,003515614 | 0,046149113 | The Moose: suction sampler vs FISH | Lipid transport and metabolism |
| gene_337016 | 8,956027977 | 5,675121906 | 1,547949703 | 3,666218543 | 0,000246164 | 0,008947252 | The Moose: suction sampler vs FISH | Lipid transport and metabolism |
| gene_1377962 | 3,709683973 | 5,761227749 | 1,978536487 | 2,911863282 | 0,003592799 | 0,046815421 | The Moose: suction sampler vs FISH | Lipid transport and metabolism |
| gene_1001641 | 5,841520772 | 5,831916797 | 1,783488443 | 3,269949307 | 0,001075668 | 0,02279521 | The Moose: suction sampler vs FISH | Lipid transport and metabolism |
| gene_1367910 | 5,42805468 | 5,86440181 | 1,814487072 | 3,231988753 | 0,001229319 | 0,024661931 | The Moose: suction sampler vs FISH | Lipid transport and metabolism |
| gene_306894 | 5,917861832 | 5,914298916 | 1,83577586 | 3,221689011 | 0,001274374 | 0,025270955 | The Moose: suction sampler vs FISH | Lipid transport and metabolism |
| gene_1823522 | 3,85514094 | 5,966334672 | 1,920106575 | 3,107293497 | 0,001888088 | 0,032069995 | The Moose: suction sampler vs FISH | Lipid transport and metabolism |
| gene_425324 | 6,750347719 | 6,003225023 | 1,768785284 | 3,393981777 | 0,000688843 | 0,017144038 | The Moose: suction sampler vs FISH | Lipid transport and metabolism |
| gene_1780066 | 7,641863493 | 6,088527804 | 1,743805672 | 3,491517376 | 0,000480285 | 0,013577723 | The Moose: suction sampler vs FISH | Lipid transport and metabolism |
| gene_846207 | 5,865207165 | 6,137091357 | 1,696627385 | 3,617229929 | 0,000297773 | 0,01015477 | The Moose: suction sampler vs FISH | Lipid transport and metabolism |
| gene_1868236 | 5,818241322 | 6,237440307 | 2,025514106 | 3,07943563 | 0,002073932 | 0,033939377 | The Moose: suction sampler vs FISH | Lipid transport and metabolism |
| gene_1069852 | 8,634158858 | 6,254564384 | 1,625380318 | 3,84806209 | 0,000119056 | 0,005615966 | The Moose: suction sampler vs FISH | Lipid transport and metabolism |
| gene_1646388 | 4,647328086 | 6,370563366 | 1,896796132 | 3,358591501 | 0,000783408 | 0,018693108 | The Moose: suction sampler vs FISH | Lipid transport and metabolism |
| gene_915976 | 11,32597287 | 6,403368452 | 1,537764679 | 4,164075649 | 3,13E-05 | 0,002268786 | The Moose: suction sampler vs FISH | Lipid transport and metabolism |
| gene_117548 | 7,733131161 | 6,439406511 | 1,737232475 | 3,70670397 | 0,000209974 | 0,008053302 | The Moose: suction sampler vs FISH | Lipid transport and metabolism |
| gene_404071 | 4,41062441 | 6,636866836 | 2,121788683 | 3,127958448 | 0,001760251 | 0,030719076 | The Moose: suction sampler vs FISH | Lipid transport and metabolism |

| | | | | | | | | |
|---|---|---|---|---|---|---|---|---|
| gene_1402134 | 5,911126107 | 7,045120014 | 1,695060575 | 4,156264453 | 3,23E-05 | 0,002327029 | The Moose: suction sampler vs FISH | Lipid transport and metabolism |
| gene_276135 | 6,771399267 | 7,925910131 | 1,802224043 | 4,397849514 | 1,09E-05 | 0,00112776 | The Moose: suction sampler vs FISH | Lipid transport and metabolism |
| gene_185966 | 56,47093681 | -3,529055777 | 0,830341926 | -4,250123552 | 2,14E-05 | 0,001740321 | The Moose: suction sampler vs FISH | Mobilome: prophages, transposons |
| gene_652921 | 23,54655412 | -3,465377194 | 1,120626205 | -3,092357807 | 0,001985733 | 0,03304831 | The Moose: suction sampler vs FISH | Mobilome: prophages, transposons |
| gene_280113 | 185,6226673 | -2,097292598 | 0,586773146 | -3,574281833 | 0,00035119 | 0,011199034 | The Moose: suction sampler vs FISH | Mobilome: prophages, transposons |
| gene_53407 | 149,9613484 | 2,106898395 | 0,615536806 | 3,422863387 | 0,000619652 | 0,01598537 | The Moose: suction sampler vs FISH | Mobilome: prophages, transposons |
| gene_159876 | 46,54900269 | 2,397299024 | 0,778330974 | 3,080050907 | 0,002069652 | 0,033907116 | The Moose: suction sampler vs FISH | Mobilome: prophages, transposons |
| gene_1113762 | 27,20454484 | 2,462536932 | 0,815175317 | 3,020867881 | 0,002520513 | 0,037965358 | The Moose: suction sampler vs FISH | Mobilome: prophages, transposons |
| gene_1054856 | 28,90111336 | 2,67814107 | 0,923737459 | 2,899244848 | 0,003740627 | 0,048009067 | The Moose: suction sampler vs FISH | Mobilome: prophages, transposons |
| gene_392742 | 32,46429661 | 2,70511548 | 0,836186487 | 3,23506242 | 0,001216161 | 0,024511793 | The Moose: suction sampler vs FISH | Mobilome: prophages, transposons |
| gene_412934 | 23,61978299 | 2,737496633 | 0,919504788 | 2,977142337 | 0,002909489 | 0,041340185 | The Moose: suction sampler vs FISH | Mobilome: prophages, transposons |
| gene_485278 | 27,4698462 | 2,764602873 | 0,873414428 | 3,165281891 | 0,001549328 | 0,028426274 | The Moose: suction sampler vs FISH | Mobilome: prophages, transposons |
| gene_1794099 | 13,45943227 | 3,403880906 | 1,122024712 | 3,033695131 | 0,002415784 | 0,036956316 | The Moose: suction sampler vs FISH | Mobilome: prophages, transposons |
| gene_180242 | 71,65762819 | 3,542436911 | 0,739441514 | 4,790692494 | 1,66E-06 | 0,000309039 | The Moose: suction sampler vs FISH | Mobilome: prophages, transposons |
| gene_422578 | 9,089026039 | 3,58163524 | 1,212358489 | 2,954270763 | 0,003134089 | 0,043275647 | The Moose: suction sampler vs FISH | Mobilome: prophages, transposons |
| gene_1302353 | 12,53404523 | 4,166183339 | 1,149452911 | 3,624492399 | 0,00028953 | 0,009951611 | The Moose: suction sampler vs FISH | Mobilome: prophages, transposons |
| gene_1882836 | 18,9543498 | 4,299151862 | 1,314155259 | 3,271418527 | 0,001070094 | 0,022715418 | The Moose: suction sampler vs FISH | Mobilome: prophages, transposons |
| gene_1319195 | 12,87998485 | 4,548741552 | 1,366713962 | 3,328232299 | 0,00087399 | 0,020098697 | The Moose: suction sampler vs FISH | Mobilome: prophages, transposons |
| gene_291620 | 6,424156999 | 4,714446894 | 1,58036415 | 2,983139611 | 0,002853078 | 0,040907188 | The Moose: suction sampler vs FISH | Mobilome: prophages, transposons |
| gene_123274 | 42,18344676 | 5,056806436 | 1,242546106 | 4,069713317 | 4,71E-05 | 0,003051307 | The Moose: suction sampler vs FISH | Mobilome: prophages, transposons |
| gene_499871 | 5,331338746 | 5,079419876 | 1,551457512 | 3,273966472 | 0,001060492 | 0,022622083 | The Moose: suction sampler vs FISH | Mobilome: prophages, transposons |
| gene_875863 | 7,929492708 | 5,097034926 | 1,577287864 | 3,231518509 | 0,001231343 | 0,024681166 | The Moose: suction sampler vs FISH | Mobilome: prophages, transposons |
| gene_188354 | 12,03691731 | 5,64765343 | 1,678419209 | 3,364864629 | 0,000765812 | 0,01840749 | The Moose: suction sampler vs FISH | Mobilome: prophages, transposons |
| gene_749811 | 3,157797922 | 5,896730445 | 2,036107675 | 2,896079867 | 0,003778562 | 0,048229356 | The Moose: suction sampler vs FISH | Mobilome: prophages, transposons |
| gene_415184 | 4,480787623 | 5,935386379 | 2,040533008 | 2,908743135 | 0,003628849 | 0,047079009 | The Moose: suction sampler vs FISH | Mobilome: prophages, transposons |
| gene_1715312 | 7,210395957 | 6,005579063 | 1,852106796 | 3,242566291 | 0,001184584 | 0,024140334 | The Moose: suction sampler vs FISH | Mobilome: prophages, transposons |
| gene_384754 | 4,088737443 | 6,054035242 | 2,101787111 | 2,880422671 | 0,003971424 | 0,049588781 | The Moose: suction sampler vs FISH | Mobilome: prophages, transposons |
| gene_285394 | 4,622335041 | 6,23761877 | 2,098034751 | 2,973076955 | 0,002948305 | 0,041690887 | The Moose: suction sampler vs FISH | Mobilome: prophages, transposons |
| gene_1925690 | 3,921342495 | 6,246030538 | 2,102208349 | 2,971175783 | 0,002966619 | 0,041835364 | The Moose: suction sampler vs FISH | Mobilome: prophages, transposons |
| gene_802702 | 6,868741386 | 6,255830085 | 1,889276796 | 3,311230042 | 0,000928868 | 0,020815977 | The Moose: suction sampler vs FISH | Mobilome: prophages, transposons |
| gene_1135325 | 7,233351665 | 6,38686504 | 1,734984889 | 3,681222285 | 0,000232119 | 0,008618779 | The Moose: suction sampler vs FISH | Mobilome: prophages, transposons |
| gene_223993 | 3,995674486 | 6,432517986 | 1,852290946 | 3,47273629 | 0,000515181 | 0,014213933 | The Moose: suction sampler vs FISH | Mobilome: prophages, transposons |
| gene_186497 | 4,061641258 | 6,442297808 | 1,891758113 | 3,40545536 | 0,000660538 | 0,016699904 | The Moose: suction sampler vs FISH | Mobilome: prophages, transposons |
| gene_429382 | 4,261492604 | 6,603097681 | 1,934461052 | 3,413404304 | 0,000641567 | 0,016372438 | The Moose: suction sampler vs FISH | Mobilome: prophages, transposons |
| gene_784155 | 4,091104836 | 7,330338933 | 2,201081941 | 3,330334413 | 0,000867417 | 0,020014914 | The Moose: suction sampler vs FISH | Mobilome: prophages, transposons |
| gene_123273 | 10,43528552 | 8,523259116 | 2,756178195 | 3,092419471 | 0,001985321 | 0,03304831 | The Moose: suction sampler vs FISH | Mobilome: prophages, transposons |
| gene_1000322 | 189,5399377 | 2,279306689 | 0,699155421 | 3,26008584 | 0,001113785 | 0,023278867 | The Moose: suction sampler vs FISH | NA |
| gene_1000589 | 3,607898878 | 6,290073792 | 1,982006326 | 3,17358916 | 0,001505666 | 0,027998858 | The Moose: suction sampler vs FISH | NA |
| gene_1000691 | 20,68727339 | 3,48144818 | 1,016112721 | 3,426242098 | 0,000611995 | 0,01587145 | The Moose: suction sampler vs FISH | NA |
| gene_100120 | 27,48837135 | 3,071301987 | 1,021825096 | 3,005702246 | 0,002649683 | 0,039038312 | The Moose: suction sampler vs FISH | NA |
| gene_1001231 | 9,000152773 | 3,904943681 | 1,345848608 | 2,901473209 | 0,003714125 | 0,047814888 | The Moose: suction sampler vs FISH | NA |
| gene_1001642 | 6,163697973 | 6,844135737 | 1,949331233 | 3,511017328 | 0,000446395 | 0,01301012 | The Moose: suction sampler vs FISH | NA |
| gene_1001707 | 9,542521872 | 4,625471252 | 1,30494208 | 3,544579735 | 0,00039324 | 0,012074197 | The Moose: suction sampler vs FISH | NA |
| gene_1001717 | 4,787815928 | 6,351165872 | 1,873011737 | 3,390884182 | 0,000696675 | 0,017245474 | The Moose: suction sampler vs FISH | NA |
| gene_1001980 | 100,3628327 | 3,722776114 | 0,771658396 | 4,824383603 | 1,40E-06 | 0,000278767 | The Moose: suction sampler vs FISH | NA |
| gene_1002186 | 57,77228916 | 2,967982761 | 0,89805457 | 3,304902463 | 0,000950095 | 0,021079479 | The Moose: suction sampler vs FISH | NA |
| gene_1003053 | 13,43198388 | 8,190507462 | 1,915169157 | 4,276649628 | 1,90E-05 | 0,001596631 | The Moose: suction sampler vs FISH | NA |
| gene_1004411 | 25,08187078 | 3,814925811 | 0,990918622 | 3,849888101 | 0,000118172 | 0,005581433 | The Moose: suction sampler vs FISH | NA |
| gene_1004456 | 21,19558259 | 2,924010746 | 0,947299731 | 3,086679591 | 0,002024056 | 0,033413536 | The Moose: suction sampler vs FISH | NA |
| gene_1004895 | 182,2999916 | 2,873104338 | 0,880504942 | 3,263018978 | 0,001102321 | 0,023164519 | The Moose: suction sampler vs FISH | NA |
| gene_1005055 | 4,947600169 | 5,912518435 | 1,887002849 | 3,133285378 | 0,001728613 | 0,030426115 | The Moose: suction sampler vs FISH | NA |
| gene_1006832 | 169,8398822 | 2,425682654 | 0,736267174 | 3,294568521 | 0,000985729 | 0,021527467 | The Moose: suction sampler vs FISH | NA |
| gene_1006834 | 114,5398262 | 2,38764839 | 0,811756667 | 2,941335116 | 0,003268008 | 0,044267582 | The Moose: suction sampler vs FISH | NA |
| gene_1006837 | 92,84357531 | 2,319731762 | 0,786454055 | 2,949608749 | 0,003181766 | 0,043646766 | The Moose: suction sampler vs FISH | NA |
| gene_1006838 | 42,47476886 | 2,722984776 | 0,876298325 | 3,107371882 | 0,001887588 | 0,032068908 | The Moose: suction sampler vs FISH | NA |
| gene_1007521 | 34,1820148 | 3,205539552 | 0,827127097 | 3,875510261 | 0,000106401 | 0,005195916 | The Moose: suction sampler vs FISH | NA |
| gene_100792 | 6,066297471 | 6,477744208 | 2,144669084 | 3,020393336 | 0,002524466 | 0,037994269 | The Moose: suction sampler vs FISH | NA |
| gene_1008050 | 20,67833912 | 3,020539762 | 0,947662806 | 3,187357088 | 0,001435794 | 0,027140573 | The Moose: suction sampler vs FISH | NA |
| gene_1010413 | 4,480610758 | 6,320561415 | 1,772852798 | 3,565192453 | 0,000363589 | 0,01147787 | The Moose: suction sampler vs FISH | NA |
| gene_1010873 | 3,418381323 | 5,523220321 | 1,875624394 | 2,944736878 | 0,003232294 | 0,044027419 | The Moose: suction sampler vs FISH | NA |
| gene_1011438 | 3,703670419 | 6,257301963 | 1,834892115 | 3,410174315 | 0,000649214 | 0,016493082 | The Moose: suction sampler vs FISH | NA |
| gene_1012781 | 18,19088842 | 3,145284975 | 0,936133932 | 3,359866432 | 0,000779802 | 0,018631272 | The Moose: suction sampler vs FISH | NA |
| gene_1012782 | 24,10443942 | 4,320191874 | 1,21790459 | 3,547233428 | 0,000389299 | 0,011988303 | The Moose: suction sampler vs FISH | NA |
| gene_1013460 | 17,97074229 | 3,680909726 | 1,069128222 | 3,442907643 | 0,000575496 | 0,015257613 | The Moose: suction sampler vs FISH | NA |
| gene_1013461 | 5,088157029 | 6,546285025 | 2,177361058 | 3,006522506 | 0,002642545 | 0,03898959 | The Moose: suction sampler vs FISH | NA |

| | | | | | | | | |
|---|---|---|---|---|---|---|---|---|
| gene_1013462 | 5,036970748 | 6,038176871 | 1,948503341 | 3,098879403 | 0,001942541 | 0,032613671 | The Moose: suction sampler vs FISH | NA |
| gene_1013881 | 2,81039515 | 5,932329587 | 2,030188732 | 2,922058178 | 0,003477266 | 0,045883541 | The Moose: suction sampler vs FISH | NA |
| gene_1014264 | 9,453654646 | 4,007294077 | 1,332530142 | 3,007282124 | 0,00263595 | 0,038961028 | The Moose: suction sampler vs FISH | NA |
| gene_1016312 | 12,94035759 | 3,66283301 | 1,247456755 | 2,936240471 | 0,003322167 | 0,044739299 | The Moose: suction sampler vs FISH | NA |
| gene_1016398 | 13,17157428 | 3,103509471 | 1,013883006 | 3,061013405 | 0,002205892 | 0,035128307 | The Moose: suction sampler vs FISH | NA |
| gene_101679 | 13,11972911 | 3,497966707 | 1,092892623 | 3,200649939 | 0,00137118 | 0,02637363 | The Moose: suction sampler vs FISH | NA |
| gene_1017073 | 16,71744733 | 5,778316983 | 1,333785796 | 4,332267594 | 1,48E-05 | 0,001361698 | The Moose: suction sampler vs FISH | NA |
| gene_1017212 | 6,26233259 | 4,864998717 | 1,676028619 | 2,902694299 | 0,003699676 | 0,047701033 | The Moose: suction sampler vs FISH | NA |
| gene_1018111 | 5,860654493 | 5,65714205 | 1,805146508 | 3,133896348 | 0,001725018 | 0,030396816 | The Moose: suction sampler vs FISH | NA |
| gene_1018781 | 12,77531446 | 4,527737449 | 1,384344631 | 3,270672163 | 0,001072922 | 0,022755058 | The Moose: suction sampler vs FISH | NA |
| gene_1019544 | 4,984634686 | 4,938392191 | 1,597060241 | 3,092176528 | 0,001986947 | 0,033053519 | The Moose: suction sampler vs FISH | NA |
| gene_1019939 | 23,34009672 | 3,710557676 | 1,263910175 | 2,935776411 | 0,00332714 | 0,044779518 | The Moose: suction sampler vs FISH | NA |
| gene_1020370 | 188,1894827 | 3,693468494 | 0,956969181 | 3,859548004 | 0,000113597 | 0,005435262 | The Moose: suction sampler vs FISH | NA |
| gene_1021068 | 15,71839097 | 4,048359706 | 1,048089942 | 3,862607153 | 0,000112183 | 0,005392215 | The Moose: suction sampler vs FISH | NA |
| gene_1022795 | 27,27100099 | 2,50849247 | 0,824981186 | 3,040666274 | 0,002360553 | 0,036414792 | The Moose: suction sampler vs FISH | NA |
| gene_1023115 | 82,93803378 | 3,52864967 | 0,873011146 | 4,041929688 | 5,30E-05 | 0,003274642 | The Moose: suction sampler vs FISH | NA |
| gene_1023455 | 22,54635217 | 3,674638826 | 1,027150583 | 3,577507415 | 0,000346886 | 0,011139952 | The Moose: suction sampler vs FISH | NA |
| gene_1023818 | 16,81624774 | 4,024796462 | 1,107433679 | 3,634345368 | 0,000278687 | 0,009705686 | The Moose: suction sampler vs FISH | NA |
| gene_1025109 | 4,947483751 | 6,625349609 | 1,754064192 | 3,777142046 | 0,000158638 | 0,00673479 | The Moose: suction sampler vs FISH | NA |
| gene_1026379 | 30,1447493 | 2,859934213 | 0,859006777 | 3,32935 | 0,000870489 | 0,020049897 | The Moose: suction sampler vs FISH | NA |
| gene_1027181 | 49,11525283 | 3,622914719 | 0,853696306 | 4,243798051 | 2,20E-05 | 0,001779589 | The Moose: suction sampler vs FISH | NA |
| gene_1027532 | 7,629003636 | 3,91666602 | 1,272351336 | 3,078289706 | 0,002081924 | 0,03403983 | The Moose: suction sampler vs FISH | NA |
| gene_1027834 | 12,8190366 | 5,304679563 | 1,37535782 | 3,856945069 | 0,000114813 | 0,005461423 | The Moose: suction sampler vs FISH | NA |
| gene_1028316 | 3,843304255 | 6,113843883 | 1,988468804 | 3,074649133 | 0,002107503 | 0,034252904 | The Moose: suction sampler vs FISH | NA |
| gene_1028412 | 26,2107134 | 5,256923497 | 1,084055809 | 4,849310759 | 1,24E-06 | 0,000254879 | The Moose: suction sampler vs FISH | NA |
| gene_1028707 | 98,61937346 | 5,79130976 | 0,980490224 | 5,906545129 | 3,49E-09 | 4,14E-06 | The Moose: suction sampler vs FISH | NA |
| gene_1028889 | 4,132502895 | 6,375931873 | 1,999597843 | 3,188607097 | 0,0014296 | 0,027061083 | The Moose: suction sampler vs FISH | NA |
| gene_1029587 | 48,07937732 | 3,66962814 | 0,725058846 | 5,061145259 | 4,17E-07 | 0,000122611 | The Moose: suction sampler vs FISH | NA |
| gene_1031156 | 247,0035435 | 2,392314407 | 0,785293097 | 3,046396835 | 0,002316019 | 0,036030135 | The Moose: suction sampler vs FISH | NA |
| gene_1031678 | 2621,82357 | 5,478432708 | 0,854336331 | 6,412501152 | 1,43E-10 | 4,72E-07 | The Moose: suction sampler vs FISH | NA |
| gene_1031875 | 58,00218977 | 9,505776296 | 1,59137814 | 5,973298273 | 2,33E-09 | 3,22E-06 | The Moose: suction sampler vs FISH | NA |
| gene_1032120 | 44,79092659 | 4,587040527 | 0,90322147 | 5,078533535 | 3,80E-07 | 0,000115915 | The Moose: suction sampler vs FISH | NA |
| gene_1032121 | 23,5706289 | 5,301380649 | 1,137057135 | 4,662369626 | 3,13E-06 | 0,00046748 | The Moose: suction sampler vs FISH | NA |
| gene_1032339 | 5,16658513 | 5,867623394 | 1,864125356 | 3,147654945 | 0,001645859 | 0,029538022 | The Moose: suction sampler vs FISH | NA |
| gene_103406 | 100,1444105 | 4,085410198 | 0,825678677 | 4,947941995 | 7,50E-07 | 0,000183618 | The Moose: suction sampler vs FISH | NA |
| gene_1034355 | 49,47059417 | 2,842986169 | 0,783668953 | 3,627789716 | 0,000285858 | 0,009875275 | The Moose: suction sampler vs FISH | NA |
| gene_103472 | 4,626842305 | 6,356787362 | 1,808528412 | 3,514894939 | 0,000439928 | 0,012913873 | The Moose: suction sampler vs FISH | NA |
| gene_1035351 | 38,73675677 | 2,19262526 | 0,723149976 | 3,032047752 | 0,002429008 | 0,037096792 | The Moose: suction sampler vs FISH | NA |
| gene_1035680 | 8,735551985 | 4,05632777 | 1,231272124 | 3,294420212 | 0,000986249 | 0,021532428 | The Moose: suction sampler vs FISH | NA |
| gene_1036167 | 8,831985317 | 4,129329742 | 1,28718621 | 3,208028264 | 0,001336484 | 0,025940286 | The Moose: suction sampler vs FISH | NA |
| gene_1037422 | 5,330337955 | 7,491969989 | 1,733119079 | 4,322824714 | 1,54E-05 | 0,001400219 | The Moose: suction sampler vs FISH | NA |
| gene_1037843 | 689,514518 | 3,034953991 | 0,818776208 | 3,706695383 | 0,000209981 | 0,008053302 | The Moose: suction sampler vs FISH | NA |
| gene_1038944 | 27,768277 | 5,052723401 | 1,036460051 | 4,874981333 | 1,09E-06 | 0,000235758 | The Moose: suction sampler vs FISH | NA |
| gene_1038945 | 78,44490339 | 2,910908826 | 0,716038819 | 4,065294714 | 4,80E-05 | 0,003082493 | The Moose: suction sampler vs FISH | NA |
| gene_1038997 | 27,37067509 | 5,32238542 | 1,180730238 | 4,507706545 | 6,55E-06 | 0,000806199 | The Moose: suction sampler vs FISH | NA |
| gene_1039031 | 9,149601742 | 4,794397043 | 1,478863185 | 3,241947662 | 0,001187158 | 0,024172995 | The Moose: suction sampler vs FISH | NA |
| gene_1039580 | 12,5437878 | 3,565773966 | 1,225293614 | 2,910138375 | 0,003612688 | 0,046951593 | The Moose: suction sampler vs FISH | NA |
| gene_1039645 | 47,00868316 | -2,679224811 | 0,877591698 | -3,05292862 | 0,002266198 | 0,035564303 | The Moose: suction sampler vs FISH | NA |
| gene_1040406 | 8,114109083 | 4,255777358 | 1,408111903 | 3,022328942 | 0,002508378 | 0,037875787 | The Moose: suction sampler vs FISH | NA |
| gene_1040907 | 2,838207783 | 6,275734264 | 2,130545381 | 2,945599901 | 0,00322329 | 0,043978184 | The Moose: suction sampler vs FISH | NA |
| gene_1041357 | 9,425692898 | 4,901332033 | 1,520734278 | 3,223003587 | 0,00126854 | 0,025191854 | The Moose: suction sampler vs FISH | NA |
| gene_1041359 | 9,84376271 | 8,402599281 | 1,906450905 | 4,407456421 | 1,05E-05 | 0,001094264 | The Moose: suction sampler vs FISH | NA |
| gene_1042055 | 5,223780977 | 7,054580454 | 1,895921676 | 3,720923994 | 0,000198495 | 0,007771044 | The Moose: suction sampler vs FISH | NA |
| gene_1042576 | 4,910309099 | 6,127189227 | 1,740159956 | 3,521049433 | 0,000429842 | 0,01272975 | The Moose: suction sampler vs FISH | NA |
| gene_1042640 | 159,3731925 | 4,799370288 | 0,768013053 | 6,249073851 | 4,13E-10 | 1,01E-06 | The Moose: suction sampler vs FISH | NA |
| gene_1045346 | 18,33357739 | 7,162279372 | 1,532349642 | 4,674050346 | 2,95E-06 | 0,000447086 | The Moose: suction sampler vs FISH | NA |
| gene_1045489 | 21,93490918 | 3,483614141 | 1,002576553 | 3,474661491 | 0,000511498 | 0,014165535 | The Moose: suction sampler vs FISH | NA |
| gene_1046720 | 16,16927072 | 3,594206494 | 1,239807037 | 2,899004754 | 0,003743492 | 0,048020094 | The Moose: suction sampler vs FISH | NA |
| gene_1048674 | 1255,447682 | 2,518767641 | 0,675257686 | 3,73008363 | 0,000191416 | 0,007614874 | The Moose: suction sampler vs FISH | NA |
| gene_1049312 | 4,091533714 | 6,377527949 | 1,832790616 | 3,479681691 | 0,00050201 | 0,014008402 | The Moose: suction sampler vs FISH | NA |
| gene_1050175 | 8,067582316 | 5,434655787 | 1,504015808 | 3,613429964 | 0,000302173 | 0,01025559 | The Moose: suction sampler vs FISH | NA |
| gene_1050638 | 3,807894822 | 6,086586832 | 2,100018545 | 2,898349087 | 0,003751328 | 0,0480412 | The Moose: suction sampler vs FISH | NA |
| gene_105164 | 6,369716885 | 6,495622314 | 1,921408943 | 3,380655815 | 0,000723131 | 0,017704153 | The Moose: suction sampler vs FISH | NA |
| gene_1053654 | 19,14497503 | -3,994045027 | 1,237701697 | -3,226985176 | 0,001251019 | 0,024933818 | The Moose: suction sampler vs FISH | NA |
| gene_1053695 | 8,956687841 | 3,901682336 | 1,213182563 | 3,216071889 | 0,001299583 | 0,025546445 | The Moose: suction sampler vs FISH | NA |

| | | | | | | | | |
|---|---|---|---|---|---|---|---|---|
| gene_1053755 | 9,377625993 | 5,022041992 | 1,436806624 | 3,495280373 | 0,000473564 | 0,013465315 | The Moose: suction sampler vs FISH | NA |
| gene_1053756 | 8,354658393 | 7,137920567 | 1,713710562 | 4,165184439 | 3,11E-05 | 0,002267936 | The Moose: suction sampler vs FISH | NA |
| gene_1054459 | 6,566730845 | 4,358399818 | 1,493211987 | 2,918808486 | 0,00351372 | 0,046134909 | The Moose: suction sampler vs FISH | NA |
| gene_1055956 | 19,06658082 | 3,626533425 | 1,071971336 | 3,383050743 | 0,000716854 | 0,017608472 | The Moose: suction sampler vs FISH | NA |
| gene_1056060 | 132,977104 | 3,437678542 | 0,744522886 | 4,617290629 | 3,89E-06 | 0,000552304 | The Moose: suction sampler vs FISH | NA |
| gene_1056396 | 9,537523672 | 5,669223894 | 1,538211937 | 3,685593484 | 0,00022817 | 0,008527735 | The Moose: suction sampler vs FISH | NA |
| gene_1056609 | 66,17638008 | -2,583037827 | 0,761089293 | -3,393869616 | 0,000689125 | 0,017145251 | The Moose: suction sampler vs FISH | NA |
| gene_1057377 | 28,86027935 | 3,520716286 | 0,924881947 | 3,806665593 | 0,000140853 | 0,006234468 | The Moose: suction sampler vs FISH | NA |
| gene_1058126 | 51,57368975 | 5,762112022 | 1,061268422 | 5,429457716 | 5,65E-08 | 3,28E-05 | The Moose: suction sampler vs FISH | NA |
| gene_1058127 | 20,85290311 | 4,572690785 | 1,047264999 | 4,366316824 | 1,26E-05 | 0,001239045 | The Moose: suction sampler vs FISH | NA |
| gene_1060379 | 177,1038214 | 12,83387577 | 4,247219484 | 3,021712398 | 0,002513492 | 0,037931428 | The Moose: suction sampler vs FISH | NA |
| gene_1060932 | 12,91463524 | 3,149056661 | 1,026982537 | 3,066319579 | 0,002167116 | 0,03478518 | The Moose: suction sampler vs FISH | NA |
| gene_1061731 | 5,049013597 | 4,650041322 | 1,486006539 | 3,129219961 | 0,001752711 | 0,030666956 | The Moose: suction sampler vs FISH | NA |
| gene_1062878 | 135,1851135 | -2,013238948 | 0,605955247 | -3,32242184 | 0,000892397 | 0,020379998 | The Moose: suction sampler vs FISH | NA |
| gene_1063956 | 4161,607121 | 3,638891299 | 0,768284595 | 4,736384566 | 2,18E-06 | 0,000366549 | The Moose: suction sampler vs FISH | NA |
| gene_1064600 | 13,842088 | 4,295084465 | 1,229808159 | 3,492483306 | 0,000478552 | 0,013571764 | The Moose: suction sampler vs FISH | NA |
| gene_1064654 | 50,95791277 | 2,561339765 | 0,853029506 | 3,002639122 | 0,002676496 | 0,039306464 | The Moose: suction sampler vs FISH | NA |
| gene_1066070 | 38,44086938 | 3,004370487 | 0,79726923 | 3,768326148 | 0,000164346 | 0,006894075 | The Moose: suction sampler vs FISH | NA |
| gene_1066447 | 4,816649778 | 6,09475071 | 1,917997275 | 3,177663904 | 0,001484667 | 0,027738846 | The Moose: suction sampler vs FISH | NA |
| gene_106648 | 25,76051964 | 5,532726169 | 1,324428111 | 4,177445438 | 2,95E-05 | 0,002186829 | The Moose: suction sampler vs FISH | NA |
| gene_1067548 | 17,18366392 | 3,797965723 | 1,059701002 | 3,583997483 | 0,000338375 | 0,010962495 | The Moose: suction sampler vs FISH | NA |
| gene_1067979 | 5,241397961 | 5,444079291 | 1,675675428 | 3,248886509 | 0,001158577 | 0,023840915 | The Moose: suction sampler vs FISH | NA |
| gene_1068307 | 4,819592751 | 6,083652704 | 1,913351286 | 3,179579594 | 0,001474889 | 0,027618763 | The Moose: suction sampler vs FISH | NA |
| gene_1070120 | 25,05490476 | 3,503848416 | 0,945144291 | 3,707210051 | 0,000209555 | 0,008049569 | The Moose: suction sampler vs FISH | NA |
| gene_1070646 | 74,78876278 | 3,332565846 | 0,827757292 | 4,02601811 | 5,67E-05 | 0,003432031 | The Moose: suction sampler vs FISH | NA |
| gene_1070663 | 19,7839389 | 2,581040865 | 0,82384873 | 3,132906285 | 0,001730847 | 0,030426115 | The Moose: suction sampler vs FISH | NA |
| gene_1070730 | 12,13850696 | 5,749582486 | 1,361899064 | 4,221739068 | 2,42E-05 | 0,001914503 | The Moose: suction sampler vs FISH | NA |
| gene_1071172 | 11,64530956 | 3,437400471 | 1,112701078 | 3,089239813 | 0,002006694 | 0,033231484 | The Moose: suction sampler vs FISH | NA |
| gene_1071440 | 4,699016308 | 7,043470968 | 2,007988082 | 3,507725484 | 0,000451955 | 0,013116056 | The Moose: suction sampler vs FISH | NA |
| gene_1073096 | 30,05118648 | 4,735464588 | 1,125698019 | 4,206691767 | 2,59E-05 | 0,002009742 | The Moose: suction sampler vs FISH | NA |
| gene_1076117 | 70,3560968 | 2,714295729 | 0,743084974 | 3,652739355 | 0,000259458 | 0,009267608 | The Moose: suction sampler vs FISH | NA |
| gene_1076645 | 8,353191692 | 4,503847478 | 1,299977771 | 3,464557302 | 0,000531105 | 0,014489972 | The Moose: suction sampler vs FISH | NA |
| gene_1076859 | 48,26537691 | 4,136943652 | 0,91091083 | 4,541546238 | 5,58E-06 | 0,000712048 | The Moose: suction sampler vs FISH | NA |
| gene_1076944 | 5,266281375 | 5,204103938 | 1,769835495 | 2,940445003 | 0,003277412 | 0,044349108 | The Moose: suction sampler vs FISH | NA |
| gene_1077725 | 3,312919493 | 5,989822554 | 2,040058405 | 2,936103466 | 0,003323634 | 0,044748732 | The Moose: suction sampler vs FISH | NA |
| gene_1078744 | 33,58247698 | 3,249690336 | 1,125417849 | 2,887541139 | 0,003882658 | 0,048963228 | The Moose: suction sampler vs FISH | NA |
| gene_1078759 | 10,35265887 | -5,08201731 | 1,548614444 | -3,281654339 | 0,001032 | 0,022168839 | The Moose: suction sampler vs FISH | NA |
| gene_107925 | 6,729325883 | 4,450641426 | 1,435913808 | 3,099518509 | 0,001938355 | 0,032577222 | The Moose: suction sampler vs FISH | NA |
| gene_1079269 | 11,1398062 | 3,543324487 | 1,17721132 | 3,009930694 | 0,002613073 | 0,038833904 | The Moose: suction sampler vs FISH | NA |
| gene_1079479 | 17,27190874 | 3,162604933 | 0,956788662 | 3,3054373 | 0,000948283 | 0,021060374 | The Moose: suction sampler vs FISH | NA |
| gene_1079508 | 63,90094252 | 2,587307148 | 0,80578292 | 3,210923293 | 0,001323092 | 0,025823392 | The Moose: suction sampler vs FISH | NA |
| gene_10799 | 15,99715665 | 3,448961316 | 1,078207652 | 3,198791355 | 0,00138005 | 0,026474467 | The Moose: suction sampler vs FISH | NA |
| gene_1080152 | 14,91299415 | 3,957470371 | 0,96950641 | 4,081943483 | 4,47E-05 | 0,002923442 | The Moose: suction sampler vs FISH | NA |
| gene_1080283 | 35,81593438 | 10,44767215 | 3,40895081 | 3,064776447 | 0,002178328 | 0,03487841 | The Moose: suction sampler vs FISH | NA |
| gene_1082139 | 7,790734375 | 6,82813114 | 1,615481395 | 4,226685098 | 2,37E-05 | 0,001878981 | The Moose: suction sampler vs FISH | NA |
| gene_1082424 | 9,813276613 | 6,180673516 | 1,727831682 | 3,57712709 | 0,000347391 | 0,011146419 | The Moose: suction sampler vs FISH | NA |
| gene_1082425 | 7,267171979 | 7,440772761 | 1,798079727 | 4,138177329 | 3,50E-05 | 0,002454294 | The Moose: suction sampler vs FISH | NA |
| gene_1083182 | 11,26034419 | 8,476242248 | 1,900371864 | 4,46030717 | 8,18E-06 | 0,000934822 | The Moose: suction sampler vs FISH | NA |
| gene_1083571 | 10,16625979 | 3,881460177 | 1,25450686 | 3,094012717 | 0,00197469 | 0,03293916 | The Moose: suction sampler vs FISH | NA |
| gene_1083858 | 32,64793395 | 2,59804375 | 0,818817675 | 3,172920943 | 0,001509136 | 0,028018373 | The Moose: suction sampler vs FISH | NA |
| gene_108484 | 10,63092843 | 3,42917945 | 1,068641394 | 3,208915048 | 0,001332369 | 0,025901484 | The Moose: suction sampler vs FISH | NA |
| gene_1085047 | 7,611738303 | 6,147753296 | 1,590100525 | 3,866267069 | 0,000110514 | 0,005332914 | The Moose: suction sampler vs FISH | NA |
| gene_108568 | 148,3294463 | -2,481152931 | 0,689200276 | -3,600046341 | 0,00031816 | 0,010580484 | The Moose: suction sampler vs FISH | NA |
| gene_108569 | 117,5623283 | -2,394030778 | 0,647339643 | -3,698260724 | 0,000217082 | 0,008226554 | The Moose: suction sampler vs FISH | NA |
| gene_1085767 | 14,89663431 | 4,385155307 | 1,322699419 | 3,315307502 | 0,000915424 | 0,02061142 | The Moose: suction sampler vs FISH | NA |
| gene_1086682 | 8,548557119 | 4,154149497 | 1,35263104 | 3,071162331 | 0,002132272 | 0,034462996 | The Moose: suction sampler vs FISH | NA |
| gene_1086683 | 7,725745515 | 7,901214004 | 1,728714513 | 4,570571915 | 4,86E-06 | 0,000647361 | The Moose: suction sampler vs FISH | NA |
| gene_1087066 | 111,3883199 | 5,469042652 | 1,057568105 | 5,171338493 | 2,32E-07 | 8,21E-05 | The Moose: suction sampler vs FISH | NA |
| gene_1087995 | 22,22322288 | 5,472005798 | 1,146764182 | 4,771692284 | 1,83E-06 | 0,00032805 | The Moose: suction sampler vs FISH | NA |
| gene_1088106 | 170,8258308 | 4,209261822 | 0,996475247 | 4,22415091 | 2,40E-05 | 0,001898199 | The Moose: suction sampler vs FISH | NA |
| gene_1088107 | 14,30472805 | -4,084112206 | 1,317887523 | -3,098983891 | 0,001941856 | 0,032613671 | The Moose: suction sampler vs FISH | NA |
| gene_1088149 | 2,990620352 | 5,711211118 | 1,933903864 | 2,953203221 | 0,003144948 | 0,043344106 | The Moose: suction sampler vs FISH | NA |
| gene_1089958 | 84,85547076 | 3,236561858 | 0,865740452 | 3,738489809 | 0,000185129 | 0,007425887 | The Moose: suction sampler vs FISH | NA |
| gene_1091957 | 59,36651696 | 2,649580813 | 0,685788445 | 3,863554182 | 0,000111749 | 0,005378383 | The Moose: suction sampler vs FISH | NA |
| gene_1092217 | 42,80343159 | 2,813838383 | 0,831526176 | 3,383944444 | 0,000714525 | 0,017574769 | The Moose: suction sampler vs FISH | NA |

| | | | | | | | | |
|---|---|---|---|---|---|---|---|---|
| gene_1093045 | 22,58858201 | 4,727565084 | 1,391139291 | 3,398340564 | 0,00067796 | 0,016970943 | The Moose: suction sampler vs FISH | NA |
| gene_109306 | 421,0348595 | 3,263735249 | 0,85057476 | 3,83709393 | 0,000124499 | 0,005787227 | The Moose: suction sampler vs FISH | NA |
| gene_1094596 | 5,861847013 | -5,607309242 | 1,772955969 | -3,162689509 | 0,00156319 | 0,028594883 | The Moose: suction sampler vs FISH | NA |
| gene_1094748 | 23,8085995 | 2,985259398 | 0,896190893 | 3,331053039 | 0,000865181 | 0,019982148 | The Moose: suction sampler vs FISH | NA |
| gene_1095329 | 91,7950602 | 5,64707392 | 0,953823364 | 5,920460888 | 3,21E-09 | 3,93E-06 | The Moose: suction sampler vs FISH | NA |
| gene_1095883 | 6,000035303 | 5,763312845 | 1,588464826 | 3,62822818 | 0,000285373 | 0,009863162 | The Moose: suction sampler vs FISH | NA |
| gene_1096026 | 13,89260451 | 8,038869621 | 1,848146298 | 4,349693327 | 1,36E-05 | 0,001300339 | The Moose: suction sampler vs FISH | NA |
| gene_1096568 | 50,33657841 | 2,914390738 | 0,714752662 | 4,077481476 | 4,55E-05 | 0,002972121 | The Moose: suction sampler vs FISH | NA |
| gene_1097070 | 328,4355298 | 4,438567145 | 0,871758676 | 5,091509001 | 3,55E-07 | 0,000110082 | The Moose: suction sampler vs FISH | NA |
| gene_1097143 | 9,934540252 | 3,25156022 | 1,020152269 | 3,187328322 | 0,001435937 | 0,027140573 | The Moose: suction sampler vs FISH | NA |
| gene_1097160 | 6,509596591 | 4,859098926 | 1,620318557 | 2,998854086 | 0,002709971 | 0,0395379 | The Moose: suction sampler vs FISH | NA |
| gene_1097185 | 12,6816607 | 7,357767845 | 1,611813862 | 4,564899223 | 5,00E-06 | 0,000658941 | The Moose: suction sampler vs FISH | NA |
| gene_1097236 | 4,463952594 | 5,927832378 | 1,952547494 | 3,035947856 | 0,002397808 | 0,036757885 | The Moose: suction sampler vs FISH | NA |
| gene_1097459 | 6,598315313 | 7,219879379 | 2,006647212 | 3,597981417 | 0,000320697 | 0,010614491 | The Moose: suction sampler vs FISH | NA |
| gene_1099169 | 31,82345399 | 3,535273239 | 0,897069827 | 3,940911992 | 8,12E-05 | 0,004338943 | The Moose: suction sampler vs FISH | NA |
| gene_1100306 | 6,346937553 | 6,290933359 | 1,790848809 | 3,512822148 | 0,000443374 | 0,012965789 | The Moose: suction sampler vs FISH | NA |
| gene_1101049 | 12,21623569 | 4,042912365 | 1,312171491 | 3,081085356 | 0,002062475 | 0,03383482 | The Moose: suction sampler vs FISH | NA |
| gene_1103512 | 20,42392657 | 2,593388901 | 0,869594896 | 2,982295448 | 0,002860958 | 0,040967641 | The Moose: suction sampler vs FISH | NA |
| gene_1103613 | 4,639559973 | 7,136613569 | 1,99670984 | 3,574186607 | 0,000351318 | 0,011199034 | The Moose: suction sampler vs FISH | NA |
| gene_1104091 | 7,729462474 | 3,935375842 | 1,241071283 | 3,170950692 | 0,001519409 | 0,02812324 | The Moose: suction sampler vs FISH | NA |
| gene_1104092 | 11,59813559 | 3,552908357 | 1,235048652 | 2,876735544 | 0,004018122 | 0,04992372 | The Moose: suction sampler vs FISH | NA |
| gene_1104195 | 13,38259239 | 2,872020786 | 0,909326818 | 3,158403261 | 0,00158636 | 0,028860583 | The Moose: suction sampler vs FISH | NA |
| gene_1104975 | 16,7522755 | 4,200223175 | 1,18279248 | 3,55110744 | 0,000383614 | 0,011875719 | The Moose: suction sampler vs FISH | NA |
| gene_1104976 | 10,65140794 | 7,657822233 | 1,703486098 | 4,495382875 | 6,94E-06 | 0,00083664 | The Moose: suction sampler vs FISH | NA |
| gene_1105101 | 14,34975036 | 3,675344167 | 1,09556482 | 3,354748255 | 0,000794372 | 0,018842546 | The Moose: suction sampler vs FISH | NA |
| gene_1106050 | 3,615520845 | 6,518089401 | 2,131802505 | 3,057548429 | 0,002231556 | 0,035292122 | The Moose: suction sampler vs FISH | NA |
| gene_1107567 | 15,70390085 | 4,235043512 | 1,019796841 | 4,152830587 | 3,28E-05 | 0,002356761 | The Moose: suction sampler vs FISH | NA |
| gene_11082 | 5,558251264 | 5,608586107 | 1,779366481 | 3,152012902 | 0,001621491 | 0,029267735 | The Moose: suction sampler vs FISH | NA |
| gene_1108441 | 17,48251683 | 8,040200204 | 1,807058926 | 4,449329288 | 8,61E-06 | 0,000962933 | The Moose: suction sampler vs FISH | NA |
| gene_1108473 | 30,08288751 | 4,047693291 | 1,006677924 | 4,020842411 | 5,80E-05 | 0,003482506 | The Moose: suction sampler vs FISH | NA |
| gene_1109508 | 375,3341243 | 13,90358758 | 3,223143399 | 4,313673286 | 1,61E-05 | 0,001441643 | The Moose: suction sampler vs FISH | NA |
| gene_111069 | 26,20453343 | 2,756387589 | 0,904075668 | 3,048846116 | 0,002297221 | 0,035836748 | The Moose: suction sampler vs FISH | NA |
| gene_1112062 | 96,0504371 | 2,153188564 | 0,731366483 | 2,9440624 | 0,003239347 | 0,044090776 | The Moose: suction sampler vs FISH | NA |
| gene_1112367 | 12,34898819 | 3,320308082 | 1,022007445 | 3,248810074 | 0,001158888 | 0,023840915 | The Moose: suction sampler vs FISH | NA |
| gene_1112593 | 4,934595607 | 6,088977054 | 1,831084055 | 3,32534 | 0,000883108 | 0,020232022 | The Moose: suction sampler vs FISH | NA |
| gene_1113036 | 28,50786209 | 3,741823021 | 0,913981822 | 4,093979694 | 4,24E-05 | 0,002813278 | The Moose: suction sampler vs FISH | NA |
| gene_1113889 | 9,11801688 | 4,405927313 | 1,39555295 | 3,157119415 | 0,001593361 | 0,028923476 | The Moose: suction sampler vs FISH | NA |
| gene_1114187 | 5,226924607 | 5,415343298 | 1,876946431 | 2,885187988 | 0,0039118 | 0,049132477 | The Moose: suction sampler vs FISH | NA |
| gene_1115477 | 7,65824699 | 4,007252965 | 1,335980759 | 2,999484039 | 0,002704373 | 0,039498414 | The Moose: suction sampler vs FISH | NA |
| gene_1115919 | 12,00811333 | 4,866156416 | 1,423621528 | 3,418153155 | 0,000630476 | 0,016173707 | The Moose: suction sampler vs FISH | NA |
| gene_1116699 | 9,790821783 | 3,671276004 | 1,151625278 | 3,187908492 | 0,001433059 | 0,027112572 | The Moose: suction sampler vs FISH | NA |
| gene_111676 | 19,69354766 | -4,904678573 | 1,543122709 | -3,178411247 | 0,001480845 | 0,027688568 | The Moose: suction sampler vs FISH | NA |
| gene_111692 | 16,0449908 | -2,93954609 | 0,963209245 | -3,051825036 | 0,002274546 | 0,035649607 | The Moose: suction sampler vs FISH | NA |
| gene_111767 | 5,583410138 | 7,556792722 | 2,115183664 | 3,572641398 | 0,000353399 | 0,011250696 | The Moose: suction sampler vs FISH | NA |
| gene_1118049 | 25,36573927 | 5,242430906 | 1,036916692 | 5,05578794 | 4,29E-07 | 0,000123936 | The Moose: suction sampler vs FISH | NA |
| gene_1118163 | 47,18081822 | 2,608256936 | 0,830334725 | 3,141211439 | 0,001682505 | 0,029964014 | The Moose: suction sampler vs FISH | NA |
| gene_1118243 | 87,02796412 | 2,402650188 | 0,789331637 | 3,043904584 | 0,002335292 | 0,036197774 | The Moose: suction sampler vs FISH | NA |
| gene_1119191 | 12,66725491 | 8,59617224 | 1,931093139 | 4,451453979 | 8,53E-06 | 0,000961318 | The Moose: suction sampler vs FISH | NA |
| gene_1119192 | 33,61500993 | 7,66337287 | 1,434387324 | 5,342610565 | 9,16E-08 | 4,55E-05 | The Moose: suction sampler vs FISH | NA |
| gene_1119194 | 7,966773701 | 5,038805927 | 1,732894687 | 2,907739267 | 0,003640517 | 0,047187461 | The Moose: suction sampler vs FISH | NA |
| gene_1119267 | 6,871664941 | 3,825993096 | 1,317793322 | 2,903333195 | 0,003692136 | 0,047631991 | The Moose: suction sampler vs FISH | NA |
| gene_1119269 | 7,423895234 | 6,693915601 | 1,734803181 | 3,858602332 | 0,000114037 | 0,005442151 | The Moose: suction sampler vs FISH | NA |
| gene_1119969 | 39,52997147 | 4,731494135 | 0,99142948 | 4,772396051 | 1,82E-06 | 0,00032805 | The Moose: suction sampler vs FISH | NA |
| gene_1120842 | 13,11706314 | 4,7381976 | 1,181692386 | 4,009670924 | 6,08E-05 | 0,003589797 | The Moose: suction sampler vs FISH | NA |
| gene_1121114 | 112,997099 | 2,814024646 | 0,937925345 | 3,000265066 | 0,002697448 | 0,03943505 | The Moose: suction sampler vs FISH | NA |
| gene_1121439 | 34,9090653 | 2,6551763 | 0,881988113 | 3,01044454 | 0,002608656 | 0,038790953 | The Moose: suction sampler vs FISH | NA |
| gene_1122984 | 14,44396929 | 4,804502381 | 1,227426022 | 3,914290797 | 9,07E-05 | 0,004696249 | The Moose: suction sampler vs FISH | NA |
| gene_1125054 | 18,75971867 | 2,429532747 | 0,81668834 | 2,974859109 | 0,002931231 | 0,041528606 | The Moose: suction sampler vs FISH | NA |
| gene_1126267 | 7,687938925 | 4,370426224 | 1,50955243 | 2,895180146 | 0,00378941 | 0,048307916 | The Moose: suction sampler vs FISH | NA |
| gene_1126645 | 7,469295984 | 3,706858777 | 1,282426948 | 2,890502873 | 0,00384626 | 0,048721728 | The Moose: suction sampler vs FISH | NA |
| gene_1126958 | 19,09934004 | 3,632029916 | 1,17489048 | 3,091377433 | 0,001992302 | 0,033090147 | The Moose: suction sampler vs FISH | NA |
| gene_1127446 | 10,73539399 | 3,61426206 | 1,185076482 | 3,049813337 | 0,002289836 | 0,035759522 | The Moose: suction sampler vs FISH | NA |
| gene_1128502 | 11,32630155 | 3,631647354 | 1,078818506 | 3,366319112 | 0,000761785 | 0,018350055 | The Moose: suction sampler vs FISH | NA |
| gene_1128664 | 6,371858706 | 6,586852498 | 1,986642295 | 3,315570455 | 0,000914563 | 0,020610638 | The Moose: suction sampler vs FISH | NA |
| gene_1128909 | 76,56239782 | -2,329502387 | 0,796602749 | -2,924296196 | 0,003452361 | 0,045700744 | The Moose: suction sampler vs FISH | NA |

| | | | | | | | | |
|---|---|---|---|---|---|---|---|---|
| gene_1130733 | 11,51585467 | 4,188179477 | 1,298440114 | 3,225546895 | 0,001257322 | 0,025025485 | The Moose: suction sampler vs FISH | NA |
| gene_1130993 | 17,5451242 | 4,192005843 | 1,038198978 | 4,03776726 | 5,40E-05 | 0,003307895 | The Moose: suction sampler vs FISH | NA |
| gene_1131613 | 6,710984411 | 6,225063954 | 1,640502459 | 3,794608122 | 0,000147877 | 0,006411342 | The Moose: suction sampler vs FISH | NA |
| gene_1132726 | 6,894579105 | 4,924848985 | 1,522396086 | 3,234932768 | 0,001216714 | 0,024516198 | The Moose: suction sampler vs FISH | NA |
| gene_1133223 | 7,309644648 | 5,454246495 | 1,528025602 | 3,569473238 | 0,0003577 | 0,011357993 | The Moose: suction sampler vs FISH | NA |
| gene_1134773 | 19,75546061 | 4,591084944 | 1,14785803 | 3,999697545 | 6,34E-05 | 0,003679414 | The Moose: suction sampler vs FISH | NA |
| gene_1135311 | 28,02924656 | 3,175641485 | 0,934593103 | 3,39788671 | 0,000679085 | 0,016986711 | The Moose: suction sampler vs FISH | NA |
| gene_1135509 | 7,144641239 | 4,702738474 | 1,488960612 | 3,158403544 | 0,001586358 | 0,028860583 | The Moose: suction sampler vs FISH | NA |
| gene_1135983 | 12,96173482 | 8,833559387 | 1,71329392 | 5,155892565 | 2,52E-07 | 8,66E-05 | The Moose: suction sampler vs FISH | NA |
| gene_1136019 | 11,79941082 | 5,673968433 | 1,408250852 | 4,029089295 | 5,60E-05 | 0,003393095 | The Moose: suction sampler vs FISH | NA |
| gene_1136054 | 86,05554019 | 3,144675856 | 0,732428847 | 4,293489897 | 1,76E-05 | 0,001530565 | The Moose: suction sampler vs FISH | NA |
| gene_1136880 | 8,136837614 | 5,606347203 | 1,437026644 | 3,901352302 | 9,57E-05 | 0,004864833 | The Moose: suction sampler vs FISH | NA |
| gene_1137096 | 798,8692573 | 3,034879392 | 0,844725501 | 3,592740349 | 0,000327219 | 0,01075887 | The Moose: suction sampler vs FISH | NA |
| gene_113772 | 68,35099254 | 2,608902299 | 0,78737685 | 3,313409965 | 0,000921658 | 0,020705711 | The Moose: suction sampler vs FISH | NA |
| gene_1138464 | 6,116440002 | 5,866567404 | 1,697744774 | 3,45550609 | 0,000549261 | 0,014796594 | The Moose: suction sampler vs FISH | NA |
| gene_113963 | 502,5618398 | 7,399727886 | 1,135018818 | 6,519475948 | 7,06E-11 | 2,88E-07 | The Moose: suction sampler vs FISH | NA |
| gene_1139634 | 8,784267335 | 4,565856564 | 1,22381221 | 3,73084737 | 0,000190837 | 0,007604997 | The Moose: suction sampler vs FISH | NA |
| gene_1140429 | 14,39169025 | 3,473353348 | 1,187874259 | 2,924007588 | 0,003455563 | 0,045712147 | The Moose: suction sampler vs FISH | NA |
| gene_114051 | 51,74945887 | 3,062645466 | 0,939604058 | 3,259506426 | 0,001116063 | 0,0233132 | The Moose: suction sampler vs FISH | NA |
| gene_1140760 | 8,823124694 | 4,656627558 | 1,529703819 | 3,044136714 | 0,002333491 | 0,036179697 | The Moose: suction sampler vs FISH | NA |
| gene_1141442 | 40,0792954 | 2,478561995 | 0,846263897 | 2,928828707 | 0,003402419 | 0,045327524 | The Moose: suction sampler vs FISH | NA |
| gene_1141766 | 97,60075242 | 3,515056475 | 0,819935032 | 4,286993892 | 1,81E-05 | 0,001555722 | The Moose: suction sampler vs FISH | NA |
| gene_1142312 | 376,8295537 | 2,686589667 | 0,837027884 | 3,20967762 | 0,001328839 | 0,025868826 | The Moose: suction sampler vs FISH | NA |
| gene_1142394 | 14,36044426 | 2,939656185 | 0,9941928 | 2,956827072 | 0,003108224 | 0,043085392 | The Moose: suction sampler vs FISH | NA |
| gene_1142566 | 8,073178445 | 3,593809016 | 1,239212393 | 2,900075109 | 0,003730733 | 0,047936258 | The Moose: suction sampler vs FISH | NA |
| gene_114273 | 318,4319229 | -3,442501103 | 1,041913409 | -3,304018426 | 0,000953096 | 0,02110344 | The Moose: suction sampler vs FISH | NA |
| gene_1143972 | 7,940087996 | 5,953148092 | 1,55708535 | 3,823263826 | 0,000131697 | 0,005963296 | The Moose: suction sampler vs FISH | NA |
| gene_1143991 | 9,492748553 | 4,739714077 | 1,252002765 | 3,785705759 | 0,000153273 | 0,0065601 | The Moose: suction sampler vs FISH | NA |
| gene_1144744 | 33,94722059 | 3,304720095 | 0,784370101 | 4,213215281 | 2,52E-05 | 0,001973381 | The Moose: suction sampler vs FISH | NA |
| gene_1145614 | 72,60497292 | 4,647581932 | 0,903029895 | 5,146653456 | 2,65E-07 | 8,97E-05 | The Moose: suction sampler vs FISH | NA |
| gene_1145782 | 7,051490844 | 5,587176112 | 1,796213794 | 3,110529565 | 0,001867522 | 0,031875471 | The Moose: suction sampler vs FISH | NA |
| gene_1146558 | 5,185363309 | 6,92556195 | 2,16162103 | 3,203874247 | 0,001355917 | 0,026192876 | The Moose: suction sampler vs FISH | NA |
| gene_1147423 | 10,96490999 | 3,631284104 | 1,050785241 | 3,455781413 | 0,0005487 | 0,014796594 | The Moose: suction sampler vs FISH | NA |
| gene_1147518 | 4,900077235 | 5,810270361 | 1,852746396 | 3,136031123 | 0,00171251 | 0,030263551 | The Moose: suction sampler vs FISH | NA |
| gene_1148500 | 28,76685856 | 3,823468262 | 0,836529053 | 4,570634156 | 4,86E-06 | 0,000647361 | The Moose: suction sampler vs FISH | NA |
| gene_114922 | 5,188469115 | 6,523303618 | 1,90619485 | 3,422159922 | 0,000621258 | 0,016004302 | The Moose: suction sampler vs FISH | NA |
| gene_1149931 | 33,73634073 | 3,744370497 | 0,948828352 | 3,946309667 | 7,94E-05 | 0,004276566 | The Moose: suction sampler vs FISH | NA |
| gene_115005 | 63,23959843 | 3,480410898 | 0,763763469 | 4,556922452 | 5,19E-06 | 0,000673573 | The Moose: suction sampler vs FISH | NA |
| gene_1150856 | 13,19402566 | 5,366380874 | 1,283296984 | 4,18171393 | 2,89E-05 | 0,002163859 | The Moose: suction sampler vs FISH | NA |
| gene_1151424 | 15,81488226 | 2,868647669 | 0,989434859 | 2,89927896 | 0,00374022 | 0,048009067 | The Moose: suction sampler vs FISH | NA |
| gene_1151585 | 5,430000994 | 6,658999219 | 1,895974692 | 3,512177272 | 0,000444451 | 0,012980234 | The Moose: suction sampler vs FISH | NA |
| gene_1152214 | 38,36335447 | 2,587945393 | 0,886986771 | 2,917682065 | 0,003526437 | 0,046244641 | The Moose: suction sampler vs FISH | NA |
| gene_1152263 | 3,060650075 | 6,224122314 | 2,014647677 | 3,089434637 | 0,002005378 | 0,033231484 | The Moose: suction sampler vs FISH | NA |
| gene_1152755 | 5,219413276 | 5,193450134 | 1,623506956 | 3,198908458 | 0,00137949 | 0,026474162 | The Moose: suction sampler vs FISH | NA |
| gene_1152756 | 5,575978958 | 5,27917747 | 1,615085338 | 3,26866782 | 0,001080551 | 0,022864031 | The Moose: suction sampler vs FISH | NA |
| gene_1153124 | 11,23167612 | 3,831537962 | 1,322062558 | 2,898151785 | 0,003753689 | 0,048063053 | The Moose: suction sampler vs FISH | NA |
| gene_1153230 | 16,65232149 | 2,923066939 | 0,912892299 | 3,201984442 | 0,001364844 | 0,026282366 | The Moose: suction sampler vs FISH | NA |
| gene_1153814 | 231,7750104 | -2,770615457 | 0,756911798 | -3,660420493 | 0,000251802 | 0,009078829 | The Moose: suction sampler vs FISH | NA |
| gene_115400 | 9,962498254 | 3,32834263 | 1,117829303 | 2,977505261 | 0,002906047 | 0,041323251 | The Moose: suction sampler vs FISH | NA |
| gene_1154186 | 17,30931777 | 4,477394269 | 1,174338567 | 3,812694561 | 0,00013746 | 0,006145279 | The Moose: suction sampler vs FISH | NA |
| gene_1155548 | 18,73043316 | 3,148700718 | 1,056885533 | 2,979225866 | 0,002889777 | 0,041179393 | The Moose: suction sampler vs FISH | NA |
| gene_1156174 | 10,19034399 | 7,971938232 | 1,708221161 | 4,66680686 | 3,06E-06 | 0,000460411 | The Moose: suction sampler vs FISH | NA |
| gene_1156204 | 11,03043365 | 3,323088913 | 1,067688281 | 3,112414899 | 0,001855635 | 0,031746358 | The Moose: suction sampler vs FISH | NA |
| gene_1156960 | 109,1077052 | 2,750369888 | 0,741829197 | 3,707551415 | 0,000209273 | 0,008049569 | The Moose: suction sampler vs FISH | NA |
| gene_1157266 | 8,607564249 | 4,152631739 | 1,190037245 | 3,489497288 | 0,00048393 | 0,013649185 | The Moose: suction sampler vs FISH | NA |
| gene_1157946 | 28,16400805 | 3,356511397 | 1,143615095 | 2,935000955 | 0,003335467 | 0,044841602 | The Moose: suction sampler vs FISH | NA |
| gene_1157947 | 14,22373523 | 4,072471794 | 1,100449812 | 3,700733783 | 0,000214977 | 0,008176582 | The Moose: suction sampler vs FISH | NA |
| gene_1158400 | 29,20533847 | 2,596049445 | 0,814494883 | 3,187312162 | 0,001436017 | 0,027140573 | The Moose: suction sampler vs FISH | NA |
| gene_1159151 | 11,71691848 | 4,684533293 | 1,414631497 | 3,311486634 | 0,000928017 | 0,020805308 | The Moose: suction sampler vs FISH | NA |
| gene_1159152 | 32,8913552 | 5,91984282 | 1,286576821 | 4,601235406 | 4,20E-06 | 0,000584211 | The Moose: suction sampler vs FISH | NA |
| gene_1159638 | 9,307979629 | 4,147864761 | 1,302796627 | 3,183816012 | 0,001453474 | 0,027366728 | The Moose: suction sampler vs FISH | NA |
| gene_1159808 | 6,99424931 | 5,530595206 | 1,854634133 | 2,982041097 | 0,002863336 | 0,040982187 | The Moose: suction sampler vs FISH | NA |
| gene_1160664 | 10,20025798 | -5,044600409 | 1,598821303 | -3,155199645 | 0,001603883 | 0,029057674 | The Moose: suction sampler vs FISH | NA |
| gene_1161855 | 3,603316988 | 6,16512298 | 1,919385747 | 3,212029156 | 0,00131801 | 0,02579303 | The Moose: suction sampler vs FISH | NA |
| gene_1162033 | 81,36523771 | -2,406005901 | 0,71590443 | -3,360792027 | 0,000777193 | 0,018587091 | The Moose: suction sampler vs FISH | NA |

| | | | | | | | | |
|---|---|---|---|---|---|---|---|---|
| gene_1162513 | 26,72967215 | 9,948069308 | 2,732762506 | 3,640297788 | 0,000272323 | 0,009545863 | The Moose: suction sampler vs FISH | NA |
| gene_1162562 | 8,445204639 | 6,309013428 | 1,606447016 | 3,927308753 | 8,59E-05 | 0,004520338 | The Moose: suction sampler vs FISH | NA |
| gene_1164627 | 38,66130683 | 3,125983276 | 0,832244044 | 3,756089692 | 0,000172589 | 0,007105263 | The Moose: suction sampler vs FISH | NA |
| gene_1166018 | 13,51064338 | 3,099025276 | 1,016267848 | 3,04941781 | 0,002292854 | 0,035783815 | The Moose: suction sampler vs FISH | NA |
| gene_1167954 | 163,6694186 | 3,563340557 | 0,915832684 | 3,890820474 | 9,99E-05 | 0,004993997 | The Moose: suction sampler vs FISH | NA |
| gene_116835 | 11,07740191 | 7,498435658 | 1,862588096 | 4,025815301 | 5,68E-05 | 0,003432163 | The Moose: suction sampler vs FISH | NA |
| gene_116836 | 147,6218696 | 4,75491792 | 0,814733542 | 5,836163207 | 5,34E-09 | 5,69E-06 | The Moose: suction sampler vs FISH | NA |
| gene_1170049 | 11,56356099 | 7,640114611 | 1,568209073 | 4,87187247 | 1,11E-06 | 0,000236834 | The Moose: suction sampler vs FISH | NA |
| gene_1170256 | 21,12469701 | 3,38652424 | 0,934760523 | 3,622878971 | 0,000291342 | 0,009984893 | The Moose: suction sampler vs FISH | NA |
| gene_1170398 | 6,259993864 | 5,1715484 | 1,636665154 | 3,15980846 | 0,001578729 | 0,028764511 | The Moose: suction sampler vs FISH | NA |
| gene_1170711 | 23,97911763 | 2,933040179 | 0,937547628 | 3,128417257 | 0,001757505 | 0,030695208 | The Moose: suction sampler vs FISH | NA |
| gene_1171032 | 64,18876534 | -2,904731625 | 0,943256517 | -3,079471569 | 0,002073682 | 0,033939377 | The Moose: suction sampler vs FISH | NA |
| gene_1171800 | 9,793373727 | 7,400989059 | 1,595106534 | 4,639808629 | 3,49E-06 | 0,000509197 | The Moose: suction sampler vs FISH | NA |
| gene_117206 | 4,957457364 | 5,930981777 | 2,023655598 | 2,930825671 | 0,003380624 | 0,04511082 | The Moose: suction sampler vs FISH | NA |
| gene_1173410 | 16,79012791 | 2,719035052 | 0,892891132 | 3,045203333 | 0,00232523 | 0,03608972 | The Moose: suction sampler vs FISH | NA |
| gene_117405 | 24,90663497 | 4,418877693 | 0,996939503 | 4,432443173 | 9,32E-06 | 0,001015276 | The Moose: suction sampler vs FISH | NA |
| gene_1174087 | 5,537852068 | -5,820308782 | 1,891110406 | -3,077720245 | 0,002085906 | 0,034074598 | The Moose: suction sampler vs FISH | NA |
| gene_1174926 | 3,74168909 | 6,008260557 | 1,817501343 | 3,305780533 | 0,000947122 | 0,021053694 | The Moose: suction sampler vs FISH | NA |
| gene_1176707 | 32,9066463 | 3,23490977 | 0,986624947 | 3,278763405 | 0,00104263 | 0,022357947 | The Moose: suction sampler vs FISH | NA |
| gene_1178500 | 20,31844367 | 7,101628151 | 1,504896243 | 4,719015136 | 2,37E-06 | 0,000390473 | The Moose: suction sampler vs FISH | NA |
| gene_117876 | 46,17283029 | 3,736603628 | 1,085222412 | 3,443168503 | 0,000574941 | 0,015255253 | The Moose: suction sampler vs FISH | NA |
| gene_1179243 | 18,34725884 | 3,61614845 | 0,949041579 | 3,810316142 | 0,000138789 | 0,00617406 | The Moose: suction sampler vs FISH | NA |
| gene_1180207 | 8,693923851 | 5,599901241 | 1,530997634 | 3,657681186 | 0,000254507 | 0,009127096 | The Moose: suction sampler vs FISH | NA |
| gene_1180648 | 4,18980309 | 6,970507342 | 2,310635376 | 3,016705887 | 0,002555376 | 0,0383176 | The Moose: suction sampler vs FISH | NA |
| gene_118090 | 90,16064527 | 2,678486165 | 0,781974056 | 3,425287761 | 0,000614149 | 0,015899239 | The Moose: suction sampler vs FISH | NA |
| gene_1180915 | 11,28243458 | 3,222924939 | 1,093923483 | 2,946206923 | 0,003216971 | 0,04392122 | The Moose: suction sampler vs FISH | NA |
| gene_1181968 | 6,326557854 | 4,924717946 | 1,580630254 | 3,115667268 | 0,001835292 | 0,031560066 | The Moose: suction sampler vs FISH | NA |
| gene_118309 | 22,86459418 | 2,897007144 | 0,941449742 | 3,077176628 | 0,002089714 | 0,034092895 | The Moose: suction sampler vs FISH | NA |
| gene_1183152 | 61,5510296 | 3,76622689 | 0,868024008 | 4,338851061 | 1,43E-05 | 0,001337845 | The Moose: suction sampler vs FISH | NA |
| gene_1183268 | 7,184720638 | 6,915575084 | 1,67201357 | 4,136075932 | 3,53E-05 | 0,002471217 | The Moose: suction sampler vs FISH | NA |
| gene_1184205 | 21,86474152 | 3,060146321 | 0,989877247 | 3,091440206 | 0,001991881 | 0,033090147 | The Moose: suction sampler vs FISH | NA |
| gene_1184600 | 4,612066274 | 6,951843257 | 2,149796086 | 3,233722166 | 0,001221883 | 0,024539704 | The Moose: suction sampler vs FISH | NA |
| gene_1185437 | 12,86442372 | 4,532559041 | 1,295119429 | 3,499722835 | 0,000465742 | 0,013354236 | The Moose: suction sampler vs FISH | NA |
| gene_1185796 | 53,73436353 | -2,77995974 | 0,958204726 | -2,901216895 | 0,003717165 | 0,047828868 | The Moose: suction sampler vs FISH | NA |
| gene_118587 | 9,676286266 | 4,340873824 | 1,398957327 | 3,102935123 | 0,001916116 | 0,032317394 | The Moose: suction sampler vs FISH | NA |
| gene_1185918 | 53,99089366 | 2,229252303 | 0,713239605 | 3,125530731 | 0,001774845 | 0,030852902 | The Moose: suction sampler vs FISH | NA |
| gene_1186403 | 5,961395683 | 5,922803105 | 1,884175165 | 3,143446116 | 0,001669712 | 0,029801206 | The Moose: suction sampler vs FISH | NA |
| gene_1186682 | 9,34897906 | 5,850868141 | 1,638699333 | 3,570434198 | 0,00035639 | 0,011331196 | The Moose: suction sampler vs FISH | NA |
| gene_118674 | 2,959243563 | 6,281166777 | 2,156028005 | 2,913304819 | 0,003576253 | 0,04666156 | The Moose: suction sampler vs FISH | NA |
| gene_1189042 | 15,47849843 | 4,019380985 | 1,009326314 | 3,982241351 | 6,83E-05 | 0,003872931 | The Moose: suction sampler vs FISH | NA |
| gene_1189378 | 18,91709699 | 3,579509547 | 0,932400951 | 3,839023911 | 0,000123524 | 0,005759871 | The Moose: suction sampler vs FISH | NA |
| gene_1189640 | 14,91898566 | 3,999434991 | 1,02485421 | 3,902442857 | 9,52E-05 | 0,004858804 | The Moose: suction sampler vs FISH | NA |
| gene_1190586 | 16,338204 | 2,890530164 | 0,939992549 | 3,075056463 | 0,002104627 | 0,034228152 | The Moose: suction sampler vs FISH | NA |
| gene_1190592 | 3,833654435 | 6,388760827 | 2,113752404 | 3,022473594 | 0,00250718 | 0,037874584 | The Moose: suction sampler vs FISH | NA |
| gene_1194279 | 27,30448763 | 5,107656316 | 1,168327344 | 4,371768186 | 1,23E-05 | 0,001216624 | The Moose: suction sampler vs FISH | NA |
| gene_1194402 | 17,45459154 | 3,158942776 | 0,869809758 | 3,631762862 | 0,000281492 | 0,009779641 | The Moose: suction sampler vs FISH | NA |
| gene_119494 | 7,253314628 | 7,43478504 | 1,755469106 | 4,235212693 | 2,28E-05 | 0,001828083 | The Moose: suction sampler vs FISH | NA |
| gene_1195431 | 52,38386058 | 4,256869556 | 0,907251177 | 4,692051841 | 2,70E-06 | 0,000427235 | The Moose: suction sampler vs FISH | NA |
| gene_1196207 | 8,07201497 | 5,544380425 | 1,636661876 | 3,387615062 | 0,000705031 | 0,017405388 | The Moose: suction sampler vs FISH | NA |
| gene_1196306 | 4,108405666 | 5,610084753 | 1,833823445 | 3,059228395 | 0,002219079 | 0,03520092 | The Moose: suction sampler vs FISH | NA |
| gene_1198581 | 8,50689973 | 6,392699203 | 1,740610588 | 3,672676271 | 0,000240023 | 0,008805457 | The Moose: suction sampler vs FISH | NA |
| gene_1200527 | 116,8448008 | 4,476865731 | 0,978641997 | 4,574569397 | 4,77E-06 | 0,000640731 | The Moose: suction sampler vs FISH | NA |
| gene_120096 | 28,64168155 | 2,962054467 | 0,89938495 | 3,29342232 | 0,000989757 | 0,021576935 | The Moose: suction sampler vs FISH | NA |
| gene_1201597 | 3,746753965 | 6,809380689 | 1,964806053 | 3,465675748 | 0,000528901 | 0,014467454 | The Moose: suction sampler vs FISH | NA |
| gene_1202409 | 4,18319963 | 6,853471428 | 2,028889567 | 3,377942073 | 0,000730305 | 0,017843388 | The Moose: suction sampler vs FISH | NA |
| gene_1202956 | 18,7052067 | 4,600740065 | 1,06849186 | 4,30582603 | 1,66E-05 | 0,001473895 | The Moose: suction sampler vs FISH | NA |
| gene_120342 | 6,939778984 | -6,049918352 | 2,062857815 | -2,932784949 | 0,003359365 | 0,044974217 | The Moose: suction sampler vs FISH | NA |
| gene_1205587 | 57,47980018 | 3,863247859 | 0,972368716 | 3,973027717 | 7,10E-05 | 0,003979153 | The Moose: suction sampler vs FISH | NA |
| gene_1206512 | 13,55592358 | 4,305600735 | 1,16292588 | 3,702386205 | 0,000213581 | 0,008156947 | The Moose: suction sampler vs FISH | NA |
| gene_1206596 | 3,553344673 | 7,154598341 | 2,212899535 | 3,233132922 | 0,001224406 | 0,024577857 | The Moose: suction sampler vs FISH | NA |
| gene_1207053 | 10,51693956 | 3,50084638 | 1,147498084 | 3,050851612 | 0,002281933 | 0,035719647 | The Moose: suction sampler vs FISH | NA |
| gene_1208048 | 5,738915868 | 7,025903726 | 1,789545353 | 3,926083077 | 8,63E-05 | 0,004532714 | The Moose: suction sampler vs FISH | NA |
| gene_1210541 | 28,45873918 | 2,675846859 | 0,841378242 | 3,180313828 | 0,001471156 | 0,027591695 | The Moose: suction sampler vs FISH | NA |
| gene_1210542 | 21,24280026 | 3,099011468 | 0,887623009 | 3,49136 | 0,000480568 | 0,013580352 | The Moose: suction sampler vs FISH | NA |
| gene_1210544 | 29,8279736 | 3,834846779 | 0,808775634 | 4,741545885 | 2,12E-06 | 0,00036142 | The Moose: suction sampler vs FISH | NA |

| | | | | | | | | |
|---|---|---|---|---|---|---|---|---|
| gene_1210545 | 15,14657529 | 3,041310577 | 0,986536066 | 3,082817427 | 0,002050509 | 0,033698734 | The Moose: suction sampler vs FISH | NA |
| gene_1210815 | 86,13740505 | 2,18943574 | 0,753451426 | 2,905875104 | 0,003662276 | 0,047388277 | The Moose: suction sampler vs FISH | NA |
| gene_1211281 | 20,90890203 | 3,378160078 | 0,874842708 | 3,861448517 | 0,000112717 | 0,005405093 | The Moose: suction sampler vs FISH | NA |
| gene_1211892 | 767,7615974 | 4,351122046 | 0,939072391 | 4,633425587 | 3,60E-06 | 0,000521016 | The Moose: suction sampler vs FISH | NA |
| gene_1212154 | 8,678588238 | 7,371828741 | 1,644872719 | 4,481701628 | 7,41E-06 | 0,000880035 | The Moose: suction sampler vs FISH | NA |
| gene_1212571 | 4,138459492 | 7,000939833 | 2,097395934 | 3,337919999 | 0,00084408 | 0,019667986 | The Moose: suction sampler vs FISH | NA |
| gene_1212746 | 5,141764319 | 7,027182541 | 1,735695478 | 4,048626403 | 5,15E-05 | 0,003223638 | The Moose: suction sampler vs FISH | NA |
| gene_1213840 | 4,015388371 | 6,900136915 | 2,092150869 | 3,298106756 | 0,000973391 | 0,02135326 | The Moose: suction sampler vs FISH | NA |
| gene_1214345 | 18,54315441 | 3,077764425 | 0,973674827 | 3,16097771 | 0,001572405 | 0,028696412 | The Moose: suction sampler vs FISH | NA |
| gene_1214346 | 9,630101344 | 6,966837564 | 1,566598794 | 4,447110255 | 8,70E-06 | 0,000967043 | The Moose: suction sampler vs FISH | NA |
| gene_1214349 | 11,04469197 | 5,147838107 | 1,345178593 | 3,826880782 | 0,000129777 | 0,005918523 | The Moose: suction sampler vs FISH | NA |
| gene_1214836 | 25,23327996 | 2,984064425 | 1,008309403 | 2,959472971 | 0,003081657 | 0,042882214 | The Moose: suction sampler vs FISH | NA |
| gene_121525 | 42,3455854 | 2,86423709 | 0,758430063 | 3,776534226 | 0,000159026 | 0,006747157 | The Moose: suction sampler vs FISH | NA |
| gene_121526 | 6,322623559 | 6,194237797 | 1,751775113 | 3,535977735 | 0,000406269 | 0,012287256 | The Moose: suction sampler vs FISH | NA |
| gene_1215445 | 16,82644576 | 3,183045963 | 1,045870596 | 3,043441489 | 0,002338889 | 0,036216894 | The Moose: suction sampler vs FISH | NA |
| gene_1215448 | 18,33567862 | 3,269051319 | 0,917281484 | 3,563847495 | 0,000365458 | 0,011490192 | The Moose: suction sampler vs FISH | NA |
| gene_121778 | 7,239569043 | 4,727734666 | 1,447198417 | 3,266818572 | 0,001087634 | 0,022967582 | The Moose: suction sampler vs FISH | NA |
| gene_1218485 | 43,70371054 | 5,313751757 | 1,170326825 | 4,540399863 | 5,61E-06 | 0,00071469 | The Moose: suction sampler vs FISH | NA |
| gene_1221071 | 34,3430566 | 2,87520019 | 0,850228661 | 3,381678744 | 0,000720443 | 0,017685616 | The Moose: suction sampler vs FISH | NA |
| gene_1221409 | 3,933493495 | 6,809465593 | 2,070767792 | 3,2883772 | 0,001007667 | 0,021818264 | The Moose: suction sampler vs FISH | NA |
| gene_1222327 | 61,71204874 | 4,173327471 | 0,881684304 | 4,733358021 | 2,21E-06 | 0,000370303 | The Moose: suction sampler vs FISH | NA |
| gene_1222632 | 33,6132409 | 3,284833607 | 0,779866992 | 4,212043389 | 2,53E-05 | 0,001981534 | The Moose: suction sampler vs FISH | NA |
| gene_1222955 | 21,31975444 | 3,564395509 | 1,025992933 | 3,474093627 | 0,000512582 | 0,014178245 | The Moose: suction sampler vs FISH | NA |
| gene_1224718 | 10,37946457 | 4,508702144 | 1,232148581 | 3,659219523 | 0,000252985 | 0,00909469 | The Moose: suction sampler vs FISH | NA |
| gene_1224783 | 95,46925282 | -2,615160755 | 0,723562427 | -3,614284901 | 0,000301178 | 0,010226537 | The Moose: suction sampler vs FISH | NA |
| gene_1225119 | 35,59234741 | -3,714100198 | 0,994201674 | -3,735761362 | 0,000187148 | 0,007482358 | The Moose: suction sampler vs FISH | NA |
| gene_1225120 | 25,68660509 | -2,740932302 | 0,937693848 | -2,923056718 | 0,003466134 | 0,045802481 | The Moose: suction sampler vs FISH | NA |
| gene_1225128 | 32,20693159 | -3,073272237 | 0,984663561 | -3,121139402 | 0,001801527 | 0,031137829 | The Moose: suction sampler vs FISH | NA |
| gene_1225442 | 17,340852 | 3,568346087 | 0,988024851 | 3,61159548 | 0,000304319 | 0,010285388 | The Moose: suction sampler vs FISH | NA |
| gene_1226069 | 7,186520312 | 6,566973646 | 1,808512761 | 3,631145871 | 0,000282166 | 0,009793788 | The Moose: suction sampler vs FISH | NA |
| gene_1226325 | 15,7599447 | 5,408303151 | 1,202218711 | 4,498601713 | 6,84E-06 | 0,00083296 | The Moose: suction sampler vs FISH | NA |
| gene_1226351 | 6,845117982 | 4,673642644 | 1,614583074 | 2,894643651 | 0,003795892 | 0,048350551 | The Moose: suction sampler vs FISH | NA |
| gene_1227402 | 4,673960129 | 5,689676164 | 1,686931069 | 3,372797067 | 0,000744088 | 0,01807791 | The Moose: suction sampler vs FISH | NA |
| gene_1227856 | 20,26107109 | 4,174627561 | 1,155529825 | 3,612738909 | 0,00030298 | 0,010271678 | The Moose: suction sampler vs FISH | NA |
| gene_122818 | 109,334095 | -2,632053311 | 0,894014404 | -2,944083785 | 0,003239123 | 0,044090776 | The Moose: suction sampler vs FISH | NA |
| gene_1228182 | 24,63910691 | 3,199143039 | 1,106840968 | 2,890336672 | 0,003848294 | 0,048731896 | The Moose: suction sampler vs FISH | NA |
| gene_1228905 | 28,3372205 | 2,850139668 | 0,952430042 | 2,992492407 | 0,002767096 | 0,040060976 | The Moose: suction sampler vs FISH | NA |
| gene_1228906 | 22,99889658 | 2,80799129 | 0,970604214 | 2,893034308 | 0,003815396 | 0,048472889 | The Moose: suction sampler vs FISH | NA |
| gene_1228907 | 103,1625348 | 2,213170012 | 0,720293706 | 3,072593853 | 0,002122071 | 0,034367254 | The Moose: suction sampler vs FISH | NA |
| gene_1228908 | 25,22764471 | 3,217453953 | 0,93914604 | 3,425935708 | 0,000612685 | 0,01587815 | The Moose: suction sampler vs FISH | NA |
| gene_122932 | 49,34599167 | 2,903992853 | 0,964566954 | 3,010670063 | 0,002606719 | 0,038790953 | The Moose: suction sampler vs FISH | NA |
| gene_1229761 | 6,194419522 | 5,137237428 | 1,672354428 | 3,071859255 | 0,0021273 | 0,034429165 | The Moose: suction sampler vs FISH | NA |
| gene_123029 | 3,617140865 | 5,865645586 | 1,873419137 | 3,130984129 | 0,001742216 | 0,030575158 | The Moose: suction sampler vs FISH | NA |
| gene_1230679 | 6,562944498 | 7,372836091 | 1,699447236 | 4,338373051 | 1,44E-05 | 0,001337845 | The Moose: suction sampler vs FISH | NA |
| gene_1230680 | 13,65220336 | 6,385389089 | 1,383535639 | 4,615268959 | 3,93E-06 | 0,000554505 | The Moose: suction sampler vs FISH | NA |
| gene_123271 | 14507,79746 | 2,497144287 | 0,844736344 | 2,956122705 | 0,003115331 | 0,043133087 | The Moose: suction sampler vs FISH | NA |
| gene_123272 | 2481748,494 | 2,94228765 | 0,817573007 | 3,598807232 | 0,00031968 | 0,010599955 | The Moose: suction sampler vs FISH | NA |
| gene_123275 | 4259592,332 | 3,292960447 | 0,803823282 | 4,096622382 | 4,19E-05 | 0,002796526 | The Moose: suction sampler vs FISH | NA |
| gene_1232816 | 19,2470009 | 3,23014452 | 1,057974072 | 3,053141476 | 0,002264591 | 0,035551536 | The Moose: suction sampler vs FISH | NA |
| gene_1233273 | 8,211819304 | 7,897288554 | 1,999516511 | 3,949599071 | 7,83E-05 | 0,004249399 | The Moose: suction sampler vs FISH | NA |
| gene_1233937 | 9,037406063 | 3,524797313 | 1,170849251 | 3,010462115 | 0,002608505 | 0,038790953 | The Moose: suction sampler vs FISH | NA |
| gene_1235524 | 15,39343676 | 3,294911811 | 0,974640049 | 3,380644798 | 0,00072316 | 0,017704153 | The Moose: suction sampler vs FISH | NA |
| gene_1235946 | 6,619713084 | 4,653284451 | 1,499214073 | 3,103815884 | 0,001910422 | 0,032281658 | The Moose: suction sampler vs FISH | NA |
| gene_1236940 | 5,303362579 | 6,499474055 | 2,039513955 | 3,186775967 | 0,001438682 | 0,027176948 | The Moose: suction sampler vs FISH | NA |
| gene_1237013 | 27,61133659 | 3,098657393 | 0,812154979 | 3,815352334 | 0,000135989 | 0,006113186 | The Moose: suction sampler vs FISH | NA |
| gene_1237014 | 25,76942297 | 4,193501739 | 0,835882052 | 5,016858212 | 5,25E-07 | 0,000143894 | The Moose: suction sampler vs FISH | NA |
| gene_1237018 | 30,28596659 | 2,346225296 | 0,745971991 | 3,14519221 | 0,001659777 | 0,029688834 | The Moose: suction sampler vs FISH | NA |
| gene_1237020 | 32,61668458 | 3,053053063 | 0,949302857 | 3,216100152 | 0,001299455 | 0,025546445 | The Moose: suction sampler vs FISH | NA |
| gene_1237657 | 2,950067655 | 6,142421711 | 2,126290029 | 2,888797683 | 0,003867178 | 0,048863897 | The Moose: suction sampler vs FISH | NA |
| gene_1238304 | 21,56534311 | 3,003217443 | 1,017061284 | 2,952838231 | 0,003148669 | 0,043362837 | The Moose: suction sampler vs FISH | NA |
| gene_1238774 | 499,2841907 | 3,256488554 | 0,687551932 | 4,73635285 | 2,18E-06 | 0,000366549 | The Moose: suction sampler vs FISH | NA |
| gene_1238829 | 50,75230069 | 10,77510088 | 2,726222975 | 3,952391636 | 7,74E-05 | 0,00422194 | The Moose: suction sampler vs FISH | NA |
| gene_1238830 | 62,59723641 | 11,04326604 | 3,06600107 | 3,601846768 | 0,000315965 | 0,010552988 | The Moose: suction sampler vs FISH | NA |
| gene_1239393 | 40,78094698 | 3,195037086 | 0,747144565 | 4,276330495 | 1,90E-05 | 0,001596631 | The Moose: suction sampler vs FISH | NA |
| gene_1239872 | 3,92567898 | 5,958729012 | 2,010498448 | 2,96380682 | 0,00303859 | 0,042472454 | The Moose: suction sampler vs FISH | NA |

| | | | | | | | | |
|---|---|---|---|---|---|---|---|---|
| gene_1239925 | 223,8144837 | 4,636249738 | 1,106540918 | 4,189858378 | 2,79E-05 | 0,002120021 | The Moose: suction sampler vs FISH | NA |
| gene_1240023 | 6,983953361 | 3,549118693 | 1,225029126 | 2,897170865 | 0,003765446 | 0,048146447 | The Moose: suction sampler vs FISH | NA |
| gene_1240057 | 2,784202153 | 5,914250073 | 2,053793369 | 2,879671423 | 0,003980898 | 0,049647999 | The Moose: suction sampler vs FISH | NA |
| gene_1242036 | 10,70223125 | 8,683674565 | 1,665841761 | 5,212784771 | 1,86E-07 | 6,83E-05 | The Moose: suction sampler vs FISH | NA |
| gene_1242037 | 29,58528463 | 9,248920151 | 1,443459046 | 6,407469733 | 1,48E-10 | 4,72E-07 | The Moose: suction sampler vs FISH | NA |
| gene_124328 | 24,66610625 | 5,597419474 | 1,084424275 | 5,161650843 | 2,45E-07 | 8,56E-05 | The Moose: suction sampler vs FISH | NA |
| gene_124329 | 37,75248767 | 4,872794275 | 1,080399468 | 4,510178334 | 6,48E-06 | 0,000800886 | The Moose: suction sampler vs FISH | NA |
| gene_1243561 | 11,5650226 | 6,860497654 | 1,541839402 | 4,449553983 | 8,60E-06 | 0,000962933 | The Moose: suction sampler vs FISH | NA |
| gene_1243697 | 6,223192003 | 4,401094235 | 1,514842997 | 2,905313781 | 0,003668851 | 0,047425091 | The Moose: suction sampler vs FISH | NA |
| gene_1244457 | 7,511821941 | 6,539506588 | 1,750834692 | 3,735079399 | 0,000187656 | 0,007498582 | The Moose: suction sampler vs FISH | NA |
| gene_1244476 | 26,31867894 | 4,445114345 | 0,919929958 | 4,832013898 | 1,35E-06 | 0,000271965 | The Moose: suction sampler vs FISH | NA |
| gene_1246830 | 4,520054904 | 6,220352129 | 1,937994096 | 3,209685799 | 0,001328801 | 0,025868826 | The Moose: suction sampler vs FISH | NA |
| gene_1247086 | 67,40920434 | 4,522708469 | 1,00752892 | 4,488911811 | 7,16E-06 | 0,000859114 | The Moose: suction sampler vs FISH | NA |
| gene_1247174 | 14,25396805 | 4,076016551 | 1,081658519 | 3,768302545 | 0,000164361 | 0,006894075 | The Moose: suction sampler vs FISH | NA |
| gene_124740 | 71,81404103 | 2,889105734 | 0,845837878 | 3,415673155 | 0,000636246 | 0,016293257 | The Moose: suction sampler vs FISH | NA |
| gene_1247781 | 52,52349926 | 2,686844512 | 0,753770709 | 3,564538233 | 0,000364497 | 0,011490192 | The Moose: suction sampler vs FISH | NA |
| gene_1248373 | 13,97365694 | 8,57975544 | 2,900815362 | 2,957704773 | 0,003099388 | 0,043012037 | The Moose: suction sampler vs FISH | NA |
| gene_1248477 | 8,81825667 | 4,842595512 | 1,428280041 | 3,390508424 | 0,000697631 | 0,017257485 | The Moose: suction sampler vs FISH | NA |
| gene_1248519 | 3,570632255 | 5,825578255 | 1,964518878 | 2,965396932 | 0,003022926 | 0,042362843 | The Moose: suction sampler vs FISH | NA |
| gene_1249748 | 5,53807388 | 5,895801809 | 2,042653257 | 2,886344899 | 0,003897448 | 0,049048664 | The Moose: suction sampler vs FISH | NA |
| gene_1249978 | 7,942058425 | 5,720092471 | 1,621889422 | 3,526807926 | 0,000420602 | 0,01257792 | The Moose: suction sampler vs FISH | NA |
| gene_1250285 | 58,84427276 | 2,730437119 | 0,843482788 | 3,237098798 | 0,001207516 | 0,02442468 | The Moose: suction sampler vs FISH | NA |
| gene_1250600 | 23,56884513 | 2,603740083 | 0,898084304 | 2,899215666 | 0,003740975 | 0,048009067 | The Moose: suction sampler vs FISH | NA |
| gene_1250748 | 8,127613262 | 4,903552786 | 1,551052978 | 3,161434752 | 0,00156994 | 0,02868807 | The Moose: suction sampler vs FISH | NA |
| gene_1251678 | 5,686360267 | -5,288460764 | 1,682478263 | -3,143256517 | 0,001670794 | 0,029806031 | The Moose: suction sampler vs FISH | NA |
| gene_1251719 | 89,87317926 | 3,486037803 | 0,850013961 | 4,101153585 | 4,11E-05 | 0,002754828 | The Moose: suction sampler vs FISH | NA |
| gene_1252390 | 13,33643325 | 3,93754373 | 1,140046911 | 3,453843603 | 0,000552658 | 0,014846361 | The Moose: suction sampler vs FISH | NA |
| gene_1252920 | 10,57836415 | 7,982945351 | 1,725629639 | 4,62610584 | 3,73E-06 | 0,000534492 | The Moose: suction sampler vs FISH | NA |
| gene_1253455 | 31,85417751 | 3,136269284 | 0,914496974 | 3,42950209 | 0,00060469 | 0,015726431 | The Moose: suction sampler vs FISH | NA |
| gene_1253800 | 4,482424641 | 5,832895381 | 1,773924529 | 3,288130518 | 0,001008551 | 0,021818264 | The Moose: suction sampler vs FISH | NA |
| gene_1253910 | 46,73450876 | 3,823360729 | 0,957910594 | 3,991354466 | 6,57E-05 | 0,003757879 | The Moose: suction sampler vs FISH | NA |
| gene_1254253 | 12,33141293 | 3,503332744 | 1,150362176 | 3,045417189 | 0,002323578 | 0,036082789 | The Moose: suction sampler vs FISH | NA |
| gene_1254618 | 12,37776935 | 3,45569825 | 1,14834327 | 3,009290289 | 0,002618588 | 0,038860817 | The Moose: suction sampler vs FISH | NA |
| gene_1255336 | 159,8073276 | 2,570865207 | 0,711981367 | 3,610860236 | 0,000305183 | 0,010291574 | The Moose: suction sampler vs FISH | NA |
| gene_1255352 | 10,59754506 | 5,336639897 | 1,461473922 | 3,65154644 | 0,000260666 | 0,009279986 | The Moose: suction sampler vs FISH | NA |
| gene_1256661 | 9,898493571 | 6,134712042 | 1,351754166 | 4,538334112 | 5,67E-06 | 0,000717993 | The Moose: suction sampler vs FISH | NA |
| gene_1256882 | 4,321572323 | 6,57600488 | 1,961924606 | 3,351813244 | 0,000802842 | 0,018965252 | The Moose: suction sampler vs FISH | NA |
| gene_125702 | 3,652797026 | 6,882018922 | 2,160940188 | 3,184733645 | 0,001448873 | 0,027306255 | The Moose: suction sampler vs FISH | NA |
| gene_1259762 | 19,72230419 | 3,875344211 | 0,975436771 | 3,972932254 | 7,10E-05 | 0,003979153 | The Moose: suction sampler vs FISH | NA |
| gene_1259913 | 3,894671901 | 6,418270056 | 2,164420231 | 2,965353014 | 0,003023358 | 0,042362843 | The Moose: suction sampler vs FISH | NA |
| gene_1261413 | 4,885332145 | 5,697465535 | 1,917372449 | 2,971496507 | 0,002963522 | 0,041808664 | The Moose: suction sampler vs FISH | NA |
| gene_1262160 | 24,1528701 | 3,243823577 | 0,962987677 | 3,368499573 | 0,000755785 | 0,018265429 | The Moose: suction sampler vs FISH | NA |
| gene_1263208 | 19,2307951 | 4,949485366 | 1,498222443 | 3,303571769 | 0,000954615 | 0,021124351 | The Moose: suction sampler vs FISH | NA |
| gene_126416 | 34,69670347 | 4,15258887 | 1,087879195 | 3,817141542 | 0,000135007 | 0,006083171 | The Moose: suction sampler vs FISH | NA |
| gene_126417 | 20,52054857 | 3,801807914 | 1,097731597 | 3,463331039 | 0,000533532 | 0,014534579 | The Moose: suction sampler vs FISH | NA |
| gene_1264560 | 5,563205067 | 4,401677169 | 1,52826036 | 2,880188013 | 0,003974381 | 0,049608838 | The Moose: suction sampler vs FISH | NA |
| gene_1265372 | 8,215235142 | 4,450171707 | 1,259834551 | 3,532346136 | 0,00041189 | 0,012407978 | The Moose: suction sampler vs FISH | NA |
| gene_1265427 | 15,41593485 | 3,733381674 | 0,99806235 | 3,740629707 | 0,00018356 | 0,007375027 | The Moose: suction sampler vs FISH | NA |
| gene_1265801 | 4,37158198 | 6,012150404 | 1,876144503 | 3,204524169 | 0,001352859 | 0,026154449 | The Moose: suction sampler vs FISH | NA |
| gene_1267174 | 7,652567177 | 4,408233741 | 1,466912973 | 3,005109249 | 0,002654855 | 0,039075312 | The Moose: suction sampler vs FISH | NA |
| gene_1267368 | 5,788014955 | 7,850339227 | 2,013988371 | 3,89790693 | 9,70E-05 | 0,004911233 | The Moose: suction sampler vs FISH | NA |
| gene_1267523 | 181,0330888 | 5,586290397 | 0,86827594 | 6,433773113 | 1,24E-10 | 4,64E-07 | The Moose: suction sampler vs FISH | NA |
| gene_1267707 | 11,88981631 | 4,113568293 | 1,170802193 | 3,513461385 | 0,000442309 | 0,012950027 | The Moose: suction sampler vs FISH | NA |
| gene_1267714 | 13,4843177 | -3,909327743 | 1,321089976 | -2,959168424 | 0,003084705 | 0,042900233 | The Moose: suction sampler vs FISH | NA |
| gene_1267718 | 16,27160926 | -4,090395621 | 1,341994853 | -3,047996505 | 0,002303726 | 0,035907717 | The Moose: suction sampler vs FISH | NA |
| gene_126840 | 25,15577334 | 5,357452034 | 1,142382588 | 4,689717868 | 2,74E-06 | 0,000428233 | The Moose: suction sampler vs FISH | NA |
| gene_126952 | 7,935666807 | 4,215278868 | 1,437058619 | 2,933268561 | 0,003354136 | 0,044952336 | The Moose: suction sampler vs FISH | NA |
| gene_1271938 | 11,31311635 | 4,570187606 | 1,324431674 | 3,450678276 | 0,00055918 | 0,014977737 | The Moose: suction sampler vs FISH | NA |
| gene_1272306 | 17,31956197 | 5,33434706 | 1,231650919 | 4,331054342 | 1,48E-05 | 0,001367506 | The Moose: suction sampler vs FISH | NA |
| gene_1272529 | 27,23424946 | 2,686275366 | 0,898943101 | 2,988259617 | 0,002805711 | 0,040470508 | The Moose: suction sampler vs FISH | NA |
| gene_127271 | 5,301658959 | 4,916476373 | 1,660507453 | 2,960827646 | 0,003068136 | 0,042750756 | The Moose: suction sampler vs FISH | NA |
| gene_1273851 | 22,42294207 | 3,646407828 | 0,851006155 | 4,284819572 | 1,83E-05 | 0,001563699 | The Moose: suction sampler vs FISH | NA |
| gene_1273852 | 37,38391587 | 2,470259313 | 0,827436373 | 2,985437182 | 0,002831733 | 0,040747771 | The Moose: suction sampler vs FISH | NA |
| gene_1273876 | 15,21496398 | 4,174131004 | 1,162287625 | 3,591306416 | 0,000329025 | 0,010797679 | The Moose: suction sampler vs FISH | NA |
| gene_1274238 | 3,725248315 | 6,21804384 | 2,014149563 | 3,08718079 | 0,002020647 | 0,033372249 | The Moose: suction sampler vs FISH | NA |

| | | | | | | | | |
|---|---|---|---|---|---|---|---|---|
| gene_1274239 | 5,839329315 | 6,782339874 | 1,803750159 | 3,760132654 | 0,000169823 | 0,007031351 | The Moose: suction sampler vs FISH | NA |
| gene_1274355 | 19,02032225 | 3,079729719 | 0,970441819 | 3,173533598 | 0,001505955 | 0,027998858 | The Moose: suction sampler vs FISH | NA |
| gene_1275107 | 13,60682982 | 4,025106783 | 1,200571491 | 3,352658974 | 0,000800393 | 0,018932314 | The Moose: suction sampler vs FISH | NA |
| gene_1275157 | 15,13156605 | 3,590036494 | 1,124304438 | 3,193117784 | 0,001407455 | 0,026779935 | The Moose: suction sampler vs FISH | NA |
| gene_1277269 | 7,243769379 | 6,325327755 | 1,811200039 | 3,492340779 | 0,000478807 | 0,013571764 | The Moose: suction sampler vs FISH | NA |
| gene_1277491 | 3,122595014 | 6,645479069 | 2,300577656 | 2,888613236 | 0,003869447 | 0,048867143 | The Moose: suction sampler vs FISH | NA |
| gene_1277636 | 12,65314029 | 3,723229447 | 1,040042077 | 3,579883476 | 0,000343747 | 0,011077899 | The Moose: suction sampler vs FISH | NA |
| gene_1278559 | 6,548915907 | 3,848385604 | 1,291041127 | 2,980838892 | 0,0028746 | 0,041074899 | The Moose: suction sampler vs FISH | NA |
| gene_1281137 | 8,190209116 | 6,015636155 | 1,719704921 | 3,498063 | 0,00046865 | 0,013392145 | The Moose: suction sampler vs FISH | NA |
| gene_1281633 | 6,755996282 | -5,363291906 | 1,7916494 | -2,9934941 | 0,002758028 | 0,039984879 | The Moose: suction sampler vs FISH | NA |
| gene_1282234 | 6,429981434 | 5,463233775 | 1,484755677 | 3,679550689 | 0,000233645 | 0,008653593 | The Moose: suction sampler vs FISH | NA |
| gene_1283377 | 8,138053176 | 4,721700428 | 1,31289781 | 3,596395997 | 0,000322657 | 0,010660194 | The Moose: suction sampler vs FISH | NA |
| gene_1283662 | 14,94196367 | 2,778377601 | 0,962137754 | 2,887712897 | 0,003880539 | 0,048953312 | The Moose: suction sampler vs FISH | NA |
| gene_1284450 | 2,853297848 | 5,857384228 | 1,955248896 | 2,995723071 | 0,00273795 | 0,039788031 | The Moose: suction sampler vs FISH | NA |
| gene_1285161 | 7,797726279 | 4,035708156 | 1,385774143 | 2,912240914 | 0,003588458 | 0,046779244 | The Moose: suction sampler vs FISH | NA |
| gene_1285343 | 11,22735871 | 4,446056055 | 1,360927651 | 3,266930501 | 0,001087204 | 0,022967582 | The Moose: suction sampler vs FISH | NA |
| gene_1285929 | 15,82524344 | 4,129160743 | 1,308973981 | 3,154501772 | 0,001607724 | 0,029115558 | The Moose: suction sampler vs FISH | NA |
| gene_128625 | 12,8264086 | -3,567461681 | 1,191241111 | -2,994743589 | 0,002746756 | 0,039845057 | The Moose: suction sampler vs FISH | NA |
| gene_1286549 | 5,363691708 | 5,701414264 | 1,875638415 | 3,039719287 | 0,002367987 | 0,036498812 | The Moose: suction sampler vs FISH | NA |
| gene_1288255 | 7,3762176 | 7,643445229 | 1,91486282 | 3,99164115 | 6,56E-05 | 0,003756263 | The Moose: suction sampler vs FISH | NA |
| gene_1288546 | 8,163750932 | 4,933139701 | 1,703799071 | 2,895376446 | 0,003787041 | 0,048288057 | The Moose: suction sampler vs FISH | NA |
| gene_1289037 | 46,25050784 | 2,363249648 | 0,773634028 | 3,054738496 | 0,002252568 | 0,035451731 | The Moose: suction sampler vs FISH | NA |
| gene_1289633 | 3,244529729 | 5,728848508 | 1,989254858 | 2,879896704 | 0,003978055 | 0,049620964 | The Moose: suction sampler vs FISH | NA |
| gene_1289830 | 7,507269962 | -4,70390898 | 1,559596361 | -3,016106665 | 0,002560432 | 0,038357374 | The Moose: suction sampler vs FISH | NA |
| gene_129010 | 11,28116693 | 3,185629666 | 1,058672351 | 3,009079874 | 0,002620402 | 0,038864183 | The Moose: suction sampler vs FISH | NA |
| gene_1290838 | 8,359359475 | 3,86488212 | 1,235313759 | 3,12866435 | 0,001756028 | 0,030685572 | The Moose: suction sampler vs FISH | NA |
| gene_1290947 | 16,06557871 | 5,906500689 | 1,294462294 | 4,562898988 | 5,05E-06 | 0,000660549 | The Moose: suction sampler vs FISH | NA |
| gene_1291091 | 255,9631017 | 4,116742558 | 0,891984295 | 4,615263497 | 3,93E-06 | 0,000554505 | The Moose: suction sampler vs FISH | NA |
| gene_1291530 | 15,82853046 | 3,71178542 | 0,956169394 | 3,88193289 | 0,000103629 | 0,005101251 | The Moose: suction sampler vs FISH | NA |
| gene_1292085 | 24,62969175 | 2,916853616 | 0,953701942 | 3,058454102 | 0,002224822 | 0,035225748 | The Moose: suction sampler vs FISH | NA |
| gene_12927 | 61,59262177 | 4,064335782 | 0,854563882 | 4,756035057 | 1,97E-06 | 0,000347141 | The Moose: suction sampler vs FISH | NA |
| gene_1292790 | 52,70009624 | 3,480527458 | 1,065971123 | 3,265123588 | 0,001094163 | 0,023059059 | The Moose: suction sampler vs FISH | NA |
| gene_1292791 | 52,8996994 | 4,103813507 | 1,170027554 | 3,507450308 | 0,000452423 | 0,0131233 | The Moose: suction sampler vs FISH | NA |
| gene_1293370 | 34,21736058 | 3,765642767 | 0,972211851 | 3,873273879 | 0,000107383 | 0,005222096 | The Moose: suction sampler vs FISH | NA |
| gene_1293593 | 56,13006102 | -2,139986854 | 0,693615177 | -3,085265326 | 0,002033706 | 0,033527625 | The Moose: suction sampler vs FISH | NA |
| gene_1294374 | 23,25023873 | -4,302675837 | 1,275492949 | -3,37334349 | 0,000742613 | 0,018054014 | The Moose: suction sampler vs FISH | NA |
| gene_1294940 | 95,3397462 | 4,606108526 | 0,830800062 | 5,544184138 | 2,95E-08 | 1,97E-05 | The Moose: suction sampler vs FISH | NA |
| gene_1295757 | 5,641728448 | 7,035888134 | 1,793575386 | 3,922828217 | 8,75E-05 | 0,004572734 | The Moose: suction sampler vs FISH | NA |
| gene_1295758 | 7,80139337 | 5,5463677 | 1,793968631 | 3,091674852 | 0,001990307 | 0,033077161 | The Moose: suction sampler vs FISH | NA |
| gene_1295837 | 18,93305223 | 3,112434352 | 1,008706874 | 3,085568695 | 0,002031633 | 0,033503497 | The Moose: suction sampler vs FISH | NA |
| gene_1296032 | 9,574807655 | 3,836210652 | 1,08165468 | 3,546613093 | 0,000390217 | 0,011996443 | The Moose: suction sampler vs FISH | NA |
| gene_1296436 | 4,230022466 | 6,546862468 | 1,902634815 | 3,440945375 | 0,000579686 | 0,015331295 | The Moose: suction sampler vs FISH | NA |
| gene_1296437 | 10,99872306 | 7,530024141 | 1,763339329 | 4,270320532 | 1,95E-05 | 0,00162723 | The Moose: suction sampler vs FISH | NA |
| gene_1296543 | 4,567441083 | 5,481307554 | 1,755380763 | 3,122574697 | 0,001792766 | 0,031046853 | The Moose: suction sampler vs FISH | NA |
| gene_129657 | 13,14593939 | 3,755172027 | 1,111206072 | 3,379366007 | 0,000726532 | 0,017768949 | The Moose: suction sampler vs FISH | NA |
| gene_1298262 | 8,061651741 | 5,911460578 | 1,572135163 | 3,760147801 | 0,000169813 | 0,007031351 | The Moose: suction sampler vs FISH | NA |
| gene_1299194 | 35,12502636 | 2,601127246 | 0,821956466 | 3,164555977 | 0,001553198 | 0,028468833 | The Moose: suction sampler vs FISH | NA |
| gene_1299468 | 272,6761421 | -2,146658697 | 0,6106303 | -3,515480146 | 0,00043896 | 0,012910361 | The Moose: suction sampler vs FISH | NA |
| gene_1300135 | 13,80926172 | 4,350334363 | 1,089390623 | 3,993364978 | 6,51E-05 | 0,003740709 | The Moose: suction sampler vs FISH | NA |
| gene_1300277 | 8,792740466 | 3,9372231 | 1,319087373 | 2,984808421 | 0,00283756 | 0,040790747 | The Moose: suction sampler vs FISH | NA |
| gene_1301044 | 24,9665619 | 3,1325322 | 0,852916025 | 3,67273226 | 0,000239971 | 0,008805457 | The Moose: suction sampler vs FISH | NA |
| gene_1301284 | 5,078681512 | 5,803639279 | 1,975152187 | 2,938325116 | 0,003299908 | 0,044559979 | The Moose: suction sampler vs FISH | NA |
| gene_130159 | 13,76627798 | -3,391022193 | 1,087354472 | -3,118598654 | 0,001817133 | 0,031347689 | The Moose: suction sampler vs FISH | NA |
| gene_130178 | 14,79083013 | 4,131422275 | 1,144229874 | 3,610657587 | 0,000305422 | 0,010291574 | The Moose: suction sampler vs FISH | NA |
| gene_1303250 | 14,18408027 | 7,66029645 | 1,697668837 | 4,512244252 | 6,41E-06 | 0,00079446 | The Moose: suction sampler vs FISH | NA |
| gene_1303287 | 6,592856297 | 5,837117178 | 1,7460173 | 3,343103861 | 0,000828469 | 0,019384165 | The Moose: suction sampler vs FISH | NA |
| gene_1304151 | 8,746265797 | 7,411997821 | 1,659806915 | 4,465578347 | 7,99E-06 | 0,000917807 | The Moose: suction sampler vs FISH | NA |
| gene_1304369 | 4,098470461 | 5,686366406 | 1,795491949 | 3,167024174 | 0,001540075 | 0,028358408 | The Moose: suction sampler vs FISH | NA |
| gene_1304519 | 13,62991788 | 3,205243017 | 1,110849242 | 2,885398753 | 0,003909182 | 0,049120591 | The Moose: suction sampler vs FISH | NA |
| gene_1304521 | 4,176592414 | 6,393737345 | 2,191600725 | 2,917382382 | 0,003529827 | 0,046261274 | The Moose: suction sampler vs FISH | NA |
| gene_1304690 | 38,33612788 | 3,170366448 | 0,876830737 | 3,615710896 | 0,000299525 | 0,010192304 | The Moose: suction sampler vs FISH | NA |
| gene_1305574 | 19,08983895 | 4,681176664 | 1,087414283 | 4,304869576 | 1,67E-05 | 0,001478388 | The Moose: suction sampler vs FISH | NA |
| gene_1305575 | 12,7842734 | 3,797004863 | 1,183334273 | 3,208733957 | 0,001333208 | 0,025904089 | The Moose: suction sampler vs FISH | NA |
| gene_1306105 | 14,35328403 | 3,553566669 | 1,038172104 | 3,422907104 | 0,000619552 | 0,01598537 | The Moose: suction sampler vs FISH | NA |
| gene_1306617 | 10,87288487 | -3,661247771 | 1,199036032 | -3,053492701 | 0,002261942 | 0,035545359 | The Moose: suction sampler vs FISH | NA |

| | | | | | | | | |
|---|---|---|---|---|---|---|---|---|
| gene_1307860 | 11,15253388 | -3,894958872 | 1,329935523 | -2,928682485 | 0,00340402 | 0,045330064 | The Moose: suction sampler vs FISH | NA |
| gene_130876 | 9,460349573 | 3,416077203 | 1,158368083 | 2,949042927 | 0,003187597 | 0,04367781 | The Moose: suction sampler vs FISH | NA |
| gene_1308843 | 23,67731125 | 5,782292141 | 1,374258605 | 4,207572083 | 2,58E-05 | 0,002007468 | The Moose: suction sampler vs FISH | NA |
| gene_1309000 | 5,164499221 | -4,964584831 | 1,697171594 | -2,925210892 | 0,003442229 | 0,045647426 | The Moose: suction sampler vs FISH | NA |
| gene_1309285 | 16,83743057 | 8,332971305 | 1,581265508 | 5,269811592 | 1,37E-07 | 5,51E-05 | The Moose: suction sampler vs FISH | NA |
| gene_1309489 | 14,81244135 | 4,158616465 | 1,222258613 | 3,402403077 | 0,00066796 | 0,016789307 | The Moose: suction sampler vs FISH | NA |
| gene_1310037 | 24,73675058 | 4,356404967 | 1,182678584 | 3,683507107 | 0,000230047 | 0,008563503 | The Moose: suction sampler vs FISH | NA |
| gene_1312034 | 6,504089932 | 6,930612061 | 1,653707623 | 4,190953688 | 2,78E-05 | 0,002114184 | The Moose: suction sampler vs FISH | NA |
| gene_1312583 | 42,12136691 | 3,599282385 | 0,827211249 | 4,351104252 | 1,35E-05 | 0,001297052 | The Moose: suction sampler vs FISH | NA |
| gene_1312584 | 12,74170104 | 4,965965962 | 1,38474544 | 3,586194124 | 0,000335539 | 0,010917666 | The Moose: suction sampler vs FISH | NA |
| gene_1313180 | 602,6309914 | 4,213565576 | 0,934375079 | 4,50950124 | 6,50E-06 | 0,000802096 | The Moose: suction sampler vs FISH | NA |
| gene_1313238 | 5,369078655 | 6,264014294 | 1,812499929 | 3,456008022 | 0,000548239 | 0,014792584 | The Moose: suction sampler vs FISH | NA |
| gene_131414 | 12,9590966 | 4,003063588 | 1,061828334 | 3,76997247 | 0,000163266 | 0,006860135 | The Moose: suction sampler vs FISH | NA |
| gene_1315717 | 11,68584903 | 4,705921247 | 1,250229204 | 3,764046809 | 0,000167186 | 0,006976673 | The Moose: suction sampler vs FISH | NA |
| gene_131639 | 366,9343635 | 13,59068419 | 2,709356938 | 5,016202922 | 5,27E-07 | 0,000143894 | The Moose: suction sampler vs FISH | NA |
| gene_131640 | 214,1973685 | 12,80982512 | 2,96460221 | 4,320925445 | 1,55E-05 | 0,001407103 | The Moose: suction sampler vs FISH | NA |
| gene_1316409 | 70,87827144 | -2,014947554 | 0,695380568 | -2,897618435 | 0,003760077 | 0,048102924 | The Moose: suction sampler vs FISH | NA |
| gene_1317115 | 26,98348287 | 3,267766544 | 0,962796805 | 3,39403551 | 0,000688708 | 0,017144038 | The Moose: suction sampler vs FISH | NA |
| gene_1317828 | 18,79991567 | 2,846737457 | 0,906834603 | 3,139202506 | 0,001694083 | 0,030059907 | The Moose: suction sampler vs FISH | NA |
| gene_1319254 | 50,82375108 | 2,365035128 | 0,7938576 | 2,979167963 | 0,002890323 | 0,041179393 | The Moose: suction sampler vs FISH | NA |
| gene_1319594 | 8,762436187 | 7,16503767 | 1,962502601 | 3,65096977 | 0,000261252 | 0,009287344 | The Moose: suction sampler vs FISH | NA |
| gene_1319596 | 24,37612813 | 5,94896818 | 1,358648678 | 4,378591963 | 1,19E-05 | 0,001193592 | The Moose: suction sampler vs FISH | NA |
| gene_1320117 | 4,961525342 | 6,422125511 | 2,130560109 | 3,014289757 | 0,002575817 | 0,038502178 | The Moose: suction sampler vs FISH | NA |
| gene_1321817 | 13,13819144 | 3,435319567 | 1,046653489 | 3,282193777 | 0,001030028 | 0,022150412 | The Moose: suction sampler vs FISH | NA |
| gene_1322955 | 18,01387688 | 5,332543244 | 1,819406328 | 2,930924864 | 0,003379545 | 0,045104614 | The Moose: suction sampler vs FISH | NA |
| gene_1323890 | 866,6266888 | -2,281080513 | 0,530699092 | -4,298255921 | 1,72E-05 | 0,001514916 | The Moose: suction sampler vs FISH | NA |
| gene_132391 | 5,914321471 | 4,728953422 | 1,634982129 | 2,892357866 | 0,003823621 | 0,048552192 | The Moose: suction sampler vs FISH | NA |
| gene_1324161 | 18,30888248 | 3,240916935 | 1,079347267 | 3,002663771 | 0,002676279 | 0,039306464 | The Moose: suction sampler vs FISH | NA |
| gene_1325082 | 8,672155712 | 4,698889126 | 1,604693818 | 2,928215385 | 0,003409138 | 0,045344949 | The Moose: suction sampler vs FISH | NA |
| gene_1325083 | 31,07148472 | 3,636729172 | 0,988055394 | 3,680693608 | 0,0002326 | 0,008627946 | The Moose: suction sampler vs FISH | NA |
| gene_1325196 | 1027,94168 | 3,178378151 | 0,954017328 | 3,331572769 | 0,000863567 | 0,019963706 | The Moose: suction sampler vs FISH | NA |
| gene_1325398 | 5,887536904 | 6,820638409 | 1,797728359 | 3,794031714 | 0,000148221 | 0,006414892 | The Moose: suction sampler vs FISH | NA |
| gene_1325711 | 9,550973439 | 6,476093014 | 1,801388904 | 3,595055459 | 0,000324322 | 0,010695092 | The Moose: suction sampler vs FISH | NA |
| gene_132593 | 9,122334483 | 6,839510985 | 1,762226517 | 3,881175841 | 0,000103953 | 0,005111938 | The Moose: suction sampler vs FISH | NA |
| gene_1328059 | 27,08950961 | 2,987478594 | 0,950154182 | 3,144204015 | 0,001665393 | 0,02976753 | The Moose: suction sampler vs FISH | NA |
| gene_1328894 | 12,65811467 | 5,404979084 | 1,485246311 | 3,639112951 | 0,000273579 | 0,009572651 | The Moose: suction sampler vs FISH | NA |
| gene_1329268 | 3,478385094 | 6,633793178 | 2,169081226 | 3,058342444 | 0,002225651 | 0,035228466 | The Moose: suction sampler vs FISH | NA |
| gene_1329575 | 23,14034986 | 3,515815423 | 0,905056611 | 3,884635922 | 0,000102483 | 0,00506993 | The Moose: suction sampler vs FISH | NA |
| gene_1329759 | 3,063130526 | 6,087634199 | 1,988034736 | 3,062136736 | 0,00219763 | 0,035034722 | The Moose: suction sampler vs FISH | NA |
| gene_1330746 | 3,582502602 | 5,547537591 | 1,864418874 | 2,975478134 | 0,002925322 | 0,041500923 | The Moose: suction sampler vs FISH | NA |
| gene_1330992 | 17,40707781 | 3,550284013 | 1,111370911 | 3,194508673 | 0,001400691 | 0,026706306 | The Moose: suction sampler vs FISH | NA |
| gene_1333306 | 15,73896096 | 4,217805346 | 0,9975435 | 4,2281919 | 2,36E-05 | 0,001869328 | The Moose: suction sampler vs FISH | NA |
| gene_1334571 | 29,74917841 | 4,918127491 | 1,120696078 | 4,388457842 | 1,14E-05 | 0,001161387 | The Moose: suction sampler vs FISH | NA |
| gene_1334572 | 21,18787549 | 3,648694655 | 0,999710884 | 3,649749858 | 0,000262496 | 0,009327046 | The Moose: suction sampler vs FISH | NA |
| gene_1336332 | 9,460027339 | 4,73504015 | 1,294929146 | 3,656601724 | 0,000255581 | 0,009161126 | The Moose: suction sampler vs FISH | NA |
| gene_1336333 | 15,06457127 | 5,162784764 | 1,307779623 | 3,947748284 | 7,89E-05 | 0,004260333 | The Moose: suction sampler vs FISH | NA |
| gene_133702 | 28,29116219 | 3,748666747 | 0,916989741 | 4,088013835 | 4,35E-05 | 0,002871034 | The Moose: suction sampler vs FISH | NA |
| gene_1337582 | 11,70118892 | 3,454488842 | 1,082001727 | 3,192683298 | 0,001409575 | 0,026813313 | The Moose: suction sampler vs FISH | NA |
| gene_1337789 | 3,176543972 | 6,125550858 | 1,85391972 | 3,304107936 | 0,000952791 | 0,021103066 | The Moose: suction sampler vs FISH | NA |
| gene_1337965 | 10,04345141 | 4,858175954 | 1,270394281 | 3,824148162 | 0,000131225 | 0,005960309 | The Moose: suction sampler vs FISH | NA |
| gene_1338108 | 5,263578461 | 7,379925622 | 2,10566164 | 3,504801285 | 0,000456948 | 0,013218023 | The Moose: suction sampler vs FISH | NA |
| gene_1338109 | 3,744901059 | 6,975315353 | 2,286201452 | 3,05105018 | 0,002280425 | 0,035717888 | The Moose: suction sampler vs FISH | NA |
| gene_1338209 | 15,37907777 | 5,472975179 | 1,260308426 | 4,342568108 | 1,41E-05 | 0,001327731 | The Moose: suction sampler vs FISH | NA |
| gene_1338639 | 4,4052806 | 7,093911408 | 1,977118933 | 3,588004389 | 0,000333219 | 0,010881835 | The Moose: suction sampler vs FISH | NA |
| gene_1339071 | 25,62180944 | 6,35080352 | 1,338173404 | 4,745874862 | 2,08E-06 | 0,000357089 | The Moose: suction sampler vs FISH | NA |
| gene_1339072 | 21,85489767 | 5,397821916 | 1,232626675 | 4,379121454 | 1,19E-05 | 0,001192318 | The Moose: suction sampler vs FISH | NA |
| gene_1340134 | 7,88134139 | 7,530693645 | 1,692155036 | 4,450356785 | 8,57E-06 | 0,000962733 | The Moose: suction sampler vs FISH | NA |
| gene_13426 | 69,43534689 | 3,098056471 | 0,752744559 | 4,115680989 | 3,86E-05 | 0,002637447 | The Moose: suction sampler vs FISH | NA |
| gene_13427 | 71,4176007 | 2,974145074 | 0,788675124 | 3,771064895 | 0,000162552 | 0,006847895 | The Moose: suction sampler vs FISH | NA |
| gene_134272 | 8,396497631 | 3,807049074 | 1,211157046 | 3,143315796 | 0,001670455 | 0,029806031 | The Moose: suction sampler vs FISH | NA |
| gene_1343310 | 5,284014775 | 5,778164586 | 1,927429953 | 2,997859703 | 0,002718828 | 0,039635634 | The Moose: suction sampler vs FISH | NA |
| gene_1343512 | 85,23628556 | 4,776930044 | 0,923725635 | 5,171373258 | 2,32E-07 | 8,21E-05 | The Moose: suction sampler vs FISH | NA |
| gene_1343621 | 14,32090644 | 4,291255162 | 1,179377466 | 3,638576525 | 0,000274149 | 0,009588041 | The Moose: suction sampler vs FISH | NA |
| gene_1343681 | 28,52127403 | 3,180433742 | 0,801291998 | 3,96913204 | 7,21E-05 | 0,004034995 | The Moose: suction sampler vs FISH | NA |
| gene_1344355 | 8,553375472 | 4,930130601 | 1,309632747 | 3,76451384 | 0,000166873 | 0,006967606 | The Moose: suction sampler vs FISH | NA |

| | | | | | | | | |
|---|---|---|---|---|---|---|---|---|
| gene_1344407 | 56,68647177 | 2,751937721 | 0,843994405 | 3,260611332 | 0,001111723 | 0,023266414 | The Moose: suction sampler vs FISH | NA |
| gene_1345721 | 18,82938231 | 8,468169919 | 2,44793324 | 3,459314078 | 0,000541553 | 0,014654547 | The Moose: suction sampler vs FISH | NA |
| gene_1346442 | 12,21451147 | 2,963726143 | 0,99517478 | 2,978096112 | 0,00290045 | 0,041275637 | The Moose: suction sampler vs FISH | NA |
| gene_134696 | 13,75110363 | 5,604011928 | 1,351510678 | 4,146479949 | 3,38E-05 | 0,002405133 | The Moose: suction sampler vs FISH | NA |
| gene_1347321 | 24,46639192 | 5,011515559 | 1,139766302 | 4,396967652 | 1,10E-05 | 0,001129178 | The Moose: suction sampler vs FISH | NA |
| gene_1347771 | 8,082824482 | 5,16364848 | 1,689169937 | 3,05691474 | 0,002236279 | 0,035359202 | The Moose: suction sampler vs FISH | NA |
| gene_1348103 | 14,78939894 | 3,217792366 | 1,015260858 | 3,169424232 | 0,001527413 | 0,0282074 | The Moose: suction sampler vs FISH | NA |
| gene_1349172 | 16,08996402 | 4,024855368 | 1,057835483 | 3,804802764 | 0,000141917 | 0,006263889 | The Moose: suction sampler vs FISH | NA |
| gene_1349753 | 16,10688093 | 3,38634098 | 1,13083718 | 2,99454337 | 0,00274856 | 0,039863344 | The Moose: suction sampler vs FISH | NA |
| gene_1350522 | 13,90014065 | 5,84627051 | 1,310287102 | 4,461824056 | 8,13E-06 | 0,000931124 | The Moose: suction sampler vs FISH | NA |
| gene_1350525 | 12,81011798 | 3,769900207 | 1,061615618 | 3,55109716 | 0,000383629 | 0,011875719 | The Moose: suction sampler vs FISH | NA |
| gene_1350592 | 6,866705937 | 6,550595887 | 1,764889395 | 3,711618362 | 0,000205938 | 0,00797319 | The Moose: suction sampler vs FISH | NA |
| gene_1350693 | 35,70413598 | 3,044554308 | 0,920407843 | 3,307831772 | 0,000940213 | 0,020938121 | The Moose: suction sampler vs FISH | NA |
| gene_1350724 | 67,00660617 | 3,926149454 | 0,888320562 | 4,419743976 | 9,88E-06 | 0,001052955 | The Moose: suction sampler vs FISH | NA |
| gene_1350748 | 5,165183225 | 6,888414923 | 1,813314917 | 3,798796811 | 0,0001454 | 0,006360286 | The Moose: suction sampler vs FISH | NA |
| gene_1351365 | 14,18064025 | 2,720653912 | 0,924929004 | 2,941473237 | 0,003266551 | 0,044256007 | The Moose: suction sampler vs FISH | NA |
| gene_1352149 | 5,840094347 | 6,937022481 | 1,730823742 | 4,007931201 | 6,13E-05 | 0,003606207 | The Moose: suction sampler vs FISH | NA |
| gene_1353129 | 7,630771706 | 5,548842248 | 1,567023738 | 3,541007142 | 0,000398603 | 0,012187919 | The Moose: suction sampler vs FISH | NA |
| gene_1353346 | 4,265393996 | 5,764004666 | 1,820343214 | 3,166438407 | 0,00154318 | 0,028370623 | The Moose: suction sampler vs FISH | NA |
| gene_1353616 | 9,205084308 | 3,771507069 | 1,212992569 | 3,109258181 | 0,001875577 | 0,031968389 | The Moose: suction sampler vs FISH | NA |
| gene_1353617 | 12,90531285 | 3,915783016 | 1,251560464 | 3,128720608 | 0,001755692 | 0,030685572 | The Moose: suction sampler vs FISH | NA |
| gene_1354765 | 82,58005586 | 2,89638917 | 0,858830754 | 3,372479568 | 0,000744946 | 0,0180868 | The Moose: suction sampler vs FISH | NA |
| gene_1355142 | 33,61602876 | 4,487065912 | 1,242444252 | 3,611482693 | 0,000304451 | 0,010285388 | The Moose: suction sampler vs FISH | NA |
| gene_135707 | 8,889518983 | 5,524346258 | 1,640250968 | 3,367988415 | 0,000757188 | 0,018293308 | The Moose: suction sampler vs FISH | NA |
| gene_135710 | 4,098666788 | 5,729325561 | 1,865181536 | 3,071725432 | 0,002128254 | 0,034435957 | The Moose: suction sampler vs FISH | NA |
| gene_135711 | 8,210826546 | 4,018428774 | 1,250497137 | 3,213464994 | 0,001311438 | 0,025713863 | The Moose: suction sampler vs FISH | NA |
| gene_1357417 | 3,87310192 | 6,235798016 | 1,877942859 | 3,32054726 | 0,000898411 | 0,020474439 | The Moose: suction sampler vs FISH | NA |
| gene_1357418 | 8,576908866 | 6,071493424 | 1,631916995 | 3,720467059 | 0,000198855 | 0,007780971 | The Moose: suction sampler vs FISH | NA |
| gene_1357419 | 6,185934884 | 6,395722984 | 1,879142167 | 3,403533323 | 0,000665203 | 0,016754401 | The Moose: suction sampler vs FISH | NA |
| gene_1358741 | 358,2805472 | 7,143205443 | 1,656506505 | 4,312210922 | 1,62E-05 | 0,001447674 | The Moose: suction sampler vs FISH | NA |
| gene_135897 | 15,02094625 | 5,403830854 | 1,364644134 | 3,959882814 | 7,50E-05 | 0,004144008 | The Moose: suction sampler vs FISH | NA |
| gene_1359213 | 6,025522314 | 5,061071092 | 1,713880456 | 2,952989559 | 0,003147126 | 0,043353321 | The Moose: suction sampler vs FISH | NA |
| gene_1361637 | 10,06768741 | 3,836313698 | 1,207607938 | 3,176787415 | 0,001489162 | 0,027780408 | The Moose: suction sampler vs FISH | NA |
| gene_1361924 | 3,442654081 | 6,55258032 | 2,105214993 | 3,112546862 | 0,001854806 | 0,031739562 | The Moose: suction sampler vs FISH | NA |
| gene_1362331 | 123,7200379 | 3,127048347 | 0,699248913 | 4,472010307 | 7,75E-06 | 0,000900488 | The Moose: suction sampler vs FISH | NA |
| gene_1362866 | 13,86214922 | 3,545028268 | 1,02404356 | 3,461794406 | 0,000536587 | 0,014579957 | The Moose: suction sampler vs FISH | NA |
| gene_1363135 | 26,48599058 | 3,814199885 | 0,948653603 | 4,020645549 | 5,80E-05 | 0,00348257 | The Moose: suction sampler vs FISH | NA |
| gene_1363429 | 10,34185658 | 5,390454101 | 1,49671988 | 3,601511661 | 0,000316372 | 0,010561799 | The Moose: suction sampler vs FISH | NA |
| gene_136359 | 118,8761481 | 4,889862777 | 1,203016643 | 4,064667604 | 4,81E-05 | 0,003086725 | The Moose: suction sampler vs FISH | NA |
| gene_136366 | 30,87066105 | 3,116395743 | 0,859068709 | 3,627644344 | 0,000286019 | 0,00987619 | The Moose: suction sampler vs FISH | NA |
| gene_1363957 | 3,223839118 | 6,731490555 | 1,983559539 | 3,393641796 | 0,000689698 | 0,017153708 | The Moose: suction sampler vs FISH | NA |
| gene_1364550 | 16,40216785 | 3,859020642 | 1,088041766 | 3,546757819 | 0,000390003 | 0,011994877 | The Moose: suction sampler vs FISH | NA |
| gene_1365414 | 23,3308086 | 2,54168283 | 0,853499657 | 2,977954132 | 0,002901794 | 0,041278765 | The Moose: suction sampler vs FISH | NA |
| gene_1366265 | 17,90883209 | 3,234581022 | 1,017966337 | 3,177493108 | 0,001485542 | 0,02774813 | The Moose: suction sampler vs FISH | NA |
| gene_1366266 | 31,83585157 | 3,178609569 | 0,837428778 | 3,795677501 | 0,000147241 | 0,006389958 | The Moose: suction sampler vs FISH | NA |
| gene_1366508 | 3,917590215 | 5,407012806 | 1,874878092 | 2,883927669 | 0,00392749 | 0,049252677 | The Moose: suction sampler vs FISH | NA |
| gene_1366509 | 12,06528574 | 3,754189779 | 1,133794713 | 3,311172416 | 0,000929059 | 0,020815977 | The Moose: suction sampler vs FISH | NA |
| gene_1366626 | 18,80055072 | 4,134566201 | 1,068783185 | 3,868479835 | 0,000109516 | 0,005293545 | The Moose: suction sampler vs FISH | NA |
| gene_1367329 | 3,306643298 | 6,414106089 | 2,11836241 | 3,027860605 | 0,002462917 | 0,037412392 | The Moose: suction sampler vs FISH | NA |
| gene_1367467 | 4,364637197 | 5,977006045 | 1,929123473 | 3,098301446 | 0,001946333 | 0,03263058 | The Moose: suction sampler vs FISH | NA |
| gene_1368702 | 31,23901755 | 2,369795225 | 0,732191648 | 3,23657779 | 0,001209722 | 0,02444536 | The Moose: suction sampler vs FISH | NA |
| gene_1369077 | 11,96232328 | 3,290105572 | 1,081843301 | 3,041203443 | 0,002356346 | 0,03638717 | The Moose: suction sampler vs FISH | NA |
| gene_1369226 | 12,55101183 | 3,934899245 | 1,316377304 | 2,989188003 | 0,0027972 | 0,04038536 | The Moose: suction sampler vs FISH | NA |
| gene_1370020 | 13,78906947 | 6,70787164 | 1,433666149 | 4,678824038 | 2,89E-06 | 0,000439641 | The Moose: suction sampler vs FISH | NA |
| gene_137026 | 19,85786225 | 4,171762747 | 0,882360037 | 4,727959759 | 2,27E-06 | 0,000378554 | The Moose: suction sampler vs FISH | NA |
| gene_137027 | 21,54185038 | 4,265380268 | 0,882930471 | 4,830935627 | 1,36E-06 | 0,000272696 | The Moose: suction sampler vs FISH | NA |
| gene_1371054 | 247,5400975 | -2,294716336 | 0,6187877 | -3,708406511 | 0,000208568 | 0,008041076 | The Moose: suction sampler vs FISH | NA |
| gene_1371055 | 197,5864405 | -2,527625344 | 0,58638569 | -4,310516758 | 1,63E-05 | 0,001451729 | The Moose: suction sampler vs FISH | NA |
| gene_1372025 | 6,94030625 | 4,823971577 | 1,61946375 | 2,978746254 | 0,002894304 | 0,041220113 | The Moose: suction sampler vs FISH | NA |
| gene_1373759 | 46,41268002 | 3,022239795 | 0,875309197 | 3,45276824 | 0,000554865 | 0,014894772 | The Moose: suction sampler vs FISH | NA |
| gene_1374111 | 71,26644898 | 3,410520863 | 0,766632374 | 4,44870446 | 8,64E-06 | 0,000963089 | The Moose: suction sampler vs FISH | NA |
| gene_137428 | 16,23336061 | 5,078607233 | 1,116910605 | 4,547013174 | 5,44E-06 | 0,00069866 | The Moose: suction sampler vs FISH | NA |
| gene_137431 | 7,340729464 | 4,215956667 | 1,455545429 | 2,896478931 | 0,00377376 | 0,048199831 | The Moose: suction sampler vs FISH | NA |
| gene_1375790 | 10,0472107 | 3,581642171 | 1,219000838 | 2,938178597 | 0,003301468 | 0,044566386 | The Moose: suction sampler vs FISH | NA |
| gene_137652 | 29,77475307 | 3,281295074 | 0,841047391 | 3,901438978 | 9,56E-05 | 0,004864833 | The Moose: suction sampler vs FISH | NA |

| | | | | | | | | |
|---|---|---|---|---|---|---|---|---|
| gene_1377062 | 12,33180456 | 3,201550465 | 1,102278299 | 2,904484711 | 0,003678582 | 0,047473806 | The Moose: suction sampler vs FISH | NA |
| gene_1377416 | 14,72849095 | 6,071835865 | 1,440899465 | 4,21392055 | 2,51E-05 | 0,001969327 | The Moose: suction sampler vs FISH | NA |
| gene_1377427 | 90,55759668 | 5,055778865 | 1,04577376 | 4,834486253 | 1,33E-06 | 0,000271584 | The Moose: suction sampler vs FISH | NA |
| gene_1377606 | 4,389946105 | 6,505613054 | 2,224663681 | 2,924313058 | 0,003452174 | 0,045700744 | The Moose: suction sampler vs FISH | NA |
| gene_1379317 | 5,061213599 | 5,382938637 | 1,835773274 | 2,93224589 | 0,003365202 | 0,044994949 | The Moose: suction sampler vs FISH | NA |
| gene_1381927 | 23,13006498 | 3,231533 | 1,043895475 | 3,095648057 | 0,001963833 | 0,032800264 | The Moose: suction sampler vs FISH | NA |
| gene_1381928 | 15,6514459 | 3,230750743 | 0,914407973 | 3,533161168 | 0,000410622 | 0,012380188 | The Moose: suction sampler vs FISH | NA |
| gene_1384337 | 7,03350611 | 5,046364168 | 1,593614986 | 3,166614404 | 0,001542247 | 0,028370623 | The Moose: suction sampler vs FISH | NA |
| gene_1384781 | 3,976942362 | 6,694291886 | 1,859455722 | 3,60013514 | 0,000318052 | 0,010580484 | The Moose: suction sampler vs FISH | NA |
| gene_1386093 | 3,604098118 | 6,158188948 | 1,915090456 | 3,215612573 | 0,001301664 | 0,02556853 | The Moose: suction sampler vs FISH | NA |
| gene_1386418 | 156,8363902 | 4,275951557 | 0,837749085 | 5,104095766 | 3,32E-07 | 0,000104323 | The Moose: suction sampler vs FISH | NA |
| gene_1387111 | 21,17160776 | 3,635925323 | 0,918262392 | 3,959571201 | 7,51E-05 | 0,004146299 | The Moose: suction sampler vs FISH | NA |
| gene_1387241 | 21,85553891 | 3,293811554 | 1,118490099 | 2,944873233 | 0,00323087 | 0,044022733 | The Moose: suction sampler vs FISH | NA |
| gene_1388677 | 69,24985742 | 11,47499062 | 2,277619063 | 5,038151816 | 4,70E-07 | 0,00013433 | The Moose: suction sampler vs FISH | NA |
| gene_1389182 | 27,24015751 | 2,797417835 | 0,787083594 | 3,554155943 | 0,000379194 | 0,011793724 | The Moose: suction sampler vs FISH | NA |
| gene_1391088 | 59,04051277 | 3,424091844 | 0,899937094 | 3,804812432 | 0,000141912 | 0,006263889 | The Moose: suction sampler vs FISH | NA |
| gene_1391163 | 6,949753319 | 7,805180951 | 2,035074745 | 3,835328885 | 0,000125396 | 0,005797411 | The Moose: suction sampler vs FISH | NA |
| gene_1392048 | 4,117861174 | 6,597704636 | 2,027687863 | 3,253806839 | 0,001138697 | 0,023598077 | The Moose: suction sampler vs FISH | NA |
| gene_1395410 | 10,05264901 | 3,976784948 | 1,225493478 | 3,245047826 | 0,001174309 | 0,024004209 | The Moose: suction sampler vs FISH | NA |
| gene_1395882 | 13,62383451 | 4,143532514 | 1,055839116 | 3,92439762 | 8,69E-05 | 0,004554796 | The Moose: suction sampler vs FISH | NA |
| gene_1396130 | 10,88518737 | 4,882515301 | 1,425905341 | 3,42415107 | 0,000616723 | 0,015934812 | The Moose: suction sampler vs FISH | NA |
| gene_1397621 | 288,1776808 | 3,563504616 | 0,685223597 | 5,200498976 | 1,99E-07 | 7,19E-05 | The Moose: suction sampler vs FISH | NA |
| gene_1399219 | 44,56424539 | 2,726775541 | 0,934609866 | 2,917554842 | 0,003527876 | 0,046245653 | The Moose: suction sampler vs FISH | NA |
| gene_1400 | 103,6293071 | 2,617554525 | 0,777551634 | 3,366406052 | 0,000761545 | 0,018350055 | The Moose: suction sampler vs FISH | NA |
| gene_1402804 | 3,904120976 | 6,471910167 | 1,837381789 | 3,522354584 | 0,000427732 | 0,012694431 | The Moose: suction sampler vs FISH | NA |
| gene_1403681 | 46,29089012 | 2,596528242 | 0,751381236 | 3,455673521 | 0,00054892 | 0,014796594 | The Moose: suction sampler vs FISH | NA |
| gene_1404162 | 2,967106033 | 6,532793608 | 2,109888949 | 3,096273674 | 0,001959694 | 0,032755968 | The Moose: suction sampler vs FISH | NA |
| gene_1406373 | 4,567886521 | 5,939007731 | 1,968476243 | 3,017058373 | 0,002552407 | 0,038288705 | The Moose: suction sampler vs FISH | NA |
| gene_1406517 | 4,121554927 | 5,139778892 | 1,774228912 | 2,896908543 | 0,003768596 | 0,04816995 | The Moose: suction sampler vs FISH | NA |
| gene_1406529 | 11,44137107 | 5,117102895 | 1,281176608 | 3,994065193 | 6,49E-05 | 0,003740709 | The Moose: suction sampler vs FISH | NA |
| gene_1406530 | 16,90184375 | 2,897414538 | 0,928322897 | 3,121127949 | 0,001801597 | 0,031137829 | The Moose: suction sampler vs FISH | NA |
| gene_1406708 | 5,934299993 | 5,42044663 | 1,656646784 | 3,271938642 | 0,001068128 | 0,022692689 | The Moose: suction sampler vs FISH | NA |
| gene_1406996 | 4,358164693 | 6,673819355 | 2,039319211 | 3,272572198 | 0,001065736 | 0,02266812 | The Moose: suction sampler vs FISH | NA |
| gene_1407639 | 37,79314082 | 4,680613397 | 0,868139453 | 5,391545539 | 6,99E-08 | 3,75E-05 | The Moose: suction sampler vs FISH | NA |
| gene_1408751 | 4,615525034 | 5,920031278 | 1,92929404 | 3,068496121 | 0,002151391 | 0,034648127 | The Moose: suction sampler vs FISH | NA |
| gene_1409357 | 15,14352334 | 2,889465485 | 0,894685148 | 3,229589193 | 0,001239682 | 0,024777463 | The Moose: suction sampler vs FISH | NA |
| gene_1412049 | 20,24094171 | 2,956934061 | 0,993614836 | 2,9759359 | 0,002920959 | 0,041457385 | The Moose: suction sampler vs FISH | NA |
| gene_1412253 | 18,28753404 | 3,103701618 | 1,077317864 | 2,880952523 | 0,003964754 | 0,049580287 | The Moose: suction sampler vs FISH | NA |
| gene_1416562 | 9,20413229 | 7,903629583 | 1,933966774 | 4,086745278 | 4,37E-05 | 0,002879652 | The Moose: suction sampler vs FISH | NA |
| gene_1417575 | 7,125077599 | 4,180962352 | 1,359227072 | 3,075985195 | 0,002098083 | 0,03417469 | The Moose: suction sampler vs FISH | NA |
| gene_1417576 | 15,06440163 | 4,951319144 | 1,027520476 | 4,81870606 | 1,44E-06 | 0,000282993 | The Moose: suction sampler vs FISH | NA |
| gene_1417747 | 29,19170265 | 6,814902042 | 1,394028758 | 4,88863806 | 1,02E-06 | 0,000225296 | The Moose: suction sampler vs FISH | NA |
| gene_1418622 | 21,38331272 | 2,835618875 | 0,874045305 | 3,24424702 | 0,001177616 | 0,024038351 | The Moose: suction sampler vs FISH | NA |
| gene_1419163 | 74,04164875 | 3,686419273 | 0,786223801 | 4,688765805 | 2,75E-06 | 0,000428597 | The Moose: suction sampler vs FISH | NA |
| gene_1419173 | 512,3756183 | 2,417461349 | 0,648671338 | 3,726789221 | 0,000193935 | 0,007660085 | The Moose: suction sampler vs FISH | NA |
| gene_1420352 | 15,16614685 | 4,030389532 | 1,082459418 | 3,723363173 | 0,000196586 | 0,007725891 | The Moose: suction sampler vs FISH | NA |
| gene_1420479 | 18,27651574 | -3,501112937 | 1,1595177 | -3,019456224 | 0,002532289 | 0,038080253 | The Moose: suction sampler vs FISH | NA |
| gene_1420481 | 36,68409856 | -3,292099776 | 1,039427038 | -3,167225457 | 0,00153901 | 0,028350277 | The Moose: suction sampler vs FISH | NA |
| gene_1421541 | 7,241388922 | 5,115729341 | 1,589529252 | 3,218392701 | 0,001289112 | 0,025437621 | The Moose: suction sampler vs FISH | NA |
| gene_1421887 | 5,140659045 | 5,804816862 | 1,903460097 | 3,049613108 | 0,002291363 | 0,035775761 | The Moose: suction sampler vs FISH | NA |
| gene_1422641 | 9,432272478 | 7,241596973 | 1,607606148 | 4,504584024 | 6,65E-06 | 0,000815412 | The Moose: suction sampler vs FISH | NA |
| gene_1423763 | 27,30477794 | 3,422578616 | 0,925970465 | 3,696207108 | 0,000218844 | 0,008266383 | The Moose: suction sampler vs FISH | NA |
| gene_1423828 | 12,64933911 | 4,124884886 | 1,106138788 | 3,72908439 | 0,000192177 | 0,007625297 | The Moose: suction sampler vs FISH | NA |
| gene_1427203 | 20,29140618 | 3,145691608 | 0,948121046 | 3,317816454 | 0,000907241 | 0,020558413 | The Moose: suction sampler vs FISH | NA |
| gene_1427768 | 6,848249886 | 5,486678675 | 1,569895696 | 3,494931981 | 0,000474183 | 0,013477686 | The Moose: suction sampler vs FISH | NA |
| gene_1428286 | 9,788620428 | 4,672940524 | 1,348870146 | 3,464336829 | 0,00053154 | 0,014496469 | The Moose: suction sampler vs FISH | NA |
| gene_1428968 | 26,24484758 | 2,984214639 | 0,833580165 | 3,579997179 | 0,000343598 | 0,011077899 | The Moose: suction sampler vs FISH | NA |
| gene_14297 | 14,25154478 | 3,389039811 | 1,014025516 | 3,342164233 | 0,000831279 | 0,019443604 | The Moose: suction sampler vs FISH | NA |
| gene_1430025 | 5,097365247 | 5,373682233 | 1,7208145 | 3,122755087 | 0,001791668 | 0,031046275 | The Moose: suction sampler vs FISH | NA |
| gene_1430239 | 13,88915228 | 3,828851877 | 0,977440401 | 3,917222853 | 8,96E-05 | 0,004662531 | The Moose: suction sampler vs FISH | NA |
| gene_1430240 | 14,44296465 | 3,889935398 | 1,074710197 | 3,619520321 | 0,00029515 | 0,01008714 | The Moose: suction sampler vs FISH | NA |
| gene_143064 | 52,93641675 | 4,195450558 | 0,902193294 | 4,650279032 | 3,31E-06 | 0,000487896 | The Moose: suction sampler vs FISH | NA |
| gene_143066 | 6,827358431 | -4,628332756 | 1,588642695 | -2,913388122 | 0,003575299 | 0,046657401 | The Moose: suction sampler vs FISH | NA |
| gene_1430786 | 18,58268905 | 3,091690012 | 0,92301042 | 3,349572166 | 0,000809365 | 0,019064718 | The Moose: suction sampler vs FISH | NA |
| gene_1432002 | 7,560778593 | 5,56953254 | 1,59502755 | 3,491809618 | 0,00047976 | 0,013577723 | The Moose: suction sampler vs FISH | NA |

| | | | | | | | | |
|---|---|---|---|---|---|---|---|---|
| gene_1433067 | 14,42923103 | 6,649595337 | 1,426341306 | 4,661994508 | 3,13E-06 | 0,00046748 | The Moose: suction sampler vs FISH | NA |
| gene_1433726 | 398,5656227 | 5,307399434 | 1,012291409 | 5,242956116 | 1,58E-07 | 6,17E-05 | The Moose: suction sampler vs FISH | NA |
| gene_1433988 | 13,22220566 | 4,847291925 | 1,06362662 | 4,557324756 | 5,18E-06 | 0,000673475 | The Moose: suction sampler vs FISH | NA |
| gene_1434263 | 289,5319266 | 4,20003591 | 0,800514598 | 5,24666998 | 1,55E-07 | 6,10E-05 | The Moose: suction sampler vs FISH | NA |
| gene_1434734 | 27,04233784 | 2,381888842 | 0,803939634 | 2,962770763 | 0,003048835 | 0,042546401 | The Moose: suction sampler vs FISH | NA |
| gene_1438198 | 8,940844137 | 4,068714627 | 1,392389644 | 2,922109228 | 0,003476696 | 0,045883541 | The Moose: suction sampler vs FISH | NA |
| gene_143925 | 82883,63821 | 3,624744408 | 0,790939163 | 4,582835921 | 4,59E-06 | 0,000621589 | The Moose: suction sampler vs FISH | NA |
| gene_1439508 | 11,07074436 | 3,535839858 | 1,163266391 | 3,039578798 | 0,002369092 | 0,036508178 | The Moose: suction sampler vs FISH | NA |
| gene_1439525 | 196,3707353 | 2,621612748 | 0,827824571 | 3,166869937 | 0,001540892 | 0,028358408 | The Moose: suction sampler vs FISH | NA |
| gene_144 | 681,6516482 | 2,191224466 | 0,75701285 | 2,894567068 | 0,003796818 | 0,048353961 | The Moose: suction sampler vs FISH | NA |
| gene_1440553 | 155,7475393 | 2,541354374 | 0,728293736 | 3,489463452 | 0,000483991 | 0,013649185 | The Moose: suction sampler vs FISH | NA |
| gene_1441284 | 3,889818079 | 6,094747165 | 1,815805121 | 3,356498501 | 0,000789361 | 0,018755954 | The Moose: suction sampler vs FISH | NA |
| gene_1441395 | 3,09257613 | 6,4688544 | 2,046718441 | 3,160598092 | 0,001574456 | 0,028708021 | The Moose: suction sampler vs FISH | NA |
| gene_1442060 | 83,13641594 | 3,488473424 | 0,770723636 | 4,526231274 | 6,00E-06 | 0,000753845 | The Moose: suction sampler vs FISH | NA |
| gene_144383 | 7,17056733 | 5,854931616 | 1,577256535 | 3,712098498 | 0,000205548 | 0,007966475 | The Moose: suction sampler vs FISH | NA |
| gene_1445079 | 12,43325402 | 4,081775219 | 1,409908934 | 2,895063022 | 0,003790824 | 0,048311137 | The Moose: suction sampler vs FISH | NA |
| gene_1445267 | 22,92967706 | 4,195085305 | 0,922723994 | 4,54641402 | 5,46E-06 | 0,000699428 | The Moose: suction sampler vs FISH | NA |
| gene_1445268 | 51,38183825 | 3,610910133 | 0,752608707 | 4,797858568 | 1,60E-06 | 0,000305934 | The Moose: suction sampler vs FISH | NA |
| gene_1445269 | 31,67746292 | 4,358774576 | 0,990844815 | 4,39904868 | 1,09E-05 | 0,001124706 | The Moose: suction sampler vs FISH | NA |
| gene_1445270 | 20,35730748 | 4,354036692 | 0,970133067 | 4,488081936 | 7,19E-06 | 0,000861059 | The Moose: suction sampler vs FISH | NA |
| gene_1446372 | 8,489685258 | 3,98250273 | 1,265947283 | 3,145867749 | 0,001655949 | 0,029656462 | The Moose: suction sampler vs FISH | NA |
| gene_1447130 | 5,209018288 | 6,460611595 | 2,021259022 | 3,196330369 | 0,001391876 | 0,026578131 | The Moose: suction sampler vs FISH | NA |
| gene_1447446 | 52,86023457 | 3,784062938 | 0,826505735 | 4,578386786 | 4,69E-06 | 0,000631459 | The Moose: suction sampler vs FISH | NA |
| gene_1447908 | 707,8435188 | 27,09924009 | 2,918318666 | 9,285908492 | 1,60E-20 | 1,18E-15 | The Moose: suction sampler vs FISH | NA |
| gene_1448095 | 8,322978424 | 5,77661113 | 1,554934634 | 3,715018627 | 0,000203189 | 0,007904235 | The Moose: suction sampler vs FISH | NA |
| gene_145029 | 28,35851879 | 2,990372669 | 0,899894053 | 3,32302748 | 0,000890461 | 0,02035479 | The Moose: suction sampler vs FISH | NA |
| gene_1450600 | 16,41525229 | 4,564074497 | 1,138625381 | 4,008407483 | 6,11E-05 | 0,003606154 | The Moose: suction sampler vs FISH | NA |
| gene_1450752 | 13,46005259 | 6,47306364 | 1,510573932 | 4,285168375 | 1,83E-05 | 0,001563067 | The Moose: suction sampler vs FISH | NA |
| gene_1451078 | 5,739145457 | 4,996338259 | 1,681124068 | 2,972022323 | 0,002958452 | 0,041777252 | The Moose: suction sampler vs FISH | NA |
| gene_145143 | 13,5230177 | 3,635136765 | 1,053057637 | 3,45198272 | 0,000556483 | 0,01492729 | The Moose: suction sampler vs FISH | NA |
| gene_145144 | 27,23277865 | 3,053605179 | 0,777376125 | 3,928092311 | 8,56E-05 | 0,004511143 | The Moose: suction sampler vs FISH | NA |
| gene_145145 | 17,13549772 | 4,827220364 | 1,169435637 | 4,127820472 | 3,66E-05 | 0,002530279 | The Moose: suction sampler vs FISH | NA |
| gene_145193 | 7,175868212 | 6,064527224 | 1,664927197 | 3,642517964 | 0,000269984 | 0,009514868 | The Moose: suction sampler vs FISH | NA |
| gene_1452637 | 8,242793043 | 4,526012481 | 1,377823004 | 3,28490123 | 0,001020181 | 0,021998595 | The Moose: suction sampler vs FISH | NA |
| gene_145285 | 11,78569753 | 5,335144475 | 1,452108614 | 3,674067093 | 0,00023872 | 0,008792774 | The Moose: suction sampler vs FISH | NA |
| gene_145286 | 11,95503025 | 2,95398256 | 0,938050539 | 3,149065469 | 0,001637935 | 0,029448747 | The Moose: suction sampler vs FISH | NA |
| gene_1453401 | 4,904229101 | 6,500818001 | 1,804264277 | 3,603029825 | 0,000314529 | 0,010514617 | The Moose: suction sampler vs FISH | NA |
| gene_145414 | 5,872950173 | 5,542633557 | 1,864315949 | 2,973011929 | 0,00294893 | 0,041690887 | The Moose: suction sampler vs FISH | NA |
| gene_1455863 | 4,112581507 | 6,073999146 | 2,047125179 | 2,967087312 | 0,003006356 | 0,042258717 | The Moose: suction sampler vs FISH | NA |
| gene_1455909 | 53,66989479 | 4,831325348 | 1,03132055 | 4,684601068 | 2,81E-06 | 0,000434636 | The Moose: suction sampler vs FISH | NA |
| gene_1455932 | 74,10544245 | 2,826281836 | 0,772963504 | 3,656423392 | 0,000255759 | 0,009163027 | The Moose: suction sampler vs FISH | NA |
| gene_145773 | 37,65441083 | 4,007830148 | 0,953159242 | 4,204785489 | 2,61E-05 | 0,002024043 | The Moose: suction sampler vs FISH | NA |
| gene_1457911 | 5,408414836 | 6,774985628 | 1,993983616 | 3,39771379 | 0,000679515 | 0,016986711 | The Moose: suction sampler vs FISH | NA |
| gene_1458181 | 76,15248661 | 2,350808217 | 0,760316863 | 3,091879623 | 0,001988935 | 0,033071614 | The Moose: suction sampler vs FISH | NA |
| gene_1459637 | 12,36114772 | 4,680857999 | 1,370946498 | 3,414325802 | 0,000639401 | 0,016345556 | The Moose: suction sampler vs FISH | NA |
| gene_1461220 | 18,7712747 | 3,176913432 | 0,928524524 | 3,421464217 | 0,000622849 | 0,016036582 | The Moose: suction sampler vs FISH | NA |
| gene_1461575 | 18,51515864 | 4,368459783 | 1,227224837 | 3,559624651 | 0,000371385 | 0,011606974 | The Moose: suction sampler vs FISH | NA |
| gene_1462217 | 13,14653089 | 4,034139861 | 1,232399561 | 3,273402546 | 0,00106261 | 0,0226344 | The Moose: suction sampler vs FISH | NA |
| gene_1462616 | 20,59516812 | 4,02585252 | 0,958914905 | 4,19834179 | 2,69E-05 | 0,002065653 | The Moose: suction sampler vs FISH | NA |
| gene_1463020 | 35,90845851 | 4,326224471 | 0,830812205 | 5,20722306 | 1,92E-07 | 7,00E-05 | The Moose: suction sampler vs FISH | NA |
| gene_1463021 | 82,72341063 | 3,059648251 | 0,772013218 | 3,963207081 | 7,39E-05 | 0,004114566 | The Moose: suction sampler vs FISH | NA |
| gene_1464226 | 16,17346362 | 4,01987887 | 1,057711534 | 3,800543665 | 0,000144379 | 0,006334476 | The Moose: suction sampler vs FISH | NA |
| gene_1464947 | 26,80223366 | 3,829218219 | 1,05800749 | 3,61927326 | 0,000295432 | 0,010092079 | The Moose: suction sampler vs FISH | NA |
| gene_1465341 | 79,43201581 | 3,376542597 | 0,925417959 | 3,64866768 | 0,000263604 | 0,009357363 | The Moose: suction sampler vs FISH | NA |
| gene_1466914 | 8929,948659 | 2,62472185 | 0,786585361 | 3,336855707 | 0,000847319 | 0,019705941 | The Moose: suction sampler vs FISH | NA |
| gene_1467155 | 34,49880515 | 3,222634684 | 0,916832291 | 3,514966387 | 0,00043981 | 0,012913873 | The Moose: suction sampler vs FISH | NA |
| gene_1467637 | 4,958406864 | 6,644377922 | 1,877196692 | 3,539521432 | 0,000400853 | 0,01221018 | The Moose: suction sampler vs FISH | NA |
| gene_1468017 | 3,469857747 | 6,877420385 | 2,229630606 | 3,084555964 | 0,002038563 | 0,033592602 | The Moose: suction sampler vs FISH | NA |
| gene_1468870 | 8,624849878 | 4,200186592 | 1,398513452 | 3,003322268 | 0,002670495 | 0,039242595 | The Moose: suction sampler vs FISH | NA |
| gene_1468872 | 9,767354041 | 4,504400002 | 1,326374394 | 3,396024547 | 0,000683722 | 0,017045479 | The Moose: suction sampler vs FISH | NA |
| gene_1468873 | 7,483519805 | 3,990210447 | 1,382429214 | 2,886375958 | 0,003897063 | 0,049048664 | The Moose: suction sampler vs FISH | NA |
| gene_1468874 | 10,63779846 | 3,759082858 | 1,274621042 | 2,949176842 | 0,003186216 | 0,043675184 | The Moose: suction sampler vs FISH | NA |
| gene_1469380 | 24,40467976 | 3,64576185 | 0,950619192 | 3,835144378 | 0,000125491 | 0,005797411 | The Moose: suction sampler vs FISH | NA |
| gene_1469575 | 3,363103803 | 7,003336631 | 2,27147065 | 3,083172847 | 0,002048062 | 0,033673581 | The Moose: suction sampler vs FISH | NA |
| gene_1470657 | 27,94598944 | 3,631252669 | 0,846341593 | 4,290528433 | 1,78E-05 | 0,001540172 | The Moose: suction sampler vs FISH | NA |

| | | | | | | | | |
|---|---|---|---|---|---|---|---|---|
| gene_1471305 | 38,56278601 | 3,1578277 | 0,882504633 | 3,57825623 | 0,000345894 | 0,011122688 | The Moose: suction sampler vs FISH | NA |
| gene_1472403 | 11,3571206 | 3,186569252 | 1,054578004 | 3,02165344 | 0,002513982 | 0,037931428 | The Moose: suction sampler vs FISH | NA |
| gene_1472464 | 35,38206992 | 3,489110718 | 1,07697274 | 3,239739122 | 0,001196391 | 0,024295584 | The Moose: suction sampler vs FISH | NA |
| gene_1473020 | 15,3184814 | 3,455646068 | 1,035049518 | 3,338628741 | 0,00084193 | 0,019624105 | The Moose: suction sampler vs FISH | NA |
| gene_1473370 | 12,00050637 | 5,085782572 | 1,313085113 | 3,873155306 | 0,000107435 | 0,005222096 | The Moose: suction sampler vs FISH | NA |
| gene_1476133 | 9,215488404 | 7,043482766 | 1,570569185 | 4,484668891 | 7,30E-06 | 0,000870696 | The Moose: suction sampler vs FISH | NA |
| gene_1476228 | 13,27851816 | 3,61795223 | 1,15326073 | 3,137150286 | 0,001705987 | 0,0301817 | The Moose: suction sampler vs FISH | NA |
| gene_1476482 | 9,734746062 | 7,148736629 | 1,981609477 | 3,607540593 | 0,000309113 | 0,010380804 | The Moose: suction sampler vs FISH | NA |
| gene_1476548 | 3,971812945 | 6,926677813 | 2,076561213 | 3,335648268 | 0,000851008 | 0,019772937 | The Moose: suction sampler vs FISH | NA |
| gene_1476914 | 5,459646733 | 6,292408025 | 1,716627628 | 3,665563761 | 0,000246794 | 0,008964297 | The Moose: suction sampler vs FISH | NA |
| gene_1476922 | 79,03322626 | 2,143566193 | 0,668561367 | 3,206237002 | 0,001344832 | 0,02605862 | The Moose: suction sampler vs FISH | NA |
| gene_1477013 | 32,21014522 | 4,248238997 | 0,927402873 | 4,580791283 | 4,63E-06 | 0,000626541 | The Moose: suction sampler vs FISH | NA |
| gene_1477090 | 15,99963583 | 4,642617277 | 1,294265017 | 3,587068501 | 0,000334416 | 0,010901562 | The Moose: suction sampler vs FISH | NA |
| gene_1477387 | 48,45072325 | 3,84683835 | 0,776488489 | 4,954147299 | 7,26E-07 | 0,000179048 | The Moose: suction sampler vs FISH | NA |
| gene_1477388 | 148,0973821 | 3,403605616 | 0,67830878 | 5,017782043 | 5,23E-07 | 0,000143786 | The Moose: suction sampler vs FISH | NA |
| gene_1478733 | 11,25253801 | 4,050506088 | 1,350465741 | 2,999340128 | 0,002705651 | 0,039498414 | The Moose: suction sampler vs FISH | NA |
| gene_1480225 | 15,75320029 | 3,680607448 | 1,206133122 | 3,051576465 | 0,002276431 | 0,035663917 | The Moose: suction sampler vs FISH | NA |
| gene_1480845 | 24,35480265 | 5,808875275 | 1,402219786 | 4,142628233 | 3,43E-05 | 0,002431747 | The Moose: suction sampler vs FISH | NA |
| gene_1482510 | 15,6943092 | 2,94441564 | 0,949206537 | 3,101975728 | 0,001922337 | 0,032396984 | The Moose: suction sampler vs FISH | NA |
| gene_1482554 | 5,053361697 | 5,914688151 | 2,051079501 | 2,88369522 | 0,003930389 | 0,049269066 | The Moose: suction sampler vs FISH | NA |
| gene_1482737 | 10,95423801 | 3,838326446 | 1,151392794 | 3,333637717 | 0,000857182 | 0,019873068 | The Moose: suction sampler vs FISH | NA |
| gene_1482845 | 17,46205003 | 3,077549766 | 0,942913691 | 3,263872184 | 0,001099007 | 0,023114717 | The Moose: suction sampler vs FISH | NA |
| gene_1484065 | 5,95106697 | 4,154871457 | 1,439092393 | 2,887147119 | 0,003887524 | 0,048991899 | The Moose: suction sampler vs FISH | NA |
| gene_1488139 | 9,757916133 | 4,343336856 | 1,228162878 | 3,536450201 | 0,000405543 | 0,012277453 | The Moose: suction sampler vs FISH | NA |
| gene_1488454 | 14,42732468 | 3,534332017 | 1,129795881 | 3,128292531 | 0,001758251 | 0,030695208 | The Moose: suction sampler vs FISH | NA |
| gene_1488789 | 540,7928964 | 2,532190791 | 0,839243418 | 3,017230445 | 0,002550958 | 0,038274793 | The Moose: suction sampler vs FISH | NA |
| gene_1488791 | 114461,399 | 3,071667331 | 0,801145606 | 3,834093714 | 0,000126028 | 0,005810504 | The Moose: suction sampler vs FISH | NA |
| gene_1488874 | 10,60628986 | 3,384147195 | 1,090332325 | 3,1037759 | 0,00191068 | 0,032281658 | The Moose: suction sampler vs FISH | NA |
| gene_1489235 | 75,5244258 | 3,042674514 | 0,82681437 | 3,679997139 | 0,000233237 | 0,008647178 | The Moose: suction sampler vs FISH | NA |
| gene_1489967 | 8,744917106 | 8,001903184 | 1,85579063 | 4,311856659 | 1,62E-05 | 0,00144823 | The Moose: suction sampler vs FISH | NA |
| gene_1491255 | 21,01486575 | 9,697949575 | 2,674060619 | 3,626675291 | 0,000287094 | 0,00989935 | The Moose: suction sampler vs FISH | NA |
| gene_1492126 | 6,816926577 | 4,97707728 | 1,545349367 | 3,220680957 | 0,001278864 | 0,025289765 | The Moose: suction sampler vs FISH | NA |
| gene_1492216 | 5,031674973 | 5,102317259 | 1,681770993 | 3,033895388 | 0,002414181 | 0,036939488 | The Moose: suction sampler vs FISH | NA |
| gene_1493095 | 3,728191922 | 5,963187353 | 1,972943083 | 3,022483216 | 0,0025071 | 0,037874584 | The Moose: suction sampler vs FISH | NA |
| gene_1493414 | 4,488973108 | 4,853370312 | 1,626984737 | 2,983046 | 0,002853951 | 0,040907188 | The Moose: suction sampler vs FISH | NA |
| gene_1497799 | 223,3710228 | 7,959488837 | 1,666094162 | 4,777334329 | 1,78E-06 | 0,000323252 | The Moose: suction sampler vs FISH | NA |
| gene_150098 | 9,823124811 | -4,437039979 | 1,442670593 | -3,075573871 | 0,002100979 | 0,034191532 | The Moose: suction sampler vs FISH | NA |
| gene_1501080 | 3,097605297 | 6,021065476 | 2,005755427 | 3,001894146 | 0,002683055 | 0,039332723 | The Moose: suction sampler vs FISH | NA |
| gene_1502379 | 15,82828617 | 3,052518545 | 1,045628019 | 2,919315942 | 0,003508005 | 0,046082171 | The Moose: suction sampler vs FISH | NA |
| gene_1502919 | 33,58035483 | 6,225780515 | 1,186716061 | 5,246225884 | 1,55E-07 | 6,10E-05 | The Moose: suction sampler vs FISH | NA |
| gene_1502922 | 30,07917879 | 3,664340711 | 1,121526429 | 3,267279857 | 0,001085863 | 0,022949984 | The Moose: suction sampler vs FISH | NA |
| gene_1503976 | 13,10892351 | 3,541054238 | 1,137253346 | 3,113689882 | 0,001847636 | 0,031668521 | The Moose: suction sampler vs FISH | NA |
| gene_1505723 | 8,303924554 | 4,908376443 | 1,684632382 | 2,913618719 | 0,00357266 | 0,046635886 | The Moose: suction sampler vs FISH | NA |
| gene_1505753 | 7,91395415 | 4,316984743 | 1,291858575 | 3,341685248 | 0,000832714 | 0,019464895 | The Moose: suction sampler vs FISH | NA |
| gene_1505861 | 4,989904926 | 6,5727881 | 1,868519334 | 3,517645217 | 0,000435394 | 0,012842376 | The Moose: suction sampler vs FISH | NA |
| gene_1505935 | 11,74481509 | 8,75017351 | 1,761351825 | 4,967873759 | 6,77E-07 | 0,000169101 | The Moose: suction sampler vs FISH | NA |
| gene_1506341 | 80,35248088 | 2,563582605 | 0,797788341 | 3,213361834 | 0,001311909 | 0,025713863 | The Moose: suction sampler vs FISH | NA |
| gene_1506639 | 6,152351078 | 7,365173447 | 1,786650552 | 4,122335752 | 3,75E-05 | 0,002583238 | The Moose: suction sampler vs FISH | NA |
| gene_1506791 | 6,638832224 | 6,169539215 | 1,713326323 | 3,600913108 | 0,000317102 | 0,010580484 | The Moose: suction sampler vs FISH | NA |
| gene_1506838 | 6,556938333 | 6,966432768 | 1,776204769 | 3,922088765 | 8,78E-05 | 0,004582062 | The Moose: suction sampler vs FISH | NA |
| gene_1506915 | 5,05839638 | 7,04624474 | 1,874717683 | 3,758563119 | 0,000170892 | 0,007059144 | The Moose: suction sampler vs FISH | NA |
| gene_1507023 | 8,448363018 | 4,21326234 | 1,463634614 | 2,878629885 | 0,003994068 | 0,049763219 | The Moose: suction sampler vs FISH | NA |
| gene_1508804 | 10,82702217 | 4,360286399 | 1,195296613 | 3,647869785 | 0,000264424 | 0,009377397 | The Moose: suction sampler vs FISH | NA |
| gene_150911 | 23,99932032 | 6,384062448 | 1,360415739 | 4,692729042 | 2,70E-06 | 0,000427235 | The Moose: suction sampler vs FISH | NA |
| gene_150942 | 13,05514586 | 3,560720812 | 1,155789345 | 3,080769717 | 0,002064663 | 0,033844316 | The Moose: suction sampler vs FISH | NA |
| gene_1510001 | 5,022199424 | 6,687456561 | 1,785235429 | 3,745980196 | 0,000179691 | 0,007263285 | The Moose: suction sampler vs FISH | NA |
| gene_1510002 | 14,04009881 | 4,234513576 | 1,346854253 | 3,144002824 | 0,001666538 | 0,029770449 | The Moose: suction sampler vs FISH | NA |
| gene_1510175 | 28,23421504 | -3,476056644 | 1,016270309 | -3,420405588 | 0,000625278 | 0,016079681 | The Moose: suction sampler vs FISH | NA |
| gene_1511775 | 9,259920652 | 3,608414492 | 1,239964363 | 2,910095321 | 0,003613185 | 0,046951593 | The Moose: suction sampler vs FISH | NA |
| gene_1512282 | 3,807914415 | 6,118545733 | 1,973705517 | 3,100029705 | 0,001935012 | 0,032541813 | The Moose: suction sampler vs FISH | NA |
| gene_1512907 | 3,568949979 | 5,752544941 | 1,855671094 | 3,099980895 | 0,001935331 | 0,032541813 | The Moose: suction sampler vs FISH | NA |
| gene_151310 | 17,68097355 | 4,107721246 | 1,38130345 | 2,973800758 | 0,00294136 | 0,041639973 | The Moose: suction sampler vs FISH | NA |
| gene_1513820 | 20,57420426 | 6,992915157 | 2,363133475 | 2,959170623 | 0,003084683 | 0,042900233 | The Moose: suction sampler vs FISH | NA |
| gene_1513821 | 473,3183179 | 8,464273595 | 1,663297414 | 5,088851532 | 3,60E-07 | 0,000111167 | The Moose: suction sampler vs FISH | NA |
| gene_1514683 | 9,080674168 | 8,214732474 | 1,930771924 | 4,254636383 | 2,09E-05 | 0,001720199 | The Moose: suction sampler vs FISH | NA |

| | | | | | | | | |
|---|---|---|---|---|---|---|---|---|
| gene_1514794 | 21,69098123 | 4,076528887 | 0,920289195 | 4,429617243 | 9,44E-06 | 0,001024113 | The Moose: suction sampler vs FISH | NA |
| gene_1514836 | 14,87366075 | 5,797863747 | 1,271149323 | 4,561119331 | 5,09E-06 | 0,000664947 | The Moose: suction sampler vs FISH | NA |
| gene_1516169 | 33,8794425 | 2,663879329 | 0,849556502 | 3,135611726 | 0,001714961 | 0,030282421 | The Moose: suction sampler vs FISH | NA |
| gene_151830 | 11,6005733 | 3,550754946 | 1,113130254 | 3,189882705 | 0,001423305 | 0,026988521 | The Moose: suction sampler vs FISH | NA |
| gene_1518867 | 3,462974008 | 6,517264087 | 2,252085278 | 2,893879797 | 0,003805138 | 0,04839279 | The Moose: suction sampler vs FISH | NA |
| gene_1520191 | 19,94379752 | 2,877902785 | 0,985761208 | 2,919472546 | 0,003506243 | 0,046082171 | The Moose: suction sampler vs FISH | NA |
| gene_1520192 | 8,87237785 | 5,168141084 | 1,475892231 | 3,501706272 | 0,000462289 | 0,013322059 | The Moose: suction sampler vs FISH | NA |
| gene_1520634 | 42,82804031 | 3,446829748 | 0,791976478 | 4,352187023 | 1,35E-05 | 0,001295729 | The Moose: suction sampler vs FISH | NA |
| gene_1520650 | 31,52943511 | 2,923390786 | 0,858918311 | 3,403572549 | 0,000665107 | 0,016754401 | The Moose: suction sampler vs FISH | NA |
| gene_1520652 | 45,07412934 | 2,439769313 | 0,811512181 | 3,00644817 | 0,002643191 | 0,03898959 | The Moose: suction sampler vs FISH | NA |
| gene_1521010 | 7,105949017 | 7,896426903 | 1,892812353 | 4,171795947 | 3,02E-05 | 0,002224016 | The Moose: suction sampler vs FISH | NA |
| gene_1521163 | 17,75989263 | 7,018850119 | 1,444153585 | 4,860182595 | 1,17E-06 | 0,000247371 | The Moose: suction sampler vs FISH | NA |
| gene_1521831 | 11,60018146 | 4,625763503 | 1,357861674 | 3,406652968 | 0,000657647 | 0,016649733 | The Moose: suction sampler vs FISH | NA |
| gene_1521832 | 5,724504894 | 5,549258566 | 1,772084275 | 3,131486829 | 0,001739236 | 0,030537456 | The Moose: suction sampler vs FISH | NA |
| gene_1524141 | 9,476483947 | 3,360226976 | 1,162663172 | 2,890112164 | 0,003851044 | 0,048731896 | The Moose: suction sampler vs FISH | NA |
| gene_1524300 | 18,33250286 | 3,010827009 | 0,84521898 | 3,562185755 | 0,00036778 | 0,011538489 | The Moose: suction sampler vs FISH | NA |
| gene_1524343 | 30,10786652 | 4,499822357 | 1,210693099 | 3,716732474 | 0,000201816 | 0,007871985 | The Moose: suction sampler vs FISH | NA |
| gene_1525093 | 12,14673481 | 5,290565987 | 1,225041255 | 4,318683934 | 1,57E-05 | 0,001417971 | The Moose: suction sampler vs FISH | NA |
| gene_1526228 | 129,5528262 | 6,369955386 | 1,58997846 | 4,006315524 | 6,17E-05 | 0,003613435 | The Moose: suction sampler vs FISH | NA |
| gene_1526229 | 110,0281094 | 8,158523588 | 1,773108529 | 4,601254494 | 4,20E-06 | 0,000584211 | The Moose: suction sampler vs FISH | NA |
| gene_1526716 | 3,704007393 | 5,351988567 | 1,855373812 | 2,884587749 | 0,003919265 | 0,049188385 | The Moose: suction sampler vs FISH | NA |
| gene_152703 | 29,55546834 | 3,252842943 | 0,915239002 | 3,554091265 | 0,000379288 | 0,011793724 | The Moose: suction sampler vs FISH | NA |
| gene_152720 | 3,728754952 | 5,652310134 | 1,916756665 | 2,948892907 | 0,003189145 | 0,04368487 | The Moose: suction sampler vs FISH | NA |
| gene_1527363 | 8,201445901 | 7,477337265 | 1,739665143 | 4,29814743 | 1,72E-05 | 0,001514916 | The Moose: suction sampler vs FISH | NA |
| gene_1527722 | 8,743163786 | 5,688925409 | 1,628718179 | 3,492885069 | 0,000477832 | 0,013565667 | The Moose: suction sampler vs FISH | NA |
| gene_1530557 | 24,89609103 | 4,088258382 | 1,121426537 | 3,645587338 | 0,000266782 | 0,009438241 | The Moose: suction sampler vs FISH | NA |
| gene_1531035 | 123,2404342 | 2,889407282 | 0,637461296 | 4,532678772 | 5,82E-06 | 0,000736224 | The Moose: suction sampler vs FISH | NA |
| gene_1531214 | 7,981709785 | 4,523280187 | 1,332296068 | 3,395101356 | 0,000686032 | 0,017091458 | The Moose: suction sampler vs FISH | NA |
| gene_1532010 | 14,71236597 | 6,763332059 | 1,480992465 | 4,566756564 | 4,95E-06 | 0,000655485 | The Moose: suction sampler vs FISH | NA |
| gene_1532942 | 46,2594575 | 3,974215376 | 0,833331514 | 4,769068863 | 1,85E-06 | 0,000330734 | The Moose: suction sampler vs FISH | NA |
| gene_153300 | 42,83658986 | 2,850032846 | 0,877939033 | 3,246276492 | 0,001169252 | 0,023920816 | The Moose: suction sampler vs FISH | NA |
| gene_153303 | 15,77479887 | 3,391800173 | 0,969364499 | 3,498993595 | 0,000467018 | 0,013372367 | The Moose: suction sampler vs FISH | NA |
| gene_1533091 | 1305,184122 | 2,824655882 | 0,699353759 | 4,038951453 | 5,37E-05 | 0,003302606 | The Moose: suction sampler vs FISH | NA |
| gene_1533674 | 45,23852211 | 2,746894387 | 0,92203197 | 2,979174776 | 0,002890259 | 0,041179393 | The Moose: suction sampler vs FISH | NA |
| gene_1534100 | 8,48290395 | 4,725594721 | 1,578841314 | 2,993077696 | 0,002761794 | 0,040015779 | The Moose: suction sampler vs FISH | NA |
| gene_1534896 | 5,58437605 | 7,229913397 | 1,832834227 | 3,944663019 | 7,99E-05 | 0,004290319 | The Moose: suction sampler vs FISH | NA |
| gene_153540 | 4,638436362 | 6,070436215 | 1,841898281 | 3,295749976 | 0,000981593 | 0,021475461 | The Moose: suction sampler vs FISH | NA |
| gene_1535901 | 4,060424523 | 6,515821931 | 2,107121119 | 3,092286377 | 0,001986211 | 0,033048775 | The Moose: suction sampler vs FISH | NA |
| gene_1536552 | 3,103095359 | 6,790261576 | 2,175474797 | 3,121277978 | 0,00180068 | 0,031137829 | The Moose: suction sampler vs FISH | NA |
| gene_1537146 | 34,07993245 | 3,356210032 | 1,09811163 | 3,056346858 | 0,002240519 | 0,03537294 | The Moose: suction sampler vs FISH | NA |
| gene_1537383 | 5,795284098 | 4,515956189 | 1,535570079 | 2,940898792 | 0,003272614 | 0,044313637 | The Moose: suction sampler vs FISH | NA |
| gene_1538121 | 28,16698105 | 3,029305011 | 0,983351118 | 3,080593447 | 0,002065885 | 0,033852953 | The Moose: suction sampler vs FISH | NA |
| gene_1538222 | 31,85592522 | 2,745205715 | 0,907939512 | 3,023555729 | 0,00249823 | 0,037784702 | The Moose: suction sampler vs FISH | NA |
| gene_1538679 | 326,304105 | 11,43670678 | 2,163769336 | 5,285548043 | 1,25E-07 | 5,32E-05 | The Moose: suction sampler vs FISH | NA |
| gene_1538680 | 498,8107255 | 13,33265583 | 1,984654012 | 6,71787412 | 1,84E-11 | 1,23E-07 | The Moose: suction sampler vs FISH | NA |
| gene_1538861 | 38,55748653 | 2,608549702 | 0,820505632 | 3,179197801 | 0,001476833 | 0,027634645 | The Moose: suction sampler vs FISH | NA |
| gene_1538971 | 10,09143895 | 7,255447361 | 1,629313212 | 4,453070966 | 8,47E-06 | 0,000959286 | The Moose: suction sampler vs FISH | NA |
| gene_1539223 | 2,710251688 | 5,813430912 | 2,014277095 | 2,886112803 | 0,003900323 | 0,049060697 | The Moose: suction sampler vs FISH | NA |
| gene_1539264 | 26,27616469 | 3,824377973 | 1,155067355 | 3,31095668 | 0,000929776 | 0,020825678 | The Moose: suction sampler vs FISH | NA |
| gene_1539523 | 15,71129022 | 6,424776099 | 1,318744339 | 4,871889045 | 1,11E-06 | 0,000236834 | The Moose: suction sampler vs FISH | NA |
| gene_1539871 | 28,88303981 | 3,391320658 | 1,060142441 | 3,198929244 | 0,00137939 | 0,026474162 | The Moose: suction sampler vs FISH | NA |
| gene_1540138 | 34,37600504 | 2,904556636 | 0,735889393 | 3,947001633 | 7,91E-05 | 0,004267813 | The Moose: suction sampler vs FISH | NA |
| gene_1541735 | 30,36635929 | 6,699595632 | 1,258165101 | 5,324893868 | 1,01E-07 | 4,64E-05 | The Moose: suction sampler vs FISH | NA |
| gene_1541870 | 14,04966542 | 2,949923874 | 0,975204182 | 3,024929475 | 0,002486911 | 0,037690906 | The Moose: suction sampler vs FISH | NA |
| gene_1542224 | 22,66726863 | -3,899228683 | 1,325486272 | -2,94173449 | 0,003263796 | 0,04423501 | The Moose: suction sampler vs FISH | NA |
| gene_1543698 | 16,35457024 | 3,699707602 | 1,050539353 | 3,521722048 | 0,000428753 | 0,01270262 | The Moose: suction sampler vs FISH | NA |
| gene_1543882 | 6,96954964 | 7,59407164 | 2,067207142 | 3,673590076 | 0,000239166 | 0,008795979 | The Moose: suction sampler vs FISH | NA |
| gene_1544492 | 33,25851344 | 4,376583963 | 1,008751247 | 4,338615666 | 1,43E-05 | 0,001337845 | The Moose: suction sampler vs FISH | NA |
| gene_1544493 | 8,275725738 | 7,738824565 | 1,946556147 | 3,975649291 | 7,02E-05 | 0,003947106 | The Moose: suction sampler vs FISH | NA |
| gene_1544701 | 3,503000877 | 6,135902187 | 1,963373155 | 3,125183907 | 0,001776939 | 0,030868708 | The Moose: suction sampler vs FISH | NA |
| gene_1545062 | 35,0562547 | -2,371055418 | 0,812656095 | -2,917661522 | 0,003526669 | 0,046244641 | The Moose: suction sampler vs FISH | NA |
| gene_1545063 | 26,3883027 | -2,451846077 | 0,815888085 | -3,005125487 | 0,002654713 | 0,039075312 | The Moose: suction sampler vs FISH | NA |
| gene_1545328 | 39,5532612 | 6,316338778 | 1,173021476 | 5,384674455 | 7,26E-08 | 3,81E-05 | The Moose: suction sampler vs FISH | NA |
| gene_1546122 | 37,29975884 | 3,277767265 | 0,860180521 | 3,810557419 | 0,000138654 | 0,006171775 | The Moose: suction sampler vs FISH | NA |
| gene_1546894 | 4,936066098 | 4,890850855 | 1,586642733 | 3,082515524 | 0,00205259 | 0,033717846 | The Moose: suction sampler vs FISH | NA |

| | | | | | | | | |
|---|---|---|---|---|---|---|---|---|
| gene_1547320 | 10,38633005 | 4,420589032 | 1,394391242 | 3,170264485 | 0,001523002 | 0,028160723 | The Moose: suction sampler vs FISH | NA |
| gene_1547573 | 3,875302398 | 6,20560454 | 2,127377262 | 2,917021184 | 0,003533918 | 0,046284899 | The Moose: suction sampler vs FISH | NA |
| gene_1548200 | 9,572774356 | 5,247422579 | 1,380963425 | 3,799827342 | 0,000144797 | 0,006349021 | The Moose: suction sampler vs FISH | NA |
| gene_1549455 | 25,02454235 | 7,991951295 | 1,281800218 | 6,234943 | 4,52E-10 | 1,06E-06 | The Moose: suction sampler vs FISH | NA |
| gene_1549459 | 3,713220672 | 6,717273754 | 2,029827772 | 3,309282614 | 0,000935354 | 0,020887457 | The Moose: suction sampler vs FISH | NA |
| gene_1549774 | 36,759332 | 3,499523323 | 1,016494785 | 3,442736131 | 0,000575861 | 0,015257613 | The Moose: suction sampler vs FISH | NA |
| gene_1549775 | 29,94121444 | 3,388500309 | 0,95050864 | 3,564933728 | 0,000363948 | 0,011482032 | The Moose: suction sampler vs FISH | NA |
| gene_1549776 | 27,73884964 | 2,929998694 | 0,948376978 | 3,089487369 | 0,002005022 | 0,033231484 | The Moose: suction sampler vs FISH | NA |
| gene_1551322 | 4,178245738 | 6,493017165 | 1,807275314 | 3,592710592 | 0,000327256 | 0,01075887 | The Moose: suction sampler vs FISH | NA |
| gene_1553858 | 17,75564821 | 3,533901573 | 1,073866905 | 3,290818962 | 0,000998962 | 0,021726014 | The Moose: suction sampler vs FISH | NA |
| gene_1554173 | 369,3208646 | 13,6909491 | 2,293787946 | 5,968707404 | 2,39E-09 | 3,22E-06 | The Moose: suction sampler vs FISH | NA |
| gene_1555532 | 7,750336157 | 5,31581784 | 1,74805549 | 3,040989185 | 0,002358023 | 0,03638717 | The Moose: suction sampler vs FISH | NA |
| gene_1555619 | 13,87880353 | 3,75067436 | 1,149603461 | 3,262580958 | 0,001104026 | 0,023187079 | The Moose: suction sampler vs FISH | NA |
| gene_1555624 | 8,773411259 | 4,181462213 | 1,374280412 | 3,042655761 | 0,002345004 | 0,036262681 | The Moose: suction sampler vs FISH | NA |
| gene_1556221 | 8,87418653 | 5,644101779 | 1,638400974 | 3,444884291 | 0,000571304 | 0,015224751 | The Moose: suction sampler vs FISH | NA |
| gene_1556384 | 9,683161395 | 6,422942114 | 1,667206817 | 3,852516706 | 0,00011691 | 0,005535866 | The Moose: suction sampler vs FISH | NA |
| gene_1556654 | 8,226459346 | 5,244987702 | 1,471712977 | 3,563865906 | 0,000365433 | 0,011490192 | The Moose: suction sampler vs FISH | NA |
| gene_1556768 | 3,651871503 | 6,974256757 | 2,367418876 | 2,945932731 | 0,003219824 | 0,043939053 | The Moose: suction sampler vs FISH | NA |
| gene_1557225 | 37,42273723 | 3,420560656 | 0,866023834 | 3,949730389 | 7,82E-05 | 0,004249399 | The Moose: suction sampler vs FISH | NA |
| gene_1557549 | 9,666071971 | 4,043929829 | 1,048888001 | 3,855444837 | 0,000115519 | 0,00549147 | The Moose: suction sampler vs FISH | NA |
| gene_1557623 | 13,62052921 | 5,802250185 | 1,603432453 | 3,618643351 | 0,000296151 | 0,010111968 | The Moose: suction sampler vs FISH | NA |
| gene_1557625 | 39,94003137 | 3,083639125 | 0,8781824 | 3,511387979 | 0,000445773 | 0,013002311 | The Moose: suction sampler vs FISH | NA |
| gene_1558986 | 40,29745646 | 3,391577179 | 0,844900327 | 4,0141743 | 5,97E-05 | 0,003539821 | The Moose: suction sampler vs FISH | NA |
| gene_1560227 | 6,63861988 | 6,768806722 | 1,857193525 | 3,644642646 | 0,000267764 | 0,009454758 | The Moose: suction sampler vs FISH | NA |
| gene_1560574 | 24,85204805 | 3,039202105 | 0,898777949 | 3,381482722 | 0,000720958 | 0,017685616 | The Moose: suction sampler vs FISH | NA |
| gene_1560951 | 356,3481193 | 5,072233014 | 0,961710951 | 5,274176205 | 1,33E-07 | 5,47E-05 | The Moose: suction sampler vs FISH | NA |
| gene_1561356 | 21,16145345 | 3,567718802 | 1,094982507 | 3,258242739 | 0,001121045 | 0,023373898 | The Moose: suction sampler vs FISH | NA |
| gene_1562506 | 59,43263566 | 3,468762594 | 0,86464633 | 4,011770448 | 6,03E-05 | 0,003569491 | The Moose: suction sampler vs FISH | NA |
| gene_1562542 | 23,9450151 | 3,405242549 | 0,890248029 | 3,82504924 | 0,000130746 | 0,005945648 | The Moose: suction sampler vs FISH | NA |
| gene_1563139 | 6,516711633 | 5,661086405 | 1,770095632 | 3,198181106 | 0,001382974 | 0,026497228 | The Moose: suction sampler vs FISH | NA |
| gene_1563808 | 55,35599405 | -2,437448255 | 0,726800308 | -3,353669817 | 0,000797475 | 0,018893717 | The Moose: suction sampler vs FISH | NA |
| gene_1564631 | 7,651484617 | 5,230605008 | 1,585927806 | 3,298135634 | 0,000973291 | 0,02135326 | The Moose: suction sampler vs FISH | NA |
| gene_1564806 | 15,46341727 | 2,972034325 | 0,92208037 | 3,223183598 | 0,001267743 | 0,025188609 | The Moose: suction sampler vs FISH | NA |
| gene_1565789 | 6,426195709 | 6,935212581 | 1,997137973 | 3,472575593 | 0,00051549 | 0,014214983 | The Moose: suction sampler vs FISH | NA |
| gene_1565915 | 22,08937543 | 3,240649523 | 0,925228021 | 3,502541479 | 0,000460842 | 0,013294007 | The Moose: suction sampler vs FISH | NA |
| gene_1565990 | 51,68813757 | 3,567055952 | 0,916160993 | 3,893481582 | 9,88E-05 | 0,00497608 | The Moose: suction sampler vs FISH | NA |
| gene_1566282 | 47,42573454 | 2,954142044 | 0,832586011 | 3,548152388 | 0,000387944 | 0,011961597 | The Moose: suction sampler vs FISH | NA |
| gene_15666 | 151,4399072 | 4,069509099 | 0,843948389 | 4,821988111 | 1,42E-06 | 0,000279769 | The Moose: suction sampler vs FISH | NA |
| gene_1567837 | 33,03630117 | 3,685699833 | 1,00343774 | 3,673072762 | 0,000239651 | 0,008804994 | The Moose: suction sampler vs FISH | NA |
| gene_1569043 | 11,00445134 | 5,622473284 | 1,485500626 | 3,784901324 | 0,00015377 | 0,006570706 | The Moose: suction sampler vs FISH | NA |
| gene_1569113 | 51,52679515 | 3,281344874 | 0,771379674 | 4,253864841 | 2,10E-05 | 0,001724211 | The Moose: suction sampler vs FISH | NA |
| gene_1570087 | 1096,01846 | 3,174366959 | 1,015464551 | 3,126024395 | 0,001771869 | 0,03083765 | The Moose: suction sampler vs FISH | NA |
| gene_1570135 | 16,01816112 | 3,627808667 | 1,048763722 | 3,459128678 | 0,000541926 | 0,014654547 | The Moose: suction sampler vs FISH | NA |
| gene_1570292 | 6,756823936 | 4,816303042 | 1,631308162 | 2,95241767 | 0,003152961 | 0,043383106 | The Moose: suction sampler vs FISH | NA |
| gene_1570321 | 34,23676189 | 3,014202968 | 0,946251641 | 3,185413729 | 0,001445472 | 0,027256146 | The Moose: suction sampler vs FISH | NA |
| gene_1570368 | 8,542735591 | 3,794642149 | 1,224679157 | 3,09847859 | 0,00194517 | 0,03263058 | The Moose: suction sampler vs FISH | NA |
| gene_1570546 | 13,20015164 | 4,557270484 | 1,373289689 | 3,3185063 | 0,000905003 | 0,020533806 | The Moose: suction sampler vs FISH | NA |
| gene_1571220 | 63,56592293 | 3,75818278 | 0,930973217 | 4,036832329 | 5,42E-05 | 0,003307895 | The Moose: suction sampler vs FISH | NA |
| gene_1571221 | 74,18710587 | 4,185138687 | 0,838023097 | 4,994061267 | 5,91E-07 | 0,000152537 | The Moose: suction sampler vs FISH | NA |
| gene_1571405 | 8,518442436 | 4,020750574 | 1,277049023 | 3,148470029 | 0,001641276 | 0,029487153 | The Moose: suction sampler vs FISH | NA |
| gene_1571842 | 19,49991041 | 3,123695401 | 1,072890312 | 2,91147694 | 0,003597245 | 0,0468273 | The Moose: suction sampler vs FISH | NA |
| gene_1572106 | 51,09662619 | -2,142834369 | 0,702181304 | -3,05168246 | 0,002275627 | 0,035658934 | The Moose: suction sampler vs FISH | NA |
| gene_1572390 | 3,694102924 | 5,728670145 | 1,894197279 | 3,024326034 | 0,002491877 | 0,037743026 | The Moose: suction sampler vs FISH | NA |
| gene_1573460 | 5,045915797 | 6,118208172 | 1,861999264 | 3,285827386 | 0,001016833 | 0,021939278 | The Moose: suction sampler vs FISH | NA |
| gene_1573471 | 4,366670346 | 5,913809355 | 1,855383998 | 3,187377578 | 0,001435692 | 0,027140573 | The Moose: suction sampler vs FISH | NA |
| gene_1573641 | 52,27910623 | 4,029309351 | 0,674162384 | 5,976763828 | 2,28E-09 | 3,21E-06 | The Moose: suction sampler vs FISH | NA |
| gene_1574833 | 4,956250952 | 7,130767323 | 2,169201153 | 3,287278044 | 0,001011609 | 0,021865104 | The Moose: suction sampler vs FISH | NA |
| gene_1576595 | 39,15490377 | 3,551357385 | 0,893653337 | 3,973976527 | 7,07E-05 | 0,003968867 | The Moose: suction sampler vs FISH | NA |
| gene_1576659 | 65,7957958 | 2,223178591 | 0,717420884 | 3,098848446 | 0,001942744 | 0,032613671 | The Moose: suction sampler vs FISH | NA |
| gene_1578220 | 78,44435862 | 2,298348198 | 0,682710333 | 3,366505655 | 0,00076127 | 0,018350055 | The Moose: suction sampler vs FISH | NA |
| gene_1578830 | 59,22319978 | 3,934553666 | 0,822392706 | 4,784275977 | 1,72E-06 | 0,000315088 | The Moose: suction sampler vs FISH | NA |
| gene_1580047 | 18,9368033 | 4,894249296 | 1,067948808 | 4,582850095 | 4,59E-06 | 0,000621589 | The Moose: suction sampler vs FISH | NA |
| gene_158065 | 4,238280436 | 6,377938283 | 1,836481235 | 3,472912307 | 0,000514844 | 0,014209953 | The Moose: suction sampler vs FISH | NA |
| gene_1581095 | 27,75469194 | 3,602590286 | 0,952103509 | 3,783822085 | 0,000154438 | 0,006590771 | The Moose: suction sampler vs FISH | NA |
| gene_1581524 | 2,93992832 | 5,607461935 | 1,931909032 | 2,902549676 | 0,003701385 | 0,047701033 | The Moose: suction sampler vs FISH | NA |

| | | | | | | | | |
|---|---|---|---|---|---|---|---|---|
| gene_1581738 | 3,437717127 | 6,118460415 | 1,8982896 | 3,223143832 | 0,001267919 | 0,025188609 | The Moose: suction sampler vs FISH | NA |
| gene_1582571 | 74,91425368 | 4,029348678 | 0,81815091 | 4,924945542 | 8,44E-07 | 0,000199922 | The Moose: suction sampler vs FISH | NA |
| gene_1585106 | 5,129055504 | 5,800026865 | 1,828190687 | 3,172550274 | 0,001511064 | 0,028032354 | The Moose: suction sampler vs FISH | NA |
| gene_1585963 | 11,65232743 | 3,873190671 | 1,340090532 | 2,89024553 | 0,00384941 | 0,048731896 | The Moose: suction sampler vs FISH | NA |
| gene_1586697 | 7,802989709 | 7,196703731 | 1,889598721 | 3,808588379 | 0,000139762 | 0,006206076 | The Moose: suction sampler vs FISH | NA |
| gene_1587580 | 3,915442445 | 6,21475824 | 1,782426357 | 3,486684437 | 0,000489048 | 0,013761737 | The Moose: suction sampler vs FISH | NA |
| gene_158779 | 21,83746124 | 3,572983988 | 0,840438238 | 4,251334394 | 2,13E-05 | 0,001736052 | The Moose: suction sampler vs FISH | NA |
| gene_158780 | 20,62815705 | 4,625772117 | 1,235946851 | 3,742695014 | 0,000182057 | 0,007327963 | The Moose: suction sampler vs FISH | NA |
| gene_1588680 | 8,635972643 | 6,909277494 | 1,636346883 | 4,222379475 | 2,42E-05 | 0,001911125 | The Moose: suction sampler vs FISH | NA |
| gene_1589004 | 14,05698632 | 3,773689837 | 1,084437109 | 3,47986048 | 0,000501675 | 0,014004378 | The Moose: suction sampler vs FISH | NA |
| gene_1589201 | 33,48524909 | 3,137204521 | 0,899068879 | 3,489392854 | 0,000484119 | 0,013649185 | The Moose: suction sampler vs FISH | NA |
| gene_1589421 | 6,761986774 | 5,430397362 | 1,512062003 | 3,591385375 | 0,000328925 | 0,010797679 | The Moose: suction sampler vs FISH | NA |
| gene_1589693 | 122,5854802 | 3,610296634 | 0,775953846 | 4,652720846 | 3,28E-06 | 0,000484092 | The Moose: suction sampler vs FISH | NA |
| gene_1590429 | 4,167152069 | 5,891697739 | 1,782357989 | 3,305563627 | 0,000947856 | 0,02105725 | The Moose: suction sampler vs FISH | NA |
| gene_1591151 | 18,2740541 | 4,733549745 | 1,14672225 | 4,127895614 | 3,66E-05 | 0,002530279 | The Moose: suction sampler vs FISH | NA |
| gene_1591732 | 77,39126664 | 2,908038542 | 0,814511243 | 3,570286558 | 0,000356591 | 0,011332678 | The Moose: suction sampler vs FISH | NA |
| gene_1591820 | 20,6242568 | 2,798531308 | 0,89067679 | 3,142027884 | 0,001677821 | 0,029902337 | The Moose: suction sampler vs FISH | NA |
| gene_1591821 | 23,41351394 | 3,321748137 | 1,002537144 | 3,313341712 | 0,000921883 | 0,020705711 | The Moose: suction sampler vs FISH | NA |
| gene_1592925 | 4,172777025 | 6,305880167 | 1,811251455 | 3,481504542 | 0,000498605 | 0,013950502 | The Moose: suction sampler vs FISH | NA |
| gene_1593712 | 10,92214339 | 3,57461284 | 1,200720594 | 2,977056325 | 0,002910305 | 0,041343787 | The Moose: suction sampler vs FISH | NA |
| gene_1594163 | 87,47405777 | 3,291829416 | 0,718816463 | 4,57951311 | 4,66E-06 | 0,000629223 | The Moose: suction sampler vs FISH | NA |
| gene_1594164 | 10,70185098 | 3,021934785 | 1,00864761 | 2,996026317 | 0,002735228 | 0,039760916 | The Moose: suction sampler vs FISH | NA |
| gene_159419 | 39,33694727 | 5,904279621 | 1,216380842 | 4,853972884 | 1,21E-06 | 0,000251778 | The Moose: suction sampler vs FISH | NA |
| gene_1594238 | 10,05775979 | 4,787329738 | 1,421035416 | 3,368902481 | 0,000754681 | 0,018252285 | The Moose: suction sampler vs FISH | NA |
| gene_1594850 | 5,559367287 | 5,87560058 | 1,75708616 | 3,343945626 | 0,000825959 | 0,019337767 | The Moose: suction sampler vs FISH | NA |
| gene_1595496 | 5,206779674 | 7,301166513 | 2,153603829 | 3,390208734 | 0,000698394 | 0,017264748 | The Moose: suction sampler vs FISH | NA |
| gene_1595498 | 3,256915639 | 6,521206363 | 2,254934103 | 2,891972034 | 0,00382832 | 0,048578262 | The Moose: suction sampler vs FISH | NA |
| gene_1596461 | 5,930787197 | 4,655576299 | 1,606214239 | 2,898477792 | 0,003749789 | 0,0480412 | The Moose: suction sampler vs FISH | NA |
| gene_1596947 | 5,74894683 | 4,551963051 | 1,393880073 | 3,265677685 | 0,001092025 | 0,023035748 | The Moose: suction sampler vs FISH | NA |
| gene_1597022 | 9,440795049 | 4,80996799 | 1,39367617 | 3,451280931 | 0,000557932 | 0,014960241 | The Moose: suction sampler vs FISH | NA |
| gene_1597393 | 6,032862183 | 6,067970444 | 1,66201155 | 3,650979709 | 0,000261242 | 0,009287344 | The Moose: suction sampler vs FISH | NA |
| gene_1597394 | 4,247192752 | 6,132635595 | 1,964063845 | 3,122421713 | 0,001793698 | 0,031050919 | The Moose: suction sampler vs FISH | NA |
| gene_159877 | 57,80250115 | 2,434003904 | 0,792616979 | 3,07084502 | 0,002134539 | 0,034477947 | The Moose: suction sampler vs FISH | NA |
| gene_1600047 | 5,669026963 | 4,964537507 | 1,679857962 | 2,955331713 | 0,00312333 | 0,043200728 | The Moose: suction sampler vs FISH | NA |
| gene_16004 | 9,034390482 | 3,479203438 | 1,193767582 | 2,914473045 | 0,003562896 | 0,046561722 | The Moose: suction sampler vs FISH | NA |
| gene_1600490 | 13,04155835 | 6,260951542 | 1,558770186 | 4,016596928 | 5,90E-05 | 0,00351991 | The Moose: suction sampler vs FISH | NA |
| gene_1600789 | 12,498081 | 4,026645633 | 1,074735204 | 3,746639747 | 0,000179219 | 0,007252205 | The Moose: suction sampler vs FISH | NA |
| gene_1602027 | 199,1107332 | 6,414059327 | 1,771195395 | 3,621316624 | 0,000293108 | 0,010031352 | The Moose: suction sampler vs FISH | NA |
| gene_1602237 | 5,281064989 | 6,115260833 | 1,855397933 | 3,295929527 | 0,000980966 | 0,021469493 | The Moose: suction sampler vs FISH | NA |
| gene_1603186 | 16,197102 | 3,148752881 | 1,005846423 | 3,130450939 | 0,001745382 | 0,030623398 | The Moose: suction sampler vs FISH | NA |
| gene_1604065 | 12,8358128 | 4,056096956 | 1,212804468 | 3,3443948 | 0,000824623 | 0,019324963 | The Moose: suction sampler vs FISH | NA |
| gene_1604437 | 4,544874784 | 6,976331148 | 1,931133897 | 3,612556933 | 0,000303193 | 0,010271678 | The Moose: suction sampler vs FISH | NA |
| gene_1604635 | 12,77440848 | 5,117147548 | 1,190997225 | 4,296523486 | 1,73E-05 | 0,001522405 | The Moose: suction sampler vs FISH | NA |
| gene_1604784 | 7,178624066 | -4,772505614 | 1,609294932 | -2,965587922 | 0,00302105 | 0,042362843 | The Moose: suction sampler vs FISH | NA |
| gene_1605153 | 27,54465914 | 5,256031707 | 1,242171162 | 4,231326463 | 2,32E-05 | 0,00185415 | The Moose: suction sampler vs FISH | NA |
| gene_1605374 | 8,963746043 | 4,590671376 | 1,48872575 | 3,08362462 | 0,002044954 | 0,033671466 | The Moose: suction sampler vs FISH | NA |
| gene_1606251 | 8,636510895 | 3,976330991 | 1,293971342 | 3,072966813 | 0,002119421 | 0,034339476 | The Moose: suction sampler vs FISH | NA |
| gene_1609436 | 135,5111217 | 2,74492462 | 0,799685589 | 3,432504795 | 0,000598033 | 0,015608584 | The Moose: suction sampler vs FISH | NA |
| gene_1609778 | 10,61395022 | 3,615534189 | 1,181782005 | 3,059391811 | 0,002217869 | 0,03520092 | The Moose: suction sampler vs FISH | NA |
| gene_1612086 | 22,81298905 | 3,626296569 | 0,973058839 | 3,726698144 | 0,000194005 | 0,007660085 | The Moose: suction sampler vs FISH | NA |
| gene_1612087 | 30,1859901 | 3,677153115 | 1,045638843 | 3,516656961 | 0,000437018 | 0,012867033 | The Moose: suction sampler vs FISH | NA |
| gene_1612647 | 19,75378648 | 4,611637982 | 1,184714943 | 3,892614009 | 9,92E-05 | 0,00497849 | The Moose: suction sampler vs FISH | NA |
| gene_1614020 | 7,247546698 | 4,452599371 | 1,302405515 | 3,418750398 | 0,000629094 | 0,016148603 | The Moose: suction sampler vs FISH | NA |
| gene_1614768 | 6,476953922 | 6,798120185 | 1,718140512 | 3,956673007 | 7,60E-05 | 0,004178056 | The Moose: suction sampler vs FISH | NA |
| gene_1615069 | 69,38522171 | 3,543242423 | 0,995525027 | 3,55916961 | 0,000372029 | 0,011615767 | The Moose: suction sampler vs FISH | NA |
| gene_1619499 | 3,528704705 | 5,622240563 | 1,824860636 | 3,08091503 | 0,002063655 | 0,033844316 | The Moose: suction sampler vs FISH | NA |
| gene_1619508 | 18,94283391 | 3,839415032 | 0,953825344 | 4,025280999 | 5,69E-05 | 0,003434312 | The Moose: suction sampler vs FISH | NA |
| gene_1620239 | 3,926248378 | 6,810752521 | 2,080407262 | 3,273759252 | 0,00106127 | 0,022622371 | The Moose: suction sampler vs FISH | NA |
| gene_1621099 | 61,36371658 | 4,353292776 | 0,868608054 | 5,011803372 | 5,39E-07 | 0,000145124 | The Moose: suction sampler vs FISH | NA |
| gene_1621377 | 16,91589485 | 3,562292339 | 0,941650808 | 3,783029027 | 0,000154931 | 0,006604139 | The Moose: suction sampler vs FISH | NA |
| gene_1621379 | 10,87625439 | 4,064208631 | 1,219384348 | 3,333000492 | 0,000859148 | 0,01988658 | The Moose: suction sampler vs FISH | NA |
| gene_1623209 | 6,592718713 | 4,125505748 | 1,398002923 | 2,950999372 | 0,003167476 | 0,043488543 | The Moose: suction sampler vs FISH | NA |
| gene_1624657 | 11,8169319 | 3,133161075 | 1,064877285 | 2,942274307 | 0,003258112 | 0,044186999 | The Moose: suction sampler vs FISH | NA |
| gene_1624776 | 12,18979721 | 3,964776308 | 1,256358496 | 3,155768293 | 0,00160076 | 0,029029085 | The Moose: suction sampler vs FISH | NA |
| gene_1624839 | 15,63241594 | 5,231603968 | 1,21339396 | 4,311546077 | 1,62E-05 | 0,001448501 | The Moose: suction sampler vs FISH | NA |

| | | | | | | | | |
|---|---|---|---|---|---|---|---|---|
| gene_162819 | 77,64260922 | 3,683373896 | 0,744211395 | 4,949365088 | 7,45E-07 | 0,00018289 | The Moose: suction sampler vs FISH | NA |
| gene_1628502 | 11,13199226 | 4,434090845 | 1,193642218 | 3,71475705 | 0,000203399 | 0,007908227 | The Moose: suction sampler vs FISH | NA |
| gene_1629380 | 28,5820181 | 3,227140954 | 0,887574628 | 3,635909423 | 0,000277002 | 0,009673983 | The Moose: suction sampler vs FISH | NA |
| gene_1632414 | 13,17824922 | 3,429205923 | 1,110015603 | 3,089331279 | 0,002006076 | 0,033231484 | The Moose: suction sampler vs FISH | NA |
| gene_1632767 | 9,975718477 | -4,57087359 | 1,551304366 | -2,946471169 | 0,003214224 | 0,043916724 | The Moose: suction sampler vs FISH | NA |
| gene_1633253 | 14,49852674 | 9,121234327 | 3,034865264 | 3,005482463 | 0,002651599 | 0,039050868 | The Moose: suction sampler vs FISH | NA |
| gene_1634068 | 7,309421743 | 4,689254708 | 1,492347197 | 3,142200902 | 0,001676829 | 0,029891925 | The Moose: suction sampler vs FISH | NA |
| gene_1634069 | 28,39341406 | 3,173641038 | 0,821925616 | 3,861226585 | 0,000112819 | 0,005405093 | The Moose: suction sampler vs FISH | NA |
| gene_163425 | 6,183944286 | 7,105534631 | 1,771748394 | 4,010464836 | 6,06E-05 | 0,003583508 | The Moose: suction sampler vs FISH | NA |
| gene_1634880 | 14,99551731 | 3,235943517 | 1,123403361 | 2,880482319 | 0,003970672 | 0,049588781 | The Moose: suction sampler vs FISH | NA |
| gene_1635093 | 3,618637151 | 5,981925598 | 1,968114982 | 3,039418759 | 0,002370351 | 0,036509811 | The Moose: suction sampler vs FISH | NA |
| gene_1635358 | 131,8573194 | 9,866276877 | 2,22206505 | 4,440138634 | 8,99E-06 | 0,000987025 | The Moose: suction sampler vs FISH | NA |
| gene_1635360 | 133,5842104 | 10,54628602 | 2,522661525 | 4,180618729 | 2,91E-05 | 0,002172096 | The Moose: suction sampler vs FISH | NA |
| gene_1635409 | 25,88172818 | 3,01970432 | 0,834541328 | 3,618399975 | 0,00029643 | 0,010116777 | The Moose: suction sampler vs FISH | NA |
| gene_1635597 | 5,204515049 | 6,597312841 | 1,937892164 | 3,404375622 | 0,000663155 | 0,016737807 | The Moose: suction sampler vs FISH | NA |
| gene_1635920 | 26,59798106 | 4,248516131 | 1,081832441 | 3,927148023 | 8,60E-05 | 0,004520338 | The Moose: suction sampler vs FISH | NA |
| gene_1636746 | 5,283185604 | 5,103170602 | 1,725381503 | 2,95770564 | 0,003099379 | 0,043012037 | The Moose: suction sampler vs FISH | NA |
| gene_1637143 | 12,61621173 | -5,03645736 | 1,721754133 | -2,925189644 | 0,003442464 | 0,045647426 | The Moose: suction sampler vs FISH | NA |
| gene_1638544 | 57,75691064 | -5,232808708 | 1,228704752 | -4,258800742 | 2,06E-05 | 0,001696795 | The Moose: suction sampler vs FISH | NA |
| gene_163913 | 21,44269154 | 9,605750197 | 1,620772354 | 5,926649832 | 3,09E-09 | 3,85E-06 | The Moose: suction sampler vs FISH | NA |
| gene_1639138 | 12,65117862 | 7,685405691 | 1,798511705 | 4,273203042 | 1,93E-05 | 0,001612963 | The Moose: suction sampler vs FISH | NA |
| gene_1639761 | 22,21688208 | 3,469098956 | 0,893410659 | 3,882983621 | 0,000103183 | 0,005092903 | The Moose: suction sampler vs FISH | NA |
| gene_1640671 | 178,9396581 | 12,62071303 | 3,049491473 | 4,138628732 | 3,49E-05 | 0,002454294 | The Moose: suction sampler vs FISH | NA |
| gene_1642997 | 11,96803235 | 4,098252831 | 1,339915873 | 3,058589657 | 0,002223815 | 0,035225748 | The Moose: suction sampler vs FISH | NA |
| gene_1644272 | 9,57886384 | 5,147335606 | 1,478813225 | 3,480720566 | 0,000500067 | 0,013970108 | The Moose: suction sampler vs FISH | NA |
| gene_1644545 | 4,822093211 | 6,438771969 | 1,832562198 | 3,513535298 | 0,000442186 | 0,012950027 | The Moose: suction sampler vs FISH | NA |
| gene_1645051 | 6,272246565 | 5,347562182 | 1,629334084 | 3,282053836 | 0,001030539 | 0,022150412 | The Moose: suction sampler vs FISH | NA |
| gene_1645639 | 10,68498068 | 6,631415128 | 1,464378539 | 4,528484235 | 5,94E-06 | 0,000747602 | The Moose: suction sampler vs FISH | NA |
| gene_1645720 | 31,44153881 | 3,184751636 | 0,937310845 | 3,397753962 | 0,000679415 | 0,016986711 | The Moose: suction sampler vs FISH | NA |
| gene_1646615 | 3,61179757 | 5,704166113 | 1,981563236 | 2,878619268 | 0,003994202 | 0,049763219 | The Moose: suction sampler vs FISH | NA |
| gene_1647234 | 5,013398414 | 5,089511967 | 1,764673568 | 2,884109593 | 0,003925221 | 0,049237895 | The Moose: suction sampler vs FISH | NA |
| gene_1647549 | 7,719534395 | 3,67564147 | 1,244729576 | 2,952963875 | 0,003147388 | 0,043353321 | The Moose: suction sampler vs FISH | NA |
| gene_1647551 | 15,79754068 | 4,493978099 | 1,204945957 | 3,729609676 | 0,000191777 | 0,007617649 | The Moose: suction sampler vs FISH | NA |
| gene_1647552 | 5,80354583 | 6,809412946 | 1,693327356 | 4,021321053 | 5,79E-05 | 0,003478546 | The Moose: suction sampler vs FISH | NA |
| gene_1647737 | 39,87473791 | 4,894535505 | 0,907890073 | 5,391110279 | 7,00E-08 | 3,75E-05 | The Moose: suction sampler vs FISH | NA |
| gene_1647738 | 98,16579266 | 3,768329329 | 0,861488115 | 4,374209305 | 1,22E-05 | 0,001206333 | The Moose: suction sampler vs FISH | NA |
| gene_1648448 | 30,32956415 | 4,446190916 | 1,027093474 | 4,32890582 | 1,50E-05 | 0,001377457 | The Moose: suction sampler vs FISH | NA |
| gene_16491 | 74,61630667 | 2,329616913 | 0,71439763 | 3,260952748 | 0,001110385 | 0,023260769 | The Moose: suction sampler vs FISH | NA |
| gene_1649916 | 7,780699833 | 4,518283818 | 1,523598697 | 2,965534053 | 0,003021579 | 0,042362843 | The Moose: suction sampler vs FISH | NA |
| gene_1650720 | 6,292240976 | 4,797290289 | 1,562630756 | 3,070008875 | 0,002140524 | 0,034529056 | The Moose: suction sampler vs FISH | NA |
| gene_165127 | 4,600561557 | 5,759857808 | 1,987036984 | 2,898716961 | 0,00374693 | 0,0480412 | The Moose: suction sampler vs FISH | NA |
| gene_165128 | 6,84440194 | 6,565716324 | 1,918562678 | 3,422205801 | 0,000621153 | 0,016004302 | The Moose: suction sampler vs FISH | NA |
| gene_165130 | 212,4933866 | 13,09992615 | 4,247188581 | 3,084375912 | 0,002039797 | 0,033605401 | The Moose: suction sampler vs FISH | NA |
| gene_1651390 | 63,91959636 | 2,768782758 | 0,866832599 | 3,194137786 | 0,001402492 | 0,026727036 | The Moose: suction sampler vs FISH | NA |
| gene_1652363 | 144,6127215 | 2,661270466 | 0,890035851 | 2,990071088 | 0,002789125 | 0,040308403 | The Moose: suction sampler vs FISH | NA |
| gene_1652824 | 102,744519 | 2,337561861 | 0,807449358 | 2,894995009 | 0,003791646 | 0,048313222 | The Moose: suction sampler vs FISH | NA |
| gene_1652938 | 36,19983874 | 2,413694662 | 0,796759174 | 3,029390488 | 0,002450477 | 0,037269684 | The Moose: suction sampler vs FISH | NA |
| gene_1652939 | 9,333035215 | 4,02526152 | 1,294842863 | 3,108687266 | 0,001879205 | 0,032001019 | The Moose: suction sampler vs FISH | NA |
| gene_1653463 | 3,658525279 | 6,967107903 | 2,109508194 | 3,302716682 | 0,000957531 | 0,021156994 | The Moose: suction sampler vs FISH | NA |
| gene_1655813 | 20,68343504 | 2,83503638 | 0,932234381 | 3,041119742 | 0,002357001 | 0,03638717 | The Moose: suction sampler vs FISH | NA |
| gene_1656106 | 482,3628161 | 2,161211972 | 0,715585316 | 3,020201679 | 0,002526064 | 0,037994426 | The Moose: suction sampler vs FISH | NA |
| gene_1656988 | 274,3421087 | 12,52709147 | 2,991573389 | 4,187459188 | 2,82E-05 | 0,002138127 | The Moose: suction sampler vs FISH | NA |
| gene_1657111 | 14,04833845 | 9,15428329 | 1,911532958 | 4,788974865 | 1,68E-06 | 0,000310126 | The Moose: suction sampler vs FISH | NA |
| gene_165822 | 52,54976929 | 5,609386333 | 1,146592145 | 4,892224633 | 9,97E-07 | 0,000221899 | The Moose: suction sampler vs FISH | NA |
| gene_165823 | 24,1389729 | 5,942370652 | 1,387826223 | 4,281782944 | 1,85E-05 | 0,001574414 | The Moose: suction sampler vs FISH | NA |
| gene_1658541 | 3,055316295 | 6,418000111 | 2,179414842 | 2,944827202 | 0,003231351 | 0,044022733 | The Moose: suction sampler vs FISH | NA |
| gene_165856 | 12,31350171 | 3,655781944 | 1,155018642 | 3,165128087 | 0,001550147 | 0,028434201 | The Moose: suction sampler vs FISH | NA |
| gene_1658787 | 6,124504047 | 5,07754951 | 1,600192954 | 3,173085782 | 0,00150828 | 0,028016083 | The Moose: suction sampler vs FISH | NA |
| gene_1659149 | 4,571802716 | 5,270275339 | 1,793833609 | 2,937995649 | 0,003303417 | 0,044574579 | The Moose: suction sampler vs FISH | NA |
| gene_1659566 | 10,26121119 | 4,314824153 | 1,202459461 | 3,588332324 | 0,0003328 | 0,010872994 | The Moose: suction sampler vs FISH | NA |
| gene_1661774 | 70,60507372 | 2,392079787 | 0,793108339 | 3,016082002 | 0,00256064 | 0,038357374 | The Moose: suction sampler vs FISH | NA |
| gene_1661910 | 3,127753654 | 6,08231736 | 2,011882239 | 3,023197502 | 0,002501189 | 0,037821672 | The Moose: suction sampler vs FISH | NA |
| gene_1662202 | 7,423484144 | 4,050164099 | 1,397095982 | 2,898987722 | 0,003743696 | 0,048020094 | The Moose: suction sampler vs FISH | NA |
| gene_1663138 | 41,03140436 | 2,436484052 | 0,790877293 | 3,08073588 | 0,002064897 | 0,033844316 | The Moose: suction sampler vs FISH | NA |
| gene_1664120 | 60,3316592 | 2,518405937 | 0,6564511 | 3,836395337 | 0,000124853 | 0,005797411 | The Moose: suction sampler vs FISH | NA |

| | | | | | | | | |
|---|---|---|---|---|---|---|---|---|
| gene_1664135 | 7,402176631 | 5,414050829 | 1,478304594 | 3,662337824 | 0,000249924 | 0,0090333 | The Moose: suction sampler vs FISH | NA |
| gene_1664136 | 10,70330593 | 3,479288946 | 1,18147676 | 2,944864482 | 0,003230961 | 0,044022733 | The Moose: suction sampler vs FISH | NA |
| gene_1665427 | 4,422187626 | 6,900470221 | 1,924208042 | 3,586135214 | 0,000335615 | 0,010917666 | The Moose: suction sampler vs FISH | NA |
| gene_1666736 | 21,53192521 | 2,989393312 | 0,980186445 | 3,049821111 | 0,002289777 | 0,035759522 | The Moose: suction sampler vs FISH | NA |
| gene_1666840 | 16,06435582 | 3,786022394 | 1,212787411 | 3,121752716 | 0,001797779 | 0,031111654 | The Moose: suction sampler vs FISH | NA |
| gene_1668402 | 8,394924951 | 6,286631101 | 1,606899008 | 3,912275178 | 9,14E-05 | 0,004725632 | The Moose: suction sampler vs FISH | NA |
| gene_1668467 | 22,07653604 | 3,244782192 | 0,862765078 | 3,760910442 | 0,000169296 | 0,007025858 | The Moose: suction sampler vs FISH | NA |
| gene_1668468 | 25,66709429 | 3,42827347 | 1,071302197 | 3,200099355 | 0,001373802 | 0,026406413 | The Moose: suction sampler vs FISH | NA |
| gene_166871 | 27,85857517 | 3,496010235 | 0,899775325 | 3,885425771 | 0,000102151 | 0,005069338 | The Moose: suction sampler vs FISH | NA |
| gene_1668939 | 22,97244861 | 3,480506518 | 1,167895137 | 2,980153277 | 0,002881042 | 0,041108821 | The Moose: suction sampler vs FISH | NA |
| gene_1669623 | 56,41554297 | 5,832673888 | 0,977420294 | 5,967416397 | 2,41E-09 | 3,22E-06 | The Moose: suction sampler vs FISH | NA |
| gene_1670003 | 23,25220363 | 3,443157448 | 1,082720457 | 3,180098266 | 0,001472251 | 0,027593959 | The Moose: suction sampler vs FISH | NA |
| gene_16703 | 41,8732868 | 5,224887591 | 1,099639717 | 4,751454056 | 2,02E-06 | 0,000350538 | The Moose: suction sampler vs FISH | NA |
| gene_1670896 | 22,86285752 | 4,699434863 | 1,029412402 | 4,565162472 | 4,99E-06 | 0,000658941 | The Moose: suction sampler vs FISH | NA |
| gene_1672782 | 8,443777584 | 5,071569083 | 1,594263323 | 3,181136396 | 0,001466985 | 0,027541602 | The Moose: suction sampler vs FISH | NA |
| gene_1672969 | 9,066210193 | 4,47479003 | 1,286483059 | 3,478312441 | 0,000504581 | 0,014044726 | The Moose: suction sampler vs FISH | NA |
| gene_1673244 | 65,67286687 | 3,074863954 | 0,988028199 | 3,112121656 | 0,001857479 | 0,031751858 | The Moose: suction sampler vs FISH | NA |
| gene_1674795 | 85,60315985 | 2,507617294 | 0,736338515 | 3,405522381 | 0,000660376 | 0,016699904 | The Moose: suction sampler vs FISH | NA |
| gene_1674796 | 67,47321112 | 2,424738589 | 0,758613877 | 3,196275025 | 0,001392143 | 0,026578131 | The Moose: suction sampler vs FISH | NA |
| gene_1674797 | 38,44624756 | 2,394282469 | 0,729561505 | 3,28181031 | 0,00103143 | 0,022163064 | The Moose: suction sampler vs FISH | NA |
| gene_167601 | 12,47754372 | 3,297356335 | 1,104921536 | 2,984244788 | 0,002842792 | 0,040818942 | The Moose: suction sampler vs FISH | NA |
| gene_1676632 | 17,44512384 | 2,874794892 | 0,87569586 | 3,282869114 | 0,001027564 | 0,022125304 | The Moose: suction sampler vs FISH | NA |
| gene_1676725 | 6,25875897 | 4,968183237 | 1,550875082 | 3,203470926 | 0,001357817 | 0,026205737 | The Moose: suction sampler vs FISH | NA |
| gene_1676797 | 15,56152411 | 3,515905451 | 1,11401691 | 3,156061114 | 0,001599154 | 0,02900712 | The Moose: suction sampler vs FISH | NA |
| gene_1677177 | 93,74576154 | 2,811571166 | 0,755240674 | 3,722748604 | 0,000197066 | 0,007730949 | The Moose: suction sampler vs FISH | NA |
| gene_1677178 | 101,1033435 | 2,803479132 | 0,865469536 | 3,239258015 | 0,001198411 | 0,024320891 | The Moose: suction sampler vs FISH | NA |
| gene_167836 | 9,510256341 | 6,02529411 | 1,617268012 | 3,725600249 | 0,000194851 | 0,007685731 | The Moose: suction sampler vs FISH | NA |
| gene_1679294 | 288,968467 | 2,734614176 | 0,684893502 | 3,992758242 | 6,53E-05 | 0,003741521 | The Moose: suction sampler vs FISH | NA |
| gene_1679956 | 3,715144834 | 6,66289468 | 1,951719002 | 3,413859614 | 0,000640496 | 0,016350784 | The Moose: suction sampler vs FISH | NA |
| gene_1680394 | 7,197119369 | 4,150113656 | 1,37284863 | 3,022994353 | 0,002502869 | 0,03783928 | The Moose: suction sampler vs FISH | NA |
| gene_1680395 | 6,881315026 | 4,079952491 | 1,285638004 | 3,173484665 | 0,001506209 | 0,027998858 | The Moose: suction sampler vs FISH | NA |
| gene_1680899 | 22,81267738 | 5,656498224 | 1,307702191 | 4,325524773 | 1,52E-05 | 0,001388331 | The Moose: suction sampler vs FISH | NA |
| gene_1681135 | 5,405916925 | 6,571734781 | 1,703069717 | 3,85875852 | 0,000113964 | 0,005442151 | The Moose: suction sampler vs FISH | NA |
| gene_168126 | 85,90180358 | 2,757764156 | 0,670130968 | 4,115261476 | 3,87E-05 | 0,002639276 | The Moose: suction sampler vs FISH | NA |
| gene_1681394 | 9,720089347 | 3,899958564 | 1,314444309 | 2,967001749 | 0,003007192 | 0,04226239 | The Moose: suction sampler vs FISH | NA |
| gene_1681395 | 38,29568434 | 3,045348983 | 0,956749814 | 3,183014973 | 0,001457501 | 0,027413476 | The Moose: suction sampler vs FISH | NA |
| gene_1681658 | 21,05866587 | 2,639686403 | 0,871441179 | 3,029104505 | 0,002452798 | 0,037289539 | The Moose: suction sampler vs FISH | NA |
| gene_168183 | 10,38130201 | 8,364917675 | 1,91698304 | 4,363584601 | 1,28E-05 | 0,001251288 | The Moose: suction sampler vs FISH | NA |
| gene_1682390 | 48,84434491 | 2,973624932 | 0,9627733 | 3,088603444 | 0,002010997 | 0,033280228 | The Moose: suction sampler vs FISH | NA |
| gene_1682689 | 20,94991151 | 5,212799344 | 1,229920933 | 4,238320695 | 2,25E-05 | 0,001813559 | The Moose: suction sampler vs FISH | NA |
| gene_168355 | 6,966612636 | 7,971350795 | 2,01457688 | 3,956836233 | 7,59E-05 | 0,004178056 | The Moose: suction sampler vs FISH | NA |
| gene_1684114 | 7,203080161 | 4,503642021 | 1,4339588 | 3,140705312 | 0,001685415 | 0,029972233 | The Moose: suction sampler vs FISH | NA |
| gene_1684930 | 49,40567697 | 2,955440823 | 0,726135348 | 4,07009634 | 4,70E-05 | 0,003048986 | The Moose: suction sampler vs FISH | NA |
| gene_1685286 | 33,80692366 | 3,047928754 | 0,811135048 | 3,75760949 | 0,000171544 | 0,007073158 | The Moose: suction sampler vs FISH | NA |
| gene_1685694 | 136,2160867 | 4,274109084 | 0,821148839 | 5,205035776 | 1,94E-07 | 7,05E-05 | The Moose: suction sampler vs FISH | NA |
| gene_1685760 | 38,49976527 | 4,452117541 | 0,929104995 | 4,791834687 | 1,65E-06 | 0,000308848 | The Moose: suction sampler vs FISH | NA |
| gene_1686263 | 5,716569322 | 5,721798902 | 1,780841051 | 3,212975631 | 0,001313674 | 0,025735613 | The Moose: suction sampler vs FISH | NA |
| gene_1686264 | 15,74926372 | 4,987799992 | 1,202252324 | 4,148713122 | 3,34E-05 | 0,002386424 | The Moose: suction sampler vs FISH | NA |
| gene_1686265 | 20,75653326 | 4,179600471 | 0,864130709 | 4,836768822 | 1,32E-06 | 0,000270735 | The Moose: suction sampler vs FISH | NA |
| gene_1686294 | 11,23501474 | 5,456637865 | 1,454499985 | 3,751555807 | 0,000175741 | 0,007176644 | The Moose: suction sampler vs FISH | NA |
| gene_1688415 | 22,73547012 | 3,342626297 | 0,927083675 | 3,60552816 | 0,000311519 | 0,010442488 | The Moose: suction sampler vs FISH | NA |
| gene_1690215 | 4,744697546 | 5,556707625 | 1,779628593 | 3,122397363 | 0,001793846 | 0,031050919 | The Moose: suction sampler vs FISH | NA |
| gene_1691111 | 15,67658779 | 3,005382092 | 1,003272103 | 2,995580245 | 0,002739232 | 0,039798795 | The Moose: suction sampler vs FISH | NA |
| gene_1691227 | 13,02722737 | 3,584464306 | 1,168126355 | 3,068558715 | 0,002150941 | 0,034648127 | The Moose: suction sampler vs FISH | NA |
| gene_169126 | 3,386274944 | 5,453111623 | 1,852778492 | 2,943207538 | 0,003248306 | 0,044160858 | The Moose: suction sampler vs FISH | NA |
| gene_1692565 | 60,72770797 | 2,510991444 | 0,727572509 | 3,451190654 | 0,000558119 | 0,014960241 | The Moose: suction sampler vs FISH | NA |
| gene_1693096 | 3,89674995 | 6,902761897 | 1,933148999 | 3,570734537 | 0,000355982 | 0,011323111 | The Moose: suction sampler vs FISH | NA |
| gene_1694993 | 59,09294025 | 4,7782011 | 1,268543453 | 3,766683031 | 0,000165431 | 0,006927062 | The Moose: suction sampler vs FISH | NA |
| gene_1695244 | 12,40773329 | 3,889447425 | 1,238127159 | 3,141395775 | 0,001681446 | 0,029952421 | The Moose: suction sampler vs FISH | NA |
| gene_1696004 | 28,57230774 | 3,983070652 | 0,953517879 | 4,177237514 | 2,95E-05 | 0,002186829 | The Moose: suction sampler vs FISH | NA |
| gene_1696402 | 3,145958096 | 6,449404879 | 2,081912896 | 3,09782647 | 0,001949455 | 0,032644267 | The Moose: suction sampler vs FISH | NA |
| gene_1696998 | 26,68865168 | 22,27197885 | 4,248689929 | 5,242081495 | 1,59E-07 | 6,17E-05 | The Moose: suction sampler vs FISH | NA |
| gene_1697741 | 31,74555048 | 3,992029331 | 0,96465903 | 4,138280168 | 3,50E-05 | 0,002454294 | The Moose: suction sampler vs FISH | NA |
| gene_1697742 | 9,25070324 | 4,975095872 | 1,38436804 | 3,593766779 | 0,000325932 | 0,010724931 | The Moose: suction sampler vs FISH | NA |
| gene_1698014 | 8,96414278 | 4,992100295 | 1,466120039 | 3,404973783 | 0,000661704 | 0,016723625 | The Moose: suction sampler vs FISH | NA |

| | | | | | | | | |
|---|---|---|---|---|---|---|---|---|
| gene_1698564 | 42,63574781 | 2,493959748 | 0,746910325 | 3,339035042 | 0,0008407 | 0,01960787 | The Moose: suction sampler vs FISH | NA |
| gene_169904 | 14,27211236 | 4,385259965 | 1,357342316 | 3,230769359 | 0,001234575 | 0,024693184 | The Moose: suction sampler vs FISH | NA |
| gene_1699118 | 6,103209702 | 5,939395665 | 1,716600805 | 3,45997488 | 0,000540226 | 0,014635516 | The Moose: suction sampler vs FISH | NA |
| gene_1699248 | 24,45341014 | 3,390925756 | 1,068956523 | 3,172183043 | 0,001512976 | 0,028048634 | The Moose: suction sampler vs FISH | NA |
| gene_1700564 | 11,96663811 | 5,651390387 | 1,367257899 | 4,133375563 | 3,57E-05 | 0,002490955 | The Moose: suction sampler vs FISH | NA |
| gene_1700989 | 7,54019477 | 5,691899666 | 1,858217838 | 3,063096021 | 0,002190598 | 0,034974172 | The Moose: suction sampler vs FISH | NA |
| gene_1701673 | 28,22211218 | 9,982958019 | 1,639656427 | 6,088445026 | 1,14E-09 | 1,95E-06 | The Moose: suction sampler vs FISH | NA |
| gene_1703183 | 58,22636584 | 2,879030911 | 0,82016883 | 3,510290571 | 0,000447617 | 0,013030222 | The Moose: suction sampler vs FISH | NA |
| gene_1703641 | 78,76006398 | 3,423537852 | 0,798459945 | 4,287676386 | 1,81E-05 | 0,001552767 | The Moose: suction sampler vs FISH | NA |
| gene_1704786 | 5,963603231 | 4,912165254 | 1,62569546 | 3,021577765 | 0,00251461 | 0,037931428 | The Moose: suction sampler vs FISH | NA |
| gene_1705304 | 5,275165067 | 6,895358726 | 1,898844212 | 3,631345152 | 0,000281948 | 0,009790854 | The Moose: suction sampler vs FISH | NA |
| gene_1705305 | 9,823443748 | 4,141917221 | 1,363761387 | 3,037127507 | 0,002388444 | 0,036660244 | The Moose: suction sampler vs FISH | NA |
| gene_1705386 | 27,6270192 | 3,362883752 | 0,888511024 | 3,784853154 | 0,000153799 | 0,006570706 | The Moose: suction sampler vs FISH | NA |
| gene_1708317 | 4,312972098 | 7,339170442 | 2,187735552 | 3,354688109 | 0,000794545 | 0,018842546 | The Moose: suction sampler vs FISH | NA |
| gene_1708357 | 27,924532 | 2,92801986 | 0,913434353 | 3,205506614 | 0,001348249 | 0,026096408 | The Moose: suction sampler vs FISH | NA |
| gene_1708793 | 44,4173076 | 2,8649062 | 0,77665388 | 3,688781158 | 0,000225331 | 0,008447897 | The Moose: suction sampler vs FISH | NA |
| gene_1708857 | 4,377851033 | 5,517636235 | 1,785260679 | 3,090661381 | 0,001997112 | 0,033147553 | The Moose: suction sampler vs FISH | NA |
| gene_1709702 | 4,38065176 | 6,744082404 | 1,879803708 | 3,587652464 | 0,000333669 | 0,010886847 | The Moose: suction sampler vs FISH | NA |
| gene_1710087 | 14,80560791 | 4,944779154 | 1,216884036 | 4,063476064 | 4,83E-05 | 0,003090397 | The Moose: suction sampler vs FISH | NA |
| gene_1710268 | 17,74970313 | 3,620524839 | 1,254759608 | 2,885433048 | 0,003908756 | 0,049120591 | The Moose: suction sampler vs FISH | NA |
| gene_1711141 | 16,41928318 | 4,919890215 | 1,24584911 | 3,949025749 | 7,85E-05 | 0,004252351 | The Moose: suction sampler vs FISH | NA |
| gene_1711670 | 12,65193789 | 5,161214123 | 1,424422945 | 3,623371934 | 0,000290787 | 0,009970526 | The Moose: suction sampler vs FISH | NA |
| gene_1712933 | 5,673868295 | 6,740458676 | 1,757335698 | 3,835612447 | 0,000125252 | 0,005797411 | The Moose: suction sampler vs FISH | NA |
| gene_171385 | 6,776081929 | 4,312596442 | 1,29885128 | 3,320315812 | 0,000899157 | 0,020477074 | The Moose: suction sampler vs FISH | NA |
| gene_1714686 | 13,235759 | 3,515633866 | 1,089119982 | 3,227958282 | 0,001246772 | 0,024884919 | The Moose: suction sampler vs FISH | NA |
| gene_1714689 | 5,679074566 | 4,926217527 | 1,679164678 | 2,933731033 | 0,003349143 | 0,04493566 | The Moose: suction sampler vs FISH | NA |
| gene_1714898 | 141,1406215 | 4,142112008 | 0,695142371 | 5,958652758 | 2,54E-09 | 3,30E-06 | The Moose: suction sampler vs FISH | NA |
| gene_171581 | 4,109793914 | 6,226229794 | 2,111915796 | 2,948143012 | 0,003196891 | 0,043756184 | The Moose: suction sampler vs FISH | NA |
| gene_1716558 | 59,513034 | 2,534726257 | 0,863193468 | 2,936452082 | 0,003319901 | 0,044733024 | The Moose: suction sampler vs FISH | NA |
| gene_1716733 | 10,14543238 | 2,991659386 | 0,976084702 | 3,064958788 | 0,002177 | 0,034864754 | The Moose: suction sampler vs FISH | NA |
| gene_1718027 | 11,10406407 | 7,651432508 | 1,72868059 | 4,426169041 | 9,59E-06 | 0,001029043 | The Moose: suction sampler vs FISH | NA |
| gene_1718162 | 5,104051713 | 6,504396356 | 2,066756603 | 3,147151603 | 0,001648695 | 0,029566792 | The Moose: suction sampler vs FISH | NA |
| gene_1719695 | 10,6579378 | 3,109762627 | 1,001275013 | 3,105802688 | 0,001897633 | 0,032172588 | The Moose: suction sampler vs FISH | NA |
| gene_1719785 | 8,98730171 | 4,991453838 | 1,306800589 | 3,819598704 | 0,000133669 | 0,006033999 | The Moose: suction sampler vs FISH | NA |
| gene_1720070 | 9,002420131 | 6,084820683 | 1,773055954 | 3,431826655 | 0,000599531 | 0,015635485 | The Moose: suction sampler vs FISH | NA |
| gene_1720508 | 7,140151477 | 6,753881564 | 1,949281645 | 3,464805397 | 0,000530615 | 0,014487375 | The Moose: suction sampler vs FISH | NA |
| gene_1720511 | 7,545039246 | 4,552372149 | 1,46447011 | 3,108545621 | 0,001880106 | 0,032001019 | The Moose: suction sampler vs FISH | NA |
| gene_1720682 | 5,712768385 | 5,826355237 | 1,783860526 | 3,26614954 | 0,001090207 | 0,023015299 | The Moose: suction sampler vs FISH | NA |
| gene_1720721 | 5,289510525 | 5,825406682 | 1,970523334 | 2,95627389 | 0,003113804 | 0,043133087 | The Moose: suction sampler vs FISH | NA |
| gene_1721953 | 4,298965462 | 6,394646548 | 1,897826725 | 3,369457529 | 0,000753163 | 0,018226048 | The Moose: suction sampler vs FISH | NA |
| gene_1722343 | 45,37727444 | 2,425020379 | 0,799510882 | 3,033129921 | 0,002420314 | 0,036994785 | The Moose: suction sampler vs FISH | NA |
| gene_1722939 | 39,47102459 | 2,689938852 | 0,79695532 | 3,375269332 | 0,000737435 | 0,017981707 | The Moose: suction sampler vs FISH | NA |
| gene_1723126 | 5,397642072 | 6,173064351 | 1,809412885 | 3,411639434 | 0,000645735 | 0,016438815 | The Moose: suction sampler vs FISH | NA |
| gene_1723519 | 6,823184622 | 5,103188756 | 1,709751084 | 2,984755385 | 0,002838052 | 0,040790747 | The Moose: suction sampler vs FISH | NA |
| gene_1723875 | 8,795906578 | 4,248843635 | 1,4128985 | 3,007182494 | 0,002636814 | 0,038965959 | The Moose: suction sampler vs FISH | NA |
| gene_1724352 | 7,417001728 | 4,711457983 | 1,637716016 | 2,87684674 | 0,004016706 | 0,049918912 | The Moose: suction sampler vs FISH | NA |
| gene_1724589 | 5,512226181 | 5,682446606 | 1,837389638 | 3,092673698 | 0,001983621 | 0,033037287 | The Moose: suction sampler vs FISH | NA |
| gene_1725193 | 3,097170898 | 6,758562643 | 2,311209572 | 2,924253484 | 0,003452834 | 0,045700744 | The Moose: suction sampler vs FISH | NA |
| gene_1727354 | 11,97310394 | 3,699298522 | 1,139379441 | 3,24676608 | 0,001167243 | 0,023920816 | The Moose: suction sampler vs FISH | NA |
| gene_1728571 | 174,4754372 | 5,971796675 | 0,95916783 | 6,226018522 | 4,78E-10 | 1,06E-06 | The Moose: suction sampler vs FISH | NA |
| gene_1729806 | 8,661545084 | 3,672435019 | 1,183382997 | 3,103335968 | 0,001913523 | 0,032292895 | The Moose: suction sampler vs FISH | NA |
| gene_1729993 | 10,61354981 | 4,933809294 | 1,234340688 | 3,997121168 | 6,41E-05 | 0,00371303 | The Moose: suction sampler vs FISH | NA |
| gene_1729994 | 10,29580831 | 4,482296 | 1,281991322 | 3,4963544 | 0,000471662 | 0,013437239 | The Moose: suction sampler vs FISH | NA |
| gene_1730380 | 14,45369444 | 3,708645614 | 1,10849636 | 3,345654299 | 0,000820887 | 0,019261989 | The Moose: suction sampler vs FISH | NA |
| gene_1731500 | 16,2325859 | 3,736856729 | 1,063768984 | 3,512846103 | 0,000443334 | 0,012965789 | The Moose: suction sampler vs FISH | NA |
| gene_1731604 | 82,4345979 | 2,533552441 | 0,748264186 | 3,385906325 | 0,000709436 | 0,017490609 | The Moose: suction sampler vs FISH | NA |
| gene_1731605 | 124,8606143 | 2,459801619 | 0,769563746 | 3,196358498 | 0,001391741 | 0,026578131 | The Moose: suction sampler vs FISH | NA |
| gene_1731607 | 65,99533033 | 2,401485758 | 0,713631778 | 3,365160902 | 0,00076499 | 0,018403117 | The Moose: suction sampler vs FISH | NA |
| gene_1731992 | 15,15797898 | 3,31814198 | 1,013853177 | 3,272803257 | 0,001064866 | 0,022656159 | The Moose: suction sampler vs FISH | NA |
| gene_1732197 | 10,16640558 | 5,146886408 | 1,373353174 | 3,747678678 | 0,000178479 | 0,007226628 | The Moose: suction sampler vs FISH | NA |
| gene_173477 | 44,62998482 | 7,248988934 | 1,133664017 | 6,394300977 | 1,61E-10 | 4,94E-07 | The Moose: suction sampler vs FISH | NA |
| gene_1735613 | 4,26025766 | 6,802621687 | 1,943731247 | 3,499774826 | 0,000465651 | 0,013354236 | The Moose: suction sampler vs FISH | NA |
| gene_1737210 | 17,52586215 | 4,065546681 | 1,18624559 | 3,427238606 | 0,000609753 | 0,015824492 | The Moose: suction sampler vs FISH | NA |
| gene_1739206 | 7,336554914 | 5,030301919 | 1,608617076 | 3,127097179 | 0,001765416 | 0,030776395 | The Moose: suction sampler vs FISH | NA |
| gene_1740258 | 14,87081423 | 4,029870029 | 1,229436667 | 3,27781832 | 0,001046127 | 0,022413302 | The Moose: suction sampler vs FISH | NA |

| | | | | | | | | |
|---|---|---|---|---|---|---|---|---|
| gene_1741503 | 10,29311447 | 4,632439733 | 1,495230201 | 3,098144841 | 0,001947362 | 0,032638983 | The Moose: suction sampler vs FISH | NA |
| gene_1741757 | 4,114077243 | 5,890358927 | 1,923015017 | 3,063085247 | 0,002190676 | 0,034974172 | The Moose: suction sampler vs FISH | NA |
| gene_1743950 | 24,74090598 | 4,179320731 | 1,021836724 | 4,090008346 | 4,31E-05 | 0,002856724 | The Moose: suction sampler vs FISH | NA |
| gene_1746348 | 41,85180769 | 3,144599476 | 0,922088303 | 3,410301883 | 0,00064891 | 0,016491073 | The Moose: suction sampler vs FISH | NA |
| gene_1746349 | 6,837803295 | 6,65080016 | 1,755391987 | 3,788783479 | 0,000151387 | 0,006498306 | The Moose: suction sampler vs FISH | NA |
| gene_1747095 | 4,26038919 | 5,214467801 | 1,778640291 | 2,931715776 | 0,003370951 | 0,045047216 | The Moose: suction sampler vs FISH | NA |
| gene_1747686 | 2,800192091 | 6,160887154 | 2,007668493 | 3,068677511 | 0,002150086 | 0,034648127 | The Moose: suction sampler vs FISH | NA |
| gene_1748509 | 30,10862922 | 3,455704525 | 1,044760738 | 3,307651598 | 0,000940818 | 0,020945244 | The Moose: suction sampler vs FISH | NA |
| gene_1749580 | 60,79704086 | 2,645440874 | 0,826899953 | 3,199227264 | 0,001377965 | 0,026458729 | The Moose: suction sampler vs FISH | NA |
| gene_1752841 | 8,907275898 | 5,736908981 | 1,511997099 | 3,794259252 | 0,000148085 | 0,006412819 | The Moose: suction sampler vs FISH | NA |
| gene_1753332 | 62,6361932 | 2,825255171 | 0,760891769 | 3,713084153 | 0,000204749 | 0,007943884 | The Moose: suction sampler vs FISH | NA |
| gene_1754991 | 7,031078273 | 5,049549764 | 1,696837533 | 2,975859307 | 0,002921689 | 0,041457385 | The Moose: suction sampler vs FISH | NA |
| gene_1755187 | 11,40794342 | 3,029572209 | 0,991112893 | 3,056737764 | 0,002237599 | 0,035360455 | The Moose: suction sampler vs FISH | NA |
| gene_1755839 | 8,869110318 | 4,379747864 | 1,389662758 | 3,1516624 | 0,001623439 | 0,029288491 | The Moose: suction sampler vs FISH | NA |
| gene_1756168 | 4,649346063 | 6,595127906 | 2,085952054 | 3,161687199 | 0,00156858 | 0,028672058 | The Moose: suction sampler vs FISH | NA |
| gene_1757753 | 4,339178862 | 7,289920686 | 2,045507729 | 3,563868562 | 0,000365429 | 0,011490192 | The Moose: suction sampler vs FISH | NA |
| gene_1757829 | 9,13052615 | 6,582129726 | 1,720859898 | 3,824907382 | 0,000130821 | 0,005945648 | The Moose: suction sampler vs FISH | NA |
| gene_1759096 | 20,5973041 | 6,255930725 | 1,259921694 | 4,965332967 | 6,86E-07 | 0,000170749 | The Moose: suction sampler vs FISH | NA |
| gene_1759097 | 39,01343281 | 3,594208784 | 1,029056945 | 3,492720982 | 0,000478126 | 0,01356876 | The Moose: suction sampler vs FISH | NA |
| gene_1759568 | 235,8048705 | 13,232323 | 2,546903258 | 5,19545568 | 2,04E-07 | 7,35E-05 | The Moose: suction sampler vs FISH | NA |
| gene_175972 | 12,78823354 | 5,058803221 | 1,260159179 | 4,01441604 | 5,96E-05 | 0,003539821 | The Moose: suction sampler vs FISH | NA |
| gene_1760606 | 191,1470224 | 3,043730956 | 0,773209366 | 3,936490024 | 8,27E-05 | 0,004397236 | The Moose: suction sampler vs FISH | NA |
| gene_1761339 | 6,218802289 | 5,90303405 | 1,650593878 | 3,576309188 | 0,000348479 | 0,011156964 | The Moose: suction sampler vs FISH | NA |
| gene_1761993 | 5,361613931 | 5,865705033 | 1,845366395 | 3,178612686 | 0,001479817 | 0,027676381 | The Moose: suction sampler vs FISH | NA |
| gene_1761995 | 5,906255079 | 7,272263569 | 2,013253449 | 3,612194764 | 0,000303616 | 0,010280823 | The Moose: suction sampler vs FISH | NA |
| gene_1761996 | 6,611987828 | 7,451433581 | 1,962895278 | 3,796144228 | 0,000146964 | 0,006389958 | The Moose: suction sampler vs FISH | NA |
| gene_1762432 | 16,78155024 | 2,931142257 | 0,907597518 | 3,229561783 | 0,001239801 | 0,024777463 | The Moose: suction sampler vs FISH | NA |
| gene_1762967 | 24,27968516 | 3,20805399 | 0,980719315 | 3,271123493 | 0,001071211 | 0,022725334 | The Moose: suction sampler vs FISH | NA |
| gene_176361 | 119,2303721 | -2,479648713 | 0,672990999 | -3,684519876 | 0,000229134 | 0,008544153 | The Moose: suction sampler vs FISH | NA |
| gene_1764558 | 3,689428488 | 5,22235514 | 1,796542779 | 2,906891614 | 0,003650397 | 0,047267873 | The Moose: suction sampler vs FISH | NA |
| gene_1764691 | 3,826107533 | 6,415040661 | 2,162293804 | 2,966775675 | 0,003009404 | 0,042267959 | The Moose: suction sampler vs FISH | NA |
| gene_1765625 | 25,65611548 | 3,151163705 | 0,986824541 | 3,193236054 | 0,001406879 | 0,026776907 | The Moose: suction sampler vs FISH | NA |
| gene_1765635 | 11,42881756 | 3,985499607 | 1,094646873 | 3,640899825 | 0,000271687 | 0,009538157 | The Moose: suction sampler vs FISH | NA |
| gene_176580 | 9,693223403 | 5,79316127 | 1,847431915 | 3,135791485 | 0,00171391 | 0,030278141 | The Moose: suction sampler vs FISH | NA |
| gene_1767270 | 31,02675807 | 10,25801551 | 3,416535547 | 3,002461226 | 0,002678061 | 0,039306464 | The Moose: suction sampler vs FISH | NA |
| gene_1767852 | 25,24261196 | 3,129995264 | 0,814249952 | 3,844022659 | 0,000121034 | 0,005669215 | The Moose: suction sampler vs FISH | NA |
| gene_1769951 | 49,09991452 | 2,90800342 | 0,977387931 | 2,97528067 | 0,002927206 | 0,041503598 | The Moose: suction sampler vs FISH | NA |
| gene_1770038 | 22,19934378 | 3,149955772 | 1,03905293 | 3,031564303 | 0,002432901 | 0,037131069 | The Moose: suction sampler vs FISH | NA |
| gene_1770440 | 9,572570566 | 4,329193447 | 1,433349783 | 3,020332858 | 0,00252497 | 0,037994269 | The Moose: suction sampler vs FISH | NA |
| gene_1770782 | 10,98151874 | 3,458177575 | 1,007675813 | 3,431835447 | 0,000599511 | 0,015635485 | The Moose: suction sampler vs FISH | NA |
| gene_1771896 | 10,6236796 | 5,809142093 | 1,584522941 | 3,666177335 | 0,000246203 | 0,008947252 | The Moose: suction sampler vs FISH | NA |
| gene_1772036 | 48,26873459 | 3,13397434 | 1,004124644 | 3,121100911 | 0,001801763 | 0,031137829 | The Moose: suction sampler vs FISH | NA |
| gene_177223 | 41,35217048 | 3,965180936 | 0,906334875 | 4,374962327 | 1,21E-05 | 0,001203799 | The Moose: suction sampler vs FISH | NA |
| gene_177227 | 17,5132107 | 5,065164122 | 1,416363229 | 3,576175955 | 0,000348657 | 0,011157784 | The Moose: suction sampler vs FISH | NA |
| gene_1772404 | 5,147262582 | 6,855005151 | 1,937246476 | 3,538530194 | 0,000402361 | 0,012231549 | The Moose: suction sampler vs FISH | NA |
| gene_1772618 | 17,7585378 | 2,768276537 | 0,81820614 | 3,383348526 | 0,000716077 | 0,017595273 | The Moose: suction sampler vs FISH | NA |
| gene_1776549 | 107,9022742 | 5,976244733 | 1,88522017 | 3,170051344 | 0,00152412 | 0,02816289 | The Moose: suction sampler vs FISH | NA |
| gene_1776840 | 4,899963397 | 5,196248948 | 1,718379309 | 3,0239243 | 0,002495189 | 0,037769306 | The Moose: suction sampler vs FISH | NA |
| gene_177757 | 65,87585213 | 2,424243156 | 0,759125131 | 3,193469769 | 0,001405741 | 0,026768114 | The Moose: suction sampler vs FISH | NA |
| gene_1777813 | 162,067011 | 2,689812254 | 0,792508273 | 3,394049433 | 0,000688673 | 0,017144038 | The Moose: suction sampler vs FISH | NA |
| gene_1778146 | 4,445399048 | 5,774028334 | 1,82055581 | 3,171574472 | 0,00151615 | 0,02807828 | The Moose: suction sampler vs FISH | NA |
| gene_1779795 | 12,00611431 | 3,34389864 | 1,14271302 | 2,926280335 | 0,003430417 | 0,045535325 | The Moose: suction sampler vs FISH | NA |
| gene_1780581 | 6,774365084 | 5,084833478 | 1,560973967 | 3,257474876 | 0,001124082 | 0,023420768 | The Moose: suction sampler vs FISH | NA |
| gene_1781822 | 6,075204914 | 4,900485462 | 1,58699044 | 3,087911143 | 0,002015687 | 0,033327815 | The Moose: suction sampler vs FISH | NA |
| gene_1782604 | 149,1616196 | 3,273351126 | 0,829057246 | 3,948281185 | 7,87E-05 | 0,004257123 | The Moose: suction sampler vs FISH | NA |
| gene_1782605 | 434,5586096 | 3,331260756 | 0,790296477 | 4,215203856 | 2,50E-05 | 0,001960256 | The Moose: suction sampler vs FISH | NA |
| gene_1783117 | 6,590355565 | 7,049887436 | 1,604073533 | 4,394990187 | 1,11E-05 | 0,001137912 | The Moose: suction sampler vs FISH | NA |
| gene_1783517 | 5,441004452 | 4,974372122 | 1,603717839 | 3,101775139 | 0,00192364 | 0,032411505 | The Moose: suction sampler vs FISH | NA |
| gene_1783594 | 6,177783708 | 5,242792785 | 1,755613928 | 2,986301659 | 0,002823739 | 0,040656645 | The Moose: suction sampler vs FISH | NA |
| gene_1784928 | 4,952642718 | 6,572321269 | 1,918603526 | 3,42557552 | 0,000613498 | 0,015893611 | The Moose: suction sampler vs FISH | NA |
| gene_1785103 | 13,60112562 | 2,950440302 | 0,980784815 | 3,008244272 | 0,002627618 | 0,038900511 | The Moose: suction sampler vs FISH | NA |
| gene_1786023 | 54,24676488 | 3,476762383 | 0,968154089 | 3,591125032 | 0,000329254 | 0,010800372 | The Moose: suction sampler vs FISH | NA |
| gene_1786722 | 10,78161937 | 3,517000037 | 1,150892634 | 3,055888911 | 0,002243944 | 0,035391185 | The Moose: suction sampler vs FISH | NA |
| gene_1786723 | 7,402998668 | 5,131761089 | 1,583474453 | 3,240823419 | 0,00119185 | 0,024234611 | The Moose: suction sampler vs FISH | NA |
| gene_178684 | 52,6110331 | 3,80079396 | 1,126564077 | 3,373792968 | 0,000741401 | 0,01803053 | The Moose: suction sampler vs FISH | NA |

| | | | | | | | | |
|---|---|---|---|---|---|---|---|---|
| gene_1787565 | 427,7705785 | 3,838625907 | 0,677537185 | 5,665557542 | 1,47E-08 | 1,16E-05 | The Moose: suction sampler vs FISH | NA |
| gene_1787566 | 582,189992 | 3,292941296 | 0,686574566 | 4,796188874 | 1,62E-06 | 0,000306899 | The Moose: suction sampler vs FISH | NA |
| gene_1787567 | 22,91897328 | 4,195230628 | 1,02708773 | 4,084588402 | 4,42E-05 | 0,002903281 | The Moose: suction sampler vs FISH | NA |
| gene_1788173 | 179,8345871 | 3,927260235 | 0,780121526 | 5,034164685 | 4,80E-07 | 0,000136096 | The Moose: suction sampler vs FISH | NA |
| gene_1788174 | 112,9010536 | 2,700580182 | 0,74858076 | 3,607600312 | 0,000309042 | 0,010380804 | The Moose: suction sampler vs FISH | NA |
| gene_1788578 | 12,1838904 | -4,76102982 | 1,611065129 | -2,9552063 | 0,0031246 | 0,043209617 | The Moose: suction sampler vs FISH | NA |
| gene_1788819 | 6,29428167 | 5,980038828 | 1,952717862 | 3,062418255 | 0,002195564 | 0,03502552 | The Moose: suction sampler vs FISH | NA |
| gene_178930 | 5,91625047 | 7,039992772 | 1,762680361 | 3,993913433 | 6,50E-05 | 0,003740709 | The Moose: suction sampler vs FISH | NA |
| gene_1789707 | 3,573881357 | 6,061950646 | 2,083732738 | 2,90917858 | 0,003623798 | 0,047056129 | The Moose: suction sampler vs FISH | NA |
| gene_1789929 | 43,72767074 | 3,889604773 | 1,016412636 | 3,826796947 | 0,000129822 | 0,005918523 | The Moose: suction sampler vs FISH | NA |
| gene_1789934 | 16,14328637 | 4,783481905 | 1,034370776 | 4,624533114 | 3,75E-06 | 0,000536467 | The Moose: suction sampler vs FISH | NA |
| gene_1789936 | 1242,292647 | 3,66322377 | 0,779513675 | 4,699370759 | 2,61E-06 | 0,000418483 | The Moose: suction sampler vs FISH | NA |
| gene_1789937 | 247,0325441 | 3,665933376 | 0,828263621 | 4,426046591 | 9,60E-06 | 0,001029043 | The Moose: suction sampler vs FISH | NA |
| gene_1790604 | 43,26977389 | 4,416276951 | 0,979714727 | 4,507717225 | 6,55E-06 | 0,000806199 | The Moose: suction sampler vs FISH | NA |
| gene_1790639 | 15,33014606 | 3,493571628 | 1,026589552 | 3,40308512 | 0,000666295 | 0,016770408 | The Moose: suction sampler vs FISH | NA |
| gene_1790735 | 45,45464085 | 2,47124601 | 0,751908887 | 3,286629606 | 0,001013941 | 0,021902617 | The Moose: suction sampler vs FISH | NA |
| gene_1792021 | 7,391894131 | 3,735373238 | 1,219490475 | 3,063060609 | 0,002190857 | 0,034974172 | The Moose: suction sampler vs FISH | NA |
| gene_1792023 | 8,874186837 | 4,683649582 | 1,530060985 | 3,061086865 | 0,002205351 | 0,035127306 | The Moose: suction sampler vs FISH | NA |
| gene_1793113 | 94,31779629 | 2,632653198 | 0,83319434 | 3,15971085 | 0,001579258 | 0,028767012 | The Moose: suction sampler vs FISH | NA |
| gene_1794217 | 17,30481772 | 3,590682231 | 1,118392351 | 3,210574738 | 0,001324698 | 0,02582757 | The Moose: suction sampler vs FISH | NA |
| gene_1794440 | 10,06983541 | 6,191933592 | 1,615510167 | 3,832803852 | 0,000126691 | 0,005828286 | The Moose: suction sampler vs FISH | NA |
| gene_1795285 | 3,733868942 | 5,353477403 | 1,816933846 | 2,94643496 | 0,0032146 | 0,043916724 | The Moose: suction sampler vs FISH | NA |
| gene_1795429 | 8,598191343 | 3,733304216 | 1,249189408 | 2,988581388 | 0,002802758 | 0,040441767 | The Moose: suction sampler vs FISH | NA |
| gene_1795516 | 4,081059538 | 6,231091616 | 1,95781878 | 3,182670266 | 0,001459237 | 0,02742417 | The Moose: suction sampler vs FISH | NA |
| gene_1797089 | 22,17838392 | 3,388624028 | 0,958232954 | 3,536325916 | 0,000405734 | 0,01227817 | The Moose: suction sampler vs FISH | NA |
| gene_1797137 | 25,00311173 | 4,057688931 | 1,0786293 | 3,761893851 | 0,000168632 | 0,007017082 | The Moose: suction sampler vs FISH | NA |
| gene_1799059 | 5,278798585 | 7,324646955 | 1,93257777 | 3,790091695 | 0,000150592 | 0,006479324 | The Moose: suction sampler vs FISH | NA |
| gene_1799425 | 5,607742 | 6,296297038 | 1,873422847 | 3,360852061 | 0,000777024 | 0,018587091 | The Moose: suction sampler vs FISH | NA |
| gene_1799863 | 25,49682996 | 3,546125658 | 0,956941626 | 3,705686494 | 0,000210819 | 0,008076994 | The Moose: suction sampler vs FISH | NA |
| gene_1800443 | 3,760537834 | 5,611546614 | 1,910320133 | 2,93749017 | 0,003308807 | 0,044639108 | The Moose: suction sampler vs FISH | NA |
| gene_1801825 | 6,522378145 | 5,453706716 | 1,687551615 | 3,231727354 | 0,001230444 | 0,024673038 | The Moose: suction sampler vs FISH | NA |
| gene_1801996 | 11,8752365 | 6,781028662 | 1,54034162 | 4,4022888 | 1,07E-05 | 0,001112737 | The Moose: suction sampler vs FISH | NA |
| gene_1802738 | 279,2752291 | 4,43451787 | 0,841320801 | 5,270900071 | 1,36E-07 | 5,51E-05 | The Moose: suction sampler vs FISH | NA |
| gene_1802815 | 56,02956541 | -2,561976228 | 0,803373606 | -3,189022155 | 0,001427549 | 0,027043168 | The Moose: suction sampler vs FISH | NA |
| gene_1802918 | 12,09888997 | 4,240490114 | 1,271146982 | 3,335955774 | 0,000850067 | 0,019757329 | The Moose: suction sampler vs FISH | NA |
| gene_1803970 | 8,362415508 | 4,295202861 | 1,258449829 | 3,413090265 | 0,000642307 | 0,01638018 | The Moose: suction sampler vs FISH | NA |
| gene_1805431 | 13,38870021 | 3,199620494 | 1,009334673 | 3,170029307 | 0,001524236 | 0,02816289 | The Moose: suction sampler vs FISH | NA |
| gene_1806577 | 3,813950622 | 6,186128353 | 1,871592394 | 3,305275429 | 0,000948831 | 0,021066172 | The Moose: suction sampler vs FISH | NA |
| gene_1807415 | 18,91502635 | 3,602471063 | 1,175504028 | 3,06461822 | 0,00217948 | 0,034889259 | The Moose: suction sampler vs FISH | NA |
| gene_1807983 | 6,993671958 | 6,883482794 | 1,841984878 | 3,736992022 | 0,000186235 | 0,007458025 | The Moose: suction sampler vs FISH | NA |
| gene_1808389 | 77,10183883 | 7,520729477 | 1,207734058 | 6,227140344 | 4,75E-10 | 1,06E-06 | The Moose: suction sampler vs FISH | NA |
| gene_1809194 | 5,278223655 | 5,491677978 | 1,852783043 | 2,964015672 | 0,003036528 | 0,042463407 | The Moose: suction sampler vs FISH | NA |
| gene_1809565 | 9,247304371 | 5,933790452 | 1,515511693 | 3,915370947 | 9,03E-05 | 0,004681876 | The Moose: suction sampler vs FISH | NA |
| gene_1809566 | 8,621262658 | 4,347811782 | 1,323106066 | 3,286064432 | 0,001015978 | 0,021927264 | The Moose: suction sampler vs FISH | NA |
| gene_1809567 | 17,2597287 | 4,735211803 | 1,062145206 | 4,458158616 | 8,27E-06 | 0,000940188 | The Moose: suction sampler vs FISH | NA |
| gene_1809568 | 4,617868637 | 5,625803317 | 1,674553069 | 3,359584967 | 0,000780596 | 0,018638136 | The Moose: suction sampler vs FISH | NA |
| gene_181032 | 6,802309198 | 4,335060826 | 1,483812462 | 2,921569227 | 0,003482729 | 0,045939118 | The Moose: suction sampler vs FISH | NA |
| gene_1810484 | 37,57226084 | 3,323338158 | 0,955632776 | 3,477630991 | 0,000505866 | 0,014057251 | The Moose: suction sampler vs FISH | NA |
| gene_1810859 | 5,773964103 | 4,154944127 | 1,431477217 | 2,902556938 | 0,003701299 | 0,047701033 | The Moose: suction sampler vs FISH | NA |
| gene_1810935 | 26,88213744 | 2,304008668 | 0,795216957 | 2,897333423 | 0,003763495 | 0,048129883 | The Moose: suction sampler vs FISH | NA |
| gene_1811413 | 13,00080907 | 3,990380354 | 1,360293672 | 2,933469762 | 0,003351963 | 0,044942491 | The Moose: suction sampler vs FISH | NA |
| gene_1811821 | 66,93093078 | 3,690301537 | 0,897048916 | 4,113824199 | 3,89E-05 | 0,002650086 | The Moose: suction sampler vs FISH | NA |
| gene_1813410 | 3,990562542 | 6,431359914 | 1,973896984 | 3,258204438 | 0,001121196 | 0,023373898 | The Moose: suction sampler vs FISH | NA |
| gene_1813663 | 57,51249902 | 3,417068993 | 0,797737763 | 4,283448964 | 1,84E-05 | 0,001569713 | The Moose: suction sampler vs FISH | NA |
| gene_181384 | 16,25108648 | 3,30068123 | 1,073541396 | 3,07457285 | 0,002108042 | 0,034252904 | The Moose: suction sampler vs FISH | NA |
| gene_1814021 | 5,225859739 | 5,656795785 | 1,765311006 | 3,204418806 | 0,001353355 | 0,026157095 | The Moose: suction sampler vs FISH | NA |
| gene_181486 | 318,5309253 | -2,003458265 | 0,585178026 | -3,423673096 | 0,000617809 | 0,015954633 | The Moose: suction sampler vs FISH | NA |
| gene_1814909 | 12,96952227 | 3,570161523 | 1,207046131 | 2,957767256 | 0,00309876 | 0,043012037 | The Moose: suction sampler vs FISH | NA |
| gene_1815989 | 12,87240642 | 6,30220319 | 1,688817126 | 3,731726244 | 0,000190172 | 0,007586741 | The Moose: suction sampler vs FISH | NA |
| gene_1816154 | 5,249204967 | 5,780114954 | 1,756094769 | 3,291459583 | 0,000996689 | 0,021695866 | The Moose: suction sampler vs FISH | NA |
| gene_1816379 | 4,181809239 | 5,957613962 | 1,794777489 | 3,319416474 | 0,000902058 | 0,02049231 | The Moose: suction sampler vs FISH | NA |
| gene_1818296 | 5,231019965 | 5,011852814 | 1,576332795 | 3,17943827 | 0,001475608 | 0,027618763 | The Moose: suction sampler vs FISH | NA |
| gene_1818376 | 6,644156719 | 5,170946852 | 1,647646563 | 3,13838354 | 0,001698824 | 0,030103022 | The Moose: suction sampler vs FISH | NA |
| gene_1821173 | 7,798684898 | 6,934898586 | 1,82634518 | 3,797145613 | 0,000146372 | 0,006372423 | The Moose: suction sampler vs FISH | NA |
| gene_182140 | 45,56948111 | 2,202870588 | 0,716207894 | 3,075741845 | 0,002099796 | 0,034184632 | The Moose: suction sampler vs FISH | NA |

| | | | | | | | | |
|---|---|---|---|---|---|---|---|---|
| gene_1822285 | 9,13813534 | 5,997823293 | 1,511393583 | 3,968405953 | 7,24E-05 | 0,004044226 | The Moose: suction sampler vs FISH | NA |
| gene_1822301 | 4,034780492 | 6,067834506 | 1,911624507 | 3,174176981 | 0,00150262 | 0,027960465 | The Moose: suction sampler vs FISH | NA |
| gene_1822348 | 13,96144161 | 5,748932191 | 1,177811143 | 4,881030568 | 1,06E-06 | 0,00023068 | The Moose: suction sampler vs FISH | NA |
| gene_1822349 | 27,37361492 | 3,802239476 | 0,932863599 | 4,075879348 | 4,58E-05 | 0,002987379 | The Moose: suction sampler vs FISH | NA |
| gene_1822650 | 8,153224963 | 4,702631495 | 1,454189139 | 3,233851339 | 0,00122133 | 0,024539704 | The Moose: suction sampler vs FISH | NA |
| gene_1824013 | 21,8920955 | 3,035717843 | 0,963618978 | 3,150330071 | 0,001630861 | 0,029386306 | The Moose: suction sampler vs FISH | NA |
| gene_1824074 | 62,06571853 | 11,14058711 | 1,643054507 | 6,78041237 | 1,20E-11 | 8,83E-08 | The Moose: suction sampler vs FISH | NA |
| gene_1825122 | 84,05577889 | 3,149383669 | 0,729372434 | 4,317936244 | 1,57E-05 | 0,001421034 | The Moose: suction sampler vs FISH | NA |
| gene_1825634 | 5,252497053 | 5,42675245 | 1,698799666 | 3,194462865 | 0,001400913 | 0,026706306 | The Moose: suction sampler vs FISH | NA |
| gene_182610 | 16,28537813 | 2,788233108 | 0,91592459 | 3,044173219 | 0,002333208 | 0,036179697 | The Moose: suction sampler vs FISH | NA |
| gene_1826969 | 93,05537037 | 3,101027688 | 0,651163658 | 4,762286179 | 1,91E-06 | 0,000340394 | The Moose: suction sampler vs FISH | NA |
| gene_1826970 | 155,9751786 | 4,226377171 | 0,738664505 | 5,721646488 | 1,05E-08 | 9,48E-06 | The Moose: suction sampler vs FISH | NA |
| gene_1827210 | 9,132626929 | 3,826220768 | 1,254226379 | 3,05066201 | 0,002283375 | 0,035723316 | The Moose: suction sampler vs FISH | NA |
| gene_1827423 | 5,287540453 | 5,891492862 | 1,775321987 | 3,318548919 | 0,000904865 | 0,020533806 | The Moose: suction sampler vs FISH | NA |
| gene_1827722 | 72,26510984 | 2,281348075 | 0,784755579 | 2,907081054 | 0,003648187 | 0,047262814 | The Moose: suction sampler vs FISH | NA |
| gene_1828073 | 56,09096153 | 3,030228024 | 0,679715572 | 4,45808239 | 8,27E-06 | 0,000940188 | The Moose: suction sampler vs FISH | NA |
| gene_1828363 | 37,77440305 | 3,004669586 | 0,831875063 | 3,611924098 | 0,000303934 | 0,010285388 | The Moose: suction sampler vs FISH | NA |
| gene_1828364 | 49,53561307 | 3,447709416 | 0,862672657 | 3,996544215 | 6,43E-05 | 0,00371408 | The Moose: suction sampler vs FISH | NA |
| gene_1828365 | 14,38035531 | 3,197146075 | 1,089472349 | 2,934582119 | 0,003339972 | 0,04486178 | The Moose: suction sampler vs FISH | NA |
| gene_1830048 | 9,886120105 | 5,21959608 | 1,401908658 | 3,723206965 | 0,000196708 | 0,007725891 | The Moose: suction sampler vs FISH | NA |
| gene_1830697 | 51,24137603 | 3,391901731 | 0,939425692 | 3,61061206 | 0,000305475 | 0,010291574 | The Moose: suction sampler vs FISH | NA |
| gene_1831081 | 74,42221689 | 3,060828557 | 0,743283327 | 4,117983608 | 3,82E-05 | 0,002619796 | The Moose: suction sampler vs FISH | NA |
| gene_183240 | 44,01253992 | 10,73359214 | 2,501513608 | 4,290838997 | 1,78E-05 | 0,001540172 | The Moose: suction sampler vs FISH | NA |
| gene_1832648 | 143,391051 | 3,666905467 | 0,968761031 | 3,785149637 | 0,000153616 | 0,006570706 | The Moose: suction sampler vs FISH | NA |
| gene_1832837 | 43,90070599 | 3,001133372 | 0,967050889 | 3,103387223 | 0,001913191 | 0,032292895 | The Moose: suction sampler vs FISH | NA |
| gene_1833050 | 3,785313861 | 5,622475081 | 1,830503515 | 3,071545634 | 0,002129536 | 0,034442585 | The Moose: suction sampler vs FISH | NA |
| gene_1834080 | 12,01440928 | 5,139618457 | 1,347488453 | 3,81422078 | 0,000136613 | 0,006129239 | The Moose: suction sampler vs FISH | NA |
| gene_1834081 | 18,67523843 | 3,99188949 | 1,256466588 | 3,177075719 | 0,001487682 | 0,027759857 | The Moose: suction sampler vs FISH | NA |
| gene_1835122 | 13,98477768 | 5,408138033 | 1,662286479 | 3,253433208 | 0,001140195 | 0,023607465 | The Moose: suction sampler vs FISH | NA |
| gene_1836212 | 7,258786241 | 6,371172687 | 1,652699811 | 3,855009026 | 0,000115725 | 0,005496257 | The Moose: suction sampler vs FISH | NA |
| gene_1839075 | 5,326754032 | 5,818981946 | 1,715604205 | 3,391797438 | 0,000694358 | 0,017217113 | The Moose: suction sampler vs FISH | NA |
| gene_1841470 | 11,73548981 | 4,897489056 | 1,336422123 | 3,664627345 | 0,000247699 | 0,008988266 | The Moose: suction sampler vs FISH | NA |
| gene_1841701 | 15,75099156 | 3,679653377 | 1,094724767 | 3,36125891 | 0,000775881 | 0,018573843 | The Moose: suction sampler vs FISH | NA |
| gene_184377 | 5,788976942 | 5,551823143 | 1,626206079 | 3,41397269 | 0,00064023 | 0,016349683 | The Moose: suction sampler vs FISH | NA |
| gene_1844567 | 25,49307326 | 2,709851733 | 0,778646206 | 3,480209258 | 0,000501022 | 0,013991476 | The Moose: suction sampler vs FISH | NA |
| gene_1844717 | 6,118789391 | 5,887154141 | 1,746892151 | 3,37007304 | 0,000751483 | 0,018203378 | The Moose: suction sampler vs FISH | NA |
| gene_1844806 | 3,803199226 | 7,144741547 | 2,348865666 | 3,041783807 | 0,002351807 | 0,036348589 | The Moose: suction sampler vs FISH | NA |
| gene_1846626 | 15,64549658 | 3,716252273 | 1,069758974 | 3,473915494 | 0,000512923 | 0,014178245 | The Moose: suction sampler vs FISH | NA |
| gene_1848171 | 8,652058586 | 3,727571126 | 1,186833328 | 3,140770516 | 0,00168504 | 0,029972233 | The Moose: suction sampler vs FISH | NA |
| gene_1848172 | 15,71887586 | 3,429656191 | 1,11152321 | 3,085546177 | 0,002031787 | 0,033503497 | The Moose: suction sampler vs FISH | NA |
| gene_1848173 | 46,96752518 | 2,202327557 | 0,748636924 | 2,941783241 | 0,003263283 | 0,04423501 | The Moose: suction sampler vs FISH | NA |
| gene_1848470 | 21,15042434 | -2,720159065 | 0,849135421 | -3,203445525 | 0,001357937 | 0,026205737 | The Moose: suction sampler vs FISH | NA |
| gene_1848892 | 16,53700922 | 3,791690266 | 1,229765834 | 3,083262001 | 0,002047448 | 0,033673581 | The Moose: suction sampler vs FISH | NA |
| gene_1849014 | 64,24979068 | 4,169515221 | 0,903615655 | 4,614257398 | 3,95E-06 | 0,000555063 | The Moose: suction sampler vs FISH | NA |
| gene_1849332 | 54,29985961 | 3,040365524 | 1,052475896 | 2,888774495 | 0,003867463 | 0,048863897 | The Moose: suction sampler vs FISH | NA |
| gene_1850784 | 8,28933957 | 5,259343935 | 1,59356838 | 3,300356609 | 0,00096562 | 0,021271744 | The Moose: suction sampler vs FISH | NA |
| gene_1851305 | 30,79594329 | 3,41017791 | 0,805316272 | 4,234582147 | 2,29E-05 | 0,001829948 | The Moose: suction sampler vs FISH | NA |
| gene_1851656 | 7,723577458 | 4,980320125 | 1,514880939 | 3,287598382 | 0,001010459 | 0,021852142 | The Moose: suction sampler vs FISH | NA |
| gene_1852232 | 6,080014469 | 6,567247192 | 1,78756912 | 3,673842381 | 0,00023893 | 0,008795979 | The Moose: suction sampler vs FISH | NA |
| gene_1854580 | 110,0997983 | 2,42813387 | 0,834329772 | 2,910280745 | 0,003611042 | 0,046940354 | The Moose: suction sampler vs FISH | NA |
| gene_1855613 | 16,26557194 | 4,642312749 | 1,424403097 | 3,259128514 | 0,00111755 | 0,023337123 | The Moose: suction sampler vs FISH | NA |
| gene_1856565 | 42215,99348 | 2,398336131 | 0,770010501 | 3,114679773 | 0,001841447 | 0,031612271 | The Moose: suction sampler vs FISH | NA |
| gene_1857233 | 3,027472081 | 5,756932839 | 1,959400304 | 2,938109598 | 0,003302203 | 0,044566386 | The Moose: suction sampler vs FISH | NA |
| gene_1859439 | 3,516286745 | 6,039771953 | 2,096638658 | 2,880692832 | 0,003968021 | 0,049588452 | The Moose: suction sampler vs FISH | NA |
| gene_1859440 | 6,947168612 | 4,849563204 | 1,573422484 | 3,082174847 | 0,002054941 | 0,033748913 | The Moose: suction sampler vs FISH | NA |
| gene_185965 | 7,044160574 | -5,351968138 | 1,638244218 | -3,266892738 | 0,001087349 | 0,022967582 | The Moose: suction sampler vs FISH | NA |
| gene_1861004 | 27,34788142 | 2,69054123 | 0,879318985 | 3,059801137 | 0,00221484 | 0,035196395 | The Moose: suction sampler vs FISH | NA |
| gene_1861662 | 9,407128536 | 4,878082994 | 1,407741629 | 3,465183449 | 0,00052987 | 0,014483176 | The Moose: suction sampler vs FISH | NA |
| gene_1861675 | 56,43568353 | 5,279537208 | 1,024788348 | 5,151831807 | 2,58E-07 | 8,79E-05 | The Moose: suction sampler vs FISH | NA |
| gene_186463 | 28,5455931 | 3,279426725 | 0,886050675 | 3,701172874 | 0,000214605 | 0,008176582 | The Moose: suction sampler vs FISH | NA |
| gene_1865337 | 28,62787871 | 4,638025453 | 1,093163896 | 4,242753962 | 2,21E-05 | 0,001783958 | The Moose: suction sampler vs FISH | NA |
| gene_1866028 | 31,61830198 | 3,794375451 | 0,860427578 | 4,409871962 | 1,03E-05 | 0,001086773 | The Moose: suction sampler vs FISH | NA |
| gene_1866789 | 16,53096919 | 5,464771953 | 1,269513549 | 4,30461885 | 1,67E-05 | 0,001478388 | The Moose: suction sampler vs FISH | NA |
| gene_18677 | 16,30722928 | 3,788688562 | 1,063064752 | 3,563930189 | 0,000365343 | 0,011490192 | The Moose: suction sampler vs FISH | NA |
| gene_1868302 | 85,06554223 | 3,451067717 | 0,885082253 | 3,899149153 | 9,65E-05 | 0,004889477 | The Moose: suction sampler vs FISH | NA |

| | | | | | | | | |
|---|---|---|---|---|---|---|---|---|
| gene_1868450 | 121,5696252 | 2,194576756 | 0,717600787 | 3,058213978 | 0,002226605 | 0,035228466 | The Moose: suction sampler vs FISH | NA |
| gene_1868782 | 11,20490766 | 3,752180585 | 1,158795136 | 3,238001669 | 0,001203701 | 0,024381088 | The Moose: suction sampler vs FISH | NA |
| gene_1869516 | 37,62534531 | 6,01898218 | 1,127446159 | 5,338598328 | 9,37E-08 | 4,59E-05 | The Moose: suction sampler vs FISH | NA |
| gene_1870152 | 34,7088517 | 3,241731454 | 0,873366523 | 3,711765185 | 0,000205819 | 0,007972768 | The Moose: suction sampler vs FISH | NA |
| gene_187046 | 4,147611667 | 6,03081853 | 1,933047452 | 3,119850226 | 0,00180943 | 0,031232338 | The Moose: suction sampler vs FISH | NA |
| gene_1871364 | 9,199379071 | 6,144981771 | 1,602117344 | 3,835537886 | 0,00012529 | 0,005797411 | The Moose: suction sampler vs FISH | NA |
| gene_1871716 | 31,45486248 | 3,024221583 | 0,783045138 | 3,862129317 | 0,000112403 | 0,005399242 | The Moose: suction sampler vs FISH | NA |
| gene_1872012 | 117,5502845 | 12,23626005 | 3,416216571 | 3,581816257 | 0,000341214 | 0,011021805 | The Moose: suction sampler vs FISH | NA |
| gene_1872248 | 9,467024275 | 3,235636614 | 1,085957889 | 2,979523099 | 0,002886975 | 0,041155639 | The Moose: suction sampler vs FISH | NA |
| gene_1873345 | 4,422649821 | 6,937459259 | 1,88263907 | 3,684965095 | 0,000228734 | 0,008540604 | The Moose: suction sampler vs FISH | NA |
| gene_1873402 | 4,74361363 | 4,775381449 | 1,620017974 | 2,947733621 | 0,003201128 | 0,043776571 | The Moose: suction sampler vs FISH | NA |
| gene_1873780 | 5,535596603 | 5,948164277 | 1,838547983 | 3,235251043 | 0,001215358 | 0,024511793 | The Moose: suction sampler vs FISH | NA |
| gene_1874856 | 7,05707975 | -4,33158141 | 1,339438618 | -3,233878248 | 0,001221215 | 0,024539704 | The Moose: suction sampler vs FISH | NA |
| gene_1875165 | 13,40306701 | 3,047644477 | 0,981978687 | 3,103574974 | 0,001911978 | 0,032281658 | The Moose: suction sampler vs FISH | NA |
| gene_1875977 | 6,606713709 | 5,71575499 | 1,95196736 | 2,928202135 | 0,003409283 | 0,045344949 | The Moose: suction sampler vs FISH | NA |
| gene_1876054 | 47,050351 | 3,613682748 | 1,21174985 | 2,982201935 | 0,002861832 | 0,040972169 | The Moose: suction sampler vs FISH | NA |
| gene_1876529 | 21,09483123 | 3,716061619 | 1,049062111 | 3,542270359 | 0,000396699 | 0,01214993 | The Moose: suction sampler vs FISH | NA |
| gene_1877424 | 13,07255167 | 3,370967585 | 1,107831146 | 3,042853233 | 0,002343466 | 0,036250181 | The Moose: suction sampler vs FISH | NA |
| gene_1877789 | 45,58682358 | 3,662650885 | 0,849831329 | 4,309856275 | 1,63E-05 | 0,001453599 | The Moose: suction sampler vs FISH | NA |
| gene_1877992 | 26,78222498 | 4,413696003 | 1,10955994 | 3,977879738 | 6,95E-05 | 0,00391627 | The Moose: suction sampler vs FISH | NA |
| gene_1878161 | 133,8535955 | 12,43393853 | 4,247278111 | 2,9275075 | 0,003416908 | 0,045400743 | The Moose: suction sampler vs FISH | NA |
| gene_1879195 | 35,068916 | 2,791612897 | 0,935627049 | 2,983681266 | 0,002848033 | 0,040865094 | The Moose: suction sampler vs FISH | NA |
| gene_1879869 | 24,03638889 | 3,244010345 | 0,941493567 | 3,445600116 | 0,000569793 | 0,015195511 | The Moose: suction sampler vs FISH | NA |
| gene_1880367 | 15,71351833 | 4,110677611 | 1,026820734 | 4,003306004 | 6,25E-05 | 0,003643865 | The Moose: suction sampler vs FISH | NA |
| gene_1881279 | 7,9510737 | 4,750637963 | 1,407770677 | 3,374582269 | 0,000739278 | 0,018002747 | The Moose: suction sampler vs FISH | NA |
| gene_1881318 | 40,03512646 | 5,249894473 | 1,18374625 | 4,43498298 | 9,21E-06 | 0,001006367 | The Moose: suction sampler vs FISH | NA |
| gene_1881908 | 45,28640104 | -2,162738696 | 0,746175258 | -2,898432603 | 0,003750329 | 0,0480412 | The Moose: suction sampler vs FISH | NA |
| gene_188398 | 10,2384286 | 3,802751543 | 1,23126383 | 3,088494482 | 0,002011734 | 0,033284935 | The Moose: suction sampler vs FISH | NA |
| gene_1884481 | 22,32228728 | 6,063275547 | 1,394452403 | 4,348140914 | 1,37E-05 | 0,001305305 | The Moose: suction sampler vs FISH | NA |
| gene_1885771 | 156,8151258 | -2,283059738 | 0,690145907 | -3,308082705 | 0,000939371 | 0,020932062 | The Moose: suction sampler vs FISH | NA |
| gene_1886568 | 12,78168713 | 5,01580343 | 1,331217916 | 3,767830474 | 0,000164672 | 0,006903179 | The Moose: suction sampler vs FISH | NA |
| gene_1887020 | 5,445616338 | 7,373471855 | 1,790829993 | 4,117348875 | 3,83E-05 | 0,002623319 | The Moose: suction sampler vs FISH | NA |
| gene_1887525 | 16,89112401 | 3,56425188 | 1,174639988 | 3,034335555 | 0,002410661 | 0,036916394 | The Moose: suction sampler vs FISH | NA |
| gene_1887854 | 9,913437525 | 5,761356659 | 1,495327188 | 3,852907046 | 0,000116724 | 0,005530821 | The Moose: suction sampler vs FISH | NA |
| gene_1887874 | 26,61263187 | 2,660920761 | 0,767292676 | 3,467934522 | 0,000524475 | 0,014377268 | The Moose: suction sampler vs FISH | NA |
| gene_1887875 | 31,40004571 | 4,153262589 | 0,753430461 | 5,512469699 | 3,54E-08 | 2,26E-05 | The Moose: suction sampler vs FISH | NA |
| gene_1888822 | 8,32780325 | 4,803728118 | 1,670348231 | 2,87588422 | 0,004028975 | 0,049994565 | The Moose: suction sampler vs FISH | NA |
| gene_1889317 | 7,510343337 | 4,920826191 | 1,532322953 | 3,211350572 | 0,001321127 | 0,025815906 | The Moose: suction sampler vs FISH | NA |
| gene_1890519 | 4,137211975 | 5,872826291 | 1,998602757 | 2,938466021 | 0,003298408 | 0,044556301 | The Moose: suction sampler vs FISH | NA |
| gene_1890580 | 7,477491177 | 3,724641769 | 1,224593801 | 3,041532437 | 0,002353772 | 0,036361491 | The Moose: suction sampler vs FISH | NA |
| gene_1890591 | 6,50305267 | 5,680699442 | 1,928190996 | 2,946129015 | 0,003217782 | 0,04392122 | The Moose: suction sampler vs FISH | NA |
| gene_1890648 | 21,29858245 | 2,991300243 | 0,909454829 | 3,289113594 | 0,001005035 | 0,021789908 | The Moose: suction sampler vs FISH | NA |
| gene_1891027 | 4,612846159 | 5,419940176 | 1,841137966 | 2,943799039 | 0,003242104 | 0,04410379 | The Moose: suction sampler vs FISH | NA |
| gene_1891194 | 6,60124966 | 6,78959291 | 1,769996769 | 3,8359352 | 0,000125087 | 0,005797411 | The Moose: suction sampler vs FISH | NA |
| gene_1891297 | 49,89665968 | 2,744759429 | 0,896410248 | 3,061945615 | 0,002199034 | 0,035049491 | The Moose: suction sampler vs FISH | NA |
| gene_1891298 | 39,40016663 | 2,894480924 | 0,826993349 | 3,500005083 | 0,000465249 | 0,013354236 | The Moose: suction sampler vs FISH | NA |
| gene_1891299 | 21,64165855 | 4,058022017 | 0,99508553 | 4,078063536 | 4,54E-05 | 0,002967358 | The Moose: suction sampler vs FISH | NA |
| gene_1891300 | 26,63413641 | 3,797265519 | 0,926365338 | 4,099101471 | 4,15E-05 | 0,002774302 | The Moose: suction sampler vs FISH | NA |
| gene_1892453 | 66,49411117 | 2,9081428 | 0,744365379 | 3,906875415 | 9,35E-05 | 0,004794295 | The Moose: suction sampler vs FISH | NA |
| gene_1892470 | 26,32496055 | 3,323131535 | 1,122240789 | 2,961157327 | 0,003064853 | 0,042713122 | The Moose: suction sampler vs FISH | NA |
| gene_1892488 | 12,1045628 | 4,618587494 | 1,528126689 | 3,022385203 | 0,002507912 | 0,037875787 | The Moose: suction sampler vs FISH | NA |
| gene_1894471 | 6,754589009 | 5,928819782 | 1,669126539 | 3,552049315 | 0,000382243 | 0,011850508 | The Moose: suction sampler vs FISH | NA |
| gene_1894474 | 4,144580528 | 6,037373272 | 1,881781753 | 3,208328097 | 0,001335091 | 0,025926958 | The Moose: suction sampler vs FISH | NA |
| gene_1896190 | 17,89698056 | 3,133662773 | 0,907531773 | 3,452951032 | 0,00055449 | 0,014890125 | The Moose: suction sampler vs FISH | NA |
| gene_1896598 | 4,035918643 | 6,22857049 | 1,771991989 | 3,515010524 | 0,000439737 | 0,012913873 | The Moose: suction sampler vs FISH | NA |
| gene_1896606 | 4,706909186 | 6,563848291 | 1,981348557 | 3,312818569 | 0,000923609 | 0,020731798 | The Moose: suction sampler vs FISH | NA |
| gene_189670 | 25,58536504 | 3,371011664 | 1,112283778 | 3,030711883 | 0,002439779 | 0,03717118 | The Moose: suction sampler vs FISH | NA |
| gene_1897097 | 9,754269816 | 3,944661185 | 1,270551548 | 3,104684095 | 0,001904823 | 0,032241111 | The Moose: suction sampler vs FISH | NA |
| gene_1897192 | 271,8775737 | 6,409758313 | 1,624546466 | 3,945567853 | 7,96E-05 | 0,004280407 | The Moose: suction sampler vs FISH | NA |
| gene_1899688 | 11,00200421 | 3,317463604 | 1,08936287 | 3,045324653 | 0,002324293 | 0,036082789 | The Moose: suction sampler vs FISH | NA |
| gene_1901705 | 74,56290621 | 11,58797905 | 3,588450325 | 3,229243267 | 0,001241183 | 0,024791586 | The Moose: suction sampler vs FISH | NA |
| gene_1903021 | 3,721500933 | 5,509839047 | 1,857830236 | 2,965738709 | 0,003019569 | 0,042355281 | The Moose: suction sampler vs FISH | NA |
| gene_1903053 | 9,113949879 | 7,431903891 | 1,586727316 | 4,683794007 | 2,82E-06 | 0,000435433 | The Moose: suction sampler vs FISH | NA |
| gene_1903596 | 28,19502803 | 2,843777195 | 0,871192545 | 3,264234998 | 0,001097601 | 0,023104986 | The Moose: suction sampler vs FISH | NA |
| gene_1904905 | 21,30982118 | 3,312701823 | 0,928951829 | 3,56606416 | 0,000362383 | 0,011447396 | The Moose: suction sampler vs FISH | NA |

| | | | | | | | | |
|---|---|---|---|---|---|---|---|---|
| gene_1904955 | 17,90853579 | 3,566266496 | 1,104105517 | 3,230005141 | 0,00123788 | 0,024752543 | The Moose: suction sampler vs FISH | NA |
| gene_1905062 | 74,33744091 | 3,611180931 | 1,039502786 | 3,473950218 | 0,000512856 | 0,014178245 | The Moose: suction sampler vs FISH | NA |
| gene_1905336 | 24,13563915 | 7,520891296 | 2,403228186 | 3,129495293 | 0,001751069 | 0,030666956 | The Moose: suction sampler vs FISH | NA |
| gene_1905614 | 52,87226878 | 2,583342644 | 0,889182259 | 2,905301606 | 0,003668994 | 0,047425091 | The Moose: suction sampler vs FISH | NA |
| gene_190567 | 22,78104146 | 4,34161343 | 0,951435301 | 4,563225082 | 5,04E-06 | 0,000660549 | The Moose: suction sampler vs FISH | NA |
| gene_1905907 | 17,40117997 | 6,247466897 | 1,323923046 | 4,718904861 | 2,37E-06 | 0,000390473 | The Moose: suction sampler vs FISH | NA |
| gene_1906227 | 114,0419353 | 3,777017884 | 0,891482892 | 4,236781119 | 2,27E-05 | 0,001822039 | The Moose: suction sampler vs FISH | NA |
| gene_1906779 | 49,58132904 | 3,84624517 | 0,990761356 | 3,882110609 | 0,000103554 | 0,005101251 | The Moose: suction sampler vs FISH | NA |
| gene_1907487 | 7,512406779 | 3,890381802 | 1,307659184 | 2,975073207 | 0,002929186 | 0,041507639 | The Moose: suction sampler vs FISH | NA |
| gene_1908078 | 18,49442203 | 4,694405756 | 1,099415591 | 4,269910118 | 1,96E-05 | 0,001628379 | The Moose: suction sampler vs FISH | NA |
| gene_190924 | 14,48114102 | 3,278327063 | 1,116173564 | 2,937112263 | 0,003312842 | 0,044685335 | The Moose: suction sampler vs FISH | NA |
| gene_1909316 | 6,867346781 | 5,17715265 | 1,477308798 | 3,504448534 | 0,000457554 | 0,013225129 | The Moose: suction sampler vs FISH | NA |
| gene_1909696 | 4,535254844 | 6,167252487 | 1,725393732 | 3,574402974 | 0,000351028 | 0,011199034 | The Moose: suction sampler vs FISH | NA |
| gene_1910286 | 8,597130743 | 5,538189009 | 1,558765703 | 3,552932297 | 0,000380962 | 0,011820782 | The Moose: suction sampler vs FISH | NA |
| gene_1910364 | 8,360758185 | 6,885256719 | 1,793447359 | 3,83911838 | 0,000123477 | 0,005759871 | The Moose: suction sampler vs FISH | NA |
| gene_1910568 | 18,53078807 | 2,872450654 | 0,952953991 | 3,014259536 | 0,002576073 | 0,038502178 | The Moose: suction sampler vs FISH | NA |
| gene_1911281 | 27,31324962 | 2,854513305 | 0,819390063 | 3,483705055 | 0,000494524 | 0,01387331 | The Moose: suction sampler vs FISH | NA |
| gene_1911514 | 32,34353791 | 3,815852411 | 0,972108293 | 3,925336751 | 8,66E-05 | 0,0045403 | The Moose: suction sampler vs FISH | NA |
| gene_1911515 | 18,03382961 | 5,299991017 | 1,281856455 | 4,134621313 | 3,56E-05 | 0,002482193 | The Moose: suction sampler vs FISH | NA |
| gene_1911597 | 3,438169855 | 5,971409457 | 2,042773248 | 2,923187614 | 0,003464677 | 0,04579971 | The Moose: suction sampler vs FISH | NA |
| gene_1911672 | 16,21175286 | 3,097224275 | 1,015640093 | 3,049529351 | 0,002292002 | 0,035778132 | The Moose: suction sampler vs FISH | NA |
| gene_1912040 | 7,153740554 | 3,799683478 | 1,316954044 | 2,885205825 | 0,003911578 | 0,049132477 | The Moose: suction sampler vs FISH | NA |
| gene_1913148 | 514,8286935 | 7,826899389 | 1,482658952 | 5,278961409 | 1,30E-07 | 5,46E-05 | The Moose: suction sampler vs FISH | NA |
| gene_1913471 | 12,79892304 | 5,501736865 | 1,480340775 | 3,716534029 | 0,000201974 | 0,00787368 | The Moose: suction sampler vs FISH | NA |
| gene_1913980 | 21,44669932 | 3,312882174 | 0,96010733 | 3,450533155 | 0,00055948 | 0,014980329 | The Moose: suction sampler vs FISH | NA |
| gene_1914701 | 10,80931925 | 2,884516295 | 0,995878775 | 2,896453231 | 0,003774069 | 0,048199831 | The Moose: suction sampler vs FISH | NA |
| gene_1915794 | 14,17279867 | 4,254359412 | 1,084364182 | 3,923367704 | 8,73E-05 | 0,00457106 | The Moose: suction sampler vs FISH | NA |
| gene_1916656 | 36,37941007 | 4,808610203 | 0,902521547 | 5,327972744 | 9,93E-08 | 4,64E-05 | The Moose: suction sampler vs FISH | NA |
| gene_1916657 | 34,77849703 | 3,118800123 | 0,877062991 | 3,555959098 | 0,000376603 | 0,011730103 | The Moose: suction sampler vs FISH | NA |
| gene_1917082 | 13,66544245 | 9,031844191 | 2,940972304 | 3,071040206 | 0,002133144 | 0,034462996 | The Moose: suction sampler vs FISH | NA |
| gene_1917914 | 47,92932261 | 2,804401153 | 0,698302071 | 4,016028689 | 5,92E-05 | 0,003522685 | The Moose: suction sampler vs FISH | NA |
| gene_1918314 | 10,13489524 | 5,688407666 | 1,674505829 | 3,397066506 | 0,000681124 | 0,017003793 | The Moose: suction sampler vs FISH | NA |
| gene_1918316 | 6,80544968 | 7,814191366 | 1,943678745 | 4,020310139 | 5,81E-05 | 0,003484685 | The Moose: suction sampler vs FISH | NA |
| gene_1922261 | 10,91896107 | 5,534239675 | 1,344489305 | 4,11623927 | 3,85E-05 | 0,00263352 | The Moose: suction sampler vs FISH | NA |
| gene_19226 | 13,3503602 | 3,303099712 | 0,943592137 | 3,500558752 | 0,000464284 | 0,013354236 | The Moose: suction sampler vs FISH | NA |
| gene_1924044 | 4,739472457 | 5,352925588 | 1,712175712 | 3,126387993 | 0,001769679 | 0,030814148 | The Moose: suction sampler vs FISH | NA |
| gene_1924265 | 7,750472221 | 7,349434589 | 1,746790225 | 4,20739393 | 2,58E-05 | 0,002007468 | The Moose: suction sampler vs FISH | NA |
| gene_1925014 | 17,44986843 | 3,901910985 | 1,075084742 | 3,629398531 | 0,000284082 | 0,00984091 | The Moose: suction sampler vs FISH | NA |
| gene_1926777 | 37,48787511 | 4,053855098 | 0,830665371 | 4,880250506 | 1,06E-06 | 0,000230908 | The Moose: suction sampler vs FISH | NA |
| gene_1926778 | 19,65013333 | 3,617571344 | 0,924735076 | 3,912008356 | 9,15E-05 | 0,004727531 | The Moose: suction sampler vs FISH | NA |
| gene_1927098 | 30,49882983 | 3,331690193 | 0,989443116 | 3,36723773 | 0,000759252 | 0,018319076 | The Moose: suction sampler vs FISH | NA |
| gene_1928492 | 9,529726866 | 3,848851536 | 1,206548591 | 3,189968118 | 0,001422885 | 0,026988521 | The Moose: suction sampler vs FISH | NA |
| gene_1928607 | 5,424606643 | 4,995492137 | 1,704215037 | 2,93125693 | 0,003375934 | 0,045076057 | The Moose: suction sampler vs FISH | NA |
| gene_1928637 | 4,399118134 | 6,631424586 | 1,986939645 | 3,337506805 | 0,000845337 | 0,019691006 | The Moose: suction sampler vs FISH | NA |
| gene_1929622 | 4,74586547 | 5,6033136 | 1,847577809 | 3,03278897 | 0,00242305 | 0,0370289 | The Moose: suction sampler vs FISH | NA |
| gene_1929757 | 7,159562531 | 7,075164563 | 1,723782082 | 4,104442571 | 4,05E-05 | 0,00273088 | The Moose: suction sampler vs FISH | NA |
| gene_193034 | 23,96847807 | 3,234193648 | 0,8992877 | 3,596394846 | 0,000322658 | 0,010660194 | The Moose: suction sampler vs FISH | NA |
| gene_193035 | 52,50100253 | 3,933193652 | 0,82390335 | 4,77385321 | 1,81E-06 | 0,000326946 | The Moose: suction sampler vs FISH | NA |
| gene_1931386 | 12,34122 | 4,637452055 | 1,083203283 | 4,281238923 | 1,86E-05 | 0,001574414 | The Moose: suction sampler vs FISH | NA |
| gene_1933990 | 6,933496479 | 6,804279353 | 1,724767838 | 3,945040718 | 7,98E-05 | 0,004286696 | The Moose: suction sampler vs FISH | NA |
| gene_1933991 | 8,501533289 | 4,639158328 | 1,266991221 | 3,661555227 | 0,000250689 | 0,009047587 | The Moose: suction sampler vs FISH | NA |
| gene_1935534 | 12,63306989 | 3,585316861 | 1,077658464 | 3,326950959 | 0,000878018 | 0,020152886 | The Moose: suction sampler vs FISH | NA |
| gene_1935607 | 4,739608348 | 6,796526632 | 1,884506856 | 3,606527941 | 0,000310321 | 0,010407105 | The Moose: suction sampler vs FISH | NA |
| gene_1939905 | 34,06591481 | 2,812321329 | 0,858386359 | 3,276288467 | 0,001051811 | 0,02250881 | The Moose: suction sampler vs FISH | NA |
| gene_1940712 | 16,06758924 | 2,822091219 | 0,879425471 | 3,209016925 | 0,001331897 | 0,025899163 | The Moose: suction sampler vs FISH | NA |
| gene_1940751 | 8,708063942 | 4,925912626 | 1,488517026 | 3,309275299 | 0,000935378 | 0,020887457 | The Moose: suction sampler vs FISH | NA |
| gene_1941138 | 21,06843524 | 4,564439743 | 1,130355521 | 4,038056753 | 5,39E-05 | 0,003306916 | The Moose: suction sampler vs FISH | NA |
| gene_1942033 | 52,30172577 | 2,355235165 | 0,782844334 | 3,008561295 | 0,002624878 | 0,038881355 | The Moose: suction sampler vs FISH | NA |
| gene_1942228 | 4,501394573 | 7,046380848 | 2,317116993 | 3,041012115 | 0,002357843 | 0,03638717 | The Moose: suction sampler vs FISH | NA |
| gene_1943078 | 16,22546978 | 3,677654164 | 1,037078253 | 3,546168434 | 0,000390876 | 0,012011677 | The Moose: suction sampler vs FISH | NA |
| gene_1943332 | 8,619040193 | 4,549362957 | 1,324840922 | 3,433893747 | 0,000594977 | 0,015556466 | The Moose: suction sampler vs FISH | NA |
| gene_1943695 | 6,725155696 | 3,685798121 | 1,235572146 | 2,983069935 | 0,002853728 | 0,040907188 | The Moose: suction sampler vs FISH | NA |
| gene_1944450 | 3,844401988 | 5,858483043 | 1,92776073 | 3,039009433 | 0,002373574 | 0,036523603 | The Moose: suction sampler vs FISH | NA |
| gene_1945307 | 8,705121176 | 3,642071748 | 1,246959723 | 2,920761337 | 0,003491772 | 0,046025341 | The Moose: suction sampler vs FISH | NA |
| gene_194541 | 23,10534902 | 3,466266343 | 1,124100378 | 3,083591476 | 0,002045182 | 0,033671466 | The Moose: suction sampler vs FISH | NA |

| | | | | | | | | |
|---|---|---|---|---|---|---|---|---|
| gene_1945996 | 10,93339692 | 4,548758199 | 1,468666333 | 3,097203291 | 0,001953559 | 0,032684422 | The Moose: suction sampler vs FISH | NA |
| gene_1946802 | 4,939080938 | 6,174550949 | 1,855216602 | 3,328210271 | 0,000874059 | 0,020098697 | The Moose: suction sampler vs FISH | NA |
| gene_1946952 | 7,98992423 | 7,63915543 | 1,765255855 | 4,327506072 | 1,51E-05 | 0,001382776 | The Moose: suction sampler vs FISH | NA |
| gene_1947150 | 4,933250805 | 5,82326769 | 2,023197979 | 2,878249065 | 0,003998893 | 0,049803793 | The Moose: suction sampler vs FISH | NA |
| gene_1947151 | 9,473324311 | 3,972332433 | 1,287728383 | 3,084759553 | 0,002037168 | 0,033577151 | The Moose: suction sampler vs FISH | NA |
| gene_1947152 | 5,403986002 | 5,520692239 | 1,714055108 | 3,220837074 | 0,001278168 | 0,025289765 | The Moose: suction sampler vs FISH | NA |
| gene_194791 | 5,700766739 | 4,265827877 | 1,40145363 | 3,043859451 | 0,002335642 | 0,036197774 | The Moose: suction sampler vs FISH | NA |
| gene_1947917 | 5,626319649 | 6,934957085 | 1,753091482 | 3,955844379 | 7,63E-05 | 0,004186299 | The Moose: suction sampler vs FISH | NA |
| gene_1949712 | 8,519145103 | 4,020061776 | 1,237811304 | 3,247717777 | 0,001163346 | 0,023888006 | The Moose: suction sampler vs FISH | NA |
| gene_1950302 | 62,67228572 | 6,624774498 | 1,200720913 | 5,51733082 | 3,44E-08 | 2,22E-05 | The Moose: suction sampler vs FISH | NA |
| gene_1951396 | 8,527828008 | 5,046332119 | 1,464772202 | 3,445130999 | 0,000570783 | 0,015216383 | The Moose: suction sampler vs FISH | NA |
| gene_1951905 | 23,74080947 | 4,498803508 | 1,068588767 | 4,210041922 | 2,55E-05 | 0,001994917 | The Moose: suction sampler vs FISH | NA |
| gene_1952298 | 2043,981342 | 3,240358111 | 0,802633451 | 4,037158068 | 5,41E-05 | 0,003307895 | The Moose: suction sampler vs FISH | NA |
| gene_1952850 | 16,70788699 | 3,189461715 | 1,076397343 | 2,963089547 | 0,00304568 | 0,042526603 | The Moose: suction sampler vs FISH | NA |
| gene_19897 | 8,429695807 | 4,088978299 | 1,308499406 | 3,124937068 | 0,001778431 | 0,030878692 | The Moose: suction sampler vs FISH | NA |
| gene_20044 | 145,837746 | 5,332371391 | 0,875085906 | 6,093540481 | 1,10E-09 | 1,93E-06 | The Moose: suction sampler vs FISH | NA |
| gene_201981 | 11,32305335 | 4,555428292 | 1,234238191 | 3,690882622 | 0,000223477 | 0,00839984 | The Moose: suction sampler vs FISH | NA |
| gene_202519 | 59,4147584 | 2,404454423 | 0,747310528 | 3,217476982 | 0,001293234 | 0,025471058 | The Moose: suction sampler vs FISH | NA |
| gene_202958 | 32,69703857 | 2,625960586 | 0,909804502 | 2,886291045 | 0,003898115 | 0,049048664 | The Moose: suction sampler vs FISH | NA |
| gene_203223 | 8,676502134 | 3,707099597 | 1,134160187 | 3,268585549 | 0,001080865 | 0,022864092 | The Moose: suction sampler vs FISH | NA |
| gene_203874 | 15,52048138 | 3,202705802 | 1,058556981 | 3,025539353 | 0,002481901 | 0,037638493 | The Moose: suction sampler vs FISH | NA |
| gene_204489 | 98,68419752 | 2,379960181 | 0,805532091 | 2,954519389 | 0,003131565 | 0,043265194 | The Moose: suction sampler vs FISH | NA |
| gene_204872 | 45,58291158 | 2,872629117 | 0,7285577 | 3,942898574 | 8,05E-05 | 0,004312561 | The Moose: suction sampler vs FISH | NA |
| gene_205608 | 6,611067807 | 6,579368192 | 1,617692906 | 4,067130522 | 4,76E-05 | 0,003066367 | The Moose: suction sampler vs FISH | NA |
| gene_208374 | 7,784266978 | -5,726875543 | 1,591304231 | -3,598856478 | 0,00031962 | 0,010599955 | The Moose: suction sampler vs FISH | NA |
| gene_208526 | 33,26132795 | 2,968834269 | 0,985195892 | 3,013445643 | 0,002582993 | 0,03856156 | The Moose: suction sampler vs FISH | NA |
| gene_208584 | 11,72284522 | 3,43954811 | 1,06293082 | 3,235909663 | 0,001212558 | 0,024471718 | The Moose: suction sampler vs FISH | NA |
| gene_209428 | 52,83694927 | -2,98784616 | 1,038494243 | -2,877094583 | 0,004013553 | 0,049896274 | The Moose: suction sampler vs FISH | NA |
| gene_209431 | 15,07305455 | -6,569193566 | 1,578521936 | -4,161610565 | 3,16E-05 | 0,002284654 | The Moose: suction sampler vs FISH | NA |
| gene_209432 | 23,42499785 | -5,513955148 | 1,375858748 | -4,007646247 | 6,13E-05 | 0,003606207 | The Moose: suction sampler vs FISH | NA |
| gene_210822 | 63,88947333 | 3,314878876 | 0,71641922 | 4,627009971 | 3,71E-06 | 0,000533257 | The Moose: suction sampler vs FISH | NA |
| gene_211125 | 25,11292861 | 2,746543676 | 0,921038408 | 2,982007757 | 0,002863647 | 0,040982187 | The Moose: suction sampler vs FISH | NA |
| gene_21146 | 977,0016036 | -2,009046265 | 0,533552296 | -3,765415841 | 0,000166272 | 0,00695436 | The Moose: suction sampler vs FISH | NA |
| gene_212592 | 7,225241295 | 5,850926089 | 1,794042515 | 3,261308492 | 0,001108993 | 0,023254434 | The Moose: suction sampler vs FISH | NA |
| gene_212743 | 52,18406532 | 5,166101391 | 1,123052406 | 4,600053712 | 4,22E-06 | 0,000585318 | The Moose: suction sampler vs FISH | NA |
| gene_213343 | 13,12273744 | -4,850606517 | 1,49148717 | -3,252194599 | 0,001145176 | 0,023685565 | The Moose: suction sampler vs FISH | NA |
| gene_213873 | 64,49851674 | 4,981192386 | 0,838192163 | 5,942780914 | 2,80E-09 | 3,55E-06 | The Moose: suction sampler vs FISH | NA |
| gene_214145 | 11,41607392 | 3,075653341 | 0,916381392 | 3,356302701 | 0,000789921 | 0,018763169 | The Moose: suction sampler vs FISH | NA |
| gene_214581 | 1180,633238 | 5,625146064 | 0,912871623 | 6,162034096 | 7,18E-10 | 1,35E-06 | The Moose: suction sampler vs FISH | NA |
| gene_21508 | 10,22231682 | 4,363177454 | 1,295549054 | 3,36782111 | 0,000757647 | 0,018294693 | The Moose: suction sampler vs FISH | NA |
| gene_216695 | 3,523864886 | 6,456842001 | 2,219059938 | 2,909719512 | 0,003617533 | 0,046991453 | The Moose: suction sampler vs FISH | NA |
| gene_217942 | 4,886133307 | 4,899870947 | 1,703069652 | 2,877081945 | 0,004013714 | 0,049896274 | The Moose: suction sampler vs FISH | NA |
| gene_218896 | 124,0966859 | 3,749609059 | 0,816888433 | 4,590111585 | 4,43E-06 | 0,00060703 | The Moose: suction sampler vs FISH | NA |
| gene_219491 | 5,810770211 | 5,893240805 | 1,996159877 | 2,952288979 | 0,003154276 | 0,043383106 | The Moose: suction sampler vs FISH | NA |
| gene_219741 | 12,34779014 | 3,866913473 | 1,132998249 | 3,41299157 | 0,000642539 | 0,01638018 | The Moose: suction sampler vs FISH | NA |
| gene_21990 | 20,62129655 | 2,820223462 | 0,878220285 | 3,21129392 | 0,001321387 | 0,025815906 | The Moose: suction sampler vs FISH | NA |
| gene_21992 | 10,8162824 | 4,204397287 | 1,236733539 | 3,399598341 | 0,000674849 | 0,016904601 | The Moose: suction sampler vs FISH | NA |
| gene_21994 | 12,44253852 | 3,342285536 | 0,983123568 | 3,399659661 | 0,000674698 | 0,016904601 | The Moose: suction sampler vs FISH | NA |
| gene_21995 | 6,857068251 | 3,826805087 | 1,238215808 | 3,090580061 | 0,001997659 | 0,03314914 | The Moose: suction sampler vs FISH | NA |
| gene_222472 | 22,34567468 | 4,452564335 | 0,958640933 | 4,644663275 | 3,41E-06 | 0,000499355 | The Moose: suction sampler vs FISH | NA |
| gene_223157 | 117,2497849 | 2,606861277 | 0,692125987 | 3,766454843 | 0,000165582 | 0,006929442 | The Moose: suction sampler vs FISH | NA |
| gene_22342 | 14,81044272 | 5,418867649 | 1,243332098 | 4,358342924 | 1,31E-05 | 0,001269942 | The Moose: suction sampler vs FISH | NA |
| gene_224230 | 27,25246554 | 2,818323931 | 0,974639201 | 2,891658707 | 0,00383214 | 0,048609933 | The Moose: suction sampler vs FISH | NA |
| gene_226326 | 11,64940726 | 4,992825262 | 1,290525657 | 3,868830686 | 0,000109359 | 0,005291066 | The Moose: suction sampler vs FISH | NA |
| gene_226604 | 86,49667878 | 2,793585466 | 0,889888629 | 3,139252909 | 0,001693792 | 0,030059907 | The Moose: suction sampler vs FISH | NA |
| gene_226605 | 138,8863907 | 2,839347006 | 0,830839958 | 3,417441565 | 0,000632127 | 0,016199072 | The Moose: suction sampler vs FISH | NA |
| gene_226606 | 193,2509432 | 3,414074258 | 0,855745601 | 3,989590191 | 6,62E-05 | 0,003777113 | The Moose: suction sampler vs FISH | NA |
| gene_22872 | 4,8299712 | 5,367851151 | 1,751148781 | 3,065331289 | 0,00217429 | 0,03482895 | The Moose: suction sampler vs FISH | NA |
| gene_230910 | 7,160119858 | 5,131273071 | 1,592738575 | 3,221666853 | 0,001274472 | 0,025270955 | The Moose: suction sampler vs FISH | NA |
| gene_231386 | 26,00833519 | 2,376913317 | 0,786647508 | 3,021573568 | 0,002514645 | 0,037931428 | The Moose: suction sampler vs FISH | NA |
| gene_232867 | 4,340205196 | 5,736704574 | 1,766745309 | 3,247046727 | 0,001166092 | 0,023920816 | The Moose: suction sampler vs FISH | NA |
| gene_234916 | 251,154821 | -2,50268212 | 0,809833756 | -3,090365275 | 0,001999105 | 0,033158141 | The Moose: suction sampler vs FISH | NA |
| gene_235806 | 28,57356932 | 3,10485452 | 0,982135507 | 3,16133008 | 0,001570504 | 0,02868807 | The Moose: suction sampler vs FISH | NA |
| gene_236162 | 33,50070485 | 10,2670466 | 3,113603738 | 3,297480177 | 0,000975566 | 0,021388184 | The Moose: suction sampler vs FISH | NA |
| gene_236743 | 83,57177189 | 3,020434298 | 0,744883724 | 4,054907097 | 5,02E-05 | 0,003168826 | The Moose: suction sampler vs FISH | NA |

| | | | | | | | | |
|---|---|---|---|---|---|---|---|---|
| gene_237302 | 13,92546051 | 3,595519032 | 1,147622888 | 3,133014397 | 0,00173021 | 0,030426115 | The Moose: suction sampler vs FISH | NA |
| gene_237667 | 6,237890573 | 5,860281166 | 1,589566536 | 3,686716495 | 0,000227166 | 0,008499348 | The Moose: suction sampler vs FISH | NA |
| gene_238045 | 5,085334013 | 7,139541342 | 1,914142206 | 3,729890767 | 0,000191563 | 0,007614874 | The Moose: suction sampler vs FISH | NA |
| gene_241162 | 21,14141922 | 3,586582233 | 1,034691183 | 3,466331107 | 0,000527613 | 0,014448376 | The Moose: suction sampler vs FISH | NA |
| gene_243645 | 21,76005769 | 2,747632072 | 0,917335146 | 2,995232531 | 0,002742357 | 0,039821806 | The Moose: suction sampler vs FISH | NA |
| gene_243837 | 91,01529998 | 2,245529539 | 0,719634166 | 3,120376497 | 0,0018062 | 0,031191247 | The Moose: suction sampler vs FISH | NA |
| gene_24437 | 10,62647853 | 7,606376389 | 1,508453313 | 5,042500369 | 4,59E-07 | 0,000132342 | The Moose: suction sampler vs FISH | NA |
| gene_244533 | 4,857354861 | 5,94252984 | 1,884048372 | 3,154128061 | 0,001609784 | 0,029135192 | The Moose: suction sampler vs FISH | NA |
| gene_244665 | 22,81334978 | 3,393994791 | 1,012669452 | 3,35153271 | 0,000803655 | 0,018965252 | The Moose: suction sampler vs FISH | NA |
| gene_244872 | 2,749125394 | 6,189539964 | 2,032289003 | 3,045600284 | 0,002322163 | 0,036082789 | The Moose: suction sampler vs FISH | NA |
| gene_244909 | 30,4293929 | 4,967280405 | 0,975157501 | 5,093823713 | 3,51E-07 | 0,000109672 | The Moose: suction sampler vs FISH | NA |
| gene_245620 | 20,12815645 | 3,154798014 | 1,071971591 | 2,942986586 | 0,003250625 | 0,044160858 | The Moose: suction sampler vs FISH | NA |
| gene_245909 | 27,77573093 | 2,644179476 | 0,843821078 | 3,133578368 | 0,001726888 | 0,030422472 | The Moose: suction sampler vs FISH | NA |
| gene_246216 | 5,412429137 | 6,414404724 | 1,809763211 | 3,544333692 | 0,000393607 | 0,012080334 | The Moose: suction sampler vs FISH | NA |
| gene_246342 | 23,46365611 | 3,342358151 | 0,980655717 | 3,40828906 | 0,000653716 | 0,016570047 | The Moose: suction sampler vs FISH | NA |
| gene_246343 | 8,731675407 | 5,548820418 | 1,467561789 | 3,780979076 | 0,000156213 | 0,00664719 | The Moose: suction sampler vs FISH | NA |
| gene_247855 | 12,19333988 | 4,377574614 | 1,160035192 | 3,773656733 | 0,000160872 | 0,006802101 | The Moose: suction sampler vs FISH | NA |
| gene_249836 | 4,144015261 | 6,156141817 | 2,08332021 | 2,954966686 | 0,003127028 | 0,043219384 | The Moose: suction sampler vs FISH | NA |
| gene_250181 | 15,92215807 | -3,82291601 | 1,326369883 | -2,882239758 | 0,003948592 | 0,049430905 | The Moose: suction sampler vs FISH | NA |
| gene_250284 | 294,8539517 | 8,314845232 | 1,092664744 | 7,609694808 | 2,75E-14 | 6,73E-10 | The Moose: suction sampler vs FISH | NA |
| gene_250475 | 47,24176543 | 2,490292187 | 0,750449285 | 3,318401706 | 0,000905342 | 0,020535154 | The Moose: suction sampler vs FISH | NA |
| gene_250477 | 65,84212389 | 2,228638828 | 0,707536738 | 3,149855982 | 0,00163351 | 0,029419597 | The Moose: suction sampler vs FISH | NA |
| gene_251070 | 18,4469187 | 3,485485446 | 0,947996658 | 3,676685374 | 0,000236284 | 0,008724932 | The Moose: suction sampler vs FISH | NA |
| gene_251329 | 24,78692078 | 3,504586642 | 0,872052251 | 4,018780568 | 5,85E-05 | 0,003498812 | The Moose: suction sampler vs FISH | NA |
| gene_252230 | 1779,677029 | 3,687818535 | 0,795710639 | 4,634622629 | 3,58E-06 | 0,000519034 | The Moose: suction sampler vs FISH | NA |
| gene_252527 | 30,95444096 | 2,956754634 | 0,867118678 | 3,409861544 | 0,000649959 | 0,016506298 | The Moose: suction sampler vs FISH | NA |
| gene_252552 | 16,16692519 | 4,177078559 | 1,365520085 | 3,058965302 | 0,002221029 | 0,035211572 | The Moose: suction sampler vs FISH | NA |
| gene_253449 | 4,126815505 | 6,373823648 | 2,039914446 | 3,124554396 | 0,001780746 | 0,030904273 | The Moose: suction sampler vs FISH | NA |
| gene_2542 | 71,54626035 | 3,163231368 | 0,905967837 | 3,491549303 | 0,000480228 | 0,013577723 | The Moose: suction sampler vs FISH | NA |
| gene_2543 | 30,45059398 | 4,40277067 | 0,992068472 | 4,437970556 | 9,08E-06 | 0,000995465 | The Moose: suction sampler vs FISH | NA |
| gene_2544 | 90,3182275 | 3,536877026 | 0,850261439 | 4,159752358 | 3,19E-05 | 0,002296276 | The Moose: suction sampler vs FISH | NA |
| gene_254460 | 35,30905645 | 2,754581134 | 0,886411437 | 3,107564975 | 0,001886355 | 0,032056506 | The Moose: suction sampler vs FISH | NA |
| gene_254597 | 12,70244475 | 4,196743613 | 1,142218084 | 3,67420519 | 0,000238591 | 0,008792429 | The Moose: suction sampler vs FISH | NA |
| gene_255162 | 5,315840378 | 6,553774515 | 1,853852706 | 3,535218571 | 0,000407438 | 0,01231452 | The Moose: suction sampler vs FISH | NA |
| gene_256944 | 6,971154221 | 4,379098956 | 1,275404478 | 3,43349818 | 0,000595846 | 0,015568098 | The Moose: suction sampler vs FISH | NA |
| gene_257636 | 4,735191155 | 6,250134502 | 1,997363414 | 3,129192443 | 0,001752875 | 0,030666956 | The Moose: suction sampler vs FISH | NA |
| gene_258795 | 14,11820643 | 4,572430672 | 1,129909118 | 4,046724291 | 5,19E-05 | 0,003223638 | The Moose: suction sampler vs FISH | NA |
| gene_258994 | 64,79152664 | 4,105790302 | 0,937026079 | 4,381724685 | 1,18E-05 | 0,001187867 | The Moose: suction sampler vs FISH | NA |
| gene_260948 | 46,06211821 | 3,400507722 | 0,95180404 | 3,5726973 | 0,000353323 | 0,011250696 | The Moose: suction sampler vs FISH | NA |
| gene_261300 | 5,975900177 | 5,103200798 | 1,709208767 | 2,985709467 | 0,002829213 | 0,040727469 | The Moose: suction sampler vs FISH | NA |
| gene_262224 | 5,117418696 | 5,190214711 | 1,775670814 | 2,922959971 | 0,003467211 | 0,045808473 | The Moose: suction sampler vs FISH | NA |
| gene_263528 | 21,25377396 | 3,838808618 | 0,951199245 | 4,035756585 | 5,44E-05 | 0,003314563 | The Moose: suction sampler vs FISH | NA |
| gene_263710 | 14,90800514 | 2,716885646 | 0,874549381 | 3,106612051 | 0,001892446 | 0,032114297 | The Moose: suction sampler vs FISH | NA |
| gene_263930 | 4,599238686 | 5,830415808 | 1,845969645 | 3,158457034 | 0,001586067 | 0,028860583 | The Moose: suction sampler vs FISH | NA |
| gene_264035 | 3,647334785 | 6,708045634 | 2,071909641 | 3,237614953 | 0,001205334 | 0,024393974 | The Moose: suction sampler vs FISH | NA |
| gene_264143 | 19,96977725 | 4,209819603 | 1,008169328 | 4,175706882 | 2,97E-05 | 0,002194943 | The Moose: suction sampler vs FISH | NA |
| gene_264144 | 9,93526075 | 4,181520508 | 1,208645001 | 3,459676334 | 0,000540825 | 0,014646348 | The Moose: suction sampler vs FISH | NA |
| gene_267218 | 118,2280164 | -2,46970211 | 0,804887888 | -3,068380266 | 0,002152226 | 0,034648127 | The Moose: suction sampler vs FISH | NA |
| gene_268029 | 10,30062025 | 3,427124063 | 1,183950361 | 2,894651815 | 0,003795793 | 0,048350551 | The Moose: suction sampler vs FISH | NA |
| gene_268030 | 11,94886931 | 2,820416319 | 0,974492229 | 2,894241982 | 0,003800751 | 0,048372106 | The Moose: suction sampler vs FISH | NA |
| gene_269364 | 271,2242866 | 6,363912035 | 0,960407215 | 6,626264292 | 3,44E-11 | 1,59E-07 | The Moose: suction sampler vs FISH | NA |
| gene_269908 | 7,397341963 | 6,201233262 | 1,759531896 | 3,524365359 | 0,000424498 | 0,012627498 | The Moose: suction sampler vs FISH | NA |
| gene_270015 | 28,50036051 | 3,848959445 | 1,168960778 | 3,292633522 | 0,000992538 | 0,021624717 | The Moose: suction sampler vs FISH | NA |
| gene_270381 | 62,89212909 | 3,589131009 | 0,853095276 | 4,207186593 | 2,59E-05 | 0,002007468 | The Moose: suction sampler vs FISH | NA |
| gene_27051 | 90,63596233 | 2,128481868 | 0,716224489 | 2,971808282 | 0,002960515 | 0,041790316 | The Moose: suction sampler vs FISH | NA |
| gene_270798 | 62,0907113 | 2,915763939 | 0,717416262 | 4,064256828 | 4,82E-05 | 0,003086725 | The Moose: suction sampler vs FISH | NA |
| gene_270799 | 41,16686149 | 3,1844435 | 0,865821855 | 3,677943079 | 0,000235122 | 0,008692925 | The Moose: suction sampler vs FISH | NA |
| gene_270800 | 79,36883197 | 2,763987199 | 0,797761504 | 3,464678583 | 0,000530866 | 0,014488822 | The Moose: suction sampler vs FISH | NA |
| gene_271469 | 5,818321621 | 6,847531602 | 1,842195032 | 3,717050304 | 0,000201562 | 0,007871985 | The Moose: suction sampler vs FISH | NA |
| gene_273389 | 4,355235898 | 6,377512979 | 2,013101061 | 3,168004379 | 0,001534892 | 0,028301954 | The Moose: suction sampler vs FISH | NA |
| gene_273390 | 68,28776499 | 4,222535067 | 0,863022719 | 4,89272759 | 9,94E-07 | 0,000221899 | The Moose: suction sampler vs FISH | NA |
| gene_27583 | 4,385256224 | 5,909047788 | 1,742867888 | 3,390416352 | 0,000697865 | 0,017257485 | The Moose: suction sampler vs FISH | NA |
| gene_276113 | 8,589623991 | 4,595074545 | 1,52394008 | 3,01525933 | 0,002567596 | 0,038434581 | The Moose: suction sampler vs FISH | NA |
| gene_276562 | 94,97828524 | 5,075845738 | 1,091278881 | 4,651281928 | 3,30E-06 | 0,000486504 | The Moose: suction sampler vs FISH | NA |
| gene_27829 | 10,10343645 | 4,397167734 | 1,108211905 | 3,967804095 | 7,25E-05 | 0,004051367 | The Moose: suction sampler vs FISH | NA |

| | | | | | | | | |
|---|---|---|---|---|---|---|---|---|
| gene_279301 | 40,24426137 | 2,527068726 | 0,778411622 | 3,246442696 | 0,00116857 | 0,023920816 | The Moose: suction sampler vs FISH | NA |
| gene_281637 | 11,09930188 | 4,185880255 | 1,194115248 | 3,505424006 | 0,000455881 | 0,013192337 | The Moose: suction sampler vs FISH | NA |
| gene_281958 | 1681,468565 | 3,989248242 | 0,995689592 | 4,006517968 | 6,16E-05 | 0,003613435 | The Moose: suction sampler vs FISH | NA |
| gene_28286 | 10,08092314 | -6,454404703 | 1,75047816 | -3,687223783 | 0,000226714 | 0,008486752 | The Moose: suction sampler vs FISH | NA |
| gene_283515 | 6,35194789 | 7,62827347 | 2,066864972 | 3,690745923 | 0,000223597 | 0,008400057 | The Moose: suction sampler vs FISH | NA |
| gene_283800 | 119,194129 | 2,707933089 | 0,831880432 | 3,255195078 | 0,001133145 | 0,023529499 | The Moose: suction sampler vs FISH | NA |
| gene_285040 | 16,27982759 | -3,11166207 | 1,072401675 | -2,901582629 | 0,003712828 | 0,047806572 | The Moose: suction sampler vs FISH | NA |
| gene_285398 | 67,08164418 | 3,590479298 | 1,121811549 | 3,200608248 | 0,001371378 | 0,02637363 | The Moose: suction sampler vs FISH | NA |
| gene_286240 | 7,36603784 | 6,149242401 | 1,952844484 | 3,148864363 | 0,001639063 | 0,029461808 | The Moose: suction sampler vs FISH | NA |
| gene_286348 | 8,332476796 | 5,74192057 | 1,629766023 | 3,523156385 | 0,00042644 | 0,012674974 | The Moose: suction sampler vs FISH | NA |
| gene_286739 | 14,21043169 | 6,023199546 | 1,400399142 | 4,301059152 | 1,70E-05 | 0,001500535 | The Moose: suction sampler vs FISH | NA |
| gene_286937 | 10,7380839 | 5,107662239 | 1,299754696 | 3,929712472 | 8,50E-05 | 0,004493751 | The Moose: suction sampler vs FISH | NA |
| gene_287016 | 7,108158687 | 7,883298893 | 1,892853377 | 4,164769966 | 3,12E-05 | 0,002267936 | The Moose: suction sampler vs FISH | NA |
| gene_287753 | 7,585908108 | 7,639062401 | 1,822002634 | 4,192673632 | 2,76E-05 | 0,00210257 | The Moose: suction sampler vs FISH | NA |
| gene_287854 | 15,07715475 | 3,640964284 | 1,053498841 | 3,456068619 | 0,000548116 | 0,014792584 | The Moose: suction sampler vs FISH | NA |
| gene_287855 | 16,23832623 | 3,771931748 | 1,018563707 | 3,703186872 | 0,000212908 | 0,008135808 | The Moose: suction sampler vs FISH | NA |
| gene_288405 | 6,281827203 | 4,323048338 | 1,486154517 | 2,908882144 | 0,003627236 | 0,047077827 | The Moose: suction sampler vs FISH | NA |
| gene_289653 | 51,06911032 | 2,93649153 | 0,947262593 | 3,099976237 | 0,001935362 | 0,032541813 | The Moose: suction sampler vs FISH | NA |
| gene_289778 | 12,81530286 | 3,319531278 | 1,05354934 | 3,150807611 | 0,001628197 | 0,029347992 | The Moose: suction sampler vs FISH | NA |
| gene_289811 | 7,940580084 | 4,781337032 | 1,611302252 | 2,967374387 | 0,00300355 | 0,042227358 | The Moose: suction sampler vs FISH | NA |
| gene_29071 | 4,285253761 | 7,323209858 | 2,201186689 | 3,32693719 | 0,000878062 | 0,020152886 | The Moose: suction sampler vs FISH | NA |
| gene_290728 | 23,1931074 | -3,999903396 | 1,248719332 | -3,203204511 | 0,001359074 | 0,026212503 | The Moose: suction sampler vs FISH | NA |
| gene_291276 | 30,33600492 | 2,992898212 | 0,774257994 | 3,865505082 | 0,00011086 | 0,005342571 | The Moose: suction sampler vs FISH | NA |
| gene_29143 | 16,77417912 | 2,899470004 | 0,943939649 | 3,071668838 | 0,002128657 | 0,034435957 | The Moose: suction sampler vs FISH | NA |
| gene_291627 | 97,18269176 | 3,134522817 | 0,82187195 | 3,813882218 | 0,000136801 | 0,006133899 | The Moose: suction sampler vs FISH | NA |
| gene_292691 | 13,43512447 | 3,531260273 | 1,070853252 | 3,297613624 | 0,000975102 | 0,021384406 | The Moose: suction sampler vs FISH | NA |
| gene_292703 | 27,37342687 | 4,340763501 | 1,156500571 | 3,753360447 | 0,00017448 | 0,00714705 | The Moose: suction sampler vs FISH | NA |
| gene_293770 | 3,14755852 | 6,333875961 | 2,069821708 | 3,060107031 | 0,002212579 | 0,035181396 | The Moose: suction sampler vs FISH | NA |
| gene_294156 | 4,702745379 | 5,295581945 | 1,756931323 | 3,014108677 | 0,002577355 | 0,038513492 | The Moose: suction sampler vs FISH | NA |
| gene_2946 | 33,29008422 | 2,573674593 | 0,882086217 | 2,917713194 | 0,003526085 | 0,046244641 | The Moose: suction sampler vs FISH | NA |
| gene_29473 | 16,01445263 | 3,577403531 | 1,170660246 | 3,05588538 | 0,00224397 | 0,035391185 | The Moose: suction sampler vs FISH | NA |
| gene_294959 | 9,057743624 | 4,226930048 | 1,294677418 | 3,264851916 | 0,001095213 | 0,023074565 | The Moose: suction sampler vs FISH | NA |
| gene_295268 | 22,79974213 | 2,960711262 | 0,83649399 | 3,539429208 | 0,000400993 | 0,01221018 | The Moose: suction sampler vs FISH | NA |
| gene_295316 | 14,25435046 | 3,45005273 | 1,08466306 | 3,180759865 | 0,001468893 | 0,027570373 | The Moose: suction sampler vs FISH | NA |
| gene_295573 | 10,11607835 | 6,109050111 | 1,666057735 | 3,666769754 | 0,000245634 | 0,00894426 | The Moose: suction sampler vs FISH | NA |
| gene_295698 | 50,03323458 | 3,738313902 | 0,89828415 | 4,161616236 | 3,16E-05 | 0,002284654 | The Moose: suction sampler vs FISH | NA |
| gene_295852 | 148,7232009 | 2,266263995 | 0,770997783 | 2,939391066 | 0,003288578 | 0,044447856 | The Moose: suction sampler vs FISH | NA |
| gene_296598 | 31,31437651 | 4,138580243 | 0,866331305 | 4,777133433 | 1,78E-06 | 0,000323252 | The Moose: suction sampler vs FISH | NA |
| gene_299891 | 30,11362589 | -5,763396185 | 1,324888863 | -4,350097843 | 1,36E-05 | 0,001299631 | The Moose: suction sampler vs FISH | NA |
| gene_299892 | 8,143722752 | -5,794757547 | 1,752429943 | -3,306698546 | 0,000944024 | 0,020997535 | The Moose: suction sampler vs FISH | NA |
| gene_299928 | 19,63308515 | -3,137193104 | 0,956962793 | -3,278281169 | 0,001044413 | 0,02238311 | The Moose: suction sampler vs FISH | NA |
| gene_299960 | 15,95069092 | 4,800359547 | 1,303631306 | 3,682298458 | 0,000231141 | 0,008595504 | The Moose: suction sampler vs FISH | NA |
| gene_301254 | 30,85739162 | 2,787527847 | 0,92073031 | 3,027518282 | 0,002465708 | 0,037439306 | The Moose: suction sampler vs FISH | NA |
| gene_302143 | 7,065497103 | -6,370276795 | 1,859365688 | -3,426048375 | 0,000612431 | 0,015877168 | The Moose: suction sampler vs FISH | NA |
| gene_303964 | 55,88128301 | 2,636453729 | 0,750556812 | 3,512663782 | 0,000443638 | 0,012965789 | The Moose: suction sampler vs FISH | NA |
| gene_304247 | 7,160437526 | 7,891615808 | 1,889185892 | 4,177257433 | 2,95E-05 | 0,002186829 | The Moose: suction sampler vs FISH | NA |
| gene_304703 | 17,15417123 | 2,575124769 | 0,862678624 | 2,985033704 | 0,002835471 | 0,040773117 | The Moose: suction sampler vs FISH | NA |
| gene_304951 | 9,784936712 | 4,052625442 | 1,323345016 | 3,062410327 | 0,002195623 | 0,03502552 | The Moose: suction sampler vs FISH | NA |
| gene_305195 | 9,545389545 | 3,50985591 | 1,13431644 | 3,094247589 | 0,001973127 | 0,032933924 | The Moose: suction sampler vs FISH | NA |
| gene_30575 | 92,7310907 | -2,336613807 | 0,804380882 | -2,904859947 | 0,003674175 | 0,047441941 | The Moose: suction sampler vs FISH | NA |
| gene_305834 | 19,6329365 | 6,694109603 | 1,5162579 | 4,414888526 | 1,01E-05 | 0,001071068 | The Moose: suction sampler vs FISH | NA |
| gene_305989 | 86,29221269 | -3,182995556 | 1,103974748 | -2,883214097 | 0,003936398 | 0,049327546 | The Moose: suction sampler vs FISH | NA |
| gene_307404 | 146,155471 | 9,929864084 | 2,783703232 | 3,567141774 | 0,000360896 | 0,01141517 | The Moose: suction sampler vs FISH | NA |
| gene_307869 | 21,19105164 | 3,66239947 | 1,102900794 | 3,320697103 | 0,000897929 | 0,020474439 | The Moose: suction sampler vs FISH | NA |
| gene_307870 | 18,15484947 | 4,414838365 | 1,021676175 | 4,321171886 | 1,55E-05 | 0,001407103 | The Moose: suction sampler vs FISH | NA |
| gene_307969 | 9,64887101 | 4,487109512 | 1,26385033 | 3,550348809 | 0,000384721 | 0,011892187 | The Moose: suction sampler vs FISH | NA |
| gene_308261 | 19,52015338 | 2,829840069 | 0,935783168 | 3,02403395 | 0,002494284 | 0,037763909 | The Moose: suction sampler vs FISH | NA |
| gene_308511 | 11,00654563 | 5,339398364 | 1,339876066 | 3,984994209 | 6,75E-05 | 0,003839038 | The Moose: suction sampler vs FISH | NA |
| gene_309150 | 44,91739285 | -2,941855184 | 0,725089684 | -4,057229403 | 4,97E-05 | 0,003144098 | The Moose: suction sampler vs FISH | NA |
| gene_30976 | 1626,877987 | 2,435302398 | 0,686387352 | 3,548000107 | 0,000388168 | 0,011963493 | The Moose: suction sampler vs FISH | NA |
| gene_30977 | 2272,494182 | 2,239701132 | 0,667999999 | 3,352846013 | 0,000799852 | 0,018928075 | The Moose: suction sampler vs FISH | NA |
| gene_312067 | 5,216361287 | 5,534886457 | 1,902944088 | 2,908591216 | 0,003630612 | 0,047086409 | The Moose: suction sampler vs FISH | NA |
| gene_312922 | 9,295565901 | 4,730228526 | 1,586774846 | 2,981033218 | 0,002872776 | 0,041064822 | The Moose: suction sampler vs FISH | NA |
| gene_316758 | 5,957214199 | 5,301508335 | 1,646001225 | 3,220841062 | 0,00127815 | 0,025289765 | The Moose: suction sampler vs FISH | NA |
| gene_31691 | 94,4954574 | -2,248470225 | 0,770113049 | -2,919662545 | 0,003504106 | 0,046082171 | The Moose: suction sampler vs FISH | NA |

| | | | | | | | | |
|---|---|---|---|---|---|---|---|---|
| gene_318706 | 48,21810986 | 2,648598172 | 0,877609659 | 3,017968348 | 0,002544755 | 0,038212946 | The Moose: suction sampler vs FISH | NA |
| gene_31871 | 739,9931321 | 3,329542981 | 0,817581277 | 4,072430562 | 4,65E-05 | 0,003026602 | The Moose: suction sampler vs FISH | NA |
| gene_319060 | 5,456225179 | 6,200879157 | 1,755670385 | 3,531915335 | 0,000412561 | 0,012408097 | The Moose: suction sampler vs FISH | NA |
| gene_319062 | 3,691782495 | 5,557890411 | 1,918860145 | 2,896454139 | 0,003774058 | 0,048199831 | The Moose: suction sampler vs FISH | NA |
| gene_319162 | 17,71008743 | 4,287659542 | 1,21926153 | 3,516603648 | 0,000437106 | 0,012867033 | The Moose: suction sampler vs FISH | NA |
| gene_319799 | 144,2169082 | 2,770263016 | 0,64328057 | 4,306461511 | 1,66E-05 | 0,001472428 | The Moose: suction sampler vs FISH | NA |
| gene_319976 | 214,1363307 | 5,802145577 | 1,014014962 | 5,721952627 | 1,05E-08 | 9,48E-06 | The Moose: suction sampler vs FISH | NA |
| gene_320233 | 14,52630294 | 4,140910712 | 1,077031475 | 3,844744382 | 0,000120678 | 0,005656494 | The Moose: suction sampler vs FISH | NA |
| gene_32103 | 10,57778914 | 4,91717766 | 1,296099836 | 3,793826311 | 0,000148343 | 0,006416423 | The Moose: suction sampler vs FISH | NA |
| gene_321293 | 623,4004216 | -2,044028262 | 0,646807149 | -3,160181926 | 0,001576707 | 0,028734792 | The Moose: suction sampler vs FISH | NA |
| gene_321393 | 30,29630249 | 3,477726198 | 0,929154885 | 3,742891799 | 0,000181915 | 0,007327963 | The Moose: suction sampler vs FISH | NA |
| gene_323036 | 6,296447177 | 6,19329959 | 2,024732984 | 3,058822887 | 0,002222085 | 0,035212473 | The Moose: suction sampler vs FISH | NA |
| gene_323497 | 76,84022576 | 2,305424917 | 0,76947487 | 2,996101636 | 0,002734553 | 0,039760916 | The Moose: suction sampler vs FISH | NA |
| gene_324724 | 76,04250588 | 3,665084163 | 0,897703645 | 4,082732853 | 4,45E-05 | 0,002917852 | The Moose: suction sampler vs FISH | NA |
| gene_324725 | 52,69574585 | 2,898783769 | 0,847417592 | 3,420726446 | 0,000624541 | 0,016066349 | The Moose: suction sampler vs FISH | NA |
| gene_327891 | 9,471189172 | 3,762414884 | 1,226430497 | 3,067776684 | 0,002156577 | 0,034681368 | The Moose: suction sampler vs FISH | NA |
| gene_328527 | 3,387155035 | 5,724335062 | 1,983203529 | 2,886408267 | 0,003896663 | 0,049048664 | The Moose: suction sampler vs FISH | NA |
| gene_328688 | 4,103464906 | 5,926154159 | 1,923133287 | 3,081509846 | 0,002059537 | 0,033794162 | The Moose: suction sampler vs FISH | NA |
| gene_329273 | 4,954658395 | 4,925414095 | 1,650050898 | 2,985007373 | 0,002835715 | 0,040773117 | The Moose: suction sampler vs FISH | NA |
| gene_329322 | 238,5546499 | 3,752770491 | 0,818606763 | 4,584338488 | 4,55E-06 | 0,000620571 | The Moose: suction sampler vs FISH | NA |
| gene_331001 | 115,2682006 | 4,733638121 | 0,890349319 | 5,316607787 | 1,06E-07 | 4,79E-05 | The Moose: suction sampler vs FISH | NA |
| gene_331886 | 7,288187608 | 6,619626252 | 1,662490059 | 3,981753886 | 6,84E-05 | 0,003873762 | The Moose: suction sampler vs FISH | NA |
| gene_331887 | 33,46320936 | 3,758363517 | 0,858447546 | 4,378093378 | 1,20E-05 | 0,001193907 | The Moose: suction sampler vs FISH | NA |
| gene_334151 | 18,00137995 | -4,126417952 | 1,378879712 | -2,992587328 | 0,002766235 | 0,040056415 | The Moose: suction sampler vs FISH | NA |
| gene_33433 | 55,95915742 | 2,366631117 | 0,782652727 | 3,023858522 | 0,002495731 | 0,037769306 | The Moose: suction sampler vs FISH | NA |
| gene_334592 | 60,83140638 | 2,354073316 | 0,714167098 | 3,296250026 | 0,000979848 | 0,021456444 | The Moose: suction sampler vs FISH | NA |
| gene_335370 | 14,68204101 | -4,125921053 | 1,161172504 | -3,553236956 | 0,000380522 | 0,01181209 | The Moose: suction sampler vs FISH | NA |
| gene_337297 | 223,0969176 | 2,323185554 | 0,759662245 | 3,058182198 | 0,002226841 | 0,035228466 | The Moose: suction sampler vs FISH | NA |
| gene_337969 | 8,294021601 | 7,81181708 | 1,743471646 | 4,480610337 | 7,44E-06 | 0,000881694 | The Moose: suction sampler vs FISH | NA |
| gene_338589 | 21,23211392 | 2,961844961 | 0,909533376 | 3,256444503 | 0,00112817 | 0,023465996 | The Moose: suction sampler vs FISH | NA |
| gene_338915 | 719,5519888 | 4,912555412 | 0,930798962 | 5,277783508 | 1,31E-07 | 5,46E-05 | The Moose: suction sampler vs FISH | NA |
| gene_339651 | 4,272120663 | 6,381274562 | 1,984752266 | 3,21514915 | 0,001303768 | 0,025589314 | The Moose: suction sampler vs FISH | NA |
| gene_340584 | 13,77057303 | 3,639288983 | 1,036231378 | 3,512042831 | 0,000444676 | 0,012980625 | The Moose: suction sampler vs FISH | NA |
| gene_340919 | 8,135124809 | 5,437326624 | 1,733570185 | 3,136490619 | 0,001709829 | 0,030233485 | The Moose: suction sampler vs FISH | NA |
| gene_341925 | 48,77853986 | 4,455059014 | 0,977067145 | 4,559624217 | 5,12E-06 | 0,000668509 | The Moose: suction sampler vs FISH | NA |
| gene_342075 | 11,14021905 | 4,468426249 | 1,486273139 | 3,006463705 | 0,002643056 | 0,03898959 | The Moose: suction sampler vs FISH | NA |
| gene_342525 | 128,2822125 | -3,671696854 | 0,784709569 | -4,679051968 | 2,88E-06 | 0,000439641 | The Moose: suction sampler vs FISH | NA |
| gene_344155 | 63,42397521 | 3,17208663 | 0,857395639 | 3,699676655 | 0,000215874 | 0,008197978 | The Moose: suction sampler vs FISH | NA |
| gene_344620 | 10,66043976 | 3,827962858 | 1,327671288 | 2,883215819 | 0,003936376 | 0,049327546 | The Moose: suction sampler vs FISH | NA |
| gene_3452 | 13,6581375 | -3,40238834 | 0,989180386 | -3,439603523 | 0,000582567 | 0,015379809 | The Moose: suction sampler vs FISH | NA |
| gene_345588 | 25,06701586 | 3,359659295 | 1,046327393 | 3,210906375 | 0,00132317 | 0,025823392 | The Moose: suction sampler vs FISH | NA |
| gene_345790 | 5,57279356 | 4,819411818 | 1,53660927 | 3,13639382 | 0,001710394 | 0,030233485 | The Moose: suction sampler vs FISH | NA |
| gene_346809 | 8,235110451 | 6,321212054 | 1,553581821 | 4,068798932 | 4,73E-05 | 0,003057907 | The Moose: suction sampler vs FISH | NA |
| gene_347378 | 11,50343439 | 6,262909998 | 1,546942957 | 4,048572036 | 5,15E-05 | 0,003223638 | The Moose: suction sampler vs FISH | NA |
| gene_350865 | 4,997745441 | 6,483085948 | 1,900113396 | 3,411946867 | 0,000645007 | 0,016426416 | The Moose: suction sampler vs FISH | NA |
| gene_351431 | 13,87049331 | -5,714405716 | 1,543767405 | -3,701597597 | 0,000214246 | 0,008171553 | The Moose: suction sampler vs FISH | NA |
| gene_351432 | 9,514892382 | -7,254730544 | 1,831115766 | -3,961918017 | 7,44E-05 | 0,004121578 | The Moose: suction sampler vs FISH | NA |
| gene_351599 | 8,71759663 | 4,945363824 | 1,496034864 | 3,305647444 | 0,000947572 | 0,02105725 | The Moose: suction sampler vs FISH | NA |
| gene_352478 | 6,084082208 | 5,279790319 | 1,800033349 | 2,933162501 | 0,003355282 | 0,044952336 | The Moose: suction sampler vs FISH | NA |
| gene_353008 | 74,97375047 | 6,201781103 | 1,783202869 | 3,477888697 | 0,00050538 | 0,014054379 | The Moose: suction sampler vs FISH | NA |
| gene_353103 | 202,0472419 | 3,526233337 | 0,795156826 | 4,434638833 | 9,22E-06 | 0,001006478 | The Moose: suction sampler vs FISH | NA |
| gene_35361 | 6,065940245 | 6,064714884 | 2,023502801 | 2,997136886 | 0,002725283 | 0,039706202 | The Moose: suction sampler vs FISH | NA |
| gene_353650 | 15,48595093 | -4,384086856 | 1,487285126 | -2,947711087 | 0,003201361 | 0,043776571 | The Moose: suction sampler vs FISH | NA |
| gene_354029 | 4,864148488 | 6,598998306 | 1,994181424 | 3,309126354 | 0,000935876 | 0,020892221 | The Moose: suction sampler vs FISH | NA |
| gene_354235 | 26,52022588 | 3,739526647 | 1,043084259 | 3,585066705 | 0,000336992 | 0,010927321 | The Moose: suction sampler vs FISH | NA |
| gene_354372 | 35,97388053 | 3,304729866 | 1,139421218 | 2,900358368 | 0,003727362 | 0,047901336 | The Moose: suction sampler vs FISH | NA |
| gene_35452 | 29,37605493 | -2,658828129 | 0,806918626 | -3,295038736 | 0,000984081 | 0,021510666 | The Moose: suction sampler vs FISH | NA |
| gene_357933 | 17,1994424 | 4,908298261 | 1,125381909 | 4,361451187 | 1,29E-05 | 0,001256859 | The Moose: suction sampler vs FISH | NA |
| gene_357934 | 44,96223284 | 2,794914839 | 0,87101785 | 3,208791689 | 0,00133294 | 0,025904089 | The Moose: suction sampler vs FISH | NA |
| gene_359116 | 7,50480988 | 5,030614345 | 1,449957938 | 3,469489847 | 0,000521448 | 0,014321278 | The Moose: suction sampler vs FISH | NA |
| gene_360540 | 46,89933817 | 2,589905181 | 0,762574929 | 3,396263218 | 0,000683126 | 0,017039886 | The Moose: suction sampler vs FISH | NA |
| gene_363079 | 11,58156301 | 3,681743585 | 1,082817902 | 3,400150273 | 0,000673488 | 0,01689814 | The Moose: suction sampler vs FISH | NA |
| gene_36519 | 73,43453413 | 2,329587863 | 0,718040039 | 3,244370421 | 0,001177106 | 0,024038351 | The Moose: suction sampler vs FISH | NA |
| gene_366011 | 4,168188248 | 6,643982824 | 2,054171222 | 3,234386088 | 0,001219045 | 0,024539704 | The Moose: suction sampler vs FISH | NA |
| gene_368249 | 12,50685918 | 3,808094671 | 1,239661643 | 3,07188231 | 0,002127136 | 0,034429165 | The Moose: suction sampler vs FISH | NA |

| | | | | | | | | |
|---|---|---|---|---|---|---|---|---|
| gene_369150 | 10,42122002 | 4,106023142 | 1,127786797 | 3,640779582 | 0,000271814 | 0,009538157 | The Moose: suction sampler vs FISH | NA |
| gene_369580 | 18,44251391 | 3,481450774 | 1,081944298 | 3,217772653 | 0,001291902 | 0,0254655 | The Moose: suction sampler vs FISH | NA |
| gene_369665 | 6,458125538 | 7,449953537 | 1,760743969 | 4,231139602 | 2,33E-05 | 0,00185415 | The Moose: suction sampler vs FISH | NA |
| gene_370467 | 403,8993673 | 2,268342902 | 0,74372698 | 3,049967209 | 0,002288664 | 0,035756412 | The Moose: suction sampler vs FISH | NA |
| gene_371202 | 7,424963042 | 5,112146026 | 1,447836689 | 3,530885815 | 0,000414171 | 0,012436121 | The Moose: suction sampler vs FISH | NA |
| gene_372872 | 4,819047926 | 7,294352431 | 2,235650284 | 3,262743052 | 0,001103395 | 0,02318045 | The Moose: suction sampler vs FISH | NA |
| gene_373952 | 5,093075518 | 5,043234168 | 1,706662288 | 2,955027603 | 0,003126411 | 0,043219384 | The Moose: suction sampler vs FISH | NA |
| gene_374217 | 3,976453741 | 5,951906429 | 1,772722427 | 3,357494856 | 0,000786522 | 0,018712703 | The Moose: suction sampler vs FISH | NA |
| gene_374451 | 5,280714297 | 7,303349478 | 1,878383645 | 3,88810321 | 0,000101031 | 0,005030642 | The Moose: suction sampler vs FISH | NA |
| gene_375509 | 4,902000306 | 6,141267426 | 1,878104459 | 3,269928568 | 0,001075746 | 0,02279521 | The Moose: suction sampler vs FISH | NA |
| gene_376046 | 5,905449724 | 6,3509972 | 1,872807893 | 3,391163196 | 0,000695966 | 0,017235719 | The Moose: suction sampler vs FISH | NA |
| gene_376283 | 16,31224465 | 3,661415179 | 1,005259361 | 3,642259223 | 0,000270256 | 0,009517412 | The Moose: suction sampler vs FISH | NA |
| gene_376579 | 31,32547788 | 2,624765223 | 0,78736155 | 3,333621285 | 0,000857233 | 0,019873068 | The Moose: suction sampler vs FISH | NA |
| gene_37664 | 15,45337134 | 6,279096499 | 1,612914949 | 3,893011535 | 9,90E-05 | 0,00497608 | The Moose: suction sampler vs FISH | NA |
| gene_376808 | 9,296511067 | 4,365305367 | 1,250632001 | 3,490479505 | 0,000482155 | 0,013614704 | The Moose: suction sampler vs FISH | NA |
| gene_376809 | 6,288432146 | 5,02050931 | 1,533001803 | 3,274953297 | 0,001056794 | 0,022573109 | The Moose: suction sampler vs FISH | NA |
| gene_376822 | 12,03328544 | 2,985923478 | 0,989490642 | 3,017636903 | 0,002547539 | 0,038239124 | The Moose: suction sampler vs FISH | NA |
| gene_376880 | 14,27804556 | 3,584541361 | 1,133958485 | 3,161086944 | 0,001571816 | 0,028695503 | The Moose: suction sampler vs FISH | NA |
| gene_376881 | 12,29165793 | 4,79200672 | 1,343450565 | 3,566939376 | 0,000361175 | 0,011419069 | The Moose: suction sampler vs FISH | NA |
| gene_377634 | 30,1463501 | 3,079625964 | 0,941872312 | 3,269685207 | 0,001076672 | 0,022801671 | The Moose: suction sampler vs FISH | NA |
| gene_378487 | 119,9735057 | -2,440959878 | 0,763603671 | -3,196631933 | 0,001390422 | 0,026578131 | The Moose: suction sampler vs FISH | NA |
| gene_378488 | 331,0040353 | -2,329191446 | 0,632160125 | -3,684495989 | 0,000229156 | 0,008544153 | The Moose: suction sampler vs FISH | NA |
| gene_378489 | 46,64748246 | -2,294366821 | 0,736637045 | -3,114650339 | 0,001841631 | 0,031612271 | The Moose: suction sampler vs FISH | NA |
| gene_379002 | 34,4438068 | 5,446755232 | 1,374842938 | 3,96172907 | 7,44E-05 | 0,004121578 | The Moose: suction sampler vs FISH | NA |
| gene_3794 | 366,9660175 | 2,134804892 | 0,66510187 | 3,209741224 | 0,001328545 | 0,025868826 | The Moose: suction sampler vs FISH | NA |
| gene_379590 | 37,91684478 | 4,123347425 | 0,877685786 | 4,697976762 | 2,63E-06 | 0,000420431 | The Moose: suction sampler vs FISH | NA |
| gene_379591 | 14,96521749 | 4,205800623 | 1,090031999 | 3,858419411 | 0,000114123 | 0,005442689 | The Moose: suction sampler vs FISH | NA |
| gene_380043 | 19,04143277 | 3,769324283 | 1,06163933 | 3,550475362 | 0,000384536 | 0,011891476 | The Moose: suction sampler vs FISH | NA |
| gene_380159 | 107,3064359 | 3,406264985 | 0,829434055 | 4,106733942 | 4,01E-05 | 0,002711404 | The Moose: suction sampler vs FISH | NA |
| gene_380423 | 6,429437156 | 6,646029976 | 1,636667622 | 4,06070841 | 4,89E-05 | 0,00312183 | The Moose: suction sampler vs FISH | NA |
| gene_381679 | 7,328215282 | 5,462737849 | 1,700857111 | 3,211755893 | 0,001319264 | 0,025804086 | The Moose: suction sampler vs FISH | NA |
| gene_381852 | 13,3719778 | 3,301556314 | 1,088765035 | 3,032386426 | 0,002426284 | 0,037070608 | The Moose: suction sampler vs FISH | NA |
| gene_381960 | 18,53799033 | 4,341980525 | 1,37894998 | 3,14875854 | 0,001639656 | 0,029465267 | The Moose: suction sampler vs FISH | NA |
| gene_383805 | 16,01981885 | 3,021923612 | 1,049721063 | 2,87878725 | 0,003992075 | 0,049753602 | The Moose: suction sampler vs FISH | NA |
| gene_383806 | 20,79273152 | 3,892577168 | 1,004935048 | 3,87346145 | 0,0001073 | 0,005222096 | The Moose: suction sampler vs FISH | NA |
| gene_384582 | 18,79580427 | 4,090125339 | 1,076805628 | 3,798387779 | 0,00014564 | 0,006362888 | The Moose: suction sampler vs FISH | NA |
| gene_384920 | 10,28662876 | 4,958171565 | 1,308161975 | 3,790181691 | 0,000150537 | 0,006479324 | The Moose: suction sampler vs FISH | NA |
| gene_385626 | 4,46694569 | 5,903776952 | 1,954502338 | 3,020603679 | 0,002522713 | 0,037990706 | The Moose: suction sampler vs FISH | NA |
| gene_388617 | 4,054399565 | 6,563657796 | 1,979042437 | 3,316582643 | 0,000911256 | 0,020592991 | The Moose: suction sampler vs FISH | NA |
| gene_389188 | 495,237443 | -2,214414656 | 0,653377601 | -3,38918055 | 0,000701018 | 0,01731796 | The Moose: suction sampler vs FISH | NA |
| gene_390068 | 9,706808403 | -4,825892518 | 1,568700213 | -3,076363782 | 0,00209542 | 0,034154056 | The Moose: suction sampler vs FISH | NA |
| gene_390069 | 19,83932481 | -4,195354971 | 1,226363889 | -3,420970732 | 0,00062398 | 0,016057549 | The Moose: suction sampler vs FISH | NA |
| gene_390318 | 39,47963009 | 3,885056368 | 0,90609274 | 4,287702786 | 1,81E-05 | 0,001552767 | The Moose: suction sampler vs FISH | NA |
| gene_39053 | 4,385758849 | 5,367160573 | 1,729023197 | 3,104157644 | 0,001908216 | 0,032262648 | The Moose: suction sampler vs FISH | NA |
| gene_390534 | 19,10042667 | 4,096654593 | 1,124287959 | 3,643776989 | 0,000268666 | 0,009477517 | The Moose: suction sampler vs FISH | NA |
| gene_390754 | 32,153148 | 3,388831343 | 0,891214864 | 3,802485215 | 0,000143252 | 0,006307632 | The Moose: suction sampler vs FISH | NA |
| gene_390861 | 2,95506919 | 6,417480413 | 2,101862923 | 3,053234511 | 0,002263889 | 0,035551536 | The Moose: suction sampler vs FISH | NA |
| gene_392523 | 33,47359354 | 2,757902659 | 0,733415022 | 3,760357472 | 0,000169671 | 0,007031351 | The Moose: suction sampler vs FISH | NA |
| gene_393568 | 43,26900156 | -2,58110521 | 0,807313729 | -3,197152628 | 0,001387915 | 0,026542102 | The Moose: suction sampler vs FISH | NA |
| gene_393668 | 53,61847807 | 4,672780364 | 0,994864344 | 4,696902036 | 2,64E-06 | 0,000421729 | The Moose: suction sampler vs FISH | NA |
| gene_396022 | 1130,553397 | 2,419945989 | 0,633929032 | 3,817376813 | 0,000134878 | 0,006081106 | The Moose: suction sampler vs FISH | NA |
| gene_39662 | 21,06772816 | 3,074539438 | 0,919424957 | 3,343980838 | 0,000825854 | 0,019337767 | The Moose: suction sampler vs FISH | NA |
| gene_398685 | 5,556570297 | 4,837741044 | 1,549258541 | 3,122616992 | 0,001792508 | 0,031046853 | The Moose: suction sampler vs FISH | NA |
| gene_399720 | 4,287599077 | 5,948479949 | 1,850060475 | 3,215289461 | 0,00130313 | 0,025586188 | The Moose: suction sampler vs FISH | NA |
| gene_401286 | 32,19977735 | 3,185316266 | 0,888232643 | 3,586128355 | 0,000335624 | 0,010917666 | The Moose: suction sampler vs FISH | NA |
| gene_401498 | 69,82917798 | 2,338714555 | 0,812774042 | 2,877447403 | 0,004009067 | 0,049863836 | The Moose: suction sampler vs FISH | NA |
| gene_401665 | 53,31830924 | 3,817544883 | 1,019808743 | 3,743392973 | 0,000181552 | 0,007326419 | The Moose: suction sampler vs FISH | NA |
| gene_402021 | 19,40415389 | 3,089475616 | 0,942571342 | 3,27771011 | 0,001046528 | 0,022415289 | The Moose: suction sampler vs FISH | NA |
| gene_402166 | 16,39093836 | 4,404951138 | 1,281644657 | 3,436951977 | 0,0005883 | 0,015458922 | The Moose: suction sampler vs FISH | NA |
| gene_402392 | 5,172091762 | 6,729217377 | 1,757090118 | 3,829750852 | 0,000128273 | 0,005883033 | The Moose: suction sampler vs FISH | NA |
| gene_402571 | 3,696316516 | 6,285181506 | 2,059452292 | 3,051870408 | 0,002274203 | 0,035649607 | The Moose: suction sampler vs FISH | NA |
| gene_402967 | 7,799380002 | 5,017338627 | 1,727355797 | 2,904635302 | 0,003676812 | 0,047459313 | The Moose: suction sampler vs FISH | NA |
| gene_403997 | 9,322833786 | 4,179536639 | 1,256221578 | 3,327069614 | 0,000877644 | 0,020152886 | The Moose: suction sampler vs FISH | NA |
| gene_405355 | 139,5052359 | -2,728182387 | 0,904754326 | -3,015384741 | 0,002566535 | 0,038429997 | The Moose: suction sampler vs FISH | NA |
| gene_40586 | 18,44741298 | 3,475515795 | 0,964103928 | 3,604918197 | 0,000312251 | 0,010457499 | The Moose: suction sampler vs FISH | NA |

| | | | | | | | | |
|---|---|---|---|---|---|---|---|---|
| gene_408821 | 12,58782773 | 2,944300734 | 1,011125098 | 2,911905501 | 0,003592313 | 0,046815421 | The Moose: suction sampler vs FISH | NA |
| gene_409290 | 16,32295575 | -3,195633757 | 1,045514749 | -3,056517146 | 0,002239247 | 0,035360455 | The Moose: suction sampler vs FISH | NA |
| gene_412480 | 4,728506938 | 5,894964876 | 1,961115236 | 3,005924776 | 0,002647745 | 0,039038312 | The Moose: suction sampler vs FISH | NA |
| gene_412811 | 39,54944012 | 3,870534359 | 0,856275879 | 4,520195483 | 6,18E-06 | 0,000769156 | The Moose: suction sampler vs FISH | NA |
| gene_412812 | 16,08459754 | 3,847297455 | 1,077999867 | 3,568922014 | 0,000358453 | 0,01136236 | The Moose: suction sampler vs FISH | NA |
| gene_412813 | 12,61547233 | 3,526637387 | 1,053526399 | 3,347459912 | 0,000815558 | 0,019179841 | The Moose: suction sampler vs FISH | NA |
| gene_41325 | 7,088946571 | -6,098615145 | 1,682963258 | -3,623736357 | 0,000290378 | 0,009965786 | The Moose: suction sampler vs FISH | NA |
| gene_413480 | 5,561057222 | 5,360821318 | 1,68872786 | 3,174473191 | 0,001501088 | 0,027946872 | The Moose: suction sampler vs FISH | NA |
| gene_413552 | 6,943062342 | 5,358956011 | 1,532108992 | 3,497764219 | 0,000469176 | 0,013392145 | The Moose: suction sampler vs FISH | NA |
| gene_415476 | 122,3642132 | -2,182517084 | 0,666852153 | -3,27286502 | 0,001064633 | 0,022656159 | The Moose: suction sampler vs FISH | NA |
| gene_416622 | 7,835324691 | 4,55628579 | 1,446716829 | 3,149397104 | 0,001636077 | 0,029448747 | The Moose: suction sampler vs FISH | NA |
| gene_417288 | 7,632437944 | 4,40647847 | 1,478610006 | 2,98014923 | 0,00288108 | 0,041108821 | The Moose: suction sampler vs FISH | NA |
| gene_418094 | 11,32101609 | 4,26393324 | 1,355011753 | 3,146786905 | 0,001650752 | 0,029592262 | The Moose: suction sampler vs FISH | NA |
| gene_41881 | 13,43891982 | 3,719262184 | 1,164277027 | 3,194482153 | 0,001400819 | 0,026706306 | The Moose: suction sampler vs FISH | NA |
| gene_420192 | 31,68146343 | 4,881663954 | 1,020664603 | 4,782828697 | 1,73E-06 | 0,000316574 | The Moose: suction sampler vs FISH | NA |
| gene_420193 | 18,85616461 | 3,6618124 | 1,059432985 | 3,456388892 | 0,000547465 | 0,014782557 | The Moose: suction sampler vs FISH | NA |
| gene_420194 | 57,51003688 | 4,761777958 | 0,851638493 | 5,591313681 | 2,25E-08 | 1,61E-05 | The Moose: suction sampler vs FISH | NA |
| gene_421234 | 6,791519999 | 4,984429367 | 1,659250021 | 3,004025495 | 0,00266433 | 0,039191211 | The Moose: suction sampler vs FISH | NA |
| gene_421521 | 29,95085792 | 2,792938849 | 0,779207641 | 3,584331957 | 0,000337942 | 0,01095329 | The Moose: suction sampler vs FISH | NA |
| gene_421522 | 167,5233007 | 2,86305493 | 0,657921895 | 4,351663854 | 1,35E-05 | 0,001296276 | The Moose: suction sampler vs FISH | NA |
| gene_421605 | 73,94057628 | 2,167045892 | 0,723038879 | 2,997136052 | 0,002725291 | 0,039706202 | The Moose: suction sampler vs FISH | NA |
| gene_422340 | 5,005816742 | 6,63487501 | 1,779451896 | 3,728605996 | 0,000192542 | 0,007635659 | The Moose: suction sampler vs FISH | NA |
| gene_422786 | 1077,336769 | -2,245833343 | 0,618373717 | -3,631838289 | 0,000281409 | 0,009779641 | The Moose: suction sampler vs FISH | NA |
| gene_423189 | 43,19073508 | 2,568180459 | 0,874758207 | 2,935874666 | 0,003326087 | 0,044773542 | The Moose: suction sampler vs FISH | NA |
| gene_423259 | 10,33920165 | 4,185418755 | 1,31686635 | 3,178317036 | 0,001481327 | 0,027690518 | The Moose: suction sampler vs FISH | NA |
| gene_423262 | 30,42746978 | 3,58545485 | 1,004798955 | 3,568330592 | 0,000359263 | 0,011381105 | The Moose: suction sampler vs FISH | NA |
| gene_425134 | 6,910230823 | 7,471682928 | 1,865766523 | 4,00461839 | 6,21E-05 | 0,003629461 | The Moose: suction sampler vs FISH | NA |
| gene_425323 | 26,7102877 | 3,834948766 | 1,004286217 | 3,818581496 | 0,000134221 | 0,006055208 | The Moose: suction sampler vs FISH | NA |
| gene_427172 | 116,6533399 | 4,195808352 | 0,87022438 | 4,821524708 | 1,42E-06 | 0,000279769 | The Moose: suction sampler vs FISH | NA |
| gene_427723 | 23,14695318 | 4,361915897 | 1,159426198 | 3,762133289 | 0,00016847 | 0,00701434 | The Moose: suction sampler vs FISH | NA |
| gene_427724 | 13,8181218 | 5,449162593 | 1,3966464 | 3,90160501 | 9,56E-05 | 0,004864833 | The Moose: suction sampler vs FISH | NA |
| gene_427827 | 8,129556128 | 4,769376085 | 1,520968865 | 3,135748662 | 0,00171416 | 0,030278141 | The Moose: suction sampler vs FISH | NA |
| gene_42849 | 5,10682182 | 6,354008557 | 2,205873276 | 2,880495732 | 0,003970503 | 0,049588781 | The Moose: suction sampler vs FISH | NA |
| gene_429050 | 16,17413718 | 3,032925123 | 1,03387211 | 2,933559279 | 0,003350997 | 0,044942491 | The Moose: suction sampler vs FISH | NA |
| gene_429287 | 110,7796426 | -2,300966574 | 0,791885253 | -2,905681807 | 0,003664539 | 0,047400856 | The Moose: suction sampler vs FISH | NA |
| gene_429291 | 32,95900043 | 2,450599047 | 0,765180614 | 3,202641315 | 0,001361735 | 0,026236538 | The Moose: suction sampler vs FISH | NA |
| gene_429292 | 81,69977955 | 2,829949914 | 0,691132528 | 4,094655942 | 4,23E-05 | 0,002811398 | The Moose: suction sampler vs FISH | NA |
| gene_429338 | 29,9721864 | 3,70769867 | 0,965965306 | 3,838335235 | 0,000123871 | 0,005765353 | The Moose: suction sampler vs FISH | NA |
| gene_429339 | 25,94205105 | 3,400355789 | 0,911620334 | 3,730013101 | 0,00019147 | 0,007614874 | The Moose: suction sampler vs FISH | NA |
| gene_430866 | 30,71141494 | 3,338657165 | 0,825000857 | 4,046852966 | 5,19E-05 | 0,003223638 | The Moose: suction sampler vs FISH | NA |
| gene_431023 | 15,80943958 | 3,45274543 | 1,132509295 | 3,048756814 | 0,002297904 | 0,035839789 | The Moose: suction sampler vs FISH | NA |
| gene_432119 | 7,212996159 | 5,587278492 | 1,797200172 | 3,108879344 | 0,001877984 | 0,031994556 | The Moose: suction sampler vs FISH | NA |
| gene_432155 | 19,10789012 | 4,804846934 | 1,236697441 | 3,885224288 | 0,000102235 | 0,005069338 | The Moose: suction sampler vs FISH | NA |
| gene_432603 | 10,90205613 | 3,597557299 | 1,222031998 | 2,943914156 | 0,003240899 | 0,044095556 | The Moose: suction sampler vs FISH | NA |
| gene_433066 | 13,62001836 | 4,346395196 | 1,070713745 | 4,059343792 | 4,92E-05 | 0,003123845 | The Moose: suction sampler vs FISH | NA |
| gene_433221 | 54,96965281 | -3,322967565 | 0,966326128 | -3,438764064 | 0,000584376 | 0,015399043 | The Moose: suction sampler vs FISH | NA |
| gene_434373 | 13,97972565 | 4,474993146 | 1,546176599 | 2,894231583 | 0,003800877 | 0,048372106 | The Moose: suction sampler vs FISH | NA |
| gene_43510 | 5,084642715 | 5,54380975 | 1,67593357 | 3,307893493 | 0,000940006 | 0,020938121 | The Moose: suction sampler vs FISH | NA |
| gene_435150 | 369,6272409 | 2,844915566 | 0,693653888 | 4,101347396 | 4,11E-05 | 0,002754828 | The Moose: suction sampler vs FISH | NA |
| gene_435493 | 18,32978175 | 5,26323295 | 1,080354297 | 4,871765644 | 1,11E-06 | 0,000236834 | The Moose: suction sampler vs FISH | NA |
| gene_436043 | 32,88057991 | -2,000333736 | 0,6934457 | -2,884629233 | 0,003918749 | 0,049188385 | The Moose: suction sampler vs FISH | NA |
| gene_436096 | 9,071417297 | 3,956056558 | 1,34278945 | 2,946148079 | 0,003217583 | 0,04392122 | The Moose: suction sampler vs FISH | NA |
| gene_437277 | 12,5124462 | 4,013523745 | 1,365113694 | 2,940065551 | 0,003281428 | 0,044378532 | The Moose: suction sampler vs FISH | NA |
| gene_438008 | 13,27446948 | 4,01484997 | 1,259197676 | 3,188419138 | 0,00143053 | 0,027071705 | The Moose: suction sampler vs FISH | NA |
| gene_438427 | 13,76398559 | 5,055341169 | 1,281095159 | 3,946109025 | 7,94E-05 | 0,004277011 | The Moose: suction sampler vs FISH | NA |
| gene_438926 | 16,4822076 | -2,703823304 | 0,865769648 | -3,123028521 | 0,001790004 | 0,03103561 | The Moose: suction sampler vs FISH | NA |
| gene_438931 | 59,77236877 | -2,491674184 | 0,637369082 | -3,909311349 | 9,26E-05 | 0,004763868 | The Moose: suction sampler vs FISH | NA |
| gene_439606 | 18,494934 | 3,511350509 | 1,176572589 | 2,984389185 | 0,002841451 | 0,040815641 | The Moose: suction sampler vs FISH | NA |
| gene_439881 | 23,72101897 | 2,903858685 | 0,904040137 | 3,212090442 | 0,001317729 | 0,02579303 | The Moose: suction sampler vs FISH | NA |
| gene_440796 | 22,80825596 | 2,851499738 | 0,762143889 | 3,741419145 | 0,000182984 | 0,007355919 | The Moose: suction sampler vs FISH | NA |
| gene_443388 | 34,41900343 | 2,457200141 | 0,734263231 | 3,346483982 | 0,000818434 | 0,019216717 | The Moose: suction sampler vs FISH | NA |
| gene_44359 | 444,3451524 | -2,719725089 | 0,595540752 | -4,566816094 | 4,95E-06 | 0,000655485 | The Moose: suction sampler vs FISH | NA |
| gene_44360 | 857,9172007 | -2,347067471 | 0,57090776 | -4,111115024 | 3,94E-05 | 0,002675225 | The Moose: suction sampler vs FISH | NA |
| gene_444043 | 6,485135843 | 5,264784999 | 1,749622361 | 3,009097915 | 0,002620247 | 0,038864183 | The Moose: suction sampler vs FISH | NA |
| gene_448150 | 22,97894431 | 3,715270422 | 1,061448358 | 3,500189523 | 0,000464927 | 0,013354236 | The Moose: suction sampler vs FISH | NA |

| | | | | | | | | |
|---|---|---|---|---|---|---|---|---|
| gene_448738 | 4,100856689 | 5,700905688 | 1,890483096 | 3,015581414 | 0,002564871 | 0,038412914 | The Moose: suction sampler vs FISH | NA |
| gene_449563 | 78,81679768 | 2,604477528 | 0,692063569 | 3,763350135 | 0,000167652 | 0,006992173 | The Moose: suction sampler vs FISH | NA |
| gene_450059 | 4,340090113 | 6,9014647 | 1,917058726 | 3,600027797 | 0,000318183 | 0,010580484 | The Moose: suction sampler vs FISH | NA |
| gene_450604 | 4,864785705 | 6,227098875 | 1,765985111 | 3,526133282 | 0,000421675 | 0,012594509 | The Moose: suction sampler vs FISH | NA |
| gene_452019 | 12,65235441 | 4,164156533 | 1,177675085 | 3,535912906 | 0,000406369 | 0,012287256 | The Moose: suction sampler vs FISH | NA |
| gene_453666 | 57,42252955 | 5,122856566 | 1,100057173 | 4,65690029 | 3,21E-06 | 0,000477252 | The Moose: suction sampler vs FISH | NA |
| gene_45392 | 16,03593186 | 4,993547092 | 1,452236301 | 3,438522428 | 0,000584898 | 0,015402596 | The Moose: suction sampler vs FISH | NA |
| gene_45393 | 4,28778279 | 6,424036685 | 2,137439228 | 3,00548273 | 0,002651597 | 0,039050868 | The Moose: suction sampler vs FISH | NA |
| gene_454523 | 14,9034721 | 3,201849674 | 1,079509248 | 2,966023383 | 0,003016776 | 0,042332266 | The Moose: suction sampler vs FISH | NA |
| gene_454960 | 5,802255868 | 6,357087025 | 1,884731773 | 3,372939915 | 0,000743702 | 0,018074513 | The Moose: suction sampler vs FISH | NA |
| gene_456138 | 11,31662984 | 4,841366367 | 1,652719272 | 2,929333765 | 0,003396895 | 0,045270354 | The Moose: suction sampler vs FISH | NA |
| gene_456404 | 42,13330697 | 2,381467348 | 0,820001412 | 2,904223472 | 0,003681653 | 0,047505092 | The Moose: suction sampler vs FISH | NA |
| gene_456959 | 31,94895859 | 4,926740496 | 1,027158182 | 4,796476904 | 1,61E-06 | 0,000306899 | The Moose: suction sampler vs FISH | NA |
| gene_457785 | 6,617609693 | 6,779442566 | 1,690258837 | 4,010890177 | 6,05E-05 | 0,00357994 | The Moose: suction sampler vs FISH | NA |
| gene_460090 | 19,00412252 | 3,566803882 | 1,17946964 | 3,024074348 | 0,002493951 | 0,037763909 | The Moose: suction sampler vs FISH | NA |
| gene_460091 | 26,72787211 | 4,337993602 | 1,020115176 | 4,252454727 | 2,11E-05 | 0,001731238 | The Moose: suction sampler vs FISH | NA |
| gene_46125 | 67,87678987 | 11,40482495 | 3,533450976 | 3,227673179 | 0,001248015 | 0,024900962 | The Moose: suction sampler vs FISH | NA |
| gene_461811 | 17,10463279 | 3,397309174 | 1,150290597 | 2,953435578 | 0,003142582 | 0,043327751 | The Moose: suction sampler vs FISH | NA |
| gene_4631 | 19,07152912 | 3,146372771 | 1,008899242 | 3,118619421 | 0,001817005 | 0,031347689 | The Moose: suction sampler vs FISH | NA |
| gene_464320 | 7,874382642 | 6,589703737 | 1,672256836 | 3,940605052 | 8,13E-05 | 0,004341339 | The Moose: suction sampler vs FISH | NA |
| gene_464804 | 6,859790922 | 4,81614123 | 1,636547889 | 2,942866055 | 0,003251891 | 0,044160858 | The Moose: suction sampler vs FISH | NA |
| gene_465191 | 50,03467633 | 3,044988054 | 0,743171114 | 4,097290644 | 4,18E-05 | 0,002791003 | The Moose: suction sampler vs FISH | NA |
| gene_466281 | 21,86854661 | 5,746968525 | 1,165714072 | 4,929998413 | 8,22E-07 | 0,000196723 | The Moose: suction sampler vs FISH | NA |
| gene_466282 | 6,739506249 | 4,771627939 | 1,516653922 | 3,146154749 | 0,001654324 | 0,02964183 | The Moose: suction sampler vs FISH | NA |
| gene_466693 | 6,876559063 | 6,956916678 | 1,702333813 | 4,086693589 | 4,38E-05 | 0,002879652 | The Moose: suction sampler vs FISH | NA |
| gene_46804 | 7,423434846 | 6,400897364 | 2,001999972 | 3,197251475 | 0,001387439 | 0,026542102 | The Moose: suction sampler vs FISH | NA |
| gene_468591 | 6,414441779 | 4,506498189 | 1,524084183 | 2,956856478 | 0,003107927 | 0,043085392 | The Moose: suction sampler vs FISH | NA |
| gene_468902 | 65,95603807 | 2,114989684 | 0,730329818 | 2,895937743 | 0,003780274 | 0,048233054 | The Moose: suction sampler vs FISH | NA |
| gene_470196 | 3,801127843 | 7,128662415 | 2,350157779 | 3,033269714 | 0,002419193 | 0,036985348 | The Moose: suction sampler vs FISH | NA |
| gene_470881 | 31,32370598 | -2,410822999 | 0,805820589 | -2,991761481 | 0,002773729 | 0,040141185 | The Moose: suction sampler vs FISH | NA |
| gene_470882 | 20,01792313 | -3,053667529 | 0,86255181 | -3,540271431 | 0,000399716 | 0,012205583 | The Moose: suction sampler vs FISH | NA |
| gene_474672 | 114,2393385 | -2,28313769 | 0,641999083 | -3,556294314 | 0,000376123 | 0,011720579 | The Moose: suction sampler vs FISH | NA |
| gene_475242 | 13,10055768 | 3,125301719 | 1,06560648 | 2,932885429 | 0,003358278 | 0,044974217 | The Moose: suction sampler vs FISH | NA |
| gene_475837 | 18,8478832 | 3,203915266 | 1,061584634 | 3,018049776 | 0,002544071 | 0,038210492 | The Moose: suction sampler vs FISH | NA |
| gene_476083 | 5,727395327 | 4,584970724 | 1,51900731 | 3,018399382 | 0,002541138 | 0,038189863 | The Moose: suction sampler vs FISH | NA |
| gene_476756 | 6,271240087 | 7,431492195 | 1,853148516 | 4,010197851 | 6,07E-05 | 0,003584677 | The Moose: suction sampler vs FISH | NA |
| gene_478363 | 45,02695068 | 3,432873508 | 0,873239079 | 3,931195466 | 8,45E-05 | 0,004469333 | The Moose: suction sampler vs FISH | NA |
| gene_478364 | 6,229876166 | 6,882336592 | 1,637729349 | 4,202365058 | 2,64E-05 | 0,002037796 | The Moose: suction sampler vs FISH | NA |
| gene_479924 | 17,0091645 | 3,051405211 | 0,905762653 | 3,368879475 | 0,000754744 | 0,018252285 | The Moose: suction sampler vs FISH | NA |
| gene_481157 | 7,611334577 | 4,752188084 | 1,568074129 | 3,030588921 | 0,002440773 | 0,03717118 | The Moose: suction sampler vs FISH | NA |
| gene_482037 | 23,04583987 | 2,886197465 | 0,833498015 | 3,462752654 | 0,00053468 | 0,014549666 | The Moose: suction sampler vs FISH | NA |
| gene_482368 | 4,259671126 | 6,424122881 | 1,800485892 | 3,567994011 | 0,000359725 | 0,011387923 | The Moose: suction sampler vs FISH | NA |
| gene_482919 | 22,05890478 | 3,382975036 | 1,004221461 | 3,368753974 | 0,000755088 | 0,018254587 | The Moose: suction sampler vs FISH | NA |
| gene_483807 | 34,27645143 | 8,630705427 | 2,356985895 | 3,66175523 | 0,000250493 | 0,00904497 | The Moose: suction sampler vs FISH | NA |
| gene_483811 | 2,957409845 | 6,115855883 | 2,054213883 | 2,977224491 | 0,002908709 | 0,041340185 | The Moose: suction sampler vs FISH | NA |
| gene_484386 | 12,22671317 | 5,677949513 | 1,575620876 | 3,603626735 | 0,000313808 | 0,010495265 | The Moose: suction sampler vs FISH | NA |
| gene_484394 | 13,65948739 | 4,763256769 | 1,185665804 | 4,017368765 | 5,89E-05 | 0,00351411 | The Moose: suction sampler vs FISH | NA |
| gene_484646 | 45,18913229 | 3,527394464 | 1,00786571 | 3,499865536 | 0,000465493 | 0,013354236 | The Moose: suction sampler vs FISH | NA |
| gene_484761 | 16,52812308 | 3,535507095 | 1,094007087 | 3,231704013 | 0,001230544 | 0,024673038 | The Moose: suction sampler vs FISH | NA |
| gene_486390 | 16,59745086 | 3,239208995 | 1,033505032 | 3,134197605 | 0,001723248 | 0,030372912 | The Moose: suction sampler vs FISH | NA |
| gene_487868 | 27,41728634 | 3,997824216 | 0,916860526 | 4,360340643 | 1,30E-05 | 0,001261584 | The Moose: suction sampler vs FISH | NA |
| gene_490344 | 10,73291675 | 4,346302627 | 1,296939481 | 3,351199259 | 0,000804624 | 0,018977392 | The Moose: suction sampler vs FISH | NA |
| gene_492753 | 9,762633072 | 4,427618875 | 1,297507347 | 3,412403703 | 0,000643927 | 0,016409854 | The Moose: suction sampler vs FISH | NA |
| gene_49346 | 340,9499116 | -3,039178662 | 0,643861386 | -4,720237504 | 2,36E-06 | 0,000389671 | The Moose: suction sampler vs FISH | NA |
| gene_494881 | 13,71847562 | 4,967137405 | 1,592987782 | 3,118126492 | 0,001820047 | 0,031351249 | The Moose: suction sampler vs FISH | NA |
| gene_49570 | 9,240296349 | 4,382858698 | 1,503758119 | 2,914603514 | 0,003561407 | 0,046550547 | The Moose: suction sampler vs FISH | NA |
| gene_496205 | 13,91017984 | 8,750870485 | 1,556821359 | 5,620985628 | 1,90E-08 | 1,41E-05 | The Moose: suction sampler vs FISH | NA |
| gene_496206 | 8,904363412 | 7,141089828 | 1,741480935 | 4,100584556 | 4,12E-05 | 0,002759094 | The Moose: suction sampler vs FISH | NA |
| gene_496771 | 27,83893363 | 3,662215657 | 0,85436222 | 4,286490636 | 1,82E-05 | 0,001557427 | The Moose: suction sampler vs FISH | NA |
| gene_496772 | 128,5034558 | 3,037092813 | 0,696136725 | 4,362782057 | 1,28E-05 | 0,001254219 | The Moose: suction sampler vs FISH | NA |
| gene_496869 | 32,76810131 | 3,395425491 | 1,003157652 | 3,384737667 | 0,000712463 | 0,017536627 | The Moose: suction sampler vs FISH | NA |
| gene_497153 | 14,78204149 | 4,364776729 | 1,332867256 | 3,274727253 | 0,00105764 | 0,022580929 | The Moose: suction sampler vs FISH | NA |
| gene_497604 | 3,512905434 | 6,399247624 | 1,936730883 | 3,30414911 | 0,000952651 | 0,021103066 | The Moose: suction sampler vs FISH | NA |
| gene_499011 | 4,176659638 | 5,021981155 | 1,717214453 | 2,924492714 | 0,003450182 | 0,045700744 | The Moose: suction sampler vs FISH | NA |
| gene_499967 | 22,53849378 | 3,023401038 | 0,979292266 | 3,087332703 | 0,002019614 | 0,0333627 | The Moose: suction sampler vs FISH | NA |

| | | | | | | | | |
|---|---|---|---|---|---|---|---|---|
| gene_504391 | 3,503989875 | 5,763122989 | 1,979207826 | 2,911833166 | 0,003593145 | 0,046815421 | The Moose: suction sampler vs FISH | NA |
| gene_504556 | 23,47134412 | 3,482605705 | 1,058354528 | 3,290585162 | 0,000999792 | 0,02173764 | The Moose: suction sampler vs FISH | NA |
| gene_505821 | 5,432272677 | -5,394079483 | 1,87518112 | -2,876564523 | 0,0040203 | 0,049935892 | The Moose: suction sampler vs FISH | NA |
| gene_506063 | 3,452824016 | 5,791067418 | 1,971898261 | 2,936798278 | 0,003316197 | 0,044714176 | The Moose: suction sampler vs FISH | NA |
| gene_506111 | 5,960522916 | 4,660851233 | 1,497573196 | 3,112269401 | 0,00185655 | 0,03174722 | The Moose: suction sampler vs FISH | NA |
| gene_509856 | 7,940411423 | 4,462759521 | 1,357786032 | 3,28679145 | 0,001013359 | 0,021896474 | The Moose: suction sampler vs FISH | NA |
| gene_510025 | 6,242909812 | 6,076338908 | 1,895796015 | 3,205164934 | 0,001349851 | 0,026116916 | The Moose: suction sampler vs FISH | NA |
| gene_510080 | 25,93550801 | 4,131010476 | 0,897951583 | 4,600482423 | 4,22E-06 | 0,000585219 | The Moose: suction sampler vs FISH | NA |
| gene_510262 | 7,322872406 | 6,821428737 | 1,680151153 | 4,060008962 | 4,91E-05 | 0,003123845 | The Moose: suction sampler vs FISH | NA |
| gene_510264 | 7,33653438 | 4,127355034 | 1,361481303 | 3,031517969 | 0,002433274 | 0,037131069 | The Moose: suction sampler vs FISH | NA |
| gene_510822 | 5,968425839 | 6,512233869 | 1,695865134 | 3,840065898 | 0,000123001 | 0,005746711 | The Moose: suction sampler vs FISH | NA |
| gene_51105 | 11,08317292 | 4,819325436 | 1,223443309 | 3,939148958 | 8,18E-05 | 0,004359781 | The Moose: suction sampler vs FISH | NA |
| gene_511627 | 22,02916368 | 6,76382119 | 1,271306865 | 5,320368649 | 1,04E-07 | 4,72E-05 | The Moose: suction sampler vs FISH | NA |
| gene_511872 | 3,88924581 | 6,78070512 | 2,12076324 | 3,197294725 | 0,001387231 | 0,026542102 | The Moose: suction sampler vs FISH | NA |
| gene_51239 | 13,39696386 | 4,273847608 | 1,139283068 | 3,751348306 | 0,000175886 | 0,007176644 | The Moose: suction sampler vs FISH | NA |
| gene_51240 | 10,25224308 | 3,574682151 | 1,207115382 | 2,961342557 | 0,003063011 | 0,042698629 | The Moose: suction sampler vs FISH | NA |
| gene_51241 | 31,55558471 | 3,42934335 | 1,002703148 | 3,420098317 | 0,000625985 | 0,01609222 | The Moose: suction sampler vs FISH | NA |
| gene_512613 | 16,81096346 | 8,41305034 | 1,60797122 | 5,232090123 | 1,68E-07 | 6,38E-05 | The Moose: suction sampler vs FISH | NA |
| gene_512937 | 85,5105057 | 4,99732835 | 1,03635124 | 4,822041174 | 1,42E-06 | 0,000279769 | The Moose: suction sampler vs FISH | NA |
| gene_513100 | 19,31934542 | 3,759906735 | 1,188041943 | 3,164792924 | 0,001551934 | 0,028459866 | The Moose: suction sampler vs FISH | NA |
| gene_51334 | 22,00001132 | 5,480951228 | 1,368482954 | 4,005129337 | 6,20E-05 | 0,003625765 | The Moose: suction sampler vs FISH | NA |
| gene_515635 | 51,48662527 | 5,171178434 | 1,104879523 | 4,680309777 | 2,86E-06 | 0,000439201 | The Moose: suction sampler vs FISH | NA |
| gene_516133 | 8,689279564 | 6,525044808 | 1,7028093 | 3,831929275 | 0,000127142 | 0,005839879 | The Moose: suction sampler vs FISH | NA |
| gene_516303 | 116,5221811 | 3,547101265 | 0,820704707 | 4,322018911 | 1,55E-05 | 0,001403607 | The Moose: suction sampler vs FISH | NA |
| gene_517201 | 44,18733127 | 3,085190912 | 0,878880268 | 3,510365433 | 0,000447491 | 0,013030222 | The Moose: suction sampler vs FISH | NA |
| gene_517565 | 40,98267454 | 10,62543965 | 3,600569809 | 2,951043922 | 0,003167019 | 0,043488543 | The Moose: suction sampler vs FISH | NA |
| gene_518705 | 109,827396 | 3,501958146 | 0,729222611 | 4,802317006 | 1,57E-06 | 0,000302009 | The Moose: suction sampler vs FISH | NA |
| gene_518785 | 267,8517601 | 3,607537431 | 0,815391237 | 4,424302428 | 9,68E-06 | 0,001034371 | The Moose: suction sampler vs FISH | NA |
| gene_51936 | 38,26194256 | 2,554754147 | 0,853626688 | 2,99282366 | 0,002764094 | 0,040036748 | The Moose: suction sampler vs FISH | NA |
| gene_520283 | 18,15228349 | 3,349488541 | 1,041381271 | 3,216390226 | 0,001298142 | 0,025541185 | The Moose: suction sampler vs FISH | NA |
| gene_520371 | 98,85309408 | 3,44695928 | 0,837917522 | 4,113721446 | 3,89E-05 | 0,002650086 | The Moose: suction sampler vs FISH | NA |
| gene_520923 | 3,662152141 | 6,147413427 | 1,925939967 | 3,191902932 | 0,001413388 | 0,026863397 | The Moose: suction sampler vs FISH | NA |
| gene_52194 | 4,351257935 | 5,895869208 | 2,035727765 | 2,896197276 | 0,003777149 | 0,048220526 | The Moose: suction sampler vs FISH | NA |
| gene_523871 | 4,2769653 | 6,222349834 | 1,779742044 | 3,496208822 | 0,000471919 | 0,013437829 | The Moose: suction sampler vs FISH | NA |
| gene_524171 | 4,826988441 | 6,975179571 | 1,883447956 | 3,703409775 | 0,000212721 | 0,008132891 | The Moose: suction sampler vs FISH | NA |
| gene_526607 | 7,590845324 | 4,874692021 | 1,447446991 | 3,367786215 | 0,000757743 | 0,018294693 | The Moose: suction sampler vs FISH | NA |
| gene_52686 | 7,238934789 | 3,792404564 | 1,235985258 | 3,06832508 | 0,002152623 | 0,034648127 | The Moose: suction sampler vs FISH | NA |
| gene_529215 | 4,730780604 | 7,377483677 | 2,041013559 | 3,614617671 | 0,000300791 | 0,010218135 | The Moose: suction sampler vs FISH | NA |
| gene_529650 | 8,609211324 | 5,596236372 | 1,558931822 | 3,589789042 | 0,000330946 | 0,010831687 | The Moose: suction sampler vs FISH | NA |
| gene_52980 | 9,640433983 | -3,532450985 | 1,223495204 | -2,887180084 | 0,003887117 | 0,048991899 | The Moose: suction sampler vs FISH | NA |
| gene_529855 | 15,37547325 | 4,022728992 | 1,12057501 | 3,589879263 | 0,000330831 | 0,010831687 | The Moose: suction sampler vs FISH | NA |
| gene_531403 | 13,67175663 | 3,002686421 | 1,025327486 | 2,92851451 | 0,003405859 | 0,045337352 | The Moose: suction sampler vs FISH | NA |
| gene_53154 | 96,6496572 | 4,227379541 | 0,883101079 | 4,786971322 | 1,69E-06 | 0,000311674 | The Moose: suction sampler vs FISH | NA |
| gene_531569 | 7,153076211 | 5,030198983 | 1,510008845 | 3,331238092 | 0,000864606 | 0,019979664 | The Moose: suction sampler vs FISH | NA |
| gene_531572 | 271,9987168 | 4,285734556 | 0,812443834 | 5,275114877 | 1,33E-07 | 5,47E-05 | The Moose: suction sampler vs FISH | NA |
| gene_531649 | 7,9168483 | 5,610925119 | 1,579091 | 3,55326268 | 0,000380484 | 0,01181209 | The Moose: suction sampler vs FISH | NA |
| gene_532092 | 16,83162995 | 3,524610588 | 1,14729301 | 3,072110227 | 0,002125512 | 0,034415398 | The Moose: suction sampler vs FISH | NA |
| gene_53312 | 14,60337119 | 3,356795689 | 1,162495008 | 2,887578584 | 0,003882196 | 0,048963228 | The Moose: suction sampler vs FISH | NA |
| gene_533120 | 12,30512178 | 3,730197984 | 1,126235823 | 3,312093176 | 0,000926007 | 0,02076659 | The Moose: suction sampler vs FISH | NA |
| gene_534126 | 5,457159661 | 5,143746477 | 1,6938693 | 3,036684399 | 0,002391958 | 0,036698835 | The Moose: suction sampler vs FISH | NA |
| gene_534196 | 35,99935564 | 2,744273677 | 0,861405327 | 3,185809968 | 0,001443494 | 0,027232994 | The Moose: suction sampler vs FISH | NA |
| gene_537110 | 3,676945131 | 6,813446842 | 2,231170965 | 3,053753813 | 0,002259975 | 0,035527361 | The Moose: suction sampler vs FISH | NA |
| gene_537544 | 8,691798427 | 6,922229369 | 1,660056065 | 4,169876857 | 3,05E-05 | 0,002238341 | The Moose: suction sampler vs FISH | NA |
| gene_53761 | 6,948248075 | 6,175043072 | 1,840443611 | 3,355192757 | 0,000793097 | 0,018820359 | The Moose: suction sampler vs FISH | NA |
| gene_539931 | 25,71754631 | -3,409809943 | 0,973233615 | -3,503588338 | 0,000459034 | 0,013262699 | The Moose: suction sampler vs FISH | NA |
| gene_539985 | 40,784928 | 3,383787823 | 0,958255557 | 3,531195617 | 0,000413686 | 0,01242854 | The Moose: suction sampler vs FISH | NA |
| gene_542090 | 13,27762075 | 4,870645747 | 1,48848708 | 3,27221231 | 0,001067094 | 0,022683339 | The Moose: suction sampler vs FISH | NA |
| gene_542284 | 6,816768916 | 7,356172661 | 1,870635189 | 3,932446426 | 8,41E-05 | 0,004452176 | The Moose: suction sampler vs FISH | NA |
| gene_543690 | 9,301881465 | 3,832134338 | 1,215530223 | 3,152644226 | 0,001617989 | 0,029225655 | The Moose: suction sampler vs FISH | NA |
| gene_544309 | 7,110548 | 4,25953235 | 1,421337889 | 2,996847113 | 0,002727875 | 0,039728092 | The Moose: suction sampler vs FISH | NA |
| gene_544464 | 4,815900154 | 5,638734643 | 1,70171491 | 3,313560109 | 0,000921163 | 0,020702209 | The Moose: suction sampler vs FISH | NA |
| gene_544825 | 14,57104273 | 4,385294292 | 1,433099876 | 3,060006051 | 0,002213325 | 0,035185643 | The Moose: suction sampler vs FISH | NA |
| gene_545284 | 10,54873906 | 3,284284185 | 1,008647059 | 3,256128251 | 0,001129427 | 0,023472208 | The Moose: suction sampler vs FISH | NA |
| gene_545779 | 15,18595295 | 4,205583509 | 1,097439642 | 3,832177505 | 0,000127014 | 0,005837638 | The Moose: suction sampler vs FISH | NA |
| gene_546389 | 151,7139881 | -2,007994502 | 0,649056597 | -3,09371249 | 0,001976689 | 0,032950058 | The Moose: suction sampler vs FISH | NA |

| | | | | | | | | |
|---|---|---|---|---|---|---|---|---|
| gene_546390 | 18,02840165 | -2,942004856 | 0,963128192 | -3,054634764 | 0,002253348 | 0,035453537 | The Moose: suction sampler vs FISH | NA |
| gene_546477 | 9,118075937 | 3,762784712 | 1,24394972 | 3,024868812 | 0,00248741 | 0,037690906 | The Moose: suction sampler vs FISH | NA |
| gene_546803 | 39,15228222 | 4,515591281 | 0,902389738 | 5,004036603 | 5,61E-07 | 0,000148858 | The Moose: suction sampler vs FISH | NA |
| gene_547896 | 40,09215606 | 2,554663071 | 0,842927603 | 3,030702827 | 0,002439853 | 0,03717118 | The Moose: suction sampler vs FISH | NA |
| gene_548754 | 2,985208756 | 6,021347819 | 2,003286483 | 3,005734762 | 0,0026494 | 0,039038312 | The Moose: suction sampler vs FISH | NA |
| gene_549562 | 108,1612244 | 11,11482765 | 1,635477224 | 6,796076083 | 1,08E-11 | 8,83E-08 | The Moose: suction sampler vs FISH | NA |
| gene_549719 | 84,66178022 | 2,482145447 | 0,784851184 | 3,162568263 | 0,001563841 | 0,028599671 | The Moose: suction sampler vs FISH | NA |
| gene_550107 | 27,28567726 | 2,928800427 | 0,932501218 | 3,140800645 | 0,001684867 | 0,029972233 | The Moose: suction sampler vs FISH | NA |
| gene_552393 | 23,08468109 | 3,643148698 | 0,938279648 | 3,88279625 | 0,000103262 | 0,005093408 | The Moose: suction sampler vs FISH | NA |
| gene_554579 | 9,473051047 | 4,040542913 | 1,157196339 | 3,491665828 | 0,000480018 | 0,013577723 | The Moose: suction sampler vs FISH | NA |
| gene_555520 | 7,930690904 | -5,353075344 | 1,647739584 | -3,248738694 | 0,001159179 | 0,023840915 | The Moose: suction sampler vs FISH | NA |
| gene_555733 | 52,52041791 | 3,68722183 | 0,8828108 | 4,176684097 | 2,96E-05 | 0,002187739 | The Moose: suction sampler vs FISH | NA |
| gene_556601 | 13,05883831 | 8,935542046 | 1,780037014 | 5,019863057 | 5,17E-07 | 0,00014331 | The Moose: suction sampler vs FISH | NA |
| gene_557228 | 55,95823056 | 3,290304225 | 0,797311674 | 4,126747836 | 3,68E-05 | 0,002539716 | The Moose: suction sampler vs FISH | NA |
| gene_55734 | 5,486168184 | 5,232860209 | 1,775036664 | 2,948029365 | 0,003198067 | 0,043764118 | The Moose: suction sampler vs FISH | NA |
| gene_557871 | 23,47771441 | 5,443325275 | 0,964639711 | 5,642858376 | 1,67E-08 | 1,28E-05 | The Moose: suction sampler vs FISH | NA |
| gene_558191 | 40,37027334 | 3,157482014 | 0,847726609 | 3,724646577 | 0,000195589 | 0,007706565 | The Moose: suction sampler vs FISH | NA |
| gene_558192 | 136,6484116 | 2,119891521 | 0,728779707 | 2,908823478 | 0,003627916 | 0,047077827 | The Moose: suction sampler vs FISH | NA |
| gene_558812 | 9,193746233 | 3,210293086 | 1,096571695 | 2,927572452 | 0,003416195 | 0,045400743 | The Moose: suction sampler vs FISH | NA |
| gene_5596 | 359,1233997 | -2,279048566 | 0,611298875 | -3,728206705 | 0,000192847 | 0,007640908 | The Moose: suction sampler vs FISH | NA |
| gene_559747 | 5,5620685 | 6,015710249 | 1,747018963 | 3,443414396 | 0,000574418 | 0,015255253 | The Moose: suction sampler vs FISH | NA |
| gene_561238 | 3,164742103 | 6,094405177 | 1,98168318 | 3,075368071 | 0,002102429 | 0,03419998 | The Moose: suction sampler vs FISH | NA |
| gene_561576 | 26,01986748 | 4,609940474 | 1,242726418 | 3,709537679 | 0,000207638 | 0,008014781 | The Moose: suction sampler vs FISH | NA |
| gene_562012 | 29,11923938 | 3,94034403 | 1,228241296 | 3,20811883 | 0,001336063 | 0,025938973 | The Moose: suction sampler vs FISH | NA |
| gene_562203 | 4,112198567 | 6,048470659 | 1,94620636 | 3,107825965 | 0,00188469 | 0,032041914 | The Moose: suction sampler vs FISH | NA |
| gene_562401 | 11,78481666 | 4,381692899 | 1,264944322 | 3,463941314 | 0,000532323 | 0,014507026 | The Moose: suction sampler vs FISH | NA |
| gene_56272 | 21,85676443 | 3,299817531 | 0,939039919 | 3,514033284 | 0,000441358 | 0,012945488 | The Moose: suction sampler vs FISH | NA |
| gene_56280 | 10,59674156 | 5,616816749 | 1,603507798 | 3,502830952 | 0,000460342 | 0,013284787 | The Moose: suction sampler vs FISH | NA |
| gene_564263 | 5,91746724 | 6,861800406 | 1,884033944 | 3,64207897 | 0,000270445 | 0,009517412 | The Moose: suction sampler vs FISH | NA |
| gene_565077 | 6,055083066 | 6,905894774 | 1,982741354 | 3,483003347 | 0,000495822 | 0,01388321 | The Moose: suction sampler vs FISH | NA |
| gene_566653 | 2056,888965 | -2,110554919 | 0,627838992 | -3,361618101 | 0,000774872 | 0,018555749 | The Moose: suction sampler vs FISH | NA |
| gene_566657 | 392,9976741 | -2,289091117 | 0,522235158 | -4,383257397 | 1,17E-05 | 0,00118116 | The Moose: suction sampler vs FISH | NA |
| gene_567868 | 9,936273311 | 4,332168996 | 1,193942013 | 3,628458459 | 0,000285119 | 0,00985901 | The Moose: suction sampler vs FISH | NA |
| gene_56822 | 5,517193048 | 5,462240604 | 1,765359203 | 3,094124185 | 0,001973948 | 0,032934267 | The Moose: suction sampler vs FISH | NA |
| gene_569082 | 9,194139527 | 4,1946534 | 1,282987151 | 3,269443032 | 0,001077594 | 0,02281462 | The Moose: suction sampler vs FISH | NA |
| gene_56948 | 7,904177438 | 4,344896111 | 1,38094108 | 3,146329828 | 0,001653334 | 0,029631319 | The Moose: suction sampler vs FISH | NA |
| gene_569562 | 17,28474223 | 3,6811139 | 1,117540726 | 3,293941611 | 0,00098793 | 0,021549906 | The Moose: suction sampler vs FISH | NA |
| gene_569926 | 15,10092555 | 3,55445406 | 1,017611757 | 3,492937296 | 0,000477739 | 0,013565667 | The Moose: suction sampler vs FISH | NA |
| gene_570420 | 18,22550459 | 3,515896221 | 1,00518275 | 3,497768164 | 0,000469169 | 0,013392145 | The Moose: suction sampler vs FISH | NA |
| gene_571115 | 46,10272063 | 2,587220741 | 0,855099134 | 3,025638361 | 0,002481089 | 0,037633943 | The Moose: suction sampler vs FISH | NA |
| gene_571442 | 555,3431477 | -2,107331821 | 0,664488582 | -3,171358963 | 0,001517275 | 0,028090819 | The Moose: suction sampler vs FISH | NA |
| gene_571776 | 78,9665966 | 4,178142653 | 0,848773184 | 4,922566747 | 8,54E-07 | 0,000201717 | The Moose: suction sampler vs FISH | NA |
| gene_57254 | 4,007845641 | 5,779679335 | 1,912899502 | 3,021423409 | 0,002515893 | 0,037936859 | The Moose: suction sampler vs FISH | NA |
| gene_572868 | 10,60766911 | -3,155300597 | 1,066080511 | -2,959720737 | 0,00307918 | 0,042855864 | The Moose: suction sampler vs FISH | NA |
| gene_576414 | 41,26778079 | 2,899226447 | 0,854943645 | 3,391131644 | 0,000696047 | 0,017235719 | The Moose: suction sampler vs FISH | NA |
| gene_578258 | 90,77839517 | 3,180595023 | 0,83217445 | 3,822029172 | 0,000132358 | 0,005985859 | The Moose: suction sampler vs FISH | NA |
| gene_578270 | 6,847688613 | 7,771071298 | 2,091208136 | 3,716067838 | 0,000202347 | 0,007879849 | The Moose: suction sampler vs FISH | NA |
| gene_578557 | 352,5433856 | 2,997357035 | 0,784363087 | 3,821389715 | 0,000132702 | 0,005997711 | The Moose: suction sampler vs FISH | NA |
| gene_579221 | 6,170974041 | 6,412118359 | 1,65520933 | 3,873901774 | 0,000107107 | 0,005218697 | The Moose: suction sampler vs FISH | NA |
| gene_580462 | 125,9327496 | 3,283500212 | 0,940382874 | 3,491663133 | 0,000480023 | 0,013577723 | The Moose: suction sampler vs FISH | NA |
| gene_582844 | 12,5322671 | 3,614026552 | 1,216202374 | 2,971566764 | 0,002962844 | 0,04180713 | The Moose: suction sampler vs FISH | NA |
| gene_584506 | 4,531812702 | 5,20610829 | 1,710749099 | 3,043174649 | 0,002340964 | 0,036228822 | The Moose: suction sampler vs FISH | NA |
| gene_584623 | 15,48009577 | 3,825353999 | 1,129378494 | 3,387131967 | 0,000706274 | 0,017424353 | The Moose: suction sampler vs FISH | NA |
| gene_584764 | 8,909402492 | 4,845938093 | 1,41582246 | 3,422701807 | 0,000620021 | 0,015989259 | The Moose: suction sampler vs FISH | NA |
| gene_58510 | 44,81219481 | 4,135441333 | 0,884835624 | 4,673683134 | 2,96E-06 | 0,000447086 | The Moose: suction sampler vs FISH | NA |
| gene_58511 | 20,58631878 | 4,578460681 | 1,121387395 | 4,082853704 | 4,45E-05 | 0,002917852 | The Moose: suction sampler vs FISH | NA |
| gene_585902 | 10,79201179 | 8,782907403 | 1,704939959 | 5,151446745 | 2,58E-07 | 8,79E-05 | The Moose: suction sampler vs FISH | NA |
| gene_586807 | 29,11144941 | 2,394491837 | 0,816842207 | 2,931400724 | 0,003374372 | 0,045068331 | The Moose: suction sampler vs FISH | NA |
| gene_587651 | 6,364207859 | 6,916396602 | 1,745833721 | 3,961658271 | 7,44E-05 | 0,004121578 | The Moose: suction sampler vs FISH | NA |
| gene_587693 | 111,5355722 | 5,971994855 | 1,012155139 | 5,900276177 | 3,63E-09 | 4,23E-06 | The Moose: suction sampler vs FISH | NA |
| gene_588274 | 6,054725025 | 4,000380531 | 1,316156796 | 3,039440699 | 0,002370179 | 0,036509811 | The Moose: suction sampler vs FISH | NA |
| gene_58893 | 32,39604843 | 7,779765214 | 1,379706763 | 5,638709199 | 1,71E-08 | 1,30E-05 | The Moose: suction sampler vs FISH | NA |
| gene_588942 | 27,43521475 | 3,329620146 | 0,845824686 | 3,93653697 | 8,27E-05 | 0,004397236 | The Moose: suction sampler vs FISH | NA |
| gene_589884 | 75,69140101 | 2,89134336 | 0,734806463 | 3,934836592 | 8,33E-05 | 0,00441802 | The Moose: suction sampler vs FISH | NA |
| gene_591145 | 5,546584656 | 4,62105501 | 1,590169969 | 2,906013256 | 0,003660659 | 0,047379514 | The Moose: suction sampler vs FISH | NA |

| | | | | | | | | |
|---|---|---|---|---|---|---|---|---|
| gene_591796 | 24,39227783 | 3,585277817 | 0,915300296 | 3,917050867 | 8,96E-05 | 0,004662553 | The Moose: suction sampler vs FISH | NA |
| gene_59197 | 127,4452359 | 2,844689206 | 0,79016625 | 3,600114791 | 0,000318077 | 0,010580484 | The Moose: suction sampler vs FISH | NA |
| gene_591998 | 11,2579316 | 3,630190973 | 1,036507011 | 3,502331324 | 0,000461206 | 0,013299275 | The Moose: suction sampler vs FISH | NA |
| gene_593627 | 22,86708481 | 3,721881858 | 1,069844212 | 3,478900776 | 0,000503475 | 0,014027966 | The Moose: suction sampler vs FISH | NA |
| gene_593628 | 51,63232424 | 3,667397596 | 0,824232666 | 4,449468876 | 8,61E-06 | 0,000962933 | The Moose: suction sampler vs FISH | NA |
| gene_596380 | 7,550363116 | 6,443516096 | 1,660615299 | 3,880197959 | 0,000104371 | 0,005127467 | The Moose: suction sampler vs FISH | NA |
| gene_596871 | 7,084740309 | 7,983056642 | 1,797416369 | 4,441406443 | 8,94E-06 | 0,000984106 | The Moose: suction sampler vs FISH | NA |
| gene_598749 | 10,13945906 | 3,682966811 | 1,231512111 | 2,990605433 | 0,00278425 | 0,040269641 | The Moose: suction sampler vs FISH | NA |
| gene_59899 | 28,69914055 | 3,176863182 | 0,782570958 | 4,05952093 | 4,92E-05 | 0,003123845 | The Moose: suction sampler vs FISH | NA |
| gene_599563 | 4,135957605 | 6,18459978 | 1,964654265 | 3,14793289 | 0,001644294 | 0,029526944 | The Moose: suction sampler vs FISH | NA |
| gene_599565 | 10,27235495 | 4,202169029 | 1,313352622 | 3,199574096 | 0,001376308 | 0,026440739 | The Moose: suction sampler vs FISH | NA |
| gene_600725 | 39,1686247 | 3,864454873 | 1,199070816 | 3,222874597 | 0,001269111 | 0,025191854 | The Moose: suction sampler vs FISH | NA |
| gene_600726 | 196,9000636 | 3,673301081 | 0,844824595 | 4,348004431 | 1,37E-05 | 0,001305305 | The Moose: suction sampler vs FISH | NA |
| gene_601403 | 3,466744958 | 6,356439537 | 2,196036301 | 2,894505675 | 0,00379756 | 0,048355032 | The Moose: suction sampler vs FISH | NA |
| gene_601553 | 5,418880999 | 5,142679598 | 1,786593065 | 2,878484027 | 0,003995915 | 0,049776118 | The Moose: suction sampler vs FISH | NA |
| gene_601703 | 4,944086968 | 5,827537448 | 1,768513208 | 3,29516196 | 0,00098365 | 0,021507637 | The Moose: suction sampler vs FISH | NA |
| gene_601832 | 23,19769127 | 4,887194959 | 1,255637697 | 3,892201524 | 9,93E-05 | 0,00498108 | The Moose: suction sampler vs FISH | NA |
| gene_601833 | 3,395874094 | 6,554951529 | 2,246482194 | 2,917873797 | 0,00352427 | 0,046237938 | The Moose: suction sampler vs FISH | NA |
| gene_602540 | 13,72086264 | 6,533073562 | 1,765940277 | 3,699487263 | 0,000216035 | 0,008199858 | The Moose: suction sampler vs FISH | NA |
| gene_604451 | 31,64914857 | 3,8350355 | 0,85158349 | 4,503416926 | 6,69E-06 | 0,000817177 | The Moose: suction sampler vs FISH | NA |
| gene_605794 | 6,671152273 | 6,344044221 | 1,624834158 | 3,904425686 | 9,44E-05 | 0,004827305 | The Moose: suction sampler vs FISH | NA |
| gene_606043 | 6,13534513 | 3,759485263 | 1,298353206 | 2,895579758 | 0,003784588 | 0,048273549 | The Moose: suction sampler vs FISH | NA |
| gene_608119 | 183,0216924 | 3,065812874 | 0,709049514 | 4,323834675 | 1,53E-05 | 0,001395548 | The Moose: suction sampler vs FISH | NA |
| gene_608413 | 172,5406297 | 2,781066948 | 0,764544059 | 3,637549616 | 0,000275244 | 0,009617178 | The Moose: suction sampler vs FISH | NA |
| gene_609275 | 44,35988502 | 2,846311657 | 0,851280838 | 3,343563641 | 0,000827097 | 0,019358239 | The Moose: suction sampler vs FISH | NA |
| gene_609534 | 3,845200601 | 5,564803126 | 1,885728735 | 2,951009348 | 0,003167373 | 0,043488543 | The Moose: suction sampler vs FISH | NA |
| gene_609535 | 4,488564909 | 5,269665269 | 1,816534126 | 2,900944823 | 0,003720394 | 0,04785852 | The Moose: suction sampler vs FISH | NA |
| gene_61105 | 9,543380353 | 3,730069011 | 1,26340721 | 2,952388574 | 0,003153259 | 0,043383106 | The Moose: suction sampler vs FISH | NA |
| gene_615198 | 5,153721585 | 6,27885987 | 1,997213393 | 3,143810217 | 0,001667636 | 0,029771389 | The Moose: suction sampler vs FISH | NA |
| gene_615363 | 4,803453222 | 6,139099389 | 1,77151341 | 3,465454652 | 0,000529336 | 0,014473964 | The Moose: suction sampler vs FISH | NA |
| gene_616559 | 16,74736375 | 2,903964939 | 0,911531626 | 3,18580821 | 0,001443503 | 0,027232994 | The Moose: suction sampler vs FISH | NA |
| gene_617565 | 105,0767705 | 3,357429219 | 0,75141337 | 4,46815209 | 7,89E-06 | 0,000909061 | The Moose: suction sampler vs FISH | NA |
| gene_617566 | 13,61594831 | 3,205294483 | 1,01020489 | 3,172915233 | 0,001509166 | 0,028018373 | The Moose: suction sampler vs FISH | NA |
| gene_618280 | 43,40911581 | -2,163662194 | 0,718682641 | -3,010594763 | 0,002607366 | 0,038790953 | The Moose: suction sampler vs FISH | NA |
| gene_618980 | 45,78416414 | -3,109133999 | 1,043295145 | -2,980109716 | 0,002881452 | 0,041108821 | The Moose: suction sampler vs FISH | NA |
| gene_618981 | 46,06050852 | -3,67693971 | 0,971908873 | -3,783214468 | 0,000154816 | 0,006603051 | The Moose: suction sampler vs FISH | NA |
| gene_619560 | 8,59551688 | 5,474000612 | 1,601010849 | 3,419090268 | 0,000628309 | 0,016135014 | The Moose: suction sampler vs FISH | NA |
| gene_619861 | 6,977925386 | 5,7084179 | 1,545549427 | 3,693455415 | 0,000221227 | 0,008340883 | The Moose: suction sampler vs FISH | NA |
| gene_619862 | 6,121266566 | 5,496463822 | 1,657265966 | 3,316585229 | 0,000911248 | 0,020592991 | The Moose: suction sampler vs FISH | NA |
| gene_620086 | 30,56917067 | 2,609077916 | 0,810834073 | 3,217770446 | 0,001291912 | 0,0254655 | The Moose: suction sampler vs FISH | NA |
| gene_621280 | 22,8862547 | 3,073550774 | 0,99376094 | 3,092847234 | 0,001982462 | 0,033031282 | The Moose: suction sampler vs FISH | NA |
| gene_621632 | 21,96341375 | 3,53843796 | 1,139039464 | 3,106510417 | 0,001893096 | 0,032117917 | The Moose: suction sampler vs FISH | NA |
| gene_621992 | 3,906633407 | 6,229981262 | 1,943884211 | 3,204913763 | 0,00135103 | 0,026125951 | The Moose: suction sampler vs FISH | NA |
| gene_622714 | 9,20618587 | 3,888630772 | 1,213549898 | 3,204343537 | 0,001353708 | 0,026157095 | The Moose: suction sampler vs FISH | NA |
| gene_624114 | 6,03900131 | 4,445751933 | 1,521555546 | 2,921846622 | 0,003479628 | 0,045906469 | The Moose: suction sampler vs FISH | NA |
| gene_624490 | 59,27530603 | 2,475354595 | 0,846358167 | 2,924712838 | 0,003447742 | 0,045690982 | The Moose: suction sampler vs FISH | NA |
| gene_624887 | 66,48805552 | 11,40945284 | 2,117727109 | 5,387593514 | 7,14E-08 | 3,77E-05 | The Moose: suction sampler vs FISH | NA |
| gene_624889 | 28,90252271 | 7,538574538 | 1,33019997 | 5,667249064 | 1,45E-08 | 1,16E-05 | The Moose: suction sampler vs FISH | NA |
| gene_625691 | 64,93076963 | 5,404037318 | 0,972688054 | 5,555776384 | 2,76E-08 | 1,90E-05 | The Moose: suction sampler vs FISH | NA |
| gene_625838 | 13,07785289 | 3,461053062 | 0,993623313 | 3,483264749 | 0,000495338 | 0,013883037 | The Moose: suction sampler vs FISH | NA |
| gene_625850 | 30,48806257 | 2,731531566 | 0,876559364 | 3,116196894 | 0,001831999 | 0,031519688 | The Moose: suction sampler vs FISH | NA |
| gene_626096 | 27,37867042 | 3,068155592 | 0,885239802 | 3,465903345 | 0,000528453 | 0,014464425 | The Moose: suction sampler vs FISH | NA |
| gene_627716 | 3,167756408 | 6,41271457 | 2,166434073 | 2,960032179 | 0,003076069 | 0,042836916 | The Moose: suction sampler vs FISH | NA |
| gene_628470 | 3,544825965 | 6,505920072 | 1,951802785 | 3,333287627 | 0,000858262 | 0,019873068 | The Moose: suction sampler vs FISH | NA |
| gene_62866 | 27,02117156 | -3,001255359 | 0,953503266 | -3,147608893 | 0,001646118 | 0,029538022 | The Moose: suction sampler vs FISH | NA |
| gene_628822 | 29,90691607 | 3,565788154 | 0,880126089 | 4,051451487 | 5,09E-05 | 0,003200698 | The Moose: suction sampler vs FISH | NA |
| gene_629207 | 8,76916636 | 5,790648542 | 1,707459201 | 3,391383255 | 0,000695408 | 0,017231521 | The Moose: suction sampler vs FISH | NA |
| gene_629602 | 12,42805071 | 4,500811442 | 1,217191849 | 3,697700938 | 0,000217561 | 0,008227995 | The Moose: suction sampler vs FISH | NA |
| gene_629670 | 47,69464217 | 3,040891147 | 0,886445061 | 3,430433853 | 0,000602617 | 0,015689187 | The Moose: suction sampler vs FISH | NA |
| gene_630395 | 3,181216392 | 6,061740278 | 2,080767929 | 2,913222659 | 0,003577194 | 0,046665549 | The Moose: suction sampler vs FISH | NA |
| gene_630636 | 61,46294756 | 3,051614307 | 0,908584644 | 3,358646138 | 0,000783253 | 0,018693108 | The Moose: suction sampler vs FISH | NA |
| gene_630960 | 6,49285331 | 4,738837394 | 1,504803319 | 3,149140711 | 0,001637513 | 0,029448747 | The Moose: suction sampler vs FISH | NA |
| gene_630961 | 28,05888173 | 3,240677785 | 0,844866962 | 3,835725544 | 0,000125194 | 0,005797411 | The Moose: suction sampler vs FISH | NA |
| gene_631638 | 5,00709112 | 5,339499463 | 1,834093992 | 2,911246363 | 0,0035999 | 0,046853569 | The Moose: suction sampler vs FISH | NA |
| gene_631768 | 27,64770992 | 4,931871325 | 1,014963466 | 4,85916143 | 1,18E-06 | 0,000247371 | The Moose: suction sampler vs FISH | NA |

| | | | | | | | | |
|---|---|---|---|---|---|---|---|---|
| gene_63576 | 13,81890647 | 8,09766326 | 1,698639245 | 4,767147165 | 1,87E-06 | 0,000333092 | The Moose: suction sampler vs FISH | NA |
| gene_636118 | 21,54754346 | 4,785621157 | 1,097132058 | 4,361937217 | 1,29E-05 | 0,001256859 | The Moose: suction sampler vs FISH | NA |
| gene_636401 | 175,0640855 | 2,922998127 | 0,746966002 | 3,913160864 | 9,11E-05 | 0,004714962 | The Moose: suction sampler vs FISH | NA |
| gene_636402 | 191,3219939 | 2,909070701 | 0,742812727 | 3,916290871 | 8,99E-05 | 0,004669667 | The Moose: suction sampler vs FISH | NA |
| gene_636713 | 4,942473437 | 7,16993877 | 2,16794736 | 3,307247631 | 0,000942176 | 0,020969118 | The Moose: suction sampler vs FISH | NA |
| gene_637558 | 4,531907951 | 6,275685581 | 1,971793246 | 3,182730032 | 0,001458936 | 0,02742417 | The Moose: suction sampler vs FISH | NA |
| gene_638264 | 5,345350044 | 6,467392827 | 1,701092431 | 3,801905592 | 0,000143587 | 0,006311059 | The Moose: suction sampler vs FISH | NA |
| gene_639811 | 11,47234815 | 7,73658245 | 1,570494347 | 4,926208403 | 8,38E-07 | 0,000199814 | The Moose: suction sampler vs FISH | NA |
| gene_641035 | 5,6488943 | 4,945131074 | 1,572919927 | 3,143917875 | 0,001667022 | 0,029770449 | The Moose: suction sampler vs FISH | NA |
| gene_642398 | 17,77125925 | 3,07419957 | 0,949518479 | 3,237640592 | 0,001205225 | 0,024393974 | The Moose: suction sampler vs FISH | NA |
| gene_642795 | 2,909439589 | 6,020593781 | 2,018394689 | 2,982862476 | 0,002855663 | 0,040923737 | The Moose: suction sampler vs FISH | NA |
| gene_644284 | 27,42689675 | 4,432141988 | 1,249577048 | 3,546913729 | 0,000389772 | 0,011992803 | The Moose: suction sampler vs FISH | NA |
| gene_645177 | 9,71108254 | 3,446214263 | 1,072146532 | 3,214312747 | 0,001307572 | 0,025650268 | The Moose: suction sampler vs FISH | NA |
| gene_646514 | 4,860128716 | 5,826039148 | 1,931524671 | 3,01629031 | 0,002558881 | 0,038346672 | The Moose: suction sampler vs FISH | NA |
| gene_647335 | 48,25508534 | 6,88428094 | 1,911895311 | 3,6007625 | 0,000317285 | 0,010580484 | The Moose: suction sampler vs FISH | NA |
| gene_647862 | 115,2014558 | 3,363790339 | 0,934206839 | 3,600691193 | 0,000317372 | 0,010580484 | The Moose: suction sampler vs FISH | NA |
| gene_647999 | 5,48544473 | 5,344073556 | 1,757998852 | 3,039861801 | 0,002366867 | 0,036489204 | The Moose: suction sampler vs FISH | NA |
| gene_648848 | 16,11929263 | 4,843196511 | 1,318452526 | 3,673394692 | 0,000239349 | 0,008798305 | The Moose: suction sampler vs FISH | NA |
| gene_648850 | 12,27411882 | 4,301465502 | 1,392997635 | 3,087920176 | 0,002015626 | 0,033327815 | The Moose: suction sampler vs FISH | NA |
| gene_649699 | 6,022119858 | 6,336578716 | 1,748515069 | 3,623977184 | 0,000290107 | 0,009961161 | The Moose: suction sampler vs FISH | NA |
| gene_650534 | 521,8240309 | 3,022402756 | 0,832806852 | 3,62917614 | 0,000284327 | 0,00984091 | The Moose: suction sampler vs FISH | NA |
| gene_651237 | 26,32546687 | 4,173256845 | 1,053423253 | 3,961614509 | 7,44E-05 | 0,004121578 | The Moose: suction sampler vs FISH | NA |
| gene_651922 | 9,272275041 | 3,225502786 | 1,025958979 | 3,143890597 | 0,001667178 | 0,029770449 | The Moose: suction sampler vs FISH | NA |
| gene_652979 | 14,79177422 | 3,279936501 | 1,044773219 | 3,139376508 | 0,001693078 | 0,03005755 | The Moose: suction sampler vs FISH | NA |
| gene_652980 | 7,61688073 | 4,408739882 | 1,266495301 | 3,481055065 | 0,000499443 | 0,013957982 | The Moose: suction sampler vs FISH | NA |
| gene_653006 | 3,060388337 | 6,501149559 | 2,115122831 | 3,073651073 | 0,002114566 | 0,034313001 | The Moose: suction sampler vs FISH | NA |
| gene_654821 | 3,46240784 | 6,447311319 | 2,113569719 | 3,050437022 | 0,002285086 | 0,035723316 | The Moose: suction sampler vs FISH | NA |
| gene_657288 | 29,63827097 | 2,328577782 | 0,740854208 | 3,143098543 | 0,001671696 | 0,02981488 | The Moose: suction sampler vs FISH | NA |
| gene_658677 | 54,80202553 | 2,546350512 | 0,828682456 | 3,072769904 | 0,002120819 | 0,034354562 | The Moose: suction sampler vs FISH | NA |
| gene_658739 | 15,94891587 | 4,040902383 | 1,132181368 | 3,569129907 | 0,000358169 | 0,011358285 | The Moose: suction sampler vs FISH | NA |
| gene_658740 | 31,32123406 | 2,775001109 | 0,932745022 | 2,975090773 | 0,002929019 | 0,041507639 | The Moose: suction sampler vs FISH | NA |
| gene_658741 | 32,09256478 | 3,480304834 | 1,034960703 | 3,362741043 | 0,000771727 | 0,018492497 | The Moose: suction sampler vs FISH | NA |
| gene_658744 | 9,841198856 | 4,564455114 | 1,208679882 | 3,776397029 | 0,000159113 | 0,006747157 | The Moose: suction sampler vs FISH | NA |
| gene_658949 | 23,3836591 | 2,397597888 | 0,76611301 | 3,12956164 | 0,001750673 | 0,030666956 | The Moose: suction sampler vs FISH | NA |
| gene_661191 | 5,749868234 | 5,559142545 | 1,683715469 | 3,301711391 | 0,000960969 | 0,021202975 | The Moose: suction sampler vs FISH | NA |
| gene_663650 | 13,56953958 | 3,918705197 | 1,132461642 | 3,460342543 | 0,000539489 | 0,014626336 | The Moose: suction sampler vs FISH | NA |
| gene_664202 | 11,49503693 | 3,349727505 | 0,940874286 | 3,560228559 | 0,000370532 | 0,011600057 | The Moose: suction sampler vs FISH | NA |
| gene_664237 | 17,4559565 | 3,04091163 | 0,838375602 | 3,627147098 | 0,00028657 | 0,009885927 | The Moose: suction sampler vs FISH | NA |
| gene_66424 | 150,1068884 | 2,361446485 | 0,792174416 | 2,980967874 | 0,002873389 | 0,041065593 | The Moose: suction sampler vs FISH | NA |
| gene_666214 | 24,29396836 | 3,382394796 | 0,868363596 | 3,895136568 | 9,81E-05 | 0,004957458 | The Moose: suction sampler vs FISH | NA |
| gene_667052 | 7,53626144 | 6,407047604 | 1,673050315 | 3,829560621 | 0,000128372 | 0,005883033 | The Moose: suction sampler vs FISH | NA |
| gene_667343 | 16,05066364 | 4,10667765 | 1,207031489 | 3,402295373 | 0,000668224 | 0,016790179 | The Moose: suction sampler vs FISH | NA |
| gene_667591 | 12,62148829 | 3,701678883 | 1,280784735 | 2,890164741 | 0,0038504 | 0,048731896 | The Moose: suction sampler vs FISH | NA |
| gene_668040 | 12,30463602 | 4,290555963 | 1,137825934 | 3,770836852 | 0,000162701 | 0,006847895 | The Moose: suction sampler vs FISH | NA |
| gene_668229 | 6,414308599 | 5,319112447 | 1,631067701 | 3,261123032 | 0,001109719 | 0,023254434 | The Moose: suction sampler vs FISH | NA |
| gene_668681 | 26,75599246 | 3,772923104 | 1,211792346 | 3,113506301 | 0,001848786 | 0,031680238 | The Moose: suction sampler vs FISH | NA |
| gene_669979 | 262,8915028 | -4,141246018 | 0,7665857 | -5,402195758 | 6,58E-08 | 3,63E-05 | The Moose: suction sampler vs FISH | NA |
| gene_670459 | 52,19248919 | 4,047932712 | 0,952020464 | 4,251938759 | 2,12E-05 | 0,001733301 | The Moose: suction sampler vs FISH | NA |
| gene_670803 | 32,24673164 | 3,323943801 | 0,853848322 | 3,892897267 | 9,91E-05 | 0,00497608 | The Moose: suction sampler vs FISH | NA |
| gene_672684 | 55,12362694 | 2,715914452 | 0,862805734 | 3,147770519 | 0,001645208 | 0,029536128 | The Moose: suction sampler vs FISH | NA |
| gene_673393 | 4,706741203 | 5,610601995 | 1,900329779 | 2,95243597 | 0,003152774 | 0,043383106 | The Moose: suction sampler vs FISH | NA |
| gene_673396 | 9,514621798 | 5,863582295 | 1,563331095 | 3,75069767 | 0,000176343 | 0,007187307 | The Moose: suction sampler vs FISH | NA |
| gene_674894 | 5,643024769 | 5,553975744 | 1,803948042 | 3,078789197 | 0,002078437 | 0,033997951 | The Moose: suction sampler vs FISH | NA |
| gene_674946 | 57,72052819 | 2,763150245 | 0,808742513 | 3,41660071 | 0,000634082 | 0,016243515 | The Moose: suction sampler vs FISH | NA |
| gene_676498 | 10,69276885 | 5,705950371 | 1,362444579 | 4,188023835 | 2,81E-05 | 0,002135019 | The Moose: suction sampler vs FISH | NA |
| gene_676499 | 7,233859629 | 5,317712277 | 1,492487798 | 3,562985428 | 0,000366661 | 0,011518144 | The Moose: suction sampler vs FISH | NA |
| gene_676590 | 3,895862215 | 6,353181166 | 1,974781147 | 3,217157088 | 0,001294677 | 0,025485807 | The Moose: suction sampler vs FISH | NA |
| gene_676818 | 11,9974021 | 8,899910874 | 2,76282553 | 3,221307598 | 0,001276071 | 0,02528218 | The Moose: suction sampler vs FISH | NA |
| gene_677033 | 32,61037706 | -5,661824382 | 1,353673649 | -4,182562308 | 2,88E-05 | 0,002160197 | The Moose: suction sampler vs FISH | NA |
| gene_677482 | 14,39078896 | 3,048446173 | 0,965212936 | 3,158314668 | 0,001586842 | 0,028862207 | The Moose: suction sampler vs FISH | NA |
| gene_678789 | 19,74975322 | 4,676607857 | 1,050547834 | 4,451589644 | 8,52E-06 | 0,000961318 | The Moose: suction sampler vs FISH | NA |
| gene_679438 | 59,66722129 | 3,577954141 | 1,049002849 | 3,410814514 | 0,000647691 | 0,016467565 | The Moose: suction sampler vs FISH | NA |
| gene_681228 | 56,25839317 | 3,261316893 | 0,741681671 | 4,397192248 | 1,10E-05 | 0,001129178 | The Moose: suction sampler vs FISH | NA |
| gene_681333 | 5,833992381 | 6,157062991 | 1,930216311 | 3,189830568 | 0,001423562 | 0,026988521 | The Moose: suction sampler vs FISH | NA |
| gene_681655 | 30,45069351 | 5,273954968 | 1,147727779 | 4,595127055 | 4,32E-06 | 0,00059483 | The Moose: suction sampler vs FISH | NA |

| | | | | | | | | |
|---|---|---|---|---|---|---|---|---|
| gene_682912 | 6,703906892 | 4,185091301 | 1,382914809 | 3,0262828 | 0,002475806 | 0,037569337 | The Moose: suction sampler vs FISH | NA |
| gene_683859 | 17,40396167 | 9,367260376 | 1,69000503 | 5,542741121 | 2,98E-08 | 1,97E-05 | The Moose: suction sampler vs FISH | NA |
| gene_684212 | 5,525492429 | 6,802941759 | 1,884039374 | 3,610827806 | 0,000305221 | 0,010291574 | The Moose: suction sampler vs FISH | NA |
| gene_684213 | 25,84779629 | 5,187119322 | 1,10239748 | 4,705307675 | 2,53E-06 | 0,000410068 | The Moose: suction sampler vs FISH | NA |
| gene_685406 | 7,171154791 | 5,032914773 | 1,586384273 | 3,172569761 | 0,001510963 | 0,028032354 | The Moose: suction sampler vs FISH | NA |
| gene_687667 | 11,89317879 | 7,65972922 | 1,64333068 | 4,661100358 | 3,15E-06 | 0,000468563 | The Moose: suction sampler vs FISH | NA |
| gene_687668 | 12,07945531 | 6,850464721 | 1,547894546 | 4,425666294 | 9,61E-06 | 0,001029355 | The Moose: suction sampler vs FISH | NA |
| gene_688242 | 6,340297128 | 4,913409605 | 1,629630925 | 3,015044405 | 0,002569416 | 0,038435586 | The Moose: suction sampler vs FISH | NA |
| gene_690130 | 7,830404557 | 4,462054969 | 1,523591719 | 2,928642176 | 0,003404461 | 0,045330064 | The Moose: suction sampler vs FISH | NA |
| gene_692884 | 13,42443822 | 8,776529987 | 1,804031768 | 4,864953124 | 1,14E-06 | 0,000243718 | The Moose: suction sampler vs FISH | NA |
| gene_694953 | 15,53304171 | 3,798442264 | 1,211693749 | 3,134820386 | 0,001719594 | 0,030330345 | The Moose: suction sampler vs FISH | NA |
| gene_694954 | 11,19016993 | 3,941889174 | 1,099371616 | 3,585583907 | 0,000336325 | 0,010927321 | The Moose: suction sampler vs FISH | NA |
| gene_695198 | 30,39137652 | 3,3156737 | 0,789640976 | 4,198963582 | 2,68E-05 | 0,002062147 | The Moose: suction sampler vs FISH | NA |
| gene_696001 | 5,890217302 | 5,322430467 | 1,712443969 | 3,108090287 | 0,001883005 | 0,032028098 | The Moose: suction sampler vs FISH | NA |
| gene_696106 | 4,279826433 | -5,262581002 | 1,819092598 | -2,892970378 | 0,003816173 | 0,048474371 | The Moose: suction sampler vs FISH | NA |
| gene_69636 | 12,87152301 | 3,334817954 | 1,007468587 | 3,310096213 | 0,000932639 | 0,020870712 | The Moose: suction sampler vs FISH | NA |
| gene_696662 | 3,398247358 | 6,119947128 | 2,088351575 | 2,930515725 | 0,003383999 | 0,045129049 | The Moose: suction sampler vs FISH | NA |
| gene_696701 | 321,6045586 | 4,764574805 | 0,746837382 | 6,379668348 | 1,77E-10 | 5,01E-07 | The Moose: suction sampler vs FISH | NA |
| gene_698168 | 22,72693734 | 3,884067354 | 1,034319545 | 3,755190912 | 0,000173209 | 0,007114857 | The Moose: suction sampler vs FISH | NA |
| gene_699878 | 39,43015562 | 3,639523197 | 0,743401134 | 4,895772999 | 9,79E-07 | 0,000220298 | The Moose: suction sampler vs FISH | NA |
| gene_700039 | 21,29532788 | 4,338165985 | 0,996214917 | 4,354648694 | 1,33E-05 | 0,001287998 | The Moose: suction sampler vs FISH | NA |
| gene_700609 | 9,520142679 | 3,905866178 | 1,200509826 | 3,253506213 | 0,001139902 | 0,023607465 | The Moose: suction sampler vs FISH | NA |
| gene_700802 | 3,991432423 | 5,748408213 | 1,893330475 | 3,036135682 | 0,002396315 | 0,036742663 | The Moose: suction sampler vs FISH | NA |
| gene_701700 | 408,8762538 | 14,04329726 | 4,247115325 | 3,30654955 | 0,000944526 | 0,021002345 | The Moose: suction sampler vs FISH | NA |
| gene_701983 | 6,025682719 | 4,885626514 | 1,671534336 | 2,922839458 | 0,003468553 | 0,045817964 | The Moose: suction sampler vs FISH | NA |
| gene_702014 | 14,44413578 | 3,470404711 | 1,176360082 | 2,95012111 | 0,003176494 | 0,043590731 | The Moose: suction sampler vs FISH | NA |
| gene_703138 | 12,00043365 | 3,356414937 | 1,124650103 | 2,984408153 | 0,002841275 | 0,040815641 | The Moose: suction sampler vs FISH | NA |
| gene_703912 | 12,78644359 | 4,505541744 | 1,273897049 | 3,536817789 | 0,000404979 | 0,012270495 | The Moose: suction sampler vs FISH | NA |
| gene_704066 | 8,681383352 | 7,345827635 | 1,676733686 | 4,381034206 | 1,18E-05 | 0,001188374 | The Moose: suction sampler vs FISH | NA |
| gene_705517 | 156,5079918 | -2,445026772 | 0,67148598 | -3,641217904 | 0,000271351 | 0,009538157 | The Moose: suction sampler vs FISH | NA |
| gene_706213 | 18,42864292 | 4,353051904 | 1,024238288 | 4,250038254 | 2,14E-05 | 0,001740321 | The Moose: suction sampler vs FISH | NA |
| gene_707007 | 21,19728509 | 4,019304459 | 1,163489712 | 3,454525137 | 0,000551263 | 0,014825155 | The Moose: suction sampler vs FISH | NA |
| gene_707008 | 25,42203085 | 5,492337126 | 1,083101251 | 5,070936 | 3,96E-07 | 0,00011867 | The Moose: suction sampler vs FISH | NA |
| gene_707754 | 3,692384558 | 6,879282831 | 2,216392871 | 3,103819237 | 0,0019104 | 0,032281658 | The Moose: suction sampler vs FISH | NA |
| gene_708316 | 12,26214683 | 4,034266222 | 1,237913305 | 3,258924681 | 0,001118354 | 0,023337123 | The Moose: suction sampler vs FISH | NA |
| gene_708676 | 148,8855606 | 2,899783158 | 0,886797411 | 3,269949959 | 0,001075665 | 0,02279521 | The Moose: suction sampler vs FISH | NA |
| gene_710063 | 4,473544042 | 5,798966216 | 1,785975603 | 3,24694593 | 0,001166505 | 0,023920816 | The Moose: suction sampler vs FISH | NA |
| gene_71020 | 14,2703835 | 3,376761732 | 0,98578654 | 3,425449217 | 0,000613784 | 0,015895395 | The Moose: suction sampler vs FISH | NA |
| gene_71021 | 11,19604816 | 3,251199047 | 1,01426694 | 3,205466846 | 0,001348436 | 0,026096408 | The Moose: suction sampler vs FISH | NA |
| gene_711721 | 3,212521005 | 6,151231896 | 2,109612071 | 2,91581186 | 0,003547644 | 0,046420218 | The Moose: suction sampler vs FISH | NA |
| gene_7122 | 10,35512959 | -4,471909572 | 1,351577593 | -3,308659151 | 0,000937439 | 0,020914398 | The Moose: suction sampler vs FISH | NA |
| gene_71279 | 6,642289055 | 5,833130196 | 1,70488507 | 3,421421361 | 0,000622947 | 0,016036582 | The Moose: suction sampler vs FISH | NA |
| gene_713 | 16,94905257 | 3,746839446 | 1,077482238 | 3,477402516 | 0,000506297 | 0,014058873 | The Moose: suction sampler vs FISH | NA |
| gene_713424 | 12,37688426 | -6,22476328 | 1,574466895 | -3,953568856 | 7,70E-05 | 0,004210597 | The Moose: suction sampler vs FISH | NA |
| gene_713436 | 75,22256386 | -2,466225237 | 0,717791427 | -3,435852178 | 0,000590693 | 0,015494093 | The Moose: suction sampler vs FISH | NA |
| gene_714223 | 3,533373892 | 5,864830741 | 2,034777065 | 2,882296464 | 0,003947881 | 0,049430905 | The Moose: suction sampler vs FISH | NA |
| gene_714313 | 120,5451532 | 2,632019281 | 0,804163037 | 3,272992118 | 0,001064154 | 0,022654152 | The Moose: suction sampler vs FISH | NA |
| gene_714453 | 12,72922376 | 5,740421686 | 1,595718232 | 3,597390549 | 0,000321426 | 0,010629045 | The Moose: suction sampler vs FISH | NA |
| gene_716380 | 17,49456581 | 3,477747509 | 1,022448793 | 3,401390401 | 0,00067044 | 0,016834349 | The Moose: suction sampler vs FISH | NA |
| gene_716381 | 9,55464169 | 5,312852859 | 1,428736093 | 3,71856838 | 0,000200355 | 0,007833937 | The Moose: suction sampler vs FISH | NA |
| gene_716382 | 9,099321094 | 4,754791989 | 1,440110208 | 3,301686192 | 0,000961055 | 0,021202975 | The Moose: suction sampler vs FISH | NA |
| gene_716819 | 49,38349058 | 2,330230405 | 0,649925215 | 3,585382367 | 0,000336585 | 0,010927321 | The Moose: suction sampler vs FISH | NA |
| gene_718265 | 4,624696736 | 6,65599756 | 1,826437051 | 3,644252374 | 0,00026817 | 0,009464565 | The Moose: suction sampler vs FISH | NA |
| gene_720696 | 4,589953376 | 7,471472048 | 2,21555002 | 3,372287684 | 0,000745465 | 0,018092798 | The Moose: suction sampler vs FISH | NA |
| gene_721476 | 19,90551867 | 3,06995229 | 1,02996428 | 2,98063957 | 0,002876471 | 0,041085654 | The Moose: suction sampler vs FISH | NA |
| gene_722131 | 11,35692732 | 3,899347049 | 1,007270019 | 3,871203326 | 0,000108299 | 0,005250198 | The Moose: suction sampler vs FISH | NA |
| gene_724045 | 4,415265314 | 6,629765789 | 1,943611463 | 3,411055098 | 0,00064712 | 0,016462673 | The Moose: suction sampler vs FISH | NA |
| gene_724356 | 21,98638575 | 3,844075875 | 1,110535456 | 3,461461634 | 0,000537251 | 0,014592595 | The Moose: suction sampler vs FISH | NA |
| gene_724868 | 4,910379627 | -6,298251819 | 2,070780464 | -3,041486979 | 0,002354127 | 0,036361491 | The Moose: suction sampler vs FISH | NA |
| gene_725178 | 30,73562403 | 3,532027341 | 0,88152024 | 4,006745595 | 6,16E-05 | 0,003613435 | The Moose: suction sampler vs FISH | NA |
| gene_725842 | 24,57731697 | 4,310006819 | 1,018721081 | 4,230801639 | 2,33E-05 | 0,001854924 | The Moose: suction sampler vs FISH | NA |
| gene_727113 | 14,3134711 | 3,934323755 | 1,203716636 | 3,268480005 | 0,001081268 | 0,022866035 | The Moose: suction sampler vs FISH | NA |
| gene_727336 | 4,493848236 | 6,473495534 | 2,150985035 | 3,009549312 | 0,002616356 | 0,038835543 | The Moose: suction sampler vs FISH | NA |
| gene_727501 | 35,51265662 | -3,65715242 | 1,095986955 | -3,336857618 | 0,000847313 | 0,019705941 | The Moose: suction sampler vs FISH | NA |
| gene_727801 | 4,517046364 | 5,657817027 | 1,934368124 | 2,924891574 | 0,003445763 | 0,045679518 | The Moose: suction sampler vs FISH | NA |

| | | | | | | | | |
|---|---|---|---|---|---|---|---|---|
| gene_728017 | 57,66173351 | 5,329461526 | 0,929444783 | 5,734027046 | 9,81E-09 | 9,12E-06 | The Moose: suction sampler vs FISH | NA |
| gene_728084 | 15,97636185 | 5,737050016 | 1,788025609 | 3,208594992 | 0,001333852 | 0,025909756 | The Moose: suction sampler vs FISH | NA |
| gene_729187 | 4,725819603 | 6,602754198 | 1,907186885 | 3,462038381 | 0,000536101 | 0,014577533 | The Moose: suction sampler vs FISH | NA |
| gene_730286 | 33,98423966 | 3,384132191 | 0,98504531 | 3,435509166 | 0,000591441 | 0,015508182 | The Moose: suction sampler vs FISH | NA |
| gene_731794 | 19,67387963 | 6,873837474 | 1,535722671 | 4,475962754 | 7,61E-06 | 0,00088776 | The Moose: suction sampler vs FISH | NA |
| gene_732021 | 17,48785189 | 3,215085467 | 1,007426405 | 3,191384951 | 0,001415925 | 0,026885368 | The Moose: suction sampler vs FISH | NA |
| gene_73225 | 20,08282094 | 2,707587026 | 0,929167627 | 2,913991994 | 0,003568391 | 0,046600364 | The Moose: suction sampler vs FISH | NA |
| gene_732825 | 11,02544449 | 4,34860474 | 1,333298892 | 3,261537803 | 0,001108097 | 0,023247448 | The Moose: suction sampler vs FISH | NA |
| gene_732827 | 17,73022621 | 3,880543275 | 1,238517431 | 3,13321652 | 0,001729018 | 0,030426115 | The Moose: suction sampler vs FISH | NA |
| gene_736744 | 5,076705521 | 4,669578321 | 1,519087788 | 3,07393579 | 0,002112549 | 0,034288654 | The Moose: suction sampler vs FISH | NA |
| gene_737102 | 9,66723887 | 4,147496547 | 1,361193083 | 3,046956819 | 0,002311709 | 0,036009221 | The Moose: suction sampler vs FISH | NA |
| gene_738430 | 144,6944155 | 3,461122843 | 0,785752242 | 4,404852649 | 1,06E-05 | 0,001104344 | The Moose: suction sampler vs FISH | NA |
| gene_738432 | 46,25809396 | 3,337097164 | 0,939867193 | 3,550605009 | 0,000384347 | 0,011890627 | The Moose: suction sampler vs FISH | NA |
| gene_739374 | 22,21426291 | -4,724782737 | 1,52605889 | -3,09606842 | 0,001961051 | 0,032771193 | The Moose: suction sampler vs FISH | NA |
| gene_739375 | 115,2542431 | -4,36990859 | 1,000144328 | -4,369277981 | 1,25E-05 | 0,001225637 | The Moose: suction sampler vs FISH | NA |
| gene_740248 | 22,22194235 | 4,481724916 | 0,919653074 | 4,873277809 | 1,10E-06 | 0,000236834 | The Moose: suction sampler vs FISH | NA |
| gene_740927 | 862,54821 | 2,576104072 | 0,703739259 | 3,660594515 | 0,000251631 | 0,009077121 | The Moose: suction sampler vs FISH | NA |
| gene_741370 | 11,16659961 | 3,191271476 | 1,063500977 | 3,000722657 | 0,002693398 | 0,039429258 | The Moose: suction sampler vs FISH | NA |
| gene_741563 | 16,31379416 | 3,158054151 | 0,922300214 | 3,424106492 | 0,000616824 | 0,015934812 | The Moose: suction sampler vs FISH | NA |
| gene_742130 | 12,78580447 | 2,882414338 | 0,970057923 | 2,97138374 | 0,002964611 | 0,041815988 | The Moose: suction sampler vs FISH | NA |
| gene_74284 | 154,0335596 | 4,150105811 | 1,053460937 | 3,939496628 | 8,17E-05 | 0,004358275 | The Moose: suction sampler vs FISH | NA |
| gene_745503 | 91,09454477 | 4,309335245 | 0,910783777 | 4,731458061 | 2,23E-06 | 0,000372935 | The Moose: suction sampler vs FISH | NA |
| gene_745709 | 68,47988435 | 3,245767854 | 0,781635574 | 4,152533433 | 3,29E-05 | 0,002356761 | The Moose: suction sampler vs FISH | NA |
| gene_747045 | 2,892261505 | 5,989710501 | 2,065583529 | 2,899766781 | 0,003734404 | 0,047958263 | The Moose: suction sampler vs FISH | NA |
| gene_748611 | 163,8895733 | 4,187956654 | 0,774802242 | 5,405194289 | 6,47E-08 | 3,60E-05 | The Moose: suction sampler vs FISH | NA |
| gene_748739 | 107,9769202 | 8,487077255 | 2,558254041 | 3,317527156 | 0,000908181 | 0,02056145 | The Moose: suction sampler vs FISH | NA |
| gene_749637 | 9,711405363 | 7,847248055 | 1,56740574 | 5,006519915 | 5,54E-07 | 0,00014856 | The Moose: suction sampler vs FISH | NA |
| gene_750070 | 21,8566773 | 5,751728971 | 1,226193181 | 4,690720075 | 2,72E-06 | 0,000427245 | The Moose: suction sampler vs FISH | NA |
| gene_750162 | 13,85159521 | 2,694613808 | 0,92548807 | 2,911559745 | 0,003596291 | 0,0468273 | The Moose: suction sampler vs FISH | NA |
| gene_750508 | 19,18851806 | 3,958373706 | 1,122912934 | 3,525094055 | 0,000423332 | 0,012611664 | The Moose: suction sampler vs FISH | NA |
| gene_750747 | 6,617796984 | 4,723430668 | 1,507024462 | 3,134276044 | 0,001722787 | 0,030372082 | The Moose: suction sampler vs FISH | NA |
| gene_751431 | 14,65365365 | -4,513081012 | 1,518013131 | -2,973018427 | 0,002948867 | 0,041690887 | The Moose: suction sampler vs FISH | NA |
| gene_751642 | 18,978623 | -2,959932561 | 0,945803135 | -3,129544036 | 0,001750778 | 0,030666956 | The Moose: suction sampler vs FISH | NA |
| gene_75169 | 3,901298405 | 6,547283661 | 2,003221448 | 3,268377377 | 0,00108166 | 0,022867745 | The Moose: suction sampler vs FISH | NA |
| gene_752046 | 30,38327309 | 3,60295568 | 0,984449714 | 3,659867671 | 0,000252346 | 0,009085058 | The Moose: suction sampler vs FISH | NA |
| gene_752786 | 51,22799655 | 4,418333858 | 0,905059871 | 4,881813898 | 1,05E-06 | 0,000230452 | The Moose: suction sampler vs FISH | NA |
| gene_753366 | 9,85926874 | 5,727340307 | 1,393597496 | 4,109752152 | 3,96E-05 | 0,002686093 | The Moose: suction sampler vs FISH | NA |
| gene_753460 | 7,203731177 | 5,403554945 | 1,677421128 | 3,221346658 | 0,001275897 | 0,02528218 | The Moose: suction sampler vs FISH | NA |
| gene_755821 | 9,832093146 | 4,553090543 | 1,320788767 | 3,447251111 | 0,000566322 | 0,015130416 | The Moose: suction sampler vs FISH | NA |
| gene_756042 | 3,629886711 | 6,678737355 | 2,249374772 | 2,969152779 | 0,002986221 | 0,042023949 | The Moose: suction sampler vs FISH | NA |
| gene_75657 | 8,630170193 | 4,403669212 | 1,464332789 | 3,007287171 | 0,002635906 | 0,038961028 | The Moose: suction sampler vs FISH | NA |
| gene_75659 | 15,71592686 | 4,114437137 | 1,258335388 | 3,269746029 | 0,001076441 | 0,022801671 | The Moose: suction sampler vs FISH | NA |
| gene_756664 | 5,183875369 | 6,940159008 | 1,736625961 | 3,996346458 | 6,43E-05 | 0,00371426 | The Moose: suction sampler vs FISH | NA |
| gene_758076 | 5253,978765 | 3,746195436 | 0,780357573 | 4,800613931 | 1,58E-06 | 0,00030333 | The Moose: suction sampler vs FISH | NA |
| gene_75865 | 159,4401656 | 2,078961315 | 0,679494736 | 3,059569418 | 0,002216554 | 0,03520092 | The Moose: suction sampler vs FISH | NA |
| gene_758844 | 27,7868502 | 3,318681753 | 1,026023233 | 3,234509363 | 0,001218519 | 0,024539113 | The Moose: suction sampler vs FISH | NA |
| gene_759362 | 22,93823553 | 4,043988001 | 0,974024988 | 4,151831884 | 3,30E-05 | 0,002358711 | The Moose: suction sampler vs FISH | NA |
| gene_759621 | 204,9221351 | 13,0326883 | 4,247195857 | 3,068539511 | 0,002151079 | 0,034648127 | The Moose: suction sampler vs FISH | NA |
| gene_76056 | 6,053125775 | 4,489914642 | 1,469368173 | 3,055677075 | 0,00224553 | 0,035391185 | The Moose: suction sampler vs FISH | NA |
| gene_761129 | 65,59985187 | 2,933263345 | 0,781038836 | 3,755592182 | 0,000172932 | 0,007107441 | The Moose: suction sampler vs FISH | NA |
| gene_761659 | 8,005618509 | 8,217099581 | 1,847103113 | 4,44864151 | 8,64E-06 | 0,000963089 | The Moose: suction sampler vs FISH | NA |
| gene_762165 | 9,57414691 | 5,257315885 | 1,487943389 | 3,533276819 | 0,000410442 | 0,012380188 | The Moose: suction sampler vs FISH | NA |
| gene_763589 | 11,62505839 | 3,809984465 | 1,150841771 | 3,310606688 | 0,00093094 | 0,02083903 | The Moose: suction sampler vs FISH | NA |
| gene_763802 | 289,0962532 | 13,54492449 | 4,247148075 | 3,189181129 | 0,001426764 | 0,027035273 | The Moose: suction sampler vs FISH | NA |
| gene_766384 | 7,464451535 | 5,999115107 | 1,604454607 | 3,739036979 | 0,000184727 | 0,007413793 | The Moose: suction sampler vs FISH | NA |
| gene_767841 | 14,06832707 | -3,50401973 | 1,088463398 | -3,219235241 | 0,00128533 | 0,025390279 | The Moose: suction sampler vs FISH | NA |
| gene_767913 | 3,409881145 | 6,90340868 | 2,092902405 | 3,298485712 | 0,000972078 | 0,021343584 | The Moose: suction sampler vs FISH | NA |
| gene_769052 | 19,7834296 | 3,120807773 | 0,985013742 | 3,168288561 | 0,001533392 | 0,028296484 | The Moose: suction sampler vs FISH | NA |
| gene_769812 | 107,8128798 | 2,180607145 | 0,700920804 | 3,111060668 | 0,001864166 | 0,031840395 | The Moose: suction sampler vs FISH | NA |
| gene_771114 | 4,223157786 | 6,802856239 | 2,112234254 | 3,220692131 | 0,001278814 | 0,025289765 | The Moose: suction sampler vs FISH | NA |
| gene_771829 | 9,834339388 | 6,078361758 | 1,454942913 | 4,177732131 | 2,94E-05 | 0,002186829 | The Moose: suction sampler vs FISH | NA |
| gene_771838 | 4,789133165 | 5,296221529 | 1,765369009 | 3,000064861 | 0,002699221 | 0,039451602 | The Moose: suction sampler vs FISH | NA |
| gene_774044 | 13,07800423 | 8,452147253 | 1,804088057 | 4,684997067 | 2,80E-06 | 0,000434636 | The Moose: suction sampler vs FISH | NA |
| gene_775643 | 5,729731323 | 4,790824278 | 1,418432633 | 3,377547984 | 0,000731352 | 0,017863033 | The Moose: suction sampler vs FISH | NA |
| gene_776515 | 87,89430904 | 6,189761834 | 0,925114931 | 6,690803087 | 2,22E-11 | 1,36E-07 | The Moose: suction sampler vs FISH | NA |

| | | | | | | | | |
|---|---|---|---|---|---|---|---|---|
| gene_776859 | 27,52977164 | 4,624227929 | 0,92259775 | 5,012182103 | 5,38E-07 | 0,000145124 | The Moose: suction sampler vs FISH | NA |
| gene_777143 | 8,651278624 | 5,035487243 | 1,479837083 | 3,402730815 | 0,00066716 | 0,016780667 | The Moose: suction sampler vs FISH | NA |
| gene_777498 | 20,6649119 | 4,16898379 | 0,988398211 | 4,217919199 | 2,47E-05 | 0,001941519 | The Moose: suction sampler vs FISH | NA |
| gene_779709 | 9,665154983 | 4,609039054 | 1,570061992 | 2,935577753 | 0,003329272 | 0,044797555 | The Moose: suction sampler vs FISH | NA |
| gene_780462 | 22,11520263 | 3,376565866 | 0,885663522 | 3,812470291 | 0,000137585 | 0,006145279 | The Moose: suction sampler vs FISH | NA |
| gene_780781 | 43,47586401 | 2,605963716 | 0,834391387 | 3,123191055 | 0,001789016 | 0,0310258 | The Moose: suction sampler vs FISH | NA |
| gene_781399 | 4,584288266 | 7,220996677 | 1,942831719 | 3,716738103 | 0,000201811 | 0,007871985 | The Moose: suction sampler vs FISH | NA |
| gene_781589 | 12,32506717 | 5,856880773 | 1,418666978 | 4,12843949 | 3,65E-05 | 0,002528235 | The Moose: suction sampler vs FISH | NA |
| gene_78221 | 616,7563943 | 7,022968176 | 1,056180466 | 6,649401689 | 2,94E-11 | 1,54E-07 | The Moose: suction sampler vs FISH | NA |
| gene_782231 | 4,767498532 | 6,754120138 | 2,250348197 | 3,001366698 | 0,002687707 | 0,039361021 | The Moose: suction sampler vs FISH | NA |
| gene_784170 | 7,395466962 | 4,206462656 | 1,294842407 | 3,248629048 | 0,001159626 | 0,023841091 | The Moose: suction sampler vs FISH | NA |
| gene_784330 | 8,85361755 | 3,574467814 | 1,160588521 | 3,079875211 | 0,002070873 | 0,033919558 | The Moose: suction sampler vs FISH | NA |
| gene_785758 | 17,62566036 | 4,11081904 | 1,245139746 | 3,301492103 | 0,00096172 | 0,021211278 | The Moose: suction sampler vs FISH | NA |
| gene_785759 | 36,09336739 | 5,817838924 | 1,06741704 | 5,450389779 | 5,03E-08 | 2,98E-05 | The Moose: suction sampler vs FISH | NA |
| gene_785781 | 6,007932087 | 6,340718927 | 1,842885449 | 3,440647345 | 0,000580324 | 0,015337144 | The Moose: suction sampler vs FISH | NA |
| gene_786191 | 141,3848899 | 3,299255867 | 0,725241016 | 4,549185435 | 5,39E-06 | 0,000695133 | The Moose: suction sampler vs FISH | NA |
| gene_786330 | 23,85024469 | 3,633663424 | 0,923860149 | 3,933131467 | 8,38E-05 | 0,004446278 | The Moose: suction sampler vs FISH | NA |
| gene_786627 | 155,9574252 | 4,081942443 | 0,735133512 | 5,552654551 | 2,81E-08 | 1,91E-05 | The Moose: suction sampler vs FISH | NA |
| gene_786707 | 4,684259514 | 6,543946003 | 1,999213062 | 3,273260928 | 0,001063143 | 0,022639178 | The Moose: suction sampler vs FISH | NA |
| gene_788039 | 1280,253495 | 2,514646806 | 0,758816797 | 3,313905035 | 0,000920027 | 0,020695687 | The Moose: suction sampler vs FISH | NA |
| gene_788279 | 7,515137615 | 4,11848071 | 1,316120925 | 3,12925707 | 0,001752489 | 0,030666956 | The Moose: suction sampler vs FISH | NA |
| gene_788960 | 63,28006349 | -2,763326651 | 0,772903533 | -3,575254263 | 0,000349888 | 0,011177686 | The Moose: suction sampler vs FISH | NA |
| gene_788989 | 16,59498231 | 3,873592871 | 1,34618522 | 2,87745907 | 0,004008919 | 0,049863836 | The Moose: suction sampler vs FISH | NA |
| gene_788990 | 6,231098962 | 6,885320952 | 1,976576856 | 3,483457237 | 0,000494982 | 0,013880858 | The Moose: suction sampler vs FISH | NA |
| gene_790172 | 35,43227218 | 4,085421417 | 1,089459524 | 3,749952456 | 0,000176868 | 0,007204702 | The Moose: suction sampler vs FISH | NA |
| gene_7913 | 178,3544752 | 2,566677979 | 0,832526236 | 3,082999513 | 0,002049255 | 0,033685659 | The Moose: suction sampler vs FISH | NA |
| gene_79131 | 8,994641865 | 4,279510214 | 1,405115498 | 3,045664374 | 0,002321668 | 0,036082789 | The Moose: suction sampler vs FISH | NA |
| gene_792356 | 59,15818693 | 3,302310137 | 0,793805356 | 4,160100599 | 3,18E-05 | 0,00229503 | The Moose: suction sampler vs FISH | NA |
| gene_792703 | 16,89285267 | 2,974752238 | 1,024872757 | 2,902557627 | 0,003701291 | 0,047701033 | The Moose: suction sampler vs FISH | NA |
| gene_792994 | 17,45107437 | 4,398369944 | 1,197671942 | 3,672432983 | 0,000240252 | 0,008809446 | The Moose: suction sampler vs FISH | NA |
| gene_793466 | 17,84791011 | 4,117716568 | 0,987726546 | 4,168883164 | 3,06E-05 | 0,002243634 | The Moose: suction sampler vs FISH | NA |
| gene_793941 | 531,4802162 | 2,001494958 | 0,640431053 | 3,125230966 | 0,001776655 | 0,030868708 | The Moose: suction sampler vs FISH | NA |
| gene_794561 | 5,659903804 | 5,654486242 | 1,73936829 | 3,250884976 | 0,001150464 | 0,023741451 | The Moose: suction sampler vs FISH | NA |
| gene_795001 | 27,47285464 | 4,188962699 | 1,052446224 | 3,980215428 | 6,89E-05 | 0,003886931 | The Moose: suction sampler vs FISH | NA |
| gene_795190 | 7,050810836 | 4,33010014 | 1,396309058 | 3,101104382 | 0,001928003 | 0,032477567 | The Moose: suction sampler vs FISH | NA |
| gene_796188 | 16,7197759 | 2,910723953 | 0,966455874 | 3,011750491 | 0,00259746 | 0,038719393 | The Moose: suction sampler vs FISH | NA |
| gene_797061 | 8,458516808 | 6,911868915 | 1,833811454 | 3,769127354 | 0,000163819 | 0,006879193 | The Moose: suction sampler vs FISH | NA |
| gene_798600 | 8,898011972 | 4,442697759 | 1,437009031 | 3,09162828 | 0,00199062 | 0,033077161 | The Moose: suction sampler vs FISH | NA |
| gene_798725 | 16,24961159 | 8,70559487 | 1,701977922 | 5,114986956 | 3,14E-07 | 0,000101071 | The Moose: suction sampler vs FISH | NA |
| gene_799435 | 7,675114407 | 4,171379635 | 1,242279811 | 3,357842249 | 0,000785534 | 0,018701511 | The Moose: suction sampler vs FISH | NA |
| gene_799593 | 8,274672801 | 3,521667058 | 1,181632553 | 2,980340251 | 0,002879284 | 0,041108821 | The Moose: suction sampler vs FISH | NA |
| gene_801644 | 208,7085386 | 3,947908448 | 0,791203321 | 4,989752121 | 6,05E-07 | 0,000155254 | The Moose: suction sampler vs FISH | NA |
| gene_802217 | 25,54380904 | 3,484727479 | 0,94552915 | 3,685478632 | 0,000228273 | 0,008527735 | The Moose: suction sampler vs FISH | NA |
| gene_802701 | 28,33263724 | 6,82509542 | 1,25908437 | 5,420681557 | 5,94E-08 | 3,35E-05 | The Moose: suction sampler vs FISH | NA |
| gene_802703 | 54,72258107 | 7,119356883 | 1,110903963 | 6,408615977 | 1,47E-10 | 4,72E-07 | The Moose: suction sampler vs FISH | NA |
| gene_803524 | 27,32851328 | 2,358497536 | 0,808517678 | 2,917063659 | 0,003533436 | 0,046284899 | The Moose: suction sampler vs FISH | NA |
| gene_803998 | 8,833893975 | 5,030845506 | 1,434610756 | 3,506766895 | 0,000453586 | 0,013137818 | The Moose: suction sampler vs FISH | NA |
| gene_805123 | 4,42124159 | 6,009613431 | 1,898260633 | 3,165852637 | 0,001546291 | 0,028391837 | The Moose: suction sampler vs FISH | NA |
| gene_807191 | 15,47798635 | -3,94457927 | 1,251174806 | -3,152700366 | 0,001617678 | 0,029225655 | The Moose: suction sampler vs FISH | NA |
| gene_808621 | 107,430071 | 3,983617097 | 0,866556554 | 4,59706534 | 4,28E-06 | 0,000590431 | The Moose: suction sampler vs FISH | NA |
| gene_808978 | 29,46562826 | 4,325671932 | 0,984795685 | 4,392456219 | 1,12E-05 | 0,001148049 | The Moose: suction sampler vs FISH | NA |
| gene_812178 | 3,466406417 | 6,076281808 | 2,008736315 | 3,024927544 | 0,002486927 | 0,037690906 | The Moose: suction sampler vs FISH | NA |
| gene_814090 | 11,53369333 | 3,569213492 | 0,960454027 | 3,716173177 | 0,000202263 | 0,007879849 | The Moose: suction sampler vs FISH | NA |
| gene_815796 | 92,26465507 | 3,021650125 | 0,709523496 | 4,258703402 | 2,06E-05 | 0,001696795 | The Moose: suction sampler vs FISH | NA |
| gene_816455 | 19,38103035 | 3,134155566 | 0,914080708 | 3,428751464 | 0,000606365 | 0,015758826 | The Moose: suction sampler vs FISH | NA |
| gene_816848 | 4,149324847 | 4,827351982 | 1,669829874 | 2,890924433 | 0,003841105 | 0,048690013 | The Moose: suction sampler vs FISH | NA |
| gene_81709 | 29,90533387 | 5,51330876 | 1,303661929 | 4,229093938 | 2,35E-05 | 0,001865013 | The Moose: suction sampler vs FISH | NA |
| gene_817102 | 3,282788258 | 6,304282299 | 2,048782476 | 3,077087184 | 0,002090342 | 0,034092895 | The Moose: suction sampler vs FISH | NA |
| gene_817112 | 122,6991346 | 2,504764383 | 0,7693796 | 3,255563812 | 0,001131675 | 0,023505612 | The Moose: suction sampler vs FISH | NA |
| gene_818448 | 9,64778661 | 3,975304047 | 1,371561934 | 2,898377353 | 0,00375099 | 0,0480412 | The Moose: suction sampler vs FISH | NA |
| gene_819875 | 14,06465726 | 5,003508521 | 1,238048906 | 4,041446583 | 5,31E-05 | 0,003277638 | The Moose: suction sampler vs FISH | NA |
| gene_821809 | 9,682729435 | 6,242530611 | 1,553060019 | 4,019503777 | 5,83E-05 | 0,003492459 | The Moose: suction sampler vs FISH | NA |
| gene_822218 | 21,50652545 | 2,972243936 | 0,984801672 | 3,018114225 | 0,00254353 | 0,03821018 | The Moose: suction sampler vs FISH | NA |
| gene_822388 | 124,1180339 | -3,088857429 | 0,837089761 | -3,689995472 | 0,000224258 | 0,008416266 | The Moose: suction sampler vs FISH | NA |
| gene_823084 | 9,532907095 | 5,034989406 | 1,547258786 | 3,254135281 | 0,001137381 | 0,02358745 | The Moose: suction sampler vs FISH | NA |

| | | | | | | | | |
|---|---|---|---|---|---|---|---|---|
| gene_823681 | 36,66772796 | 3,970504725 | 0,893372616 | 4,444399409 | 8,81E-06 | 0,000973567 | The Moose: suction sampler vs FISH | NA |
| gene_823687 | 12,67257102 | 3,668330756 | 1,134609955 | 3,233120543 | 0,001224459 | 0,024577857 | The Moose: suction sampler vs FISH | NA |
| gene_824159 | 3,731972151 | 5,885417605 | 1,913404491 | 3,075887839 | 0,002098768 | 0,03417827 | The Moose: suction sampler vs FISH | NA |
| gene_824278 | 4,177666666 | 5,734007336 | 1,845547511 | 3,106941058 | 0,001890341 | 0,032093408 | The Moose: suction sampler vs FISH | NA |
| gene_824414 | 14,79780759 | 4,223981724 | 1,009699871 | 4,183403251 | 2,87E-05 | 0,002158827 | The Moose: suction sampler vs FISH | NA |
| gene_824841 | 4,689210685 | 5,67611279 | 1,790258091 | 3,170555585 | 0,001521477 | 0,028147328 | The Moose: suction sampler vs FISH | NA |
| gene_824842 | 5,481672676 | 5,809133167 | 1,699237284 | 3,418670965 | 0,000629278 | 0,016148603 | The Moose: suction sampler vs FISH | NA |
| gene_824903 | 16,95310972 | 3,32431299 | 0,902412604 | 3,683806026 | 0,000229777 | 0,0085578 | The Moose: suction sampler vs FISH | NA |
| gene_824988 | 6,868193519 | 5,144812514 | 1,608121015 | 3,199269498 | 0,001377763 | 0,026458729 | The Moose: suction sampler vs FISH | NA |
| gene_825334 | 46,8404981 | 3,981193052 | 1,166841586 | 3,411939634 | 0,000645024 | 0,016426416 | The Moose: suction sampler vs FISH | NA |
| gene_827422 | 94,70884713 | 4,372761567 | 0,998300541 | 4,380205548 | 1,19E-05 | 0,001189643 | The Moose: suction sampler vs FISH | NA |
| gene_827526 | 3,747131594 | 7,200980819 | 2,318374341 | 3,106047497 | 0,001896063 | 0,032160812 | The Moose: suction sampler vs FISH | NA |
| gene_82757 | 11,6848615 | 4,786622323 | 1,178246254 | 4,062497382 | 4,86E-05 | 0,003100688 | The Moose: suction sampler vs FISH | NA |
| gene_828015 | 7,556744249 | 4,714652885 | 1,408267091 | 3,347839991 | 0,00081444 | 0,019159693 | The Moose: suction sampler vs FISH | NA |
| gene_828016 | 7,112685651 | 5,58251578 | 1,551068854 | 3,599141175 | 0,00031927 | 0,010599955 | The Moose: suction sampler vs FISH | NA |
| gene_828337 | 4,880220157 | 6,505750003 | 2,195176225 | 2,963657281 | 0,003040067 | 0,042472454 | The Moose: suction sampler vs FISH | NA |
| gene_828338 | 10,50804903 | 6,852062016 | 1,989867309 | 3,443476851 | 0,000574286 | 0,015255253 | The Moose: suction sampler vs FISH | NA |
| gene_828537 | 5,117133728 | 4,980705758 | 1,556321553 | 3,200306355 | 0,001372816 | 0,026394361 | The Moose: suction sampler vs FISH | NA |
| gene_82986 | 8,639024475 | 3,763238656 | 1,306356791 | 2,880712745 | 0,003967771 | 0,049588452 | The Moose: suction sampler vs FISH | NA |
| gene_830478 | 6,809812431 | 6,781486567 | 1,667923835 | 4,065825084 | 4,79E-05 | 0,003078183 | The Moose: suction sampler vs FISH | NA |
| gene_831556 | 16,51176036 | 9,250115594 | 2,628183875 | 3,519584638 | 0,000432223 | 0,012764372 | The Moose: suction sampler vs FISH | NA |
| gene_831585 | 15,00049074 | 3,309270718 | 1,097696508 | 3,014741046 | 0,002571987 | 0,038456763 | The Moose: suction sampler vs FISH | NA |
| gene_832745 | 42,68748577 | 3,455528579 | 0,838364796 | 4,121748187 | 3,76E-05 | 0,00258576 | The Moose: suction sampler vs FISH | NA |
| gene_833688 | 5,681647768 | 4,527524015 | 1,560915703 | 2,900556389 | 0,003725008 | 0,047887838 | The Moose: suction sampler vs FISH | NA |
| gene_834181 | 6,055233484 | 5,00009675 | 1,506275818 | 3,319509409 | 0,000901758 | 0,02049231 | The Moose: suction sampler vs FISH | NA |
| gene_835033 | 15,64444977 | 3,810041113 | 1,01663271 | 3,747706595 | 0,000178459 | 0,007226628 | The Moose: suction sampler vs FISH | NA |
| gene_835786 | 13,95490727 | 3,064471555 | 0,961483906 | 3,187231253 | 0,001436419 | 0,027141183 | The Moose: suction sampler vs FISH | NA |
| gene_837919 | 12,79028856 | 3,170925423 | 1,056997669 | 2,999936059 | 0,002700363 | 0,039460436 | The Moose: suction sampler vs FISH | NA |
| gene_839395 | 42,6680075 | 2,748395801 | 0,849811861 | 3,234122665 | 0,00122017 | 0,024539704 | The Moose: suction sampler vs FISH | NA |
| gene_84076 | 31,76628417 | 3,671671346 | 0,926051429 | 3,964867643 | 7,34E-05 | 0,004095336 | The Moose: suction sampler vs FISH | NA |
| gene_84123 | 3,36937863 | 6,157938755 | 1,952131874 | 3,154468629 | 0,001607907 | 0,029115558 | The Moose: suction sampler vs FISH | NA |
| gene_842232 | 9,731382353 | 3,077921244 | 1,053316909 | 2,922122694 | 0,003476546 | 0,045883541 | The Moose: suction sampler vs FISH | NA |
| gene_842460 | 13,82188335 | 3,283717269 | 1,119706649 | 2,932658541 | 0,003360733 | 0,044979901 | The Moose: suction sampler vs FISH | NA |
| gene_84292 | 5,320709679 | 6,687512117 | 2,155326858 | 3,10278327 | 0,0019171 | 0,032323552 | The Moose: suction sampler vs FISH | NA |
| gene_84440 | 6,897769004 | 5,17993982 | 1,557605276 | 3,325579273 | 0,00088235 | 0,02022603 | The Moose: suction sampler vs FISH | NA |
| gene_845203 | 28,48808504 | -3,425305678 | 1,117876725 | -3,064117538 | 0,002183131 | 0,034917263 | The Moose: suction sampler vs FISH | NA |
| gene_845209 | 30,65854959 | 9,940324207 | 2,553038666 | 3,893526698 | 9,88E-05 | 0,00497608 | The Moose: suction sampler vs FISH | NA |
| gene_846813 | 49,13060149 | 3,086484737 | 0,989827997 | 3,118203109 | 0,001819574 | 0,031351249 | The Moose: suction sampler vs FISH | NA |
| gene_848046 | 17,76522857 | 3,154789004 | 1,017533473 | 3,100427737 | 0,001932414 | 0,032529479 | The Moose: suction sampler vs FISH | NA |
| gene_848135 | 4,927949161 | 6,87700866 | 1,782169224 | 3,858785443 | 0,000113952 | 0,005442151 | The Moose: suction sampler vs FISH | NA |
| gene_848136 | 8,368448997 | 5,14728584 | 1,576985136 | 3,26400403 | 0,001098496 | 0,023110582 | The Moose: suction sampler vs FISH | NA |
| gene_849112 | 11,34442857 | 4,461091732 | 1,312798642 | 3,398153829 | 0,000678423 | 0,016976746 | The Moose: suction sampler vs FISH | NA |
| gene_850844 | 25,84009994 | 3,493347483 | 1,019028123 | 3,428116855 | 0,000607784 | 0,015790124 | The Moose: suction sampler vs FISH | NA |
| gene_851428 | 12,98509316 | 5,332949476 | 1,280649443 | 4,164253929 | 3,12E-05 | 0,002268786 | The Moose: suction sampler vs FISH | NA |
| gene_851429 | 12,28326328 | 6,380280508 | 1,488817753 | 4,28546778 | 1,82E-05 | 0,001562785 | The Moose: suction sampler vs FISH | NA |
| gene_851537 | 109,7127408 | -2,301022717 | 0,691425239 | -3,327941457 | 0,000874902 | 0,020111804 | The Moose: suction sampler vs FISH | NA |
| gene_851793 | 13,54694757 | 4,409049639 | 1,159283839 | 3,803252915 | 0,000142808 | 0,006299438 | The Moose: suction sampler vs FISH | NA |
| gene_851961 | 5,034097883 | 7,031186672 | 1,888531574 | 3,723097231 | 0,000196794 | 0,007725891 | The Moose: suction sampler vs FISH | NA |
| gene_852018 | 4,783175313 | 5,408292624 | 1,675688211 | 3,227505324 | 0,001248747 | 0,024902046 | The Moose: suction sampler vs FISH | NA |
| gene_853544 | 8,65746577 | 5,322196277 | 1,561360537 | 3,408691428 | 0,000652753 | 0,016559171 | The Moose: suction sampler vs FISH | NA |
| gene_853841 | 1250,504243 | 5,166911741 | 0,924428848 | 5,589301711 | 2,28E-08 | 1,61E-05 | The Moose: suction sampler vs FISH | NA |
| gene_854172 | 3,49364986 | 6,357307847 | 2,198836361 | 2,891214626 | 0,003837559 | 0,04865923 | The Moose: suction sampler vs FISH | NA |
| gene_854350 | 11,18964716 | 3,86709392 | 1,317028824 | 2,93622573 | 0,003322325 | 0,044739299 | The Moose: suction sampler vs FISH | NA |
| gene_855199 | 40,16032122 | 3,281502675 | 0,862884813 | 3,80294406 | 0,000142987 | 0,006299733 | The Moose: suction sampler vs FISH | NA |
| gene_855200 | 31,57623057 | 4,383699969 | 1,066092696 | 4,111931342 | 3,92E-05 | 0,00266825 | The Moose: suction sampler vs FISH | NA |
| gene_855201 | 92,04722957 | 3,564704212 | 0,883048931 | 4,036813915 | 5,42E-05 | 0,003307895 | The Moose: suction sampler vs FISH | NA |
| gene_856332 | 11,05333181 | 3,67608663 | 1,127359008 | 3,260795013 | 0,001111003 | 0,023266414 | The Moose: suction sampler vs FISH | NA |
| gene_857810 | 3,66775007 | 6,198831023 | 2,017171434 | 3,073031334 | 0,002118962 | 0,034339476 | The Moose: suction sampler vs FISH | NA |
| gene_857851 | 4,584714174 | 5,968276966 | 1,705374364 | 3,499687278 | 0,000465804 | 0,013354236 | The Moose: suction sampler vs FISH | NA |
| gene_858120 | 87,6310529 | 2,38048827 | 0,817704768 | 2,911183062 | 0,00360063 | 0,04685476 | The Moose: suction sampler vs FISH | NA |
| gene_858163 | 8,760386477 | 3,661103903 | 1,233157729 | 2,96888534 | 0,002988821 | 0,042044428 | The Moose: suction sampler vs FISH | NA |
| gene_858195 | 13,444836 | 4,322535788 | 1,232637903 | 3,506736063 | 0,000453639 | 0,013137818 | The Moose: suction sampler vs FISH | NA |
| gene_858848 | 6,109923804 | 5,129452972 | 1,746875746 | 2,936358229 | 0,003320906 | 0,044736597 | The Moose: suction sampler vs FISH | NA |
| gene_860005 | 5,127504639 | 5,350936541 | 1,671641992 | 3,201006296 | 0,001369485 | 0,026354286 | The Moose: suction sampler vs FISH | NA |
| gene_860436 | 13,35479355 | 3,497690091 | 1,129456322 | 3,096790927 | 0,001956278 | 0,03272121 | The Moose: suction sampler vs FISH | NA |

| | | | | | | | | |
|---|---|---|---|---|---|---|---|---|
| gene_860697 | 85,33755892 | 6,406892277 | 1,018523989 | 6,290369539 | 3,17E-10 | 8,02E-07 | The Moose: suction sampler vs FISH | NA |
| gene_861296 | 11,93964194 | 3,40282245 | 1,090243385 | 3,121158537 | 0,00180141 | 0,031137829 | The Moose: suction sampler vs FISH | NA |
| gene_861678 | 66,88738473 | 2,893640409 | 0,786503708 | 3,679118583 | 0,000234041 | 0,008663898 | The Moose: suction sampler vs FISH | NA |
| gene_86170 | 5,959187181 | 6,269297752 | 1,690899131 | 3,707671048 | 0,000209174 | 0,008049569 | The Moose: suction sampler vs FISH | NA |
| gene_86171 | 8,302004415 | 4,324599662 | 1,361113581 | 3,177251129 | 0,001486782 | 0,027756766 | The Moose: suction sampler vs FISH | NA |
| gene_861773 | 9,827023517 | 3,350688537 | 1,118390609 | 2,995991301 | 0,002735542 | 0,039760916 | The Moose: suction sampler vs FISH | NA |
| gene_862145 | 34,22501746 | 5,491354732 | 1,250829452 | 4,390170636 | 1,13E-05 | 0,001156955 | The Moose: suction sampler vs FISH | NA |
| gene_862225 | 7,000171251 | 6,76762632 | 1,796465264 | 3,76719019 | 0,000165095 | 0,006916953 | The Moose: suction sampler vs FISH | NA |
| gene_863319 | 16,00242196 | 3,487054772 | 0,84410726 | 4,131056487 | 3,61E-05 | 0,002511456 | The Moose: suction sampler vs FISH | NA |
| gene_863371 | 3,293153045 | 6,22559845 | 2,15611724 | 2,887411842 | 0,003884254 | 0,048974947 | The Moose: suction sampler vs FISH | NA |
| gene_864738 | 4,797036623 | 6,474698127 | 1,960297554 | 3,302915985 | 0,000956851 | 0,021154692 | The Moose: suction sampler vs FISH | NA |
| gene_865782 | 8,787063777 | 4,230021255 | 1,432411271 | 2,953077332 | 0,003146231 | 0,043353321 | The Moose: suction sampler vs FISH | NA |
| gene_866000 | 69,17492839 | 3,090946259 | 0,84922054 | 3,639745052 | 0,000272908 | 0,009558291 | The Moose: suction sampler vs FISH | NA |
| gene_866438 | 5,993789218 | 6,971278416 | 1,71934902 | 4,054603419 | 5,02E-05 | 0,003168826 | The Moose: suction sampler vs FISH | NA |
| gene_866707 | 5,181100012 | 6,85645118 | 1,860003621 | 3,6862569 | 0,000227577 | 0,008510369 | The Moose: suction sampler vs FISH | NA |
| gene_866946 | 43,51597369 | 2,526857402 | 0,802209305 | 3,149872965 | 0,001633415 | 0,029419597 | The Moose: suction sampler vs FISH | NA |
| gene_867769 | 6,449037094 | 6,65988036 | 1,662209878 | 4,006642271 | 6,16E-05 | 0,003613435 | The Moose: suction sampler vs FISH | NA |
| gene_868024 | 21,10829188 | 3,812857565 | 0,979625958 | 3,892156524 | 9,94E-05 | 0,00498108 | The Moose: suction sampler vs FISH | NA |
| gene_868601 | 11,67504937 | 4,54257553 | 1,215480509 | 3,737267276 | 0,000186031 | 0,007453932 | The Moose: suction sampler vs FISH | NA |
| gene_869702 | 84,56984003 | 3,955190074 | 1,007550795 | 3,92554906 | 8,65E-05 | 0,004539537 | The Moose: suction sampler vs FISH | NA |
| gene_872373 | 177,4874528 | 2,640703405 | 0,753506246 | 3,504554104 | 0,000457373 | 0,013225091 | The Moose: suction sampler vs FISH | NA |
| gene_873965 | 18,53880105 | 2,958240558 | 0,879312093 | 3,364266889 | 0,000767473 | 0,018432651 | The Moose: suction sampler vs FISH | NA |
| gene_87549 | 8,699093337 | 5,906970093 | 1,686124274 | 3,503282756 | 0,000459561 | 0,013272703 | The Moose: suction sampler vs FISH | NA |
| gene_875512 | 31,74593156 | 2,575479752 | 0,843273841 | 3,054144012 | 0,002257037 | 0,035496378 | The Moose: suction sampler vs FISH | NA |
| gene_87661 | 1297,703657 | -2,214624707 | 0,637776666 | -3,472414129 | 0,0005158 | 0,014214983 | The Moose: suction sampler vs FISH | NA |
| gene_876892 | 28,60407137 | 2,943440384 | 0,840282055 | 3,502919485 | 0,000460189 | 0,013284787 | The Moose: suction sampler vs FISH | NA |
| gene_877221 | 10,9280272 | 4,749739881 | 1,265728486 | 3,752574058 | 0,000175028 | 0,007161526 | The Moose: suction sampler vs FISH | NA |
| gene_878853 | 7,641553849 | 5,070528568 | 1,465409483 | 3,460144503 | 0,000539886 | 0,014631696 | The Moose: suction sampler vs FISH | NA |
| gene_878876 | 78,52959617 | 9,979881149 | 2,825461334 | 3,532124481 | 0,000412235 | 0,012407978 | The Moose: suction sampler vs FISH | NA |
| gene_8794 | 32,38918976 | 2,677066447 | 0,840621206 | 3,184628735 | 0,001449398 | 0,027309147 | The Moose: suction sampler vs FISH | NA |
| gene_8795 | 147,6449785 | 2,242248086 | 0,673437301 | 3,329557305 | 0,000869842 | 0,020049897 | The Moose: suction sampler vs FISH | NA |
| gene_879918 | 17,32823352 | 9,373454125 | 3,105392874 | 3,018443883 | 0,002540765 | 0,038189863 | The Moose: suction sampler vs FISH | NA |
| gene_880 | 27,33902746 | 3,128210449 | 1,076822401 | 2,905038423 | 0,00367208 | 0,0474274 | The Moose: suction sampler vs FISH | NA |
| gene_880969 | 9,106478149 | 4,333937512 | 1,370271859 | 3,162830415 | 0,001562433 | 0,028588655 | The Moose: suction sampler vs FISH | NA |
| gene_881285 | 9,382607262 | 3,274064932 | 1,115572867 | 2,934873219 | 0,00333684 | 0,04484432 | The Moose: suction sampler vs FISH | NA |
| gene_882426 | 3,761572721 | 6,746543254 | 2,126796748 | 3,172161731 | 0,001513087 | 0,028048634 | The Moose: suction sampler vs FISH | NA |
| gene_88309 | 18,04467579 | 8,915975698 | 2,659682131 | 3,352271159 | 0,000801515 | 0,018940557 | The Moose: suction sampler vs FISH | NA |
| gene_88310 | 25,25867843 | 9,714682677 | 3,025579551 | 3,21085019 | 0,001323429 | 0,025823392 | The Moose: suction sampler vs FISH | NA |
| gene_883434 | 18,06208916 | 3,522512111 | 0,994073597 | 3,543512392 | 0,000394835 | 0,012102938 | The Moose: suction sampler vs FISH | NA |
| gene_883700 | 3,747518356 | 5,781129077 | 1,997967482 | 2,893505089 | 0,003809681 | 0,048430347 | The Moose: suction sampler vs FISH | NA |
| gene_883701 | 4,234873413 | 6,550239665 | 2,03459689 | 3,219428722 | 0,001284463 | 0,025379978 | The Moose: suction sampler vs FISH | NA |
| gene_886723 | 53,1302052 | 3,620431457 | 0,804723012 | 4,498978409 | 6,83E-06 | 0,00083296 | The Moose: suction sampler vs FISH | NA |
| gene_886938 | 24,64916785 | 2,812964369 | 0,818581469 | 3,436389017 | 0,000589524 | 0,015474474 | The Moose: suction sampler vs FISH | NA |
| gene_887528 | 25,52654655 | 2,858355491 | 0,900428205 | 3,1744402 | 0,001501258 | 0,027946872 | The Moose: suction sampler vs FISH | NA |
| gene_887796 | 21,04687376 | 2,648228527 | 0,867358911 | 3,053209568 | 0,002264078 | 0,035551536 | The Moose: suction sampler vs FISH | NA |
| gene_889007 | 29,06301787 | 3,203476528 | 0,922695087 | 3,471869062 | 0,000516848 | 0,014233194 | The Moose: suction sampler vs FISH | NA |
| gene_88945 | 2,873511836 | 5,960168891 | 2,036194494 | 2,927111781 | 0,003421259 | 0,04544662 | The Moose: suction sampler vs FISH | NA |
| gene_889554 | 5,871956923 | 7,097402016 | 1,827326575 | 3,8840359 | 0,000102737 | 0,005077725 | The Moose: suction sampler vs FISH | NA |
| gene_889562 | 32,90351386 | 2,718318972 | 0,921181815 | 2,950903859 | 0,003168455 | 0,043488543 | The Moose: suction sampler vs FISH | NA |
| gene_889720 | 504,0269046 | 2,947841847 | 0,791777432 | 3,723068794 | 0,000196816 | 0,007725891 | The Moose: suction sampler vs FISH | NA |
| gene_890031 | 7,741188613 | 4,289383037 | 1,483611141 | 2,891177425 | 0,003838014 | 0,04865923 | The Moose: suction sampler vs FISH | NA |
| gene_890771 | 17,95345784 | 3,616606253 | 1,115260084 | 3,242836631 | 0,00118346 | 0,024130832 | The Moose: suction sampler vs FISH | NA |
| gene_890772 | 4,184933302 | 5,604838135 | 1,852602131 | 3,025386855 | 0,002483153 | 0,037649704 | The Moose: suction sampler vs FISH | NA |
| gene_891124 | 23,33884086 | 3,273608482 | 0,845855294 | 3,870175555 | 0,000108757 | 0,005268904 | The Moose: suction sampler vs FISH | NA |
| gene_892322 | 5,583525759 | 5,277053243 | 1,806635581 | 2,920928437 | 0,0034899 | 0,04601718 | The Moose: suction sampler vs FISH | NA |
| gene_89271 | 12,92888263 | 4,459669112 | 1,260388779 | 3,538328162 | 0,000402669 | 0,012232174 | The Moose: suction sampler vs FISH | NA |
| gene_893317 | 10,51746703 | 7,663809746 | 1,865537509 | 4,108097376 | 3,99E-05 | 0,00270042 | The Moose: suction sampler vs FISH | NA |
| gene_893606 | 5,766762579 | 7,856068828 | 1,967246138 | 3,993434617 | 6,51E-05 | 0,003740709 | The Moose: suction sampler vs FISH | NA |
| gene_89535 | 14,0389672 | 2,854899252 | 0,97305741 | 2,933947394 | 0,003346809 | 0,044929334 | The Moose: suction sampler vs FISH | NA |
| gene_897565 | 16,23863494 | 5,201423237 | 1,262768402 | 4,119063503 | 3,80E-05 | 0,00261362 | The Moose: suction sampler vs FISH | NA |
| gene_898353 | 14,9081383 | 4,476061443 | 1,249542866 | 3,582159175 | 0,000340766 | 0,011015763 | The Moose: suction sampler vs FISH | NA |
| gene_898556 | 14,81290371 | 7,83057592 | 1,544822686 | 5,068915669 | 4,00E-07 | 0,000119264 | The Moose: suction sampler vs FISH | NA |
| gene_899537 | 12,87401869 | 3,903490988 | 1,148091389 | 3,399982811 | 0,000673901 | 0,01689814 | The Moose: suction sampler vs FISH | NA |
| gene_90014 | 396,0923031 | 3,177930261 | 0,840152412 | 3,782563992 | 0,000155221 | 0,006612656 | The Moose: suction sampler vs FISH | NA |
| gene_900851 | 13,33685025 | 4,230120349 | 1,278236296 | 3,309341444 | 0,000935157 | 0,020887457 | The Moose: suction sampler vs FISH | NA |

| | | | | | | | | |
|---|---|---|---|---|---|---|---|---|
| gene_900986 | 31,77898095 | 3,569921171 | 1,03091023 | 3,462882672 | 0,000534421 | 0,014548027 | The Moose: suction sampler vs FISH | NA |
| gene_902571 | 11,69019553 | 5,343251555 | 1,372472672 | 3,893156975 | 9,89E-05 | 0,00497608 | The Moose: suction sampler vs FISH | NA |
| gene_902778 | 28,2566749 | 3,187752198 | 0,979799851 | 3,253472835 | 0,001140036 | 0,023607465 | The Moose: suction sampler vs FISH | NA |
| gene_902779 | 12,45828213 | 4,119987408 | 1,187035148 | 3,470821748 | 0,000518868 | 0,014277216 | The Moose: suction sampler vs FISH | NA |
| gene_902780 | 14,42501306 | 4,414526725 | 1,265750498 | 3,487675281 | 0,000487239 | 0,013721357 | The Moose: suction sampler vs FISH | NA |
| gene_90336 | 11,65110151 | 4,658406542 | 1,195823385 | 3,895564013 | 9,80E-05 | 0,004952128 | The Moose: suction sampler vs FISH | NA |
| gene_903487 | 3,253376715 | 6,260299854 | 2,045371372 | 3,060715497 | 0,002208088 | 0,035148033 | The Moose: suction sampler vs FISH | NA |
| gene_903778 | 34,73422179 | 10,34493713 | 2,681518666 | 3,857865045 | 0,000114382 | 0,005451505 | The Moose: suction sampler vs FISH | NA |
| gene_904342 | 48,94524371 | -2,808807699 | 0,741831258 | -3,786316184 | 0,000152897 | 0,00654783 | The Moose: suction sampler vs FISH | NA |
| gene_905055 | 3,796339384 | 6,008388001 | 1,90064591 | 3,161234804 | 0,001571018 | 0,02868807 | The Moose: suction sampler vs FISH | NA |
| gene_90520 | 13,14291981 | 3,588810983 | 1,224565378 | 2,930681404 | 0,003382195 | 0,04511538 | The Moose: suction sampler vs FISH | NA |
| gene_906478 | 20,39755882 | 5,073149402 | 1,242341363 | 4,083538997 | 4,44E-05 | 0,002913821 | The Moose: suction sampler vs FISH | NA |
| gene_90669 | 37,94276108 | 6,097492547 | 1,008043791 | 6,048836967 | 1,46E-09 | 2,33E-06 | The Moose: suction sampler vs FISH | NA |
| gene_906875 | 10,2310685 | 4,465255716 | 1,342915133 | 3,325046838 | 0,000884037 | 0,020245739 | The Moose: suction sampler vs FISH | NA |
| gene_907927 | 26,67961463 | 2,566742331 | 0,868458884 | 2,955513932 | 0,003121486 | 0,043192287 | The Moose: suction sampler vs FISH | NA |
| gene_908963 | 75,56546285 | 2,102940346 | 0,725639246 | 2,89805211 | 0,003754882 | 0,048064598 | The Moose: suction sampler vs FISH | NA |
| gene_909243 | 131,5503312 | 12,40647343 | 4,247282761 | 2,921037786 | 0,003488675 | 0,046009292 | The Moose: suction sampler vs FISH | NA |
| gene_909555 | 7,764482919 | 4,033212274 | 1,370741065 | 2,942358974 | 0,003257221 | 0,044186664 | The Moose: suction sampler vs FISH | NA |
| gene_909575 | 18,7594842 | 5,713370414 | 1,060128572 | 5,389318397 | 7,07E-08 | 3,76E-05 | The Moose: suction sampler vs FISH | NA |
| gene_909853 | 411,7293432 | 3,016817809 | 0,831262865 | 3,629198338 | 0,000284303 | 0,00984091 | The Moose: suction sampler vs FISH | NA |
| gene_912716 | 72,98823839 | 2,657253208 | 0,685751195 | 3,874952356 | 0,000106646 | 0,005204374 | The Moose: suction sampler vs FISH | NA |
| gene_913498 | 5,046396813 | 5,090446769 | 1,663869049 | 3,059403486 | 0,002217782 | 0,03520092 | The Moose: suction sampler vs FISH | NA |
| gene_91379 | 33,15719519 | -4,299010791 | 1,152106707 | -3,731434567 | 0,000190392 | 0,007591409 | The Moose: suction sampler vs FISH | NA |
| gene_914767 | 339,2556186 | 4,07957456 | 0,804363626 | 5,07180388 | 3,94E-07 | 0,000118614 | The Moose: suction sampler vs FISH | NA |
| gene_915455 | 3,763172573 | 5,587665282 | 1,942537888 | 2,876476859 | 0,004021417 | 0,049941318 | The Moose: suction sampler vs FISH | NA |
| gene_915807 | 6,147715701 | 4,545638028 | 1,574866299 | 2,886364406 | 0,003897206 | 0,049048664 | The Moose: suction sampler vs FISH | NA |
| gene_916550 | 3,866641554 | 5,571625646 | 1,820534834 | 3,060433418 | 0,002210169 | 0,035159134 | The Moose: suction sampler vs FISH | NA |
| gene_921318 | 38,37670306 | -2,722199586 | 0,917270136 | -2,967718538 | 0,003000189 | 0,042188186 | The Moose: suction sampler vs FISH | NA |
| gene_921548 | 7,124463789 | 5,126574926 | 1,522778213 | 3,366593298 | 0,000761028 | 0,018350055 | The Moose: suction sampler vs FISH | NA |
| gene_921549 | 40,42582642 | 4,506732375 | 0,900828686 | 5,002873959 | 5,65E-07 | 0,00014922 | The Moose: suction sampler vs FISH | NA |
| gene_92210 | 3,088763388 | 5,967881698 | 2,017514275 | 2,958036913 | 0,00309605 | 0,043002044 | The Moose: suction sampler vs FISH | NA |
| gene_922651 | 6,27269451 | 6,013025534 | 1,838082642 | 3,271357553 | 0,001070325 | 0,022715418 | The Moose: suction sampler vs FISH | NA |
| gene_923838 | 62,36332073 | 4,616585191 | 0,873421753 | 5,28563111 | 1,25E-07 | 5,32E-05 | The Moose: suction sampler vs FISH | NA |
| gene_924518 | 8,349921315 | 3,945101128 | 1,202014291 | 3,282075061 | 0,001030462 | 0,022150412 | The Moose: suction sampler vs FISH | NA |
| gene_924933 | 9,922685943 | 4,113903147 | 1,194606886 | 3,443729645 | 0,000573749 | 0,015255253 | The Moose: suction sampler vs FISH | NA |
| gene_925179 | 8,339523908 | 5,135407225 | 1,463718562 | 3,508466285 | 0,000450698 | 0,013094092 | The Moose: suction sampler vs FISH | NA |
| gene_925347 | 170,8521449 | 2,260416621 | 0,64124933 | 3,525019855 | 0,000423451 | 0,012611664 | The Moose: suction sampler vs FISH | NA |
| gene_925427 | 5,503853458 | 4,723052067 | 1,634154997 | 2,890210583 | 0,003849838 | 0,048731896 | The Moose: suction sampler vs FISH | NA |
| gene_925519 | 5,718427246 | 6,814448895 | 1,986935176 | 3,429628192 | 0,000604409 | 0,015724693 | The Moose: suction sampler vs FISH | NA |
| gene_92558 | 10,40374984 | -5,925911472 | 1,673958795 | -3,540058148 | 0,000400039 | 0,012206421 | The Moose: suction sampler vs FISH | NA |
| gene_925693 | 7,746544993 | 5,330937881 | 1,532237611 | 3,479184848 | 0,000502942 | 0,014018422 | The Moose: suction sampler vs FISH | NA |
| gene_925796 | 17,25594672 | 4,525514765 | 1,081937742 | 4,182786669 | 2,88E-05 | 0,002160197 | The Moose: suction sampler vs FISH | NA |
| gene_925797 | 33,74370266 | 3,418901195 | 0,894473247 | 3,822250926 | 0,000132239 | 0,005984162 | The Moose: suction sampler vs FISH | NA |
| gene_925867 | 28,42045862 | 3,975241555 | 1,263784563 | 3,145505706 | 0,001658 | 0,029678718 | The Moose: suction sampler vs FISH | NA |
| gene_926592 | 6,51981687 | 5,775576074 | 1,792544859 | 3,221998069 | 0,001273 | 0,025255397 | The Moose: suction sampler vs FISH | NA |
| gene_927029 | 43,97502421 | 3,249186651 | 0,918596994 | 3,537118748 | 0,000404518 | 0,012270495 | The Moose: suction sampler vs FISH | NA |
| gene_928162 | 5,457270889 | 6,719399288 | 1,728422419 | 3,887590912 | 0,000101244 | 0,005037312 | The Moose: suction sampler vs FISH | NA |
| gene_928335 | 5,116619876 | 6,535752384 | 1,974898431 | 3,309411908 | 0,000934922 | 0,020887457 | The Moose: suction sampler vs FISH | NA |
| gene_929064 | 11,90024885 | 3,122329646 | 1,044870969 | 2,988244231 | 0,002805852 | 0,040470508 | The Moose: suction sampler vs FISH | NA |
| gene_929857 | 7,849281857 | 5,576784582 | 1,609522136 | 3,464869763 | 0,000530488 | 0,014487375 | The Moose: suction sampler vs FISH | NA |
| gene_931149 | 20,38284446 | 4,2740507 | 0,970842139 | 4,402415724 | 1,07E-05 | 0,001112737 | The Moose: suction sampler vs FISH | NA |
| gene_931188 | 4,774457537 | 6,811692591 | 2,207166013 | 3,086171384 | 0,002027519 | 0,033448147 | The Moose: suction sampler vs FISH | NA |
| gene_93161 | 3,308978083 | 6,713038428 | 2,277995722 | 2,946905633 | 0,003209711 | 0,043874419 | The Moose: suction sampler vs FISH | NA |
| gene_932008 | 5,378056194 | 6,739485342 | 1,848754966 | 3,645418384 | 0,000266957 | 0,009439897 | The Moose: suction sampler vs FISH | NA |
| gene_932009 | 8,005332938 | 7,599951998 | 1,833513103 | 4,145021919 | 3,40E-05 | 0,002418143 | The Moose: suction sampler vs FISH | NA |
| gene_933342 | 7,712427211 | 8,154820304 | 1,917469981 | 4,252906373 | 2,11E-05 | 0,001729677 | The Moose: suction sampler vs FISH | NA |
| gene_933416 | 8,738100992 | 3,141010205 | 1,065537 | 2,947818991 | 0,003200244 | 0,043776571 | The Moose: suction sampler vs FISH | NA |
| gene_934120 | 4,159835371 | 6,112079903 | 1,993667703 | 3,06574656 | 0,002171273 | 0,034799623 | The Moose: suction sampler vs FISH | NA |
| gene_934510 | 431,0042019 | 5,744419047 | 0,996951656 | 5,761983557 | 8,31E-09 | 8,03E-06 | The Moose: suction sampler vs FISH | NA |
| gene_935075 | 14,52652616 | 4,422184172 | 1,109007332 | 3,987515721 | 6,68E-05 | 0,00380437 | The Moose: suction sampler vs FISH | NA |
| gene_935076 | 23,17340419 | 5,117080794 | 1,011579481 | 5,058505919 | 4,23E-07 | 0,000123153 | The Moose: suction sampler vs FISH | NA |
| gene_935077 | 14,05056368 | 3,603454391 | 1,025628705 | 3,513410237 | 0,000442394 | 0,012950027 | The Moose: suction sampler vs FISH | NA |
| gene_935377 | 5,479881014 | 5,882977424 | 1,774241997 | 3,315769457 | 0,000913912 | 0,020610638 | The Moose: suction sampler vs FISH | NA |
| gene_936881 | 13,81154315 | 3,651612847 | 1,043552549 | 3,499213193 | 0,000466633 | 0,013371785 | The Moose: suction sampler vs FISH | NA |
| gene_937464 | 31,19908281 | 2,495313538 | 0,799643705 | 3,120531708 | 0,001805249 | 0,031182146 | The Moose: suction sampler vs FISH | NA |

| | | | | | | | | |
|---|---|---|---|---|---|---|---|---|
| gene_937927 | 9,168570353 | -5,454441512 | 1,637783612 | -3,330379832 | 0,000867276 | 0,020014914 | The Moose: suction sampler vs FISH | NA |
| gene_937928 | 10,7834806 | -6,485666896 | 1,568578524 | -4,134741612 | 3,55E-05 | 0,002482193 | The Moose: suction sampler vs FISH | NA |
| gene_937932 | 332,2051772 | -3,895899605 | 1,003272035 | -3,883193657 | 0,000103093 | 0,005091926 | The Moose: suction sampler vs FISH | NA |
| gene_937934 | 153,4815359 | -3,095904741 | 0,870525559 | -3,556362832 | 0,000376025 | 0,011720579 | The Moose: suction sampler vs FISH | NA |
| gene_937935 | 49,36507957 | -3,2328438 | 1,059775072 | -3,050499947 | 0,002284607 | 0,035723316 | The Moose: suction sampler vs FISH | NA |
| gene_939151 | 4,062830243 | 6,487008845 | 2,090143853 | 3,103618364 | 0,001911697 | 0,032281658 | The Moose: suction sampler vs FISH | NA |
| gene_939198 | 7,321813507 | 4,544493909 | 1,485641563 | 3,058943705 | 0,002221189 | 0,035211572 | The Moose: suction sampler vs FISH | NA |
| gene_939200 | 25,13061128 | 3,345886786 | 1,080851332 | 3,095603149 | 0,00196413 | 0,032800264 | The Moose: suction sampler vs FISH | NA |
| gene_93930 | 3,229796009 | 6,316984837 | 2,156178253 | 2,92971364 | 0,003392745 | 0,045231468 | The Moose: suction sampler vs FISH | NA |
| gene_939928 | 72,56152677 | 2,884604799 | 0,732306943 | 3,939065208 | 8,18E-05 | 0,004359781 | The Moose: suction sampler vs FISH | NA |
| gene_941138 | 5,049495766 | 6,744934035 | 1,773585719 | 3,802992978 | 0,000142958 | 0,006299733 | The Moose: suction sampler vs FISH | NA |
| gene_943078 | 8,368780077 | 4,300310405 | 1,458010473 | 2,949437253 | 0,003183532 | 0,043653564 | The Moose: suction sampler vs FISH | NA |
| gene_946011 | 4,317430348 | 6,671325337 | 1,920867365 | 3,47307964 | 0,000514523 | 0,014209953 | The Moose: suction sampler vs FISH | NA |
| gene_946116 | 9,371010633 | 7,691296426 | 1,930197588 | 3,984719737 | 6,76E-05 | 0,003840504 | The Moose: suction sampler vs FISH | NA |
| gene_946628 | 16,53625466 | 5,824385197 | 1,438844266 | 4,047960808 | 5,17E-05 | 0,003223638 | The Moose: suction sampler vs FISH | NA |
| gene_946752 | 14,43290879 | 4,304513232 | 1,200407749 | 3,585875914 | 0,000335949 | 0,010922422 | The Moose: suction sampler vs FISH | NA |
| gene_949054 | 6,37401565 | 4,508788762 | 1,537511709 | 2,932523204 | 0,003362198 | 0,044987544 | The Moose: suction sampler vs FISH | NA |
| gene_949903 | 11,76306355 | 3,964259702 | 1,272630202 | 3,115013061 | 0,001839368 | 0,031605076 | The Moose: suction sampler vs FISH | NA |
| gene_950471 | 186,5343996 | -2,848540255 | 0,573957363 | -4,962982341 | 6,94E-07 | 0,000172246 | The Moose: suction sampler vs FISH | NA |
| gene_951429 | 3,09765721 | 6,54830489 | 2,20290856 | 2,972572267 | 0,002953157 | 0,041726549 | The Moose: suction sampler vs FISH | NA |
| gene_951760 | 98,44808743 | 3,784483062 | 0,751682413 | 5,034683526 | 4,79E-07 | 0,000136096 | The Moose: suction sampler vs FISH | NA |
| gene_951761 | 117,9887346 | 4,116126115 | 0,848750258 | 4,849631649 | 1,24E-06 | 0,000254879 | The Moose: suction sampler vs FISH | NA |
| gene_951762 | 11,51657845 | 4,058096972 | 1,160204098 | 3,497744044 | 0,000469211 | 0,013392145 | The Moose: suction sampler vs FISH | NA |
| gene_95216 | 11,02263376 | 5,461772311 | 1,517162855 | 3,599990793 | 0,000318228 | 0,010580484 | The Moose: suction sampler vs FISH | NA |
| gene_952591 | 91,75498771 | 4,049015987 | 0,758666833 | 5,337014629 | 9,45E-08 | 4,60E-05 | The Moose: suction sampler vs FISH | NA |
| gene_953312 | 132,3879252 | 3,207659426 | 0,803166375 | 3,99376708 | 6,50E-05 | 0,003740709 | The Moose: suction sampler vs FISH | NA |
| gene_95343 | 8,008013344 | 4,885799177 | 1,319143561 | 3,703766081 | 0,000212422 | 0,0081257 | The Moose: suction sampler vs FISH | NA |
| gene_9535 | 5,611971212 | 5,238391056 | 1,631045683 | 3,211676478 | 0,001319629 | 0,025804086 | The Moose: suction sampler vs FISH | NA |
| gene_953585 | 7,333099975 | 7,179462075 | 1,640912138 | 4,375287322 | 1,21E-05 | 0,001203799 | The Moose: suction sampler vs FISH | NA |
| gene_953600 | 7,902582845 | 5,596571626 | 1,723518344 | 3,247178451 | 0,001165553 | 0,023920816 | The Moose: suction sampler vs FISH | NA |
| gene_954336 | 13,68285851 | 3,587515886 | 1,115516207 | 3,216014132 | 0,001299844 | 0,025546445 | The Moose: suction sampler vs FISH | NA |
| gene_954337 | 16,45090728 | 4,065031796 | 1,043696877 | 3,894839476 | 9,83E-05 | 0,004960126 | The Moose: suction sampler vs FISH | NA |
| gene_954541 | 62,15764828 | 2,535319451 | 0,681321978 | 3,721176671 | 0,000198297 | 0,007767412 | The Moose: suction sampler vs FISH | NA |
| gene_955475 | 26,2805267 | 2,587408373 | 0,863590715 | 2,996104901 | 0,002734523 | 0,039760916 | The Moose: suction sampler vs FISH | NA |
| gene_955591 | 16,00070693 | 3,439453535 | 1,017361601 | 3,380758162 | 0,000722861 | 0,017704153 | The Moose: suction sampler vs FISH | NA |
| gene_956247 | 3,790086799 | 6,71098347 | 2,251898613 | 2,980144591 | 0,002881124 | 0,041108821 | The Moose: suction sampler vs FISH | NA |
| gene_957794 | 14,37844764 | 3,994758179 | 1,120504803 | 3,565141505 | 0,00036366 | 0,01147787 | The Moose: suction sampler vs FISH | NA |
| gene_959580 | 39,93257099 | -2,103239539 | 0,729229754 | -2,884193257 | 0,003924179 | 0,049233224 | The Moose: suction sampler vs FISH | NA |
| gene_962486 | 11,82640422 | 4,123873029 | 1,278906213 | 3,224531233 | 0,001261791 | 0,025094025 | The Moose: suction sampler vs FISH | NA |
| gene_962787 | 25,24909123 | 4,978177487 | 1,102560135 | 4,515107458 | 6,33E-06 | 0,000786454 | The Moose: suction sampler vs FISH | NA |
| gene_964586 | 4,001783676 | 6,484305371 | 2,214413348 | 2,928227189 | 0,003409009 | 0,045344949 | The Moose: suction sampler vs FISH | NA |
| gene_965158 | 3,770142756 | 6,291465143 | 1,954609664 | 3,218783401 | 0,001287357 | 0,025413125 | The Moose: suction sampler vs FISH | NA |
| gene_967848 | 245,3845248 | 4,220081322 | 0,796884753 | 5,29572351 | 1,19E-07 | 5,15E-05 | The Moose: suction sampler vs FISH | NA |
| gene_96846 | 42,60548384 | 2,788011148 | 0,918740766 | 3,034600455 | 0,002408545 | 0,036899376 | The Moose: suction sampler vs FISH | NA |
| gene_968469 | 128,6773308 | 4,100332805 | 0,871960773 | 4,702428057 | 2,57E-06 | 0,000414072 | The Moose: suction sampler vs FISH | NA |
| gene_970693 | 30,31849498 | 3,513239127 | 0,993284926 | 3,536990277 | 0,000404715 | 0,012270495 | The Moose: suction sampler vs FISH | NA |
| gene_971390 | 18,82246343 | 3,144783547 | 1,032316604 | 3,046336303 | 0,002316486 | 0,036030135 | The Moose: suction sampler vs FISH | NA |
| gene_972528 | 3,841393075 | 6,411908074 | 2,049153219 | 3,129052534 | 0,00175371 | 0,030674255 | The Moose: suction sampler vs FISH | NA |
| gene_972931 | 11,75530361 | 3,411391189 | 1,030820708 | 3,309393342 | 0,000934984 | 0,020887457 | The Moose: suction sampler vs FISH | NA |
| gene_973029 | 7,920504131 | 5,246266048 | 1,451092738 | 3,615389914 | 0,000299896 | 0,010197155 | The Moose: suction sampler vs FISH | NA |
| gene_973272 | 11,78700415 | 5,473723299 | 1,236154673 | 4,428024598 | 9,51E-06 | 0,001027151 | The Moose: suction sampler vs FISH | NA |
| gene_97366 | 16,67516635 | 3,682507803 | 1,092017375 | 3,372206237 | 0,000745686 | 0,018092798 | The Moose: suction sampler vs FISH | NA |
| gene_974558 | 15,96336666 | 3,02752993 | 1,02579665 | 2,951393856 | 0,003163433 | 0,043476479 | The Moose: suction sampler vs FISH | NA |
| gene_975155 | 5,368213587 | 5,38596942 | 1,848175893 | 2,914208243 | 0,00356592 | 0,046576375 | The Moose: suction sampler vs FISH | NA |
| gene_975392 | 11,618966 | 3,956405382 | 1,180903353 | 3,350321066 | 0,000807179 | 0,019019343 | The Moose: suction sampler vs FISH | NA |
| gene_975537 | 11,9319649 | 4,080902606 | 1,316221539 | 3,100467882 | 0,001932152 | 0,032529479 | The Moose: suction sampler vs FISH | NA |
| gene_975932 | 7,628527646 | 6,098257073 | 1,647723833 | 3,701018916 | 0,000214735 | 0,008176582 | The Moose: suction sampler vs FISH | NA |
| gene_975955 | 3,998863315 | 6,356007512 | 2,009580409 | 3,162853043 | 0,001562312 | 0,028588655 | The Moose: suction sampler vs FISH | NA |
| gene_977043 | 55,93512475 | 2,603984261 | 0,744409727 | 3,498052439 | 0,000468669 | 0,013392145 | The Moose: suction sampler vs FISH | NA |
| gene_977369 | 72,53036065 | 2,776766632 | 0,907612982 | 3,05941705 | 0,002217682 | 0,03520092 | The Moose: suction sampler vs FISH | NA |
| gene_977370 | 133,7774738 | 2,472708035 | 0,801976045 | 3,083269194 | 0,002047399 | 0,033673581 | The Moose: suction sampler vs FISH | NA |
| gene_977371 | 33,76562409 | 2,755677907 | 0,830117654 | 3,319623302 | 0,00090139 | 0,02049231 | The Moose: suction sampler vs FISH | NA |
| gene_978202 | 32,53984709 | 2,664095865 | 0,901913139 | 2,953827536 | 0,003138594 | 0,043306684 | The Moose: suction sampler vs FISH | NA |
| gene_979947 | 11,26913294 | 4,755730459 | 1,519973335 | 3,128824928 | 0,001755068 | 0,030685572 | The Moose: suction sampler vs FISH | NA |
| gene_979948 | 18,39612287 | 5,15237094 | 1,410847502 | 3,651968716 | 0,000260238 | 0,009273726 | The Moose: suction sampler vs FISH | NA |

| | | | | | | | | |
|---|---|---|---|---|---|---|---|---|
| gene_981052 | 49,74518802 | 3,791373124 | 0,746126153 | 5,081410308 | 3,75E-07 | 0,000114649 | The Moose: suction sampler vs FISH | NA |
| gene_981053 | 22,31380268 | 3,47432802 | 0,999403754 | 3,47640081 | 0,000508192 | 0,014095232 | The Moose: suction sampler vs FISH | NA |
| gene_981054 | 22,41763089 | 2,503867569 | 0,864585975 | 2,89603075 | 0,003779154 | 0,048229356 | The Moose: suction sampler vs FISH | NA |
| gene_981055 | 17,04944053 | 5,29529257 | 1,123428315 | 4,713511759 | 2,43E-06 | 0,000397392 | The Moose: suction sampler vs FISH | NA |
| gene_981722 | 3,450974094 | 6,735465751 | 2,277297215 | 2,957657747 | 0,003099861 | 0,043012037 | The Moose: suction sampler vs FISH | NA |
| gene_981970 | 9,029093222 | 5,669504821 | 1,552383442 | 3,65212915 | 0,000260075 | 0,00927342 | The Moose: suction sampler vs FISH | NA |
| gene_983489 | 278,143249 | 6,351669875 | 0,936839042 | 6,779894506 | 1,20E-11 | 8,83E-08 | The Moose: suction sampler vs FISH | NA |
| gene_984190 | 7,816129812 | 7,914622648 | 1,941153989 | 4,077277069 | 4,56E-05 | 0,002972121 | The Moose: suction sampler vs FISH | NA |
| gene_984351 | 10,37404241 | 3,68961388 | 1,281017169 | 2,880222037 | 0,003973952 | 0,049608838 | The Moose: suction sampler vs FISH | NA |
| gene_985907 | 168,4052753 | 4,23706748 | 0,890272446 | 4,759293065 | 1,94E-06 | 0,000344645 | The Moose: suction sampler vs FISH | NA |
| gene_98699 | 38,31297037 | -2,312191565 | 0,761265101 | -3,037301408 | 0,002387066 | 0,036646758 | The Moose: suction sampler vs FISH | NA |
| gene_988182 | 5,393508083 | 4,986403209 | 1,707442385 | 2,920393246 | 0,003495899 | 0,046046689 | The Moose: suction sampler vs FISH | NA |
| gene_988358 | 5,843219519 | 5,078280459 | 1,622776656 | 3,129377318 | 0,001751772 | 0,030666956 | The Moose: suction sampler vs FISH | NA |
| gene_988501 | 5,067097893 | 5,515666962 | 1,809515201 | 3,048146244 | 0,002302578 | 0,035897447 | The Moose: suction sampler vs FISH | NA |
| gene_98904 | 47,33405566 | -3,236562241 | 1,102738986 | -2,935021145 | 0,00333525 | 0,044841602 | The Moose: suction sampler vs FISH | NA |
| gene_990006 | 25,1902706 | 2,805815947 | 0,876551609 | 3,200970619 | 0,001369655 | 0,026354286 | The Moose: suction sampler vs FISH | NA |
| gene_990007 | 10,00158474 | 4,855248122 | 1,357060838 | 3,577767471 | 0,000346542 | 0,011138618 | The Moose: suction sampler vs FISH | NA |
| gene_990700 | 31,6982561 | -2,246074592 | 0,736193324 | -3,050930397 | 0,002281335 | 0,035717888 | The Moose: suction sampler vs FISH | NA |
| gene_990701 | 47,07514165 | -2,480385521 | 0,665561221 | -3,726757876 | 0,000193959 | 0,007660085 | The Moose: suction sampler vs FISH | NA |
| gene_990702 | 134,56757 | -2,329460461 | 0,562887323 | -4,138413439 | 3,50E-05 | 0,002454294 | The Moose: suction sampler vs FISH | NA |
| gene_991624 | 49,96670569 | 3,10973464 | 0,89929764 | 3,457959303 | 0,000544284 | 0,014710516 | The Moose: suction sampler vs FISH | NA |
| gene_991987 | 9,732828391 | 5,606810811 | 1,602098215 | 3,499667347 | 0,000465839 | 0,013354236 | The Moose: suction sampler vs FISH | NA |
| gene_992574 | 13,01029578 | 3,824788042 | 1,105632135 | 3,459367653 | 0,000541445 | 0,014654547 | The Moose: suction sampler vs FISH | NA |
| gene_99290 | 24,5911357 | 4,510847946 | 1,075441574 | 4,194414698 | 2,74E-05 | 0,002092916 | The Moose: suction sampler vs FISH | NA |
| gene_993464 | 391,0684465 | 3,565058796 | 1,002863132 | 3,554880703 | 0,000378151 | 0,011773325 | The Moose: suction sampler vs FISH | NA |
| gene_993933 | 37,10328787 | -3,322638682 | 0,802424744 | -4,140748034 | 3,46E-05 | 0,002444695 | The Moose: suction sampler vs FISH | NA |
| gene_994479 | 16,48279482 | 4,561827336 | 1,400615049 | 3,257017221 | 0,001125896 | 0,023426104 | The Moose: suction sampler vs FISH | NA |
| gene_995804 | 70,7583157 | 4,189638776 | 0,808229458 | 5,183724414 | 2,17E-07 | 7,75E-05 | The Moose: suction sampler vs FISH | NA |
| gene_995873 | 96,12152476 | 5,472641362 | 1,01323066 | 5,401180181 | 6,62E-08 | 3,63E-05 | The Moose: suction sampler vs FISH | NA |
| gene_997792 | 7,670418834 | 3,797328871 | 1,299415786 | 2,922335493 | 0,003474171 | 0,045875671 | The Moose: suction sampler vs FISH | NA |
| gene_997905 | 45,66893077 | 3,439449601 | 0,932002721 | 3,690385791 | 0,000223914 | 0,008407658 | The Moose: suction sampler vs FISH | NA |
| gene_999241 | 15,90404208 | 5,727677096 | 1,32057567 | 4,337257779 | 1,44E-05 | 0,00133788 | The Moose: suction sampler vs FISH | NA |
| gene_384115 | 6,534576378 | -5,626621832 | 1,735839839 | -3,24144066 | 0,001189272 | 0,024188888 | The Moose: suction sampler vs FISH | Nucleotide transport and metabolism |
| gene_367798 | 4,186254462 | -5,404983048 | 1,744505901 | -3,098288773 | 0,001946417 | 0,03263058 | The Moose: suction sampler vs FISH | Nucleotide transport and metabolism |
| gene_280482 | 6,306823804 | -5,179306435 | 1,693144163 | -3,058987267 | 0,002220866 | 0,035211572 | The Moose: suction sampler vs FISH | Nucleotide transport and metabolism |
| gene_813022 | 19,78921565 | -4,726725441 | 1,23629573 | -3,823296745 | 0,000131679 | 0,005963296 | The Moose: suction sampler vs FISH | Nucleotide transport and metabolism |
| gene_122813 | 22,69415038 | -3,986375369 | 1,274284756 | -3,128323832 | 0,001758064 | 0,030695208 | The Moose: suction sampler vs FISH | Nucleotide transport and metabolism |
| gene_472246 | 16,37097004 | -2,975582719 | 0,967013418 | -3,077085244 | 0,002090355 | 0,034092895 | The Moose: suction sampler vs FISH | Nucleotide transport and metabolism |
| gene_813023 | 57,67760774 | -2,97163572 | 1,017827981 | -2,919585408 | 0,003504973 | 0,046082171 | The Moose: suction sampler vs FISH | Nucleotide transport and metabolism |
| gene_2477 | 32,11291201 | -2,969661518 | 0,80896553 | -3,670937026 | 0,000241663 | 0,008843513 | The Moose: suction sampler vs FISH | Nucleotide transport and metabolism |
| gene_988785 | 152,221089 | -2,942125007 | 0,662622122 | -4,440124935 | 8,99E-06 | 0,000987025 | The Moose: suction sampler vs FISH | Nucleotide transport and metabolism |
| gene_1665173 | 62,8737103 | -2,89863048 | 0,853490445 | -3,3962073 | 0,000683266 | 0,017039886 | The Moose: suction sampler vs FISH | Nucleotide transport and metabolism |
| gene_988777 | 69,57768231 | -2,817865904 | 0,745330631 | -3,78069247 | 0,000156393 | 0,006650994 | The Moose: suction sampler vs FISH | Nucleotide transport and metabolism |
| gene_209433 | 59,2211689 | -2,736113516 | 0,813173777 | -3,364734076 | 0,000766175 | 0,01840749 | The Moose: suction sampler vs FISH | Nucleotide transport and metabolism |
| gene_186477 | 43,46117433 | -2,686208626 | 0,864412369 | -3,107554592 | 0,001886421 | 0,032056506 | The Moose: suction sampler vs FISH | Nucleotide transport and metabolism |
| gene_1128907 | 531,6513663 | -2,616293726 | 0,644677887 | -4,058296054 | 4,94E-05 | 0,003132475 | The Moose: suction sampler vs FISH | Nucleotide transport and metabolism |
| gene_788961 | 17,49704896 | -2,61400743 | 0,90735613 | -2,880905682 | 0,003965343 | 0,049580287 | The Moose: suction sampler vs FISH | Nucleotide transport and metabolism |
| gene_131511 | 59,45415219 | -2,512354304 | 0,770301715 | -3,261519812 | 0,001108167 | 0,023247448 | The Moose: suction sampler vs FISH | Nucleotide transport and metabolism |
| gene_326077 | 56,05958849 | -2,47341249 | 0,710116353 | -3,483108759 | 0,000495627 | 0,013883037 | The Moose: suction sampler vs FISH | Nucleotide transport and metabolism |
| gene_693371 | 560,0277677 | -2,440677701 | 0,637933277 | -3,82591376 | 0,000130288 | 0,005936101 | The Moose: suction sampler vs FISH | Nucleotide transport and metabolism |
| gene_1274177 | 269,5787647 | -2,405921567 | 0,586274034 | -4,103749149 | 4,07E-05 | 0,002736572 | The Moose: suction sampler vs FISH | Nucleotide transport and metabolism |
| gene_44352 | 8345,128879 | -2,403141604 | 0,547965683 | -4,385569529 | 1,16E-05 | 0,001171906 | The Moose: suction sampler vs FISH | Nucleotide transport and metabolism |
| gene_42543 | 36,68409952 | -2,381746658 | 0,805681795 | -2,956187757 | 0,003114674 | 0,043133087 | The Moose: suction sampler vs FISH | Nucleotide transport and metabolism |
| gene_54436 | 4004,144822 | -2,316596477 | 0,617516525 | -3,751472849 | 0,000175799 | 0,007176644 | The Moose: suction sampler vs FISH | Nucleotide transport and metabolism |
| gene_124685 | 64,68858408 | -2,315792146 | 0,695432912 | -3,330000791 | 0,000868457 | 0,020026327 | The Moose: suction sampler vs FISH | Nucleotide transport and metabolism |
| gene_501017 | 28,48253698 | -2,289934737 | 0,734313973 | -3,118468152 | 0,001817938 | 0,031347689 | The Moose: suction sampler vs FISH | Nucleotide transport and metabolism |
| gene_1323882 | 2617,957643 | -2,278923242 | 0,613710459 | -3,713352461 | 0,000204532 | 0,007939658 | The Moose: suction sampler vs FISH | Nucleotide transport and metabolism |
| gene_474107 | 22,76452119 | -2,258387737 | 0,782515281 | -2,886062154 | 0,003900951 | 0,049060697 | The Moose: suction sampler vs FISH | Nucleotide transport and metabolism |
| gene_566654 | 983,3060517 | -2,221686114 | 0,57307809 | -3,876759821 | 0,000105857 | 0,005176198 | The Moose: suction sampler vs FISH | Nucleotide transport and metabolism |
| gene_1598913 | 8872,050195 | -2,210684685 | 0,559319357 | -3,952455168 | 7,74E-05 | 0,00422194 | The Moose: suction sampler vs FISH | Nucleotide transport and metabolism |
| gene_431768 | 5323,440798 | -2,179793223 | 0,583664628 | -3,734667339 | 0,000187963 | 0,0075048 | The Moose: suction sampler vs FISH | Nucleotide transport and metabolism |
| gene_1165027 | 948,8298892 | -2,137673771 | 0,620816186 | -3,443328026 | 0,000574602 | 0,015255253 | The Moose: suction sampler vs FISH | Nucleotide transport and metabolism |
| gene_1593736 | 113,5304435 | -2,133966116 | 0,599212829 | -3,561282423 | 0,000369048 | 0,011558516 | The Moose: suction sampler vs FISH | Nucleotide transport and metabolism |
| gene_70964 | 57,66393509 | -2,075920822 | 0,695787434 | -2,983556068 | 0,002849198 | 0,04087097 | The Moose: suction sampler vs FISH | Nucleotide transport and metabolism |
| gene_1191563 | 31,92968939 | 2,670901303 | 0,757406845 | 3,526375977 | 0,000421288 | 0,012588091 | The Moose: suction sampler vs FISH | Nucleotide transport and metabolism |

| | | | | | | | | |
|---|---|---|---|---|---|---|---|---|
| gene_1243037 | 51,83445183 | 2,729759827 | 0,934586607 | 2,920820613 | 0,003491108 | 0,046024845 | The Moose: suction sampler vs FISH | Nucleotide transport and metabolism |
| gene_322362 | 31,62311657 | 2,767164527 | 0,838843333 | 3,298785864 | 0,00097104 | 0,021339916 | The Moose: suction sampler vs FISH | Nucleotide transport and metabolism |
| gene_116385 | 35,17809188 | 3,273918814 | 0,99251372 | 3,298613156 | 0,000971637 | 0,021343394 | The Moose: suction sampler vs FISH | Nucleotide transport and metabolism |
| gene_328008 | 22,80584686 | 3,27686352 | 1,088946264 | 3,009205898 | 0,002619315 | 0,038863761 | The Moose: suction sampler vs FISH | Nucleotide transport and metabolism |
| gene_1389855 | 7,942553672 | 3,765703192 | 1,249881496 | 3,012848181 | 0,002588084 | 0,038598972 | The Moose: suction sampler vs FISH | Nucleotide transport and metabolism |
| gene_1706866 | 247,2445885 | 4,02390976 | 0,84567028 | 4,758248996 | 1,95E-06 | 0,000344767 | The Moose: suction sampler vs FISH | Nucleotide transport and metabolism |
| gene_565292 | 97,80411075 | 4,227879531 | 0,867035448 | 4,876247613 | 1,08E-06 | 0,000234943 | The Moose: suction sampler vs FISH | Nucleotide transport and metabolism |
| gene_1201658 | 15,15532389 | 4,427110725 | 1,28666532 | 3,440733233 | 0,000580076 | 0,015336096 | The Moose: suction sampler vs FISH | Nucleotide transport and metabolism |
| gene_1529412 | 22,75148124 | 4,44085085 | 1,053093306 | 4,216958578 | 2,48E-05 | 0,001947153 | The Moose: suction sampler vs FISH | Nucleotide transport and metabolism |
| gene_118488 | 7,180900467 | 4,709720563 | 1,31706484 | 3,575921564 | 0,000348996 | 0,011160923 | The Moose: suction sampler vs FISH | Nucleotide transport and metabolism |
| gene_689598 | 5,02410287 | 4,979994964 | 1,617104854 | 3,079574556 | 0,002072965 | 0,033938676 | The Moose: suction sampler vs FISH | Nucleotide transport and metabolism |
| gene_1201656 | 20,56920392 | 5,304964328 | 1,251985031 | 4,237242618 | 2,26E-05 | 0,00182029 | The Moose: suction sampler vs FISH | Nucleotide transport and metabolism |
| gene_151327 | 6,201079244 | 5,315284311 | 1,662947993 | 3,19630219 | 0,001392012 | 0,026578131 | The Moose: suction sampler vs FISH | Nucleotide transport and metabolism |
| gene_1211695 | 17,09718865 | 5,838631684 | 1,318027658 | 4,429824859 | 9,43E-06 | 0,001024113 | The Moose: suction sampler vs FISH | Nucleotide transport and metabolism |
| gene_888130 | 3,304745292 | 5,984071078 | 1,839343489 | 3,253373343 | 0,001140436 | 0,023607465 | The Moose: suction sampler vs FISH | Nucleotide transport and metabolism |
| gene_393788 | 3,748243359 | 6,278450093 | 1,863065732 | 3,369956296 | 0,000751801 | 0,018205087 | The Moose: suction sampler vs FISH | Nucleotide transport and metabolism |
| gene_665281 | 3,568522906 | 6,982994495 | 2,400730611 | 2,908695571 | 0,003629401 | 0,047079009 | The Moose: suction sampler vs FISH | Nucleotide transport and metabolism |
| gene_1310989 | 8,128786863 | -6,250897414 | 1,629912792 | -3,835111575 | 0,000125507 | 0,005797411 | The Moose: suction sampler vs FISH | Posttranslational modification, protein turnover, chaperones |
| gene_1114199 | 13,14654573 | -6,160698734 | 1,554323672 | -3,963588051 | 7,38E-05 | 0,004111117 | The Moose: suction sampler vs FISH | Posttranslational modification, protein turnover, chaperones |
| gene_1443594 | 6,376307254 | -6,096443679 | 1,677919807 | -3,633334355 | 0,000279782 | 0,009729449 | The Moose: suction sampler vs FISH | Posttranslational modification, protein turnover, chaperones |
| gene_1948232 | 5,840598355 | -5,637885373 | 1,63317959 | -3,452091496 | 0,000556259 | 0,014926725 | The Moose: suction sampler vs FISH | Posttranslational modification, protein turnover, chaperones |
| gene_1412779 | 4,200717538 | -5,175895466 | 1,790182885 | -2,891266311 | 0,003836928 | 0,04865923 | The Moose: suction sampler vs FISH | Posttranslational modification, protein turnover, chaperones |
| gene_418059 | 9,865482417 | -5,169579898 | 1,443060764 | -3,5823716 | 0,000340489 | 0,011015763 | The Moose: suction sampler vs FISH | Posttranslational modification, protein turnover, chaperones |
| gene_131504 | 51,97910109 | -3,736327498 | 0,909765015 | -4,106914904 | 4,01E-05 | 0,002711404 | The Moose: suction sampler vs FISH | Posttranslational modification, protein turnover, chaperones |
| gene_1770517 | 43,18237901 | -3,542274242 | 0,812851397 | -4,357837431 | 1,31E-05 | 0,001271052 | The Moose: suction sampler vs FISH | Posttranslational modification, protein turnover, chaperones |
| gene_377144 | 777,9648668 | -3,405153513 | 0,584552877 | -5,82522753 | 5,70E-09 | 5,90E-06 | The Moose: suction sampler vs FISH | Posttranslational modification, protein turnover, chaperones |
| gene_494878 | 22,73081654 | -3,037300363 | 0,940063809 | -3,230951275 | 0,00123379 | 0,024690922 | The Moose: suction sampler vs FISH | Posttranslational modification, protein turnover, chaperones |
| gene_1353220 | 73,97852866 | -3,035153848 | 0,767416768 | -3,955026753 | 7,65E-05 | 0,004194372 | The Moose: suction sampler vs FISH | Posttranslational modification, protein turnover, chaperones |
| gene_1114202 | 17,79730065 | -3,018761551 | 0,984385836 | -3,066644644 | 0,00216476 | 0,034774897 | The Moose: suction sampler vs FISH | Posttranslational modification, protein turnover, chaperones |
| gene_790062 | 1268,955357 | -2,844957078 | 0,566357206 | -5,023255727 | 5,08E-07 | 0,000142073 | The Moose: suction sampler vs FISH | Posttranslational modification, protein turnover, chaperones |
| gene_1424971 | 942,2083629 | -2,844227472 | 0,664722652 | -4,278818336 | 1,88E-05 | 0,001586143 | The Moose: suction sampler vs FISH | Posttranslational modification, protein turnover, chaperones |
| gene_1023468 | 99,55132797 | -2,736451752 | 0,737068368 | -3,712615911 | 0,000205128 | 0,007954398 | The Moose: suction sampler vs FISH | Posttranslational modification, protein turnover, chaperones |
| gene_851535 | 297,7859551 | -2,733165119 | 0,580099834 | -4,711542669 | 2,46E-06 | 0,000400363 | The Moose: suction sampler vs FISH | Posttranslational modification, protein turnover, chaperones |
| gene_1579769 | 83,78394748 | -2,729742895 | 0,755786265 | -3,611792145 | 0,000304088 | 0,010285388 | The Moose: suction sampler vs FISH | Posttranslational modification, protein turnover, chaperones |
| gene_746924 | 152,4258879 | -2,716786234 | 0,706109619 | -3,847541743 | 0,000119309 | 0,005620684 | The Moose: suction sampler vs FISH | Posttranslational modification, protein turnover, chaperones |
| gene_439155 | 1927,341932 | -2,642596362 | 0,649817251 | -4,066676218 | 4,77E-05 | 0,003069655 | The Moose: suction sampler vs FISH | Posttranslational modification, protein turnover, chaperones |
| gene_950470 | 190,4831343 | -2,642184955 | 0,627912846 | -4,207884857 | 2,58E-05 | 0,002007468 | The Moose: suction sampler vs FISH | Posttranslational modification, protein turnover, chaperones |
| gene_851536 | 253,7186679 | -2,630572343 | 0,67646019 | -3,888731934 | 0,000100769 | 0,005024443 | The Moose: suction sampler vs FISH | Posttranslational modification, protein turnover, chaperones |
| gene_72104 | 45,53526024 | -2,565704535 | 0,79797708 | -3,215260938 | 0,00130326 | 0,025586188 | The Moose: suction sampler vs FISH | Posttranslational modification, protein turnover, chaperones |
| gene_829539 | 511,9784552 | -2,486325674 | 0,681982964 | -3,645729885 | 0,000266634 | 0,009438241 | The Moose: suction sampler vs FISH | Posttranslational modification, protein turnover, chaperones |
| gene_1661545 | 35,56862288 | -2,348071321 | 0,804811113 | -2,917543364 | 0,003528006 | 0,046245653 | The Moose: suction sampler vs FISH | Posttranslational modification, protein turnover, chaperones |
| gene_589834 | 33,26020312 | -2,206530159 | 0,766032689 | -2,880464752 | 0,003970894 | 0,049588781 | The Moose: suction sampler vs FISH | Posttranslational modification, protein turnover, chaperones |
| gene_554756 | 341,0409833 | -2,127373456 | 0,56001168 | -3,79880194 | 0,000145397 | 0,006360286 | The Moose: suction sampler vs FISH | Posttranslational modification, protein turnover, chaperones |
| gene_468422 | 83,42266368 | -2,08681437 | 0,720475339 | -2,89644108 | 0,003774215 | 0,048199831 | The Moose: suction sampler vs FISH | Posttranslational modification, protein turnover, chaperones |
| gene_1424972 | 11262,13613 | -2,033740165 | 0,652235322 | -3,118107986 | 0,001820161 | 0,031351249 | The Moose: suction sampler vs FISH | Posttranslational modification, protein turnover, chaperones |
| gene_405404 | 47,25427621 | 2,254205792 | 0,763171431 | 2,953734508 | 0,00313954 | 0,043306684 | The Moose: suction sampler vs FISH | Posttranslational modification, protein turnover, chaperones |
| gene_768739 | 73,13651619 | 2,261093934 | 0,784901114 | 2,880737324 | 0,003967461 | 0,049588452 | The Moose: suction sampler vs FISH | Posttranslational modification, protein turnover, chaperones |
| gene_27052 | 110,2899639 | 2,265553863 | 0,769691576 | 2,943456748 | 0,003245692 | 0,044144413 | The Moose: suction sampler vs FISH | Posttranslational modification, protein turnover, chaperones |
| gene_1457397 | 554,5073265 | 2,320532389 | 0,675306334 | 3,436266288 | 0,000589791 | 0,015475955 | The Moose: suction sampler vs FISH | Posttranslational modification, protein turnover, chaperones |
| gene_1329480 | 57,31273471 | 2,344828349 | 0,68108014 | 3,442808286 | 0,000575707 | 0,015257613 | The Moose: suction sampler vs FISH | Posttranslational modification, protein turnover, chaperones |
| gene_190380 | 2670,619916 | 2,354163783 | 0,713327126 | 3,300258322 | 0,000965959 | 0,021272815 | The Moose: suction sampler vs FISH | Posttranslational modification, protein turnover, chaperones |
| gene_1481827 | 46,38440031 | 2,373903785 | 0,819537697 | 2,896637695 | 0,003771851 | 0,048199831 | The Moose: suction sampler vs FISH | Posttranslational modification, protein turnover, chaperones |
| gene_262454 | 1655,528746 | 2,391614141 | 0,659118024 | 3,628506658 | 0,000285065 | 0,00985901 | The Moose: suction sampler vs FISH | Posttranslational modification, protein turnover, chaperones |
| gene_1592063 | 68,69220926 | 2,395520592 | 0,780448482 | 3,069415403 | 0,002144781 | 0,03458254 | The Moose: suction sampler vs FISH | Posttranslational modification, protein turnover, chaperones |
| gene_1034356 | 33,5482422 | 2,427110419 | 0,721294517 | 3,364936739 | 0,000765612 | 0,01840749 | The Moose: suction sampler vs FISH | Posttranslational modification, protein turnover, chaperones |
| gene_11156 | 187,0013526 | 2,432774912 | 0,69003535 | 3,525580121 | 0,000422556 | 0,012610585 | The Moose: suction sampler vs FISH | Posttranslational modification, protein turnover, chaperones |
| gene_1329730 | 72,60783954 | 2,433689788 | 0,732794437 | 3,321108439 | 0,000896607 | 0,020466154 | The Moose: suction sampler vs FISH | Posttranslational modification, protein turnover, chaperones |
| gene_99150 | 71,46543086 | 2,562228028 | 0,755826879 | 3,389966801 | 0,000699011 | 0,017274177 | The Moose: suction sampler vs FISH | Posttranslational modification, protein turnover, chaperones |
| gene_463046 | 47,60845509 | 2,565773248 | 0,873757017 | 2,936483711 | 0,003319563 | 0,044733024 | The Moose: suction sampler vs FISH | Posttranslational modification, protein turnover, chaperones |
| gene_1913971 | 89,69717343 | 2,569609897 | 0,86342948 | 2,976050687 | 0,002919866 | 0,041455549 | The Moose: suction sampler vs FISH | Posttranslational modification, protein turnover, chaperones |
| gene_1621800 | 45,29338953 | 2,572957676 | 0,717652508 | 3,585241669 | 0,000336766 | 0,010927321 | The Moose: suction sampler vs FISH | Posttranslational modification, protein turnover, chaperones |
| gene_284756 | 32,97589935 | 2,591140124 | 0,897602102 | 2,886735803 | 0,003892609 | 0,049030393 | The Moose: suction sampler vs FISH | Posttranslational modification, protein turnover, chaperones |
| gene_53885 | 97,32831929 | 2,610188204 | 0,729802272 | 3,576569028 | 0,000348133 | 0,011150745 | The Moose: suction sampler vs FISH | Posttranslational modification, protein turnover, chaperones |
| gene_1209228 | 27,75125513 | 2,615267598 | 0,793558126 | 3,295621973 | 0,000982041 | 0,021478849 | The Moose: suction sampler vs FISH | Posttranslational modification, protein turnover, chaperones |
| gene_1830345 | 36,77740341 | 2,615832171 | 0,907571579 | 2,882232357 | 0,003948684 | 0,049430905 | The Moose: suction sampler vs FISH | Posttranslational modification, protein turnover, chaperones |

| | | | | | | | | |
|---|---|---|---|---|---|---|---|---|
| gene_771487 | 15,24336633 | 2,647518367 | 0,905467074 | 2,92392561 | 0,003456473 | 0,045715953 | The Moose: suction sampler vs FISH | Posttranslational modification, protein turnover, chaperones |
| gene_642100 | 28,54316477 | 2,695820379 | 0,871683777 | 3,092658655 | 0,001983722 | 0,033037287 | The Moose: suction sampler vs FISH | Posttranslational modification, protein turnover, chaperones |
| gene_1229780 | 44,15113615 | 2,70371137 | 0,846093371 | 3,195523643 | 0,001395773 | 0,026640484 | The Moose: suction sampler vs FISH | Posttranslational modification, protein turnover, chaperones |
| gene_1477870 | 38,42723776 | 2,703973069 | 0,822127573 | 3,288994505 | 0,00100546 | 0,021789908 | The Moose: suction sampler vs FISH | Posttranslational modification, protein turnover, chaperones |
| gene_1378315 | 40,0307327 | 2,727026661 | 0,868191883 | 3,141041414 | 0,001683482 | 0,029966882 | The Moose: suction sampler vs FISH | Posttranslational modification, protein turnover, chaperones |
| gene_494914 | 40,58552955 | 2,73830832 | 0,895097355 | 3,059229596 | 0,00221907 | 0,03520092 | The Moose: suction sampler vs FISH | Posttranslational modification, protein turnover, chaperones |
| gene_1932422 | 382,6694959 | 2,752308515 | 0,643308629 | 4,278364053 | 1,88E-05 | 0,001587558 | The Moose: suction sampler vs FISH | Posttranslational modification, protein turnover, chaperones |
| gene_1799905 | 47,37864836 | 2,786242812 | 0,862279909 | 3,231250992 | 0,001232497 | 0,024687463 | The Moose: suction sampler vs FISH | Posttranslational modification, protein turnover, chaperones |
| gene_794849 | 25,97581002 | 2,788925483 | 0,905040649 | 3,081547205 | 0,002059278 | 0,033794162 | The Moose: suction sampler vs FISH | Posttranslational modification, protein turnover, chaperones |
| gene_795412 | 31,52648301 | 2,803897372 | 0,900335878 | 3,114279283 | 0,001843949 | 0,031629226 | The Moose: suction sampler vs FISH | Posttranslational modification, protein turnover, chaperones |
| gene_976213 | 18,84536803 | 2,804583624 | 0,89599078 | 3,130147861 | 0,001747183 | 0,030638783 | The Moose: suction sampler vs FISH | Posttranslational modification, protein turnover, chaperones |
| gene_1892050 | 28,35244763 | 2,810293313 | 0,783321228 | 3,587663927 | 0,000333654 | 0,010886847 | The Moose: suction sampler vs FISH | Posttranslational modification, protein turnover, chaperones |
| gene_273439 | 22,97340059 | 2,828986688 | 0,979648365 | 2,887757268 | 0,003879992 | 0,048953312 | The Moose: suction sampler vs FISH | Posttranslational modification, protein turnover, chaperones |
| gene_1898118 | 60,18811642 | 2,831730458 | 0,772066063 | 3,667730771 | 0,000244713 | 0,008928426 | The Moose: suction sampler vs FISH | Posttranslational modification, protein turnover, chaperones |
| gene_368317 | 58,33308031 | 2,841616819 | 0,850833596 | 3,339803261 | 0,000838378 | 0,01956614 | The Moose: suction sampler vs FISH | Posttranslational modification, protein turnover, chaperones |
| gene_1314268 | 15,65145467 | 2,858486542 | 0,979128064 | 2,919420499 | 0,003506828 | 0,046082171 | The Moose: suction sampler vs FISH | Posttranslational modification, protein turnover, chaperones |
| gene_1109528 | 32,02594328 | 2,859512824 | 0,93795527 | 3,048666513 | 0,002298595 | 0,035842951 | The Moose: suction sampler vs FISH | Posttranslational modification, protein turnover, chaperones |
| gene_746802 | 13,77684697 | 2,867150562 | 0,989213538 | 2,898414197 | 0,003750549 | 0,0480412 | The Moose: suction sampler vs FISH | Posttranslational modification, protein turnover, chaperones |
| gene_654911 | 131,1576395 | 2,874653851 | 0,78955514 | 3,640852559 | 0,000271737 | 0,009538157 | The Moose: suction sampler vs FISH | Posttranslational modification, protein turnover, chaperones |
| gene_1756979 | 20,11966217 | 2,881443602 | 0,926119739 | 3,111307836 | 0,001862607 | 0,031821154 | The Moose: suction sampler vs FISH | Posttranslational modification, protein turnover, chaperones |
| gene_1083989 | 1451,80497 | 2,899308274 | 0,735044203 | 3,944399888 | 8,00E-05 | 0,004291895 | The Moose: suction sampler vs FISH | Posttranslational modification, protein turnover, chaperones |
| gene_16752 | 117,4085461 | 2,912785804 | 0,761046425 | 3,82734313 | 0,000129534 | 0,005912751 | The Moose: suction sampler vs FISH | Posttranslational modification, protein turnover, chaperones |
| gene_891652 | 17,34447139 | 2,921713472 | 0,951633591 | 3,070208428 | 0,002139094 | 0,034518587 | The Moose: suction sampler vs FISH | Posttranslational modification, protein turnover, chaperones |
| gene_282898 | 33,80618508 | 2,92880455 | 1,002236648 | 2,922268465 | 0,003474919 | 0,045877298 | The Moose: suction sampler vs FISH | Posttranslational modification, protein turnover, chaperones |
| gene_1820372 | 18,55535219 | 2,929966439 | 0,971786299 | 3,015031642 | 0,002569524 | 0,038435586 | The Moose: suction sampler vs FISH | Posttranslational modification, protein turnover, chaperones |
| gene_56491 | 12,40489089 | 2,944792987 | 0,995085633 | 2,95933625 | 0,003083025 | 0,042893118 | The Moose: suction sampler vs FISH | Posttranslational modification, protein turnover, chaperones |
| gene_972751 | 17,68603498 | 2,947072395 | 0,980249954 | 3,00644992 | 0,002643176 | 0,03898959 | The Moose: suction sampler vs FISH | Posttranslational modification, protein turnover, chaperones |
| gene_189473 | 56,36586564 | 2,9492946 | 0,768872052 | 3,83587177 | 0,00012512 | 0,005797411 | The Moose: suction sampler vs FISH | Posttranslational modification, protein turnover, chaperones |
| gene_425417 | 44,96813917 | 2,955827301 | 0,982801932 | 3,007551374 | 0,002633616 | 0,038947916 | The Moose: suction sampler vs FISH | Posttranslational modification, protein turnover, chaperones |
| gene_995803 | 33,00282082 | 2,964819209 | 0,837599991 | 3,539660028 | 0,000400643 | 0,01221018 | The Moose: suction sampler vs FISH | Posttranslational modification, protein turnover, chaperones |
| gene_1791473 | 14,96285843 | 2,97141506 | 0,916912981 | 3,24067291 | 0,001192479 | 0,024240697 | The Moose: suction sampler vs FISH | Posttranslational modification, protein turnover, chaperones |
| gene_417352 | 19,56968314 | 3,043718956 | 1,002468422 | 3,036224272 | 0,002395611 | 0,036739539 | The Moose: suction sampler vs FISH | Posttranslational modification, protein turnover, chaperones |
| gene_972750 | 23,4028995 | 3,059489692 | 0,958993264 | 3,190313016 | 0,001421188 | 0,026978323 | The Moose: suction sampler vs FISH | Posttranslational modification, protein turnover, chaperones |
| gene_115006 | 1612,348687 | 3,06219752 | 0,699927564 | 4,375020614 | 1,21E-05 | 0,001203799 | The Moose: suction sampler vs FISH | Posttranslational modification, protein turnover, chaperones |
| gene_581036 | 24,38788781 | 3,123767506 | 0,989249192 | 3,157715499 | 0,001590107 | 0,028878856 | The Moose: suction sampler vs FISH | Posttranslational modification, protein turnover, chaperones |
| gene_823685 | 65,48273953 | 3,125623975 | 0,962151635 | 3,248577316 | 0,001159837 | 0,023841091 | The Moose: suction sampler vs FISH | Posttranslational modification, protein turnover, chaperones |
| gene_1527163 | 11,21613108 | 3,126552851 | 0,976197196 | 3,202788191 | 0,00136104 | 0,026236538 | The Moose: suction sampler vs FISH | Posttranslational modification, protein turnover, chaperones |
| gene_117747 | 10,29768684 | 3,141781341 | 1,074423691 | 2,924154936 | 0,003453928 | 0,045706277 | The Moose: suction sampler vs FISH | Posttranslational modification, protein turnover, chaperones |
| gene_978080 | 18,75528485 | 3,148571055 | 0,969353223 | 3,248115323 | 0,001161722 | 0,023866477 | The Moose: suction sampler vs FISH | Posttranslational modification, protein turnover, chaperones |
| gene_1873948 | 12,47958246 | 3,151109088 | 1,045665861 | 3,013495235 | 0,002582571 | 0,03856156 | The Moose: suction sampler vs FISH | Posttranslational modification, protein turnover, chaperones |
| gene_1859892 | 18,27467688 | 3,160027898 | 1,063637282 | 2,970963836 | 0,002968667 | 0,041849093 | The Moose: suction sampler vs FISH | Posttranslational modification, protein turnover, chaperones |
| gene_795459 | 36,64448036 | 3,1733222 | 0,960603579 | 3,303466976 | 0,000954972 | 0,021125883 | The Moose: suction sampler vs FISH | Posttranslational modification, protein turnover, chaperones |
| gene_1431786 | 17,03970345 | 3,176599884 | 1,058057878 | 3,002293118 | 0,00267954 | 0,039306464 | The Moose: suction sampler vs FISH | Posttranslational modification, protein turnover, chaperones |
| gene_633547 | 17,55711779 | 3,184779261 | 1,0681546 | 2,981571452 | 0,002867731 | 0,041011106 | The Moose: suction sampler vs FISH | Posttranslational modification, protein turnover, chaperones |
| gene_1268426 | 29,98045344 | 3,229400497 | 0,808767433 | 3,992990278 | 6,52E-05 | 0,003741521 | The Moose: suction sampler vs FISH | Posttranslational modification, protein turnover, chaperones |
| gene_452990 | 335,5231742 | 3,233203873 | 0,6825627 | 4,736859883 | 2,17E-06 | 0,000366549 | The Moose: suction sampler vs FISH | Posttranslational modification, protein turnover, chaperones |
| gene_1146920 | 74,85987518 | 3,239865308 | 0,855274587 | 3,788099587 | 0,000151804 | 0,00650481 | The Moose: suction sampler vs FISH | Posttranslational modification, protein turnover, chaperones |
| gene_924376 | 20,90574053 | 3,242263764 | 1,02511625 | 3,162825448 | 0,00156246 | 0,028588655 | The Moose: suction sampler vs FISH | Posttranslational modification, protein turnover, chaperones |
| gene_313689 | 231,9979221 | 3,24393482 | 0,747841194 | 4,337732188 | 1,44E-05 | 0,001337845 | The Moose: suction sampler vs FISH | Posttranslational modification, protein turnover, chaperones |
| gene_51145 | 43,75191524 | 3,249128189 | 0,788885298 | 4,118631943 | 3,81E-05 | 0,002616072 | The Moose: suction sampler vs FISH | Posttranslational modification, protein turnover, chaperones |
| gene_1467279 | 20,25097274 | 3,255117861 | 1,00391485 | 3,242424256 | 0,001185174 | 0,02414567 | The Moose: suction sampler vs FISH | Posttranslational modification, protein turnover, chaperones |
| gene_1722724 | 18,49648676 | 3,273486957 | 0,943950762 | 3,46785774 | 0,000524625 | 0,014377268 | The Moose: suction sampler vs FISH | Posttranslational modification, protein turnover, chaperones |
| gene_646092 | 13,50426885 | 3,275072966 | 1,118359668 | 2,928461263 | 0,003406443 | 0,045337352 | The Moose: suction sampler vs FISH | Posttranslational modification, protein turnover, chaperones |
| gene_1540237 | 1516,138857 | 3,286799804 | 0,892671565 | 3,681981071 | 0,000231429 | 0,008601859 | The Moose: suction sampler vs FISH | Posttranslational modification, protein turnover, chaperones |
| gene_179107 | 419,2594578 | 3,292480089 | 0,742301372 | 4,43550317 | 9,19E-06 | 0,001005436 | The Moose: suction sampler vs FISH | Posttranslational modification, protein turnover, chaperones |
| gene_1365554 | 11,89996367 | 3,304056537 | 1,085210501 | 3,04462271 | 0,002329724 | 0,036151818 | The Moose: suction sampler vs FISH | Posttranslational modification, protein turnover, chaperones |
| gene_1091747 | 26,52661689 | 3,308827898 | 0,932548629 | 3,548155877 | 0,000387939 | 0,011961597 | The Moose: suction sampler vs FISH | Posttranslational modification, protein turnover, chaperones |
| gene_549369 | 23,86258041 | 3,309522034 | 0,960803266 | 3,44453662 | 0,000572039 | 0,01523329 | The Moose: suction sampler vs FISH | Posttranslational modification, protein turnover, chaperones |
| gene_1273875 | 23,19898753 | 3,317967865 | 1,056307565 | 3,141100164 | 0,001683145 | 0,029966882 | The Moose: suction sampler vs FISH | Posttranslational modification, protein turnover, chaperones |
| gene_822883 | 22,52964896 | 3,326934264 | 0,996975756 | 3,337026246 | 0,0008468 | 0,019705941 | The Moose: suction sampler vs FISH | Posttranslational modification, protein turnover, chaperones |
| gene_962885 | 28,22235111 | 3,364107138 | 1,023615595 | 3,286494611 | 0,001014427 | 0,021906675 | The Moose: suction sampler vs FISH | Posttranslational modification, protein turnover, chaperones |
| gene_1697090 | 14,33003872 | 3,382066773 | 1,105799995 | 3,058479642 | 0,002224632 | 0,035225748 | The Moose: suction sampler vs FISH | Posttranslational modification, protein turnover, chaperones |
| gene_84767 | 12,50850746 | 3,389726383 | 1,061364831 | 3,193742889 | 0,001404411 | 0,026755825 | The Moose: suction sampler vs FISH | Posttranslational modification, protein turnover, chaperones |
| gene_1758060 | 13,95935954 | 3,40025884 | 1,034857025 | 3,285728134 | 0,001017191 | 0,021940563 | The Moose: suction sampler vs FISH | Posttranslational modification, protein turnover, chaperones |
| gene_532325 | 36,50364023 | 3,421452207 | 0,936562766 | 3,653201184 | 0,000258991 | 0,009256255 | The Moose: suction sampler vs FISH | Posttranslational modification, protein turnover, chaperones |

| | | | | | | | | |
|---|---|---|---|---|---|---|---|---|
| gene_1379019 | 13,52468205 | 3,426784918 | 1,151706849 | 2,97539684 | 0,002926097 | 0,041503598 | The Moose: suction sampler vs FISH | Posttranslational modification, protein turnover, chaperones |
| gene_823682 | 34,35929512 | 3,432549028 | 0,900636963 | 3,811246006 | 0,000138268 | 0,006163751 | The Moose: suction sampler vs FISH | Posttranslational modification, protein turnover, chaperones |
| gene_1789935 | 268,3188807 | 3,468475124 | 0,755470764 | 4,59114408 | 4,41E-06 | 0,000605163 | The Moose: suction sampler vs FISH | Posttranslational modification, protein turnover, chaperones |
| gene_823684 | 16,878193 | 3,482170085 | 1,050869899 | 3,313607219 | 0,000921008 | 0,020702209 | The Moose: suction sampler vs FISH | Posttranslational modification, protein turnover, chaperones |
| gene_637951 | 18,08999098 | 3,499843371 | 1,02542729 | 3,413058541 | 0,000642381 | 0,01638018 | The Moose: suction sampler vs FISH | Posttranslational modification, protein turnover, chaperones |
| gene_1035196 | 9,301249467 | 3,50470333 | 1,163823053 | 3,01137129 | 0,002600706 | 0,038749472 | The Moose: suction sampler vs FISH | Posttranslational modification, protein turnover, chaperones |
| gene_773236 | 287,67047 | 3,513726331 | 0,868220048 | 4,047045838 | 5,19E-05 | 0,003223638 | The Moose: suction sampler vs FISH | Posttranslational modification, protein turnover, chaperones |
| gene_1762125 | 10,55303626 | 3,516912241 | 1,09518855 | 3,211239052 | 0,001321639 | 0,025815906 | The Moose: suction sampler vs FISH | Posttranslational modification, protein turnover, chaperones |
| gene_990221 | 19,67329522 | 3,520236694 | 0,99948696 | 3,522043642 | 0,000428234 | 0,012697465 | The Moose: suction sampler vs FISH | Posttranslational modification, protein turnover, chaperones |
| gene_773237 | 41,80432071 | 3,525998915 | 1,003088522 | 3,515142321 | 0,000439518 | 0,012913873 | The Moose: suction sampler vs FISH | Posttranslational modification, protein turnover, chaperones |
| gene_918224 | 45,7083687 | 3,544720713 | 0,793124044 | 4,469314405 | 7,85E-06 | 0,000907991 | The Moose: suction sampler vs FISH | Posttranslational modification, protein turnover, chaperones |
| gene_1094512 | 9,505522456 | 3,559868197 | 1,222009278 | 2,913126979 | 0,003578291 | 0,04667156 | The Moose: suction sampler vs FISH | Posttranslational modification, protein turnover, chaperones |
| gene_1768841 | 63,86608925 | 3,563038945 | 0,812177855 | 4,387018092 | 1,15E-05 | 0,001165739 | The Moose: suction sampler vs FISH | Posttranslational modification, protein turnover, chaperones |
| gene_989587 | 51,89095502 | 3,574647309 | 0,736775997 | 4,85174235 | 1,22E-06 | 0,000253907 | The Moose: suction sampler vs FISH | Posttranslational modification, protein turnover, chaperones |
| gene_823688 | 178,8650216 | 3,579241613 | 0,753434074 | 4,750570404 | 2,03E-06 | 0,000350538 | The Moose: suction sampler vs FISH | Posttranslational modification, protein turnover, chaperones |
| gene_1464687 | 13,30779904 | 3,589608296 | 1,193018255 | 3,008846076 | 0,002622419 | 0,038870554 | The Moose: suction sampler vs FISH | Posttranslational modification, protein turnover, chaperones |
| gene_1009555 | 11,00258216 | 3,595725573 | 1,162480346 | 3,093149561 | 0,001980443 | 0,033005136 | The Moose: suction sampler vs FISH | Posttranslational modification, protein turnover, chaperones |
| gene_937370 | 47,7211097 | 3,599032958 | 0,988860005 | 3,639577835 | 0,000273085 | 0,009559941 | The Moose: suction sampler vs FISH | Posttranslational modification, protein turnover, chaperones |
| gene_1750531 | 12,61064775 | 3,623717267 | 1,031609383 | 3,512683507 | 0,000443606 | 0,012965789 | The Moose: suction sampler vs FISH | Posttranslational modification, protein turnover, chaperones |
| gene_1432925 | 8,243013865 | 3,644807678 | 1,256066313 | 2,901763736 | 0,003710683 | 0,047787322 | The Moose: suction sampler vs FISH | Posttranslational modification, protein turnover, chaperones |
| gene_821812 | 11,78979955 | 3,650391796 | 1,044293417 | 3,495561435 | 0,000473066 | 0,013456351 | The Moose: suction sampler vs FISH | Posttranslational modification, protein turnover, chaperones |
| gene_760034 | 11,85426416 | 3,653192733 | 1,189420078 | 3,071406646 | 0,002130528 | 0,034451037 | The Moose: suction sampler vs FISH | Posttranslational modification, protein turnover, chaperones |
| gene_1224816 | 19,0114717 | 3,655565518 | 0,974590702 | 3,750872556 | 0,00017622 | 0,007186284 | The Moose: suction sampler vs FISH | Posttranslational modification, protein turnover, chaperones |
| gene_1813864 | 56,96133669 | 3,656321036 | 0,761736004 | 4,799984528 | 1,59E-06 | 0,000303492 | The Moose: suction sampler vs FISH | Posttranslational modification, protein turnover, chaperones |
| gene_1155295 | 201,7992412 | 3,664024739 | 0,799410674 | 4,583407326 | 4,57E-06 | 0,000621589 | The Moose: suction sampler vs FISH | Posttranslational modification, protein turnover, chaperones |
| gene_1755134 | 52,90142388 | 3,664049851 | 1,047855751 | 3,49671207 | 0,00047103 | 0,013424445 | The Moose: suction sampler vs FISH | Posttranslational modification, protein turnover, chaperones |
| gene_1166019 | 44,92291745 | 3,669153176 | 0,759098997 | 4,833563464 | 1,34E-06 | 0,000271629 | The Moose: suction sampler vs FISH | Posttranslational modification, protein turnover, chaperones |
| gene_1616838 | 51,25178812 | 3,679312169 | 1,042520902 | 3,529245467 | 0,000416746 | 0,012493036 | The Moose: suction sampler vs FISH | Posttranslational modification, protein turnover, chaperones |
| gene_86924 | 20,25860147 | 3,687841429 | 0,890072674 | 4,143303728 | 3,42E-05 | 0,002429277 | The Moose: suction sampler vs FISH | Posttranslational modification, protein turnover, chaperones |
| gene_867027 | 13,82805888 | 3,692880693 | 1,176953216 | 3,137661414 | 0,001703015 | 0,030146522 | The Moose: suction sampler vs FISH | Posttranslational modification, protein turnover, chaperones |
| gene_533058 | 32,99665188 | 3,70053161 | 0,903536201 | 4,095609678 | 4,21E-05 | 0,002806231 | The Moose: suction sampler vs FISH | Posttranslational modification, protein turnover, chaperones |
| gene_1429881 | 10,73275391 | 3,751077003 | 1,085816733 | 3,45461337 | 0,000551082 | 0,014825155 | The Moose: suction sampler vs FISH | Posttranslational modification, protein turnover, chaperones |
| gene_1186591 | 12,9598589 | 3,756884058 | 1,26634009 | 2,96672599 | 0,003009891 | 0,042267959 | The Moose: suction sampler vs FISH | Posttranslational modification, protein turnover, chaperones |
| gene_785948 | 11,59885416 | 3,761886175 | 1,286036611 | 2,925178135 | 0,003442591 | 0,045647426 | The Moose: suction sampler vs FISH | Posttranslational modification, protein turnover, chaperones |
| gene_1823056 | 12,13181113 | 3,768216311 | 1,163717654 | 3,238084682 | 0,001203351 | 0,024380718 | The Moose: suction sampler vs FISH | Posttranslational modification, protein turnover, chaperones |
| gene_319747 | 34,67778236 | 3,812839813 | 0,838363419 | 4,547955844 | 5,42E-06 | 0,000696756 | The Moose: suction sampler vs FISH | Posttranslational modification, protein turnover, chaperones |
| gene_1731729 | 9,633591896 | 3,824037204 | 1,282616671 | 2,981434196 | 0,002869017 | 0,041019068 | The Moose: suction sampler vs FISH | Posttranslational modification, protein turnover, chaperones |
| gene_1194250 | 20,35179175 | 3,829948246 | 1,075421899 | 3,561344854 | 0,00036896 | 0,011558516 | The Moose: suction sampler vs FISH | Posttranslational modification, protein turnover, chaperones |
| gene_461717 | 8,60784684 | 3,847478325 | 1,286076485 | 2,991640365 | 0,002774829 | 0,040149201 | The Moose: suction sampler vs FISH | Posttranslational modification, protein turnover, chaperones |
| gene_54558 | 69,26648556 | 3,888002359 | 0,873596089 | 4,450572076 | 8,56E-06 | 0,000962733 | The Moose: suction sampler vs FISH | Posttranslational modification, protein turnover, chaperones |
| gene_983641 | 17,68332657 | 3,893309065 | 1,016848801 | 3,828798404 | 0,00012877 | 0,005892551 | The Moose: suction sampler vs FISH | Posttranslational modification, protein turnover, chaperones |
| gene_532328 | 20,76537721 | 3,913278742 | 1,015555294 | 3,853338922 | 0,000116518 | 0,005524635 | The Moose: suction sampler vs FISH | Posttranslational modification, protein turnover, chaperones |
| gene_867025 | 8,461936451 | 3,922517081 | 1,308549446 | 2,997607077 | 0,002721082 | 0,039660628 | The Moose: suction sampler vs FISH | Posttranslational modification, protein turnover, chaperones |
| gene_91715 | 11,16321249 | 3,945660471 | 1,283842528 | 3,07332121 | 0,002116905 | 0,034323674 | The Moose: suction sampler vs FISH | Posttranslational modification, protein turnover, chaperones |
| gene_863048 | 6,835435265 | 3,966916591 | 1,368542351 | 2,898643648 | 0,003747806 | 0,0480412 | The Moose: suction sampler vs FISH | Posttranslational modification, protein turnover, chaperones |
| gene_1228928 | 6,035275503 | 4,024945549 | 1,345921775 | 2,990475097 | 0,002785438 | 0,040278898 | The Moose: suction sampler vs FISH | Posttranslational modification, protein turnover, chaperones |
| gene_1155253 | 9,855569617 | 4,027244267 | 1,236485872 | 3,257007911 | 0,001125933 | 0,023426104 | The Moose: suction sampler vs FISH | Posttranslational modification, protein turnover, chaperones |
| gene_1496144 | 15,40713641 | 4,040748359 | 1,403359332 | 2,879339786 | 0,003985087 | 0,049691805 | The Moose: suction sampler vs FISH | Posttranslational modification, protein turnover, chaperones |
| gene_1840914 | 21,72734921 | 4,04472429 | 0,931714487 | 4,34116282 | 1,42E-05 | 0,001332831 | The Moose: suction sampler vs FISH | Posttranslational modification, protein turnover, chaperones |
| gene_1756978 | 9,44222226 | 4,069295187 | 1,179468569 | 3,450109051 | 0,00056036 | 0,014998415 | The Moose: suction sampler vs FISH | Posttranslational modification, protein turnover, chaperones |
| gene_1628917 | 17,0806281 | 4,075109321 | 1,184470774 | 3,440447338 | 0,000580753 | 0,015337446 | The Moose: suction sampler vs FISH | Posttranslational modification, protein turnover, chaperones |
| gene_1390643 | 15,97637727 | 4,078641784 | 1,15475978 | 3,532026189 | 0,000412389 | 0,012407978 | The Moose: suction sampler vs FISH | Posttranslational modification, protein turnover, chaperones |
| gene_304950 | 10,88109409 | 4,093763323 | 1,362193126 | 3,005273807 | 0,002653419 | 0,039069835 | The Moose: suction sampler vs FISH | Posttranslational modification, protein turnover, chaperones |
| gene_1533855 | 22,88209884 | 4,097693455 | 0,951554822 | 4,306313583 | 1,66E-05 | 0,001472428 | The Moose: suction sampler vs FISH | Posttranslational modification, protein turnover, chaperones |
| gene_372299 | 18,52316718 | 4,105547636 | 1,081071856 | 3,797663968 | 0,000146066 | 0,006362888 | The Moose: suction sampler vs FISH | Posttranslational modification, protein turnover, chaperones |
| gene_1863753 | 14,90073308 | 4,106843442 | 1,071405084 | 3,833137909 | 0,000126519 | 0,005828286 | The Moose: suction sampler vs FISH | Posttranslational modification, protein turnover, chaperones |
| gene_277656 | 64,44565218 | 4,130544482 | 0,940581952 | 4,391477505 | 1,13E-05 | 0,001151624 | The Moose: suction sampler vs FISH | Posttranslational modification, protein turnover, chaperones |
| gene_1725649 | 13,72579074 | 4,134612671 | 1,207382542 | 3,42444298 | 0,000616061 | 0,015926296 | The Moose: suction sampler vs FISH | Posttranslational modification, protein turnover, chaperones |
| gene_1075278 | 16,1431333 | 4,143307178 | 1,009811034 | 4,103051994 | 4,08E-05 | 0,00274232 | The Moose: suction sampler vs FISH | Posttranslational modification, protein turnover, chaperones |
| gene_848564 | 11,42000353 | 4,153530636 | 1,26879635 | 3,273599136 | 0,001061871 | 0,022625222 | The Moose: suction sampler vs FISH | Posttranslational modification, protein turnover, chaperones |
| gene_467737 | 5,866898855 | 4,153966814 | 1,389758191 | 2,988985308 | 0,002799056 | 0,040404218 | The Moose: suction sampler vs FISH | Posttranslational modification, protein turnover, chaperones |
| gene_952679 | 30,77901501 | 4,165105137 | 1,123492338 | 3,707283973 | 0,000209494 | 0,008049569 | The Moose: suction sampler vs FISH | Posttranslational modification, protein turnover, chaperones |
| gene_1505282 | 7,090732197 | 4,169971288 | 1,417061368 | 2,942689274 | 0,003253748 | 0,044172188 | The Moose: suction sampler vs FISH | Posttranslational modification, protein turnover, chaperones |
| gene_338901 | 25,29369593 | 4,182217565 | 1,107070133 | 3,777734977 | 0,000158261 | 0,006722667 | The Moose: suction sampler vs FISH | Posttranslational modification, protein turnover, chaperones |
| gene_1234637 | 5,564898151 | 4,20179537 | 1,355710112 | 3,099331733 | 0,001939577 | 0,03259031 | The Moose: suction sampler vs FISH | Posttranslational modification, protein turnover, chaperones |

| | | | | | | | | |
|---|---|---|---|---|---|---|---|---|
| gene_334248 | 8,327515342 | 4,289263723 | 1,443269525 | 2,971907637 | 0,002959557 | 0,041784826 | The Moose: suction sampler vs FISH | Posttranslational modification, protein turnover, chaperones |
| gene_10493 | 12,58538441 | 4,298270234 | 1,491717506 | 2,881423739 | 0,00395883 | 0,04952415 | The Moose: suction sampler vs FISH | Posttranslational modification, protein turnover, chaperones |
| gene_1753797 | 12,08239789 | 4,308341808 | 1,286766352 | 3,348192778 | 0,000813404 | 0,019147584 | The Moose: suction sampler vs FISH | Posttranslational modification, protein turnover, chaperones |
| gene_1342036 | 12,09328114 | 4,316832028 | 1,307262718 | 3,302191647 | 0,000959325 | 0,021190263 | The Moose: suction sampler vs FISH | Posttranslational modification, protein turnover, chaperones |
| gene_1054532 | 9,357425863 | 4,332538277 | 1,303570578 | 3,323593176 | 0,000888657 | 0,020332534 | The Moose: suction sampler vs FISH | Posttranslational modification, protein turnover, chaperones |
| gene_10490 | 6,680562555 | 4,335072827 | 1,376973496 | 3,148261633 | 0,001642446 | 0,029500968 | The Moose: suction sampler vs FISH | Posttranslational modification, protein turnover, chaperones |
| gene_90242 | 10,99630532 | 4,347173455 | 1,234894845 | 3,520278242 | 0,000431094 | 0,012746265 | The Moose: suction sampler vs FISH | Posttranslational modification, protein turnover, chaperones |
| gene_1212139 | 12,36377472 | 4,352397378 | 1,289637663 | 3,374899402 | 0,000738427 | 0,017993947 | The Moose: suction sampler vs FISH | Posttranslational modification, protein turnover, chaperones |
| gene_140580 | 30,86297347 | 4,422266063 | 1,002608341 | 4,410761296 | 1,03E-05 | 0,00108387 | The Moose: suction sampler vs FISH | Posttranslational modification, protein turnover, chaperones |
| gene_372300 | 24,69489981 | 4,423616759 | 1,057058116 | 4,184837799 | 2,85E-05 | 0,002154058 | The Moose: suction sampler vs FISH | Posttranslational modification, protein turnover, chaperones |
| gene_84766 | 13,29998955 | 4,429876848 | 1,189772798 | 3,723296461 | 0,000196638 | 0,007725891 | The Moose: suction sampler vs FISH | Posttranslational modification, protein turnover, chaperones |
| gene_758405 | 15,87570599 | 4,432831661 | 1,411825332 | 3,139787593 | 0,001690704 | 0,030029924 | The Moose: suction sampler vs FISH | Posttranslational modification, protein turnover, chaperones |
| gene_637949 | 16,42051697 | 4,436338294 | 1,058236484 | 4,192199344 | 2,76E-05 | 0,002104786 | The Moose: suction sampler vs FISH | Posttranslational modification, protein turnover, chaperones |
| gene_1084454 | 13,75937013 | 4,440849089 | 1,163624387 | 3,816393967 | 0,000135416 | 0,006097882 | The Moose: suction sampler vs FISH | Posttranslational modification, protein turnover, chaperones |
| gene_700029 | 9,716685327 | 4,44511266 | 1,341992396 | 3,312323284 | 0,000925246 | 0,020762195 | The Moose: suction sampler vs FISH | Posttranslational modification, protein turnover, chaperones |
| gene_1512825 | 18,60655929 | 4,446965289 | 1,078787427 | 4,122188651 | 3,75E-05 | 0,002583238 | The Moose: suction sampler vs FISH | Posttranslational modification, protein turnover, chaperones |
| gene_1510940 | 22,53285951 | 4,447560661 | 1,004273798 | 4,428633575 | 9,48E-06 | 0,001025764 | The Moose: suction sampler vs FISH | Posttranslational modification, protein turnover, chaperones |
| gene_112223 | 1047,599456 | 4,475520405 | 0,835093396 | 5,359305232 | 8,35E-08 | 4,26E-05 | The Moose: suction sampler vs FISH | Posttranslational modification, protein turnover, chaperones |
| gene_1793337 | 21,85383701 | 4,493695853 | 1,035282377 | 4,340550898 | 1,42E-05 | 0,001334839 | The Moose: suction sampler vs FISH | Posttranslational modification, protein turnover, chaperones |
| gene_1951269 | 294,1876538 | 4,495599907 | 0,938058218 | 4,792452984 | 1,65E-06 | 0,000308683 | The Moose: suction sampler vs FISH | Posttranslational modification, protein turnover, chaperones |
| gene_590546 | 26,11000674 | 4,546926289 | 1,016840849 | 4,471620406 | 7,76E-06 | 0,000900706 | The Moose: suction sampler vs FISH | Posttranslational modification, protein turnover, chaperones |
| gene_590545 | 22,55918044 | 4,555904756 | 1,139835603 | 3,996984078 | 6,42E-05 | 0,00371303 | The Moose: suction sampler vs FISH | Posttranslational modification, protein turnover, chaperones |
| gene_1316223 | 8,921110778 | 4,56752304 | 1,336378042 | 3,41783754 | 0,000631208 | 0,01618237 | The Moose: suction sampler vs FISH | Posttranslational modification, protein turnover, chaperones |
| gene_832910 | 30,86048532 | 4,578591186 | 1,098771453 | 4,167009594 | 3,09E-05 | 0,002259891 | The Moose: suction sampler vs FISH | Posttranslational modification, protein turnover, chaperones |
| gene_1232343 | 6,696843013 | 4,58381767 | 1,463599686 | 3,131879376 | 0,001736912 | 0,030507633 | The Moose: suction sampler vs FISH | Posttranslational modification, protein turnover, chaperones |
| gene_338900 | 14,31237434 | 4,602855506 | 1,1406848 | 4,035168616 | 5,46E-05 | 0,003320122 | The Moose: suction sampler vs FISH | Posttranslational modification, protein turnover, chaperones |
| gene_837573 | 9,082041963 | 4,603206862 | 1,353160019 | 3,401820034 | 0,000669387 | 0,016813656 | The Moose: suction sampler vs FISH | Posttranslational modification, protein turnover, chaperones |
| gene_1516831 | 74,02407083 | 4,622923648 | 0,792476287 | 5,83351669 | 5,43E-09 | 5,69E-06 | The Moose: suction sampler vs FISH | Posttranslational modification, protein turnover, chaperones |
| gene_102368 | 40,0585148 | 4,632784452 | 0,941505941 | 4,920610958 | 8,63E-07 | 0,000202263 | The Moose: suction sampler vs FISH | Posttranslational modification, protein turnover, chaperones |
| gene_1288495 | 40,07118821 | 4,670036796 | 0,9532377 | 4,89913145 | 9,63E-07 | 0,000218207 | The Moose: suction sampler vs FISH | Posttranslational modification, protein turnover, chaperones |
| gene_1391274 | 9,226488617 | 4,684643895 | 1,441568214 | 3,249685898 | 0,001155325 | 0,023808326 | The Moose: suction sampler vs FISH | Posttranslational modification, protein turnover, chaperones |
| gene_1512030 | 10,37770942 | 4,685178266 | 1,347214042 | 3,477679209 | 0,000505775 | 0,014057251 | The Moose: suction sampler vs FISH | Posttranslational modification, protein turnover, chaperones |
| gene_911790 | 58,0707792 | 4,693473161 | 0,85044277 | 5,518858328 | 3,41E-08 | 2,22E-05 | The Moose: suction sampler vs FISH | Posttranslational modification, protein turnover, chaperones |
| gene_1091746 | 8,667536405 | 4,73300594 | 1,386010037 | 3,414842472 | 0,000638189 | 0,016320264 | The Moose: suction sampler vs FISH | Posttranslational modification, protein turnover, chaperones |
| gene_1223072 | 11,03298889 | 4,733840568 | 1,322908928 | 3,578357111 | 0,000345761 | 0,011122688 | The Moose: suction sampler vs FISH | Posttranslational modification, protein turnover, chaperones |
| gene_438818 | 28,23097 | 4,765059407 | 1,214719529 | 3,922765127 | 8,75E-05 | 0,004572734 | The Moose: suction sampler vs FISH | Posttranslational modification, protein turnover, chaperones |
| gene_1019221 | 5,269382877 | 4,772626153 | 1,643059991 | 2,904718136 | 0,00367584 | 0,047455096 | The Moose: suction sampler vs FISH | Posttranslational modification, protein turnover, chaperones |
| gene_1805392 | 18,06517219 | 4,789634738 | 1,312413898 | 3,649484926 | 0,000262767 | 0,009332155 | The Moose: suction sampler vs FISH | Posttranslational modification, protein turnover, chaperones |
| gene_546351 | 7,726848358 | 4,853010275 | 1,49301356 | 3,250479705 | 0,001152105 | 0,023755291 | The Moose: suction sampler vs FISH | Posttranslational modification, protein turnover, chaperones |
| gene_91714 | 26,28066379 | 4,854975509 | 0,957883691 | 5,068439472 | 4,01E-07 | 0,000119264 | The Moose: suction sampler vs FISH | Posttranslational modification, protein turnover, chaperones |
| gene_304949 | 6,188885468 | 4,880677235 | 1,566756072 | 3,115148123 | 0,001838526 | 0,031600869 | The Moose: suction sampler vs FISH | Posttranslational modification, protein turnover, chaperones |
| gene_917642 | 27,40370126 | 4,906572586 | 1,215550356 | 4,03650294 | 5,43E-05 | 0,003308096 | The Moose: suction sampler vs FISH | Posttranslational modification, protein turnover, chaperones |
| gene_506501 | 8,86803719 | 4,908817493 | 1,396964741 | 3,513916529 | 0,000441552 | 0,012946009 | The Moose: suction sampler vs FISH | Posttranslational modification, protein turnover, chaperones |
| gene_87929 | 7,919999362 | 4,918695381 | 1,579409191 | 3,114262859 | 0,001844051 | 0,031629226 | The Moose: suction sampler vs FISH | Posttranslational modification, protein turnover, chaperones |
| gene_1496721 | 329,0132228 | 4,938293176 | 0,86885848 | 5,683656534 | 1,32E-08 | 1,08E-05 | The Moose: suction sampler vs FISH | Posttranslational modification, protein turnover, chaperones |
| gene_1426789 | 7,431003386 | 4,944499898 | 1,615601137 | 3,060470673 | 0,002209894 | 0,035159134 | The Moose: suction sampler vs FISH | Posttranslational modification, protein turnover, chaperones |
| gene_133462 | 7,503526428 | 4,949048911 | 1,601382539 | 3,090485121 | 0,001998298 | 0,03315225 | The Moose: suction sampler vs FISH | Posttranslational modification, protein turnover, chaperones |
| gene_85390 | 18,78059348 | 4,955564083 | 1,21239183 | 4,087427809 | 4,36E-05 | 0,00287571 | The Moose: suction sampler vs FISH | Posttranslational modification, protein turnover, chaperones |
| gene_1898117 | 6,450781515 | 4,970758262 | 1,497864194 | 3,318564048 | 0,000904816 | 0,020533806 | The Moose: suction sampler vs FISH | Posttranslational modification, protein turnover, chaperones |
| gene_1621603 | 13,46423416 | 5,058344682 | 1,405719503 | 3,598402578 | 0,000320178 | 0,010602098 | The Moose: suction sampler vs FISH | Posttranslational modification, protein turnover, chaperones |
| gene_1368724 | 749,9224307 | 5,059430025 | 0,949106138 | 5,330731541 | 9,78E-08 | 4,64E-05 | The Moose: suction sampler vs FISH | Posttranslational modification, protein turnover, chaperones |
| gene_860609 | 7,166404997 | 5,062579161 | 1,376607233 | 3,677577045 | 0,00023546 | 0,008698872 | The Moose: suction sampler vs FISH | Posttranslational modification, protein turnover, chaperones |
| gene_303098 | 914,4125481 | 5,074864796 | 0,851853656 | 5,957437362 | 2,56E-09 | 3,30E-06 | The Moose: suction sampler vs FISH | Posttranslational modification, protein turnover, chaperones |
| gene_762381 | 26,64791648 | 5,076825016 | 1,068452649 | 4,751567625 | 2,02E-06 | 0,000350538 | The Moose: suction sampler vs FISH | Posttranslational modification, protein turnover, chaperones |
| gene_886851 | 10,92830806 | 5,091979889 | 1,37001879 | 3,716722665 | 0,000201824 | 0,007871985 | The Moose: suction sampler vs FISH | Posttranslational modification, protein turnover, chaperones |
| gene_855420 | 7,423324482 | 5,118034694 | 1,53005291 | 3,345005039 | 0,000822811 | 0,019294807 | The Moose: suction sampler vs FISH | Posttranslational modification, protein turnover, chaperones |
| gene_853285 | 15,49862949 | 5,119792915 | 1,171531906 | 4,370169425 | 1,24E-05 | 0,001223921 | The Moose: suction sampler vs FISH | Posttranslational modification, protein turnover, chaperones |
| gene_1880556 | 10,92139299 | 5,137153492 | 1,304795136 | 3,937134153 | 8,25E-05 | 0,004391815 | The Moose: suction sampler vs FISH | Posttranslational modification, protein turnover, chaperones |
| gene_596450 | 7,418579521 | 5,144249779 | 1,773079128 | 2,901308631 | 0,003716077 | 0,047824342 | The Moose: suction sampler vs FISH | Posttranslational modification, protein turnover, chaperones |
| gene_1919565 | 5,135478224 | 5,180851047 | 1,677919816 | 3,087663069 | 0,002017371 | 0,033340635 | The Moose: suction sampler vs FISH | Posttranslational modification, protein turnover, chaperones |
| gene_918222 | 8,837846029 | 5,202521587 | 1,517829003 | 3,427607178 | 0,000608926 | 0,015808611 | The Moose: suction sampler vs FISH | Posttranslational modification, protein turnover, chaperones |
| gene_1461567 | 8,226030073 | 5,224402448 | 1,784032614 | 2,928423173 | 0,00340686 | 0,045337352 | The Moose: suction sampler vs FISH | Posttranslational modification, protein turnover, chaperones |
| gene_1502618 | 74,02908822 | 5,312269477 | 1,738012146 | 3,056520342 | 0,002239223 | 0,035360455 | The Moose: suction sampler vs FISH | Posttranslational modification, protein turnover, chaperones |
| gene_1409279 | 723,3742844 | 5,321758553 | 0,914808896 | 5,817344558 | 5,98E-09 | 6,10E-06 | The Moose: suction sampler vs FISH | Posttranslational modification, protein turnover, chaperones |
| gene_1513621 | 32,31498603 | 5,323105721 | 1,143299531 | 4,65591525 | 3,23E-06 | 0,000477607 | The Moose: suction sampler vs FISH | Posttranslational modification, protein turnover, chaperones |

| | | | | | | | | |
|---|---|---|---|---|---|---|---|---|
| gene_604214 | 9,339452806 | 5,324999368 | 1,430198879 | 3,723257966 | 0,000196668 | 0,007725891 | The Moose: suction sampler vs FISH | Posttranslational modification, protein turnover, chaperones |
| gene_1118783 | 6,116435299 | 5,357185394 | 1,718731077 | 3,116942183 | 0,001827374 | 0,03145336 | The Moose: suction sampler vs FISH | Posttranslational modification, protein turnover, chaperones |
| gene_569822 | 4,715997554 | 5,362894815 | 1,646950487 | 3,256257463 | 0,001128913 | 0,023472208 | The Moose: suction sampler vs FISH | Posttranslational modification, protein turnover, chaperones |
| gene_1255597 | 17,52806567 | 5,378082092 | 1,196229614 | 4,495861021 | 6,93E-06 | 0,00083664 | The Moose: suction sampler vs FISH | Posttranslational modification, protein turnover, chaperones |
| gene_1420530 | 6,81545792 | 5,401026689 | 1,641966723 | 3,289364281 | 0,00100414 | 0,021787013 | The Moose: suction sampler vs FISH | Posttranslational modification, protein turnover, chaperones |
| gene_866572 | 5,338118179 | 5,40306405 | 1,820586814 | 2,96775963 | 0,002999788 | 0,042188186 | The Moose: suction sampler vs FISH | Posttranslational modification, protein turnover, chaperones |
| gene_284848 | 4,471176158 | 5,407337916 | 1,872276902 | 2,888108009 | 0,003875668 | 0,048917066 | The Moose: suction sampler vs FISH | Posttranslational modification, protein turnover, chaperones |
| gene_69734 | 9,714761763 | 5,416504813 | 1,422330835 | 3,808189123 | 0,000139988 | 0,006212346 | The Moose: suction sampler vs FISH | Posttranslational modification, protein turnover, chaperones |
| gene_1580819 | 3,603691134 | 5,439266485 | 1,805374331 | 3,01281922 | 0,002588331 | 0,038598972 | The Moose: suction sampler vs FISH | Posttranslational modification, protein turnover, chaperones |
| gene_1756661 | 9,223104661 | 5,439923749 | 1,391303066 | 3,909948796 | 9,23E-05 | 0,004757983 | The Moose: suction sampler vs FISH | Posttranslational modification, protein turnover, chaperones |
| gene_1764740 | 4,958066621 | 5,479148299 | 1,691844181 | 3,238565561 | 0,001201324 | 0,024359814 | The Moose: suction sampler vs FISH | Posttranslational modification, protein turnover, chaperones |
| gene_1737129 | 11,32424545 | 5,480380233 | 1,472178121 | 3,722633935 | 0,000197155 | 0,007730949 | The Moose: suction sampler vs FISH | Posttranslational modification, protein turnover, chaperones |
| gene_1915948 | 7,131168298 | 5,487127611 | 1,599014469 | 3,431568456 | 0,000600102 | 0,015635485 | The Moose: suction sampler vs FISH | Posttranslational modification, protein turnover, chaperones |
| gene_1773705 | 4,808054144 | 5,489277997 | 1,86154732 | 2,948771669 | 0,003190396 | 0,043691708 | The Moose: suction sampler vs FISH | Posttranslational modification, protein turnover, chaperones |
| gene_211759 | 8,400161049 | 5,499445408 | 1,799926162 | 3,055372784 | 0,002247809 | 0,035414478 | The Moose: suction sampler vs FISH | Posttranslational modification, protein turnover, chaperones |
| gene_720862 | 21,19997212 | 5,505783765 | 1,145308651 | 4,80724891 | 1,53E-06 | 0,000295754 | The Moose: suction sampler vs FISH | Posttranslational modification, protein turnover, chaperones |
| gene_1908039 | 14,6641717 | 5,564272278 | 1,35894478 | 4,094553627 | 4,23E-05 | 0,002811398 | The Moose: suction sampler vs FISH | Posttranslational modification, protein turnover, chaperones |
| gene_1768840 | 9,065237208 | 5,689862526 | 1,567292999 | 3,63037577 | 0,000283009 | 0,009813781 | The Moose: suction sampler vs FISH | Posttranslational modification, protein turnover, chaperones |
| gene_1841288 | 336,7630312 | 5,719063201 | 0,89535913 | 6,387451704 | 1,69E-10 | 4,96E-07 | The Moose: suction sampler vs FISH | Posttranslational modification, protein turnover, chaperones |
| gene_945835 | 5,345992758 | 5,738526981 | 1,728633921 | 3,319688982 | 0,000901178 | 0,02049231 | The Moose: suction sampler vs FISH | Posttranslational modification, protein turnover, chaperones |
| gene_288298 | 5,345374666 | 5,742069285 | 1,947333173 | 2,948683545 | 0,003191306 | 0,04369602 | The Moose: suction sampler vs FISH | Posttranslational modification, protein turnover, chaperones |
| gene_1594326 | 3,504600299 | 5,752334448 | 1,851389832 | 3,107035778 | 0,001889735 | 0,032090545 | The Moose: suction sampler vs FISH | Posttranslational modification, protein turnover, chaperones |
| gene_568167 | 7,197736566 | 5,766905985 | 1,771061295 | 3,25618656 | 0,001129195 | 0,023472208 | The Moose: suction sampler vs FISH | Posttranslational modification, protein turnover, chaperones |
| gene_1030166 | 6,619174272 | 5,795814633 | 1,659595712 | 3,49230514 | 0,000478871 | 0,013571764 | The Moose: suction sampler vs FISH | Posttranslational modification, protein turnover, chaperones |
| gene_1775553 | 57,29044581 | 5,8094845 | 1,289788573 | 4,504214585 | 6,66E-06 | 0,00081547 | The Moose: suction sampler vs FISH | Posttranslational modification, protein turnover, chaperones |
| gene_201088 | 31,6579623 | 5,810358918 | 1,235577492 | 4,70254513 | 2,57E-06 | 0,000414072 | The Moose: suction sampler vs FISH | Posttranslational modification, protein turnover, chaperones |
| gene_819333 | 6,404337705 | 5,81070431 | 1,563737541 | 3,715907663 | 0,000202476 | 0,007880665 | The Moose: suction sampler vs FISH | Posttranslational modification, protein turnover, chaperones |
| gene_1268988 | 4,611625739 | 5,885813757 | 1,965345074 | 2,994799149 | 0,002746256 | 0,039845057 | The Moose: suction sampler vs FISH | Posttranslational modification, protein turnover, chaperones |
| gene_289810 | 270,1855249 | 5,903704013 | 0,977831031 | 6,03755028 | 1,56E-09 | 2,39E-06 | The Moose: suction sampler vs FISH | Posttranslational modification, protein turnover, chaperones |
| gene_1807594 | 4,843968441 | 5,945676663 | 1,893503554 | 3,140039876 | 0,001689248 | 0,030018594 | The Moose: suction sampler vs FISH | Posttranslational modification, protein turnover, chaperones |
| gene_850648 | 4,823885262 | 5,989402693 | 1,765220619 | 3,393005175 | 0,000691303 | 0,017165926 | The Moose: suction sampler vs FISH | Posttranslational modification, protein turnover, chaperones |
| gene_1264877 | 26,40947314 | 6,014507511 | 1,051656439 | 5,719080193 | 1,07E-08 | 9,48E-06 | The Moose: suction sampler vs FISH | Posttranslational modification, protein turnover, chaperones |
| gene_1288496 | 12,96628948 | 6,034371484 | 1,491079848 | 4,046980777 | 5,19E-05 | 0,003223638 | The Moose: suction sampler vs FISH | Posttranslational modification, protein turnover, chaperones |
| gene_1353677 | 3,233698801 | 6,041701289 | 1,904960874 | 3,171561878 | 0,001516216 | 0,02807828 | The Moose: suction sampler vs FISH | Posttranslational modification, protein turnover, chaperones |
| gene_1816176 | 10,19742366 | 6,159783567 | 1,478419285 | 4,166465921 | 3,09E-05 | 0,002260777 | The Moose: suction sampler vs FISH | Posttranslational modification, protein turnover, chaperones |
| gene_984427 | 4,129929331 | 6,166137209 | 1,857025084 | 3,320438298 | 0,000898762 | 0,020474439 | The Moose: suction sampler vs FISH | Posttranslational modification, protein turnover, chaperones |
| gene_1071959 | 9,048592031 | 6,207070925 | 1,566637311 | 3,962034404 | 7,43E-05 | 0,004121578 | The Moose: suction sampler vs FISH | Posttranslational modification, protein turnover, chaperones |
| gene_1323082 | 4,631826927 | 6,215701727 | 1,940840604 | 3,20258228 | 0,001362014 | 0,026236538 | The Moose: suction sampler vs FISH | Posttranslational modification, protein turnover, chaperones |
| gene_266581 | 4,316263823 | 6,246731194 | 1,799548047 | 3,471277805 | 0,000517988 | 0,014259226 | The Moose: suction sampler vs FISH | Posttranslational modification, protein turnover, chaperones |
| gene_950932 | 3,231705301 | 6,302072899 | 1,93951215 | 3,249308285 | 0,00115686 | 0,023833269 | The Moose: suction sampler vs FISH | Posttranslational modification, protein turnover, chaperones |
| gene_1572590 | 3,618720732 | 6,406846357 | 2,089675905 | 3,065952161 | 0,00216978 | 0,034797766 | The Moose: suction sampler vs FISH | Posttranslational modification, protein turnover, chaperones |
| gene_1889426 | 4,357394911 | 6,458782213 | 1,93946266 | 3,330191577 | 0,000867863 | 0,020018895 | The Moose: suction sampler vs FISH | Posttranslational modification, protein turnover, chaperones |
| gene_983488 | 15,72487081 | 6,478298752 | 1,242738678 | 5,212921161 | 1,86E-07 | 6,83E-05 | The Moose: suction sampler vs FISH | Posttranslational modification, protein turnover, chaperones |
| gene_588047 | 4,240604144 | 6,526329654 | 2,023796259 | 3,224795789 | 0,001260626 | 0,025077639 | The Moose: suction sampler vs FISH | Posttranslational modification, protein turnover, chaperones |
| gene_1589607 | 3,695953095 | 6,528691524 | 2,088325675 | 3,126280351 | 0,001770327 | 0,030818124 | The Moose: suction sampler vs FISH | Posttranslational modification, protein turnover, chaperones |
| gene_1475258 | 7,92597532 | 6,539656681 | 1,814248051 | 3,604610008 | 0,000312622 | 0,010465143 | The Moose: suction sampler vs FISH | Posttranslational modification, protein turnover, chaperones |
| gene_1496146 | 3,114320758 | 6,539771651 | 2,178268561 | 3,002279778 | 0,002679658 | 0,039306464 | The Moose: suction sampler vs FISH | Posttranslational modification, protein turnover, chaperones |
| gene_473459 | 6,337018802 | 6,559374363 | 1,955517063 | 3,354291551 | 0,000795685 | 0,018860863 | The Moose: suction sampler vs FISH | Posttranslational modification, protein turnover, chaperones |
| gene_1073633 | 4,219229216 | 6,56149464 | 1,954049731 | 3,357895418 | 0,000785383 | 0,018701511 | The Moose: suction sampler vs FISH | Posttranslational modification, protein turnover, chaperones |
| gene_1091622 | 4,512968708 | 6,590383862 | 2,035707571 | 3,237392225 | 0,001206275 | 0,024406298 | The Moose: suction sampler vs FISH | Posttranslational modification, protein turnover, chaperones |
| gene_1376944 | 6,10098404 | 6,606601296 | 1,689331089 | 3,910779442 | 9,20E-05 | 0,004744978 | The Moose: suction sampler vs FISH | Posttranslational modification, protein turnover, chaperones |
| gene_94546 | 4,548913555 | 6,626827065 | 1,810893707 | 3,659423544 | 0,000252783 | 0,00909469 | The Moose: suction sampler vs FISH | Posttranslational modification, protein turnover, chaperones |
| gene_403808 | 32,113819 | 6,667853894 | 1,35734351 | 4,9124292 | 9,00E-07 | 0,000207759 | The Moose: suction sampler vs FISH | Posttranslational modification, protein turnover, chaperones |
| gene_1776564 | 6,339059899 | 6,706462751 | 1,786205997 | 3,754585285 | 0,000173629 | 0,007124112 | The Moose: suction sampler vs FISH | Posttranslational modification, protein turnover, chaperones |
| gene_533059 | 9,376329501 | 6,722737661 | 1,933641536 | 3,476723858 | 0,00050758 | 0,014088798 | The Moose: suction sampler vs FISH | Posttranslational modification, protein turnover, chaperones |
| gene_715584 | 8,387933418 | 6,724723709 | 1,692684893 | 3,972814868 | 7,10E-05 | 0,003979153 | The Moose: suction sampler vs FISH | Posttranslational modification, protein turnover, chaperones |
| gene_986487 | 4,170288855 | 6,742343551 | 1,923178257 | 3,505833911 | 0,000455179 | 0,01317723 | The Moose: suction sampler vs FISH | Posttranslational modification, protein turnover, chaperones |
| gene_846698 | 4,815521264 | 6,780025718 | 1,924878933 | 3,522312806 | 0,000427799 | 0,012694431 | The Moose: suction sampler vs FISH | Posttranslational modification, protein turnover, chaperones |
| gene_839508 | 19,10807693 | 6,896631918 | 1,426961522 | 4,833088917 | 1,34E-06 | 0,000271629 | The Moose: suction sampler vs FISH | Posttranslational modification, protein turnover, chaperones |
| gene_507223 | 10,40799315 | 6,979172533 | 1,636146623 | 4,265615584 | 1,99E-05 | 0,001652527 | The Moose: suction sampler vs FISH | Posttranslational modification, protein turnover, chaperones |
| gene_866573 | 4,68586711 | 6,983143599 | 1,873875273 | 3,726578658 | 0,000194097 | 0,007660085 | The Moose: suction sampler vs FISH | Posttranslational modification, protein turnover, chaperones |
| gene_1767578 | 3,260078729 | 7,007588232 | 2,289478754 | 3,060778886 | 0,002207621 | 0,035148033 | The Moose: suction sampler vs FISH | Posttranslational modification, protein turnover, chaperones |
| gene_1690258 | 8,782270035 | 7,025049533 | 1,744233422 | 4,027585668 | 5,64E-05 | 0,003412046 | The Moose: suction sampler vs FISH | Posttranslational modification, protein turnover, chaperones |
| gene_853991 | 25,35225372 | 7,026290764 | 1,500735255 | 4,681898915 | 2,84E-06 | 0,000437636 | The Moose: suction sampler vs FISH | Posttranslational modification, protein turnover, chaperones |
| gene_1404348 | 10,36608945 | 7,05448644 | 1,808576528 | 3,900573921 | 9,60E-05 | 0,004867502 | The Moose: suction sampler vs FISH | Posttranslational modification, protein turnover, chaperones |

| | | | | | | | | |
|---|---|---|---|---|---|---|---|---|
| gene_1710771 | 7,315954654 | 7,100612431 | 1,848707157 | 3,840853001 | 0,000122608 | 0,005735611 | The Moose: suction sampler vs FISH | Posttranslational modification, protein turnover, chaperones |
| gene_1316224 | 6,773681446 | 7,177625604 | 1,689788806 | 4,247646558 | 2,16E-05 | 0,00175705 | The Moose: suction sampler vs FISH | Posttranslational modification, protein turnover, chaperones |
| gene_1397582 | 4,004107345 | 7,284605067 | 2,164208977 | 3,365943467 | 0,000762823 | 0,018363017 | The Moose: suction sampler vs FISH | Posttranslational modification, protein turnover, chaperones |
| gene_778185 | 7,774996322 | 7,337672114 | 1,752433857 | 4,187132133 | 2,83E-05 | 0,002139001 | The Moose: suction sampler vs FISH | Posttranslational modification, protein turnover, chaperones |
| gene_1439369 | 7,444643307 | 7,361756741 | 1,712927363 | 4,297763525 | 1,73E-05 | 0,001515726 | The Moose: suction sampler vs FISH | Posttranslational modification, protein turnover, chaperones |
| gene_943212 | 6,985878062 | 7,540491651 | 1,796708923 | 4,196835422 | 2,71E-05 | 0,002077262 | The Moose: suction sampler vs FISH | Posttranslational modification, protein turnover, chaperones |
| gene_647334 | 17,68957489 | 7,545857067 | 1,806044708 | 4,178112 | 2,94E-05 | 0,002186829 | The Moose: suction sampler vs FISH | Posttranslational modification, protein turnover, chaperones |
| gene_1161082 | 7,433547191 | 7,548744043 | 1,718310072 | 4,393120988 | 1,12E-05 | 0,001146141 | The Moose: suction sampler vs FISH | Posttranslational modification, protein turnover, chaperones |
| gene_445921 | 8,962655621 | 8,197456473 | 1,996744031 | 4,105411784 | 4,04E-05 | 0,002724454 | The Moose: suction sampler vs FISH | Posttranslational modification, protein turnover, chaperones |
| gene_1468661 | 11,30888407 | 8,712559922 | 1,698260275 | 5,130285415 | 2,89E-07 | 9,38E-05 | The Moose: suction sampler vs FISH | Posttranslational modification, protein turnover, chaperones |
| gene_452931 | 4,269953831 | -6,000276139 | 2,05168261 | -2,924563531 | 0,003449397 | 0,045700744 | The Moose: suction sampler vs FISH | Replication, recombination and repair |
| gene_209419 | 60,94128571 | -3,464527261 | 1,032021054 | -3,357031572 | 0,000787841 | 0,018738016 | The Moose: suction sampler vs FISH | Replication, recombination and repair |
| gene_77873 | 25,90125845 | -3,380702154 | 0,886902579 | -3,811807785 | 0,000137954 | 0,006155554 | The Moose: suction sampler vs FISH | Replication, recombination and repair |
| gene_72262 | 49,11798852 | -2,979925042 | 0,667932285 | -4,461417886 | 8,14E-06 | 0,000931438 | The Moose: suction sampler vs FISH | Replication, recombination and repair |
| gene_921319 | 66,42635243 | -2,968365295 | 0,766267438 | -3,873798032 | 0,000107152 | 0,005218697 | The Moose: suction sampler vs FISH | Replication, recombination and repair |
| gene_1090382 | 20,15408192 | -2,797284591 | 0,938197094 | -2,981553248 | 0,002867902 | 0,041011106 | The Moose: suction sampler vs FISH | Replication, recombination and repair |
| gene_468661 | 75,15368436 | -2,711147082 | 0,766982998 | -3,534820314 | 0,000408053 | 0,012328021 | The Moose: suction sampler vs FISH | Replication, recombination and repair |
| gene_1184071 | 25,93190294 | -2,676248685 | 0,847791447 | -3,156729988 | 0,00159549 | 0,028954972 | The Moose: suction sampler vs FISH | Replication, recombination and repair |
| gene_1161020 | 108,9372146 | -2,593229624 | 0,640628663 | -4,047945046 | 5,17E-05 | 0,003223638 | The Moose: suction sampler vs FISH | Replication, recombination and repair |
| gene_220290 | 50,26902052 | -2,507283612 | 0,760541255 | -3,296709541 | 0,000978246 | 0,021434155 | The Moose: suction sampler vs FISH | Replication, recombination and repair |
| gene_637233 | 22,15534446 | -2,396292412 | 0,803700178 | -2,981575066 | 0,002867697 | 0,041011106 | The Moose: suction sampler vs FISH | Replication, recombination and repair |
| gene_1529674 | 48,20516092 | -2,233216714 | 0,640528546 | -3,486521762 | 0,000489346 | 0,013764835 | The Moose: suction sampler vs FISH | Replication, recombination and repair |
| gene_237310 | 233,9098163 | -2,169116253 | 0,724191904 | -2,99522301 | 0,002742443 | 0,039821806 | The Moose: suction sampler vs FISH | Replication, recombination and repair |
| gene_97132 | 261,6086575 | -2,163401844 | 0,601814998 | -3,594795493 | 0,000324646 | 0,010695092 | The Moose: suction sampler vs FISH | Replication, recombination and repair |
| gene_409287 | 251,8796767 | -2,157564514 | 0,653970436 | -3,299177446 | 0,000969686 | 0,021322929 | The Moose: suction sampler vs FISH | Replication, recombination and repair |
| gene_1252385 | 106,568795 | -2,14470305 | 0,649772264 | -3,300699596 | 0,000964441 | 0,021258511 | The Moose: suction sampler vs FISH | Replication, recombination and repair |
| gene_478787 | 42,59868503 | 2,095635503 | 0,723352309 | 2,897115935 | 0,003766106 | 0,048146496 | The Moose: suction sampler vs FISH | Replication, recombination and repair |
| gene_273587 | 32,40073631 | 2,613626542 | 0,884275608 | 2,95566961 | 0,003119911 | 0,043185425 | The Moose: suction sampler vs FISH | Replication, recombination and repair |
| gene_198022 | 27,28008563 | 2,623558485 | 0,863015899 | 3,03998859 | 0,002365871 | 0,03648239 | The Moose: suction sampler vs FISH | Replication, recombination and repair |
| gene_1945392 | 141,5107448 | 2,632785168 | 0,755448901 | 3,485060558 | 0,000492026 | 0,013823223 | The Moose: suction sampler vs FISH | Replication, recombination and repair |
| gene_614040 | 45,58581373 | 2,637281111 | 0,8294657 | 3,179493872 | 0,001475325 | 0,027618763 | The Moose: suction sampler vs FISH | Replication, recombination and repair |
| gene_216507 | 33,94597713 | 2,6475879 | 0,911295416 | 2,905301456 | 0,003668995 | 0,047425091 | The Moose: suction sampler vs FISH | Replication, recombination and repair |
| gene_776543 | 113,2564132 | 2,742773487 | 0,739699042 | 3,707958685 | 0,000208937 | 0,008046858 | The Moose: suction sampler vs FISH | Replication, recombination and repair |
| gene_1298769 | 29,56011567 | 2,778200779 | 0,862425806 | 3,221379464 | 0,001275751 | 0,02528218 | The Moose: suction sampler vs FISH | Replication, recombination and repair |
| gene_636403 | 164,7250764 | 2,816673079 | 0,844976426 | 3,33343392 | 0,00085781 | 0,019873068 | The Moose: suction sampler vs FISH | Replication, recombination and repair |
| gene_1059860 | 23,73442575 | 2,842120488 | 0,879517575 | 3,231453888 | 0,001231622 | 0,024681166 | The Moose: suction sampler vs FISH | Replication, recombination and repair |
| gene_776542 | 87,19679125 | 2,844064731 | 0,771993848 | 3,684051031 | 0,000229556 | 0,008553911 | The Moose: suction sampler vs FISH | Replication, recombination and repair |
| gene_1190081 | 54,56708148 | 2,894317882 | 0,782680483 | 3,697955866 | 0,000217343 | 0,008227995 | The Moose: suction sampler vs FISH | Replication, recombination and repair |
| gene_1245875 | 26,18230999 | 2,90580464 | 0,958867815 | 3,030453827 | 0,002441865 | 0,037177195 | The Moose: suction sampler vs FISH | Replication, recombination and repair |
| gene_1190077 | 69,70345531 | 2,921886706 | 0,767591738 | 3,806563516 | 0,000140911 | 0,006234468 | The Moose: suction sampler vs FISH | Replication, recombination and repair |
| gene_1537066 | 90,84725872 | 3,005102837 | 0,771341843 | 3,895941683 | 9,78E-05 | 0,004947822 | The Moose: suction sampler vs FISH | Replication, recombination and repair |
| gene_1884896 | 163,5629263 | 3,017010045 | 0,824568835 | 3,658894101 | 0,000253306 | 0,009097339 | The Moose: suction sampler vs FISH | Replication, recombination and repair |
| gene_1085133 | 21,07719087 | 3,08626322 | 0,947028118 | 3,25889291 | 0,001118479 | 0,023337123 | The Moose: suction sampler vs FISH | Replication, recombination and repair |
| gene_1477869 | 55,58342985 | 3,120701176 | 0,733214461 | 4,256191525 | 2,08E-05 | 0,001712113 | The Moose: suction sampler vs FISH | Replication, recombination and repair |
| gene_732839 | 11,88367413 | 3,209312268 | 1,096171248 | 2,927747169 | 0,003414276 | 0,045397242 | The Moose: suction sampler vs FISH | Replication, recombination and repair |
| gene_931758 | 70,5310671 | 3,211022812 | 0,839800922 | 3,82355238 | 0,000131543 | 0,005963296 | The Moose: suction sampler vs FISH | Replication, recombination and repair |
| gene_1190076 | 21,2403819 | 3,231978334 | 0,857026545 | 3,771153125 | 0,000162495 | 0,006847895 | The Moose: suction sampler vs FISH | Replication, recombination and repair |
| gene_776539 | 63,99819718 | 3,2357537 | 0,949390101 | 3,408244618 | 0,000653823 | 0,016570047 | The Moose: suction sampler vs FISH | Replication, recombination and repair |
| gene_221459 | 20,35789001 | 3,301259894 | 0,960505833 | 3,437001403 | 0,000588193 | 0,015458922 | The Moose: suction sampler vs FISH | Replication, recombination and repair |
| gene_130849 | 24,2876451 | 3,313623743 | 0,957189126 | 3,461827608 | 0,000536521 | 0,014579957 | The Moose: suction sampler vs FISH | Replication, recombination and repair |
| gene_478025 | 12,27738738 | 3,353429542 | 1,117143738 | 3,001788783 | 0,002683983 | 0,039338159 | The Moose: suction sampler vs FISH | Replication, recombination and repair |
| gene_478237 | 11,76882244 | 3,359749369 | 1,110214688 | 3,026215924 | 0,002476354 | 0,037569885 | The Moose: suction sampler vs FISH | Replication, recombination and repair |
| gene_1335822 | 11,86595269 | 3,663167721 | 1,17905828 | 3,10685891 | 0,001890866 | 0,032094908 | The Moose: suction sampler vs FISH | Replication, recombination and repair |
| gene_1412247 | 11,93849027 | 3,695038325 | 1,105856374 | 3,341336554 | 0,000833761 | 0,019483159 | The Moose: suction sampler vs FISH | Replication, recombination and repair |
| gene_1551319 | 10,76356722 | 3,724841153 | 1,219257825 | 3,05500697 | 0,002250553 | 0,035432339 | The Moose: suction sampler vs FISH | Replication, recombination and repair |
| gene_1471052 | 23,26575308 | 3,819925887 | 1,013002213 | 3,770895895 | 0,000162663 | 0,006847895 | The Moose: suction sampler vs FISH | Replication, recombination and repair |
| gene_1335825 | 10,59844369 | 3,847471375 | 1,158644318 | 3,320666502 | 0,000898028 | 0,020474439 | The Moose: suction sampler vs FISH | Replication, recombination and repair |
| gene_1125122 | 24,90656698 | 3,963148954 | 1,04344969 | 3,798121743 | 0,000145797 | 0,006362888 | The Moose: suction sampler vs FISH | Replication, recombination and repair |
| gene_674367 | 92,81307555 | 4,03134343 | 0,928428402 | 4,34211558 | 1,41E-05 | 0,001328764 | The Moose: suction sampler vs FISH | Replication, recombination and repair |
| gene_1799341 | 6,928770139 | 4,057846601 | 1,337255949 | 3,034457693 | 0,002409686 | 0,036909145 | The Moose: suction sampler vs FISH | Replication, recombination and repair |
| gene_98368 | 39,54673115 | 4,075088691 | 0,873537734 | 4,665040251 | 3,09E-06 | 0,000463434 | The Moose: suction sampler vs FISH | Replication, recombination and repair |
| gene_555578 | 23,5301404 | 4,08853954 | 1,035360951 | 3,948902588 | 7,85E-05 | 0,004252351 | The Moose: suction sampler vs FISH | Replication, recombination and repair |
| gene_1335827 | 15,05540929 | 4,115286317 | 1,09472279 | 3,759204024 | 0,000170455 | 0,007047063 | The Moose: suction sampler vs FISH | Replication, recombination and repair |
| gene_680710 | 11,08436022 | 4,174965419 | 1,184366846 | 3,525061035 | 0,000423385 | 0,012611664 | The Moose: suction sampler vs FISH | Replication, recombination and repair |
| gene_1270420 | 9,07083364 | 4,228246373 | 1,284249621 | 3,292386701 | 0,000993409 | 0,021630873 | The Moose: suction sampler vs FISH | Replication, recombination and repair |

| | | | | | | | | |
|---|---|---|---|---|---|---|---|---|
| gene_1767513 | 9,422551507 | 4,485689464 | 1,337788551 | 3,353063127 | 0,000799225 | 0,018928075 | The Moose: suction sampler vs FISH | Replication, recombination and repair |
| gene_1317749 | 11,04380656 | 4,546766402 | 1,358060639 | 3,347984818 | 0,000814015 | 0,019155819 | The Moose: suction sampler vs FISH | Replication, recombination and repair |
| gene_1182555 | 6,488747205 | 4,57832914 | 1,557230516 | 2,940045865 | 0,003281637 | 0,044378532 | The Moose: suction sampler vs FISH | Replication, recombination and repair |
| gene_208819 | 24,69198083 | 4,756953249 | 1,000222538 | 4,755894881 | 1,98E-06 | 0,000347141 | The Moose: suction sampler vs FISH | Replication, recombination and repair |
| gene_1909511 | 7,790884197 | 4,824724821 | 1,452761634 | 3,321071198 | 0,000896727 | 0,020466154 | The Moose: suction sampler vs FISH | Replication, recombination and repair |
| gene_1522966 | 19,43155781 | 4,877457531 | 1,198711034 | 4,068918525 | 4,72E-05 | 0,003057907 | The Moose: suction sampler vs FISH | Replication, recombination and repair |
| gene_492751 | 7,518917728 | 4,916587917 | 1,431211124 | 3,435263906 | 0,000591977 | 0,015512882 | The Moose: suction sampler vs FISH | Replication, recombination and repair |
| gene_581798 | 12,85876602 | 4,96678213 | 1,354355212 | 3,667266968 | 0,000245157 | 0,00893132 | The Moose: suction sampler vs FISH | Replication, recombination and repair |
| gene_1634533 | 8,178724909 | 5,046127293 | 1,467314782 | 3,439021643 | 0,000583821 | 0,015396302 | The Moose: suction sampler vs FISH | Replication, recombination and repair |
| gene_753461 | 5,491117193 | 5,105181143 | 1,675591727 | 3,046792999 | 0,002312969 | 0,036021211 | The Moose: suction sampler vs FISH | Replication, recombination and repair |
| gene_1936866 | 5,725907434 | 5,258391563 | 1,662267251 | 3,163385166 | 0,001559459 | 0,028562204 | The Moose: suction sampler vs FISH | Replication, recombination and repair |
| gene_1811459 | 12,42328061 | 5,58599636 | 1,47452135 | 3,788345525 | 0,000151654 | 0,006502169 | The Moose: suction sampler vs FISH | Replication, recombination and repair |
| gene_1164781 | 3,323332447 | 5,795599724 | 1,982514183 | 2,923358518 | 0,003462776 | 0,045782817 | The Moose: suction sampler vs FISH | Replication, recombination and repair |
| gene_598797 | 6,130145366 | 5,83981584 | 1,798848778 | 3,246418437 | 0,001168669 | 0,023920816 | The Moose: suction sampler vs FISH | Replication, recombination and repair |
| gene_1920030 | 3,952945557 | 5,951950291 | 2,015107528 | 2,953663865 | 0,003140258 | 0,043306684 | The Moose: suction sampler vs FISH | Replication, recombination and repair |
| gene_225738 | 4,897806434 | 6,053062001 | 1,968600371 | 3,074804866 | 0,002106403 | 0,034249454 | The Moose: suction sampler vs FISH | Replication, recombination and repair |
| gene_1791938 | 4,801155232 | 6,058363393 | 1,946760636 | 3,112022752 | 0,001858102 | 0,031751858 | The Moose: suction sampler vs FISH | Replication, recombination and repair |
| gene_1804451 | 6,153641117 | 6,179835576 | 1,879092301 | 3,288734445 | 0,001006389 | 0,021803617 | The Moose: suction sampler vs FISH | Replication, recombination and repair |
| gene_1572153 | 7,649860751 | 6,215118528 | 1,741201668 | 3,569442093 | 0,000357742 | 0,011357993 | The Moose: suction sampler vs FISH | Replication, recombination and repair |
| gene_503183 | 7,40608635 | 6,231014942 | 2,082283905 | 2,992394519 | 0,002767983 | 0,040065927 | The Moose: suction sampler vs FISH | Replication, recombination and repair |
| gene_1148899 | 3,505135424 | 6,269797149 | 2,068775473 | 3,030680338 | 0,002440034 | 0,03717118 | The Moose: suction sampler vs FISH | Replication, recombination and repair |
| gene_1548414 | 3,513018729 | 6,322776565 | 1,952177817 | 3,238832299 | 0,001200201 | 0,024350491 | The Moose: suction sampler vs FISH | Replication, recombination and repair |
| gene_1363389 | 5,436046011 | 6,441535729 | 2,017250655 | 3,193225252 | 0,001406932 | 0,026776907 | The Moose: suction sampler vs FISH | Replication, recombination and repair |
| gene_186496 | 7,188788409 | 6,598115892 | 1,679265036 | 3,929168863 | 8,52E-05 | 0,004497449 | The Moose: suction sampler vs FISH | Replication, recombination and repair |
| gene_606282 | 5,053971203 | 6,653732073 | 1,924209852 | 3,457903548 | 0,000544396 | 0,014710516 | The Moose: suction sampler vs FISH | Replication, recombination and repair |
| gene_1519189 | 3,845181285 | 6,6858159 | 1,978689787 | 3,378910602 | 0,000727737 | 0,017792484 | The Moose: suction sampler vs FISH | Replication, recombination and repair |
| gene_1145718 | 3,397596525 | 6,991011669 | 2,144384058 | 3,260149059 | 0,001113537 | 0,023278867 | The Moose: suction sampler vs FISH | Replication, recombination and repair |
| gene_1815767 | 4,784558001 | 7,01084319 | 2,043049342 | 3,431558429 | 0,000600124 | 0,015635485 | The Moose: suction sampler vs FISH | Replication, recombination and repair |
| gene_190551 | 6,03343295 | 7,026992894 | 1,946892487 | 3,609337928 | 0,000306979 | 0,01032331 | The Moose: suction sampler vs FISH | Replication, recombination and repair |
| gene_1550518 | 7,132720169 | 7,265297135 | 1,691669575 | 4,294749544 | 1,75E-05 | 0,001529148 | The Moose: suction sampler vs FISH | Replication, recombination and repair |
| gene_1200857 | 8,262889812 | 7,508873798 | 1,711556667 | 4,387160496 | 1,15E-05 | 0,001165739 | The Moose: suction sampler vs FISH | Replication, recombination and repair |
| gene_408823 | 9,139606453 | 7,578837153 | 1,615526901 | 4,691247882 | 2,72E-06 | 0,000427245 | The Moose: suction sampler vs FISH | Replication, recombination and repair |
| gene_1603083 | 72,89187957 | 2,274085343 | 0,646559393 | 3,517210278 | 0,000436108 | 0,012853115 | The Moose: suction sampler vs FISH | RNA processing and modification |
| gene_1252078 | 67,97743005 | 2,337505491 | 0,736606764 | 3,17334242 | 0,001506947 | 0,028005493 | The Moose: suction sampler vs FISH | RNA processing and modification |
| gene_1241592 | 25,86318638 | 2,488443171 | 0,848995959 | 2,931042421 | 0,003378266 | 0,045095743 | The Moose: suction sampler vs FISH | RNA processing and modification |
| gene_1283803 | 67,7929269 | 2,638797325 | 0,772934342 | 3,413999329 | 0,000640167 | 0,016349683 | The Moose: suction sampler vs FISH | RNA processing and modification |
| gene_628534 | 10,98928639 | 2,967445009 | 1,030375482 | 2,879964694 | 0,003977197 | 0,049618692 | The Moose: suction sampler vs FISH | RNA processing and modification |
| gene_1532513 | 27,57502861 | 3,022737655 | 1,022020916 | 2,95760841 | 0,003100357 | 0,043012037 | The Moose: suction sampler vs FISH | RNA processing and modification |
| gene_808625 | 106,0231844 | 3,068080689 | 0,793494456 | 3,866543321 | 0,000110389 | 0,005330381 | The Moose: suction sampler vs FISH | RNA processing and modification |
| gene_205909 | 25,71247331 | 3,082611373 | 1,017173718 | 3,030565299 | 0,002440964 | 0,03717718 | The Moose: suction sampler vs FISH | RNA processing and modification |
| gene_88591 | 36,95010964 | 3,178790519 | 0,970392833 | 3,275776995 | 0,001053717 | 0,022529876 | The Moose: suction sampler vs FISH | RNA processing and modification |
| gene_1190075 | 43,57483657 | 3,179117714 | 1,012279779 | 3,140552424 | 0,001686295 | 0,029973367 | The Moose: suction sampler vs FISH | RNA processing and modification |
| gene_814558 | 15,31829564 | 3,320253789 | 1,053631903 | 3,151246445 | 0,001625753 | 0,029323034 | The Moose: suction sampler vs FISH | RNA processing and modification |
| gene_1797092 | 37,22991425 | 3,386870368 | 1,008628704 | 3,357896075 | 0,000785381 | 0,018701511 | The Moose: suction sampler vs FISH | RNA processing and modification |
| gene_173937 | 32,88485563 | 3,439203416 | 0,929456646 | 3,700230056 | 0,000215404 | 0,008184353 | The Moose: suction sampler vs FISH | RNA processing and modification |
| gene_1101985 | 36,07321732 | 3,500264504 | 0,824856797 | 4,243481436 | 2,20E-05 | 0,00178014 | The Moose: suction sampler vs FISH | RNA processing and modification |
| gene_796077 | 297,2816398 | 3,505650327 | 0,819129192 | 4,279728229 | 1,87E-05 | 0,00158149 | The Moose: suction sampler vs FISH | RNA processing and modification |
| gene_1918624 | 15,57922227 | 3,564964223 | 1,02591509 | 3,474911576 | 0,000511022 | 0,014163017 | The Moose: suction sampler vs FISH | RNA processing and modification |
| gene_655356 | 39,39332326 | 3,573095065 | 0,936737425 | 3,814404091 | 0,000136512 | 0,006128435 | The Moose: suction sampler vs FISH | RNA processing and modification |
| gene_1614276 | 16,06209584 | 3,60286907 | 1,169024533 | 3,081944791 | 0,00205653 | 0,033767455 | The Moose: suction sampler vs FISH | RNA processing and modification |
| gene_1620279 | 27,95648927 | 3,633403734 | 0,878033657 | 4,138114417 | 3,50E-05 | 0,002454294 | The Moose: suction sampler vs FISH | RNA processing and modification |
| gene_344388 | 42,68679209 | 3,676792347 | 0,894619963 | 4,109893027 | 3,96E-05 | 0,002686093 | The Moose: suction sampler vs FISH | RNA processing and modification |
| gene_434350 | 62,45181209 | 3,767891902 | 0,714927198 | 5,270315511 | 1,36E-07 | 5,51E-05 | The Moose: suction sampler vs FISH | RNA processing and modification |
| gene_1764432 | 161,9139101 | 3,795037588 | 0,836059665 | 4,539194684 | 5,65E-06 | 0,000716304 | The Moose: suction sampler vs FISH | RNA processing and modification |
| gene_1573379 | 54,20926112 | 3,803029417 | 0,87707828 | 4,336020519 | 1,45E-05 | 0,001343732 | The Moose: suction sampler vs FISH | RNA processing and modification |
| gene_1899503 | 20,26143194 | 3,808929689 | 1,053008194 | 3,617189032 | 0,00029782 | 0,01015477 | The Moose: suction sampler vs FISH | RNA processing and modification |
| gene_397571 | 40,18159019 | 3,852933005 | 0,888259568 | 4,337620603 | 1,44E-05 | 0,001337845 | The Moose: suction sampler vs FISH | RNA processing and modification |
| gene_308095 | 53,3110255 | 3,856993565 | 0,939594609 | 4,104954977 | 4,04E-05 | 0,002727336 | The Moose: suction sampler vs FISH | RNA processing and modification |
| gene_1943079 | 61,10005454 | 3,859322657 | 0,789858125 | 4,886096042 | 1,03E-06 | 0,000226853 | The Moose: suction sampler vs FISH | RNA processing and modification |
| gene_23586 | 16,1522955 | 3,988137691 | 1,131486684 | 3,524688136 | 0,000423982 | 0,012618579 | The Moose: suction sampler vs FISH | RNA processing and modification |
| gene_666828 | 20,5787834 | 4,005866937 | 1,02678369 | 3,901373753 | 9,56E-05 | 0,004864833 | The Moose: suction sampler vs FISH | RNA processing and modification |
| gene_1759100 | 12,06610799 | 4,05081105 | 1,066574841 | 3,797962313 | 0,00014589 | 0,006362888 | The Moose: suction sampler vs FISH | RNA processing and modification |
| gene_1481069 | 8,062077093 | 4,0805701 | 1,330929682 | 3,06595469 | 0,002169762 | 0,034797766 | The Moose: suction sampler vs FISH | RNA processing and modification |
| gene_808936 | 26,43871591 | 4,082008672 | 1,056608805 | 3,863311239 | 0,00011186 | 0,005380211 | The Moose: suction sampler vs FISH | RNA processing and modification |
| gene_501624 | 17,19774151 | 4,241495088 | 1,219732678 | 3,477397272 | 0,000506307 | 0,014058873 | The Moose: suction sampler vs FISH | RNA processing and modification |

| | | | | | | | | |
|---|---|---|---|---|---|---|---|---|
| gene_490345 | 13,46011392 | 4,37475477 | 1,345404144 | 3,251628733 | 0,001147458 | 0,023706063 | The Moose: suction sampler vs FISH | RNA processing and modification |
| gene_16160 | 6,667760606 | 4,49530065 | 1,520500998 | 2,956460177 | 0,003111924 | 0,043115501 | The Moose: suction sampler vs FISH | RNA processing and modification |
| gene_1723027 | 26,0297478 | 4,58895031 | 1,073929936 | 4,27304441 | 1,93E-05 | 0,001612963 | The Moose: suction sampler vs FISH | RNA processing and modification |
| gene_732982 | 8,374656391 | 4,597132492 | 1,353112658 | 3,39744992 | 0,00068017 | 0,016997316 | The Moose: suction sampler vs FISH | RNA processing and modification |
| gene_472154 | 15,73161515 | 4,682973444 | 1,268157598 | 3,692737756 | 0,000221853 | 0,00835194 | The Moose: suction sampler vs FISH | RNA processing and modification |
| gene_256254 | 6,425334053 | 4,751268808 | 1,650868873 | 2,878041306 | 0,004001527 | 0,049803793 | The Moose: suction sampler vs FISH | RNA processing and modification |
| gene_1616931 | 7,833614686 | 4,833818205 | 1,441721435 | 3,352810112 | 0,000799956 | 0,018928075 | The Moose: suction sampler vs FISH | RNA processing and modification |
| gene_1863964 | 108,9982348 | 4,848273886 | 0,893086899 | 5,428669812 | 5,68E-08 | 3,28E-05 | The Moose: suction sampler vs FISH | RNA processing and modification |
| gene_819088 | 8,947001414 | 4,857297523 | 1,426837367 | 3,40424048 | 0,000663483 | 0,016737807 | The Moose: suction sampler vs FISH | RNA processing and modification |
| gene_523594 | 34,36598681 | 4,997489156 | 0,934419672 | 5,348227685 | 8,88E-08 | 4,50E-05 | The Moose: suction sampler vs FISH | RNA processing and modification |
| gene_1592078 | 20,48373243 | 5,025769945 | 1,088389328 | 4,617621484 | 3,88E-06 | 0,000552304 | The Moose: suction sampler vs FISH | RNA processing and modification |
| gene_1578979 | 8,007461608 | 5,217929323 | 1,625784678 | 3,209483639 | 0,001329736 | 0,025870505 | The Moose: suction sampler vs FISH | RNA processing and modification |
| gene_1335826 | 14,32010995 | 5,244817318 | 1,358285857 | 3,861350165 | 0,000112762 | 0,005405093 | The Moose: suction sampler vs FISH | RNA processing and modification |
| gene_1853221 | 4,021600915 | 5,435385967 | 1,778108457 | 3,056836013 | 0,002236866 | 0,035360455 | The Moose: suction sampler vs FISH | RNA processing and modification |
| gene_1886897 | 5,229729293 | 5,581655633 | 1,937715414 | 2,880534259 | 0,003970018 | 0,049588781 | The Moose: suction sampler vs FISH | RNA processing and modification |
| gene_1001999 | 3,258793031 | 5,670523925 | 1,879598613 | 3,016880246 | 0,002553907 | 0,038303387 | The Moose: suction sampler vs FISH | RNA processing and modification |
| gene_1835538 | 6,644458787 | 5,693189088 | 1,720351744 | 3,309316892 | 0,000935239 | 0,020887457 | The Moose: suction sampler vs FISH | RNA processing and modification |
| gene_958314 | 3,630482061 | 5,909779021 | 2,027461591 | 2,914866081 | 0,003558412 | 0,046527013 | The Moose: suction sampler vs FISH | RNA processing and modification |
| gene_1467799 | 4,20193145 | 6,414316257 | 1,944388929 | 3,298885404 | 0,000970695 | 0,021338736 | The Moose: suction sampler vs FISH | RNA processing and modification |
| gene_423568 | 3,22257861 | 6,431130642 | 2,078450558 | 3,094194671 | 0,001973479 | 0,032933924 | The Moose: suction sampler vs FISH | RNA processing and modification |
| gene_471037 | 3,878858209 | 6,564789615 | 2,103365009 | 3,121089105 | 0,001801835 | 0,031137829 | The Moose: suction sampler vs FISH | RNA processing and modification |
| gene_1478910 | 4,892060147 | 6,750545196 | 2,086822174 | 3,234844483 | 0,00121709 | 0,024517054 | The Moose: suction sampler vs FISH | RNA processing and modification |
| gene_1128713 | 7,021932885 | 6,909474881 | 1,779905621 | 3,881933288 | 0,000103629 | 0,005101251 | The Moose: suction sampler vs FISH | RNA processing and modification |
| gene_995495 | 11,08124864 | 8,078044855 | 1,722627357 | 4,689374532 | 2,74E-06 | 0,000428233 | The Moose: suction sampler vs FISH | RNA processing and modification |
| gene_832033 | 25,3792973 | -7,093838963 | 1,414513034 | -5,01503966 | 5,30E-07 | 0,000144231 | The Moose: suction sampler vs FISH | Secondary metabolites biosynthesis, transport and catabolism |
| gene_315635 | 1217,748824 | -2,74168654 | 0,949058229 | -2,888849656 | 0,003866539 | 0,048863897 | The Moose: suction sampler vs FISH | Secondary metabolites biosynthesis, transport and catabolism |
| gene_100143 | 557,9583262 | -2,616012415 | 0,759531747 | -3,444243675 | 0,000572659 | 0,015238755 | The Moose: suction sampler vs FISH | Secondary metabolites biosynthesis, transport and catabolism |
| gene_1565185 | 28,88893053 | -2,592763568 | 0,842618779 | -3,077030366 | 0,00209074 | 0,034092895 | The Moose: suction sampler vs FISH | Secondary metabolites biosynthesis, transport and catabolism |
| gene_142780 | 35,21123066 | -2,244430815 | 0,773176723 | -2,902869097 | 0,003697612 | 0,047685879 | The Moose: suction sampler vs FISH | Secondary metabolites biosynthesis, transport and catabolism |
| gene_108567 | 178,5925898 | -2,191357448 | 0,601047252 | -3,645898788 | 0,000266459 | 0,009438241 | The Moose: suction sampler vs FISH | Secondary metabolites biosynthesis, transport and catabolism |
| gene_1406758 | 306,3717638 | -2,038997938 | 0,68650743 | -2,970103235 | 0,002976997 | 0,041936634 | The Moose: suction sampler vs FISH | Secondary metabolites biosynthesis, transport and catabolism |
| gene_1254141 | 119,7920422 | 2,733773096 | 0,75207837 | 3,634957744 | 0,000278026 | 0,009695933 | The Moose: suction sampler vs FISH | Secondary metabolites biosynthesis, transport and catabolism |
| gene_183022 | 71,42139148 | 2,791980008 | 0,900734482 | 3,099670394 | 0,001937361 | 0,032567975 | The Moose: suction sampler vs FISH | Secondary metabolites biosynthesis, transport and catabolism |
| gene_634861 | 729,5621614 | 2,850680861 | 0,751038277 | 3,795653229 | 0,000147255 | 0,006389958 | The Moose: suction sampler vs FISH | Secondary metabolites biosynthesis, transport and catabolism |
| gene_1544382 | 54,43691309 | 3,125732832 | 0,812863894 | 3,845333583 | 0,000120389 | 0,0056498 | The Moose: suction sampler vs FISH | Secondary metabolites biosynthesis, transport and catabolism |
| gene_902903 | 28,29264443 | 3,13175044 | 0,961287614 | 3,257870376 | 0,001122517 | 0,023394789 | The Moose: suction sampler vs FISH | Secondary metabolites biosynthesis, transport and catabolism |
| gene_1516539 | 33,89346045 | 3,274422054 | 0,858330178 | 3,814874669 | 0,000136252 | 0,00612051 | The Moose: suction sampler vs FISH | Secondary metabolites biosynthesis, transport and catabolism |
| gene_1874519 | 23,8172207 | 3,416679786 | 0,942937713 | 3,623441652 | 0,000290709 | 0,009970526 | The Moose: suction sampler vs FISH | Secondary metabolites biosynthesis, transport and catabolism |
| gene_1325780 | 18,22918902 | 3,443974813 | 0,906514971 | 3,799137271 | 0,000145201 | 0,006359131 | The Moose: suction sampler vs FISH | Secondary metabolites biosynthesis, transport and catabolism |
| gene_696700 | 13,42704986 | 3,9865908 | 0,983228098 | 4,054594056 | 5,02E-05 | 0,003168826 | The Moose: suction sampler vs FISH | Secondary metabolites biosynthesis, transport and catabolism |
| gene_165440 | 28,21580233 | 4,270498637 | 1,070303475 | 3,989988574 | 6,61E-05 | 0,003773707 | The Moose: suction sampler vs FISH | Secondary metabolites biosynthesis, transport and catabolism |
| gene_267830 | 8,156801532 | 4,310217534 | 1,493954012 | 2,885107238 | 0,003912804 | 0,049132477 | The Moose: suction sampler vs FISH | Secondary metabolites biosynthesis, transport and catabolism |
| gene_469530 | 345,8591291 | 4,393307811 | 1,498168339 | 2,932452714 | 0,003362961 | 0,044989562 | The Moose: suction sampler vs FISH | Secondary metabolites biosynthesis, transport and catabolism |
| gene_1898487 | 5,625789694 | 4,948249542 | 1,615539855 | 3,062907749 | 0,002191976 | 0,03498255 | The Moose: suction sampler vs FISH | Secondary metabolites biosynthesis, transport and catabolism |
| gene_175026 | 7,289561829 | 5,079660123 | 1,602676765 | 3,169485099 | 0,001527093 | 0,0282074 | The Moose: suction sampler vs FISH | Secondary metabolites biosynthesis, transport and catabolism |
| gene_696699 | 109,5374352 | 5,301966582 | 0,750690508 | 7,062786226 | 1,63E-12 | 1,71E-08 | The Moose: suction sampler vs FISH | Secondary metabolites biosynthesis, transport and catabolism |
| gene_398103 | 4,396302333 | 5,514330319 | 1,778453033 | 3,100633088 | 0,001931074 | 0,032521839 | The Moose: suction sampler vs FISH | Secondary metabolites biosynthesis, transport and catabolism |
| gene_1829195 | 3,524980276 | 6,102927521 | 1,955973122 | 3,12014897 | 0,001807596 | 0,031208013 | The Moose: suction sampler vs FISH | Secondary metabolites biosynthesis, transport and catabolism |
| gene_149191 | 3,912978131 | 6,162206622 | 1,986402767 | 3,102193937 | 0,001920921 | 0,032380542 | The Moose: suction sampler vs FISH | Secondary metabolites biosynthesis, transport and catabolism |
| gene_1406868 | 10,90176652 | 6,407194455 | 1,451203383 | 4,415090628 | 1,01E-05 | 0,001071068 | The Moose: suction sampler vs FISH | Secondary metabolites biosynthesis, transport and catabolism |
| gene_1096903 | 3,71510388 | 6,706397995 | 2,126623614 | 3,153542523 | 0,001613017 | 0,029181652 | The Moose: suction sampler vs FISH | Secondary metabolites biosynthesis, transport and catabolism |
| gene_1319595 | 51,81352318 | 6,708197778 | 1,314211197 | 5,104352932 | 3,32E-07 | 0,000104323 | The Moose: suction sampler vs FISH | Secondary metabolites biosynthesis, transport and catabolism |
| gene_1157559 | 8,711396418 | 6,735148058 | 1,784960629 | 3,773275414 | 0,000161118 | 0,006804675 | The Moose: suction sampler vs FISH | Secondary metabolites biosynthesis, transport and catabolism |
| gene_1154454 | 4,627319767 | 6,766877298 | 1,933895333 | 3,499091799 | 0,000466846 | 0,013372367 | The Moose: suction sampler vs FISH | Secondary metabolites biosynthesis, transport and catabolism |
| gene_958642 | 3,382126324 | 6,881626612 | 2,103541139 | 3,271448552 | 0,001069981 | 0,022715418 | The Moose: suction sampler vs FISH | Secondary metabolites biosynthesis, transport and catabolism |
| gene_752635 | 6,036976561 | 7,331070168 | 1,901746167 | 3,854915179 | 0,00011577 | 0,005496257 | The Moose: suction sampler vs FISH | Secondary metabolites biosynthesis, transport and catabolism |
| gene_1945092 | 5,644058872 | -5,68939423 | 1,896855127 | -2,999382582 | 0,002705274 | 0,039498414 | The Moose: suction sampler vs FISH | Signal transduction mechanisms |
| gene_299079 | 4,805152794 | -5,659295901 | 1,772031274 | -3,193677214 | 0,001404731 | 0,026755825 | The Moose: suction sampler vs FISH | Signal transduction mechanisms |
| gene_904163 | 6,420412961 | -5,520971955 | 1,601710423 | -3,446922662 | 0,000567011 | 0,015137809 | The Moose: suction sampler vs FISH | Signal transduction mechanisms |
| gene_1702505 | 6,968290055 | -4,848718567 | 1,608078558 | -3,015224935 | 0,002567887 | 0,038434581 | The Moose: suction sampler vs FISH | Signal transduction mechanisms |
| gene_1353217 | 24,73914528 | -4,462957113 | 1,06713452 | -4,182187934 | 2,89E-05 | 0,002161553 | The Moose: suction sampler vs FISH | Signal transduction mechanisms |
| gene_210094 | 9,746961588 | -3,803299348 | 1,268241739 | -2,998875712 | 0,002709778 | 0,0395379 | The Moose: suction sampler vs FISH | Signal transduction mechanisms |
| gene_548857 | 19,07744226 | -3,37342279 | 1,120443456 | -3,010792531 | 0,002605668 | 0,038790953 | The Moose: suction sampler vs FISH | Signal transduction mechanisms |
| gene_703248 | 44,3099952 | -2,63503142 | 0,811552923 | -3,246900288 | 0,001166693 | 0,023920816 | The Moose: suction sampler vs FISH | Signal transduction mechanisms |
| gene_54431 | 840,6035872 | -2,634845869 | 0,603808056 | -4,363714331 | 1,28E-05 | 0,001251288 | The Moose: suction sampler vs FISH | Signal transduction mechanisms |

| | | | | | | | | |
|---|---|---|---|---|---|---|---|---|
| gene_291635 | 85,78522928 | -2,537238914 | 0,878301437 | -2,888801962 | 0,003867125 | 0,048863897 | The Moose: suction sampler vs FISH | Signal transduction mechanisms |
| gene_1278445 | 201,0116695 | -2,410571501 | 0,561139669 | -4,295849385 | 1,74E-05 | 0,001523399 | The Moose: suction sampler vs FISH | Signal transduction mechanisms |
| gene_1430523 | 44,61764781 | -2,379040394 | 0,826316155 | -2,879092196 | 0,003988217 | 0,049722393 | The Moose: suction sampler vs FISH | Signal transduction mechanisms |
| gene_791211 | 37,73852232 | -2,330796374 | 0,760280364 | -3,065706396 | 0,002171564 | 0,034799623 | The Moose: suction sampler vs FISH | Signal transduction mechanisms |
| gene_1885232 | 475,7094185 | -2,325181656 | 0,535316235 | -4,343566483 | 1,40E-05 | 0,001323407 | The Moose: suction sampler vs FISH | Signal transduction mechanisms |
| gene_591557 | 929,7518128 | -2,300913136 | 0,6201978 | -3,709966617 | 0,000207287 | 0,008008502 | The Moose: suction sampler vs FISH | Signal transduction mechanisms |
| gene_97142 | 259,167368 | -2,24255756 | 0,540799517 | -4,146744757 | 3,37E-05 | 0,002404687 | The Moose: suction sampler vs FISH | Signal transduction mechanisms |
| gene_1465603 | 107,7146815 | -2,133149592 | 0,641768992 | -3,323858926 | 0,000887811 | 0,0203195 | The Moose: suction sampler vs FISH | Signal transduction mechanisms |
| gene_1444290 | 706,8734003 | -2,116401247 | 0,62125362 | -3,406662236 | 0,000657625 | 0,016649733 | The Moose: suction sampler vs FISH | Signal transduction mechanisms |
| gene_1444288 | 2938,739008 | -2,114495472 | 0,604838379 | -3,495967757 | 0,000472346 | 0,013441083 | The Moose: suction sampler vs FISH | Signal transduction mechanisms |
| gene_566431 | 87,97297109 | -2,076139034 | 0,676693875 | -3,0680624 | 0,002154516 | 0,034663405 | The Moose: suction sampler vs FISH | Signal transduction mechanisms |
| gene_1825066 | 199,9148528 | 2,015963837 | 0,68296117 | 2,951798616 | 0,003159289 | 0,043435789 | The Moose: suction sampler vs FISH | Signal transduction mechanisms |
| gene_893691 | 56,54931679 | 2,044182175 | 0,698392554 | 2,926981629 | 0,003422691 | 0,045457421 | The Moose: suction sampler vs FISH | Signal transduction mechanisms |
| gene_193092 | 45,51229907 | 2,072011212 | 0,692042177 | 2,994053371 | 0,002752978 | 0,039919537 | The Moose: suction sampler vs FISH | Signal transduction mechanisms |
| gene_309826 | 97,46976969 | 2,144890413 | 0,716137893 | 2,995080185 | 0,002743727 | 0,039832582 | The Moose: suction sampler vs FISH | Signal transduction mechanisms |
| gene_295851 | 176,1127375 | 2,182453449 | 0,741955672 | 2,941487655 | 0,003266399 | 0,044256007 | The Moose: suction sampler vs FISH | Signal transduction mechanisms |
| gene_1352683 | 157,7580321 | 2,19683273 | 0,759111569 | 2,893952377 | 0,003804259 | 0,048389984 | The Moose: suction sampler vs FISH | Signal transduction mechanisms |
| gene_480783 | 46,28251814 | 2,203853118 | 0,763622074 | 2,886052138 | 0,003901075 | 0,049060697 | The Moose: suction sampler vs FISH | Signal transduction mechanisms |
| gene_1290293 | 61,96227591 | 2,222037019 | 0,727145316 | 3,055836255 | 0,002244338 | 0,035391185 | The Moose: suction sampler vs FISH | Signal transduction mechanisms |
| gene_1770783 | 54,30071833 | 2,267378648 | 0,745004906 | 3,043441231 | 0,002338891 | 0,036216894 | The Moose: suction sampler vs FISH | Signal transduction mechanisms |
| gene_253196 | 33,15051666 | 2,272358264 | 0,729568509 | 3,114660563 | 0,001841567 | 0,031612271 | The Moose: suction sampler vs FISH | Signal transduction mechanisms |
| gene_1882254 | 466,9546737 | 2,272952741 | 0,762591264 | 2,980564881 | 0,002877173 | 0,041087684 | The Moose: suction sampler vs FISH | Signal transduction mechanisms |
| gene_477154 | 103,3778523 | 2,358126118 | 0,6756988 | 3,489907216 | 0,000483188 | 0,013638648 | The Moose: suction sampler vs FISH | Signal transduction mechanisms |
| gene_701355 | 48,70528813 | 2,375657677 | 0,705679307 | 3,366483406 | 0,000761332 | 0,018350055 | The Moose: suction sampler vs FISH | Signal transduction mechanisms |
| gene_1115274 | 37,21760785 | 2,400971308 | 0,809634355 | 2,965500776 | 0,003021906 | 0,042362843 | The Moose: suction sampler vs FISH | Signal transduction mechanisms |
| gene_281303 | 91,98281335 | 2,412603825 | 0,690990632 | 3,491514519 | 0,00048029 | 0,013577723 | The Moose: suction sampler vs FISH | Signal transduction mechanisms |
| gene_898493 | 32,8461384 | 2,416239118 | 0,824005714 | 2,932308692 | 0,003364521 | 0,044994041 | The Moose: suction sampler vs FISH | Signal transduction mechanisms |
| gene_657203 | 205,2783876 | 2,425550922 | 0,746147448 | 3,250766225 | 0,001150945 | 0,023744697 | The Moose: suction sampler vs FISH | Signal transduction mechanisms |
| gene_131093 | 30,99348916 | 2,438578444 | 0,772177911 | 3,158052579 | 0,001588269 | 0,028868131 | The Moose: suction sampler vs FISH | Signal transduction mechanisms |
| gene_405568 | 42,4226581 | 2,447688381 | 0,801115255 | 3,055351106 | 0,002247972 | 0,035414478 | The Moose: suction sampler vs FISH | Signal transduction mechanisms |
| gene_475241 | 319,5935161 | 2,459619315 | 0,747049496 | 3,292444915 | 0,000993204 | 0,021630873 | The Moose: suction sampler vs FISH | Signal transduction mechanisms |
| gene_1455933 | 38,30384564 | 2,466994843 | 0,815652807 | 3,024564892 | 0,002489911 | 0,037721014 | The Moose: suction sampler vs FISH | Signal transduction mechanisms |
| gene_1582609 | 34,9398066 | 2,501576504 | 0,857671703 | 2,916706353 | 0,003537486 | 0,046312066 | The Moose: suction sampler vs FISH | Signal transduction mechanisms |
| gene_1189342 | 78,76069639 | 2,507133945 | 0,668864662 | 3,748342659 | 0,000178007 | 0,007226628 | The Moose: suction sampler vs FISH | Signal transduction mechanisms |
| gene_111062 | 183,1039735 | 2,517273943 | 0,703491838 | 3,578256072 | 0,000345894 | 0,011122688 | The Moose: suction sampler vs FISH | Signal transduction mechanisms |
| gene_610634 | 59,17019703 | 2,54811391 | 0,866352965 | 2,941196041 | 0,003269475 | 0,044279296 | The Moose: suction sampler vs FISH | Signal transduction mechanisms |
| gene_1352682 | 319,733107 | 2,554324929 | 0,831737658 | 3,071070431 | 0,002132928 | 0,034462996 | The Moose: suction sampler vs FISH | Signal transduction mechanisms |
| gene_309825 | 66,08033584 | 2,563279443 | 0,777414477 | 3,297185117 | 0,000976591 | 0,021404278 | The Moose: suction sampler vs FISH | Signal transduction mechanisms |
| gene_833677 | 28,57436001 | 2,566170563 | 0,882744856 | 2,90703542 | 0,003648719 | 0,047262814 | The Moose: suction sampler vs FISH | Signal transduction mechanisms |
| gene_1881264 | 91,81202158 | 2,567016047 | 0,884266336 | 2,90298958 | 0,003696188 | 0,047675894 | The Moose: suction sampler vs FISH | Signal transduction mechanisms |
| gene_274710 | 114,8551893 | 2,567868382 | 0,774185644 | 3,316863859 | 0,00091034 | 0,020591283 | The Moose: suction sampler vs FISH | Signal transduction mechanisms |
| gene_111067 | 96,7859376 | 2,568595337 | 0,803835666 | 3,195423448 | 0,001396258 | 0,026642812 | The Moose: suction sampler vs FISH | Signal transduction mechanisms |
| gene_1571625 | 79,60168211 | 2,574234931 | 0,817904701 | 3,147353143 | 0,001647558 | 0,029556651 | The Moose: suction sampler vs FISH | Signal transduction mechanisms |
| gene_1115615 | 41,21870676 | 2,578591312 | 0,890429022 | 2,895897648 | 0,003780757 | 0,048233054 | The Moose: suction sampler vs FISH | Signal transduction mechanisms |
| gene_51164 | 38,64005102 | 2,581049303 | 0,847207073 | 3,046538899 | 0,002314925 | 0,036030135 | The Moose: suction sampler vs FISH | Signal transduction mechanisms |
| gene_859304 | 314,7298308 | 2,597793267 | 0,72882179 | 3,564373763 | 0,000364726 | 0,011490192 | The Moose: suction sampler vs FISH | Signal transduction mechanisms |
| gene_616265 | 32,1459011 | 2,62018042 | 0,887101069 | 2,953643629 | 0,003140464 | 0,043306684 | The Moose: suction sampler vs FISH | Signal transduction mechanisms |
| gene_1339613 | 17,61300302 | 2,645721042 | 0,919012007 | 2,878875382 | 0,00399096 | 0,049748143 | The Moose: suction sampler vs FISH | Signal transduction mechanisms |
| gene_754768 | 69,43710936 | 2,657490171 | 0,856252474 | 3,103629188 | 0,001911628 | 0,032281658 | The Moose: suction sampler vs FISH | Signal transduction mechanisms |
| gene_696982 | 76,60466971 | 2,663235465 | 0,801385512 | 3,323288758 | 0,000889628 | 0,020348399 | The Moose: suction sampler vs FISH | Signal transduction mechanisms |
| gene_1903246 | 207,2733802 | 2,678128835 | 0,931267127 | 2,875790156 | 0,004030175 | 0,049999364 | The Moose: suction sampler vs FISH | Signal transduction mechanisms |
| gene_84726 | 39,40719264 | 2,679864936 | 0,903914661 | 2,964732238 | 0,003029465 | 0,042414374 | The Moose: suction sampler vs FISH | Signal transduction mechanisms |
| gene_700499 | 350,5159904 | 2,684217062 | 0,696419308 | 3,854311662 | 0,000116056 | 0,005506272 | The Moose: suction sampler vs FISH | Signal transduction mechanisms |
| gene_1252753 | 91,01940862 | 2,687118244 | 0,802878172 | 3,346856767 | 0,000817334 | 0,019203173 | The Moose: suction sampler vs FISH | Signal transduction mechanisms |
| gene_1008752 | 15,27742033 | 2,691733582 | 0,906419755 | 2,96963252 | 0,002981562 | 0,041974469 | The Moose: suction sampler vs FISH | Signal transduction mechanisms |
| gene_223606 | 21,68134577 | 2,692961659 | 0,926238437 | 2,907417303 | 0,003644267 | 0,047221803 | The Moose: suction sampler vs FISH | Signal transduction mechanisms |
| gene_1795850 | 35,98671472 | 2,706257774 | 0,824944845 | 3,280531772 | 0,001036116 | 0,022244237 | The Moose: suction sampler vs FISH | Signal transduction mechanisms |
| gene_1235962 | 45,01114282 | 2,712841148 | 0,70828491 | 3,830155225 | 0,000128062 | 0,005878467 | The Moose: suction sampler vs FISH | Signal transduction mechanisms |
| gene_256016 | 19,44354035 | 2,732588876 | 0,882243827 | 3,097317081 | 0,001952809 | 0,032684422 | The Moose: suction sampler vs FISH | Signal transduction mechanisms |
| gene_1009779 | 16,43371794 | 2,735861217 | 0,901945681 | 3,033288228 | 0,002419044 | 0,036985348 | The Moose: suction sampler vs FISH | Signal transduction mechanisms |
| gene_1802434 | 26,46089614 | 2,765700281 | 0,85418013 | 3,237841976 | 0,001204375 | 0,024388013 | The Moose: suction sampler vs FISH | Signal transduction mechanisms |
| gene_1901345 | 223,5568422 | 2,767265079 | 0,664147365 | 4,166643168 | 3,09E-05 | 0,002260777 | The Moose: suction sampler vs FISH | Signal transduction mechanisms |
| gene_803734 | 22,01819295 | 2,786996865 | 0,965984728 | 2,885135536 | 0,003912452 | 0,049132477 | The Moose: suction sampler vs FISH | Signal transduction mechanisms |
| gene_1937494 | 100,3441563 | 2,793045856 | 0,713539117 | 3,914355623 | 9,06E-05 | 0,004696249 | The Moose: suction sampler vs FISH | Signal transduction mechanisms |
| gene_160208 | 922,6809985 | 2,805085431 | 0,745840738 | 3,76097106 | 0,000169255 | 0,007025858 | The Moose: suction sampler vs FISH | Signal transduction mechanisms |

| | | | | | | | | |
|---|---|---|---|---|---|---|---|---|
| gene_1382270 | 26,41420743 | 2,814444256 | 0,888301646 | 3,168342949 | 0,001533106 | 0,028296484 | The Moose: suction sampler vs FISH | Signal transduction mechanisms |
| gene_265975 | 31,55944238 | 2,814583086 | 0,825727006 | 3,408612125 | 0,000652942 | 0,016559171 | The Moose: suction sampler vs FISH | Signal transduction mechanisms |
| gene_844680 | 116,9242545 | 2,829999812 | 0,780552576 | 3,625636373 | 0,000288251 | 0,009920607 | The Moose: suction sampler vs FISH | Signal transduction mechanisms |
| gene_111064 | 82,80232198 | 2,836543065 | 0,795911329 | 3,563893317 | 0,000365395 | 0,011490192 | The Moose: suction sampler vs FISH | Signal transduction mechanisms |
| gene_1818234 | 177,9455115 | 2,841455055 | 0,652979657 | 4,351521556 | 1,35E-05 | 0,001296276 | The Moose: suction sampler vs FISH | Signal transduction mechanisms |
| gene_1601172 | 32,73601793 | 2,846867104 | 0,807025193 | 3,527606235 | 0,000419335 | 0,01255131 | The Moose: suction sampler vs FISH | Signal transduction mechanisms |
| gene_1154504 | 30,26441633 | 2,849193312 | 0,859192959 | 3,316127399 | 0,000912742 | 0,020601212 | The Moose: suction sampler vs FISH | Signal transduction mechanisms |
| gene_470936 | 75,87872889 | 2,850639006 | 0,817430956 | 3,487314719 | 0,000487897 | 0,013734605 | The Moose: suction sampler vs FISH | Signal transduction mechanisms |
| gene_1329859 | 138,0870355 | 2,867186598 | 0,745524502 | 3,845865011 | 0,000120128 | 0,005644781 | The Moose: suction sampler vs FISH | Signal transduction mechanisms |
| gene_1154134 | 43,72767883 | 2,885644502 | 0,756647523 | 3,813723582 | 0,000136889 | 0,006134095 | The Moose: suction sampler vs FISH | Signal transduction mechanisms |
| gene_1938 | 30,3559073 | 2,894442754 | 0,836402679 | 3,460585226 | 0,000539003 | 0,014618554 | The Moose: suction sampler vs FISH | Signal transduction mechanisms |
| gene_1091449 | 34,90861662 | 2,912996223 | 0,749332932 | 3,887452559 | 0,000101302 | 0,005037312 | The Moose: suction sampler vs FISH | Signal transduction mechanisms |
| gene_1682391 | 108,0019017 | 2,918781603 | 0,767819719 | 3,801389218 | 0,000143887 | 0,006317896 | The Moose: suction sampler vs FISH | Signal transduction mechanisms |
| gene_1600256 | 44,93944078 | 2,921873593 | 0,847426576 | 3,447937171 | 0,000564885 | 0,015103025 | The Moose: suction sampler vs FISH | Signal transduction mechanisms |
| gene_1597113 | 36,61483741 | 2,936922548 | 0,781444152 | 3,758326863 | 0,000171053 | 0,007061839 | The Moose: suction sampler vs FISH | Signal transduction mechanisms |
| gene_971724 | 32,71006195 | 2,957956114 | 0,826972484 | 3,576849499 | 0,00034776 | 0,011150745 | The Moose: suction sampler vs FISH | Signal transduction mechanisms |
| gene_1822656 | 18,87300824 | 2,963323868 | 0,966095092 | 3,067321108 | 0,002159867 | 0,034711473 | The Moose: suction sampler vs FISH | Signal transduction mechanisms |
| gene_738431 | 298,5742029 | 2,971757591 | 0,688579564 | 4,315779537 | 1,59E-05 | 0,001431463 | The Moose: suction sampler vs FISH | Signal transduction mechanisms |
| gene_1280880 | 23,54246678 | 2,972118755 | 0,848826623 | 3,501443845 | 0,000462744 | 0,013327042 | The Moose: suction sampler vs FISH | Signal transduction mechanisms |
| gene_1391476 | 12,47554679 | 3,016985041 | 0,929243929 | 3,246709446 | 0,001167475 | 0,023920816 | The Moose: suction sampler vs FISH | Signal transduction mechanisms |
| gene_1162542 | 30,84707917 | 3,039347749 | 0,838992594 | 3,622615707 | 0,000291639 | 0,009987411 | The Moose: suction sampler vs FISH | Signal transduction mechanisms |
| gene_1575060 | 10,79941721 | 3,052131097 | 1,040269287 | 2,933981745 | 0,003346439 | 0,044929334 | The Moose: suction sampler vs FISH | Signal transduction mechanisms |
| gene_1809997 | 37,05837312 | 3,054437268 | 0,827885319 | 3,689444898 | 0,000224744 | 0,008430192 | The Moose: suction sampler vs FISH | Signal transduction mechanisms |
| gene_1278532 | 127,0918482 | 3,054835918 | 0,717252612 | 4,25907953 | 2,05E-05 | 0,001696795 | The Moose: suction sampler vs FISH | Signal transduction mechanisms |
| gene_253197 | 22,04931452 | 3,082400913 | 0,948102691 | 3,251125582 | 0,001149491 | 0,023728034 | The Moose: suction sampler vs FISH | Signal transduction mechanisms |
| gene_595717 | 321,623871 | 3,107603407 | 0,763319311 | 4,071170953 | 4,68E-05 | 0,003037636 | The Moose: suction sampler vs FISH | Signal transduction mechanisms |
| gene_127592 | 15,69919106 | 3,130318057 | 1,063317482 | 2,943916668 | 0,003240872 | 0,044095556 | The Moose: suction sampler vs FISH | Signal transduction mechanisms |
| gene_940710 | 13,44673613 | 3,147259856 | 1,073433911 | 2,931954937 | 0,003368356 | 0,045028926 | The Moose: suction sampler vs FISH | Signal transduction mechanisms |
| gene_981832 | 48,50794726 | 3,150684782 | 1,032821599 | 3,05056051 | 0,002284146 | 0,035723316 | The Moose: suction sampler vs FISH | Signal transduction mechanisms |
| gene_776 | 22,17098945 | 3,15619195 | 1,038991167 | 3,037746661 | 0,002383542 | 0,036615619 | The Moose: suction sampler vs FISH | Signal transduction mechanisms |
| gene_1098881 | 134,0369726 | 3,188438449 | 0,711668199 | 4,480231732 | 7,46E-06 | 0,000881837 | The Moose: suction sampler vs FISH | Signal transduction mechanisms |
| gene_1660614 | 36,81919592 | 3,189613147 | 0,844985424 | 3,774755228 | 0,000160165 | 0,006783914 | The Moose: suction sampler vs FISH | Signal transduction mechanisms |
| gene_1663642 | 74,66957006 | 3,192843399 | 0,789148747 | 4,04593356 | 5,21E-05 | 0,003227308 | The Moose: suction sampler vs FISH | Signal transduction mechanisms |
| gene_79431 | 35,15934126 | 3,225699293 | 0,858905124 | 3,755594422 | 0,000172931 | 0,007107441 | The Moose: suction sampler vs FISH | Signal transduction mechanisms |
| gene_410008 | 44,0973757 | 3,234620217 | 0,915181974 | 3,534401143 | 0,0004087 | 0,012342513 | The Moose: suction sampler vs FISH | Signal transduction mechanisms |
| gene_1344284 | 21,00419942 | 3,244228878 | 1,096658538 | 2,958285342 | 0,003093556 | 0,042988748 | The Moose: suction sampler vs FISH | Signal transduction mechanisms |
| gene_678360 | 16,28764426 | 3,247163651 | 1,015921314 | 3,196274757 | 0,001392145 | 0,026578131 | The Moose: suction sampler vs FISH | Signal transduction mechanisms |
| gene_324581 | 135,8928613 | 3,249717167 | 0,75287836 | 4,316390722 | 1,59E-05 | 0,001429259 | The Moose: suction sampler vs FISH | Signal transduction mechanisms |
| gene_162663 | 19,07939246 | 3,276318631 | 1,074077706 | 3,050355307 | 0,002285708 | 0,035725434 | The Moose: suction sampler vs FISH | Signal transduction mechanisms |
| gene_679865 | 8,969590411 | 3,292440355 | 1,141680764 | 2,883853753 | 0,003928411 | 0,049252677 | The Moose: suction sampler vs FISH | Signal transduction mechanisms |
| gene_1103622 | 156,811193 | 3,301532563 | 0,773324468 | 4,269272084 | 1,96E-05 | 0,001631194 | The Moose: suction sampler vs FISH | Signal transduction mechanisms |
| gene_1899963 | 27,96782486 | 3,303334147 | 0,993628701 | 3,32451563 | 0,000885723 | 0,020278024 | The Moose: suction sampler vs FISH | Signal transduction mechanisms |
| gene_416656 | 40,50449181 | 3,308314459 | 1,063526653 | 3,110701974 | 0,001866432 | 0,031864273 | The Moose: suction sampler vs FISH | Signal transduction mechanisms |
| gene_1827970 | 15,9879877 | 3,323597903 | 1,049927762 | 3,165549121 | 0,001547905 | 0,028414371 | The Moose: suction sampler vs FISH | Signal transduction mechanisms |
| gene_945563 | 16,66872936 | 3,331810448 | 1,010891884 | 3,295911758 | 0,000981028 | 0,021469493 | The Moose: suction sampler vs FISH | Signal transduction mechanisms |
| gene_1679144 | 14,21557644 | 3,331907212 | 1,118207585 | 2,979685754 | 0,002885442 | 0,041141779 | The Moose: suction sampler vs FISH | Signal transduction mechanisms |
| gene_1060329 | 82,7777752 | 3,33261011 | 0,956306236 | 3,484877528 | 0,000492362 | 0,013823223 | The Moose: suction sampler vs FISH | Signal transduction mechanisms |
| gene_1148605 | 15,19160417 | 3,347877799 | 0,961509787 | 3,481896747 | 0,000497876 | 0,013935395 | The Moose: suction sampler vs FISH | Signal transduction mechanisms |
| gene_47247 | 22,70174476 | 3,348983641 | 0,969180476 | 3,455479887 | 0,000549314 | 0,014796594 | The Moose: suction sampler vs FISH | Signal transduction mechanisms |
| gene_142858 | 46,90196509 | 3,352341387 | 0,978774885 | 3,425038215 | 0,000614713 | 0,015908243 | The Moose: suction sampler vs FISH | Signal transduction mechanisms |
| gene_975091 | 71,94164521 | 3,372284691 | 0,788755821 | 4,275448243 | 1,91E-05 | 0,001601127 | The Moose: suction sampler vs FISH | Signal transduction mechanisms |
| gene_923976 | 56,57387483 | 3,374094889 | 0,899265028 | 3,752058384 | 0,000175389 | 0,007168289 | The Moose: suction sampler vs FISH | Signal transduction mechanisms |
| gene_1344524 | 27,15336867 | 3,380545971 | 0,849232308 | 3,980708151 | 6,87E-05 | 0,003884854 | The Moose: suction sampler vs FISH | Signal transduction mechanisms |
| gene_1647548 | 9,836637903 | 3,387934394 | 1,14251454 | 2,965331534 | 0,003023569 | 0,042362843 | The Moose: suction sampler vs FISH | Signal transduction mechanisms |
| gene_292692 | 159,2489537 | 3,399477761 | 0,715471479 | 4,751381236 | 2,02E-06 | 0,000350538 | The Moose: suction sampler vs FISH | Signal transduction mechanisms |
| gene_1258064 | 30,50002789 | 3,400124042 | 0,810492511 | 4,195133201 | 2,73E-05 | 0,002090739 | The Moose: suction sampler vs FISH | Signal transduction mechanisms |
| gene_1648233 | 12,76319325 | 3,417180803 | 1,160274059 | 2,945149704 | 0,003227984 | 0,044005316 | The Moose: suction sampler vs FISH | Signal transduction mechanisms |
| gene_1907757 | 57,30557485 | 3,417795719 | 0,754277609 | 4,531217258 | 5,86E-06 | 0,000740063 | The Moose: suction sampler vs FISH | Signal transduction mechanisms |
| gene_1518146 | 9,686900923 | 3,418606764 | 1,098137406 | 3,113095634 | 0,00185136 | 0,031687986 | The Moose: suction sampler vs FISH | Signal transduction mechanisms |
| gene_1258065 | 20,24810396 | 3,418850667 | 1,039714593 | 3,288258807 | 0,001008091 | 0,021818264 | The Moose: suction sampler vs FISH | Signal transduction mechanisms |
| gene_1167363 | 28,49982427 | 3,419591262 | 0,833739138 | 4,101512222 | 4,10E-05 | 0,002754828 | The Moose: suction sampler vs FISH | Signal transduction mechanisms |
| gene_1685745 | 14,30492793 | 3,460759149 | 1,089254399 | 3,177181706 | 0,001487138 | 0,027756766 | The Moose: suction sampler vs FISH | Signal transduction mechanisms |
| gene_618883 | 15,04292198 | 3,461992019 | 1,002061284 | 3,454870549 | 0,000550557 | 0,014817027 | The Moose: suction sampler vs FISH | Signal transduction mechanisms |
| gene_1812389 | 10,60332676 | 3,482228385 | 1,157638962 | 3,008043526 | 0,002629355 | 0,038918372 | The Moose: suction sampler vs FISH | Signal transduction mechanisms |
| gene_198053 | 36,92786994 | 3,506588901 | 0,802532274 | 4,369405462 | 1,25E-05 | 0,001225637 | The Moose: suction sampler vs FISH | Signal transduction mechanisms |

| | | | | | | | | |
|---|---|---|---|---|---|---|---|---|
| gene_844676 | 40,59300419 | 3,508789214 | 0,985866766 | 3,559090676 | 0,000372141 | 0,011615767 | The Moose: suction sampler vs FISH | Signal transduction mechanisms |
| gene_787111 | 22,54503857 | 3,540075002 | 1,022793307 | 3,461183192 | 0,000537807 | 0,014596901 | The Moose: suction sampler vs FISH | Signal transduction mechanisms |
| gene_563366 | 12,59141959 | 3,54313427 | 1,050176251 | 3,373847264 | 0,000741255 | 0,01803053 | The Moose: suction sampler vs FISH | Signal transduction mechanisms |
| gene_1949403 | 12,21481362 | 3,591870556 | 1,136591912 | 3,160211257 | 0,001576548 | 0,028734792 | The Moose: suction sampler vs FISH | Signal transduction mechanisms |
| gene_1484722 | 30,73634831 | 3,628932283 | 0,877783199 | 4,134201116 | 3,56E-05 | 0,002484376 | The Moose: suction sampler vs FISH | Signal transduction mechanisms |
| gene_1449232 | 99,17330313 | 3,635013957 | 0,875386576 | 4,1524671 | 3,29E-05 | 0,002356761 | The Moose: suction sampler vs FISH | Signal transduction mechanisms |
| gene_634855 | 13,28636312 | 3,650542066 | 0,961786788 | 3,795583504 | 0,000147297 | 0,006389958 | The Moose: suction sampler vs FISH | Signal transduction mechanisms |
| gene_672035 | 10,4901863 | 3,653039767 | 1,193674907 | 3,060330536 | 0,002210928 | 0,035162762 | The Moose: suction sampler vs FISH | Signal transduction mechanisms |
| gene_306973 | 19,66843643 | 3,655799144 | 1,104997239 | 3,30842378 | 0,000938227 | 0,020925629 | The Moose: suction sampler vs FISH | Signal transduction mechanisms |
| gene_789219 | 17,26451647 | 3,667223068 | 0,997803729 | 3,675295013 | 0,000237575 | 0,008763775 | The Moose: suction sampler vs FISH | Signal transduction mechanisms |
| gene_1523566 | 509,3142836 | 3,673636905 | 0,788939673 | 4,656423082 | 3,22E-06 | 0,000477393 | The Moose: suction sampler vs FISH | Signal transduction mechanisms |
| gene_1591107 | 41,21317747 | 3,694556493 | 1,22952762 | 3,004858479 | 0,002657044 | 0,039099706 | The Moose: suction sampler vs FISH | Signal transduction mechanisms |
| gene_1148499 | 15,52475388 | 3,73246447 | 0,998938646 | 3,736430145 | 0,000186651 | 0,007470628 | The Moose: suction sampler vs FISH | Signal transduction mechanisms |
| gene_1295228 | 54,78892145 | 3,74635314 | 0,863606071 | 4,338034742 | 1,44E-05 | 0,001337845 | The Moose: suction sampler vs FISH | Signal transduction mechanisms |
| gene_1547049 | 12,71360304 | 3,7583873 | 1,139482296 | 3,298328821 | 0,000972622 | 0,021349132 | The Moose: suction sampler vs FISH | Signal transduction mechanisms |
| gene_1109782 | 43,54308081 | 3,763853455 | 0,742090287 | 5,071961622 | 3,94E-07 | 0,000118614 | The Moose: suction sampler vs FISH | Signal transduction mechanisms |
| gene_375162 | 13,16505098 | 3,765699094 | 1,025281882 | 3,672842717 | 0,000239867 | 0,008805457 | The Moose: suction sampler vs FISH | Signal transduction mechanisms |
| gene_1183703 | 16,01348436 | 3,767332082 | 1,066615719 | 3,532042531 | 0,000412363 | 0,012407978 | The Moose: suction sampler vs FISH | Signal transduction mechanisms |
| gene_1944613 | 572,700759 | 3,774041947 | 0,806068083 | 4,682038688 | 2,84E-06 | 0,000437636 | The Moose: suction sampler vs FISH | Signal transduction mechanisms |
| gene_1573902 | 9,37673297 | 3,779935118 | 1,167291782 | 3,238209312 | 0,001202825 | 0,024376794 | The Moose: suction sampler vs FISH | Signal transduction mechanisms |
| gene_1647553 | 10,46311216 | 3,803190712 | 1,21201101 | 3,137917628 | 0,001701527 | 0,03013471 | The Moose: suction sampler vs FISH | Signal transduction mechanisms |
| gene_269095 | 9,661005462 | 3,818491168 | 1,290890105 | 2,95802962 | 0,003096124 | 0,043002044 | The Moose: suction sampler vs FISH | Signal transduction mechanisms |
| gene_1401102 | 55,94687618 | 3,825782382 | 0,779946914 | 4,90518305 | 9,33E-07 | 0,00021423 | The Moose: suction sampler vs FISH | Signal transduction mechanisms |
| gene_1761821 | 22,97309027 | 3,872480116 | 0,92123569 | 4,203571526 | 2,63E-05 | 0,002029091 | The Moose: suction sampler vs FISH | Signal transduction mechanisms |
| gene_1049920 | 22,26994425 | 3,890036779 | 0,990299682 | 3,928140995 | 8,56E-05 | 0,004511143 | The Moose: suction sampler vs FISH | Signal transduction mechanisms |
| gene_862516 | 24,73458301 | 3,89259693 | 1,006093563 | 3,869020807 | 0,000109273 | 0,005290428 | The Moose: suction sampler vs FISH | Signal transduction mechanisms |
| gene_1321819 | 7,569268822 | 3,973173152 | 1,32184793 | 3,005771739 | 0,002649078 | 0,039038312 | The Moose: suction sampler vs FISH | Signal transduction mechanisms |
| gene_940913 | 10,63295148 | 3,98256698 | 1,304274312 | 3,053473447 | 0,002262087 | 0,035545359 | The Moose: suction sampler vs FISH | Signal transduction mechanisms |
| gene_1762527 | 11,0332342 | 3,989109228 | 1,202758031 | 3,316634872 | 0,000911086 | 0,020592991 | The Moose: suction sampler vs FISH | Signal transduction mechanisms |
| gene_1332844 | 12,33743579 | 4,006214116 | 1,124622571 | 3,562274329 | 0,000367656 | 0,011538489 | The Moose: suction sampler vs FISH | Signal transduction mechanisms |
| gene_47488 | 13,74207493 | 4,010052622 | 1,366036792 | 2,935537788 | 0,003329701 | 0,044797555 | The Moose: suction sampler vs FISH | Signal transduction mechanisms |
| gene_1546741 | 10,98224694 | 4,024744041 | 1,177117984 | 3,419150923 | 0,000628169 | 0,016135014 | The Moose: suction sampler vs FISH | Signal transduction mechanisms |
| gene_316039 | 8,907349041 | 4,0500328 | 1,315604427 | 3,078457869 | 0,002080749 | 0,0340282 | The Moose: suction sampler vs FISH | Signal transduction mechanisms |
| gene_1364495 | 21,04369797 | 4,050774349 | 1,052889955 | 3,847291285 | 0,000119431 | 0,005622824 | The Moose: suction sampler vs FISH | Signal transduction mechanisms |
| gene_265976 | 12,34780597 | 4,060193717 | 1,081868968 | 3,752944062 | 0,00017477 | 0,007154946 | The Moose: suction sampler vs FISH | Signal transduction mechanisms |
| gene_548123 | 9,712573338 | 4,060524148 | 1,127977085 | 3,59982858 | 0,000318427 | 0,010582296 | The Moose: suction sampler vs FISH | Signal transduction mechanisms |
| gene_860334 | 47,72763994 | 4,065804107 | 0,844651851 | 4,813585744 | 1,48E-06 | 0,000288804 | The Moose: suction sampler vs FISH | Signal transduction mechanisms |
| gene_137429 | 17,27045962 | 4,081080445 | 1,163210117 | 3,508463678 | 0,000450703 | 0,013094092 | The Moose: suction sampler vs FISH | Signal transduction mechanisms |
| gene_618234 | 18,7533223 | 4,084816138 | 1,114160269 | 3,666273382 | 0,000246111 | 0,008947252 | The Moose: suction sampler vs FISH | Signal transduction mechanisms |
| gene_1699793 | 8,794388256 | 4,087586168 | 1,280788416 | 3,191460915 | 0,001415553 | 0,026885368 | The Moose: suction sampler vs FISH | Signal transduction mechanisms |
| gene_449703 | 36,11056976 | 4,103011175 | 1,419566641 | 2,890326566 | 0,003848418 | 0,048731896 | The Moose: suction sampler vs FISH | Signal transduction mechanisms |
| gene_1792120 | 20,66581365 | 4,106903774 | 0,968892662 | 4,238760324 | 2,25E-05 | 0,001811999 | The Moose: suction sampler vs FISH | Signal transduction mechanisms |
| gene_1359103 | 12,0605691 | 4,126180462 | 1,225228949 | 3,367681171 | 0,000758032 | 0,01829565 | The Moose: suction sampler vs FISH | Signal transduction mechanisms |
| gene_1058479 | 16,8554937 | 4,128721414 | 1,036470677 | 3,983442567 | 6,79E-05 | 0,003858225 | The Moose: suction sampler vs FISH | Signal transduction mechanisms |
| gene_549517 | 27,62901857 | 4,139941777 | 1,099533475 | 3,765180297 | 0,000166429 | 0,006956957 | The Moose: suction sampler vs FISH | Signal transduction mechanisms |
| gene_448089 | 10,43153509 | 4,140304665 | 1,179615635 | 3,509876049 | 0,000448316 | 0,013040214 | The Moose: suction sampler vs FISH | Signal transduction mechanisms |
| gene_1563429 | 50,00848823 | 4,142026171 | 0,832681212 | 4,974324041 | 6,55E-07 | 0,000164687 | The Moose: suction sampler vs FISH | Signal transduction mechanisms |
| gene_1879550 | 8,396663638 | 4,14449303 | 1,372200103 | 3,020327006 | 0,002525019 | 0,037994269 | The Moose: suction sampler vs FISH | Signal transduction mechanisms |
| gene_346130 | 6,733233291 | 4,150702076 | 1,382501851 | 3,00231213 | 0,002679373 | 0,039306464 | The Moose: suction sampler vs FISH | Signal transduction mechanisms |
| gene_189017 | 30,57790222 | 4,163628899 | 0,779119033 | 5,344021546 | 9,09E-08 | 4,54E-05 | The Moose: suction sampler vs FISH | Signal transduction mechanisms |
| gene_552382 | 10,70862556 | 4,196107161 | 1,285954966 | 3,263028079 | 0,001102286 | 0,023164519 | The Moose: suction sampler vs FISH | Signal transduction mechanisms |
| gene_1900922 | 7,15937134 | 4,202182601 | 1,396708264 | 3,008633019 | 0,002624259 | 0,038881355 | The Moose: suction sampler vs FISH | Signal transduction mechanisms |
| gene_291387 | 14,03273383 | 4,206273696 | 1,2941296 | 3,250272381 | 0,001152945 | 0,023765945 | The Moose: suction sampler vs FISH | Signal transduction mechanisms |
| gene_1321818 | 10,37123493 | 4,209840341 | 1,148745612 | 3,664728116 | 0,000247601 | 0,008988266 | The Moose: suction sampler vs FISH | Signal transduction mechanisms |
| gene_119501 | 15,10032155 | 4,233294441 | 1,110409954 | 3,812370762 | 0,00013764 | 0,006145279 | The Moose: suction sampler vs FISH | Signal transduction mechanisms |
| gene_1060344 | 14,27992493 | 4,262505591 | 1,151765514 | 3,700844954 | 0,000214883 | 0,008176582 | The Moose: suction sampler vs FISH | Signal transduction mechanisms |
| gene_1648229 | 8,532770345 | 4,287090403 | 1,259741519 | 3,403150836 | 0,000666135 | 0,016770408 | The Moose: suction sampler vs FISH | Signal transduction mechanisms |
| gene_1912117 | 7,636159337 | 4,315397435 | 1,267746266 | 3,403991438 | 0,000664088 | 0,016737807 | The Moose: suction sampler vs FISH | Signal transduction mechanisms |
| gene_1552679 | 11,15508794 | 4,317328717 | 1,138184939 | 3,793169782 | 0,000148736 | 0,006426454 | The Moose: suction sampler vs FISH | Signal transduction mechanisms |
| gene_251360 | 11,18890887 | 4,345363614 | 1,390179051 | 3,125758232 | 0,001773473 | 0,030843646 | The Moose: suction sampler vs FISH | Signal transduction mechanisms |
| gene_56344 | 18,69253613 | 4,34703192 | 1,230511843 | 3,53270222 | 0,000411336 | 0,012396609 | The Moose: suction sampler vs FISH | Signal transduction mechanisms |
| gene_1554426 | 10,09611248 | 4,386657441 | 1,292861453 | 3,392983395 | 0,000691358 | 0,017165926 | The Moose: suction sampler vs FISH | Signal transduction mechanisms |
| gene_278730 | 18,10174459 | 4,386773649 | 1,115459879 | 3,932704107 | 8,40E-05 | 0,004450979 | The Moose: suction sampler vs FISH | Signal transduction mechanisms |
| gene_1601780 | 8,650409701 | 4,394559431 | 1,25124249 | 3,512156491 | 0,000444486 | 0,012980234 | The Moose: suction sampler vs FISH | Signal transduction mechanisms |
| gene_1822654 | 13,36292238 | 4,436895971 | 1,059900675 | 4,186143169 | 2,84E-05 | 0,002146125 | The Moose: suction sampler vs FISH | Signal transduction mechanisms |

| | | | | | | | | |
|---|---|---|---|---|---|---|---|---|
| gene_406543 | 6,349833576 | 4,443496036 | 1,529583206 | 2,905037149 | 0,003672095 | 0,0474274 | The Moose: suction sampler vs FISH | Signal transduction mechanisms |
| gene_1477251 | 231,1171892 | 4,446648059 | 0,813157287 | 5,468373865 | 4,54E-08 | 2,78E-05 | The Moose: suction sampler vs FISH | Signal transduction mechanisms |
| gene_1563899 | 12,29892924 | 4,45032504 | 1,096380429 | 4,059106605 | 4,93E-05 | 0,003124319 | The Moose: suction sampler vs FISH | Signal transduction mechanisms |
| gene_1006724 | 9,33620383 | 4,469016951 | 1,540612803 | 2,90080476 | 0,003722057 | 0,047866657 | The Moose: suction sampler vs FISH | Signal transduction mechanisms |
| gene_1306222 | 9,49914312 | 4,477643272 | 1,28469446 | 3,485376025 | 0,000491446 | 0,013813335 | The Moose: suction sampler vs FISH | Signal transduction mechanisms |
| gene_1433805 | 5,608640935 | 4,516597992 | 1,54643995 | 2,920642337 | 0,003493106 | 0,046026402 | The Moose: suction sampler vs FISH | Signal transduction mechanisms |
| gene_1552678 | 12,92705234 | 4,517585149 | 1,055179939 | 4,281341012 | 1,86E-05 | 0,001574414 | The Moose: suction sampler vs FISH | Signal transduction mechanisms |
| gene_1827969 | 12,40105474 | 4,518807503 | 1,285434858 | 3,515392069 | 0,000439105 | 0,012910361 | The Moose: suction sampler vs FISH | Signal transduction mechanisms |
| gene_882758 | 34,08891741 | 4,52489173 | 0,977934139 | 4,626990255 | 3,71E-06 | 0,000533257 | The Moose: suction sampler vs FISH | Signal transduction mechanisms |
| gene_1275291 | 10,28414751 | 4,544035238 | 1,319712047 | 3,443202059 | 0,00057487 | 0,015255253 | The Moose: suction sampler vs FISH | Signal transduction mechanisms |
| gene_549518 | 17,95791371 | 4,545500225 | 0,994528643 | 4,570507102 | 4,87E-06 | 0,000647361 | The Moose: suction sampler vs FISH | Signal transduction mechanisms |
| gene_1930914 | 42,5248908 | 4,553843413 | 1,028729067 | 4,426669334 | 9,57E-06 | 0,001029043 | The Moose: suction sampler vs FISH | Signal transduction mechanisms |
| gene_788278 | 16,7334146 | 4,602076978 | 1,370498328 | 3,357958841 | 0,000785203 | 0,018701511 | The Moose: suction sampler vs FISH | Signal transduction mechanisms |
| gene_1156207 | 9,108790941 | 4,623223747 | 1,342619843 | 3,443434694 | 0,000574375 | 0,015255253 | The Moose: suction sampler vs FISH | Signal transduction mechanisms |
| gene_480706 | 10,58537479 | 4,627569656 | 1,203613676 | 3,844730039 | 0,000120685 | 0,005656494 | The Moose: suction sampler vs FISH | Signal transduction mechanisms |
| gene_843811 | 10,16534765 | 4,638384976 | 1,296855389 | 3,576640091 | 0,000348039 | 0,011150745 | The Moose: suction sampler vs FISH | Signal transduction mechanisms |
| gene_1342352 | 150,0840419 | 4,639823064 | 0,75640277 | 6,134064086 | 8,57E-10 | 1,57E-06 | The Moose: suction sampler vs FISH | Signal transduction mechanisms |
| gene_1298123 | 4,908966087 | 4,669379773 | 1,623381482 | 2,876329332 | 0,004023297 | 0,049950818 | The Moose: suction sampler vs FISH | Signal transduction mechanisms |
| gene_696520 | 14,57425174 | 4,669790873 | 1,28213518 | 3,642198534 | 0,000270319 | 0,009517412 | The Moose: suction sampler vs FISH | Signal transduction mechanisms |
| gene_1643269 | 7,355630956 | 4,691511204 | 1,545797062 | 3,035011076 | 0,002405269 | 0,036864555 | The Moose: suction sampler vs FISH | Signal transduction mechanisms |
| gene_824160 | 6,564783977 | 4,698124003 | 1,334041315 | 3,521723014 | 0,000428752 | 0,01270262 | The Moose: suction sampler vs FISH | Signal transduction mechanisms |
| gene_1936449 | 8,799680059 | 4,719341902 | 1,466709644 | 3,217638829 | 0,001292505 | 0,025470352 | The Moose: suction sampler vs FISH | Signal transduction mechanisms |
| gene_107683 | 12,50897934 | 4,767665037 | 1,441727735 | 3,306910815 | 0,000943309 | 0,020987987 | The Moose: suction sampler vs FISH | Signal transduction mechanisms |
| gene_312575 | 240,4865263 | 4,796264981 | 0,960436701 | 4,993837674 | 5,92E-07 | 0,000152537 | The Moose: suction sampler vs FISH | Signal transduction mechanisms |
| gene_47714 | 14,07393707 | 4,798392493 | 1,195434897 | 4,013930417 | 5,97E-05 | 0,003539821 | The Moose: suction sampler vs FISH | Signal transduction mechanisms |
| gene_398243 | 8,77044945 | 4,799110808 | 1,414674563 | 3,39237796 | 0,000692888 | 0,017192281 | The Moose: suction sampler vs FISH | Signal transduction mechanisms |
| gene_763861 | 6,936735782 | 4,812555101 | 1,510421002 | 3,186234233 | 0,001441378 | 0,027206899 | The Moose: suction sampler vs FISH | Signal transduction mechanisms |
| gene_412993 | 5,915356847 | 4,833449844 | 1,532384197 | 3,154202356 | 0,001609375 | 0,029134955 | The Moose: suction sampler vs FISH | Signal transduction mechanisms |
| gene_102648 | 4,310468721 | 4,836949285 | 1,649769678 | 2,931893676 | 0,003369021 | 0,045029613 | The Moose: suction sampler vs FISH | Signal transduction mechanisms |
| gene_477076 | 5,198058275 | 4,873033369 | 1,588302631 | 3,068076116 | 0,002154417 | 0,034663405 | The Moose: suction sampler vs FISH | Signal transduction mechanisms |
| gene_1240146 | 134,3908744 | 4,895153358 | 0,974551088 | 5,022982804 | 5,09E-07 | 0,000142073 | The Moose: suction sampler vs FISH | Signal transduction mechanisms |
| gene_853712 | 22,82479721 | 4,906683804 | 1,091525744 | 4,495252475 | 6,95E-06 | 0,00083664 | The Moose: suction sampler vs FISH | Signal transduction mechanisms |
| gene_695409 | 9,337512515 | 4,923013646 | 1,374231517 | 3,582375737 | 0,000340484 | 0,011015763 | The Moose: suction sampler vs FISH | Signal transduction mechanisms |
| gene_1605306 | 29,97439971 | 4,952859959 | 0,96420537 | 5,136727207 | 2,80E-07 | 9,23E-05 | The Moose: suction sampler vs FISH | Signal transduction mechanisms |
| gene_1137277 | 19,18361778 | 5,000327586 | 1,043021927 | 4,794077148 | 1,63E-06 | 0,000308493 | The Moose: suction sampler vs FISH | Signal transduction mechanisms |
| gene_994678 | 5,470703544 | 5,006498485 | 1,72793718 | 2,897384548 | 0,003762882 | 0,048129883 | The Moose: suction sampler vs FISH | Signal transduction mechanisms |
| gene_1552455 | 6,921829136 | 5,031878412 | 1,653231811 | 3,043661741 | 0,002337178 | 0,036208609 | The Moose: suction sampler vs FISH | Signal transduction mechanisms |
| gene_975090 | 11,98145048 | 5,044220336 | 1,260470356 | 4,001855587 | 6,28E-05 | 0,003657564 | The Moose: suction sampler vs FISH | Signal transduction mechanisms |
| gene_800238 | 11,76831038 | 5,055557451 | 1,193333687 | 4,2364994 | 2,27E-05 | 0,001822056 | The Moose: suction sampler vs FISH | Signal transduction mechanisms |
| gene_1743688 | 5,901135192 | 5,082187892 | 1,689633343 | 3,007864347 | 0,002630905 | 0,038933478 | The Moose: suction sampler vs FISH | Signal transduction mechanisms |
| gene_1469695 | 6,209059514 | 5,092834456 | 1,719385255 | 2,962008917 | 0,003056389 | 0,042636836 | The Moose: suction sampler vs FISH | Signal transduction mechanisms |
| gene_1025733 | 6,384659591 | 5,268421437 | 1,643023276 | 3,206540963 | 0,001343412 | 0,026040348 | The Moose: suction sampler vs FISH | Signal transduction mechanisms |
| gene_670802 | 5,67027026 | 5,305155757 | 1,544425165 | 3,435035817 | 0,000592476 | 0,015518675 | The Moose: suction sampler vs FISH | Signal transduction mechanisms |
| gene_640589 | 800,7717004 | 5,311937863 | 0,825927296 | 6,431483603 | 1,26E-10 | 4,64E-07 | The Moose: suction sampler vs FISH | Signal transduction mechanisms |
| gene_548125 | 3,7399206 | 5,330706249 | 1,786064491 | 2,984610172 | 0,002839399 | 0,040802129 | The Moose: suction sampler vs FISH | Signal transduction mechanisms |
| gene_690762 | 12,90431147 | 5,423364496 | 1,292849959 | 4,194890874 | 2,73E-05 | 0,002090793 | The Moose: suction sampler vs FISH | Signal transduction mechanisms |
| gene_1490643 | 6,938870981 | 5,44900769 | 1,688978375 | 3,226215191 | 0,00125439 | 0,024987436 | The Moose: suction sampler vs FISH | Signal transduction mechanisms |
| gene_953061 | 4,72453673 | 5,458900745 | 1,79191995 | 3,046397662 | 0,002316013 | 0,036030135 | The Moose: suction sampler vs FISH | Signal transduction mechanisms |
| gene_723185 | 11,37778219 | 5,545299094 | 1,497812583 | 3,702264995 | 0,000213683 | 0,008156947 | The Moose: suction sampler vs FISH | Signal transduction mechanisms |
| gene_1300345 | 9,863024118 | 5,587992671 | 1,660385144 | 3,365479806 | 0,000764107 | 0,01838788 | The Moose: suction sampler vs FISH | Signal transduction mechanisms |
| gene_1905542 | 92,57820788 | 5,686999307 | 0,919100574 | 6,187570186 | 6,11E-10 | 1,21E-06 | The Moose: suction sampler vs FISH | Signal transduction mechanisms |
| gene_890908 | 2,941315838 | 5,808868206 | 1,980952049 | 2,932361846 | 0,003363946 | 0,044994041 | The Moose: suction sampler vs FISH | Signal transduction mechanisms |
| gene_1628808 | 9,961424236 | 5,832754522 | 1,516733418 | 3,84560296 | 0,000120256 | 0,005647205 | The Moose: suction sampler vs FISH | Signal transduction mechanisms |
| gene_845364 | 3,789432966 | 5,85625238 | 1,994043376 | 2,936873115 | 0,003315397 | 0,044711596 | The Moose: suction sampler vs FISH | Signal transduction mechanisms |
| gene_952567 | 498,3880856 | 5,87046201 | 1,656345547 | 3,54422543 | 0,000393768 | 0,012080334 | The Moose: suction sampler vs FISH | Signal transduction mechanisms |
| gene_685403 | 8,220087276 | 5,872353442 | 1,751607816 | 3,352550376 | 0,000800707 | 0,0189324 | The Moose: suction sampler vs FISH | Signal transduction mechanisms |
| gene_1589003 | 7,345490223 | 5,900800757 | 1,663039292 | 3,54820285 | 0,000387869 | 0,011961597 | The Moose: suction sampler vs FISH | Signal transduction mechanisms |
| gene_1126727 | 3,606275269 | 5,903709101 | 1,813923556 | 3,25466257 | 0,001135272 | 0,023553679 | The Moose: suction sampler vs FISH | Signal transduction mechanisms |
| gene_1602142 | 3,641827881 | 5,948673862 | 1,85738881 | 3,202707926 | 0,00136142 | 0,026236538 | The Moose: suction sampler vs FISH | Signal transduction mechanisms |
| gene_1782007 | 11,1398702 | 6,084982259 | 1,521107746 | 4,000362416 | 6,32E-05 | 0,003674897 | The Moose: suction sampler vs FISH | Signal transduction mechanisms |
| gene_923150 | 2,996729373 | 6,093736519 | 2,09192905 | 2,912974758 | 0,003580035 | 0,046686026 | The Moose: suction sampler vs FISH | Signal transduction mechanisms |
| gene_769501 | 2,710751982 | 6,103020929 | 2,054992232 | 2,969851094 | 0,002979441 | 0,041955589 | The Moose: suction sampler vs FISH | Signal transduction mechanisms |
| gene_1328401 | 3,128693717 | 6,104409131 | 2,080863774 | 2,933593832 | 0,003350624 | 0,044942491 | The Moose: suction sampler vs FISH | Signal transduction mechanisms |
| gene_602198 | 3,194370333 | 6,121050326 | 2,068818607 | 2,958717747 | 0,003089219 | 0,042944271 | The Moose: suction sampler vs FISH | Signal transduction mechanisms |
| gene_886913 | 4,38006619 | 6,121964912 | 1,846577981 | 3,315302671 | 0,000915439 | 0,02061142 | The Moose: suction sampler vs FISH | Signal transduction mechanisms |

| | | | | | | | | |
|---|---|---|---|---|---|---|---|---|
| gene_271283 | 2,838072107 | 6,122622659 | 2,031938803 | 3,01319245 | 0,002585149 | 0,038582866 | The Moose: suction sampler vs FISH | Signal transduction mechanisms |
| gene_523966 | 4,401625526 | 6,160940877 | 2,05719651 | 2,994823706 | 0,002746035 | 0,039845057 | The Moose: suction sampler vs FISH | Signal transduction mechanisms |
| gene_297491 | 4,009906414 | 6,172921712 | 1,896502007 | 3,254898592 | 0,001134329 | 0,023545921 | The Moose: suction sampler vs FISH | Signal transduction mechanisms |
| gene_1460734 | 6,781489699 | 6,228404202 | 1,862485492 | 3,344135688 | 0,000825394 | 0,019336852 | The Moose: suction sampler vs FISH | Signal transduction mechanisms |
| gene_1361874 | 4,808485401 | 6,229829351 | 1,9420478 | 3,207866125 | 0,001337237 | 0,025948057 | The Moose: suction sampler vs FISH | Signal transduction mechanisms |
| gene_1466405 | 4,152276335 | 6,23940953 | 1,929310409 | 3,234010194 | 0,001220651 | 0,024539704 | The Moose: suction sampler vs FISH | Signal transduction mechanisms |
| gene_1394011 | 26,6387529 | 6,252522167 | 1,223869023 | 5,108816426 | 3,24E-07 | 0,00010352 | The Moose: suction sampler vs FISH | Signal transduction mechanisms |
| gene_580355 | 3,23453046 | 6,292537046 | 2,053352844 | 3,064518143 | 0,002180209 | 0,034893329 | The Moose: suction sampler vs FISH | Signal transduction mechanisms |
| gene_992571 | 3,069079315 | 6,303985121 | 2,050341527 | 3,074602469 | 0,002107833 | 0,034252904 | The Moose: suction sampler vs FISH | Signal transduction mechanisms |
| gene_1557720 | 3,595490753 | 6,36638352 | 2,110639701 | 3,016328897 | 0,002558556 | 0,038346672 | The Moose: suction sampler vs FISH | Signal transduction mechanisms |
| gene_747620 | 5,540182994 | 6,384750892 | 1,741392735 | 3,666462346 | 0,000245929 | 0,008947252 | The Moose: suction sampler vs FISH | Signal transduction mechanisms |
| gene_583817 | 6,158350394 | 6,396990631 | 1,779865822 | 3,594085888 | 0,000325532 | 0,010716598 | The Moose: suction sampler vs FISH | Signal transduction mechanisms |
| gene_707753 | 4,904051082 | 6,459866122 | 1,786422457 | 3,616090973 | 0,000299085 | 0,010192304 | The Moose: suction sampler vs FISH | Signal transduction mechanisms |
| gene_356923 | 6,540446019 | 6,553456343 | 1,873665896 | 3,497665382 | 0,00046935 | 0,013392145 | The Moose: suction sampler vs FISH | Signal transduction mechanisms |
| gene_1477250 | 14,91149545 | 6,67100474 | 1,490189691 | 4,476614475 | 7,58E-06 | 0,000886907 | The Moose: suction sampler vs FISH | Signal transduction mechanisms |
| gene_1668466 | 4,32130672 | 6,672920072 | 1,884682141 | 3,540607684 | 0,000399207 | 0,012196229 | The Moose: suction sampler vs FISH | Signal transduction mechanisms |
| gene_1207606 | 7,129102234 | 6,819431272 | 1,679597157 | 4,060158856 | 4,90E-05 | 0,003123845 | The Moose: suction sampler vs FISH | Signal transduction mechanisms |
| gene_1774127 | 3,818082962 | 6,898698197 | 2,200442071 | 3,135141928 | 0,00171771 | 0,030313307 | The Moose: suction sampler vs FISH | Signal transduction mechanisms |
| gene_1654279 | 6,359466343 | 6,914393297 | 1,840274939 | 3,757261022 | 0,000171783 | 0,007076065 | The Moose: suction sampler vs FISH | Signal transduction mechanisms |
| gene_1017689 | 5,407209301 | 6,943745449 | 1,998561656 | 3,474371395 | 0,000512052 | 0,014175515 | The Moose: suction sampler vs FISH | Signal transduction mechanisms |
| gene_406541 | 6,152304629 | 7,107864498 | 1,95007551 | 3,644917574 | 0,000267478 | 0,0094492 | The Moose: suction sampler vs FISH | Signal transduction mechanisms |
| gene_988178 | 5,121636474 | 7,20856773 | 1,87988645 | 3,834576142 | 0,000125781 | 0,005802755 | The Moose: suction sampler vs FISH | Signal transduction mechanisms |
| gene_1904513 | 4,818282951 | 7,262884618 | 1,860135315 | 3,904492624 | 9,44E-05 | 0,004827305 | The Moose: suction sampler vs FISH | Signal transduction mechanisms |
| gene_1119313 | 8,158564891 | 7,368369803 | 1,602181552 | 4,598960581 | 4,25E-06 | 0,000587289 | The Moose: suction sampler vs FISH | Signal transduction mechanisms |
| gene_752780 | 7,690047865 | 7,383839669 | 1,724608812 | 4,281457696 | 1,86E-05 | 0,001574414 | The Moose: suction sampler vs FISH | Signal transduction mechanisms |
| gene_115977 | 9,085583674 | 7,435812909 | 1,660764093 | 4,477344457 | 7,56E-06 | 0,000886907 | The Moose: suction sampler vs FISH | Signal transduction mechanisms |
| gene_290262 | 6,884149136 | 7,841268375 | 2,067718682 | 3,79223172 | 0,0001493 | 0,006446386 | The Moose: suction sampler vs FISH | Signal transduction mechanisms |
| gene_1399174 | 65,28048314 | 9,685203379 | 1,533537343 | 6,315596698 | 2,69E-10 | 7,06E-07 | The Moose: suction sampler vs FISH | Signal transduction mechanisms |
| gene_1456066 | 6,003329406 | -6,045443029 | 1,707006262 | -3,541547071 | 0,000397788 | 0,012173138 | The Moose: suction sampler vs FISH | Transcription |
| gene_482999 | 5,040230707 | -5,70145186 | 1,83653319 | -3,104464374 | 0,001906239 | 0,032244061 | The Moose: suction sampler vs FISH | Transcription |
| gene_1029702 | 4,201857817 | -5,421018558 | 1,769481549 | -3,063619716 | 0,002186767 | 0,034937371 | The Moose: suction sampler vs FISH | Transcription |
| gene_51865 | 5,994984038 | -5,228121073 | 1,739986322 | -3,004690904 | 0,002658509 | 0,039113415 | The Moose: suction sampler vs FISH | Transcription |
| gene_89569 | 15,17541346 | -4,149731074 | 1,160185338 | -3,576782896 | 0,000347849 | 0,011150745 | The Moose: suction sampler vs FISH | Transcription |
| gene_806635 | 16,11989615 | -3,428115403 | 0,945901535 | -3,624177864 | 0,000289882 | 0,009958086 | The Moose: suction sampler vs FISH | Transcription |
| gene_345670 | 14,6517261 | -3,395461252 | 1,064444988 | -3,189888901 | 0,001423275 | 0,026988521 | The Moose: suction sampler vs FISH | Transcription |
| gene_1018642 | 14,31751465 | -2,910332575 | 1,003935244 | -2,898924598 | 0,003744449 | 0,048020094 | The Moose: suction sampler vs FISH | Transcription |
| gene_1166780 | 164,054286 | -2,811891371 | 0,573726505 | -4,901100694 | 9,53E-07 | 0,0002167 | The Moose: suction sampler vs FISH | Transcription |
| gene_1099998 | 58,30659227 | -2,761585006 | 0,728805504 | -3,789193401 | 0,000151137 | 0,006491388 | The Moose: suction sampler vs FISH | Transcription |
| gene_1133908 | 319,4878799 | -2,267052169 | 0,537191444 | -4,220194116 | 2,44E-05 | 0,0019256 | The Moose: suction sampler vs FISH | Transcription |
| gene_1557575 | 95,72982922 | -2,241530468 | 0,604663205 | -3,707072716 | 0,000209669 | 0,008049724 | The Moose: suction sampler vs FISH | Transcription |
| gene_313702 | 73,85668384 | -2,156853306 | 0,708702486 | -3,043383292 | 0,002339342 | 0,036216894 | The Moose: suction sampler vs FISH | Transcription |
| gene_1109862 | 107,6358592 | 2,128897893 | 0,696074184 | 3,058435355 | 0,002224961 | 0,035225748 | The Moose: suction sampler vs FISH | Transcription |
| gene_389440 | 90,77766035 | 2,162445132 | 0,705074943 | 3,066972033 | 0,002162391 | 0,034744432 | The Moose: suction sampler vs FISH | Transcription |
| gene_1342732 | 162,7855056 | 2,217917225 | 0,66500643 | 3,335181624 | 0,000852437 | 0,01978737 | The Moose: suction sampler vs FISH | Transcription |
| gene_1116872 | 28,69125631 | 2,369461666 | 0,719290039 | 3,294167216 | 0,000987138 | 0,021545414 | The Moose: suction sampler vs FISH | Transcription |
| gene_559591 | 45,71892863 | 2,401720155 | 0,749936974 | 3,202562667 | 0,001362107 | 0,026236538 | The Moose: suction sampler vs FISH | Transcription |
| gene_400530 | 136,9228151 | 2,431089716 | 0,660432351 | 3,681057889 | 0,000232268 | 0,008619981 | The Moose: suction sampler vs FISH | Transcription |
| gene_802955 | 101,588309 | 2,443512033 | 0,639493074 | 3,82101407 | 0,000132904 | 0,006003161 | The Moose: suction sampler vs FISH | Transcription |
| gene_330663 | 662,4282625 | 2,446352044 | 0,762184524 | 3,20965851 | 0,001328928 | 0,025868826 | The Moose: suction sampler vs FISH | Transcription |
| gene_1587070 | 26,55463685 | 2,540944108 | 0,851566909 | 2,983845522 | 0,002846504 | 0,040863967 | The Moose: suction sampler vs FISH | Transcription |
| gene_1605393 | 20,72385623 | 2,55347881 | 0,850162076 | 3,003520015 | 0,00266876 | 0,03922495 | The Moose: suction sampler vs FISH | Transcription |
| gene_1599443 | 98,20984361 | 2,55751535 | 0,726849022 | 3,518633542 | 0,000433775 | 0,012799774 | The Moose: suction sampler vs FISH | Transcription |
| gene_1475087 | 35,33295897 | 2,603489034 | 0,795069258 | 3,274543703 | 0,001058328 | 0,022589037 | The Moose: suction sampler vs FISH | Transcription |
| gene_1894170 | 52,5375207 | 2,650496689 | 0,739239293 | 3,58543805 | 0,000336513 | 0,010927321 | The Moose: suction sampler vs FISH | Transcription |
| gene_46338 | 530,2880947 | 2,661673866 | 0,774038973 | 3,438681977 | 0,000584554 | 0,015399043 | The Moose: suction sampler vs FISH | Transcription |
| gene_1113639 | 20,95607495 | 2,672557866 | 0,887778029 | 3,010389735 | 0,002609127 | 0,038790953 | The Moose: suction sampler vs FISH | Transcription |
| gene_1560282 | 21,70709302 | 2,752371418 | 0,870746538 | 3,160932946 | 0,001572647 | 0,028696412 | The Moose: suction sampler vs FISH | Transcription |
| gene_538131 | 429,9934751 | 2,79096421 | 0,761743347 | 3,663916753 | 0,000248388 | 0,008995475 | The Moose: suction sampler vs FISH | Transcription |
| gene_321298 | 42,00967567 | 2,829523551 | 0,882519729 | 3,206187303 | 0,001345064 | 0,02605862 | The Moose: suction sampler vs FISH | Transcription |
| gene_1548198 | 17,44679102 | 2,830355139 | 0,929131364 | 3,046237862 | 0,002317244 | 0,036034303 | The Moose: suction sampler vs FISH | Transcription |
| gene_184962 | 29,80299502 | 2,844175454 | 0,901391684 | 3,155315833 | 0,001603244 | 0,029057674 | The Moose: suction sampler vs FISH | Transcription |
| gene_1309490 | 95,42533722 | 2,847880923 | 0,871237428 | 3,268777064 | 0,001080134 | 0,022861792 | The Moose: suction sampler vs FISH | Transcription |
| gene_1927845 | 25,32324468 | 2,859630327 | 0,933197697 | 3,064334958 | 0,002181545 | 0,034899494 | The Moose: suction sampler vs FISH | Transcription |
| gene_1593719 | 29,76672723 | 2,868670955 | 0,834519935 | 3,437510399 | 0,000587088 | 0,015449191 | The Moose: suction sampler vs FISH | Transcription |
| gene_250179 | 29,5467203 | 2,935592156 | 0,805230132 | 3,645656118 | 0,00026671 | 0,009438241 | The Moose: suction sampler vs FISH | Transcription |

| | | | | | | | | |
|---|---|---|---|---|---|---|---|---|
| gene_519024 | 108,2273966 | 2,949215623 | 0,863622456 | 3,414936239 | 0,000637969 | 0,016320264 | The Moose: suction sampler vs FISH | Transcription |
| gene_750738 | 31,99323668 | 2,949335502 | 0,816414878 | 3,612545021 | 0,000303206 | 0,010271678 | The Moose: suction sampler vs FISH | Transcription |
| gene_1233498 | 30,6544729 | 2,961399422 | 0,958650858 | 3,089132397 | 0,00200742 | 0,033236008 | The Moose: suction sampler vs FISH | Transcription |
| gene_1605394 | 26,80401992 | 2,967295325 | 0,878683612 | 3,37697811 | 0,000732869 | 0,017888186 | The Moose: suction sampler vs FISH | Transcription |
| gene_1762910 | 50,68668778 | 3,019482546 | 0,708975934 | 4,258935177 | 2,05E-05 | 0,001696795 | The Moose: suction sampler vs FISH | Transcription |
| gene_860531 | 27,3003771 | 3,034282096 | 0,880102277 | 3,447647136 | 0,000565492 | 0,015113749 | The Moose: suction sampler vs FISH | Transcription |
| gene_65082 | 15,66771144 | 3,081903103 | 0,997717365 | 3,088954056 | 0,002008625 | 0,03324847 | The Moose: suction sampler vs FISH | Transcription |
| gene_1201874 | 14,20741427 | 3,11366309 | 1,012779571 | 3,074373909 | 0,002109449 | 0,034252904 | The Moose: suction sampler vs FISH | Transcription |
| gene_615591 | 35,51645449 | 3,119737741 | 0,778951519 | 4,00504738 | 6,20E-05 | 0,003625765 | The Moose: suction sampler vs FISH | Transcription |
| gene_965969 | 16,90376187 | 3,119810178 | 0,865596994 | 3,60422945 | 0,00031308 | 0,010475712 | The Moose: suction sampler vs FISH | Transcription |
| gene_436076 | 46,42558978 | 3,124370322 | 0,731332201 | 4,272162934 | 1,94E-05 | 0,001617512 | The Moose: suction sampler vs FISH | Transcription |
| gene_511192 | 31,36764382 | 3,131285421 | 0,805261969 | 3,888530118 | 0,000100853 | 0,00502521 | The Moose: suction sampler vs FISH | Transcription |
| gene_21989 | 16,95259308 | 3,138402978 | 0,913995153 | 3,433719499 | 0,00059536 | 0,015560929 | The Moose: suction sampler vs FISH | Transcription |
| gene_1213467 | 16,40173578 | 3,146652614 | 1,052640625 | 2,989294293 | 0,002796227 | 0,04037925 | The Moose: suction sampler vs FISH | Transcription |
| gene_225127 | 14,51597745 | 3,157656851 | 1,026518936 | 3,076082417 | 0,002097399 | 0,034171127 | The Moose: suction sampler vs FISH | Transcription |
| gene_1546406 | 24,19721866 | 3,166265629 | 1,09323704 | 2,896229741 | 0,003776758 | 0,048220526 | The Moose: suction sampler vs FISH | Transcription |
| gene_434517 | 93,45655662 | 3,208647291 | 0,714206953 | 4,492601586 | 7,04E-06 | 0,000845739 | The Moose: suction sampler vs FISH | Transcription |
| gene_1287918 | 72,93491503 | 3,226348159 | 0,910571826 | 3,543211053 | 0,000395286 | 0,012111721 | The Moose: suction sampler vs FISH | Transcription |
| gene_863320 | 11,14503655 | 3,228565461 | 1,079495084 | 2,990810714 | 0,002782379 | 0,040250508 | The Moose: suction sampler vs FISH | Transcription |
| gene_593722 | 21,8459439 | 3,25641439 | 1,048555055 | 3,105620801 | 0,0018988 | 0,032177526 | The Moose: suction sampler vs FISH | Transcription |
| gene_563538 | 28,24879004 | 3,269060191 | 0,943063588 | 3,466426054 | 0,000527427 | 0,014448376 | The Moose: suction sampler vs FISH | Transcription |
| gene_972198 | 128,1970345 | 3,274227555 | 0,940017174 | 3,483157165 | 0,000495537 | 0,013883037 | The Moose: suction sampler vs FISH | Transcription |
| gene_1197229 | 36,89770643 | 3,296734638 | 1,042584966 | 3,162077667 | 0,001566478 | 0,028640767 | The Moose: suction sampler vs FISH | Transcription |
| gene_1270090 | 25,39257283 | 3,307545592 | 0,886780004 | 3,729837816 | 0,000191603 | 0,007614874 | The Moose: suction sampler vs FISH | Transcription |
| gene_1394790 | 29,28523821 | 3,310984795 | 0,890412439 | 3,718484436 | 0,000200422 | 0,007833937 | The Moose: suction sampler vs FISH | Transcription |
| gene_995522 | 34,05673735 | 3,315479916 | 0,836033482 | 3,965726241 | 7,32E-05 | 0,004083721 | The Moose: suction sampler vs FISH | Transcription |
| gene_1851537 | 82,50163273 | 3,317139023 | 0,811161629 | 4,089368759 | 4,33E-05 | 0,002862031 | The Moose: suction sampler vs FISH | Transcription |
| gene_1256662 | 36,40531935 | 3,342893934 | 0,912414038 | 3,663790554 | 0,00024851 | 0,008995475 | The Moose: suction sampler vs FISH | Transcription |
| gene_406016 | 19,24478458 | 3,360541009 | 1,085028342 | 3,097191914 | 0,001953634 | 0,032684422 | The Moose: suction sampler vs FISH | Transcription |
| gene_689976 | 11,93528202 | 3,364338849 | 1,019669428 | 3,299440738 | 0,000968777 | 0,021309318 | The Moose: suction sampler vs FISH | Transcription |
| gene_888916 | 13,22753408 | 3,41474654 | 1,091243801 | 3,129224228 | 0,001752685 | 0,030666956 | The Moose: suction sampler vs FISH | Transcription |
| gene_866012 | 18,91611894 | 3,433761035 | 1,043278062 | 3,291319122 | 0,000997187 | 0,021700271 | The Moose: suction sampler vs FISH | Transcription |
| gene_1775316 | 10,99561717 | 3,504958148 | 1,185668969 | 2,95610178 | 0,003115543 | 0,043133087 | The Moose: suction sampler vs FISH | Transcription |
| gene_1905533 | 13,99845586 | 3,512897748 | 1,197864958 | 2,932632534 | 0,003361014 | 0,044979901 | The Moose: suction sampler vs FISH | Transcription |
| gene_343942 | 43,59113504 | 3,53961275 | 1,115923479 | 3,171913503 | 0,001514381 | 0,028058455 | The Moose: suction sampler vs FISH | Transcription |
| gene_1619509 | 54,33402695 | 3,545326658 | 0,814584346 | 4,352313761 | 1,35E-05 | 0,001295729 | The Moose: suction sampler vs FISH | Transcription |
| gene_525687 | 28,45825326 | 3,561924821 | 0,961611135 | 3,704121857 | 0,000212124 | 0,008118534 | The Moose: suction sampler vs FISH | Transcription |
| gene_891583 | 77,01236753 | 3,570083516 | 0,945567565 | 3,775598539 | 0,000159624 | 0,006764903 | The Moose: suction sampler vs FISH | Transcription |
| gene_1611963 | 10,94489143 | 3,589194113 | 1,205039893 | 2,978485719 | 0,002896765 | 0,041247175 | The Moose: suction sampler vs FISH | Transcription |
| gene_1576926 | 43,18005625 | 3,595965537 | 0,743720714 | 4,835102032 | 1,33E-06 | 0,000271496 | The Moose: suction sampler vs FISH | Transcription |
| gene_724295 | 51,89076972 | 3,624888414 | 0,809898819 | 4,475729966 | 7,62E-06 | 0,00088776 | The Moose: suction sampler vs FISH | Transcription |
| gene_1853238 | 16,40251677 | 3,642692897 | 1,09822769 | 3,316883128 | 0,000910277 | 0,020591283 | The Moose: suction sampler vs FISH | Transcription |
| gene_250787 | 22,70365085 | 3,690429181 | 0,973638186 | 3,790349674 | 0,000150435 | 0,006479324 | The Moose: suction sampler vs FISH | Transcription |
| gene_930082 | 13,82877346 | 3,712930076 | 1,123372913 | 3,305162544 | 0,000949213 | 0,02106829 | The Moose: suction sampler vs FISH | Transcription |
| gene_1812269 | 26,82931376 | 3,718149181 | 1,174346849 | 3,166142255 | 0,001544752 | 0,028381761 | The Moose: suction sampler vs FISH | Transcription |
| gene_436077 | 16,37318419 | 3,77307429 | 0,962044831 | 3,921931876 | 8,78E-05 | 0,004582062 | The Moose: suction sampler vs FISH | Transcription |
| gene_1805432 | 21,89653429 | 3,793923205 | 1,035327249 | 3,664467642 | 0,000247854 | 0,008989435 | The Moose: suction sampler vs FISH | Transcription |
| gene_884297 | 8,038474183 | 3,794342365 | 1,275277103 | 2,975308157 | 0,002926944 | 0,041503598 | The Moose: suction sampler vs FISH | Transcription |
| gene_40544 | 20,77152647 | 3,796873216 | 0,979193967 | 3,877549643 | 0,000105514 | 0,005169754 | The Moose: suction sampler vs FISH | Transcription |
| gene_1467186 | 32,55623201 | 3,800040963 | 0,950002664 | 4,000031904 | 6,33E-05 | 0,003677124 | The Moose: suction sampler vs FISH | Transcription |
| gene_917361 | 19,46300832 | 3,831058505 | 1,220717029 | 3,138367381 | 0,001698918 | 0,030103022 | The Moose: suction sampler vs FISH | Transcription |
| gene_1134566 | 87,68541541 | 3,876004632 | 0,802885692 | 4,827592109 | 1,38E-06 | 0,000275805 | The Moose: suction sampler vs FISH | Transcription |
| gene_1088722 | 12,22126618 | 3,903989687 | 1,283561308 | 3,041529581 | 0,002353794 | 0,036361491 | The Moose: suction sampler vs FISH | Transcription |
| gene_952406 | 87,15904345 | 3,918170173 | 0,817455841 | 4,793127625 | 1,64E-06 | 0,000308493 | The Moose: suction sampler vs FISH | Transcription |
| gene_1748510 | 15,44355795 | 3,919265561 | 1,062723555 | 3,687944567 | 0,000226073 | 0,008467069 | The Moose: suction sampler vs FISH | Transcription |
| gene_1605261 | 23,77439521 | 3,944523367 | 0,952288445 | 4,142151873 | 3,44E-05 | 0,002432114 | The Moose: suction sampler vs FISH | Transcription |
| gene_346808 | 7,83947692 | 3,950147803 | 1,323926994 | 2,983659841 | 0,002848232 | 0,040865094 | The Moose: suction sampler vs FISH | Transcription |
| gene_896839 | 27,67743096 | 3,960230051 | 1,001926686 | 3,952614605 | 7,73E-05 | 0,00422194 | The Moose: suction sampler vs FISH | Transcription |
| gene_309815 | 11,04858606 | 4,00691453 | 1,132690448 | 3,537519483 | 0,000403904 | 0,012258164 | The Moose: suction sampler vs FISH | Transcription |
| gene_1169204 | 12,80449626 | 4,027153295 | 1,12455782 | 3,581099366 | 0,000342151 | 0,011041 | The Moose: suction sampler vs FISH | Transcription |
| gene_157466 | 8,099726795 | 4,048196088 | 1,315383721 | 3,077578066 | 0,002086902 | 0,034075699 | The Moose: suction sampler vs FISH | Transcription |
| gene_1582552 | 103,7410405 | 4,048259091 | 0,902186246 | 4,487165604 | 7,22E-06 | 0,000862102 | The Moose: suction sampler vs FISH | Transcription |
| gene_1030249 | 13,2328546 | 4,054747843 | 1,15119292 | 3,522214021 | 0,000427958 | 0,012694431 | The Moose: suction sampler vs FISH | Transcription |
| gene_530055 | 10,0287761 | 4,069524637 | 1,155938653 | 3,520536861 | 0,000430674 | 0,012738969 | The Moose: suction sampler vs FISH | Transcription |
| gene_450760 | 17,24717949 | 4,187505563 | 1,168870095 | 3,582524336 | 0,00034029 | 0,011015763 | The Moose: suction sampler vs FISH | Transcription |

| | | | | | | | | |
|---|---|---|---|---|---|---|---|---|
| gene_1859673 | 30,13277018 | 4,201615357 | 0,928863505 | 4,523393734 | 6,09E-06 | 0,000761424 | The Moose: suction sampler vs FISH | Transcription |
| gene_680711 | 7,486314122 | 4,273702652 | 1,440685251 | 2,96643743 | 0,003012717 | 0,042287372 | The Moose: suction sampler vs FISH | Transcription |
| gene_42268 | 9,550444893 | 4,35447077 | 1,513414614 | 2,877249056 | 0,004011588 | 0,049886744 | The Moose: suction sampler vs FISH | Transcription |
| gene_384581 | 81,69799844 | 4,397413596 | 0,92827446 | 4,737191191 | 2,17E-06 | 0,000366549 | The Moose: suction sampler vs FISH | Transcription |
| gene_930233 | 12,02897947 | 4,411306148 | 1,27835796 | 3,450759714 | 0,000559011 | 0,014977737 | The Moose: suction sampler vs FISH | Transcription |
| gene_564264 | 10,42297077 | 4,471971984 | 1,463957521 | 3,054714306 | 0,00225275 | 0,035451731 | The Moose: suction sampler vs FISH | Transcription |
| gene_258129 | 23,74128303 | 4,536117795 | 1,033483594 | 4,389153174 | 1,14E-05 | 0,001160767 | The Moose: suction sampler vs FISH | Transcription |
| gene_104773 | 10,64614511 | 4,731502095 | 1,261901759 | 3,749501148 | 0,000177187 | 0,00720569 | The Moose: suction sampler vs FISH | Transcription |
| gene_651171 | 10,27730385 | 4,934426068 | 1,436546253 | 3,434923211 | 0,000592722 | 0,01551959 | The Moose: suction sampler vs FISH | Transcription |
| gene_1226324 | 19,83597005 | 5,099164729 | 1,139056934 | 4,476654835 | 7,58E-06 | 0,000886907 | The Moose: suction sampler vs FISH | Transcription |
| gene_190641 | 4,206481143 | 5,099911122 | 1,746308549 | 2,92039521 | 0,003495877 | 0,046046689 | The Moose: suction sampler vs FISH | Transcription |
| gene_91178 | 16,66173527 | 5,145405644 | 1,27205633 | 4,044951094 | 5,23E-05 | 0,003238142 | The Moose: suction sampler vs FISH | Transcription |
| gene_1652414 | 6,184366259 | 5,147966699 | 1,604022082 | 3,209411364 | 0,00133007 | 0,025870505 | The Moose: suction sampler vs FISH | Transcription |
| gene_1085915 | 4,403215545 | 5,194767969 | 1,801114581 | 2,884196277 | 0,003924141 | 0,049233224 | The Moose: suction sampler vs FISH | Transcription |
| gene_1212574 | 16,46527293 | 5,210935632 | 1,287272463 | 4,048044049 | 5,16E-05 | 0,003223638 | The Moose: suction sampler vs FISH | Transcription |
| gene_1030463 | 27,76603647 | 5,245318018 | 1,200606091 | 4,368891725 | 1,25E-05 | 0,001226165 | The Moose: suction sampler vs FISH | Transcription |
| gene_1797691 | 9,424853656 | 5,30048378 | 1,625361413 | 3,261110875 | 0,001109766 | 0,023254434 | The Moose: suction sampler vs FISH | Transcription |
| gene_211941 | 6,643356226 | 5,30568725 | 1,612551587 | 3,290243421 | 0,001001007 | 0,021744748 | The Moose: suction sampler vs FISH | Transcription |
| gene_1014229 | 6,908338415 | 5,389136304 | 1,732979404 | 3,109752079 | 0,001872444 | 0,031937221 | The Moose: suction sampler vs FISH | Transcription |
| gene_931510 | 4,213028827 | 5,436951987 | 1,794525844 | 3,029742929 | 0,00244762 | 0,037241647 | The Moose: suction sampler vs FISH | Transcription |
| gene_658155 | 5,147304403 | 5,465087074 | 1,663759606 | 3,28478168 | 0,001020614 | 0,022001471 | The Moose: suction sampler vs FISH | Transcription |
| gene_1587068 | 4,473566311 | 5,684347652 | 1,759417404 | 3,230812449 | 0,001234389 | 0,024693184 | The Moose: suction sampler vs FISH | Transcription |
| gene_516382 | 3,715968962 | 5,757960649 | 1,988837774 | 2,895138419 | 0,003789914 | 0,048307916 | The Moose: suction sampler vs FISH | Transcription |
| gene_1076062 | 163,3351647 | 5,795313666 | 0,868226909 | 6,674883724 | 2,47E-11 | 1,40E-07 | The Moose: suction sampler vs FISH | Transcription |
| gene_448239 | 6,156756893 | 5,835986453 | 1,812968176 | 3,219023108 | 0,001286281 | 0,025402238 | The Moose: suction sampler vs FISH | Transcription |
| gene_1899690 | 33,56828511 | 5,85300972 | 1,222697032 | 4,786966489 | 1,69E-06 | 0,000311674 | The Moose: suction sampler vs FISH | Transcription |
| gene_1216339 | 3,894593785 | 5,926152092 | 1,995279447 | 2,970086271 | 0,002977161 | 0,041936634 | The Moose: suction sampler vs FISH | Transcription |
| gene_1067032 | 3,952735442 | 6,002167435 | 2,007167071 | 2,99036763 | 0,002786419 | 0,040285142 | The Moose: suction sampler vs FISH | Transcription |
| gene_564265 | 21,38311639 | 6,008143368 | 1,317920771 | 4,558804673 | 5,14E-06 | 0,000669933 | The Moose: suction sampler vs FISH | Transcription |
| gene_1939284 | 4,210070574 | 6,018291666 | 1,980525783 | 3,038734319 | 0,002375743 | 0,036549313 | The Moose: suction sampler vs FISH | Transcription |
| gene_1592081 | 4,015157217 | 6,048082248 | 1,880400084 | 3,216380545 | 0,001298186 | 0,025541185 | The Moose: suction sampler vs FISH | Transcription |
| gene_1827154 | 5,60665518 | 6,063202603 | 1,761706904 | 3,441663644 | 0,000578149 | 0,015301668 | The Moose: suction sampler vs FISH | Transcription |
| gene_221938 | 13,61428996 | 6,133332988 | 1,519502066 | 4,036409772 | 5,43E-05 | 0,003308096 | The Moose: suction sampler vs FISH | Transcription |
| gene_475917 | 4,277770363 | 6,214688495 | 1,971277114 | 3,152620426 | 0,001618121 | 0,029225655 | The Moose: suction sampler vs FISH | Transcription |
| gene_161517 | 5,375809765 | 6,226594287 | 2,095905365 | 2,970837515 | 0,002969889 | 0,041850244 | The Moose: suction sampler vs FISH | Transcription |
| gene_252400 | 8,05576825 | 6,227832163 | 1,596988139 | 3,899736017 | 9,63E-05 | 0,004881007 | The Moose: suction sampler vs FISH | Transcription |
| gene_385515 | 6,554748334 | 6,280411907 | 1,671260576 | 3,757889104 | 0,000171353 | 0,007070228 | The Moose: suction sampler vs FISH | Transcription |
| gene_1629497 | 5,819133952 | 6,324672725 | 1,850500503 | 3,41781735 | 0,000631255 | 0,01618237 | The Moose: suction sampler vs FISH | Transcription |
| gene_1074660 | 3,616189432 | 6,37775885 | 1,834806135 | 3,475985135 | 0,00050898 | 0,014111768 | The Moose: suction sampler vs FISH | Transcription |
| gene_686920 | 3,555229226 | 6,450099853 | 2,118832211 | 3,044176796 | 0,00233318 | 0,036179697 | The Moose: suction sampler vs FISH | Transcription |
| gene_1024890 | 5,561821121 | 6,594430213 | 1,99538297 | 3,304844389 | 0,000950291 | 0,021079479 | The Moose: suction sampler vs FISH | Transcription |
| gene_1226329 | 4,54025499 | 6,983586714 | 2,086432887 | 3,347141793 | 0,000816494 | 0,019195719 | The Moose: suction sampler vs FISH | Transcription |
| gene_1072447 | 5,07858743 | 7,08012563 | 2,252793009 | 3,142821201 | 0,00167328 | 0,029835897 | The Moose: suction sampler vs FISH | Transcription |
| gene_1641279 | 7,115378492 | 7,099852802 | 1,71396326 | 4,142359972 | 3,44E-05 | 0,002432114 | The Moose: suction sampler vs FISH | Transcription |
| gene_475918 | 4,172434447 | 7,122409504 | 1,956124439 | 3,641082009 | 0,000271495 | 0,009538157 | The Moose: suction sampler vs FISH | Transcription |
| gene_914588 | 6,947239654 | 7,263851585 | 1,69310028 | 4,290266603 | 1,78E-05 | 0,001540177 | The Moose: suction sampler vs FISH | Transcription |
| gene_1737426 | 5,313333003 | 7,313469038 | 1,871563387 | 3,907679051 | 9,32E-05 | 0,004789446 | The Moose: suction sampler vs FISH | Transcription |
| gene_1030462 | 8,696863953 | 8,447612817 | 1,783977439 | 4,735268861 | 2,19E-06 | 0,00036767 | The Moose: suction sampler vs FISH | Transcription |
| gene_800375 | 14,58040614 | 8,799381218 | 1,663863249 | 5,288524295 | 1,23E-07 | 5,30E-05 | The Moose: suction sampler vs FISH | Transcription |
| gene_1134952 | 241,3022917 | 13,28424339 | 4,247170278 | 3,12778686 | 0,001761279 | 0,030726153 | The Moose: suction sampler vs FISH | Transcription |
| gene_969842 | 14,54026623 | 19,98127775 | 3,325979769 | 6,007636589 | 1,88E-09 | 2,82E-06 | The Moose: suction sampler vs FISH | Transcription |
| gene_1614277 | 10,6995357 | -6,779700839 | 1,650111113 | -4,10863292 | 3,98E-05 | 0,002696652 | The Moose: suction sampler vs FISH | Translation, ribosomal structure and biogenesis |
| gene_72261 | 31,30475423 | -5,57965183 | 1,051219343 | -5,307790298 | 1,11E-07 | 4,94E-05 | The Moose: suction sampler vs FISH | Translation, ribosomal structure and biogenesis |
| gene_1247826 | 5,056673253 | -5,362567735 | 1,649140427 | -3,251735054 | 0,001147029 | 0,023706063 | The Moose: suction sampler vs FISH | Translation, ribosomal structure and biogenesis |
| gene_806636 | 4,256741978 | -5,35088033 | 1,839214775 | -2,909328699 | 0,003622058 | 0,047041921 | The Moose: suction sampler vs FISH | Translation, ribosomal structure and biogenesis |
| gene_696757 | 30,66352937 | -4,791909486 | 0,966878146 | -4,956063498 | 7,19E-07 | 0,00017789 | The Moose: suction sampler vs FISH | Translation, ribosomal structure and biogenesis |
| gene_1132833 | 15,16207669 | -4,724976376 | 1,26949172 | -3,721943437 | 0,000197695 | 0,00774799 | The Moose: suction sampler vs FISH | Translation, ribosomal structure and biogenesis |
| gene_1501762 | 27,48070898 | -4,310710843 | 1,09880753 | -3,923080908 | 8,74E-05 | 0,004572734 | The Moose: suction sampler vs FISH | Translation, ribosomal structure and biogenesis |
| gene_819731 | 10,44854217 | -4,205495499 | 1,410084883 | -2,982441376 | 0,002859594 | 0,04096409 | The Moose: suction sampler vs FISH | Translation, ribosomal structure and biogenesis |
| gene_696758 | 40,21595376 | -3,951863874 | 0,790838043 | -4,997058384 | 5,82E-07 | 0,000151608 | The Moose: suction sampler vs FISH | Translation, ribosomal structure and biogenesis |
| gene_219120 | 34,95421501 | -3,764766044 | 1,035609535 | -3,635314194 | 0,000277642 | 0,009687135 | The Moose: suction sampler vs FISH | Translation, ribosomal structure and biogenesis |
| gene_730492 | 62,52440462 | -3,71526097 | 0,664210297 | -5,59350101 | 2,23E-08 | 1,61E-05 | The Moose: suction sampler vs FISH | Translation, ribosomal structure and biogenesis |
| gene_696756 | 21,45445677 | -3,537266916 | 0,961767921 | -3,677879911 | 0,000235181 | 0,008692925 | The Moose: suction sampler vs FISH | Translation, ribosomal structure and biogenesis |
| gene_1065120 | 38,44951958 | -3,511050512 | 0,840586866 | -4,176903843 | 2,96E-05 | 0,002187739 | The Moose: suction sampler vs FISH | Translation, ribosomal structure and biogenesis |
| gene_767840 | 119,2634189 | -3,3058639 | 0,564346867 | -5,857858159 | 4,69E-09 | 5,27E-06 | The Moose: suction sampler vs FISH | Translation, ribosomal structure and biogenesis |

| | | | | | | | | |
|---|---|---|---|---|---|---|---|---|
| gene_1710086 | 10,32202582 | -3,279868108 | 1,140151561 | -2,876694835 | 0,00401864 | 0,04992372 | The Moose: suction sampler vs FISH | Translation, ribosomal structure and biogenesis |
| gene_547342 | 14,63668135 | -3,181836345 | 1,082423191 | -2,939549309 | 0,0032869 | 0,044433342 | The Moose: suction sampler vs FISH | Translation, ribosomal structure and biogenesis |
| gene_767843 | 44,30681567 | -3,178986386 | 0,900828619 | -3,528958028 | 0,000417199 | 0,01250151 | The Moose: suction sampler vs FISH | Translation, ribosomal structure and biogenesis |
| gene_767842 | 76,96586428 | -3,15192594 | 0,678754381 | -4,643691488 | 3,42E-06 | 0,000500711 | The Moose: suction sampler vs FISH | Translation, ribosomal structure and biogenesis |
| gene_923007 | 41,17435313 | -3,114830009 | 0,746133677 | -4,174627288 | 2,98E-05 | 0,002200948 | The Moose: suction sampler vs FISH | Translation, ribosomal structure and biogenesis |
| gene_1842570 | 13,25980374 | -3,070833317 | 1,063624308 | -2,887140971 | 0,0038876 | 0,048991899 | The Moose: suction sampler vs FISH | Translation, ribosomal structure and biogenesis |
| gene_923009 | 30,45495878 | -3,063985009 | 0,774097354 | -3,958139104 | 7,55E-05 | 0,004163874 | The Moose: suction sampler vs FISH | Translation, ribosomal structure and biogenesis |
| gene_377145 | 1629,361261 | -3,062990681 | 0,59543749 | -5,144101158 | 2,69E-07 | 9,01E-05 | The Moose: suction sampler vs FISH | Translation, ribosomal structure and biogenesis |
| gene_1385216 | 16,52289051 | -3,05365113 | 1,00496352 | -3,038569132 | 0,002377046 | 0,036561699 | The Moose: suction sampler vs FISH | Translation, ribosomal structure and biogenesis |
| gene_923006 | 13,90314965 | -3,020402281 | 0,925224858 | -3,264506195 | 0,001096551 | 0,023091595 | The Moose: suction sampler vs FISH | Translation, ribosomal structure and biogenesis |
| gene_1577082 | 34,63024568 | -3,015630805 | 0,777228108 | -3,879981661 | 0,000104464 | 0,005128599 | The Moose: suction sampler vs FISH | Translation, ribosomal structure and biogenesis |
| gene_1842571 | 13,59453275 | -2,99260445 | 1,028174808 | -2,910598885 | 0,003607368 | 0,046911539 | The Moose: suction sampler vs FISH | Translation, ribosomal structure and biogenesis |
| gene_1445905 | 19,58418802 | -2,973563346 | 1,007672372 | -2,950922768 | 0,003168261 | 0,043488543 | The Moose: suction sampler vs FISH | Translation, ribosomal structure and biogenesis |
| gene_124686 | 90,584743 | -2,943860863 | 0,678493574 | -4,338819077 | 1,43E-05 | 0,001337845 | The Moose: suction sampler vs FISH | Translation, ribosomal structure and biogenesis |
| gene_600135 | 28,63756011 | -2,934202561 | 0,971198082 | -3,021219478 | 0,002517588 | 0,037936859 | The Moose: suction sampler vs FISH | Translation, ribosomal structure and biogenesis |
| gene_1020191 | 111,8786677 | -2,863906202 | 0,656634093 | -4,361494828 | 1,29E-05 | 0,001256859 | The Moose: suction sampler vs FISH | Translation, ribosomal structure and biogenesis |
| gene_950182 | 42,33413869 | -2,860073043 | 0,754727926 | -3,789541824 | 0,000150925 | 0,006486083 | The Moose: suction sampler vs FISH | Translation, ribosomal structure and biogenesis |
| gene_950185 | 21,21143844 | -2,822437562 | 0,910452015 | -3,100039887 | 0,001934946 | 0,032541813 | The Moose: suction sampler vs FISH | Translation, ribosomal structure and biogenesis |
| gene_754878 | 76,72769874 | -2,741176148 | 0,833543826 | -3,28858071 | 0,001006939 | 0,021809095 | The Moose: suction sampler vs FISH | Translation, ribosomal structure and biogenesis |
| gene_696759 | 842,2692395 | -2,716527259 | 0,518824118 | -5,235930955 | 1,64E-07 | 6,31E-05 | The Moose: suction sampler vs FISH | Translation, ribosomal structure and biogenesis |
| gene_456395 | 60,18618924 | -2,618207207 | 0,723317415 | -3,619720956 | 0,000294921 | 0,010084015 | The Moose: suction sampler vs FISH | Translation, ribosomal structure and biogenesis |
| gene_1133909 | 150,8788831 | -2,618003937 | 0,617994803 | -4,236287948 | 2,27E-05 | 0,001822056 | The Moose: suction sampler vs FISH | Translation, ribosomal structure and biogenesis |
| gene_1371330 | 62,9108037 | -2,609694466 | 0,831192479 | -3,139699325 | 0,001691213 | 0,03003171 | The Moose: suction sampler vs FISH | Translation, ribosomal structure and biogenesis |
| gene_923008 | 38,7465989 | -2,595079119 | 0,809250712 | -3,206767793 | 0,001342353 | 0,026026694 | The Moose: suction sampler vs FISH | Translation, ribosomal structure and biogenesis |
| gene_70965 | 57,29203709 | -2,584489099 | 0,739241628 | -3,496135772 | 0,000472048 | 0,013437829 | The Moose: suction sampler vs FISH | Translation, ribosomal structure and biogenesis |
| gene_2510 | 123,2599005 | -2,573875421 | 0,674615623 | -3,815321399 | 0,000136006 | 0,006113186 | The Moose: suction sampler vs FISH | Translation, ribosomal structure and biogenesis |
| gene_685603 | 110,6116733 | -2,535680669 | 0,762191265 | -3,326829874 | 0,0008784 | 0,02015435 | The Moose: suction sampler vs FISH | Translation, ribosomal structure and biogenesis |
| gene_950183 | 27,90938877 | -2,513542739 | 0,751332464 | -3,345446733 | 0,000821501 | 0,019270255 | The Moose: suction sampler vs FISH | Translation, ribosomal structure and biogenesis |
| gene_950184 | 32,81543538 | -2,473516428 | 0,767482254 | -3,222897226 | 0,001269011 | 0,025191854 | The Moose: suction sampler vs FISH | Translation, ribosomal structure and biogenesis |
| gene_805847 | 168,1705387 | -2,470536102 | 0,690565943 | -3,577552771 | 0,000346826 | 0,011139952 | The Moose: suction sampler vs FISH | Translation, ribosomal structure and biogenesis |
| gene_962431 | 338,6238523 | -2,413412251 | 0,633245618 | -3,811178766 | 0,000138306 | 0,006163751 | The Moose: suction sampler vs FISH | Translation, ribosomal structure and biogenesis |
| gene_805845 | 68,68754233 | -2,401446984 | 0,730129693 | -3,289069059 | 0,001005194 | 0,021789969 | The Moose: suction sampler vs FISH | Translation, ribosomal structure and biogenesis |
| gene_212759 | 32,45421951 | -2,376486321 | 0,729062758 | -3,259645751 | 0,001115514 | 0,023308381 | The Moose: suction sampler vs FISH | Translation, ribosomal structure and biogenesis |
| gene_1133903 | 118,0921013 | -2,375757724 | 0,675038996 | -3,519437748 | 0,000432463 | 0,012766161 | The Moose: suction sampler vs FISH | Translation, ribosomal structure and biogenesis |
| gene_1371323 | 115,7801782 | -2,368521388 | 0,650631062 | -3,640344775 | 0,000272273 | 0,009545863 | The Moose: suction sampler vs FISH | Translation, ribosomal structure and biogenesis |
| gene_326083 | 79,91118926 | -2,360076638 | 0,77490313 | -3,045640864 | 0,00232185 | 0,036082789 | The Moose: suction sampler vs FISH | Translation, ribosomal structure and biogenesis |
| gene_1371317 | 295,494477 | -2,339484783 | 0,61630832 | -3,795964951 | 0,00014707 | 0,006389958 | The Moose: suction sampler vs FISH | Translation, ribosomal structure and biogenesis |
| gene_754877 | 171,7379086 | -2,333279728 | 0,617356563 | -3,779468575 | 0,000157163 | 0,0066799 | The Moose: suction sampler vs FISH | Translation, ribosomal structure and biogenesis |
| gene_220351 | 58,92719436 | -2,300195966 | 0,696919245 | -3,300520085 | 0,000965058 | 0,021265733 | The Moose: suction sampler vs FISH | Translation, ribosomal structure and biogenesis |
| gene_1371324 | 128,6496806 | -2,296763776 | 0,659219771 | -3,484063856 | 0,000493862 | 0,013860019 | The Moose: suction sampler vs FISH | Translation, ribosomal structure and biogenesis |
| gene_1133910 | 195,8224197 | -2,293046136 | 0,637211554 | -3,598563338 | 0,00031998 | 0,010602098 | The Moose: suction sampler vs FISH | Translation, ribosomal structure and biogenesis |
| gene_1371332 | 528,9678609 | -2,271983444 | 0,549861488 | -4,131919572 | 3,60E-05 | 0,002504414 | The Moose: suction sampler vs FISH | Translation, ribosomal structure and biogenesis |
| gene_1133907 | 150,3202561 | -2,255274531 | 0,570748381 | -3,951433954 | 7,77E-05 | 0,004232585 | The Moose: suction sampler vs FISH | Translation, ribosomal structure and biogenesis |
| gene_1278446 | 54,42618724 | -2,253346987 | 0,632521289 | -3,562484023 | 0,000367362 | 0,011535238 | The Moose: suction sampler vs FISH | Translation, ribosomal structure and biogenesis |
| gene_1409574 | 50,33963587 | -2,243088378 | 0,66751223 | -3,360370458 | 0,00077838 | 0,018603364 | The Moose: suction sampler vs FISH | Translation, ribosomal structure and biogenesis |
| gene_1371322 | 167,698375 | -2,240934767 | 0,619783411 | -3,615674004 | 0,000299567 | 0,010192304 | The Moose: suction sampler vs FISH | Translation, ribosomal structure and biogenesis |
| gene_1133911 | 212,8482877 | -2,234175923 | 0,580630796 | -3,84784262 | 0,000119163 | 0,00561739 | The Moose: suction sampler vs FISH | Translation, ribosomal structure and biogenesis |
| gene_895176 | 2699,401485 | -2,232669778 | 0,589017533 | -3,790497995 | 0,000150346 | 0,006479324 | The Moose: suction sampler vs FISH | Translation, ribosomal structure and biogenesis |
| gene_1809571 | 48,91126082 | -2,210508899 | 0,720992392 | -3,065925416 | 0,002169974 | 0,034797766 | The Moose: suction sampler vs FISH | Translation, ribosomal structure and biogenesis |
| gene_1371326 | 417,4225537 | -2,21020414 | 0,560154683 | -3,945703228 | 7,96E-05 | 0,004280407 | The Moose: suction sampler vs FISH | Translation, ribosomal structure and biogenesis |
| gene_858469 | 68,48826656 | -2,197403377 | 0,684496618 | -3,210247239 | 0,001326208 | 0,025850154 | The Moose: suction sampler vs FISH | Translation, ribosomal structure and biogenesis |
| gene_1371325 | 441,9522725 | -2,192616332 | 0,606406798 | -3,615751572 | 0,000299477 | 0,010192304 | The Moose: suction sampler vs FISH | Translation, ribosomal structure and biogenesis |
| gene_1283911 | 119,1227255 | -2,185642929 | 0,645597543 | -3,385457321 | 0,000710598 | 0,017507494 | The Moose: suction sampler vs FISH | Translation, ribosomal structure and biogenesis |
| gene_1133905 | 111,4357516 | -2,18182443 | 0,604198636 | -3,611104525 | 0,000304896 | 0,010290932 | The Moose: suction sampler vs FISH | Translation, ribosomal structure and biogenesis |
| gene_1371320 | 313,327095 | -2,179395266 | 0,565026536 | -3,857155601 | 0,000114714 | 0,005461423 | The Moose: suction sampler vs FISH | Translation, ribosomal structure and biogenesis |
| gene_87662 | 1416,684 | -2,177230291 | 0,645742683 | -3,371668544 | 0,000747143 | 0,018122167 | The Moose: suction sampler vs FISH | Translation, ribosomal structure and biogenesis |
| gene_720385 | 60,62856281 | -2,170266718 | 0,665980731 | -3,258753022 | 0,00111903 | 0,023342002 | The Moose: suction sampler vs FISH | Translation, ribosomal structure and biogenesis |
| gene_121837 | 32,91154524 | -2,167782456 | 0,692312194 | -3,131220964 | 0,001740811 | 0,030557812 | The Moose: suction sampler vs FISH | Translation, ribosomal structure and biogenesis |
| gene_1044605 | 204,583795 | -2,158840197 | 0,601776969 | -3,58744237 | 0,000333938 | 0,010890782 | The Moose: suction sampler vs FISH | Translation, ribosomal structure and biogenesis |
| gene_1371318 | 218,347838 | -2,145525506 | 0,618305148 | -3,470010742 | 0,000520438 | 0,014310571 | The Moose: suction sampler vs FISH | Translation, ribosomal structure and biogenesis |
| gene_97149 | 1190,208685 | -2,135655405 | 0,652840378 | -3,271328606 | 0,001070435 | 0,022715418 | The Moose: suction sampler vs FISH | Translation, ribosomal structure and biogenesis |
| gene_1371319 | 281,2844329 | -2,12399125 | 0,604828854 | -3,511722755 | 0,000445212 | 0,012991104 | The Moose: suction sampler vs FISH | Translation, ribosomal structure and biogenesis |
| gene_858468 | 91,45466802 | -2,119117963 | 0,61784263 | -3,429866864 | 0,000603877 | 0,015716435 | The Moose: suction sampler vs FISH | Translation, ribosomal structure and biogenesis |
| gene_656195 | 1523,667251 | -2,118345996 | 0,695348507 | -3,046452209 | 0,002315593 | 0,036030135 | The Moose: suction sampler vs FISH | Translation, ribosomal structure and biogenesis |
| gene_54435 | 2971,868456 | -2,116104225 | 0,600276581 | -3,525215361 | 0,000423139 | 0,012611664 | The Moose: suction sampler vs FISH | Translation, ribosomal structure and biogenesis |

| | | | | | | | | |
|---|---|---|---|---|---|---|---|---|
| gene_1371329 | 509,4279649 | -2,115534947 | 0,550964876 | -3,839691127 | 0,000123189 | 0,005751831 | The Moose: suction sampler vs FISH | Translation, ribosomal structure and biogenesis |
| gene_1921819 | 60,58588023 | -2,11516428 | 0,591534798 | -3,575722489 | 0,000349262 | 0,011162553 | The Moose: suction sampler vs FISH | Translation, ribosomal structure and biogenesis |
| gene_1371321 | 172,539611 | -2,114960408 | 0,628783333 | -3,363575807 | 0,000769397 | 0,01845682 | The Moose: suction sampler vs FISH | Translation, ribosomal structure and biogenesis |
| gene_1133913 | 170,4817458 | -2,112421891 | 0,533115402 | -3,962410174 | 7,42E-05 | 0,004121578 | The Moose: suction sampler vs FISH | Translation, ribosomal structure and biogenesis |
| gene_1044604 | 165,9960129 | -2,091905237 | 0,558391107 | -3,746308299 | 0,000179456 | 0,007257791 | The Moose: suction sampler vs FISH | Translation, ribosomal structure and biogenesis |
| gene_87638 | 4921,596334 | -2,086199117 | 0,611071224 | -3,414003204 | 0,000640158 | 0,016349683 | The Moose: suction sampler vs FISH | Translation, ribosomal structure and biogenesis |
| gene_1371316 | 327,2691347 | -2,063553043 | 0,548460169 | -3,762448325 | 0,000168258 | 0,007009481 | The Moose: suction sampler vs FISH | Translation, ribosomal structure and biogenesis |
| gene_1133912 | 296,3734416 | -2,060751427 | 0,551023689 | -3,739859949 | 0,000184123 | 0,007393603 | The Moose: suction sampler vs FISH | Translation, ribosomal structure and biogenesis |
| gene_70988 | 478,5175804 | -2,054614902 | 0,612062319 | -3,356872069 | 0,000788296 | 0,018742759 | The Moose: suction sampler vs FISH | Translation, ribosomal structure and biogenesis |
| gene_177934 | 4917,45925 | -2,051517454 | 0,573293385 | -3,578477459 | 0,000345602 | 0,011122688 | The Moose: suction sampler vs FISH | Translation, ribosomal structure and biogenesis |
| gene_340648 | 826,119719 | -2,028165108 | 0,654566645 | -3,098485269 | 0,001945126 | 0,03263058 | The Moose: suction sampler vs FISH | Translation, ribosomal structure and biogenesis |
| gene_70990 | 354,2255715 | -2,024552453 | 0,632002128 | -3,203394992 | 0,001358176 | 0,026205737 | The Moose: suction sampler vs FISH | Translation, ribosomal structure and biogenesis |
| gene_70989 | 457,1775068 | -2,01814028 | 0,622559633 | -3,241681878 | 0,001188266 | 0,024188521 | The Moose: suction sampler vs FISH | Translation, ribosomal structure and biogenesis |
| gene_447714 | 5843,00267 | -2,017704813 | 0,608886514 | -3,313761705 | 0,000920499 | 0,020699958 | The Moose: suction sampler vs FISH | Translation, ribosomal structure and biogenesis |
| gene_70987 | 251,205436 | -2,012094664 | 0,653834349 | -3,077376811 | 0,002088311 | 0,034091137 | The Moose: suction sampler vs FISH | Translation, ribosomal structure and biogenesis |
| gene_1371314 | 274,9044715 | -2,011613779 | 0,576022372 | -3,492249393 | 0,000478971 | 0,013571764 | The Moose: suction sampler vs FISH | Translation, ribosomal structure and biogenesis |
| gene_1044606 | 172,1625645 | -2,00188532 | 0,588370187 | -3,402424808 | 0,000667907 | 0,016789307 | The Moose: suction sampler vs FISH | Translation, ribosomal structure and biogenesis |
| gene_177304 | 68,82925495 | 2,132524523 | 0,716458367 | 2,976480731 | 0,002915774 | 0,041405457 | The Moose: suction sampler vs FISH | Translation, ribosomal structure and biogenesis |
| gene_592280 | 91,65581036 | 2,268527974 | 0,762711201 | 2,97429482 | 0,002936628 | 0,041589014 | The Moose: suction sampler vs FISH | Translation, ribosomal structure and biogenesis |
| gene_8309 | 103,0130562 | 2,42841384 | 0,834757832 | 2,909123757 | 0,003624434 | 0,047056129 | The Moose: suction sampler vs FISH | Translation, ribosomal structure and biogenesis |
| gene_264398 | 28,0807088 | 2,467015358 | 0,841848444 | 2,930474452 | 0,003384448 | 0,045129049 | The Moose: suction sampler vs FISH | Translation, ribosomal structure and biogenesis |
| gene_1002888 | 46,14451969 | 2,577032256 | 0,705629907 | 3,652101802 | 0,000260103 | 0,00927342 | The Moose: suction sampler vs FISH | Translation, ribosomal structure and biogenesis |
| gene_346652 | 37,13749179 | 2,70163817 | 0,897142427 | 3,011381571 | 0,002600618 | 0,038749472 | The Moose: suction sampler vs FISH | Translation, ribosomal structure and biogenesis |
| gene_108232 | 29,78837578 | 2,705876166 | 0,935002459 | 2,893977595 | 0,003803953 | 0,048389984 | The Moose: suction sampler vs FISH | Translation, ribosomal structure and biogenesis |
| gene_1662150 | 43,84633809 | 2,7803296 | 0,82509579 | 3,369705233 | 0,000752486 | 0,018215675 | The Moose: suction sampler vs FISH | Translation, ribosomal structure and biogenesis |
| gene_1236594 | 43,86890515 | 2,83697646 | 0,933876621 | 3,037849322 | 0,002382731 | 0,036610806 | The Moose: suction sampler vs FISH | Translation, ribosomal structure and biogenesis |
| gene_1054909 | 74,64652602 | 2,849132564 | 0,82485321 | 3,454108595 | 0,000552115 | 0,014837207 | The Moose: suction sampler vs FISH | Translation, ribosomal structure and biogenesis |
| gene_402501 | 35,26702733 | 2,863577005 | 0,951828327 | 3,008501559 | 0,002625395 | 0,038881355 | The Moose: suction sampler vs FISH | Translation, ribosomal structure and biogenesis |
| gene_1832285 | 20,09537402 | 2,885449606 | 0,865613011 | 3,333417555 | 0,000857861 | 0,019873068 | The Moose: suction sampler vs FISH | Translation, ribosomal structure and biogenesis |
| gene_404883 | 41,06513408 | 2,906867983 | 0,804958833 | 3,611200804 | 0,000304783 | 0,010290932 | The Moose: suction sampler vs FISH | Translation, ribosomal structure and biogenesis |
| gene_959432 | 34,36107458 | 2,923059522 | 0,81284857 | 3,596068971 | 0,000323062 | 0,010665961 | The Moose: suction sampler vs FISH | Translation, ribosomal structure and biogenesis |
| gene_1911728 | 1675,891684 | 2,953043794 | 0,678840561 | 4,35012868 | 1,36E-05 | 0,001299631 | The Moose: suction sampler vs FISH | Translation, ribosomal structure and biogenesis |
| gene_784584 | 108,2649374 | 2,953610062 | 0,814224103 | 3,627514896 | 0,000286162 | 0,009876498 | The Moose: suction sampler vs FISH | Translation, ribosomal structure and biogenesis |
| gene_618717 | 140,5475648 | 2,965710657 | 0,763458648 | 3,884572745 | 0,00010251 | 0,00506993 | The Moose: suction sampler vs FISH | Translation, ribosomal structure and biogenesis |
| gene_1201828 | 16,85387475 | 2,966143693 | 0,932627697 | 3,180415618 | 0,00147064 | 0,027589049 | The Moose: suction sampler vs FISH | Translation, ribosomal structure and biogenesis |
| gene_1870151 | 27,85289908 | 3,020795313 | 0,910497712 | 3,317740697 | 0,000907487 | 0,020558413 | The Moose: suction sampler vs FISH | Translation, ribosomal structure and biogenesis |
| gene_69276 | 46,97369037 | 3,052973697 | 0,831885124 | 3,669946257 | 0,000242601 | 0,008869022 | The Moose: suction sampler vs FISH | Translation, ribosomal structure and biogenesis |
| gene_1492668 | 79,88420951 | 3,073892485 | 0,795564458 | 3,863788101 | 0,000111642 | 0,005376756 | The Moose: suction sampler vs FISH | Translation, ribosomal structure and biogenesis |
| gene_693335 | 68,36289922 | 3,109446438 | 0,981546517 | 3,167905324 | 0,001535415 | 0,028301954 | The Moose: suction sampler vs FISH | Translation, ribosomal structure and biogenesis |
| gene_1093672 | 20,99775134 | 3,12909822 | 0,96960971 | 3,22717294 | 0,001250199 | 0,024924225 | The Moose: suction sampler vs FISH | Translation, ribosomal structure and biogenesis |
| gene_146756 | 21,62400879 | 3,154016516 | 0,990122263 | 3,185481869 | 0,001445132 | 0,027256146 | The Moose: suction sampler vs FISH | Translation, ribosomal structure and biogenesis |
| gene_682390 | 18,74884068 | 3,180653209 | 0,930120556 | 3,419613929 | 0,000627101 | 0,016115258 | The Moose: suction sampler vs FISH | Translation, ribosomal structure and biogenesis |
| gene_1329837 | 35,33735603 | 3,197400358 | 0,8943999 | 3,574911356 | 0,000350347 | 0,011182618 | The Moose: suction sampler vs FISH | Translation, ribosomal structure and biogenesis |
| gene_516871 | 16,04222341 | 3,208668335 | 0,958887649 | 3,346240135 | 0,000819154 | 0,019227479 | The Moose: suction sampler vs FISH | Translation, ribosomal structure and biogenesis |
| gene_1588917 | 72,50775418 | 3,301073202 | 0,810817841 | 4,071288316 | 4,68E-05 | 0,003037636 | The Moose: suction sampler vs FISH | Translation, ribosomal structure and biogenesis |
| gene_1064873 | 15,30551406 | 3,301416868 | 1,147052468 | 2,878174241 | 0,003999841 | 0,049803793 | The Moose: suction sampler vs FISH | Translation, ribosomal structure and biogenesis |
| gene_965255 | 119,1830799 | 3,353442346 | 0,842423336 | 3,980709224 | 6,87E-05 | 0,003884854 | The Moose: suction sampler vs FISH | Translation, ribosomal structure and biogenesis |
| gene_384975 | 14,78871627 | 3,357289575 | 1,036259535 | 3,239815375 | 0,001196071 | 0,024295534 | The Moose: suction sampler vs FISH | Translation, ribosomal structure and biogenesis |
| gene_1950560 | 62,74347021 | 3,369311089 | 1,039432773 | 3,241490142 | 0,001189065 | 0,024188888 | The Moose: suction sampler vs FISH | Translation, ribosomal structure and biogenesis |
| gene_715471 | 25,95114064 | 3,37841906 | 1,061392762 | 3,183005556 | 0,001457548 | 0,027413476 | The Moose: suction sampler vs FISH | Translation, ribosomal structure and biogenesis |
| gene_270408 | 54,12297501 | 3,400609034 | 0,878425607 | 3,871254443 | 0,000108277 | 0,005250198 | The Moose: suction sampler vs FISH | Translation, ribosomal structure and biogenesis |
| gene_509855 | 13,9796135 | 3,418560692 | 1,097461542 | 3,114970831 | 0,001839631 | 0,031605076 | The Moose: suction sampler vs FISH | Translation, ribosomal structure and biogenesis |
| gene_724278 | 98,69069078 | 3,477607035 | 0,810426877 | 4,291080583 | 1,78E-05 | 0,001540172 | The Moose: suction sampler vs FISH | Translation, ribosomal structure and biogenesis |
| gene_38541 | 20,48364674 | 3,484188015 | 0,951035827 | 3,663571779 | 0,000248722 | 0,00899873 | The Moose: suction sampler vs FISH | Translation, ribosomal structure and biogenesis |
| gene_1531178 | 17,42031287 | 3,486301192 | 1,117971084 | 3,118418035 | 0,001818247 | 0,031347689 | The Moose: suction sampler vs FISH | Translation, ribosomal structure and biogenesis |
| gene_420196 | 76,30126034 | 3,547044972 | 0,806486569 | 4,398145125 | 1,09E-05 | 0,00112776 | The Moose: suction sampler vs FISH | Translation, ribosomal structure and biogenesis |
| gene_1191877 | 15,67434721 | 3,566680277 | 1,174458435 | 3,036872289 | 0,002390467 | 0,036683631 | The Moose: suction sampler vs FISH | Translation, ribosomal structure and biogenesis |
| gene_1105099 | 10,12999418 | 3,567612724 | 1,180983333 | 3,020883212 | 0,002520386 | 0,037965358 | The Moose: suction sampler vs FISH | Translation, ribosomal structure and biogenesis |
| gene_10615 | 12,5625629 | 3,569158014 | 1,188699413 | 3,002574052 | 0,002677068 | 0,039306464 | The Moose: suction sampler vs FISH | Translation, ribosomal structure and biogenesis |
| gene_1689859 | 12,72248811 | 3,569406893 | 1,234242293 | 2,891982322 | 0,003828195 | 0,048578262 | The Moose: suction sampler vs FISH | Translation, ribosomal structure and biogenesis |
| gene_391875 | 35,56974014 | 3,583943111 | 0,922410785 | 3,885408939 | 0,000102158 | 0,005069338 | The Moose: suction sampler vs FISH | Translation, ribosomal structure and biogenesis |
| gene_1039025 | 27,12363044 | 3,605371979 | 1,091289942 | 3,303770924 | 0,000953937 | 0,021115713 | The Moose: suction sampler vs FISH | Translation, ribosomal structure and biogenesis |
| gene_258384 | 7,929948164 | 3,66375538 | 1,23963796 | 2,955504347 | 0,003121583 | 0,043192287 | The Moose: suction sampler vs FISH | Translation, ribosomal structure and biogenesis |
| gene_759363 | 11,3533218 | 3,665208461 | 1,167970371 | 3,138100547 | 0,001700465 | 0,030123173 | The Moose: suction sampler vs FISH | Translation, ribosomal structure and biogenesis |
| gene_143846 | 14,79618323 | 3,722509133 | 1,186477942 | 3,137444872 | 0,001704273 | 0,030161529 | The Moose: suction sampler vs FISH | Translation, ribosomal structure and biogenesis |

| | | | | | | | | |
|---|---|---|---|---|---|---|---|---|
| gene_1708244 | 10,02594822 | 3,725802406 | 1,170260627 | 3,183737298 | 0,001453869 | 0,027366728 | The Moose: suction sampler vs FISH | Translation, ribosomal structure and biogenesis |
| gene_1206580 | 19,52655391 | 3,732525958 | 1,253542115 | 2,977583213 | 0,002905308 | 0,041320744 | The Moose: suction sampler vs FISH | Translation, ribosomal structure and biogenesis |
| gene_459080 | 10,17670857 | 3,739031008 | 1,19683171 | 3,124107572 | 0,001783452 | 0,030943932 | The Moose: suction sampler vs FISH | Translation, ribosomal structure and biogenesis |
| gene_179450 | 18,89209109 | 3,764336067 | 1,298363549 | 2,899292783 | 0,003740055 | 0,048009067 | The Moose: suction sampler vs FISH | Translation, ribosomal structure and biogenesis |
| gene_968621 | 12,04721683 | 3,788183344 | 1,273839407 | 2,973831179 | 0,002941068 | 0,041639973 | The Moose: suction sampler vs FISH | Translation, ribosomal structure and biogenesis |
| gene_1761864 | 14,13726724 | 3,798910362 | 1,009684174 | 3,762473911 | 0,000168241 | 0,007009481 | The Moose: suction sampler vs FISH | Translation, ribosomal structure and biogenesis |
| gene_1059302 | 2562,291248 | 3,819763835 | 0,79409008 | 4,81023996 | 1,51E-06 | 0,000292899 | The Moose: suction sampler vs FISH | Translation, ribosomal structure and biogenesis |
| gene_1856372 | 106,7586277 | 3,824725335 | 0,796484059 | 4,802011155 | 1,57E-06 | 0,000302009 | The Moose: suction sampler vs FISH | Translation, ribosomal structure and biogenesis |
| gene_1397082 | 95,14165802 | 3,847833565 | 0,870707869 | 4,419201551 | 9,91E-06 | 0,001052955 | The Moose: suction sampler vs FISH | Translation, ribosomal structure and biogenesis |
| gene_750413 | 22,61111516 | 3,859872849 | 1,23119754 | 3,135055686 | 0,001718215 | 0,030313307 | The Moose: suction sampler vs FISH | Translation, ribosomal structure and biogenesis |
| gene_1836897 | 332,0914574 | 3,881027251 | 0,791462826 | 4,903612807 | 9,41E-07 | 0,000215278 | The Moose: suction sampler vs FISH | Translation, ribosomal structure and biogenesis |
| gene_230071 | 18,20710953 | 3,890562251 | 1,10381211 | 3,524659872 | 0,000424027 | 0,012618579 | The Moose: suction sampler vs FISH | Translation, ribosomal structure and biogenesis |
| gene_1439909 | 10,63922627 | 3,910015244 | 1,133993302 | 3,448005589 | 0,000564742 | 0,015103025 | The Moose: suction sampler vs FISH | Translation, ribosomal structure and biogenesis |
| gene_24775 | 14,91755347 | 3,921306502 | 1,197814841 | 3,273716746 | 0,00106143 | 0,022622371 | The Moose: suction sampler vs FISH | Translation, ribosomal structure and biogenesis |
| gene_1874358 | 65,37976264 | 3,962776745 | 0,840152279 | 4,716736293 | 2,40E-06 | 0,000393774 | The Moose: suction sampler vs FISH | Translation, ribosomal structure and biogenesis |
| gene_1383818 | 80,31263429 | 3,96874684 | 0,873773808 | 4,542075771 | 5,57E-06 | 0,000711497 | The Moose: suction sampler vs FISH | Translation, ribosomal structure and biogenesis |
| gene_1805986 | 11,19073811 | 3,976683261 | 1,367984651 | 2,906964825 | 0,003649542 | 0,047265145 | The Moose: suction sampler vs FISH | Translation, ribosomal structure and biogenesis |
| gene_1383819 | 277,2812151 | 3,982449205 | 0,896199836 | 4,44370669 | 8,84E-06 | 0,0009751 | The Moose: suction sampler vs FISH | Translation, ribosomal structure and biogenesis |
| gene_1506395 | 8,389627204 | 4,011152939 | 1,318089222 | 3,043157376 | 0,002341099 | 0,036228822 | The Moose: suction sampler vs FISH | Translation, ribosomal structure and biogenesis |
| gene_354414 | 257,1200883 | 4,016600769 | 0,777969794 | 5,162926374 | 2,43E-07 | 8,54E-05 | The Moose: suction sampler vs FISH | Translation, ribosomal structure and biogenesis |
| gene_1352150 | 23,73514705 | 4,016897814 | 1,02769414 | 3,908651082 | 9,28E-05 | 0,00477356 | The Moose: suction sampler vs FISH | Translation, ribosomal structure and biogenesis |
| gene_852264 | 67,25008501 | 4,018869418 | 0,856768811 | 4,69072796 | 2,72E-06 | 0,000427245 | The Moose: suction sampler vs FISH | Translation, ribosomal structure and biogenesis |
| gene_1188198 | 65,96355461 | 4,023162685 | 0,913543988 | 4,403906915 | 1,06E-05 | 0,001107598 | The Moose: suction sampler vs FISH | Translation, ribosomal structure and biogenesis |
| gene_1633537 | 26,66682485 | 4,025957305 | 0,966597816 | 4,165080076 | 3,11E-05 | 0,002267936 | The Moose: suction sampler vs FISH | Translation, ribosomal structure and biogenesis |
| gene_1304729 | 13,51204321 | 4,026865837 | 1,198291142 | 3,360507056 | 0,000777995 | 0,01860022 | The Moose: suction sampler vs FISH | Translation, ribosomal structure and biogenesis |
| gene_281746 | 24,40212321 | 4,029456214 | 1,051329122 | 3,832725765 | 0,000126731 | 0,005828286 | The Moose: suction sampler vs FISH | Translation, ribosomal structure and biogenesis |
| gene_1686221 | 31,04814409 | 4,051969779 | 0,916325697 | 4,421975497 | 9,78E-06 | 0,001044056 | The Moose: suction sampler vs FISH | Translation, ribosomal structure and biogenesis |
| gene_304495 | 25,70248313 | 4,064536125 | 0,959736101 | 4,235055993 | 2,28E-05 | 0,001828083 | The Moose: suction sampler vs FISH | Translation, ribosomal structure and biogenesis |
| gene_827533 | 49,02317977 | 4,065209735 | 1,021406931 | 3,980009937 | 6,89E-05 | 0,003887304 | The Moose: suction sampler vs FISH | Translation, ribosomal structure and biogenesis |
| gene_545983 | 24,67040272 | 4,06537666 | 0,992985006 | 4,094096726 | 4,24E-05 | 0,002813278 | The Moose: suction sampler vs FISH | Translation, ribosomal structure and biogenesis |
| gene_1667722 | 32,8436884 | 4,066378574 | 0,94774717 | 4,290573162 | 1,78E-05 | 0,001540172 | The Moose: suction sampler vs FISH | Translation, ribosomal structure and biogenesis |
| gene_1376071 | 47,0331243 | 4,075635871 | 0,805459462 | 5,060013647 | 4,19E-07 | 0,00012267 | The Moose: suction sampler vs FISH | Translation, ribosomal structure and biogenesis |
| gene_844917 | 114,0866495 | 4,080864304 | 0,863626425 | 4,725265678 | 2,30E-06 | 0,000382738 | The Moose: suction sampler vs FISH | Translation, ribosomal structure and biogenesis |
| gene_863979 | 220,0601439 | 4,098131512 | 0,833049722 | 4,91943206 | 8,68E-07 | 0,000202372 | The Moose: suction sampler vs FISH | Translation, ribosomal structure and biogenesis |
| gene_24763 | 8,087556718 | 4,10026292 | 1,390732856 | 2,948275005 | 0,003195527 | 0,043745657 | The Moose: suction sampler vs FISH | Translation, ribosomal structure and biogenesis |
| gene_329323 | 159,4236093 | 4,104120444 | 0,838434291 | 4,894981619 | 9,83E-07 | 0,000220298 | The Moose: suction sampler vs FISH | Translation, ribosomal structure and biogenesis |
| gene_1351953 | 19,72952397 | 4,150618552 | 1,042961485 | 3,979647006 | 6,90E-05 | 0,003890255 | The Moose: suction sampler vs FISH | Translation, ribosomal structure and biogenesis |
| gene_1376074 | 15,63796698 | 4,153764729 | 0,991237728 | 4,190482879 | 2,78E-05 | 0,002116382 | The Moose: suction sampler vs FISH | Translation, ribosomal structure and biogenesis |
| gene_752580 | 177,9679771 | 4,157896363 | 0,828513843 | 5,01849957 | 5,21E-07 | 0,000143786 | The Moose: suction sampler vs FISH | Translation, ribosomal structure and biogenesis |
| gene_1477895 | 85,93082579 | 4,166467721 | 0,838201732 | 4,970721916 | 6,67E-07 | 0,000167204 | The Moose: suction sampler vs FISH | Translation, ribosomal structure and biogenesis |
| gene_698762 | 67,11538419 | 4,17735128 | 0,918511572 | 4,547957161 | 5,42E-06 | 0,000696756 | The Moose: suction sampler vs FISH | Translation, ribosomal structure and biogenesis |
| gene_423260 | 10,82857011 | 4,181574705 | 1,324477351 | 3,157150782 | 0,001593189 | 0,028923476 | The Moose: suction sampler vs FISH | Translation, ribosomal structure and biogenesis |
| gene_1247085 | 167,2431952 | 4,195137345 | 0,947197571 | 4,428999265 | 9,47E-06 | 0,001025536 | The Moose: suction sampler vs FISH | Translation, ribosomal structure and biogenesis |
| gene_912071 | 125,9067451 | 4,223939534 | 0,877105943 | 4,815768911 | 1,47E-06 | 0,000286424 | The Moose: suction sampler vs FISH | Translation, ribosomal structure and biogenesis |
| gene_444110 | 7,861090606 | 4,22623292 | 1,308665063 | 3,229422897 | 0,001240403 | 0,024782757 | The Moose: suction sampler vs FISH | Translation, ribosomal structure and biogenesis |
| gene_326389 | 13,84399936 | 4,236380702 | 1,211037572 | 3,498141428 | 0,000468513 | 0,013392145 | The Moose: suction sampler vs FISH | Translation, ribosomal structure and biogenesis |
| gene_10670 | 10,59177429 | 4,243814557 | 1,299996928 | 3,26448045 | 0,00109665 | 0,023091595 | The Moose: suction sampler vs FISH | Translation, ribosomal structure and biogenesis |
| gene_230715 | 66,15980957 | 4,251644864 | 0,86395799 | 4,921124537 | 8,60E-07 | 0,000202263 | The Moose: suction sampler vs FISH | Translation, ribosomal structure and biogenesis |
| gene_1760261 | 8,497401081 | 4,255785523 | 1,473308272 | 2,888591344 | 0,003869716 | 0,048867143 | The Moose: suction sampler vs FISH | Translation, ribosomal structure and biogenesis |
| gene_1869856 | 10,01482858 | 4,266827553 | 1,240826456 | 3,43869808 | 0,000584519 | 0,015399043 | The Moose: suction sampler vs FISH | Translation, ribosomal structure and biogenesis |
| gene_189822 | 14,19287924 | 4,276226366 | 1,243936573 | 3,437656274 | 0,000586772 | 0,015446405 | The Moose: suction sampler vs FISH | Translation, ribosomal structure and biogenesis |
| gene_743833 | 73,1246623 | 4,281178853 | 0,811201214 | 5,277579446 | 1,31E-07 | 5,46E-05 | The Moose: suction sampler vs FISH | Translation, ribosomal structure and biogenesis |
| gene_1862836 | 6,120724754 | 4,283294227 | 1,348933959 | 3,175317961 | 0,001496724 | 0,027878999 | The Moose: suction sampler vs FISH | Translation, ribosomal structure and biogenesis |
| gene_1443291 | 4419,266321 | 4,303053964 | 0,812525777 | 5,295898403 | 1,18E-07 | 5,15E-05 | The Moose: suction sampler vs FISH | Translation, ribosomal structure and biogenesis |
| gene_1685761 | 106,068441 | 4,364298171 | 0,805035436 | 5,421249765 | 5,92E-08 | 3,35E-05 | The Moose: suction sampler vs FISH | Translation, ribosomal structure and biogenesis |
| gene_590388 | 24,46510364 | 4,382293415 | 0,918845832 | 4,769345699 | 1,85E-06 | 0,000330734 | The Moose: suction sampler vs FISH | Translation, ribosomal structure and biogenesis |
| gene_270608 | 145,8216088 | 4,417255331 | 0,872826031 | 5,060865709 | 4,17E-07 | 0,000122611 | The Moose: suction sampler vs FISH | Translation, ribosomal structure and biogenesis |
| gene_615126 | 139,8358641 | 4,427442228 | 0,889786698 | 4,975846728 | 6,50E-07 | 0,000164687 | The Moose: suction sampler vs FISH | Translation, ribosomal structure and biogenesis |
| gene_1456957 | 7,520307037 | 4,441114091 | 1,435207477 | 3,094405625 | 0,001972077 | 0,032925476 | The Moose: suction sampler vs FISH | Translation, ribosomal structure and biogenesis |
| gene_722690 | 68,63399959 | 4,451533686 | 0,821692645 | 5,417516775 | 6,04E-08 | 3,39E-05 | The Moose: suction sampler vs FISH | Translation, ribosomal structure and biogenesis |
| gene_1506401 | 15,17154509 | 4,45761525 | 1,209838575 | 3,684471088 | 0,000229178 | 0,008544153 | The Moose: suction sampler vs FISH | Translation, ribosomal structure and biogenesis |
| gene_1068392 | 12,67084121 | 4,469519814 | 1,153236072 | 3,875633031 | 0,000106348 | 0,005195916 | The Moose: suction sampler vs FISH | Translation, ribosomal structure and biogenesis |
| gene_1665800 | 6479,919831 | 4,502738624 | 0,819414915 | 5,495065492 | 3,91E-08 | 2,46E-05 | The Moose: suction sampler vs FISH | Translation, ribosomal structure and biogenesis |
| gene_820691 | 63,80727279 | 4,503480927 | 0,975915618 | 4,614621229 | 3,94E-06 | 0,000555063 | The Moose: suction sampler vs FISH | Translation, ribosomal structure and biogenesis |
| gene_420195 | 157,8767854 | 4,509157034 | 0,808678304 | 5,575958962 | 2,46E-08 | 1,72E-05 | The Moose: suction sampler vs FISH | Translation, ribosomal structure and biogenesis |

| | | | | | | | | |
|---|---|---|---|---|---|---|---|---|
| gene_20045 | 141,8608624 | 4,553980407 | 0,854916728 | 5,326811672 | 1,00E-07 | 4,64E-05 | The Moose: suction sampler vs FISH | Translation, ribosomal structure and biogenesis |
| gene_1254106 | 146,092546 | 4,554662075 | 0,858667608 | 5,304336662 | 1,13E-07 | 5,00E-05 | The Moose: suction sampler vs FISH | Translation, ribosomal structure and biogenesis |
| gene_607163 | 34,92259225 | 4,558468968 | 0,888307041 | 5,131636647 | 2,87E-07 | 9,38E-05 | The Moose: suction sampler vs FISH | Translation, ribosomal structure and biogenesis |
| gene_1648661 | 5,911806923 | 4,584596312 | 1,389395049 | 3,299706815 | 0,000967859 | 0,021301886 | The Moose: suction sampler vs FISH | Translation, ribosomal structure and biogenesis |
| gene_1701481 | 13,11337341 | 4,593430579 | 1,208961345 | 3,79948507 | 0,000144997 | 0,006354002 | The Moose: suction sampler vs FISH | Translation, ribosomal structure and biogenesis |
| gene_1034808 | 18,0570802 | 4,603276825 | 1,100536858 | 4,182755709 | 2,88E-05 | 0,002160197 | The Moose: suction sampler vs FISH | Translation, ribosomal structure and biogenesis |
| gene_621709 | 111,4215884 | 4,6037116 | 0,866530721 | 5,31280829 | 1,08E-07 | 4,86E-05 | The Moose: suction sampler vs FISH | Translation, ribosomal structure and biogenesis |
| gene_36213 | 112,0056825 | 4,610808581 | 0,883512378 | 5,218725504 | 1,80E-07 | 6,68E-05 | The Moose: suction sampler vs FISH | Translation, ribosomal structure and biogenesis |
| gene_597265 | 47,22979246 | 4,617782859 | 0,883978696 | 5,223862156 | 1,75E-07 | 6,53E-05 | The Moose: suction sampler vs FISH | Translation, ribosomal structure and biogenesis |
| gene_1272151 | 7,762306469 | 4,634607524 | 1,479159584 | 3,133270793 | 0,001728699 | 0,030426115 | The Moose: suction sampler vs FISH | Translation, ribosomal structure and biogenesis |
| gene_1350523 | 115,1190086 | 4,649317228 | 0,814731875 | 5,706561106 | 1,15E-08 | 9,85E-06 | The Moose: suction sampler vs FISH | Translation, ribosomal structure and biogenesis |
| gene_435792 | 10,26808369 | 4,658622827 | 1,331751434 | 3,498117373 | 0,000468555 | 0,013392145 | The Moose: suction sampler vs FISH | Translation, ribosomal structure and biogenesis |
| gene_1397083 | 74,53454292 | 4,660332507 | 0,906415946 | 5,141494399 | 2,73E-07 | 9,10E-05 | The Moose: suction sampler vs FISH | Translation, ribosomal structure and biogenesis |
| gene_1405380 | 70,68637824 | 4,704569705 | 0,891924562 | 5,27462737 | 1,33E-07 | 5,47E-05 | The Moose: suction sampler vs FISH | Translation, ribosomal structure and biogenesis |
| gene_60764 | 137,6789518 | 4,706581313 | 0,860487591 | 5,469667855 | 4,51E-08 | 2,78E-05 | The Moose: suction sampler vs FISH | Translation, ribosomal structure and biogenesis |
| gene_149058 | 5,126745872 | 4,716461995 | 1,593302652 | 2,960179593 | 0,003074598 | 0,042824543 | The Moose: suction sampler vs FISH | Translation, ribosomal structure and biogenesis |
| gene_1841699 | 9,587860387 | 4,732548497 | 1,263523167 | 3,745517788 | 0,000180022 | 0,007272675 | The Moose: suction sampler vs FISH | Translation, ribosomal structure and biogenesis |
| gene_24759 | 16,45294152 | 4,736314041 | 1,308065316 | 3,62085439 | 0,000293632 | 0,010044613 | The Moose: suction sampler vs FISH | Translation, ribosomal structure and biogenesis |
| gene_118121 | 110,5445847 | 4,74134121 | 0,86830254 | 5,46047143 | 4,75E-08 | 2,88E-05 | The Moose: suction sampler vs FISH | Translation, ribosomal structure and biogenesis |
| gene_832545 | 62,32228786 | 4,754836348 | 0,984904632 | 4,827712443 | 1,38E-06 | 0,000275805 | The Moose: suction sampler vs FISH | Translation, ribosomal structure and biogenesis |
| gene_841268 | 21,4131115 | 4,774548429 | 0,983300972 | 4,855632774 | 1,20E-06 | 0,0002511 | The Moose: suction sampler vs FISH | Translation, ribosomal structure and biogenesis |
| gene_1301555 | 8,69691842 | 4,77848453 | 1,438061418 | 3,322865401 | 0,000890979 | 0,020360283 | The Moose: suction sampler vs FISH | Translation, ribosomal structure and biogenesis |
| gene_976916 | 369,2741378 | 4,789607022 | 0,817869124 | 5,856202269 | 4,74E-09 | 5,27E-06 | The Moose: suction sampler vs FISH | Translation, ribosomal structure and biogenesis |
| gene_929245 | 36,01391995 | 4,800746612 | 0,973274678 | 4,932571166 | 8,12E-07 | 0,000194783 | The Moose: suction sampler vs FISH | Translation, ribosomal structure and biogenesis |
| gene_781971 | 57,2516092 | 4,803874895 | 1,030164149 | 4,663213043 | 3,11E-06 | 0,000466616 | The Moose: suction sampler vs FISH | Translation, ribosomal structure and biogenesis |
| gene_545586 | 658,3032524 | 4,815567757 | 0,830046118 | 5,801566508 | 6,57E-09 | 6,61E-06 | The Moose: suction sampler vs FISH | Translation, ribosomal structure and biogenesis |
| gene_1466325 | 10,18784645 | 4,843657848 | 1,362907581 | 3,553915112 | 0,000379542 | 0,011793971 | The Moose: suction sampler vs FISH | Translation, ribosomal structure and biogenesis |
| gene_20046 | 259,2885191 | 4,843740431 | 0,847858227 | 5,712913173 | 1,11E-08 | 9,60E-06 | The Moose: suction sampler vs FISH | Translation, ribosomal structure and biogenesis |
| gene_1376070 | 11,66649377 | 4,876912371 | 1,361587689 | 3,581783539 | 0,000341256 | 0,011021805 | The Moose: suction sampler vs FISH | Translation, ribosomal structure and biogenesis |
| gene_985495 | 8,958689106 | 4,931788565 | 1,657641478 | 2,975184098 | 0,002928128 | 0,041507639 | The Moose: suction sampler vs FISH | Translation, ribosomal structure and biogenesis |
| gene_1721581 | 4,233960476 | 4,964836811 | 1,669130914 | 2,974504139 | 0,002934625 | 0,041568666 | The Moose: suction sampler vs FISH | Translation, ribosomal structure and biogenesis |
| gene_1252961 | 40,08053978 | 4,999492761 | 1,118964765 | 4,467962634 | 7,90E-06 | 0,000909061 | The Moose: suction sampler vs FISH | Translation, ribosomal structure and biogenesis |
| gene_876669 | 16,03651834 | 5,015332125 | 1,12670413 | 4,451330205 | 8,53E-06 | 0,000961318 | The Moose: suction sampler vs FISH | Translation, ribosomal structure and biogenesis |
| gene_1342511 | 229,8094405 | 5,035998976 | 0,860976196 | 5,8491733 | 4,94E-09 | 5,34E-06 | The Moose: suction sampler vs FISH | Translation, ribosomal structure and biogenesis |
| gene_1821079 | 120,5060857 | 5,049856609 | 0,814916637 | 6,196776919 | 5,76E-10 | 1,18E-06 | The Moose: suction sampler vs FISH | Translation, ribosomal structure and biogenesis |
| gene_547431 | 56,53832055 | 5,05463755 | 0,880458964 | 5,740912135 | 9,42E-09 | 8,87E-06 | The Moose: suction sampler vs FISH | Translation, ribosomal structure and biogenesis |
| gene_747022 | 6,856128321 | 5,104725476 | 1,612137086 | 3,166433873 | 0,001543204 | 0,028370623 | The Moose: suction sampler vs FISH | Translation, ribosomal structure and biogenesis |
| gene_1103851 | 38,30080981 | 5,119034915 | 1,035982524 | 4,941236747 | 7,76E-07 | 0,000188789 | The Moose: suction sampler vs FISH | Translation, ribosomal structure and biogenesis |
| gene_1263195 | 162,1782455 | 5,137977513 | 0,919262593 | 5,589238105 | 2,28E-08 | 1,61E-05 | The Moose: suction sampler vs FISH | Translation, ribosomal structure and biogenesis |
| gene_44666 | 153,9376373 | 5,181564001 | 0,836228593 | 6,196348752 | 5,78E-10 | 1,18E-06 | The Moose: suction sampler vs FISH | Translation, ribosomal structure and biogenesis |
| gene_1719129 | 58,8563521 | 5,235814306 | 0,86399614 | 6,059997335 | 1,36E-09 | 2,22E-06 | The Moose: suction sampler vs FISH | Translation, ribosomal structure and biogenesis |
| gene_1508276 | 4,739643779 | 5,244419082 | 1,786745229 | 2,935180123 | 0,003333541 | 0,044841013 | The Moose: suction sampler vs FISH | Translation, ribosomal structure and biogenesis |
| gene_1668063 | 31,56400125 | 5,255221439 | 0,945574453 | 5,557702435 | 2,73E-08 | 1,89E-05 | The Moose: suction sampler vs FISH | Translation, ribosomal structure and biogenesis |
| gene_1334879 | 14,39162483 | 5,272467621 | 1,18632564 | 4,444367924 | 8,82E-06 | 0,000973567 | The Moose: suction sampler vs FISH | Translation, ribosomal structure and biogenesis |
| gene_1765578 | 25,90343177 | 5,280536177 | 1,023866339 | 5,157446805 | 2,50E-07 | 8,66E-05 | The Moose: suction sampler vs FISH | Translation, ribosomal structure and biogenesis |
| gene_24769 | 10,6913026 | 5,307466111 | 1,510601878 | 3,513477766 | 0,000442281 | 0,012950027 | The Moose: suction sampler vs FISH | Translation, ribosomal structure and biogenesis |
| gene_90670 | 56,83724798 | 5,35371935 | 0,927937864 | 5,769480435 | 7,95E-09 | 7,79E-06 | The Moose: suction sampler vs FISH | Translation, ribosomal structure and biogenesis |
| gene_1532627 | 8,350198338 | 5,443522094 | 1,471046763 | 3,700441231 | 0,000215225 | 0,008181778 | The Moose: suction sampler vs FISH | Translation, ribosomal structure and biogenesis |
| gene_1124716 | 217,5127778 | 5,502033302 | 0,885431638 | 6,213956069 | 5,17E-10 | 1,12E-06 | The Moose: suction sampler vs FISH | Translation, ribosomal structure and biogenesis |
| gene_1838646 | 4,713238813 | 5,5267426 | 1,784323365 | 3,097388461 | 0,001952339 | 0,032684422 | The Moose: suction sampler vs FISH | Translation, ribosomal structure and biogenesis |
| gene_423258 | 7,628667658 | 5,558711481 | 1,512561567 | 3,675031551 | 0,00023782 | 0,008768419 | The Moose: suction sampler vs FISH | Translation, ribosomal structure and biogenesis |
| gene_1755348 | 197,5735107 | 5,679490527 | 0,92967778 | 6,109095699 | 1,00E-09 | 1,79E-06 | The Moose: suction sampler vs FISH | Translation, ribosomal structure and biogenesis |
| gene_1767678 | 9,182667814 | 5,689772972 | 1,582682412 | 3,595018765 | 0,000324368 | 0,010695092 | The Moose: suction sampler vs FISH | Translation, ribosomal structure and biogenesis |
| gene_1898383 | 7,805897388 | 5,752250853 | 1,600191877 | 3,594725693 | 0,000324734 | 0,010695092 | The Moose: suction sampler vs FISH | Translation, ribosomal structure and biogenesis |
| gene_1064136 | 8,858230339 | 5,753450733 | 1,632898485 | 3,523458921 | 0,000425953 | 0,012665638 | The Moose: suction sampler vs FISH | Translation, ribosomal structure and biogenesis |
| gene_1249210 | 4,557835234 | 5,765812419 | 1,947405666 | 2,960765966 | 0,00306875 | 0,042751208 | The Moose: suction sampler vs FISH | Translation, ribosomal structure and biogenesis |
| gene_420459 | 7,040229866 | 5,802191608 | 1,638558904 | 3,54103328 | 0,000398563 | 0,012187919 | The Moose: suction sampler vs FISH | Translation, ribosomal structure and biogenesis |
| gene_108752 | 4,567106728 | 5,83120662 | 1,826907688 | 3,191845246 | 0,001413671 | 0,026863397 | The Moose: suction sampler vs FISH | Translation, ribosomal structure and biogenesis |
| gene_449561 | 4,628817009 | 5,941897984 | 1,754223728 | 3,387195081 | 0,000706112 | 0,017424353 | The Moose: suction sampler vs FISH | Translation, ribosomal structure and biogenesis |
| gene_1738827 | 4,083171635 | 5,988529935 | 1,989176749 | 3,010556974 | 0,00260769 | 0,038790953 | The Moose: suction sampler vs FISH | Translation, ribosomal structure and biogenesis |
| gene_765338 | 2,922801529 | 5,993391314 | 2,077115076 | 2,885440188 | 0,003908667 | 0,049120591 | The Moose: suction sampler vs FISH | Translation, ribosomal structure and biogenesis |
| gene_1408202 | 7,369714876 | 6,045470192 | 1,793474344 | 3,370814983 | 0,000749462 | 0,018166408 | The Moose: suction sampler vs FISH | Translation, ribosomal structure and biogenesis |
| gene_1431188 | 3,527487804 | 6,107980194 | 2,071148442 | 2,949078911 | 0,003187226 | 0,04367781 | The Moose: suction sampler vs FISH | Translation, ribosomal structure and biogenesis |
| gene_1433066 | 27,56567742 | 6,208281332 | 1,123117752 | 5,527720776 | 3,24E-08 | 2,13E-05 | The Moose: suction sampler vs FISH | Translation, ribosomal structure and biogenesis |
| gene_621708 | 3,474583727 | 6,313972705 | 2,121718977 | 2,975876057 | 0,002921529 | 0,041457385 | The Moose: suction sampler vs FISH | Translation, ribosomal structure and biogenesis |

| | | | | | | | | |
|---|---|---|---|---|---|---|---|---|
| gene_87695 | 4,329055467 | 6,367679801 | 2,002465214 | 3,179920309 | 0,001473156 | 0,027593959 | The Moose: suction sampler vs FISH | Translation, ribosomal structure and biogenesis |
| gene_1443802 | 6,277307706 | 6,377494135 | 1,863497137 | 3,422325696 | 0,000620879 | 0,016004302 | The Moose: suction sampler vs FISH | Translation, ribosomal structure and biogenesis |
| gene_1588627 | 4,928575679 | 6,443528952 | 1,744985597 | 3,692597213 | 0,000221975 | 0,00835194 | The Moose: suction sampler vs FISH | Translation, ribosomal structure and biogenesis |
| gene_562333 | 4,045404733 | 6,524850654 | 1,991655158 | 3,276094573 | 0,001052533 | 0,022513889 | The Moose: suction sampler vs FISH | Translation, ribosomal structure and biogenesis |
| gene_1905004 | 6,902952228 | 6,564055624 | 1,740821856 | 3,77066476 | 0,000162813 | 0,006848694 | The Moose: suction sampler vs FISH | Translation, ribosomal structure and biogenesis |
| gene_1835760 | 4,760850583 | 6,580716547 | 1,854078312 | 3,549319629 | 0,000386228 | 0,011923713 | The Moose: suction sampler vs FISH | Translation, ribosomal structure and biogenesis |
| gene_788907 | 6,06588336 | 6,583419351 | 1,833627582 | 3,590379756 | 0,000330197 | 0,010821635 | The Moose: suction sampler vs FISH | Translation, ribosomal structure and biogenesis |
| gene_501526 | 4,400457648 | 6,620210666 | 1,922235626 | 3,444016216 | 0,000573141 | 0,015246057 | The Moose: suction sampler vs FISH | Translation, ribosomal structure and biogenesis |
| gene_1802606 | 3,537277584 | 6,642816669 | 1,964441775 | 3,381528918 | 0,000720836 | 0,017685616 | The Moose: suction sampler vs FISH | Translation, ribosomal structure and biogenesis |
| gene_907217 | 4,206443189 | 6,696453225 | 2,069461652 | 3,235843108 | 0,00121284 | 0,024471718 | The Moose: suction sampler vs FISH | Translation, ribosomal structure and biogenesis |
| gene_266490 | 7,765865009 | 6,719297651 | 1,792696007 | 3,748152293 | 0,000178142 | 0,007226628 | The Moose: suction sampler vs FISH | Translation, ribosomal structure and biogenesis |
| gene_1455877 | 5,291523953 | 6,735166921 | 1,988913082 | 3,386355583 | 0,000708275 | 0,017467861 | The Moose: suction sampler vs FISH | Translation, ribosomal structure and biogenesis |
| gene_814476 | 5,754706297 | 6,763316362 | 1,733823188 | 3,900810884 | 9,59E-05 | 0,0048661 | The Moose: suction sampler vs FISH | Translation, ribosomal structure and biogenesis |
| gene_1816317 | 10,92684115 | 6,816876573 | 1,605979731 | 4,244684065 | 2,19E-05 | 0,001774529 | The Moose: suction sampler vs FISH | Translation, ribosomal structure and biogenesis |
| gene_216694 | 17,99780912 | 6,852059608 | 1,49909581 | 4,570794984 | 4,86E-06 | 0,000647361 | The Moose: suction sampler vs FISH | Translation, ribosomal structure and biogenesis |
| gene_156912 | 7,341515526 | 7,004710634 | 1,736109551 | 4,03471695 | 5,47E-05 | 0,003323761 | The Moose: suction sampler vs FISH | Translation, ribosomal structure and biogenesis |
| gene_1064144 | 3,367566662 | 7,01460438 | 2,174418877 | 3,225967385 | 0,001255477 | 0,024995522 | The Moose: suction sampler vs FISH | Translation, ribosomal structure and biogenesis |
| gene_1149138 | 5,573268544 | 7,051235445 | 1,744460973 | 4,042071193 | 5,30E-05 | 0,003274642 | The Moose: suction sampler vs FISH | Translation, ribosomal structure and biogenesis |
| gene_1064164 | 3,411375861 | 7,064768867 | 2,286529586 | 3,08973429 | 0,002003356 | 0,033216015 | The Moose: suction sampler vs FISH | Translation, ribosomal structure and biogenesis |
| gene_1049756 | 4,197129609 | 7,225108452 | 2,238580909 | 3,227539564 | 0,001248598 | 0,024902046 | The Moose: suction sampler vs FISH | Translation, ribosomal structure and biogenesis |
| gene_1255983 | 4,228249482 | 7,238259119 | 2,137910449 | 3,38566993 | 0,000710047 | 0,017499807 | The Moose: suction sampler vs FISH | Translation, ribosomal structure and biogenesis |
| gene_194068 | 7,913178868 | 7,292438804 | 1,815672861 | 4,016383657 | 5,91E-05 | 0,003520238 | The Moose: suction sampler vs FISH | Translation, ribosomal structure and biogenesis |
| gene_193454 | 5,268125909 | 7,519442845 | 2,170201267 | 3,464859669 | 0,000530508 | 0,014487375 | The Moose: suction sampler vs FISH | Translation, ribosomal structure and biogenesis |
| gene_485633 | 7,221425467 | 7,743575817 | 1,924492816 | 4,02369692 | 5,73E-05 | 0,003454667 | The Moose: suction sampler vs FISH | Translation, ribosomal structure and biogenesis |
| gene_1429642 | 8,360620053 | 7,813517078 | 1,930514119 | 4,047376292 | 5,18E-05 | 0,003223638 | The Moose: suction sampler vs FISH | Translation, ribosomal structure and biogenesis |
| gene_1327606 | 7,860609982 | 7,863886844 | 1,937100579 | 4,059617208 | 4,92E-05 | 0,003123845 | The Moose: suction sampler vs FISH | Translation, ribosomal structure and biogenesis |
| gene_475925 | 9,43588462 | 8,443864679 | 1,850553427 | 4,562886191 | 5,05E-06 | 0,000660549 | The Moose: suction sampler vs FISH | Translation, ribosomal structure and biogenesis |
